# A molecular grammar for environmental sensitivity in intrinsically disordered protein regions

**DOI:** 10.64898/2026.04.20.719774

**Authors:** C Enriquez-Toledo, CA Ponce-Diego, GR Lopez-Herrera, K Nguyen, EE Rodríguez, S Biswas, S Barron-Huerta, TC Boothby, CL Cuevas-Velazquez

## Abstract

Cells must continuously sense and respond to changes in their physicochemical environment, yet the molecular mechanisms underlying this process remain incompletely understood. Intrinsically disordered protein regions (IDRs) constitute a substantial fraction of the eukaryotic proteome and participate in nearly every major cellular process. IDRs exist as dynamic conformational ensembles whose properties are strongly influenced by both amino acid sequence and the physicochemical environment. Although this intrinsic sensitivity has suggested that IDRs may contribute to environmental sensing of the cellular environment, the sequence-encoded principles that enable such sensing remain poorly understood. Here we identify a molecular grammar that captures key sequence features underlying variation in environmental ensemble sensitivity across diverse IDRs. Using 188 IDRs derived from organisms across all kingdoms of life, we systematically quantify changes in ensemble properties in response to hyperosmotic stress in living cells. We show that ensemble sensitivity is significantly associated with specific sequence features and predicted ensemble dimensions, with homogeneous patterning of oppositely charged residues emerging as a strong predictor of high sensitivity in charged IDRs. This sequence organization is necessary for high sensitivity in naturally-occurring IDRs and sufficient to confer high responsiveness in *de novo* designed low-complexity sequences. We further demonstrate that the presence of additional structured domains within IDR-containing proteins can substantially modulate ensemble sensitivity relative to isolated IDRs, revealing an additional layer of modulation at the multi-domain protein level. Applying the molecular grammar to the with-no-lysine (WNK) kinase crowding-sensing system, we find that regions within the disordered C-terminal domain – the domain necessary for crowding-induced phase separation and cell volume recovery – are predicted as hypersensitive, a property that is conserved across distant WNK homologues despite sequence divergence, and that is shared by heterologous IDRs capable of functionally replacing it. Together, our results establish environmental ensemble sensitivity as a sequence-encoded, context-dependent, and quantitatively tunable property of IDRs, provide a framework for understanding how physicochemical changes can be transduced in cells without dedicated receptor architectures as exemplified by the WNK crowding-sensing system, and offer design principles for engineering synthetic protein-based environmental sensors.

## MAIN

Cells are constantly exposed to fluctuations in their physicochemical environment, yet how such changes influence protein conformations and their functions in living systems remains incompletely understood. Protein structure and conformational biases dictate biological activity, but in cells proteins operate within a heterogeneous, crowded, and dynamic milieu that is frequently perturbed, particularly under stress. Changes in osmolarity, hydration and macromolecular crowding can directly alter protein conformations and thereby modulate function^1–4^. Although these effects have been extensively explored *in silico* and *in vitro*^5–9^, how physicochemical perturbations influence protein ensembles within living cells has only recently begun to emerge^10–17^. This gap is especially relevant during stress, when alterations in the cellular environment are amplified and can broadly impact proteome behavior.

Intrinsically disordered proteins (IDPs) and protein regions (IDRs) lack a stable three-dimensional structure and instead populate dynamic ensembles of rapidly interconverting conformations^18^. Unlike folded domains, the conformational properties of IDR ensembles are highly sensitive to the physicochemical characteristics of their environment, raising the possibility that environmental responsiveness may directly emerge from sequence-encoded ensemble properties that bias ensemble behavior under changing conditions^19–21^. Consistent with this view, IDR-based sensing has been reported for multiple physicochemical parameters– including pH, temperature, macromolecular crowding, and water availability– often manifested through environment-dependent phase behavior and condensate formation^22–32^. Such observations, spanning diverse organisms across the tree of life, suggest that environmental sensitivity may represent a broadly conserved feature of the disordered proteome^21^. Importantly, these behaviors appear to arise not from dedicated receptor architectures, but from intrinsic ensemble properties encoded in the sequence.

Recent evidence indicates that the sensitivity of IDRs ensembles to physicochemical perturbations is influenced by sequence-encoded conformational biases^15^. This implies that IDRs populate different regions of sequence-ensemble space and vary in their propensity to respond to environmental changes. Given that IDRs constitute approximately one third of the eukaryotic proteome, variability in ensemble responsiveness to the environment could have widespread consequences for cellular behavior during homeostasis and stress^18^. However, most studies linking IDR sequences, ensemble properties, and environmental responsiveness have been conducted in simplified *in vitro* systems, limiting our understanding of how these relationships operate in living cells^21^. As a result, the sequence-level principles that bias environmental sensitivity of IDRs *in vivo* remain largely undefined.

Here, we systematically characterize the ensemble sensitivity of 188 naturally occurring IDRs derived from organisms spanning all kingdoms of life to hyperosmotic stress in living cells. Using ensemble FRET, we identify sequence-encoded ensemble properties that are necessary for elevated levels of sensitivity and sufficient for *de novo* design of environmentally responsive low-complexity IDRs. We further show that structured domain context modulates the observed ensemble sensitivity relative to isolated IDRs. Finally, we show that the molecular grammar identifies sensitive disordered regions within a physiological crowding sensor. Together, our results define a molecular grammar that biases environmental ensemble sensitivity according to sequence organization in IDRs and establish a framework for understanding how physicochemical perturbations reshape the conformational behavior of the disordered proteome in cells.

## RESULTS

### IDRs are differentially sensitive to hyperosmotic stress in living cells

To systematically interrogate ensemble sensitivity of IDRs to hyperosmotic stress *in vivo*, we used ensemble Förster resonance energy transfer (FRET) in live yeast cells. We previously used this approach to estimate the ensemble sensitivity of AtLEA4-5, a plant IDP, to hyperosmotic stress in bacteria, yeast, plant, and human cell lines^20^. In this method, energy transfer (a proxy for the global dimensions of IDRs) is contrasted before and after exposure to increasing levels of hyperosmotic stress stimulated with sodium chloride (NaCl)^33^. Global dimensions of the IDR ensembles can be tracked with FRET because each IDR sequence is sandwiched by two fluorescent proteins (FPs)^20,33^. We assembled a diverse library of 200 naturally occurring IDRs derived from proteins spanning all kingdoms of life and generated constructs in the FRET context (Methods, **Supplementary Table 1, Supplementary Table 2**). From this library, 188 IDRs were robustly expressed in living yeast cells and included in subsequent quantitative analyses, representing a valuable toolkit for dynamically testing the ensemble sensitivity of IDRs to perturbations of the cellular environment (Methods, **Supplementary Table 1**). The sequence-encoded parameters of the IDRs in the library cover a wide range of values, except for certain properties that are intrinsically biased for IDRs, such as hydropathy or the fraction of polar residues (**Extended Data Fig. 1**). Comparison of sequence features between IDRs of the library and the 33,592 IDRs of the human proteome (human IDRome) confirms the diverse representation of the IDRs from our library (**Extended Data Fig. 1**).

We individually subjected cells expressing each IDR of the library to hyperosmotic stress by treating them with increasing concentrations of NaCl (**Fig. 1a**). We used the highly sensitive previously characterized AtLEA4-5 IDP as a reference^20^. For each IDR, we quantified the normalized (to the untreated condition) acceptor-to-donor emission ratio (FRET ratio, DxAm/DxDm), and verified a typical FRET behavior by comparing the normalized fluorescence emission spectra under every NaCl treatment. To simplify access to data, we created a detailed catalog of ensemble sensitivity for all IDRs, which can be found in **Extended File 1**. We found large variability in ensemble sensitivity levels across IDRs (**Fig. 1, Extended File 1**). For example, we found very sensitive IDRs ensembles, such as the C-terminal region of human TAR DNA-binding protein 43 (TDP-43, IDR001, residues 266-414) or the C-terminal region of human cellular tumor antigen p53 (p53, IDR008, residues 320-393) (**Fig. 1b,e**). In contrast, we also found IDRs with very low sensitivity, such as the C-terminal region of the *S. cerevisiae* transcriptional regulatory protein Ash1 (Ash1, IDR067, residues 417-500) (**Fig. 1f)**.

**Fig. 1.**
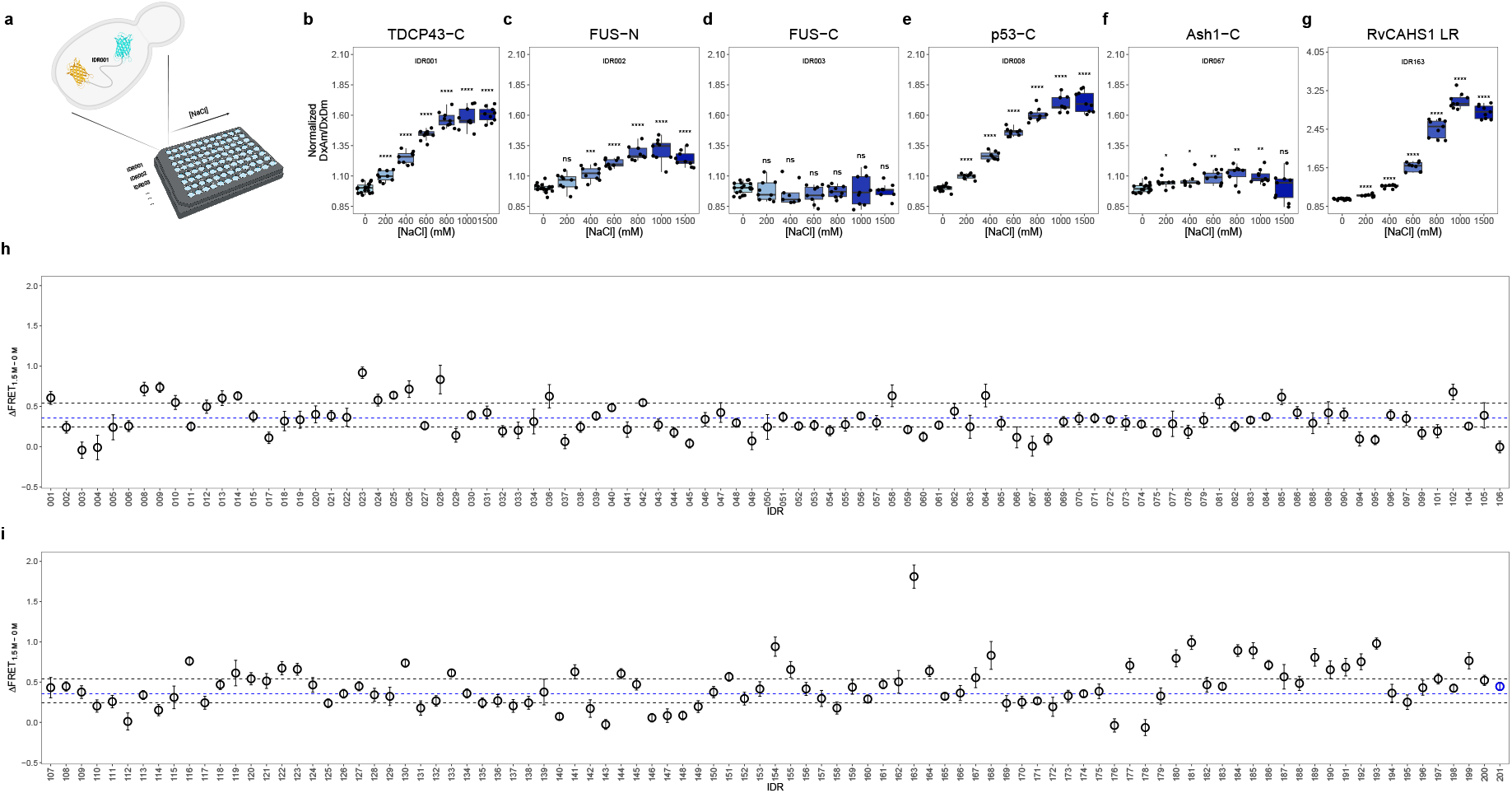
Ensemble sensitivity of 188 IDRs to hyperosmotic stress in living cells. **a**, Yeast cells expressing each IDR sandwiched by a FRET pair were subjected to different levels of hyperosmotic stress. **b-g**, Normalized FRET ratio (Normalized DxAm/DxDm) after the treatment with different NaCl concentrations in yeast cells expressing IDR001 (**b**, TDP-43_266-414_), IDR002 (**c**, FUS_1-200_), IDR003 (**d**, FUS_361-526_), IDR008 (**e**, p53_320-393_), IDR067 (**f**, Ash1_417-500_), and IDR163 (**g**, RvCAHS1 LR_901-190_). One-way ANOVA. *\*P* < 0.05, *\*\*P* < 0.01, *\*\*\*P* < 0.001, *\*\*\*\*P* < 0.0001. Boxes represent 25th-75th percentile (line at median) with whiskers at 1.5*IQR. **h**, ΔFRET_1.5M-0M_ for IDR001 to IDR106. **i**, ΔFRET_1.5M-0M_ for IDR0107-IDR201. IDR201 (blue) is AtLEA4-5 used for reference^20^. Open circles indicate the mean ΔFRET_1.5M-0M_ ± SD for each IDR (*n*= 3 independent replicates). The dashed blue line represents the median. Dashed black lines represent the 25th and 75th percentiles.

There were cases where IDRs of the same protein showed contrasting ensemble sensitivity levels. The N-terminal IDR of the human RNA-binding protein fused in sarcoma (FUS, IDR002, residues 1-200) was sensitive, displaying significant differences in normalized FRET ratio starting from the 400 mM NaCl treatment (**Fig 1c**). In contrast, the C-terminal IDR of FUS (IDR003, residues 361-526) was insensitive to the treatment, since we did not observe significant changes in normalized FRET ratio even at the most severe condition tested (1500 mM NaCl) (**Fig. 1d**).

To compare ensemble sensitivity levels across all IDRs, we calculated the FRET ratio difference between the highest and the lowest NaCl treatments (ΔFRET_1.5M-0M_). We found that the median ensemble sensitivity for the library was ΔFRET_1.5M-0M_ = 0.36 (**Fig. 1h,i**). The 25th percentile value was ΔFRET_1.5M-0M_ = 0.24 and the 75th percentile value was ΔFRET_1.5M-0M_ = 0.54 (**Fig. 1h,i**). Based on this analysis, we classified IDRs according to their position within the distribution of sensitivities. The 47 IDRs in the highest 25% of the distribution displayed very high sensitivity (hypersensitive group), while the 47 IDRs in the lowest 25% of the distribution showed very low sensitivity (insensitive group) (**Supplementary Table 3**). The group of hypersensitive IDRs had a mean ΔFRET_1.5M-0M_ = 0.72 ± 0.20, while the group of insensitive IDRs had a mean ΔFRET_1.5M-0M_ = 0.13 ± 0.09. Previously reported AtLEA4-5 showed a mean ΔFRET_1.5M-0M_ = 0.45 ± 0.05, being classified into the group of 94 IDRs within the interquartile range (average group, mean ΔFRET_1.5M-0M_ = 0.36 ± 0.08) (**Supplementary Table 3**). Among the hypersensitive group, the IDR from the cytosolic abundant heat soluble protein (RvCAHS1 LR, IDR163, residues 90-190) from the water bear (tardigrade) *Ramazzottius varieornatus* was by far the most sensitive of the library, with a mean ΔFRET_1.5M-0M_ = 1.81 ± 0.14 (**Fig. 1g**). IDR163 is the linker region (LR) of RvCAHS1, a protein that plays a key role in the desiccation tolerance of tardigrades^34,35^. Interestingly, 21 IDRs of the hypersitive group (44.7%) belong to proteins involved in stress response in their respective organisms, including several LEA proteins (IDR042, IDR130, IDR193, among others), the transcriptional co-activator yes associated protein (YAP, IDR180 and IDR181), and the mRNA decapping enzyme 1A (DCP1A, IDR009)^28,36–38^. In contrast, only 2 IDRs (4.2%) from the insensitive group participate in stress response, suggesting that high sensitivity might be involved in stress perception and response^20,21^.

These data reveal broad diversity in ensemble sensitivity across 188 IDRs from all kingdoms of life. Strikingly, even IDRs within the same protein can display contrasting sensitivities, indicating that ensemble sensitivity is not a global property of the protein context but instead arises from physicochemical features and conformational biases encoded within each individual sequence. This observation raises the possibility that specific aspects of sequence organization and intrinsic ensemble dimension systematically bias sensitivity.

### Sequence organization and predicted ensemble dimensions bias ensemble sensitivity

We leveraged the diversity of IDR sequences and ensemble sensitivity levels in our dataset to identify sequence features associated with the upper and lower ends of the sensitivity continuum. To this end, we used the previously described operational categories related to the extremes in responsiveness, hereafter termed “hypersensitive” (47 IDRs) and “insensitive” (47 IDRs), corresponding to the upper and lower tails of the continuous sensitivity distribution (**Supplementary Table 3**). First, we compared different sequence-encoded parameters including the fraction of charged residues (FCR), charge patterning (kappa, *κ*), sequence charge decoration (SCD), the absolute net charge per residue (absolute NCPR; |NCPR|), hydropathy, sequence hydropathy decoration (SHD), isoelectric point (pI), and the fraction of aromatic, aliphatic, or polar residues. Additionally, we classified the IDRs based on the region they occupy in the Das–Pappu diagram of states^39^ (**Extended Data Fig. 2**). We found that FCR was lower in hypersensitive IDRs (mean FCR = 0.25) than insensitive IDRs (mean FCR = 0.35) (difference of means = 0.10, 99%CI [0.03, 0.17], *P* = 1.3×10^-5^) (**Fig. 2a**). Hypersensitive IDRs showed lower *κ* values (mean *κ* = 0.15) than insensitive IDRs (mean *κ* = 0.25) (difference of means = 0.10, 99%CI [0.03, 0.14], *P* = 4.9×10^-4^) (**Fig. 2b**). Hydropathy was higher in hypersensitive IDRs (mean hydropathy = 3.6) than insensitive IDRs (mean hydropathy = 3.2) (difference of means = 0.4, 99%CI [0.11, 0.54], *P* = 1×10^-4^) (**Fig. 2c**). This is consistent with the higher SHD in hypersensitive IDRs (mean SHD = 4.4) than insensitive IDRs (mean SHD = 3.5) (difference of means = 0.9, 99%CI [0.50, 1.19], *P* = 6.1×10^-9^) (**Fig. 2d**). Finally, we found that hypersensitive IDRs have an |NCPR| close to zero (mean |NCPR| = 0.04), as opposed to insensitive IDRs (mean |NCPR| = 0.11) (difference of means = 0.07, 99%CI [0.04, 0.11], *P* = 1.2×10^-6^) (**Fig. 2e**). The fraction of polar, aliphatic, or aromatic residues, SCD, and pI did not show strong significant differences between the two groups (*P* ≥0.01) (**Extended Data Fig. 3a-e**). Interestingly, we found that the group of hypersensitive IDRs was enriched in IDRs that fall into the region 1 of the Das-Pappu diagram-of-states (weak polyampholytes and polyelectrolytes: globules and tadpoles), whereas the group of insensitive IDRs was enriched in IDRs that fall into the region 3 (strong polyampholytes: coils, hairpins and chimeras) (**Fig. 2f**).

**Fig. 2.**
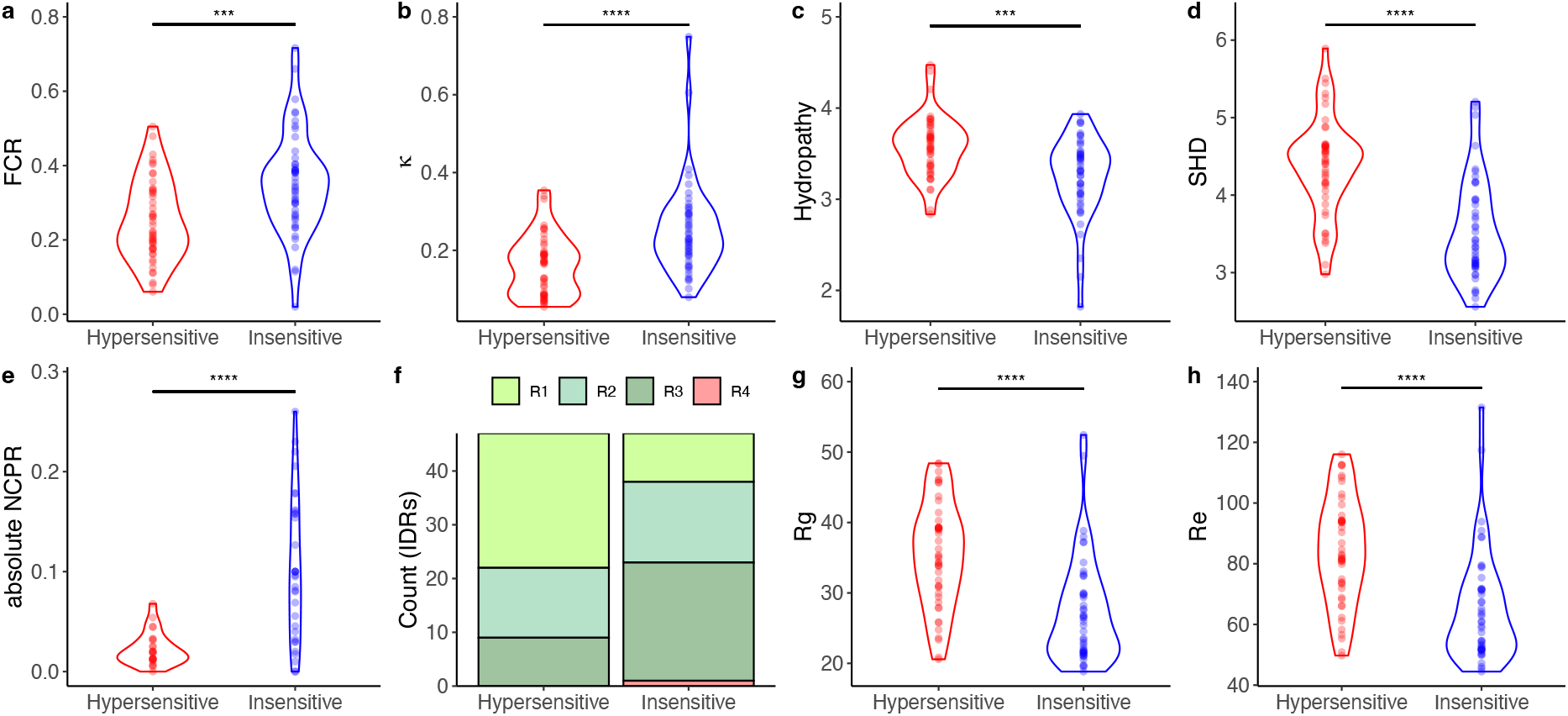
Sequence features and predicted IDR ensemble dimensions associated with ensemble sensitivity to hyperosmotic stress in cells. **a-e**, Sequence features of IDRs belonging to the hypersensitive (red) or insensitive (blue) groups. **a**, fraction of charged residues (FCR); **b**, kappa (*κ*); **c**, hydropathy; **d**, sequence hydropathy decoration (SHD); **e**, absolute net charge per residue (|NCPR|). Each point represents the corresponding value for an IDR. Color intensity indicates the representation of values within the same range. **f**, proportion of hypersensitive or insensitive IDRs categorized into the R1, R2, R3, and R4 regions of the Das-Pappu diagram-of-states. **g-h**, predicted ensemble dimensions of IDRs belonging to the hypersensitive (red) or insensitive (blue) groups. **g**, predicted radius of gyration (*R*g); **h**, predicted end-to-end distance (*R*e). Each point represents the corresponding value for an IDR. Color intensity indicates the representation of values of the same range. Mann-Whitney U test *\*P* < 0.05, *\*\*P* < 0.01, *\*\*\*P* < 0.001, *\*\*\*\*P* < 0.0001.

To determine whether the sequence features identified above translate into differences in global conformational behavior, we predicted the average ensemble dimensions of hypersensitive and insensitive IDRs from their sequence using the deep-learning model ALBATROSS^40^. The properties predicted include the radius of gyration (*R*g), end-to-end distance (*R*e), asphericity, and polymer scaling exponent (ν) and prefactor (ρ_0_). We found that *R*g was larger in hypersensitive IDRs (mean *R*g = 35.4 Å) than insensitive IDRs (mean *R*g = 27.3 Å) (difference of means = 8.1 Å, 99%CI [4.08, 12.16], *P* = 8.3×10^-7^) (**Fig. 2g**). Similarly, *R*e was larger in hypersensitive IDRs (mean *R*e = 84.1 Å) than insensitive IDRs (mean *R*e = 65.0 Å) (difference of means = 19.1 Å, 99%CI [9.4, 28.8], *P* = 1.3×10^-6^) (**Fig. 2h**). We found no significant differences between the two groups for asphericity, ν, and ρ_0_ (**Extended Data Fig. 3g**,**h**). These results indicate that hypersensitive IDRs tend to adopt more expanded conformational ensembles under basal conditions. Together with the differences in charge patterning and net charge balance, this suggests that global chain expansion contributes to biasing ensemble sensitivity across the continuum of responses.

To formally evaluate whether these sequence features collectively bias the probability of high ensemble sensitivity, we fitted a binomial logistic regression model using IDRs from the high- and low-sensitivity extremes of the response distribution. The model estimates the probability that an IDR belongs to the high-sensitivity extreme. We included FCR, *κ, R*e, and |NCPR| as predictor variables, selected to minimize multicollinearity among predictors. We found that *κ* (OR = 0.90, *P* = 0.01) and |NCPR| (OR = 0.83, *P* = 0.002) showed a significant inverse association with the probability of belonging to the hypersensitive group, while *R*e showed a significant direct association with the same group (OR = 1.04, *P* = 0.03). The model had an accuracy of 0.82, a true positive rate (sensitivity) of 0.85, a true negative rate (specificity) of 0.79, and an Akaike information criterion (AIC) of 85.9. These results indicate that IDRs with low *κ* values, |NCPR| close to zero, and relatively expanded (high Re) have a greater probability of being classified as hypersensitive IDRs. In contrast, FCR did not show a significant association with sensitivity (OR = 0.96, *P* = 0.13). To test the model, we predicted the probabilities in the rest of the IDRs from the average response group (94 IDRs). We expected that the IDRs between Q2 (median) and Q3 (75th percentile) should have a higher probability to be predicted as hypersensitive, while the IDRs between Q1 (25th percentile) and Q2 should have a higher probability to be predicted as insensitive. Using a probability threshold of 0.5, the model predicted 27 out of 47 IDRs from the Q2-Q3 group as hypersensitive (true positive rate of 0.57), and predicted 36 out of 47 IDRs from the Q1-Q2 group as insensitive (true negative rate of 0.77). Consistent with the multifactorial nature of environmental ensemble sensitivity, our results show that the associated sequence features do not define a binary switch between sensitive and insensitive IDRs, but instead shift the probability of sensitivity across the continuum of responses.

### Evolutionary conserved charged organization is associated with high ensemble sensitivity

The binomial logistic regression analysis identified *κ* and |NCPR| as key sequence features associated with high ensemble sensitivity to hyperosmotic stress in cells. To experimentally evaluate whether low *κ* and near neutral NCPR bias ensemble sensitivity at the level of individual sequences, we focused our attention on the most sensitive IDR of the library: RvCAHS1 LR (IDR163, residues 90-190) from the tardigrade *R. varieornatus*. Despite its high content of charged residues (FCR = 0.5), RvCAHS1 LR has a NCPR close to neutrality (NCPR = -0.03) because its content of positively and negatively charged residues are very similar (fraction of positive = 0.24, fraction of negative = 0.26). In addition, the *κ* value of RvCAHS1 LR is 0.06, indicating that residues with opposite charges are well-mixed across its sequence. Closer inspection of NCPR across the RvCAHS1 LR sequence revealed such charge patterning (**Fig. 3a**). The well-mixed charge patterning of the LR of CAHS proteins is conserved across tardigrade species^41^. Additionally, the extremely low *κ* value of RvCAHS1 LR makes its charged residues more well-mixed than over 99% of all tardigrade IDRs^32^. If *κ* and NCPR contribute to the sensitivity of IDRs, we hypothesized that CAHS LR homologues with low *κ* values and NCPR close to zero would show high sensitivity in our *in vivo* FRET approach. The CAHS LR system therefore provides a naturally occurring test case to evaluate whether evolutionary conserved charge organization aligns with the sequence features statistically associated with high ensemble sensitivity.

**Fig. 3.**
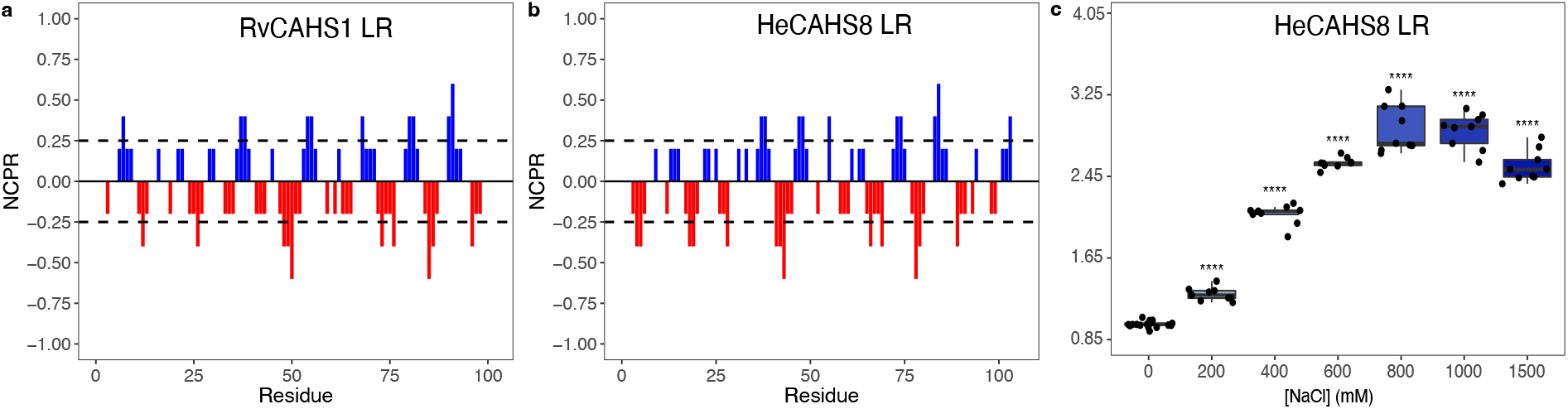
Evolutionary conserved charged organization is associated with high ensemble sensitivity. **a**, Net charge per residue across the sequence of RvCAHS1 LR. **b**, Net charge per residue across the sequence of HeCAHS8 LR. **c**, Normalized FRET ratio (Normalized DxAm/DxDm) after the treatment of different concentrations of NaCl for yeast cells expressing HeCAHS8 LR. One-way ANOVA. *\*P* < 0.05, *\*\*P* < 0.01, *\*\*\*P* < 0.001, *\*\*\*\*P* < 0.0001. Boxes represent 25th-75th percentile (line at median) with whiskers at 1.5*IQR.

To test this, we cloned the ORF that codes for the LR of the CAHS protein from the tardigrade *Hypsibius exemplaris* in our FRET system and expressed it in yeast. Despite having only a sequence identity of 57.5% and a sequence similarity of 69% (**Extended Data Fig. 4**), HeCAHS8 LR has very similar FCR, NCPR and *κ* values to RvCAHS1 LR (FCR = 0.52, NCPR = -0.01, and a *κ* = 0.06) (**Fig. 3b**). In agreement with our hypothesis, HeCAHS8 LR showed extremely high ensemble sensitivity to hyperosmotic stress, with a mean ΔFRET_1.5M-0M_ = 1.79 ± 0.29 (**Fig. 3c**). This finding further supports the idea that conformational ensembles encoded by low *κ* and near-neutral NCPR bias IDRs towards high sensitivity to hyperosmotic stress in cells. Together, these results indicate that charge organization features statistically associated with high ensemble sensitivity are not only predictive across diverse IDRs but are also conserved within homologous sequences, supporting their functional relevance. The evolutionary conservation of low *κ* and NCPR values, together with their extreme sensitivity, suggests that ensemble sensitivity to hyperosmotic stress may be functionally relevant for this class of IDRs.

### Homogeneous charge mixing is necessary and functionally sufficient to encode high ensemble sensitivity

The conservation of homogeneous charge mixing and extreme ensemble sensitivity in CAHS LR homologues suggested that charge patterning may not only correlate with, but directly encode ensemble sensitivity. To directly test causality, we generated three HeCAHS8 LR variants in which amino acid composition was preserved (parameters such as FCR and NCPR remain the same), while charge patterning (*κ* value) was systematically perturbed (**Extended Data Fig. 5**). We named these variants V1 (*κ* = 0.174), V2 (*κ* = 0.323), and V3 (*κ* = 0.69). We found that increasing charge segregation (*κ* → 1) abolished the high ensemble sensitivity of HeCAHS8 LR, following a logarithmic fit (R^2^ = 0.83) (**Fig. 4a**). These results confirmed that the homogeneous distribution of residues with opposite charges across the sequence of highly charged IDRs is necessary for their extreme ensemble sensitivity to hyperosmotic stress in cells.

**Fig. 4.**
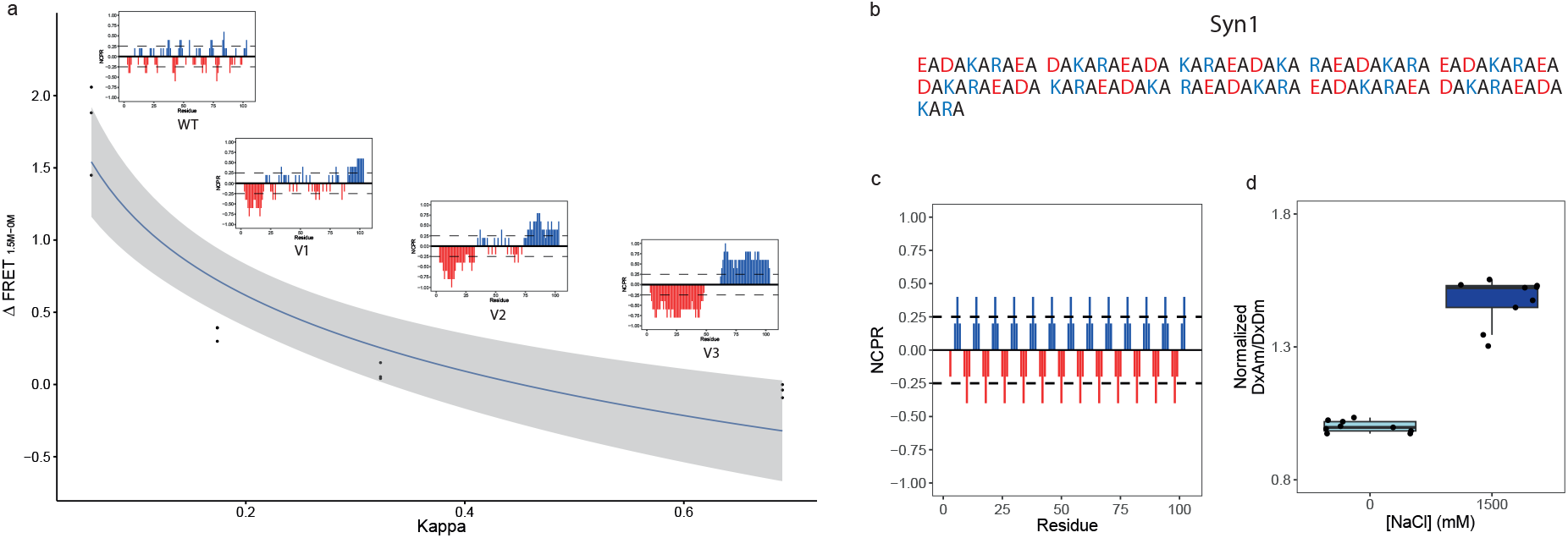
Homogeneous charge mixing is necessary and functionally sufficient to encode high ensemble sensitivity. **a**, ΔFRET_1.5M-0M_ for wild type (WT) HeCAHS8 LR and V1, V2, and V3 *κ* variants. Plots for net charge per residue across the sequence of the variants are shown above each ΔFRET_1.5M-0M_ value. The continuous line was smoothed using R with a logarithmic method smoothing function. Shaded region indicates 95% confidence interval **b**, Amino acid sequence of Syn1. Positively charged residues in blue, negatively charged residues in red, aliphatic residues in black. **c**,Net charge per residue across the sequence of Syn1. **d**, Normalized FRET ratio (Normalized DxAm/DxDm) after the treatment of 0 mM NaCl or 1,500 mM NaCl for yeast cells expressing Syn1. One-way ANOVA. *\*P* < 0.05, *\*\*P* < 0.01, *\*\*\*P* < 0.001, *\*\*\*\*P* < 0.0001. Boxes represent 25th-75th percentile (line at median) with whiskers at 1.5*IQR.

We next asked if we could design sensitive IDRs *de novo* by using charge patterning as the main design principle. We designed a low complexity synthetic IDR with specific sequence-encoded properties similar to the wild type (WT) CAHS LR proteins, including length, FCR, NCPR, and *κ*. The synthetic IDR, named Syn1, was composed of only 5 types of amino acids: Arg (12.5%), Lys (12.5%), Asp (12.5%), Glu (12.5%), and Ala (50%). Syn1 consists of 13 repetitions of the EADAKARA sequence domain (**Fig. 4b**). Syn1 is predicted to be disordered (**Extended Data Fig. 6**), has a length of 104 residues, FCR = 0.5, and NCPR = 0. Syn1 has a charge patterning that resembles that of CAHS LRs (*κ* = 0.047) (**Fig. 4c**). Strikingly, we found that the normalized FRET ratio of Syn1 increased in response to hyperosmotic stress in cells, demonstrating that the homogeneous charge patterning is sufficient for designing synthetic IDRs that are sensitive to changes in the physicochemical environment (**Fig. 4d**). Syn1 was as sensitive (mean ΔFRET_1.5M-0M_ = 0.47 ± 0.29) as the previously reported AtLEA4-5 (mean ΔFRET_1.5M-0M_ = 0.45 ± 0.05), which was used to generate SED1 hyperosmotic stress FRET biosensor^20^ (**Fig. 1i, 4d**). However, Syn1 was less sensitive than WT RvCAHS1 LR or HeCAHS8 LR (**Fig. 1g, 3c, 4d**), being classified into the group of IDRs with average sensitivity of our library (**Fig. 1h,i, Supplementary Table 3**). This result highlights that charge patterning acts in concert with other sequence-encoded properties to modulate the magnitude of environmental ensemble sensitivity.

Altogether, these results indicate that the homogeneous charge mixing is necessary for extreme ensemble sensitivity in highly charged IDRs and sufficient to confer high sensitivity when embedded in a compatible compositional background. Thus, charge patterning constitutes a tunable sequence determinant that can be exploited to engineer environmentally responsive IDRs.

### Protein domain context modulates ensemble sensitivity of IDRs

While charge patterning encodes sensitivity at the level of isolated IDRs, the protein domain context might have an impact in how this encoded responsiveness is manifested in the cellular environment. So far, we have evaluated the sensitivity of fully disordered IDPs or IDRs in isolation. To test the impact of the presence of other protein domains on the ensemble sensitivity readout of our FRET system, we used HeCAHS8 as a model. The presumed function of HeCAHS8 is preventing the inactivation of labile proteins upon desiccation^16,17,34,41,42^, likely through the formation of reversible gels which increase protection to cells experiencing hyperosmotic shock^32,43^. In cells, the gelation capacity of HeCAHS8 is dependent on the interaction between its protein domains^32,43^. HeCAHS8 is composed of three different domains: the N-terminus, the middle LR, and the C-terminus (**Fig. 5a**). *In silico* simulations of HeCAHS8 suggest that it adopts a dumbbell-like conformational ensemble where the N- and C-termini domains are relatively collapsed and are separated by the extended LR^32^. As described previously, the LR of HeCAHS8 displays extreme sensitivity to hyperosmotic stress (**Fig. 5b**). However, when we analyzed the full-length HeCAHS8 (FL_HeCAHS8), we found that the sensitivity was diminished when compared to the LR alone (**Fig. 5c,h**), indicating that the N- and/or the C-terminal collapsed domains of FL_HeCAHS8 negatively impact sensitivity. The N- and C-terminal domains of FL_HeCAHS8 contribute to its gel formation ability^32^. While FL_HeCAHS8 gelates *in vitro* and *in vivo*, the LR alone is unable to form gels under the conditions tested^32^. We hypothesized that gel formation constrains or redistributes the conformational ensemble of HeCAHS8, thereby limiting the intramolecular compaction that underlies the FRET sensitivity readout. To test this hypothesis, we characterized the sensitivity of two different variants of HeCAHS8 with divergent gel formation properties: 2X Linker Region (2X_LR) and FL_Proline (**Fig. 5a, Extended Data Fig. 7**)^32^. 2X_LR was designed to maintain the N- and C-terminal domains held apart by a tandem duplication of the LR^32^. On the other hand, FL_Proline keeps the full-length context, but the β-structure propensity is reduced through the insertion of four prolines in the C-terminal domain^32^. 2X_LR gelates at a lower concentration than FL_HeCAHS8 *in vitro* (increased gelation capacity), while FL_Proline is defective in gel formation^32^. We found that 2X_LR was insensitive to hyperosmotic stress treatment (**Fig. 5d,h**). In contrast, FL_Proline showed a sensitivity level compared to that of the isolated LR, despite having the full-length context (**Fig. 5e,h**). These results suggest that gelation prevents the hyperosmotic stress-induced compaction of HeCAHS8 ensemble. In accordance with this hypothesis, two additional variants that do not gel *in vitro* or *in vivo*, where the C-terminal domain is replaced with the N-terminal domain or vice versa (NLN and CLC, respectively, **Extended Data Fig. 7**)^32^, displayed sensitivity levels similar to LR and FL_Proline (**Fig. 5f-h**).

**Fig. 5.**
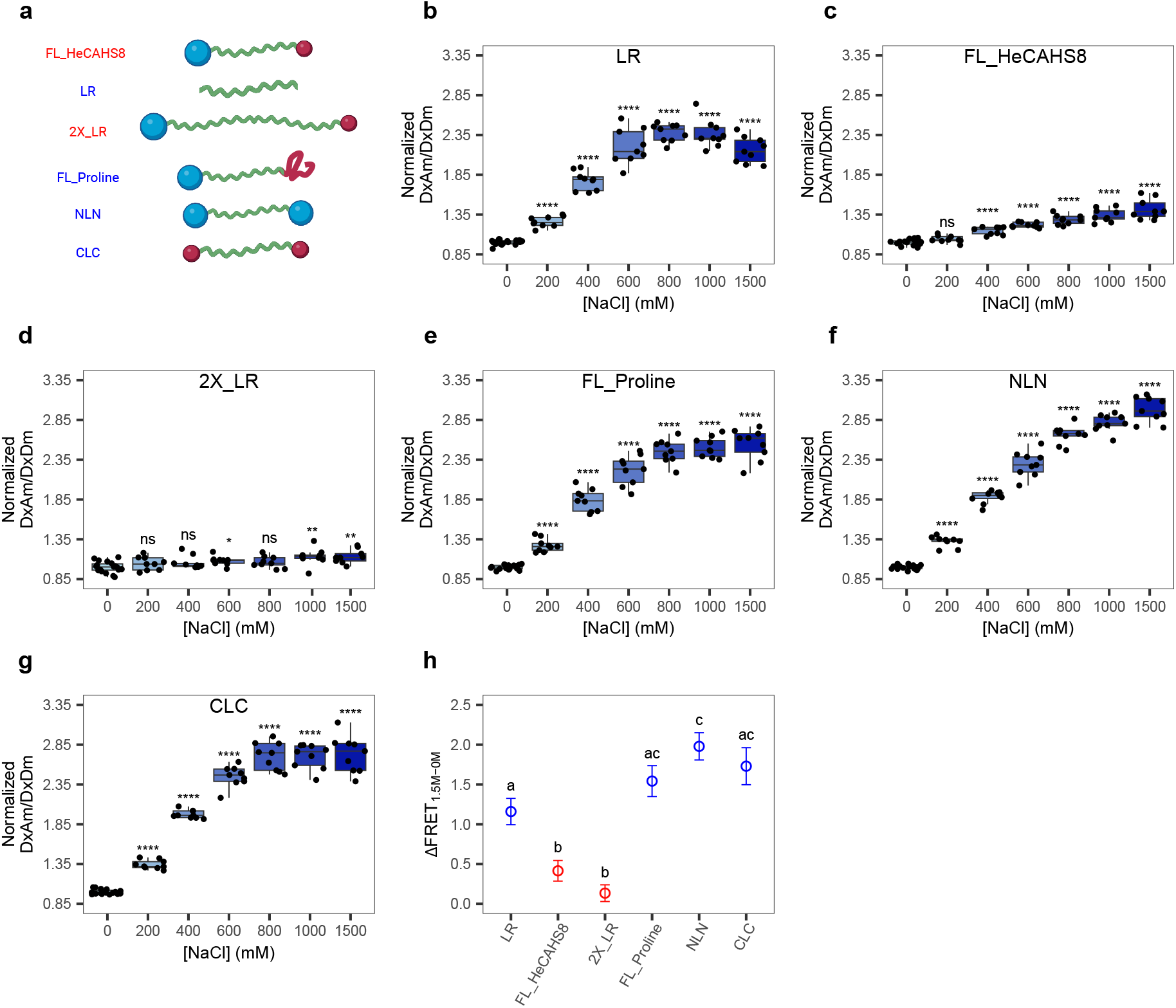
Protein domain context modulates ensemble sensitivity of IDRs. **a**, HeCAHS8 variants used in this study. Blue and red spheres represent the N- and C-terminal domains, respectively. The green segment represents the disordered linker region. Variants that form gels are labeled in red while those that do not gel are labeled in blue. **b-g**, Normalized FRET ratio (Normalized DxAm/DxDm) after the treatment of different concentrations of NaCl for yeast cells expressing HeCAHS8 LR (**b**, LR), FL_HeCAHS8 (**c**), 2X_LR (**d**), FL_Proline (**e**), NLN (**f**), or CLC (**g**). One-way ANOVA. *\*P* < 0.05, *\*\*P* < 0.01, *\*\*\*P* < 0.001, *\*\*\*\*P* < 0.0001. Boxes represent 25th-75th percentile (line at median) with whiskers at 1.5*IQR. **h**, ΔFRET_1.5M-0M_ for the indicated HeCAHS8 variants. Variants that form gels are labeled in red while those that do not gel are labeled in blue. One-way ANOVA with Tukey’s multiple comparison test, *P* < 0.01.

Altogether, these results indicate that environmental ensemble sensitivity encoded by an IDR can be modulated by its full-length protein context, particularly when supramolecular assembly competes with intramolecular compaction. In the case of HeCAHS8, gelation appears to restrict the conformational rearrangements required for hyperosmotic stress-induced compaction, thereby dampening the FRET sensitivity readout. These findings highlight that ensemble sensitivity is not solely a property of linear sequence organization, but can be dynamically influenced by higher-order protein architecture and functional state. Given the intrinsic limitations of FRET-based measurements and the diverse functional modes of IDR-containing proteins, evaluating ensemble sensitivity in full-length proteins will require context-specific analyses.

Together, our findings suggest that ensemble sensitivity to hyperosmotic stress emerges from a hierarchical organization of molecular determinants – from linear charge patterning to higher-order domain architecture – and may be dynamically tuned by functional states such as gelation. This layered regulation raises the possibility that environmental ensemble sensitivity is an evolvable feature contributing to stress adaptation in IDR containing proteins.

### The molecular grammar identifies functional crowding-responsive IDRs in WNK signaling

Having established ensemble sensitivity as a sequence-encoded and context-dependent property, we next asked whether this grammar identifies functionally responsive IDRs in a physiological system where disorder-driven environmental sensing has been directly demonstrated. The with-no-lysine (WNK) kinases are physiological macromolecular crowding sensors that activate upon hyperosmotic stress to promote cell volume recovery through the coordination of ion transporters, channels, and pumps – a response termed regulatory volume increase (RVI) (**Fig 6a**)^30^. The long C-terminal domain (CTD) of WNK kinases (**Fig 6b**) drives macromolecular crowding sensing through the formation of biomolecular condensates that concentrate and promote the activity of the signaling pathway (**Fig 6c**), ultimately resulting in RVI^30^. We analyzed the rat WNK1 CTD sequence (residues 495 - 2,126) using our sensitivity logistic regression model to calculate the predicted probability that the ensemble of this sequence is hypersensitive to hyperosmotic stress. The model identified that 70% of the rat WNK1 CTD amino acids are predicted as hypersensitive with a probability greater than 0.5 (*P* > 0.5), including a long region of 800 consecutive residues (residues 974 - 1,774 of the original full-length sequence) with a mean probability of 0.75 (*P* = 0.75) (**Fig 6d**). We next analyzed the CTD sequence of WNK homologues, from humans to protists. The CTD of these homologues has very poor sequence identity, and their amino acid composition is different between mammals, invertebrates, and protists^30^. We found that our model also predicted large regions of the CTD with high probability to be classified as hypersensitive to hyperosmotic stress in all the homologues (**Fig 6e-j**), including the fruit fly dmWNK, which also form CTD-dependent condensates during hyperosmotic stress in *Drosophila* S2 cells^30^. These findings suggest that ensemble sensitivity to hyperosmotic stress of the disordered CTD from WNK kinases is evolutionary conserved from humans to protists, highlighting the role of the molecular grammar for ensemble sensitivity of IDRs in physiological sensors of the physicochemical environment.

**Fig. 6.**
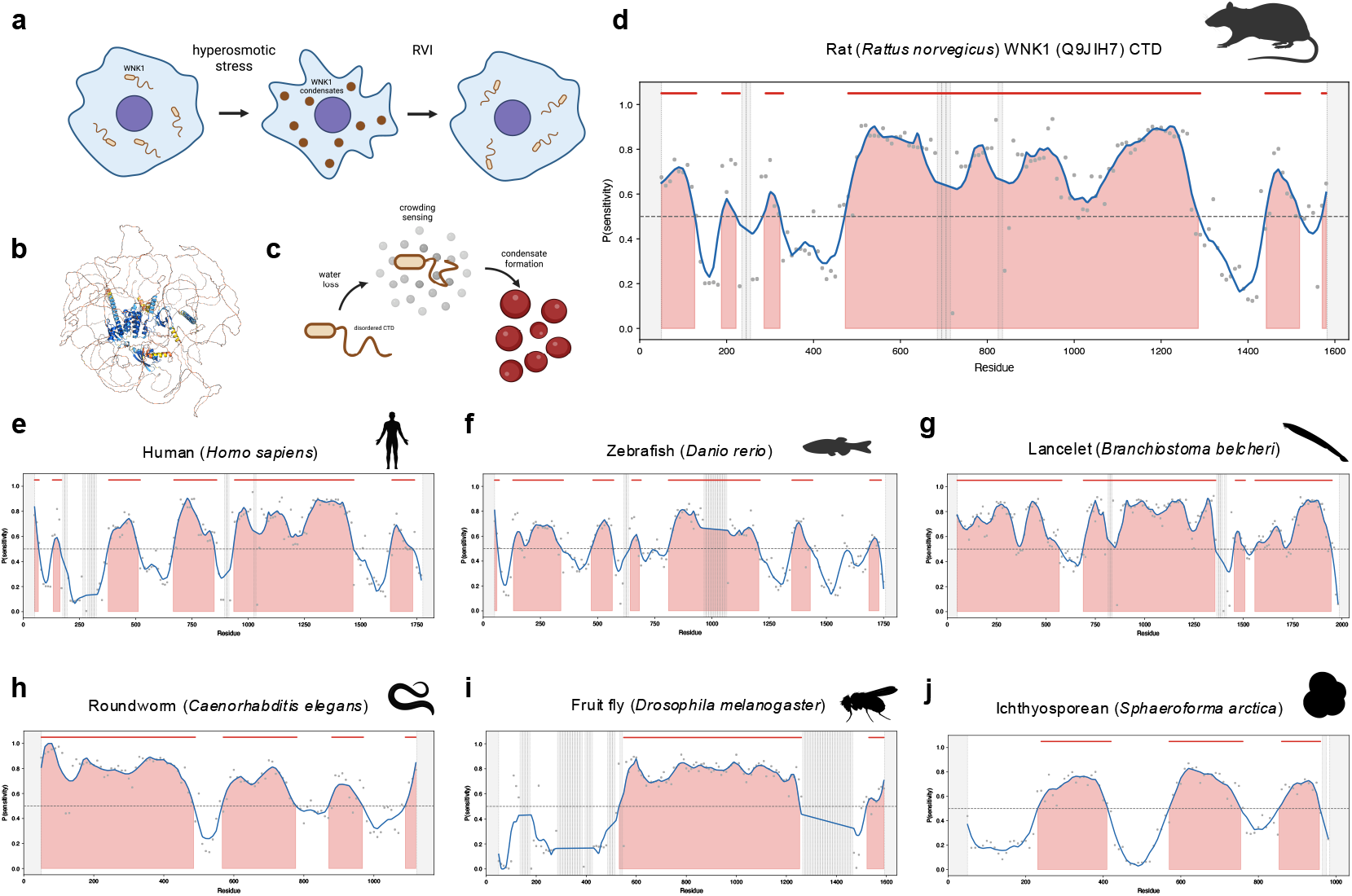
The molecular grammar identifies functional crowding-responsive IDRs in WNK signaling. **a**, in animal cells, hyperosmotic stress causes water loss, cell volume contraction, and increased macromolecular crowding. The WNK1 kinase senses changes macromolecular crowding, self-assembles into biomolecular condensates that concentrate signaling to regulate ion transporters, and thereby drives water uptake and cell volume recovery – a process termed regulatory volume increase (RVI). **b**, AlphaFold2 model of rat WNK1 showcasing the disordered regions, specially the long CTD. Colors represent standard pLDDT. **c**, Water loss causes an increase in macromolecular crowding (gray spheres) that is sensed by unknown regions within the long (1,631 residues) CTD of WNK, promoting the formation of condensates. **d**, Ensemble sensitivity profile across the CTD of rat WNK1 predicted by sliding window analysis using the logistic regression model. The probability of being classified as hypersensitive to hyperosmotic stress (crowding) is shown across the primary sequence. **e-j**, Ensemble sensitivity profile predicted across the WNK CTD of human (**e**), zebrafish (**f**), lancelet (**g**), roundworm (**h**), fruit fly (**i**), and ichthyosporea (**j**). Each data point (gray circles) represents the model output for a 100-residue window centered at the indicated position. The blue line shows the Savitzky-Golay-smoothed profile. Shaded orange areas indicate continuous regions above a probability threshold of *P* = 0.5. Horizontal red bars above the plot mark continuous segments where the smoothed probability equals or exceeds 0.5. Gray shaded regions indicate positions for which no probability score is available, either due to the center-based score assignment inherent to the sliding window approach (sequence first and last 50 residues) or to windows excluded from prediction by the model (see Methods section).

Phase separation of WNK1 kinase driven by the disordered CTD was shown to mediate RVI following cell shrinkage however, what features of the CTD sequence enable macromolecular crowding sensing in the first place were not identified (**Fig. 6b**)^30^. Boyd-Shiwarski et al. approached this by showing that two different IDRs with the ability to phase separate can orthogonally replace WNK1 CTD to rescue RVI: the N-terminal IDR from FUS (residues 1-267) and the C-terminal IDR from TDP-43 (residues 274-414)^30^. Strikingly, we showed here that the ensembles of both IDRs are sensitive to hyperosmotic stress (**Fig 1b,c**), so the authors serendipitously used sensitive IDRs for the orthogonal replacement. Furthermore, they observed that TDP-43-C, which we show is more sensitive than FUS-N, rescued volume almost as effectively as the full-length WNK1^30^, suggesting that the magnitude of ensemble sensitivity scales with functional outputs, providing a direct link between conformational responsiveness and physiological efficacy.

## DISCUSSION

The physicochemical properties of the cytoplasm are dynamic and can change substantially under stress conditions. Hyperosmotic stress induces rapid water efflux, increases intracellular macromolecular crowding, and alters mesoscale cellular organization, thereby reshaping the physical landscape in which proteins function^44,45^. While such changes are known to influence translation, degradation, phase separation, and protein stability, how individual proteins sense and respond to these perturbations at the level of conformational ensembles remains incompletely understood^44,46–48^. IDRs, whose ensembles are highly sensitive to solvent conditions *in vitro*, have been proposed to function as intracellular physicochemical sensors ^20,21^. By systematically quantifying the ensemble sensitivity of 188 naturally occurring IDRs directly in living cells, we show that environmental responsiveness is not a uniform property of disordered proteins, but instead varies widely and is biased by specific sequence-encoded features.

Our analyses define a probabilistic molecular grammar of environmental ensemble sensitivity. In this context, grammar denotes a set of sequence constraints that bias conformational ensembles toward higher or lower responsiveness, rather than a deterministic rule set. We find that charge patterning, net charge balance, and ensemble expansion collectively shape the likelihood that an IDR undergoes compaction under hyperosmotic stress. Importantly, no single parameter is sufficient to predict sensitivity across all sequences; instead, responsiveness emerges from the combined and context-dependent effects of these features, positioning sensitivity along a continuum rather than as a binary attribute. These findings are consistent with work showing that structural biases observed in IDP ensembles *in vitro* are preserved inside cells and can vary in a sequence-dependent manner in response to changes in the intracellular environment^15^. Likewise, a recent study showed that sequence–ensemble relationships tune IDR responsiveness to changes in cell volume for 32 synthetic low-complexity IDRs in human-derived cell lines^49^. Our results extend this framework by systematically mapping ensemble sensitivity across a large and diverse set of natural IDRs and by defining quantitative sequence features that bias responsiveness in crowded cellular environments.

The convergence of these studies suggests that physicochemical sensitivity of IDRs may be widespread and subject to evolutionary tuning. The evolutionary conservation of homogeneous charge organization in CAHS LR homologues, together with their extreme ensemble sensitivity, indicates that this molecular grammar is not merely descriptive but may carry functional relevance. Proteins such as CAHS and LEA proteins operate under dehydration and osmotic stress, conditions characterized by increased crowding and reduced water availability^34,50^. The ability of their disordered regions to compact in response to such perturbations may therefore contribute to protective roles during stress adaptation^41,51,52^. Our data support a model in which sequence-encoded ensemble responsiveness constitutes a physical layer of environmental sensing that can precede and potentially enable downstream functional outcomes^53^. In agreement with this, the molecular grammar for environmental sensitivity identifies long regions within the disordered CTD of WNK1, a known macromolecular crowding sensor with profound implications in organismal physiology^30^. The evolutionary conservation of high predicted sensitivity across WNK suggests that this property is under selective pressure and functionally relevant across species, a hypothesis that might extend to other signaling pathways. We envision that next steps in connecting the molecular grammar for ensemble sensitivity with functional signaling pathways will focuse on identifying new physiological sensors of the physicochemical environment within the disordered proteome.

We found that ensemble sensitivity is modulated by protein context. While isolated IDRs reveal intrinsic sequence-encoded responses, the presence of additional domains can reshape or mask these behaviors. In the case of HeCAHS8, full-length protein context and gelation capacity attenuate the hyperosmotic stress-induced compaction observed for the isolated linker region. These findings indicate that intermolecular interactions, folded domains, and stress-induced phase transitions can constrain the conformational degrees of freedom required for ensemble rearrangement, thereby influencing FRET-based sensitivity readouts. Ensemble sensitivity is therefore not an isolated property of a sequence fragment, but an emergent feature shaped by the interplay between sequence grammar and functional context. Our study specifically probes sensitivity to hyperosmotic stress, a perturbation that primarily increases intracellular crowding and alters water activity. Therefore, our findings have implications in physiological and pathological processes related to cell size, growth, and proliferation, including cancer^54^. Other environmental challenges—including shifts in pH, temperature, ionic strength, or redox state—reshape the intracellular milieu through distinct physical mechanisms^10^. We therefore do not expect that the sequence principles identified here will universally predict responsiveness across all environmental stresses. Rather, our findings define a molecular grammar of sensitivity to crowding-associated perturbations. Whether distinct environmental perturbations engage overlapping or orthogonal sequence grammars within the disordered proteome remains an open question that requires systematic investigation.

Together, our results support a model in which ensemble sensitivity arises from probabilistic sequence constraints rather than deterministic rules. Charge balance, charge patterning, and ensemble dimensions collectively bias IDRs toward sensitivity to crowding-associated perturbations, but no single parameter alone defines sensitivity. The ability to tune sensitivity through rational modification of charge organization further suggests the feasibility of designing synthetic IDRs with tailored environmental responsiveness, including biosensors optimized for specific physicochemical perturbations^55,56^. More broadly, these findings position the disordered proteome as a reservoir of tunable physical sensors that operate upstream of canonical signaling pathways, providing cells with a rapid and adaptable mechanism to navigate fluctuations in their physicochemical environment.

## METHODS

### Selection of IDRs

IDRs were selected by surveying the literature and curated disorder databases such as DispProt and MobiDB^55,56^. Only IDRs ranging between 50 and 210 residues were considered, based on previous ensemble FRET measurements *in vitro* and *in vivo*^15,19,20^. A total of 200 naturally occurring IDRs were selected from proteins spanning all kingdoms of life (**Supplementary Table 1**). The resulting library comprised sequences from animals (n = 90), higher plants (n = 74), fungi (n = 23), bacteria (n = 10), and algae (n = 3), representing 37 species. Species selection included commonly studied model organisms such as *Homo sapiens, Arabidopsis thaliana, Saccharomyces cerevisiae, Mus musculus, Drosophila melanogaster, Rattus norvegicus*, and *Escherichia coli*. The library included IDRs derived from proteins implicated in diverse cellular processes, including stress-responsive proteins and proteins from extremophiles or extremotolerant organisms (e.g. late embryogenesis abundant proteins and tardigrade disordered proteins)^34,38^.

### Genetic constructs and expression of IDRs in yeast

Open reading frames (ORFs) coding for each IDR were gene synthesized (GenScript, **Supplementary Table 2**) and fused between the coding sequences of the FRET pair fluorescent proteins (mCerulean3 and Citrine) of the pDRFLIP38-AtLEA4-5 plasmid (Addgene Plasmid #178189)^57^. Each construct was transformed into *Saccharomyces cerevisiae* (protease-deficient strain BJ5465) using lithium acetate transformation method^57^. Transformed colonies were selected in plates containing 6.8 g/L YNB media (Sigma-Aldrich) supplemented with 5 g/L glucose and 1.92 g/L synthetic drop-out medium without uracil (Sigma-Aldrich). Positive clones were genotyped by colony PCR. Correct expression of each construct was assessed by measuring fluorescence emission spectra following independent excitation of mCerulean3 or Citrine. Twelve constructs (IDR007, IDR016, IDR035, IDR076, IDR080, IDR087, IDR091, IDR092, IDR093, IDR098, IDR100, and IDR103) were excluded due to growth defects, undetectable fluorescence, or absence of acceptor fluorescence.

For generation of CAHS LR variants, each construct was either gene synthesized (GenScript) (V1, V2, and V3 *κ* variants) or PCR amplified (HeCAHS8 LR, FL_HeCAHS8, 2X_LR, FL_Proline, NLN, and CLC) from plasmids generated in a previous work^32^. DNA fragments were cloned using the Gibson Assembly cloning method (NEB) in between the coding sequences of the FRET pair using the pDRFLIP38-AtLEA4-5 plasmid digested with SacI and BglII. For Syn1, the coding sequence was gene synthesized (GenScript) and cloned in the same way as the other variants. Each construct was transformed into *Saccharomyces cerevisiae* (protease-deficient strain BJ5465) as described before.

### In vivo FRET measurements

Yeast growth and FRET measurements were performed as described previously^58^. Briefly, 2 mL of liquid YNB media (6.8 g/L) (Sigma-Aldrich) supplemented with 5 g/L glucose and 1.92 g/L synthetic drop-out medium without uracil (Sigma-Aldrich) were inoculated with each construct and grown at 30 °C for at least 12 hours until saturation (OD600 ∼ 1.0 - 2.0). Cells were centrifuged and washed twice with 50 mM MES (pH 6) and resuspended in 2 mL of the same buffer. 50 μL of the cell suspension were loaded into individual wells of a 96-well black F-bottom clear microplate (Greiner). 150 μL of treatment solution (0 M, 0.2 M, 0.4 M, 0.6 M, 0.8 M, 1.0 M, 1.5 M NaCl) were added to the cell suspension and mixed well by pipetting up and down. Fluorescence measurements were performed immediately after addition of the treatment. Fluorescence readings were acquired using a CLARIOstar (BMG Labtech) microplate reader with the following optical settings: Excitation wavelength 433 nm; excitation bandwidth 10 nm; emission wavelength 460 - 550 nm; step width 1 nm; emission bandwidth 10 nm; focal height 5 mm (obtained by autofocus). The wavelength used for gain calculation was 490 nm. For generation of fluorescence emission spectra, each fluorescence emission value was normalized to the isosbestic point of mCerulean3-Citrine FRET pair (515 nm). For calculations of FRET ratio, DxAm values (excitation = 433 nm, emission = 525 nm) were divided by DxDm values (excitation = 433 nm, emission = 475 nm) and normalized to the mean DxAm/DxDm of the non-stress condition (0 M NaCl). Experiments were repeated three times, each with three technical replicates for every condition.

### Calculation of parameters and ensemble predictions of IDRs

Sequence encoded parameters for each IDR of the library, CAHS LR variants, and the human IDRome were calculated with localCIDER^58^. The parameters calculated were the fraction of charged residues (FCR), charge patterning (kappa, *κ*), sequence charge decoration (SCD), absolute net charge per residue (|NCPR|), hydropathy, sequence hydropathy decoration (SHD), isoelectric point (pI), fraction of aromatic residues, fraction of aliphatic residues, and fraction of polar residues. NCPR plots were also generated with localCIDER^58^. Predicted ensemble properties including the radius of gyration (*R*g), end-to-end distance (*R*e), asphericity, polymer scaling exponent (ν), and prefactor (ρ_0_), were calculated with the IDR analysis tool of ALBATROSS^40^. The human IDRome was obtained with the IDRome constructor tool of ALBATROSS^40^.

### Binomial logistic regression analysis

A binomial logistic regression model was fitted using the hypersensitive and insensitive IDR groups. The response variable (Target) was encoded as a binary factor (hypersensitive = 1; insensitive = 0). Four predictor variables were included in the model: charge patterning (kappa, *κ*), fraction of charged residues (FCR), predicted end-to-end distance (*R*e), and absolute net charge per residue (|NCPR|). To ensure comparability of odds ratios (OR) and avoid distortion due to scale differences among predictors, *κ*, FCR, and |NCPR| were linearly rescaled to a 0 - 100 range prior to model fitting (denoted as r*κ*, rFCR, and r|NCPR|). This rescaling does not alter statistical significance or model fit but allows odds ratios to be interpreted per standardized relative unit increase. *R*e was retained in its original scale. Model fitting was performed using a generalized linear model with binomial error distribution and logit link function. Multicollinearity among predictors was assessed using variance inflation factors (VIF), and only variables meeting the absence-of-multicollinearity assumption were retained. Model selection was guided by Akaike’s Information Criterion (AIC) using backward stepwise elimination. Odds ratios (ORs) were obtained by exponentiating regression coefficients. Model performance was evaluated using overall classification accuracy, sensitivity (true positive rate), specificity (true negative rate), and AIC. Predicted probabilities were calculated for each IDR using the fitted model. Receiver operating characteristic (ROC) analysis was performed to visualize the trade-off between sensitivity and specificity across thresholds. A probability threshold of 0.5 was selected for classification, and confusion matrices were generated accordingly. To assess model generalizability across the continuum of ensemble responses, the fitted model was applied to IDRs from the intermediate (average-response) group. This group was divided into two subsets based on ΔFRET quartiles: Q1–Q2 (25th–50th percentile) and Q2–Q3 (50th–75th percentile). Predicted probabilities were computed without refitting the model. Using the same 0.5 probability threshold, classification performance was evaluated separately for each subset by calculating true positive and true negative rates. All analyses were performed in R using base statistical functions and standard packages for regression diagnostics and ROC analysis.

### Sliding window sensitivity analysis

A sliding window approach was applied to calculate the probability to be classified as hypersensitive for IDRs larger than 200 residues. A window of 100 amino acid residues was defined as the analytical unit, consistent with the training range of the binomial logistic regression model (50-200 residues). The window was advanced along each sequence in steps of 10 residues (stride = 10), yielding a positional resolution of one score per 10 residues. For each window position, the central residue was used as the positional reference to which the predicted score was assigned, such that the first and last 50 residues of each sequence fall outside the scored region due to the structural blind zone inherent to center-based assignment. For each window, the corresponding sequence was extracted and evaluated with the model, which outputs a probability P(sensitivity) ∈ [0,1], representing the likelihood that the region is classified as hypersensitive to hyperosmotic stress. Windows for which the model could not produce a valid prediction were assigned a missing value (NaN) and excluded from subsequent analysis. This included sequences where kappa value is either 1 or -1 because the sequence has only positive or negative residues or has no charged residues. The resulting positional probability profiles were smoothed using a Savitzky-Golay filter (window length = 11, polynomial order = 3) to reduce local noise while preserving the shape of genuine peaks. Regions where the smoothed probability equaled or exceeded a threshold of 0.5 were defined as putative environment-sensitive segments. The boundaries and lengths of these segments were determined directly from the smoothed profile to ensure consistency with the visual representation of the data. All analyses were performed in Python 3, using NumPy for numerical operations, SciPy for signal smoothing, and Matplotlib for visualization.

### Statistical analysis

Data were analyzed using one-way ANOVA followed by Tukey’s multiple comparisons test for experiments involving more than two groups, after verifying normality and homogeneity of variance. For comparisons between two groups, two-sided unpaired Student’s t-tests were performed when assumptions were met; otherwise, two-sided Mann-Whitney U tests were used. Statistical significance was defined as \**P* < 0.05, \*\**P* < 0.01, \*\*\**P* < 0.001, \*\*\*\**P* < 0.0001).

## ACKNOWLEDGEMENTS

Support for this project came from the Secretaría de Ciencia, Humanidades, Tecnología e Innovación (SECIHTI), Ciencia de Frontera project 252952; the Programa de Apoyo a Proyectos de Investigación e Innovación Tecnológica, Dirección General de Asuntos del Personal Académico, Universidad Nacional Autónoma de México (UNAM-PAPIIT), projects IA209920 and IA203422; and Programa de Apoyo a la Investigación y el Posgrado, Facultad de Química, Universidad Nacional Autónoma de México, grant 5000-9182. We acknowledge the support from the Global Perspectives Grant program from the College of Agriculture, Life Sciences, and Natural Resources of the University of Wyoming. C.E.-T. (CVU 1083636) and C.A.P.-D. (CVU 1269643) acknowledge SECIHTI for their MSc fellowships. E.E.R. (CVU 104360) acknowledge SECIHTI for his Postdoctoral fellowship. T.C.B. is part of the Water and Life Interface Institute (WALII), supported by NSF grant 2213983. NSF IntBIO grant 2128069 to T.C.B. also helped support this work.

