## Extended Data Figures for "A molecular grammar for environmental sensitivity in intrinsically disordered protein regions"

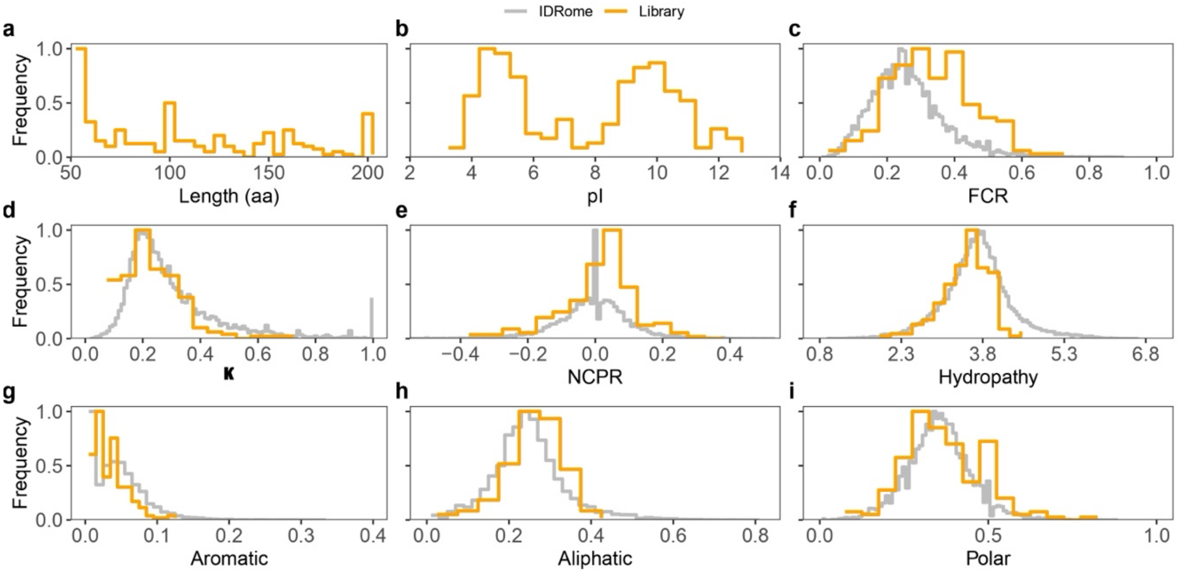


**Extended Data Fig. 1. Distributions of sequence-encoded properties of 188 naturally-occurring IDRs.** **a**, Length distribution of IDRs in the library. **b**, Distribution of isoelectric points (pI) of IDRs in the library. **c**, Distribution of fraction of charged residues (FCR) of IDRs in the library (orange) compared to the human IDRome (gray). **d**, Distribution of kappa values ($\kappa$) of IDRs in the library (orange) compared to the human IDRome (gray). **e**, Distribution of net charge per residue (NCPR) of IDRs in the library (orange) compared to the human IDRome (gray). **f**, Distribution of Hydropathy of IDRs in the library (orange) compared to the human IDRome (gray). **g**, Distribution of fraction of aromatic residues (Aromatic) of IDRs in the library (orange) compared to the human IDRome (gray). **h**, Distribution of fraction of aliphatic residues (Aliphatic) of IDRs in the library (orange) compared to the human IDRome (gray). **i**, Distribution of fraction of polar residues (Polar) of IDRs in the library (orange) compared to the human IDRome (gray).

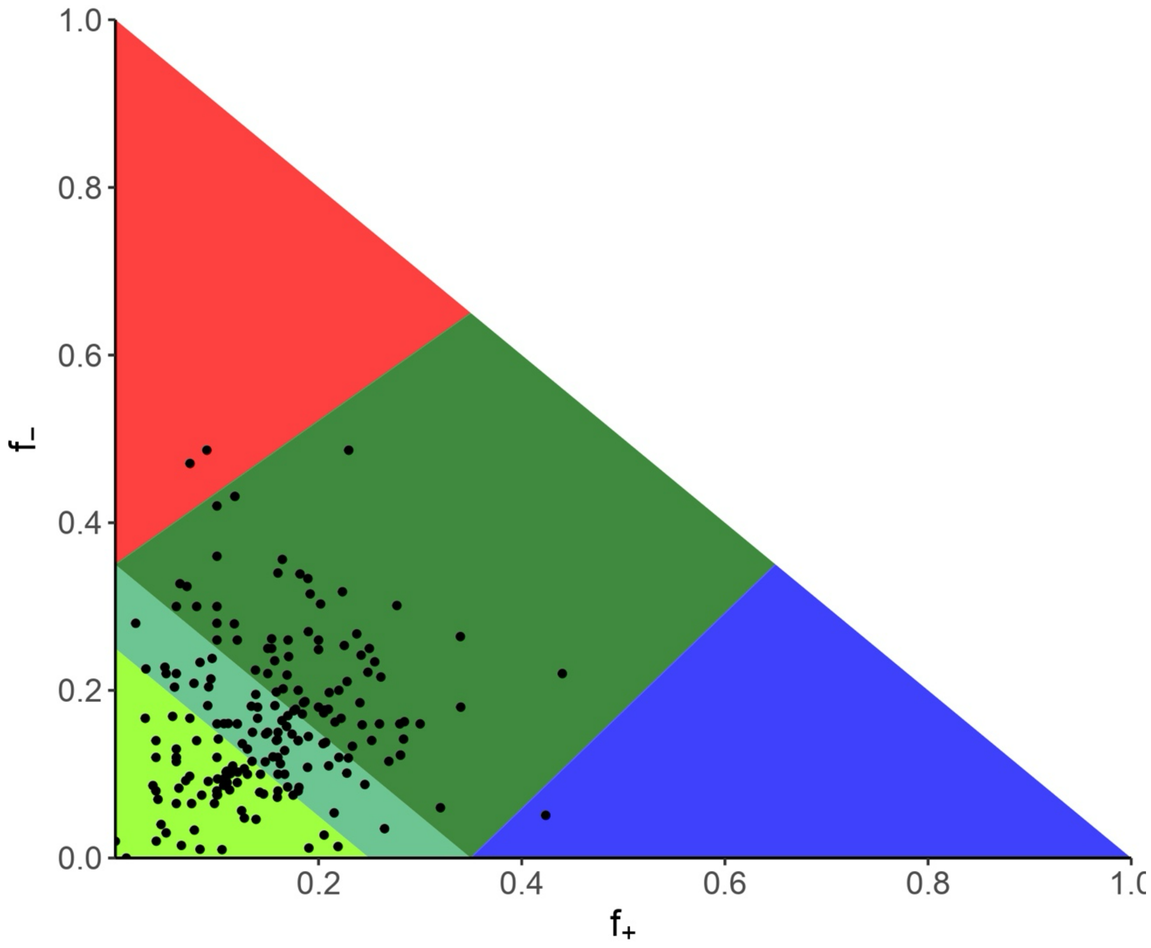


**Extended Data Fig. 2. Das-Pappu diagram-of-states for 188 naturally-occurring IDRs.** Each dot represents an individual IDR. $f_+$ is the fraction of positively charged residues. $f_-$ is the fraction of negatively charged residues. R1 (green yellow) weak polyampholytes and polyelectrolytes (globules and tadpoles). R2 (light green): boundary region. R3 (dark green): strong polyampholytes (coils, hairpins and chimeras). R4 (red): negatively charged strong polyampholytes (swollen coils). R5 (blue): positively charged strong polyampholytes (swollen coils).

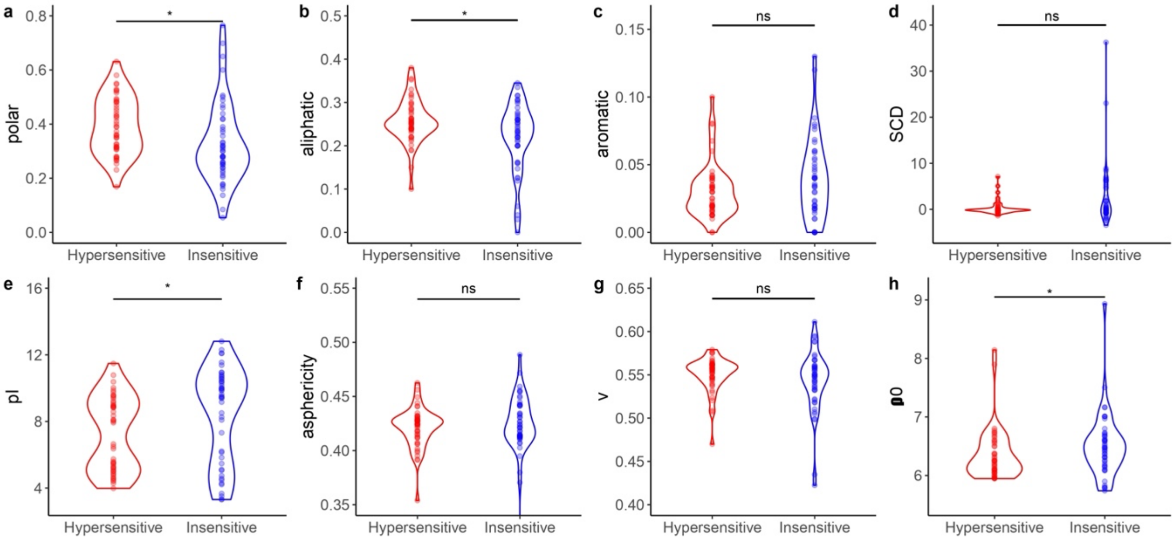


**Extended Data Fig. 3. Sequence features and predicted IDR ensemble properties of hypersensitive and insensitive IDRs. a-e**, Sequence features of IDRs belonging to the hypersensitive (red) or insensitive (blue) groups. **a**, fraction of polar residues (polar); **b**, fraction of aliphatic residues (aliphatic); **c**, fraction of aromatic residues (aromatic); **d**, sequence charge decoration (SCD); **e**, isoelectric point (pI). **f-h**, predicted ensemble properties for IDRs of the hypersensitive (red) or insensitive (blue) groups. **f**, predicted asphericity (asphericity); **g**, predicted polymer scaling exponent (v). **h**, predicted prefactor (ρ0). Each point represents the corresponding value for an IDR. Color intensity indicates the representation of values within the same range. Mann-Whitney U test *$P < 0.05$, **$P < 0.01$, ***$P < 0.001$, ****$P < 0.0001$.

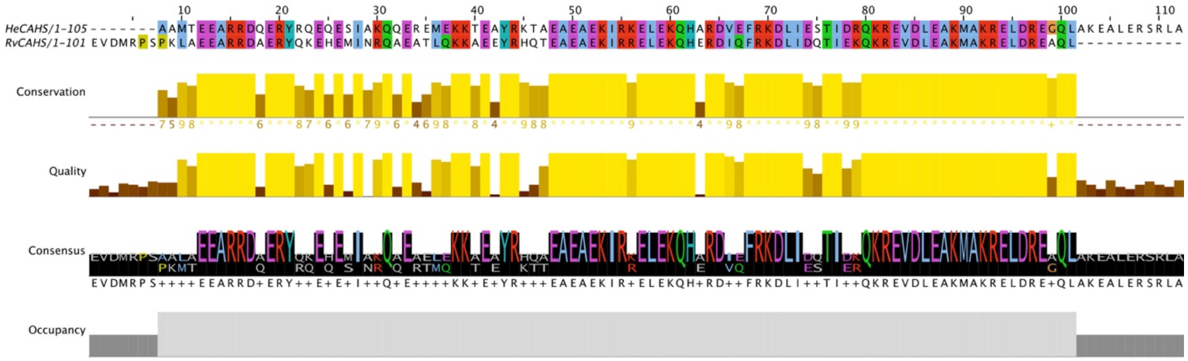


**Extended Data Fig. 4. Sequence alignment of HeCAHS8 LR and RvCAHS LR. a-e**, ClustalW multiple sequence alignment of *Hypsibius exemplaris* HeCAHS8 LR with *Ramazzottius varieornatus* RvCAHS1 LR. Sequences were aligned with T-Coffee and visualized with Jalview.

>V1
ILLTEEIEETDYQEDEQESIAKQQEREMEKEKTAYREKTAAAEKAEKLEKQHAREMKEFRDLADSQIDKQEKE
AKLEARMAKERELDRGQVRARAVRSRRLRARR

>V2
EALIEEQEEIEEEEDQEYQEQESEIAQQEEMEKEKTAYREKTAAAEKAEKLEKQHARDVKDFRDAMESTIDR
QDRVKLARKMAKRRRLRGTKLAKRALRSRLARR

>V3
AAMTEEDQEEQESQEEMEEAEEAEEEDVEDLESDEVDLEAEDEAEELLGAALAQIHATIQAFYRRKRRAKQ
RKTYRKTKIRKQRKRKIRQKRKMKRLRAKRSRLA

**Extended Data Fig. 5. Amino acid sequences of HeCAHS8 LR $\kappa$ variants.** Positively charged residues in blue, negatively charged residues in red, polar residues in green, aliphatic residues in black, aromatic residues in yellow.

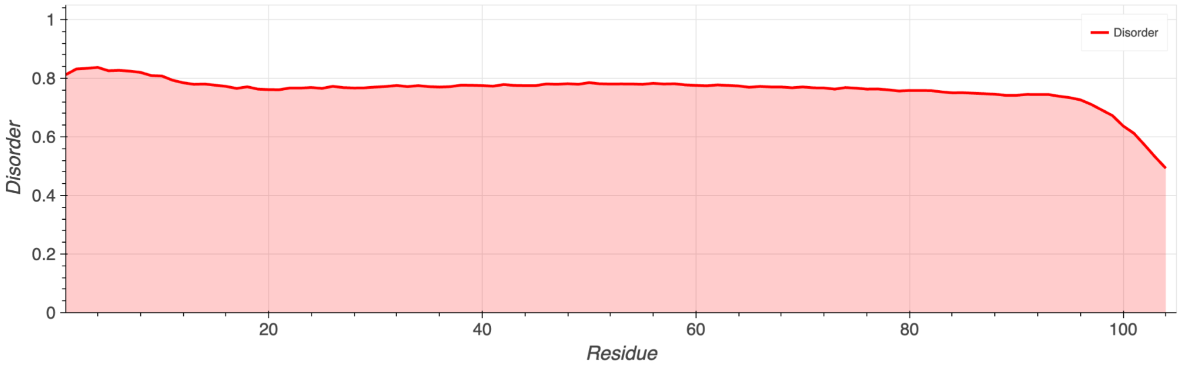


**Extended Data Fig. 6. Syn1 disorder prediction.** Disorder prediction across the sequence of Syn1 with Metapredict. The red shaded region indicates reliable disorder prediction.

>FL_HeCAHS8
MSGRNVESHMERNEKVVVNNSGHADVKKQQQQVEHTEFTHTEVKAPLIHP
APPIISTGAAGLAEEIVGQGFTASAARISGGTAEVHLQPSAAMTEEARRD
QERYRQEQESIAKQQEREMEKKTEAYRKTAEAEAEKIRKELEKQHARDVE
FRKDLIESTIDRQKREVDLEAKMAKRELDREGQLAKEALERSRLATNVEV
NFDSAAGHTVSGGTTVSTSDKMEIKRN

>LR
AAMTEEARRDQERYRQEQESIAKQQEREMEKKTEAYRKTAEAEAEKIRKE
LEKQHARDVEFRKDLIESTIDRQKREVDLEAKMAKRELDREGQLAKEALE
RSRLA

>2X_LR
MSGRNVESHMERNEKVVVNNSGHADVKKQQQQVEHTEFTHTEVKAPLIHP
APPIISTGAAGLAEEIVGQGFTASAARISGGTAEVHLQPSAAMTEEARRD
QERYRQEQESIAKQQEREMEKKTEAYRKTAEAEAEKIRKELEKQHARDVE
FRKDLIESTIDRQKREVDLEAKMAKRELDREGQLAKEALERSRLATNMTE
EARRDQERYRQEQESIAKQQEREMEKKTEAYRKTAEAEAEKIRKELEKQH
ARDVEFRKDLIESTIDRQKREVDLEAKMAKRELDREGQLAKEALERSRLA
TNVEVNFDSAAGHTVSGGTTVSTSDKMEIKRN

>FL_Proline
MSGRNVESHMERNEKVVVNNSGHADVKKQQQQVEHTEFTHTEVKAPLIHP
APPIISTGAAGLAEEIVGQGFTASAARISGGTAEVHLQPSAAMTEEARRD
QERYRQEQESIAKQQEREMEKKTEAYRKTAEAEAEKIRKELEKQHARDVE
FRKDLIESTIDRQKREVDLEAKMAKRELDREGQLAKEALERSRLATNVPV
NFPSAAGHTVSGGTTPSTSDKMPIKRN

>NLN
MSGRNVESHMERNEKVVVNNSGHADVKKQQQQVEHTEFTHTEVKAPLIHP
APPIISTGAAGLAEEIVGQGFTASAARISGGTAEVHLQPSAAMTEEARRD
QERYRQEQESIAKQQEREMEKKTEAYRKTAEAEAEKIRKELEKQHARDVE
FRKDLIESTIDRQKREVDLEAKMAKRELDREGQLAKEALERSRLAMSGRN
VESHMERNEKVVVNNSGHADVKKQQQQVEHTEFTHTEVKAPLIHPAPPII
STGAAGLAEEIVGQGFTASAARISGGTAEVHLQPS

>CLC
MTNVEVNFDSAAGHTVSGGTTVSTSDKMEIKRNAAMTEEARRDQERYRQE
QESIAKQQEREMEKKTEAYRKTAEAEAEKIRKELEKQHARDVEFRKDLIE
STIDRQKREVDLEAKMAKRELDREGQLAKEALERSRLATNVEVNFDSAAG
HTVSGGTTVSTSDKMEIKRN

**Extended Data Fig. 7. Amino acid sequences of HeCAHS8 variants.** Positively charged residues in blue, negatively charged residues in red, polar residues in green, aliphatic residues in black, aromatic residues in yellow, prolines in magenta.
