## Extended File 1 for "A molecular grammar for environmental sensitivity in intrinsically disordered protein regions"

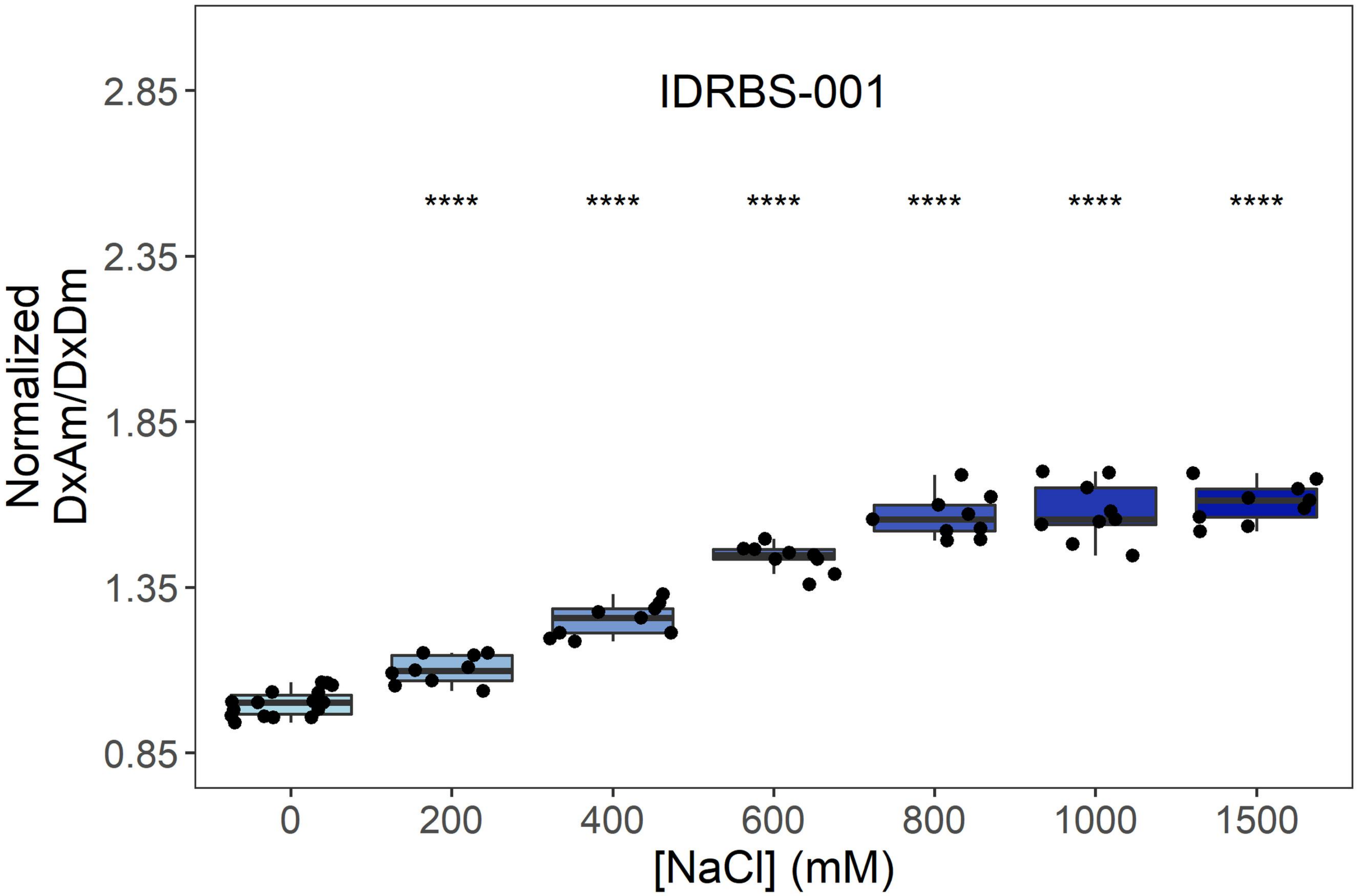
IDRBS-001
****
****
****
****
****
****
Normalized DxAm/DxDm
2.85
2.35
1.85
1.35
0.85
0
200
400
600
800
1000
1500
[NaCl] (mM)

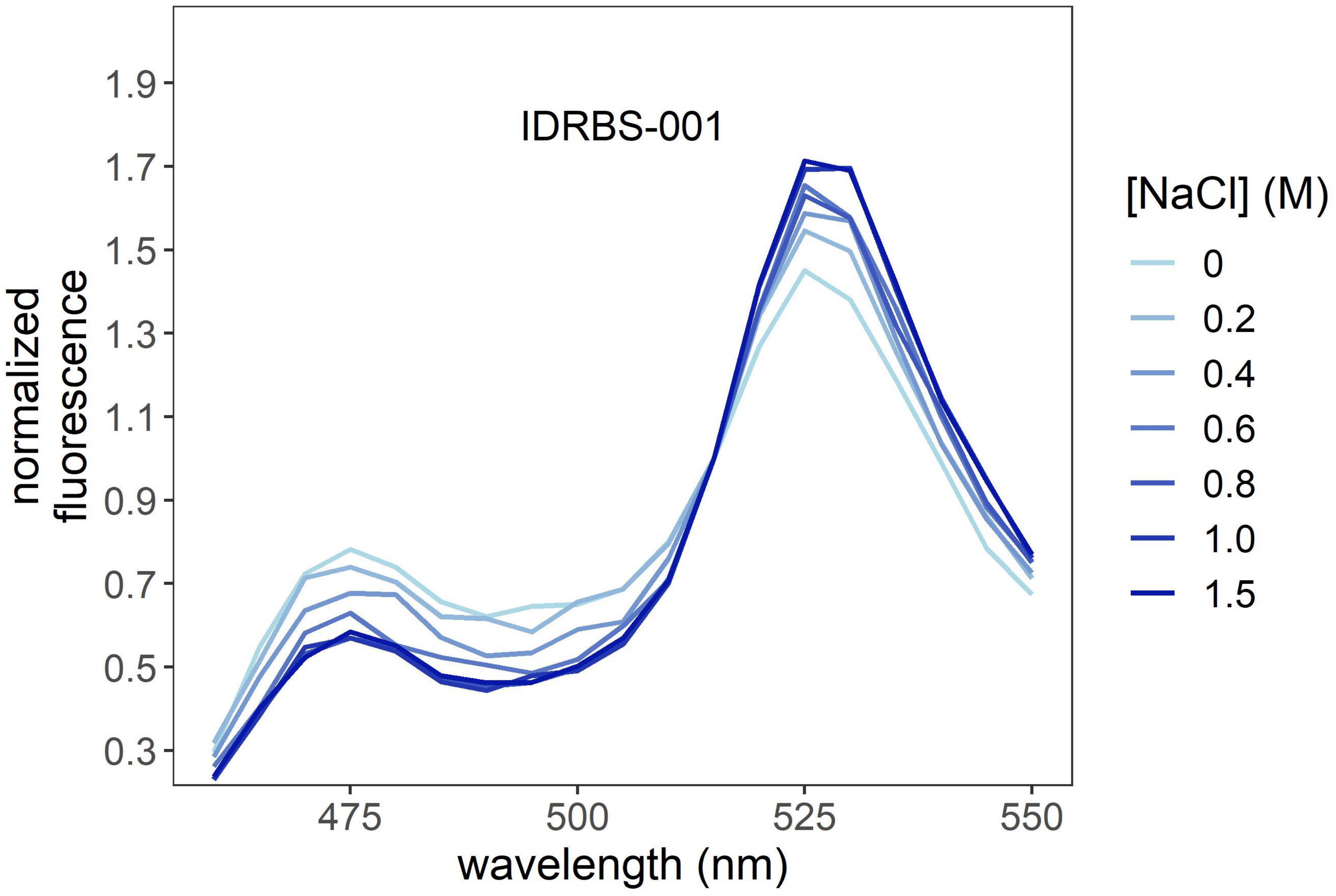
IDRBS-001
normalized fluorescence
1.9
1.7
1.5
1.3
1.1
0.9
0.7
0.5
0.3
475
500
525
550
wavelength (nm)
[NaCl] (M)
0
0.2
0.4
0.6
0.8
1.0
1.5

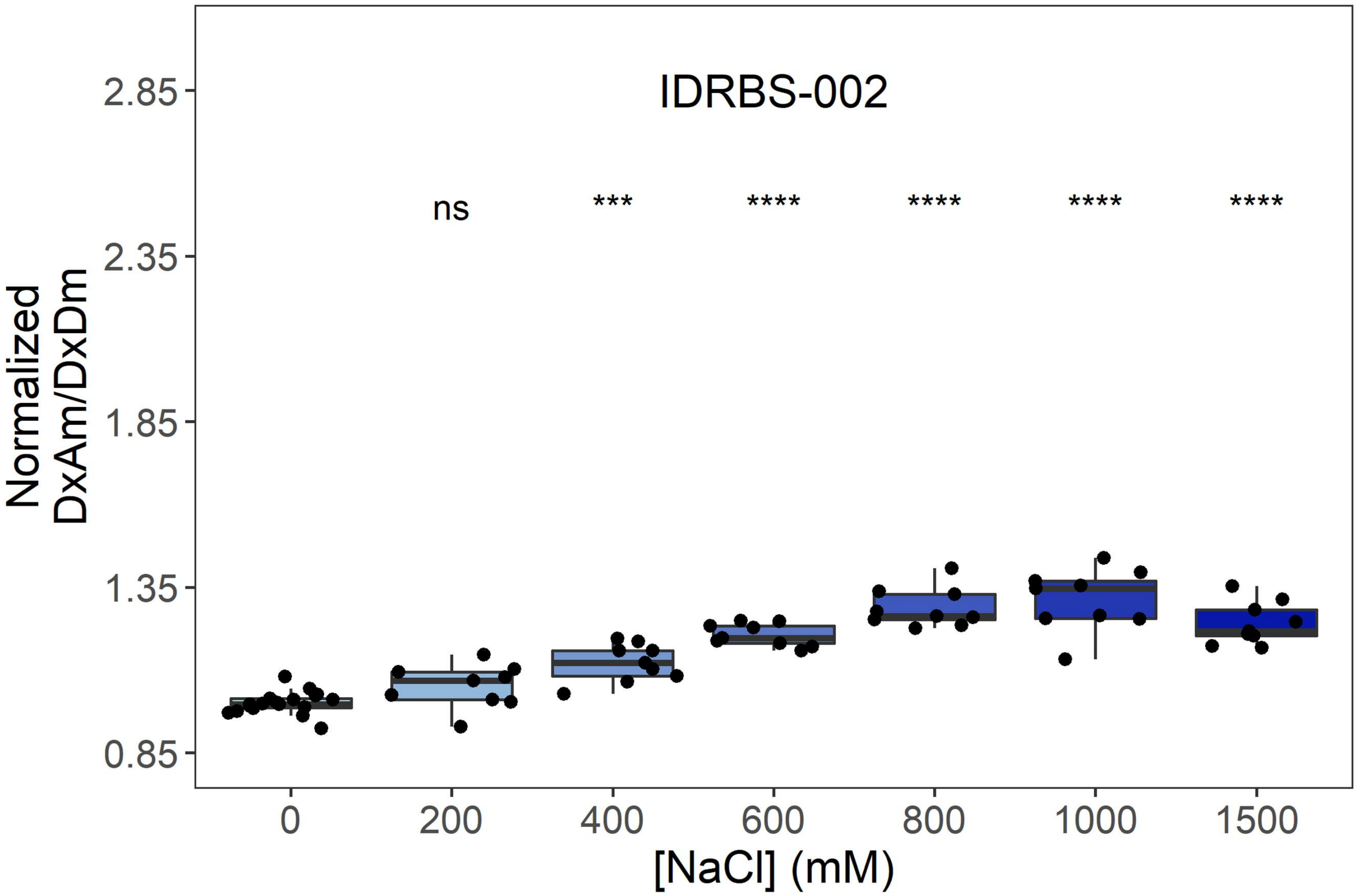

IDRBS-002
ns
***
****
****
****
****
2.85
2.35
1.85
1.35
0.85
Normalized
DxAm/DxDm
0
200
400
600
800
1000
1500
[NaCl] (mM)


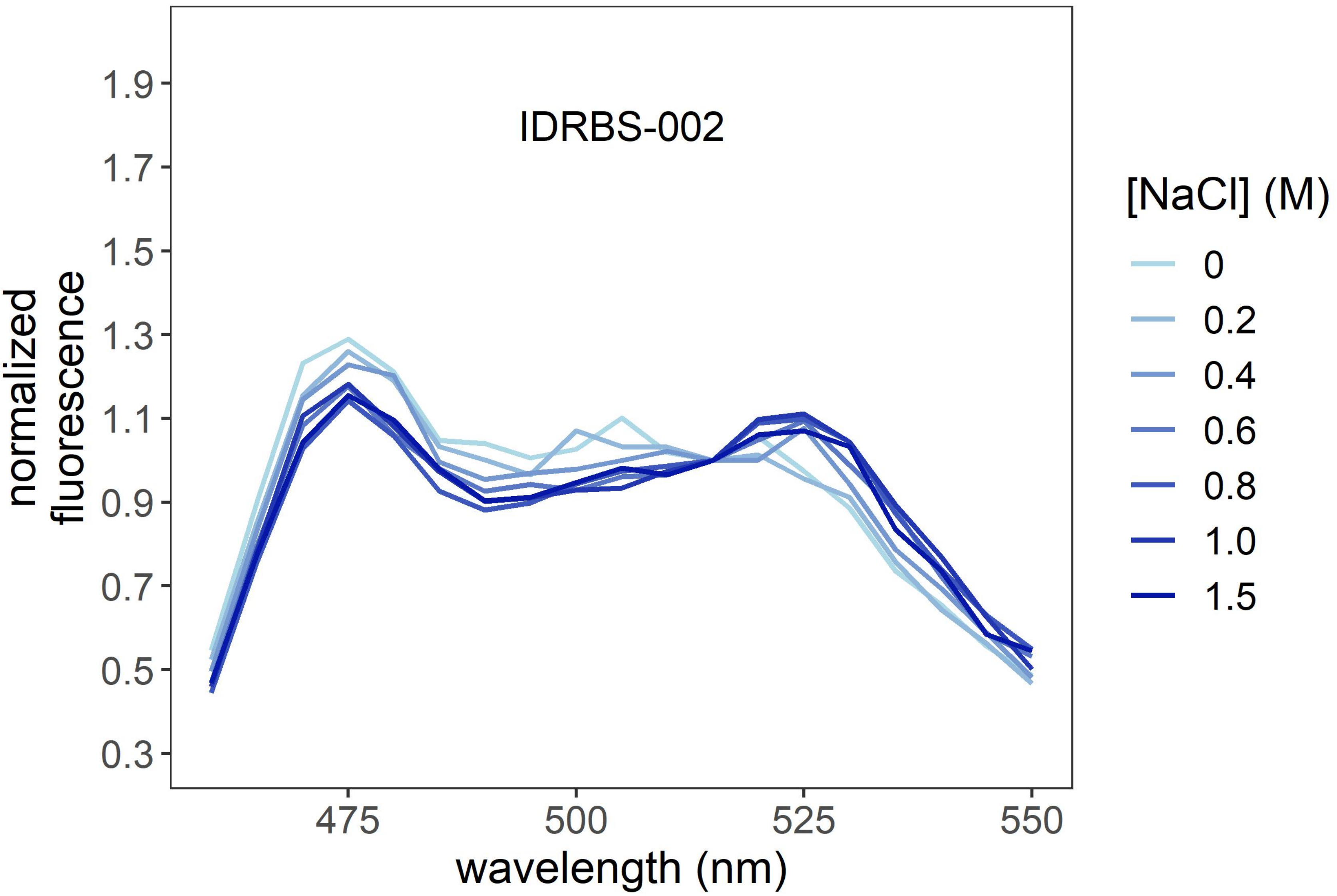

IDRBS-002
1.9
1.7
1.5
1.3
1.1
0.9
0.7
0.5
0.3
normalized
fluorescence
475
500
525
550
wavelength (nm)
[NaCl] (M)
0
0.2
0.4
0.6
0.8
1.0
1.5

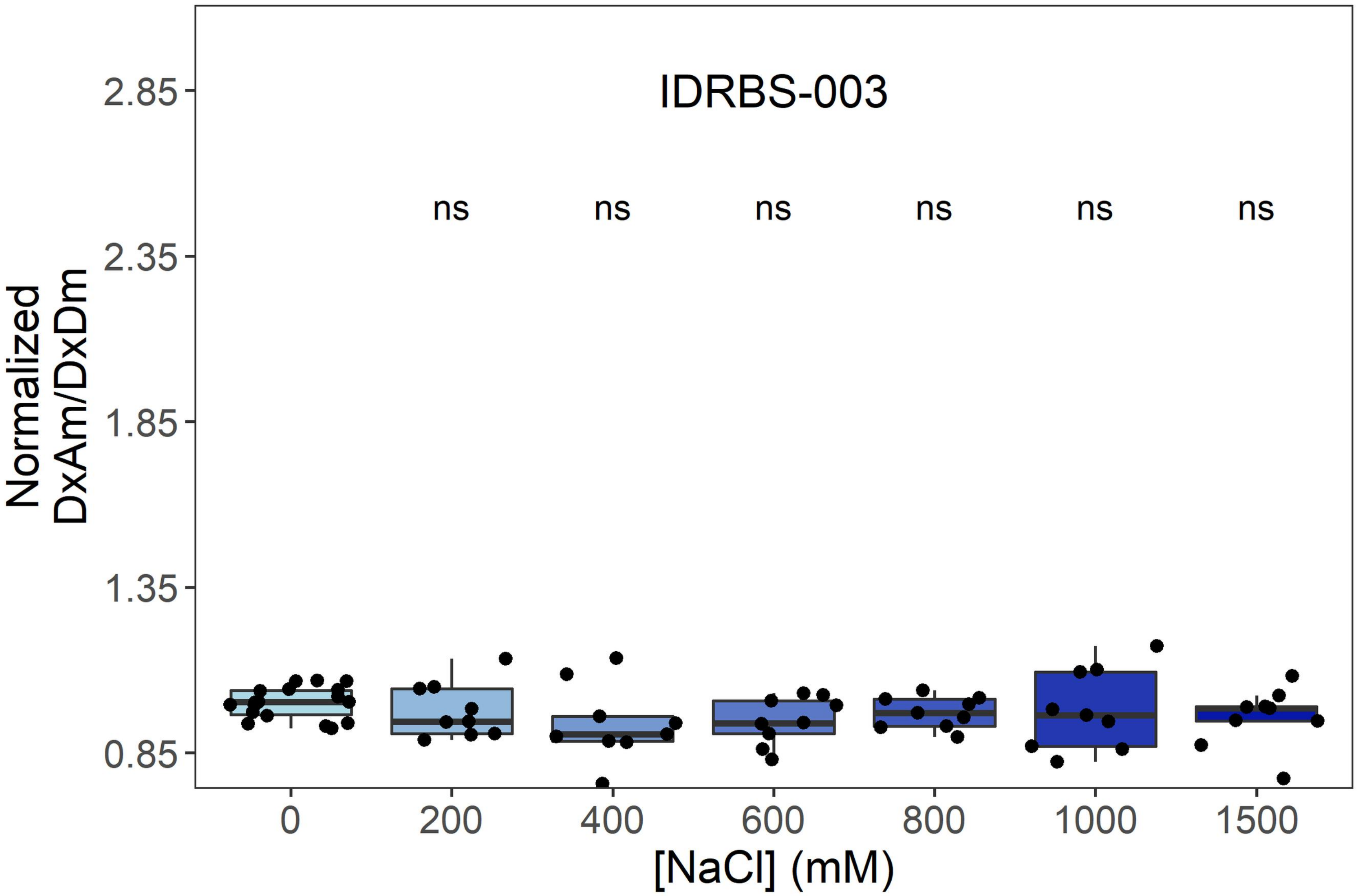

IDRBS-003
ns
ns
ns
ns
ns
ns
2.85
2.35
1.85
1.35
0.85
Normalized DxAm/DxDm
0
200
400
600
800
1000
1500
[NaCl] (mM)


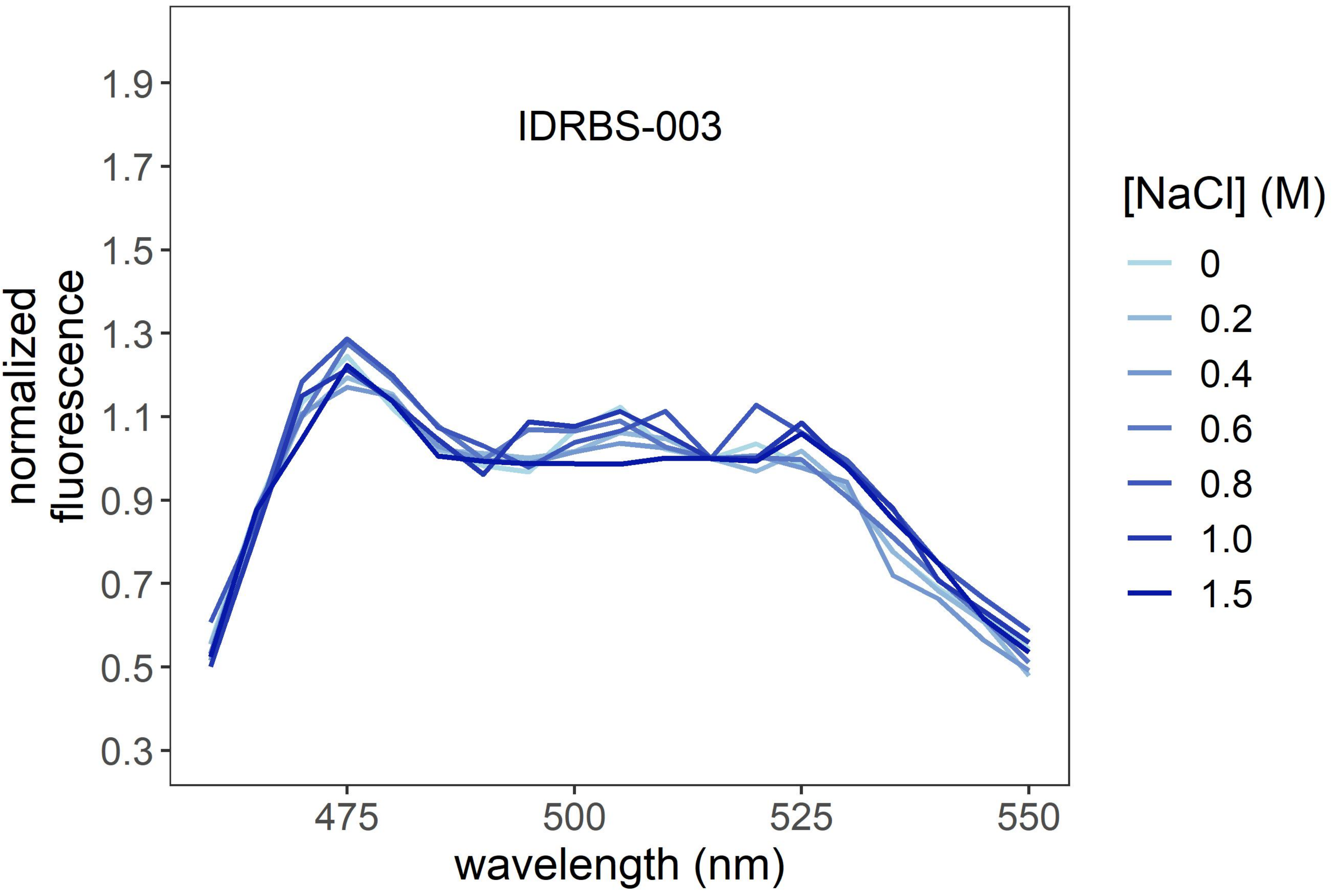

IDRBS-003
1.9
1.7
1.5
1.3
1.1
0.9
0.7
0.5
0.3
normalized fluorescence
475
500
525
550
wavelength (nm)
[NaCl] (M)
0
0.2
0.4
0.6
0.8
1.0
1.5

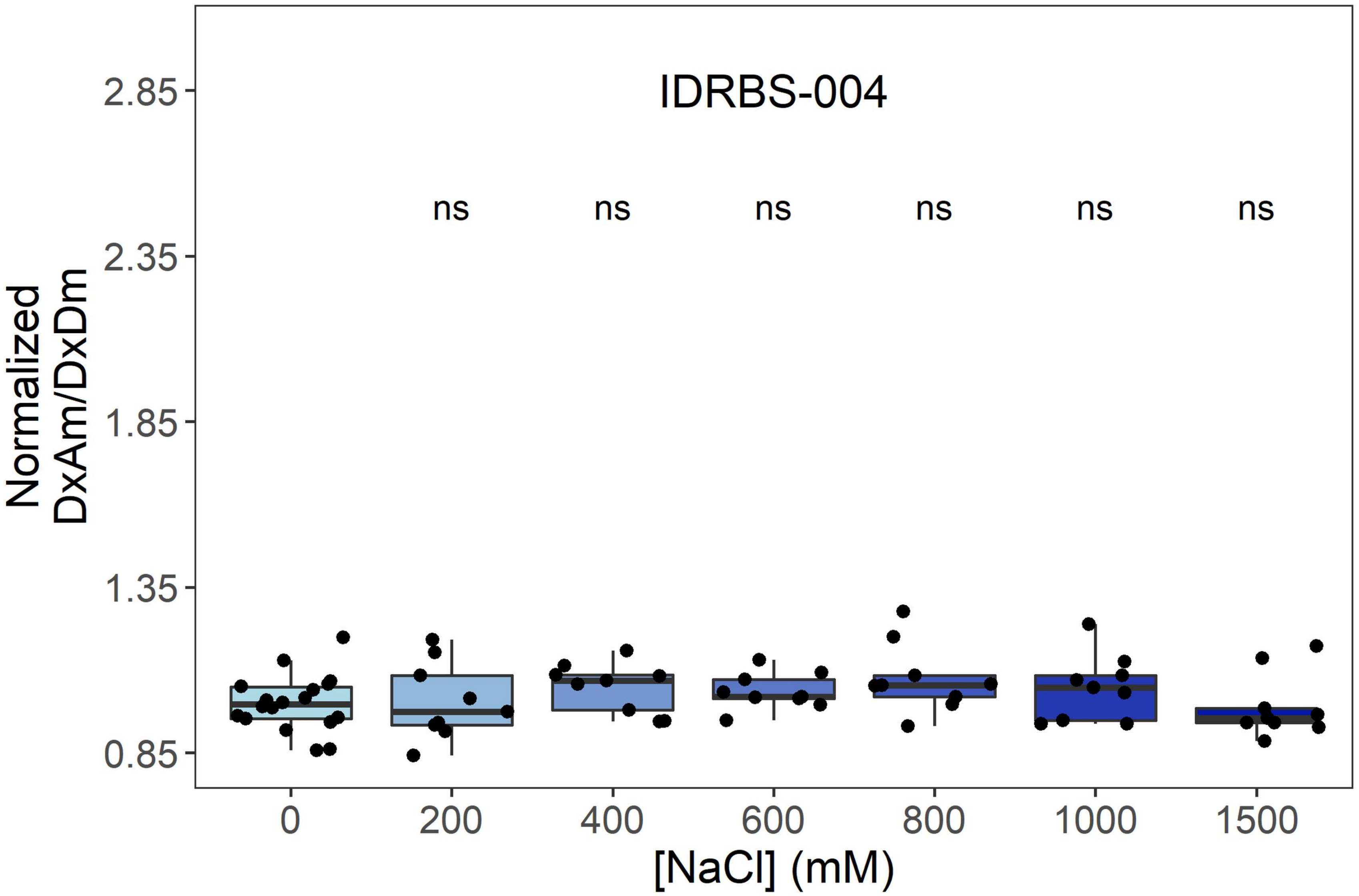
IDRBS-004
ns
ns
ns
ns
ns
ns
Normalized DxAm/DxDm
2.85
2.35
1.85
1.35
0.85
0
200
400
600
800
1000
1500
[NaCl] (mM)

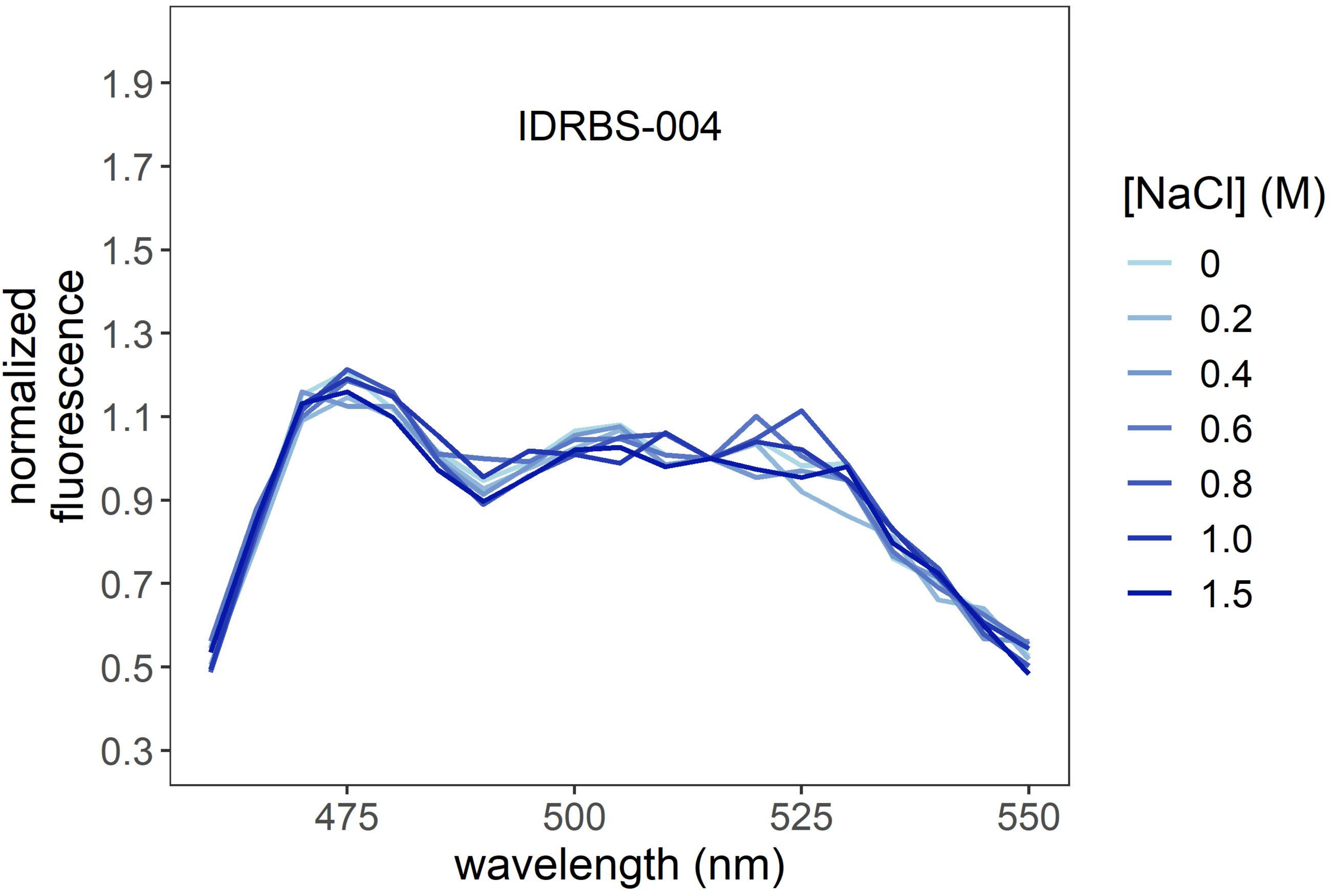
IDRBS-004
normalized fluorescence
1.9
1.7
1.5
1.3
1.1
0.9
0.7
0.5
0.3
475
500
525
550
wavelength (nm)
[NaCl] (M)
0
0.2
0.4
0.6
0.8
1.0
1.5

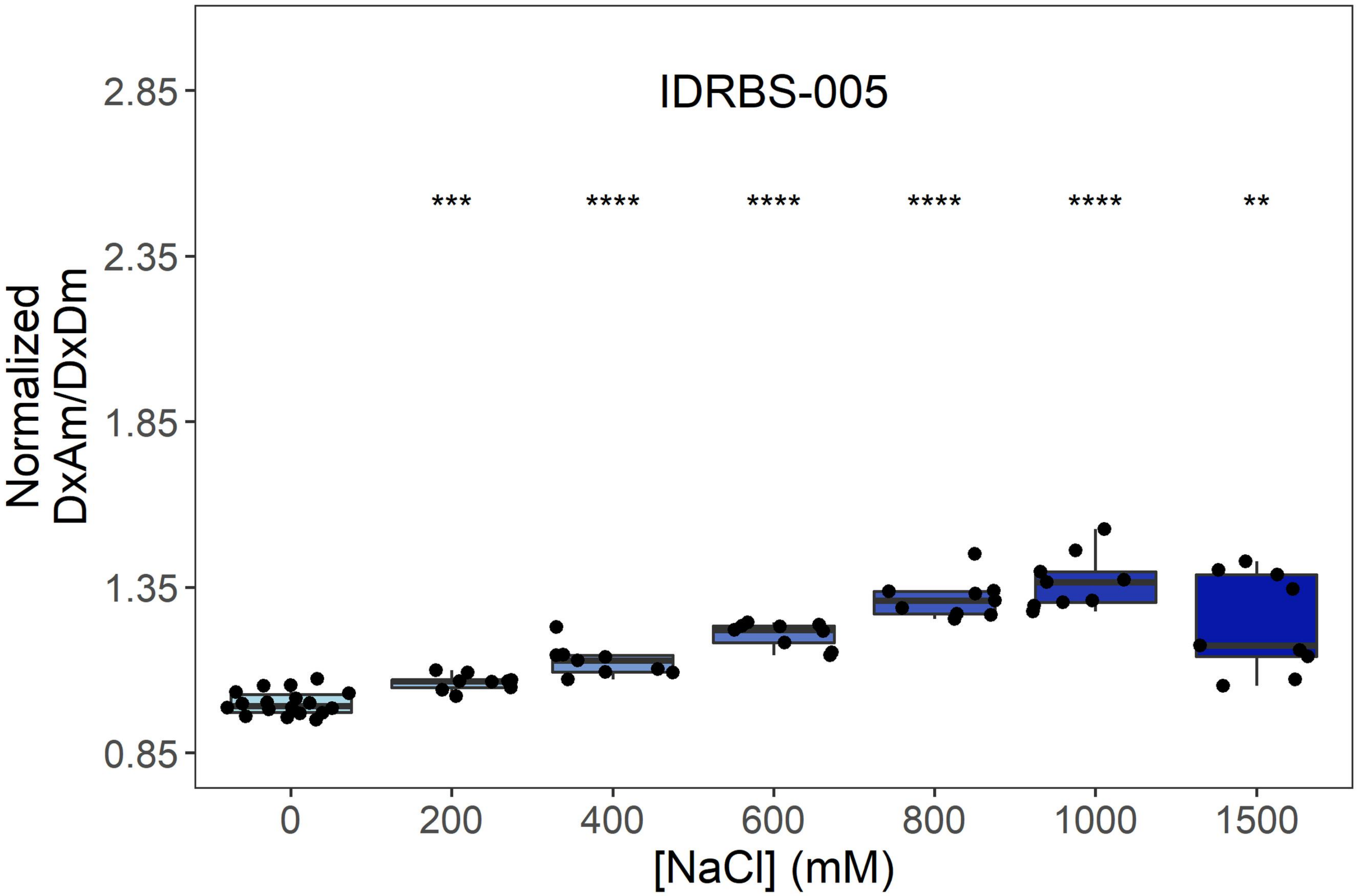
IDRBS-005
Normalized DxAm/DxDm
2.85
2.35
1.85
1.35
0.85
***
****
****
****
****
**
0
200
400
600
800
1000
1500
[NaCl] (mM)

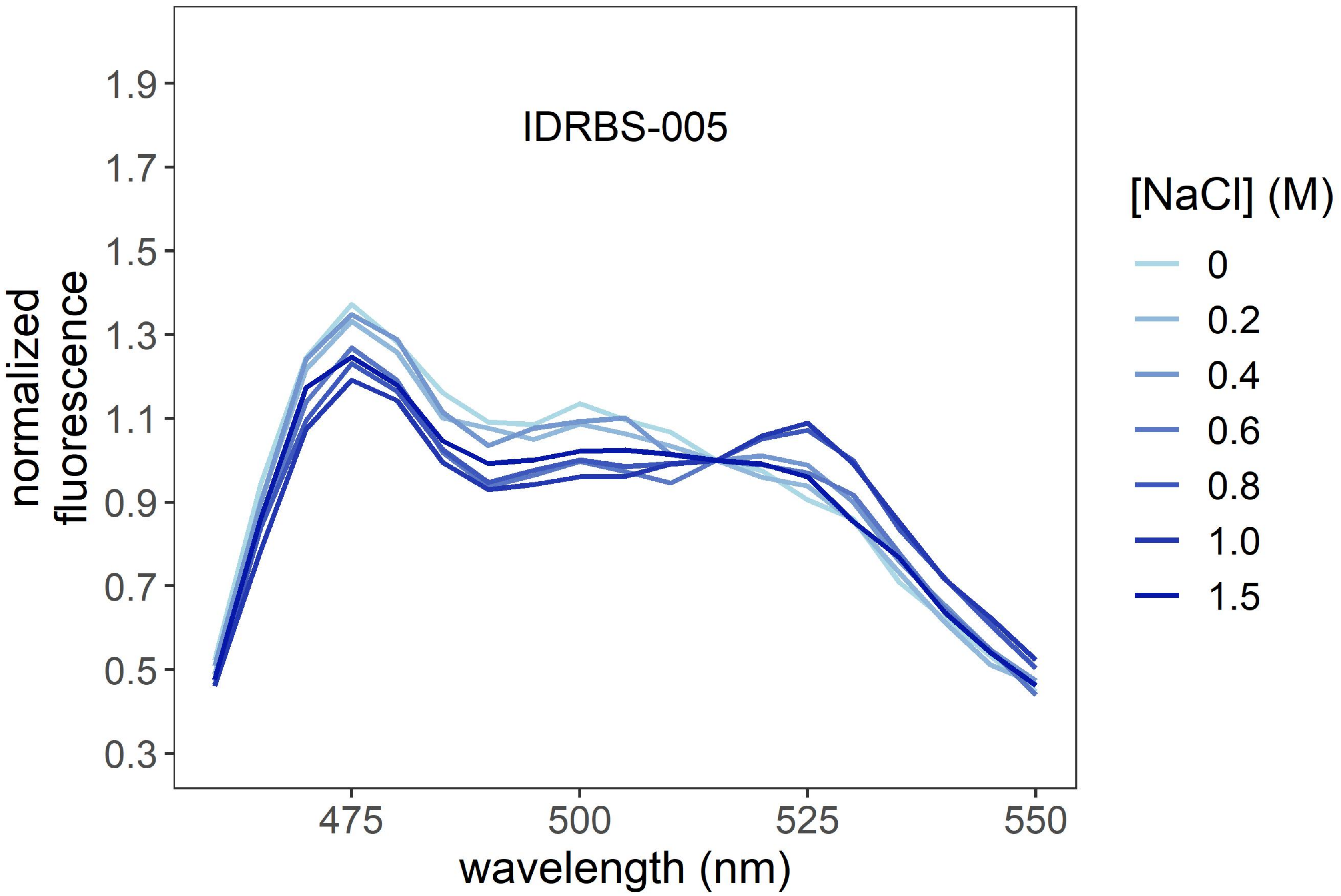
IDRBS-005
normalized fluorescence
1.9
1.7
1.5
1.3
1.1
0.9
0.7
0.5
0.3
475
500
525
550
wavelength (nm)
[NaCl] (M)
0
0.2
0.4
0.6
0.8
1.0
1.5

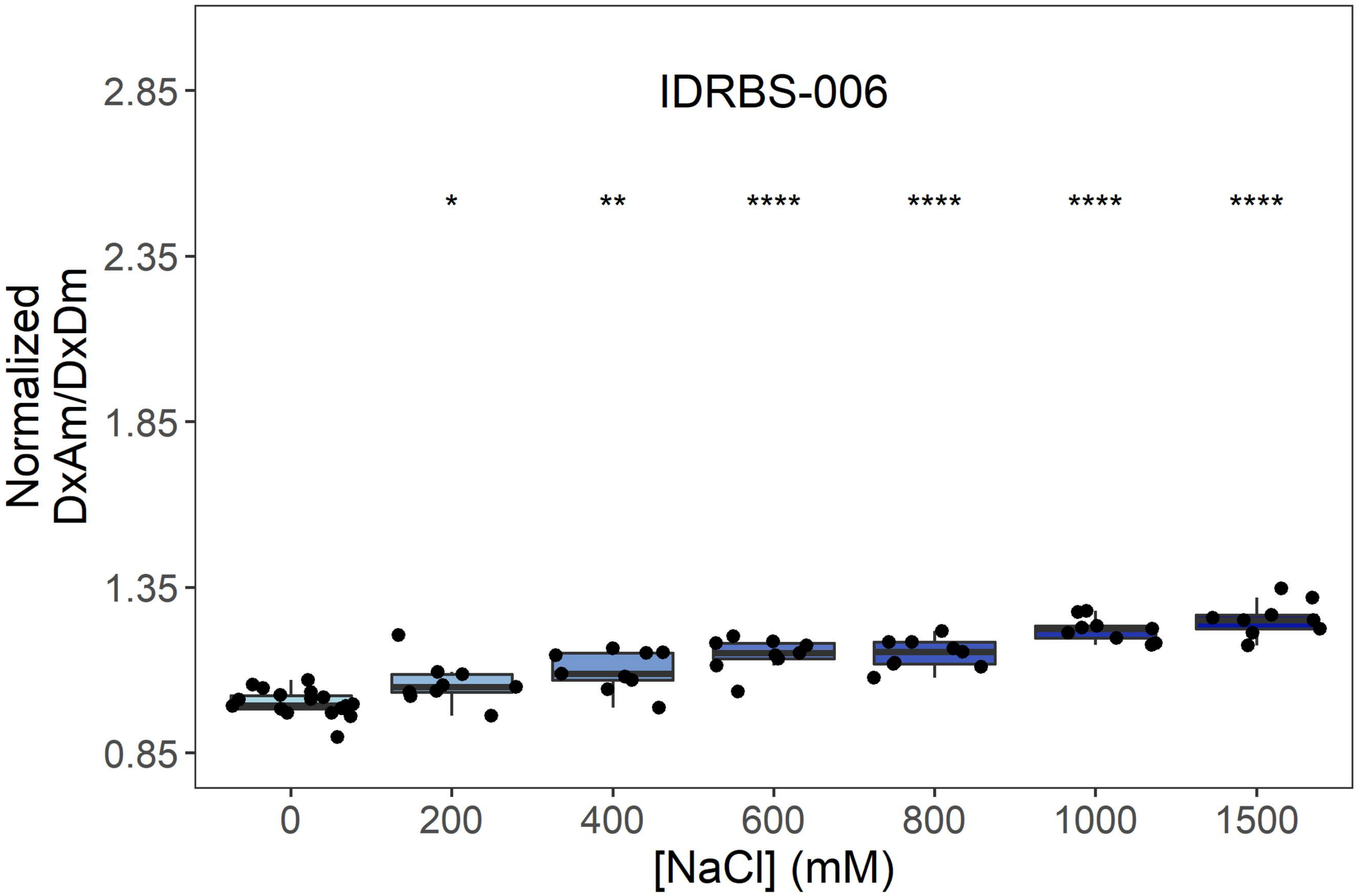
IDRBS-006
Normalized DxAm/DxDm
2.85
2.35
1.85
1.35
0.85
* ** **** **** **** ****
0 200 400 600 800 1000 1500
[NaCl] (mM)

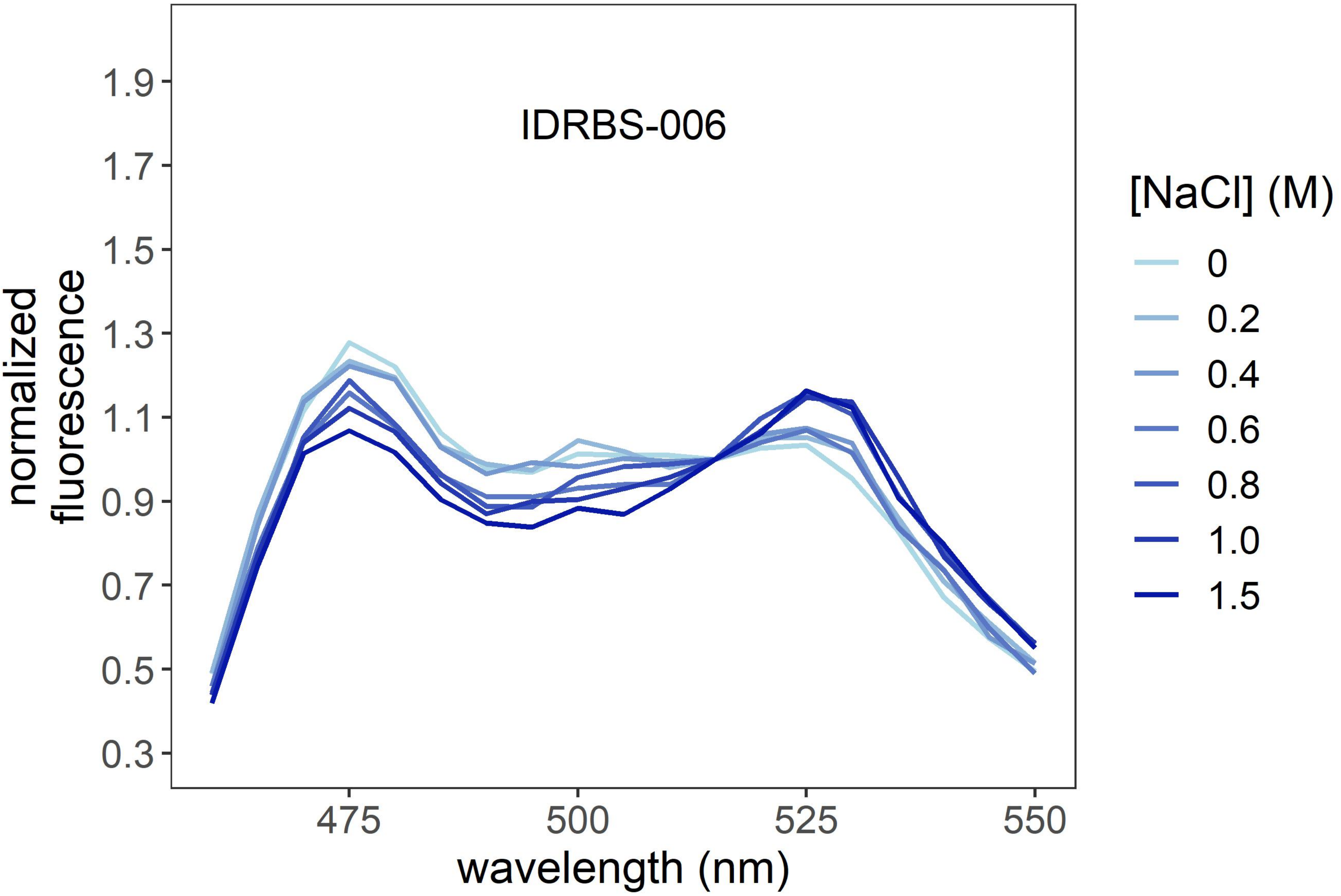
IDRBS-006
normalized fluorescence
1.9
1.7
1.5
1.3
1.1
0.9
0.7
0.5
0.3
475 500 525 550
wavelength (nm)
[NaCl] (M)
0
0.2
0.4
0.6
0.8
1.0
1.5

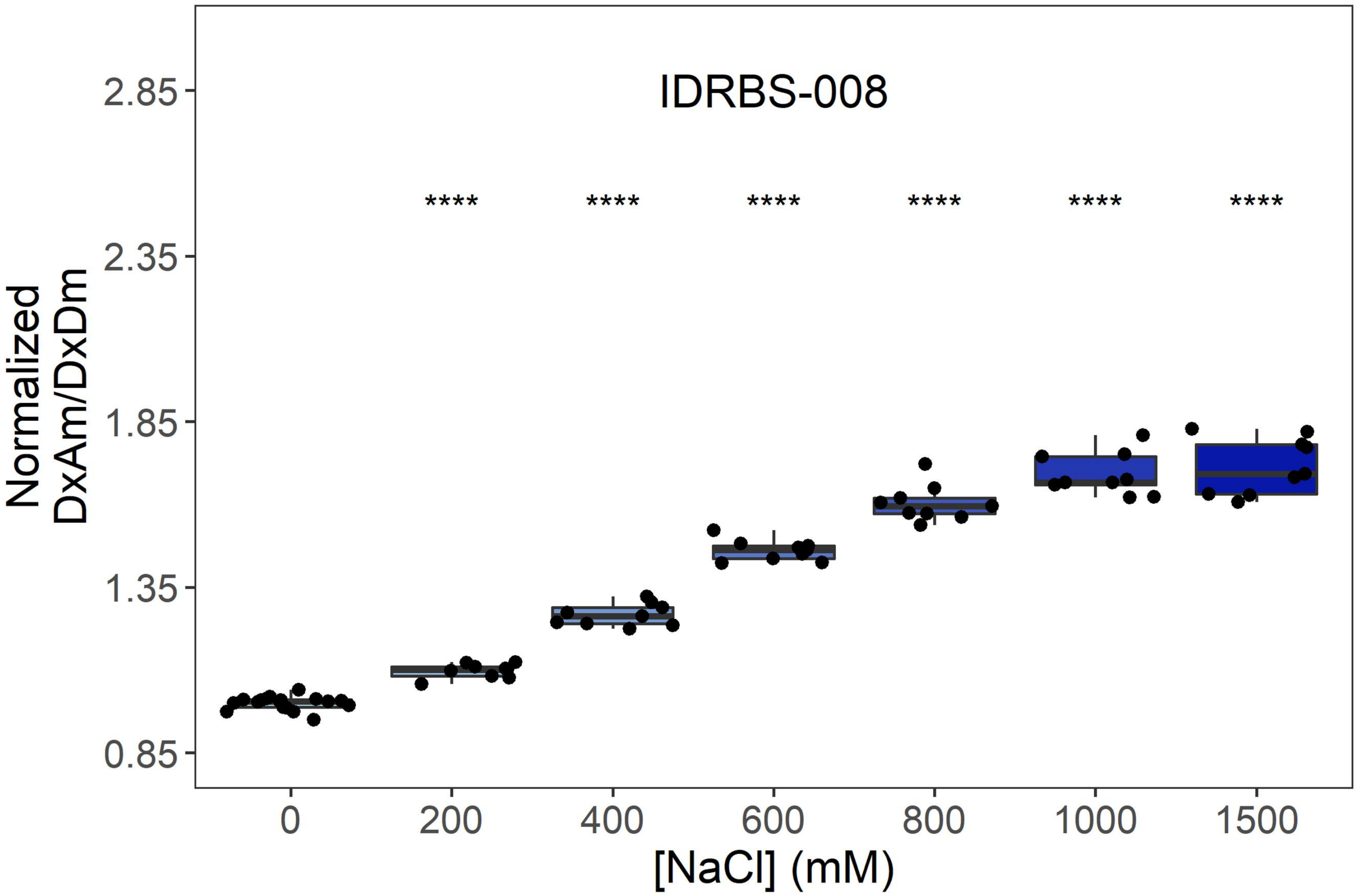

IDRBS-008
****
****
****
****
****
****
Normalized
DxAm/DxDm
2.85
2.35
1.85
1.35
0.85
0
200
400
600
800
1000
1500
[NaCl] (mM)


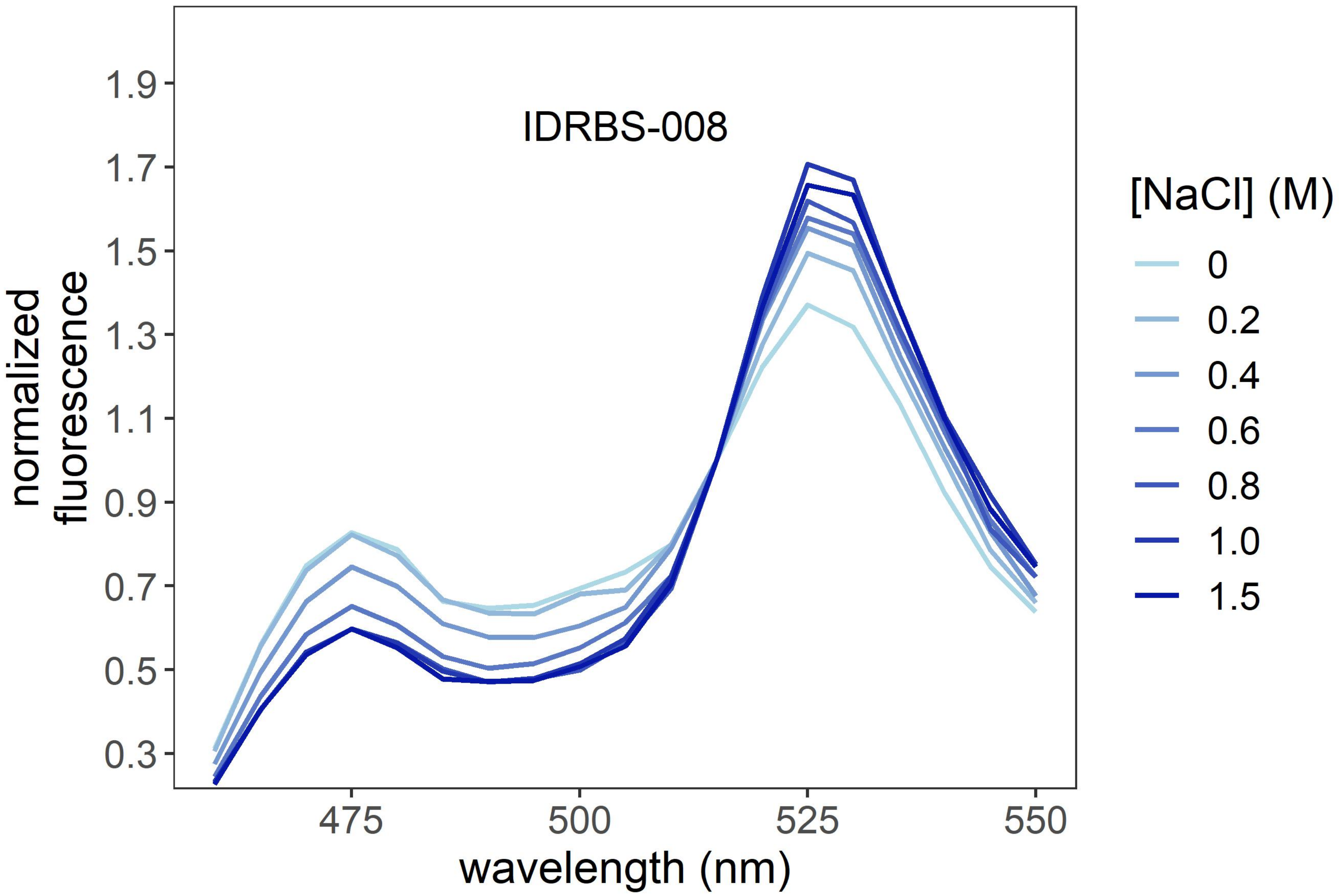

IDRBS-008
normalized
fluorescence
1.9
1.7
1.5
1.3
1.1
0.9
0.7
0.5
0.3
475
500
525
550
wavelength (nm)
[NaCl] (M)
0
0.2
0.4
0.6
0.8
1.0
1.5

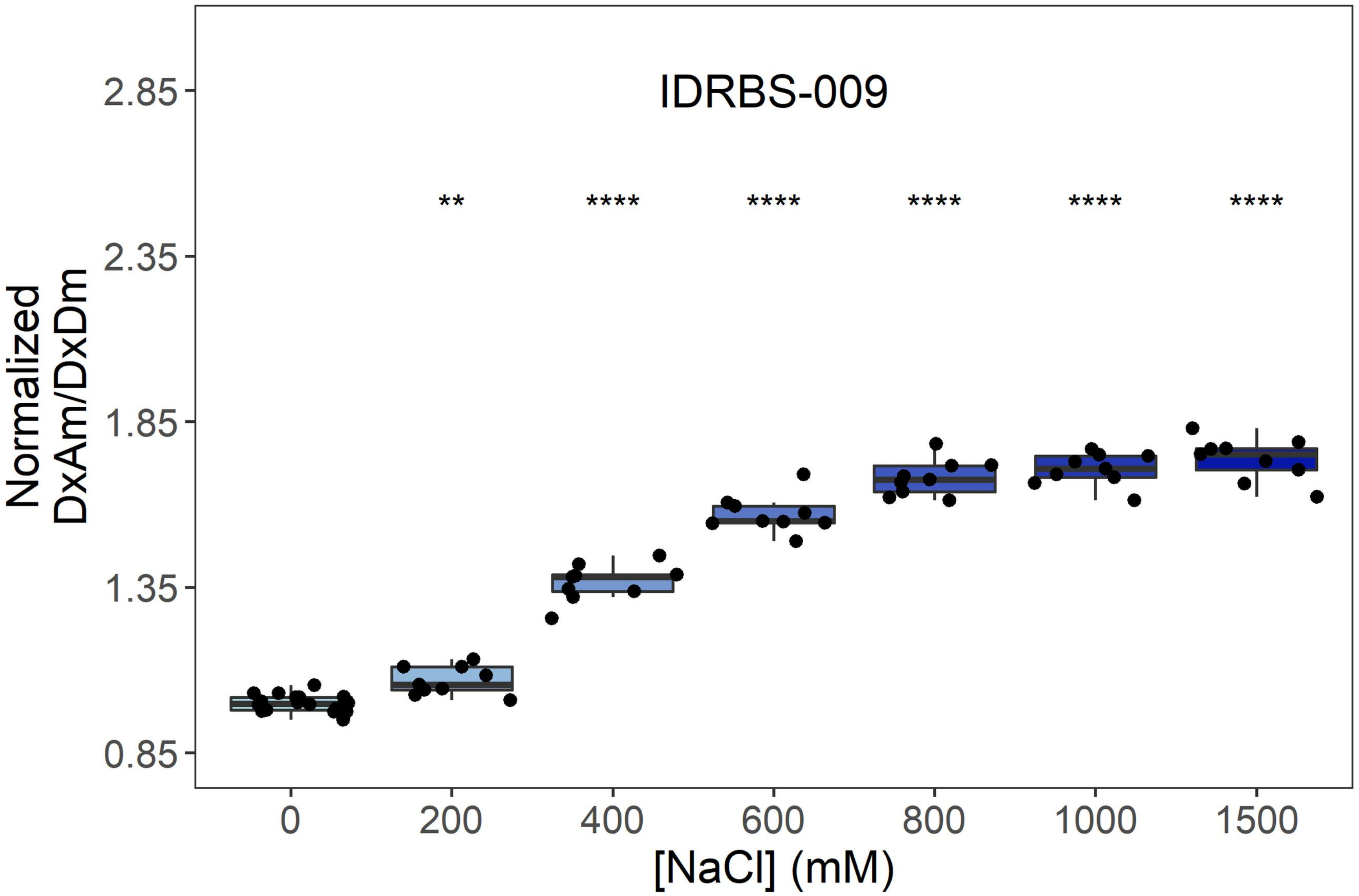
IDRBS-009
**
****
****
****
****
****
Normalized DxAm/DxDm
2.85
2.35
1.85
1.35
0.85
0
200
400
600
800
1000
1500
[NaCl] (mM)

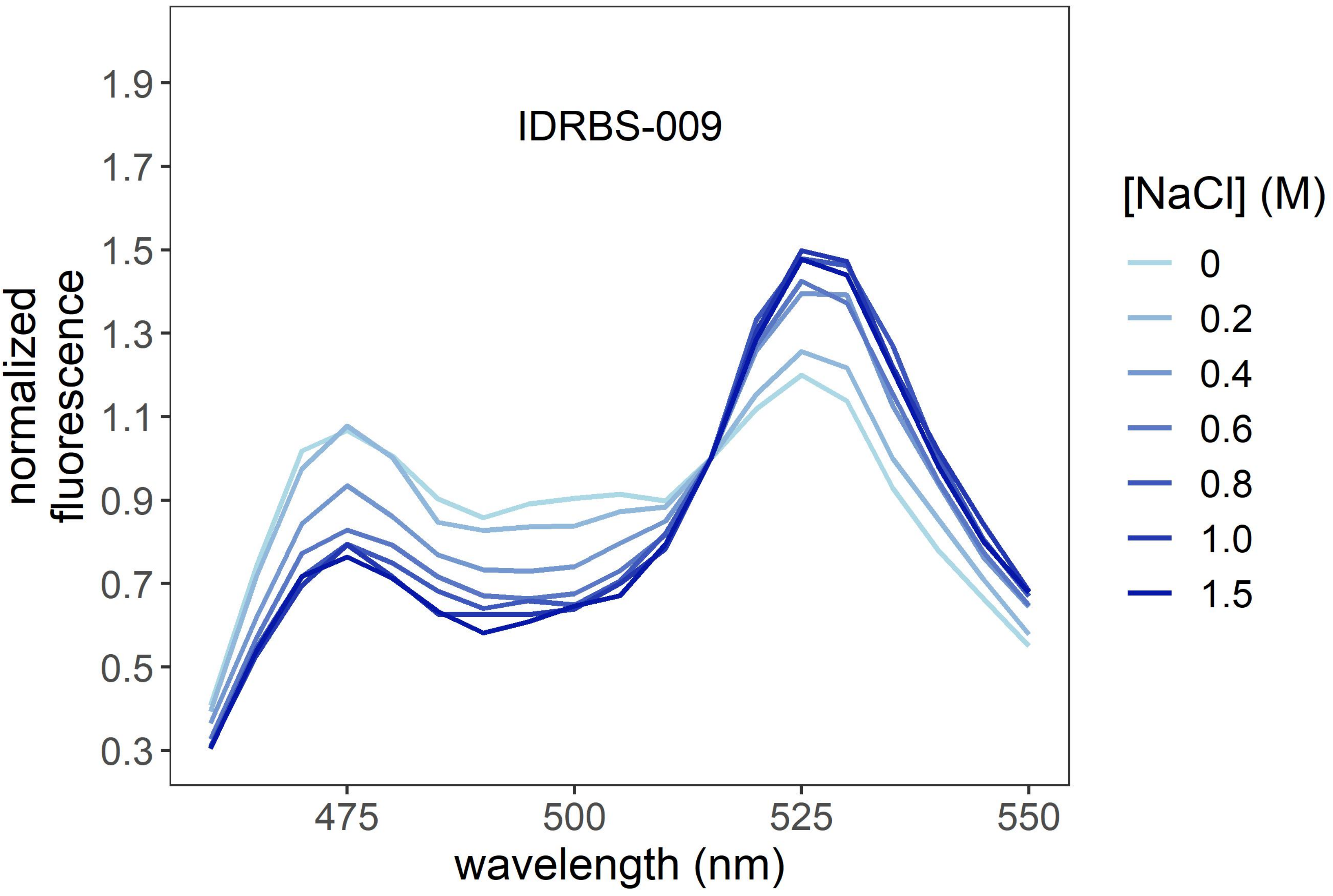
IDRBS-009
normalized fluorescence
1.9
1.7
1.5
1.3
1.1
0.9
0.7
0.5
0.3
475
500
525
550
wavelength (nm)
[NaCl] (M)
0
0.2
0.4
0.6
0.8
1.0
1.5

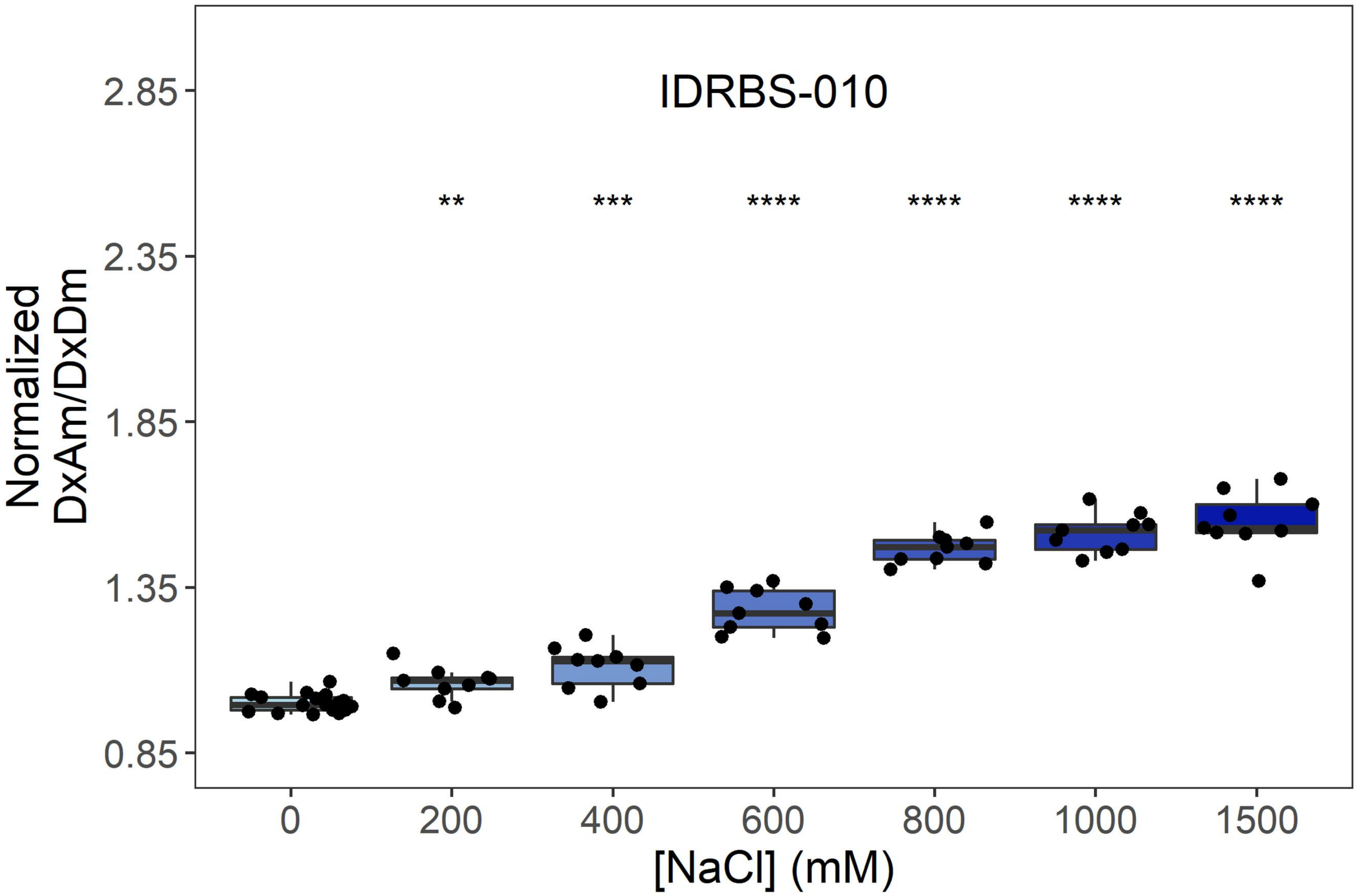
IDRBS-010
**
***
****
****
****
****
2.85
2.35
1.85
1.35
0.85
Normalized
DxAm/DxDm
0
200
400
600
800
1000
1500
[NaCl] (mM)

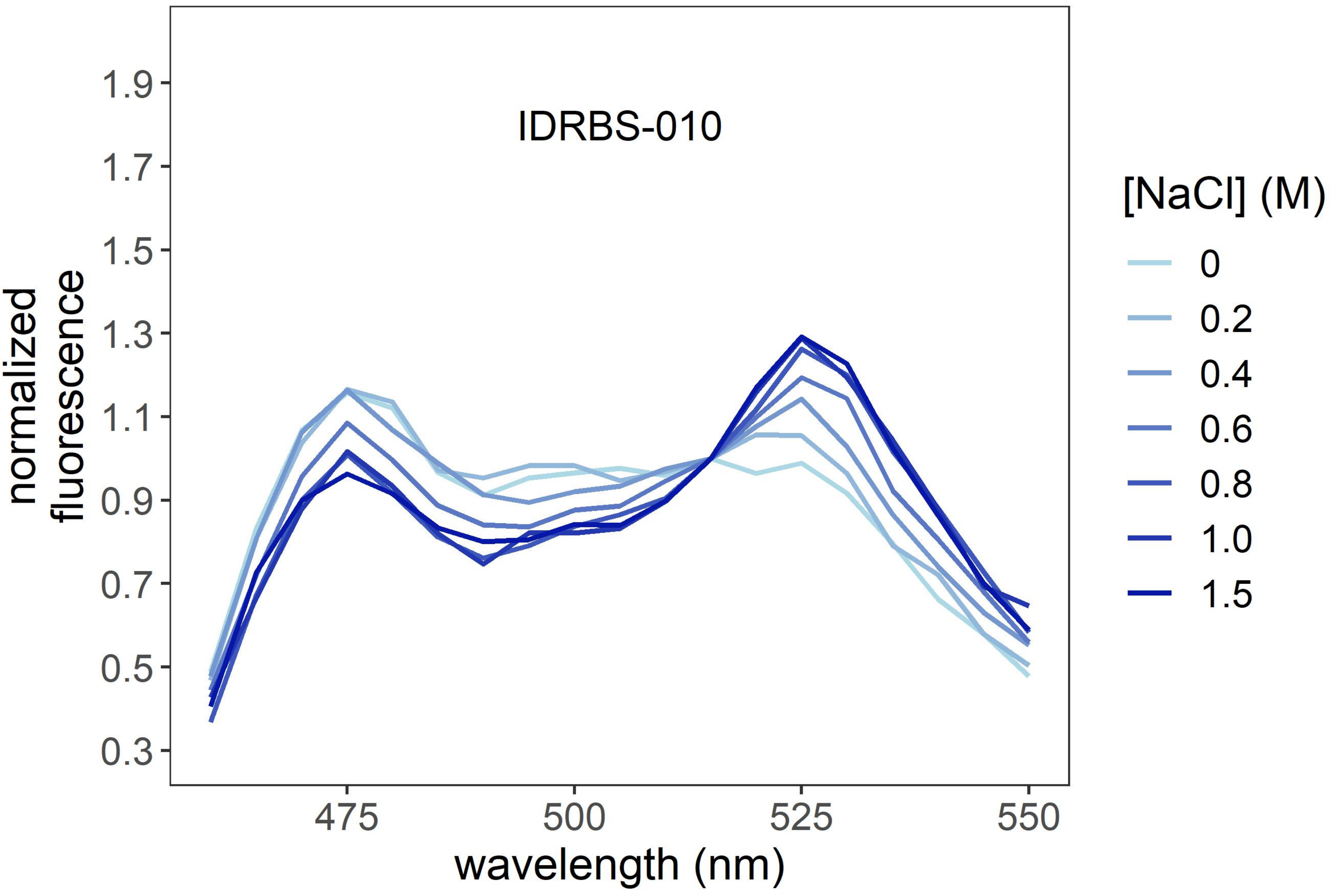
IDRBS-010
1.9
1.7
1.5
1.3
1.1
0.9
0.7
0.5
0.3
normalized
fluorescence
475
500
525
550
wavelength (nm)
[NaCl] (M)
0
0.2
0.4
0.6
0.8
1.0
1.5

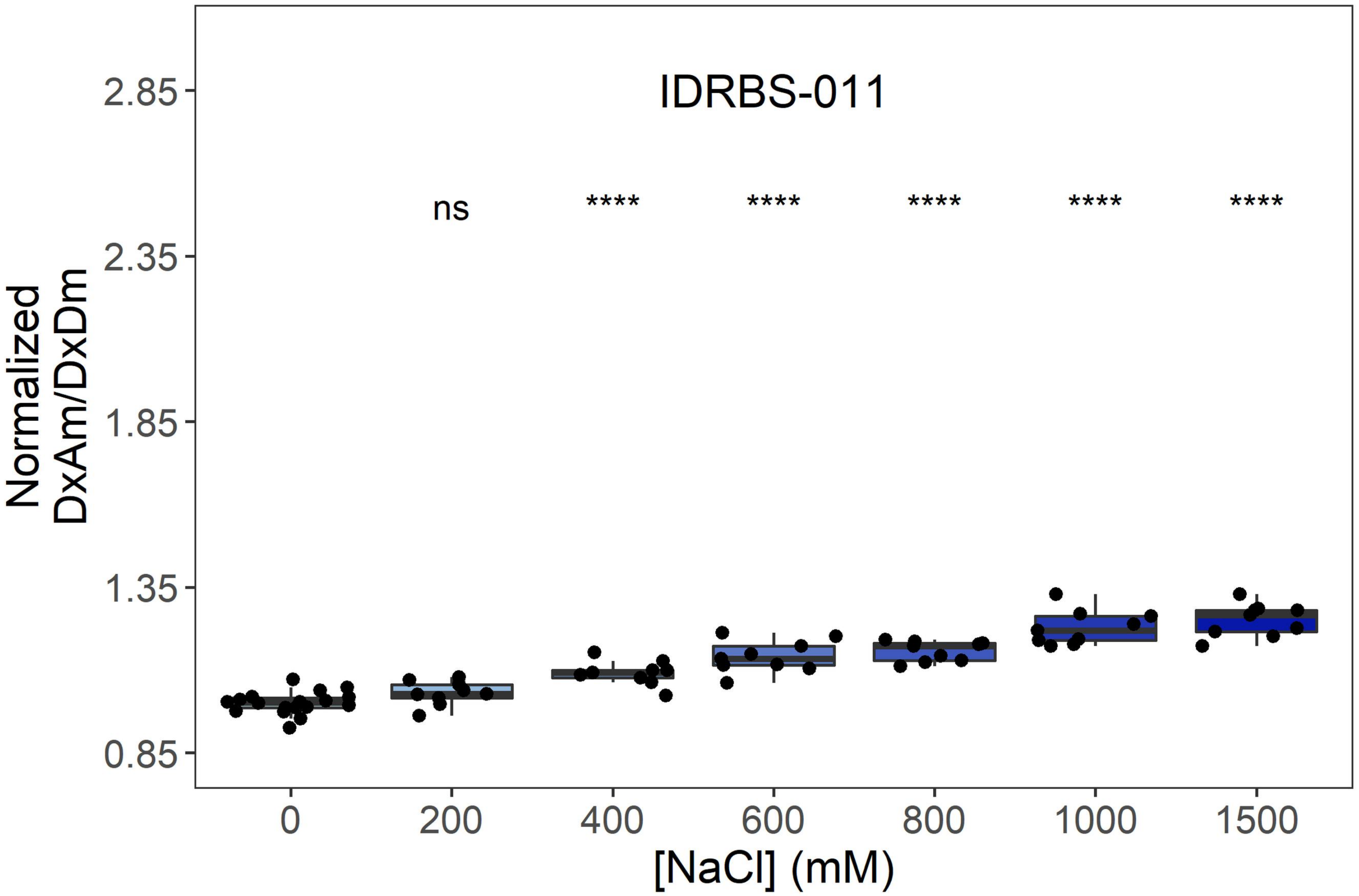
IDRBS-011
ns
****
****
****
****
****
2.85
2.35
1.85
1.35
0.85
Normalized
DxAm/DxDm
0
200
400
600
800
1000
1500
[NaCl] (mM)

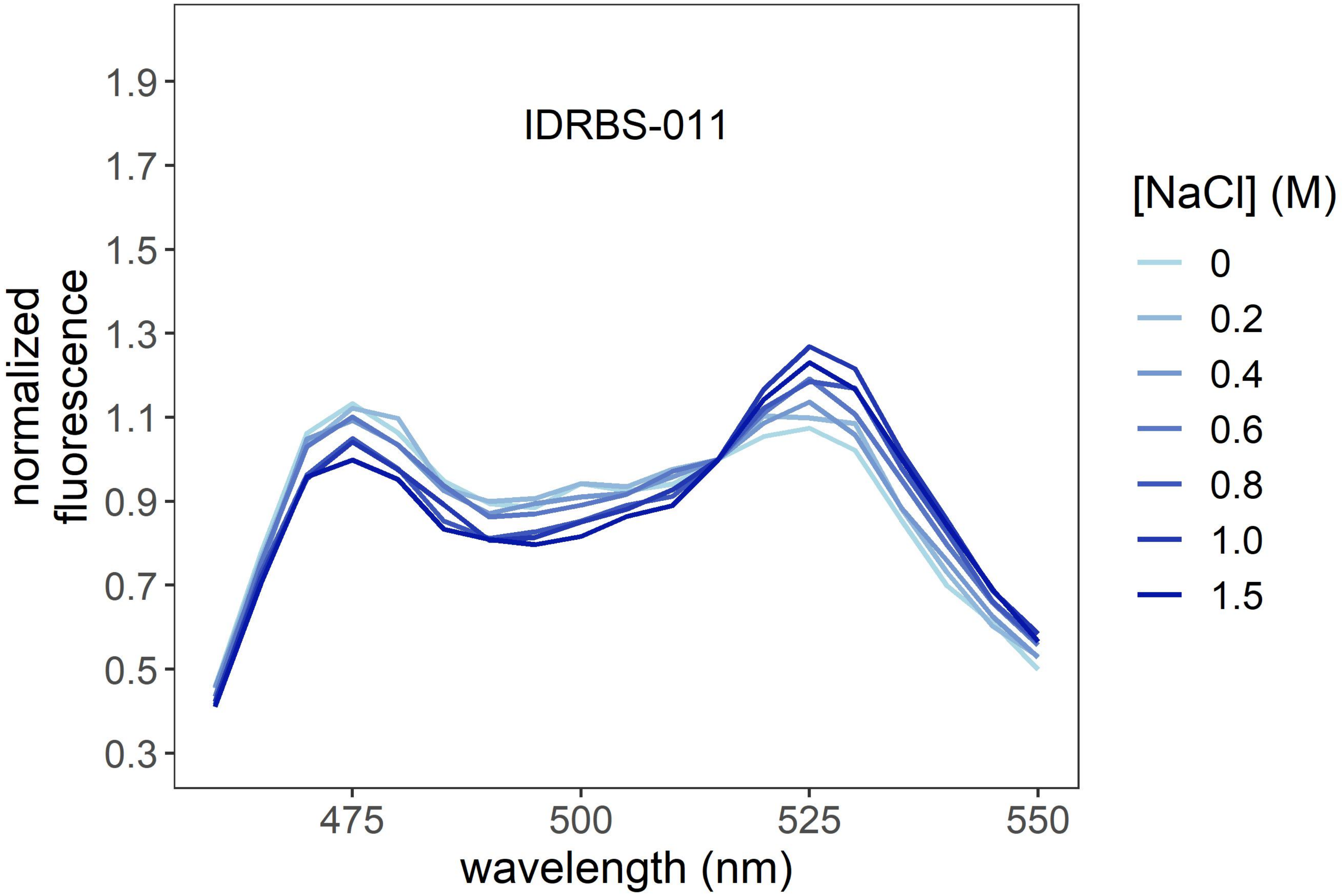
IDRBS-011
1.9
1.7
1.5
1.3
1.1
0.9
0.7
0.5
0.3
normalized
fluorescence
475
500
525
550
wavelength (nm)
[NaCl] (M)
0
0.2
0.4
0.6
0.8
1.0
1.5

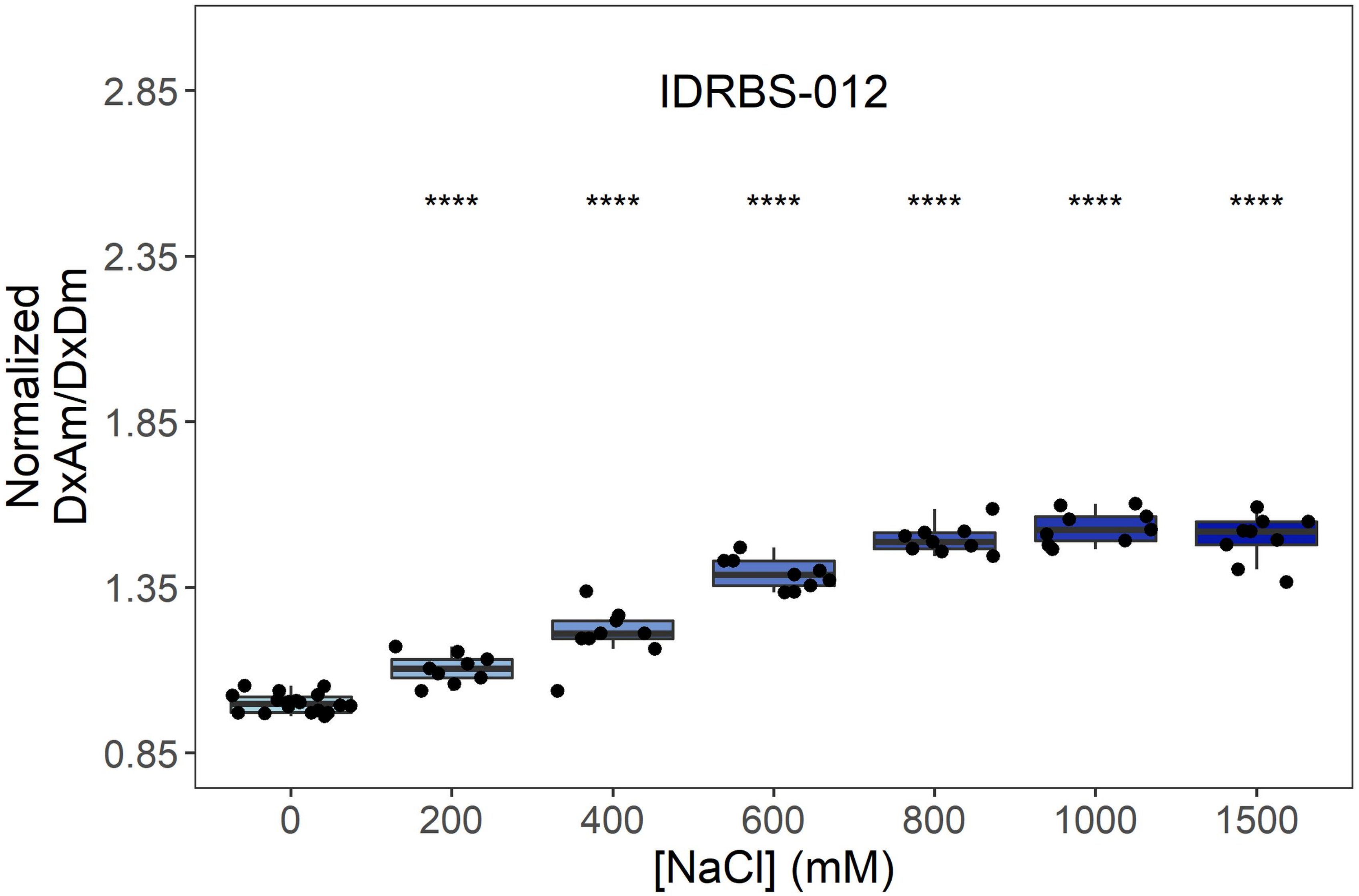
IDRBS-012
Normalized
DxAm/DxDm
2.85
2.35
1.85
1.35
0.85
****
****
****
****
****
****
0
200
400
600
800
1000
1500
[NaCl] (mM)

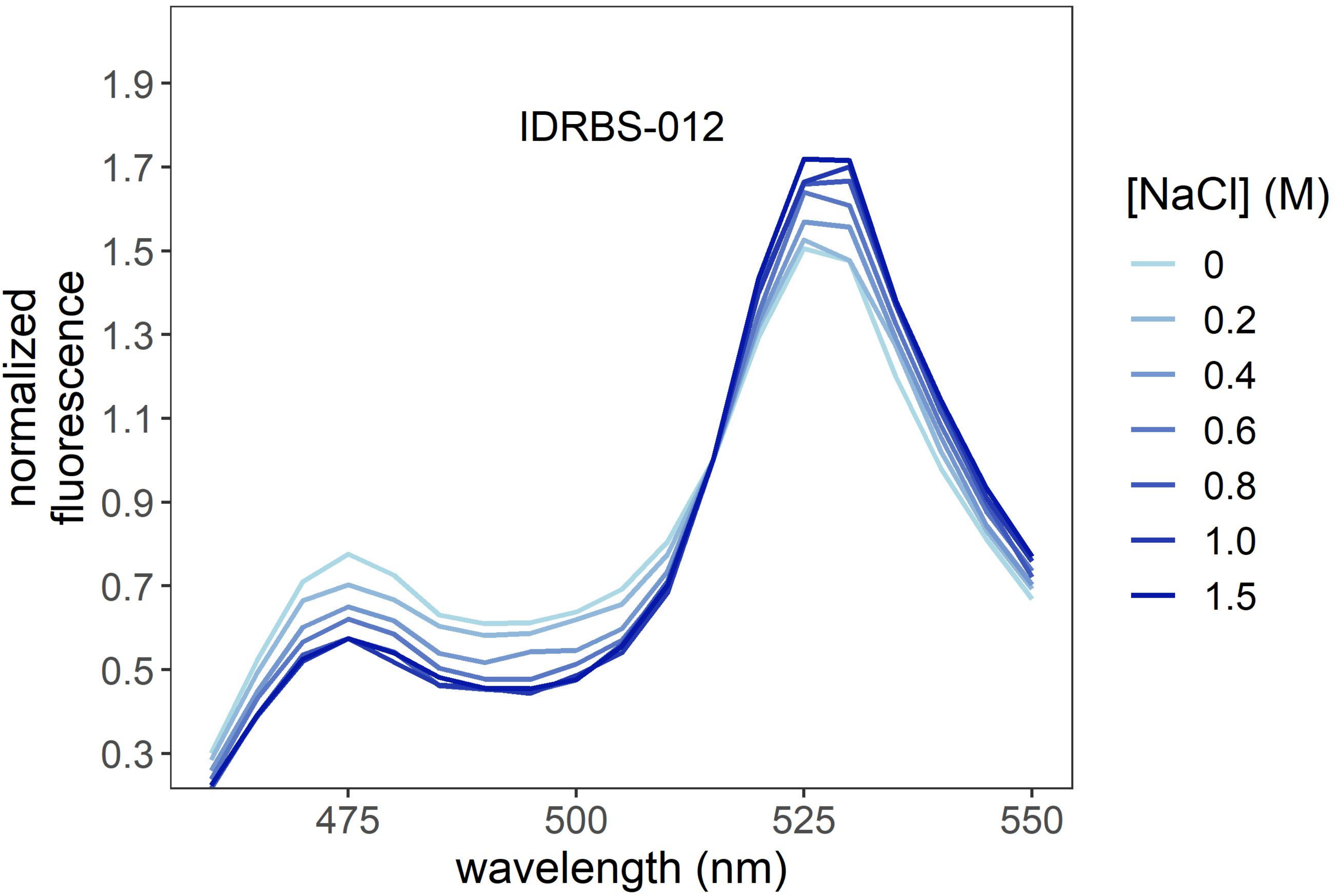
IDRBS-012
normalized
fluorescence
1.9
1.7
1.5
1.3
1.1
0.9
0.7
0.5
0.3
475
500
525
550
wavelength (nm)
[NaCl] (M)
0
0.2
0.4
0.6
0.8
1.0
1.5

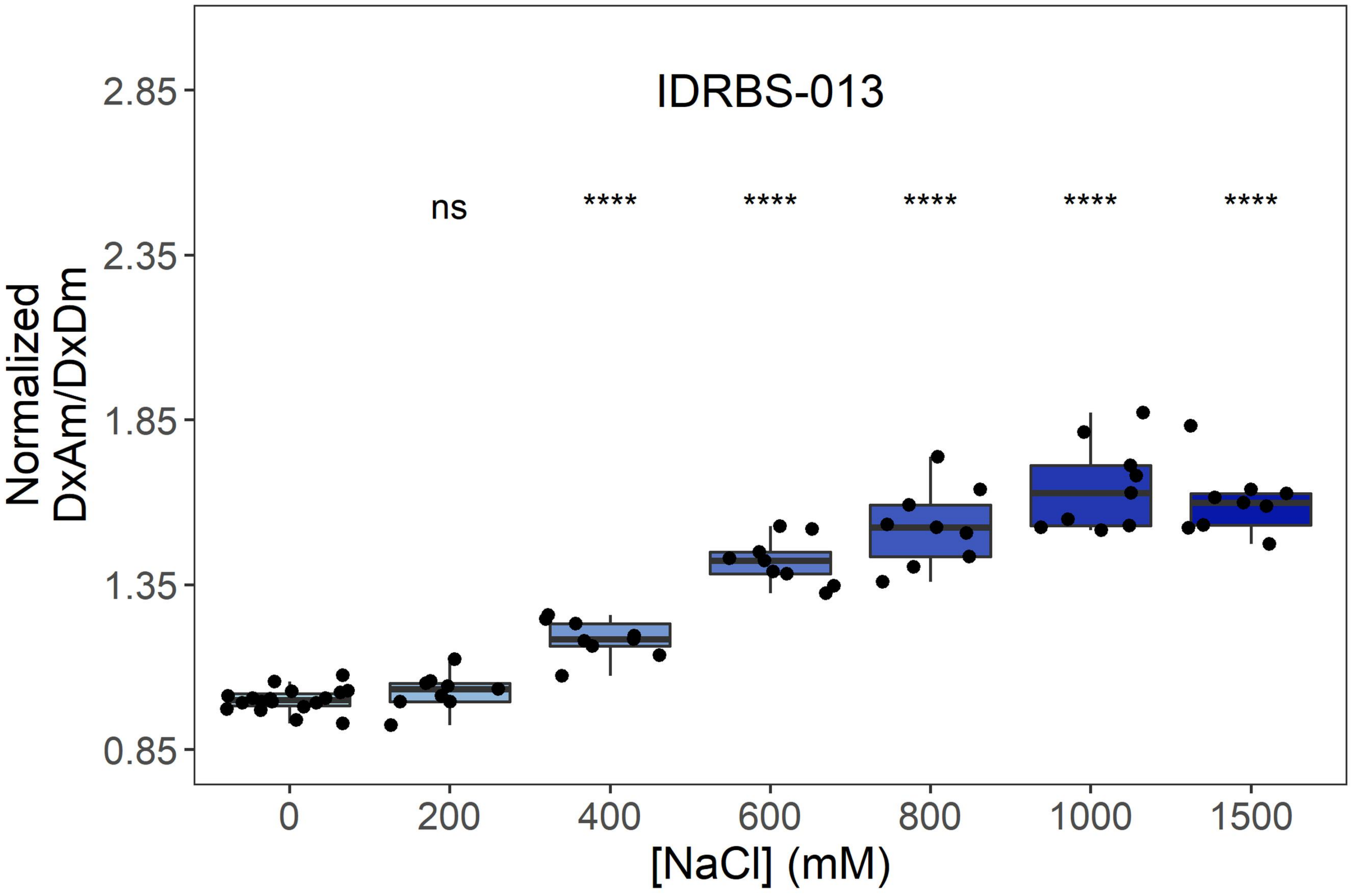

IDRBS-013
ns
****
****
****
****
****
Normalized DxAm/DxDm
2.85
2.35
1.85
1.35
0.85
0
200
400
600
800
1000
1500
[NaCl] (mM)


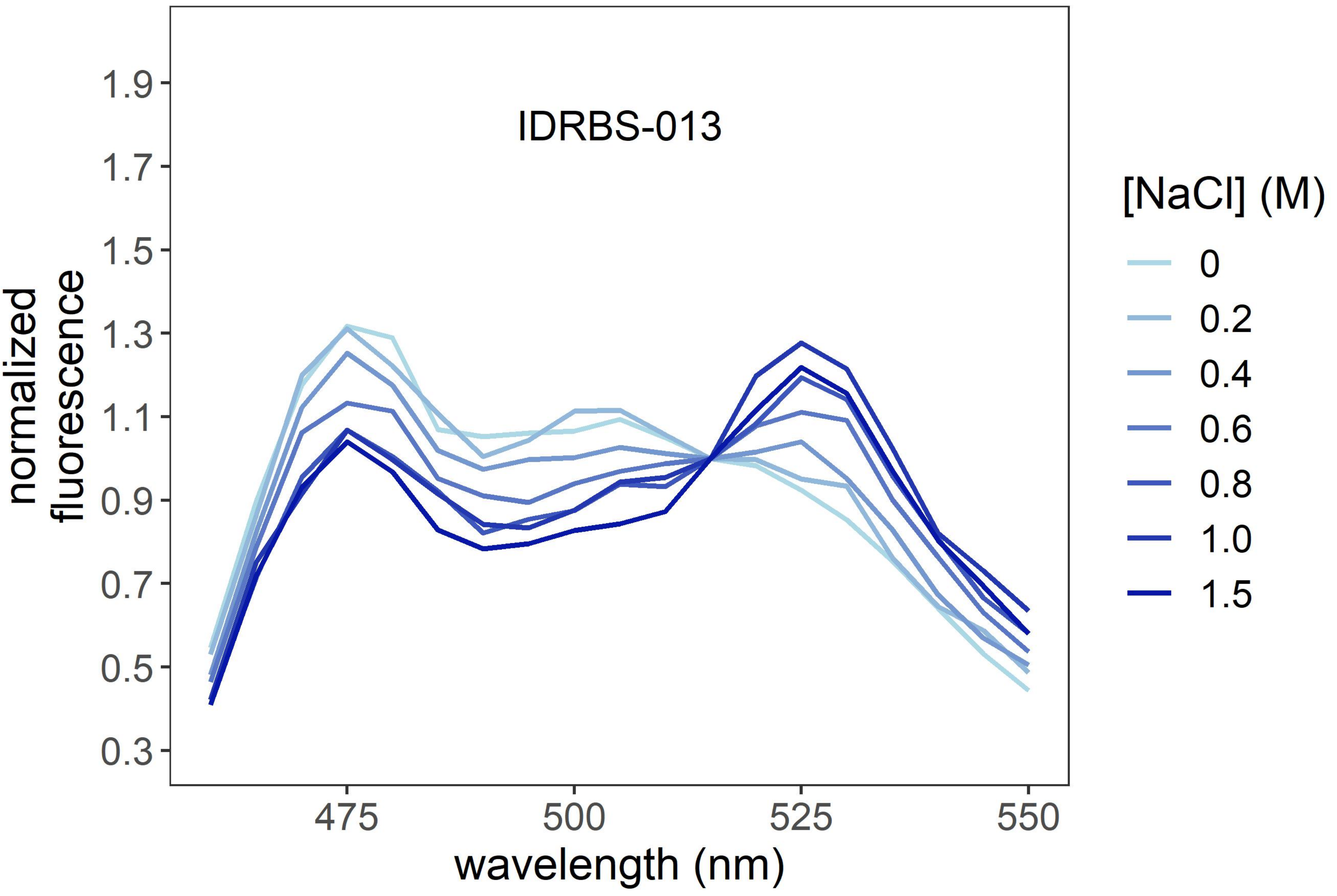

IDRBS-013
normalized fluorescence
1.9
1.7
1.5
1.3
1.1
0.9
0.7
0.5
0.3
475
500
525
550
wavelength (nm)
[NaCl] (M)
0
0.2
0.4
0.6
0.8
1.0
1.5

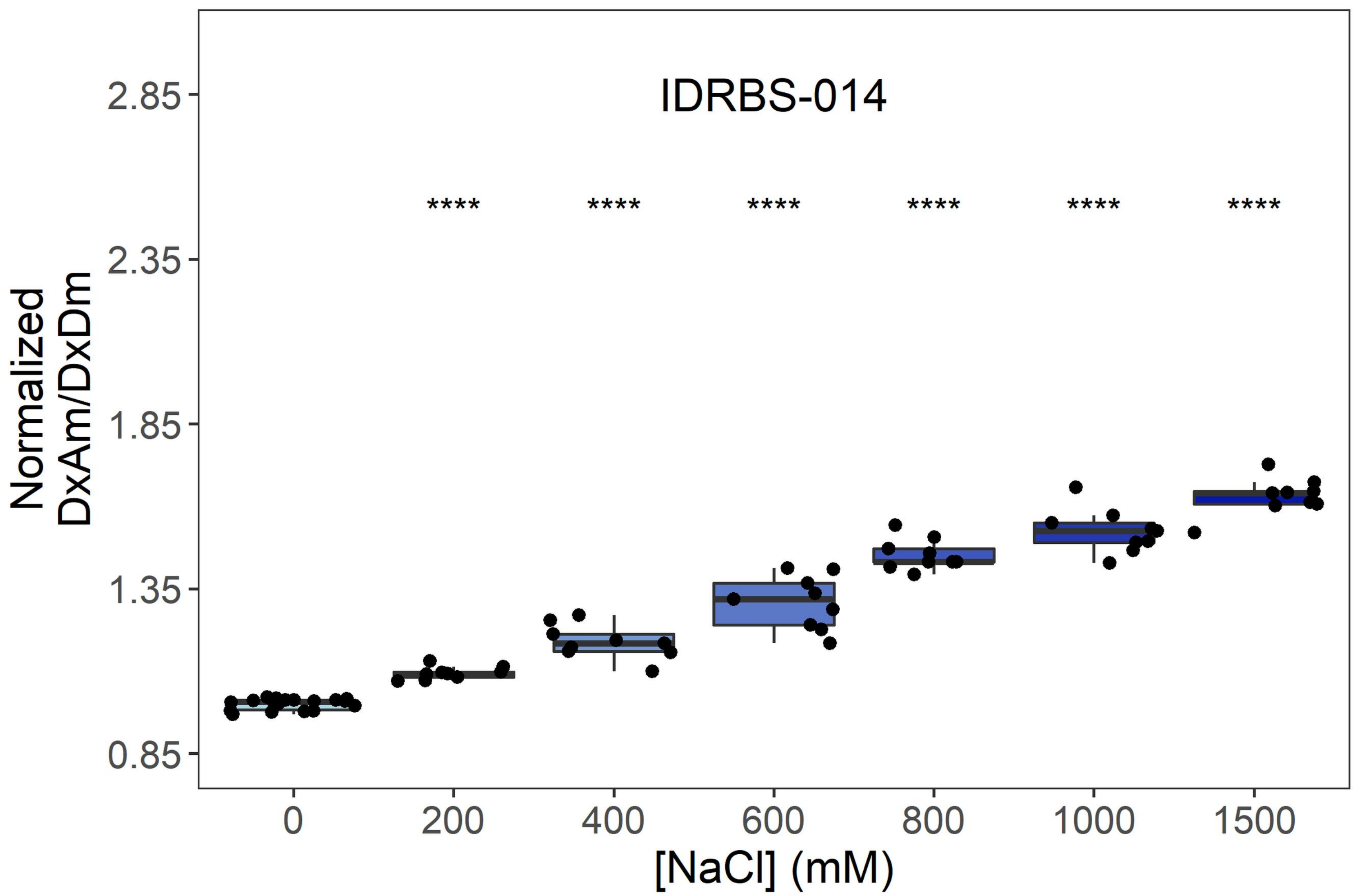
IDRBS-014
Normalized DxAm/DxDm
[NaCl] (mM)
****
2.85
2.35
1.85
1.35
0.85
0
200
400
600
800
1000
1500

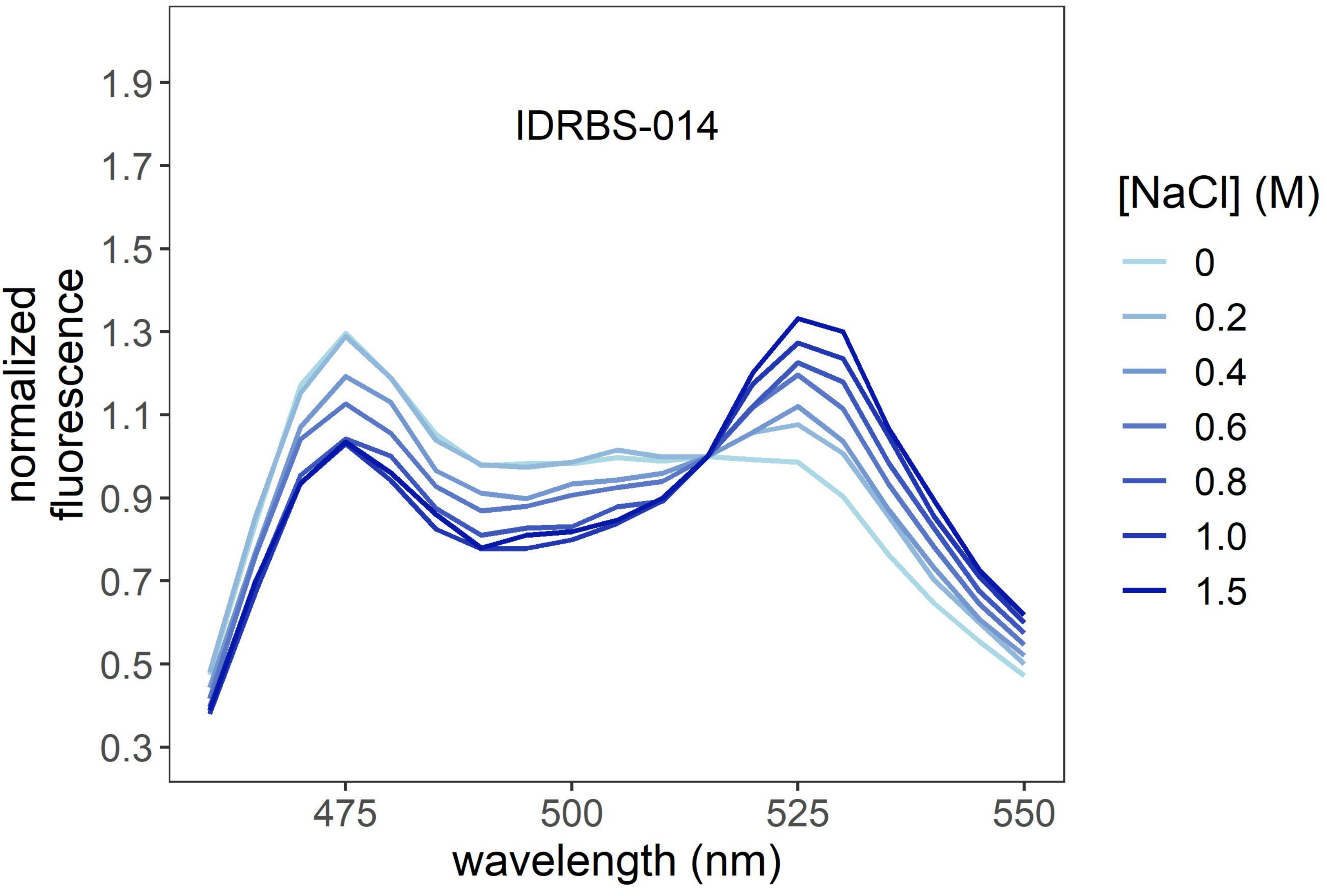
IDRBS-014
normalized fluorescence
wavelength (nm)
[NaCl] (M)
0
0.2
0.4
0.6
0.8
1.0
1.5
1.9
1.7
1.5
1.3
1.1
0.9
0.7
0.5
0.3
475
500
525
550

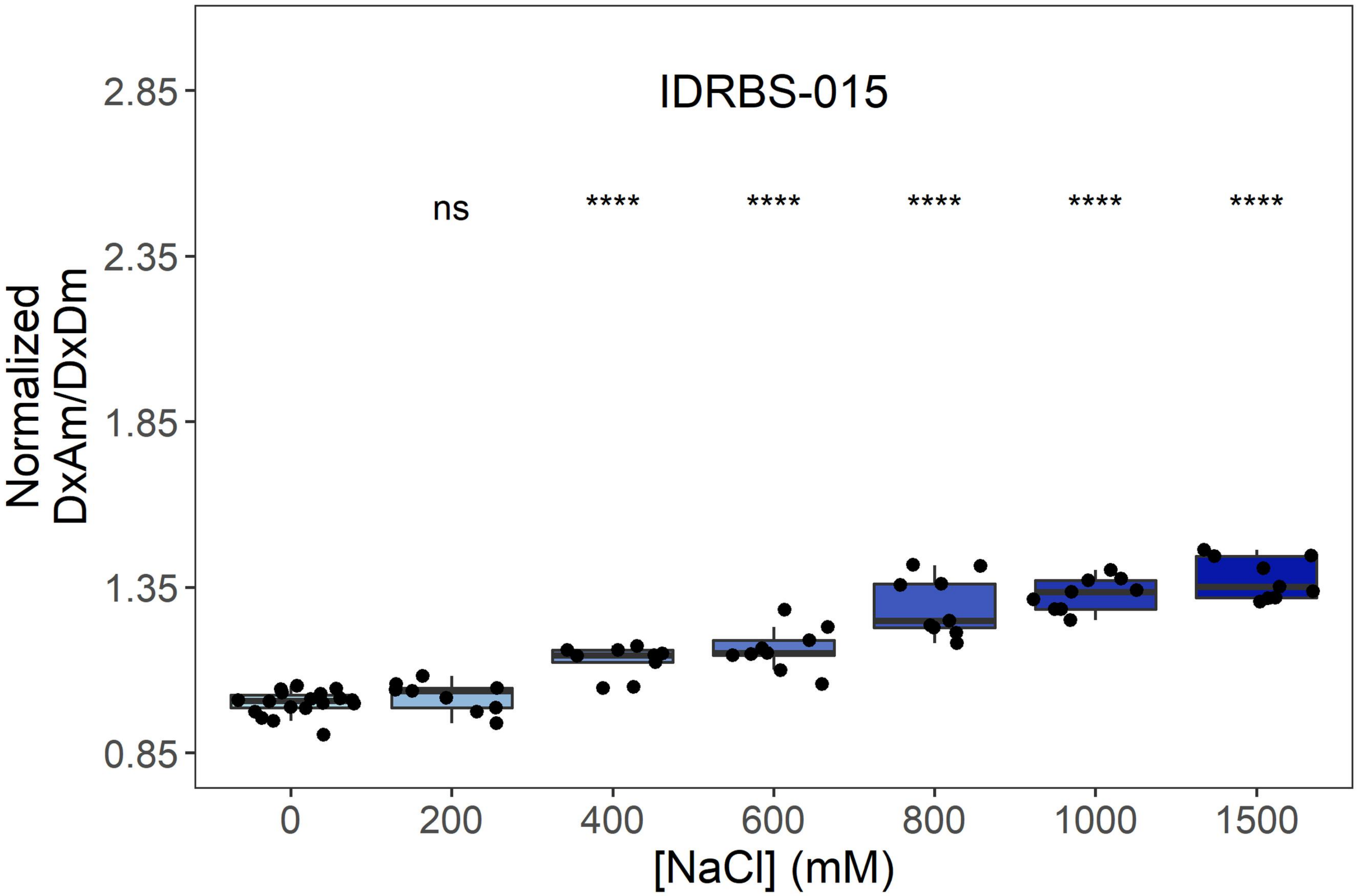
IDRBS-015
ns
****
****
****
****
****
2.85
2.35
1.85
1.35
0.85
Normalized
DxAm/DxDm
0
200
400
600
800
1000
1500
[NaCl] (mM)

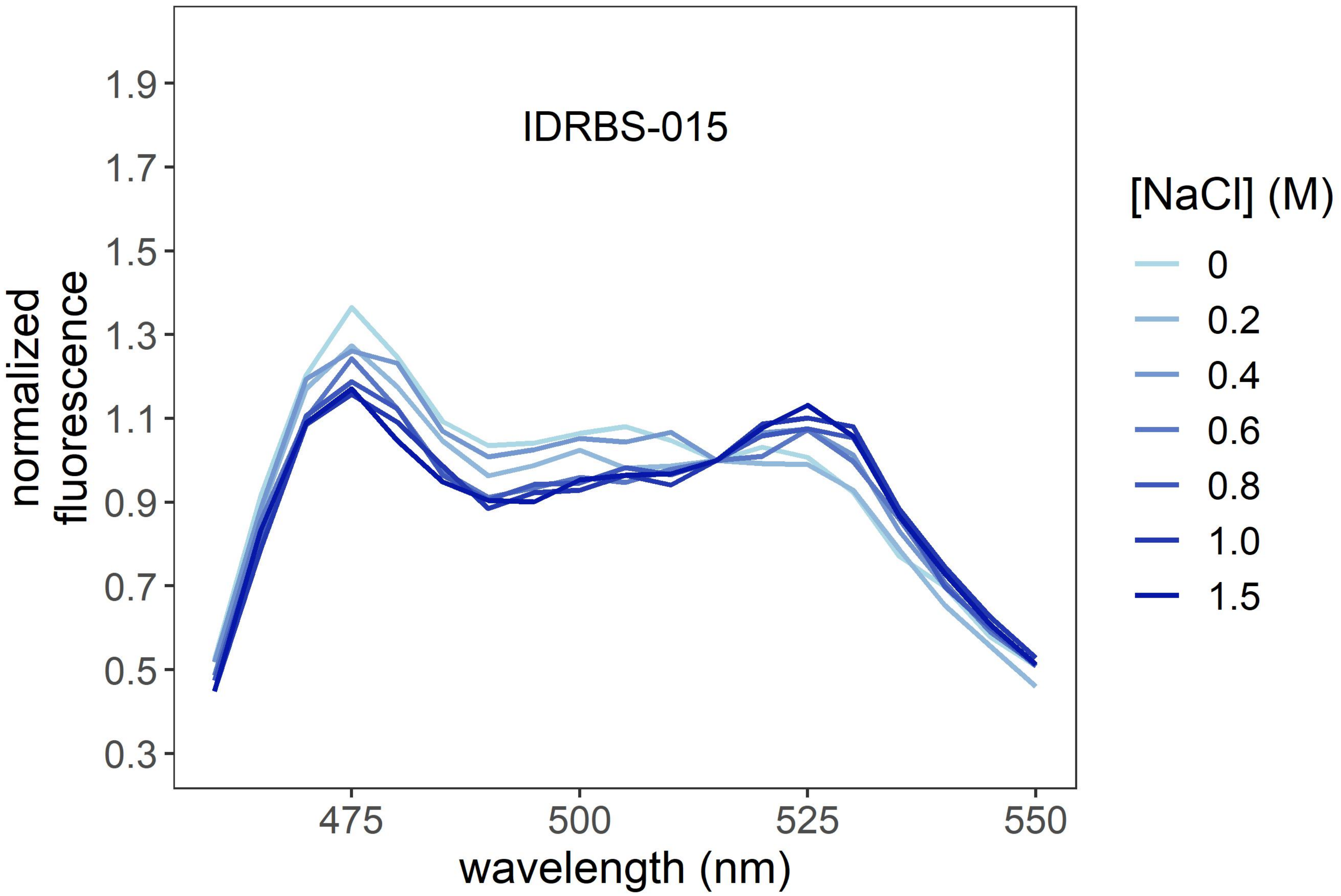
IDRBS-015
1.9
1.7
1.5
1.3
1.1
0.9
0.7
0.5
0.3
normalized
fluorescence
475
500
525
550
wavelength (nm)
[NaCl] (M)
0
0.2
0.4
0.6
0.8
1.0
1.5

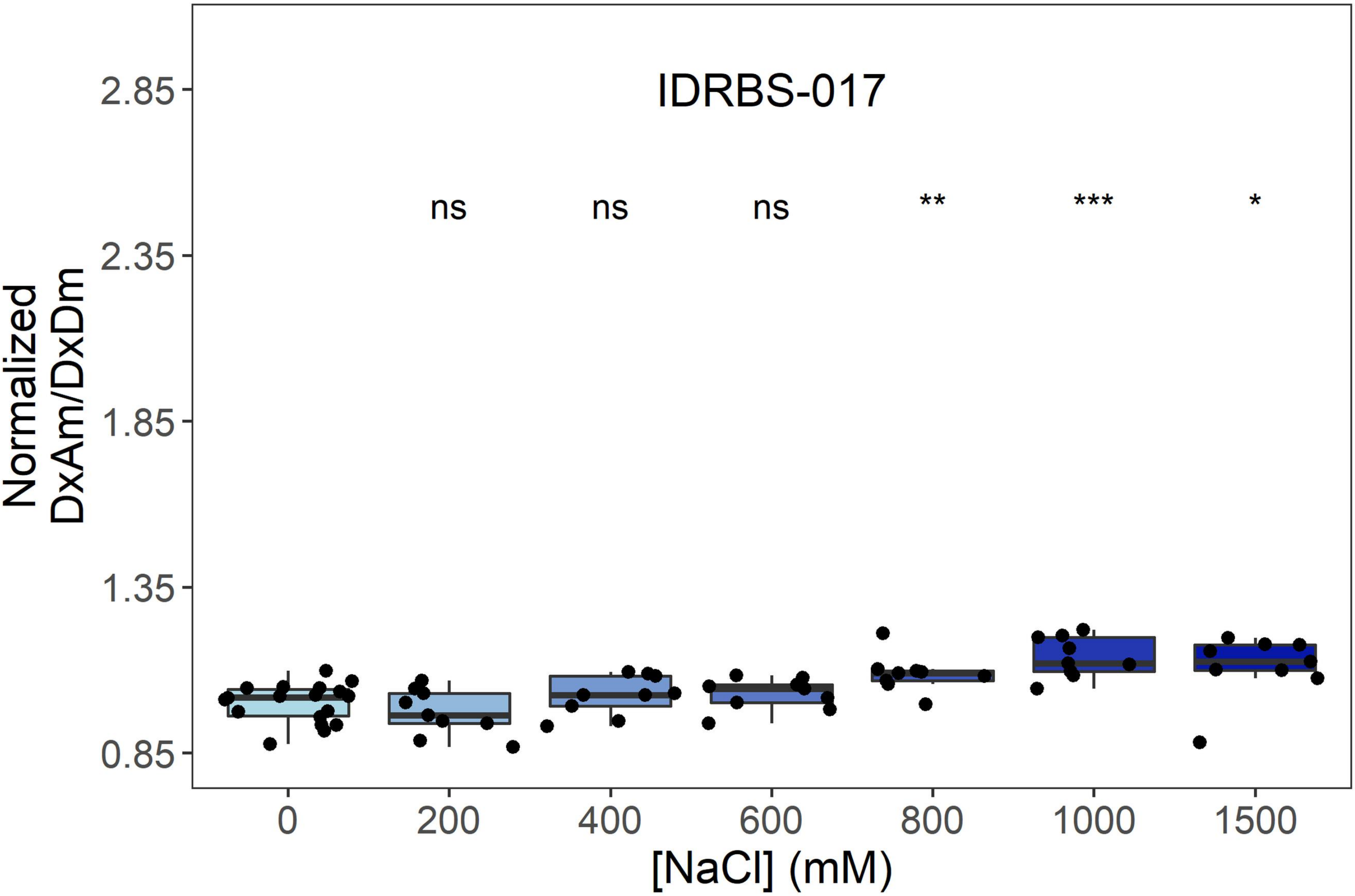
IDRBS-017
ns
ns
ns
**
***
*
Normalized
DxAm/DxDm
2.85
2.35
1.85
1.35
0.85
0
200
400
600
800
1000
1500
[NaCl] (mM)

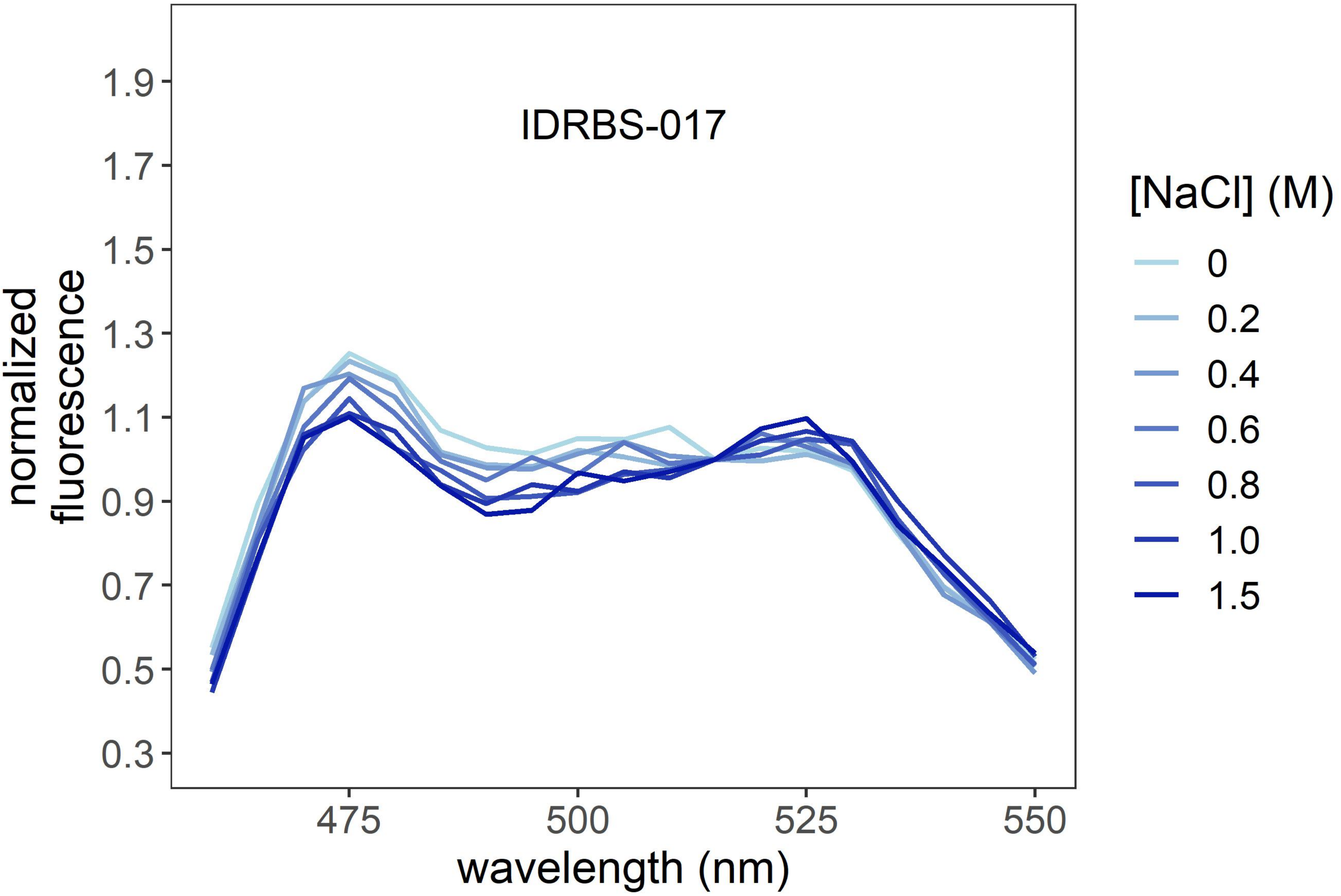
IDRBS-017
normalized
fluorescence
1.9
1.7
1.5
1.3
1.1
0.9
0.7
0.5
0.3
475
500
525
550
wavelength (nm)
[NaCl] (M)
0
0.2
0.4
0.6
0.8
1.0
1.5

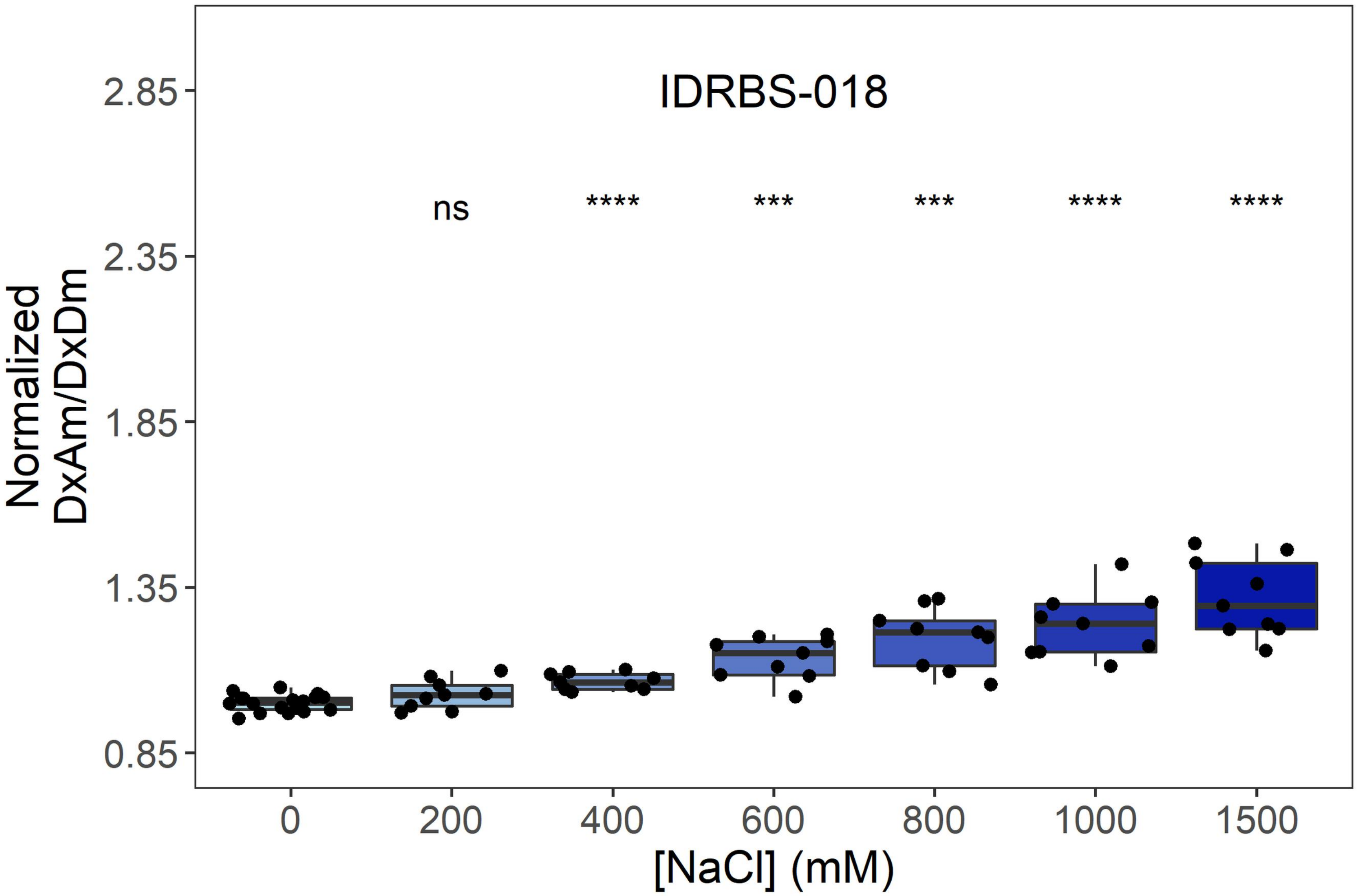

IDRBS-018
ns
****
***
***
****
****
2.85
2.35
1.85
1.35
0.85
Normalized
DxAm/DxDm
0
200
400
600
800
1000
1500
[NaCl] (mM)


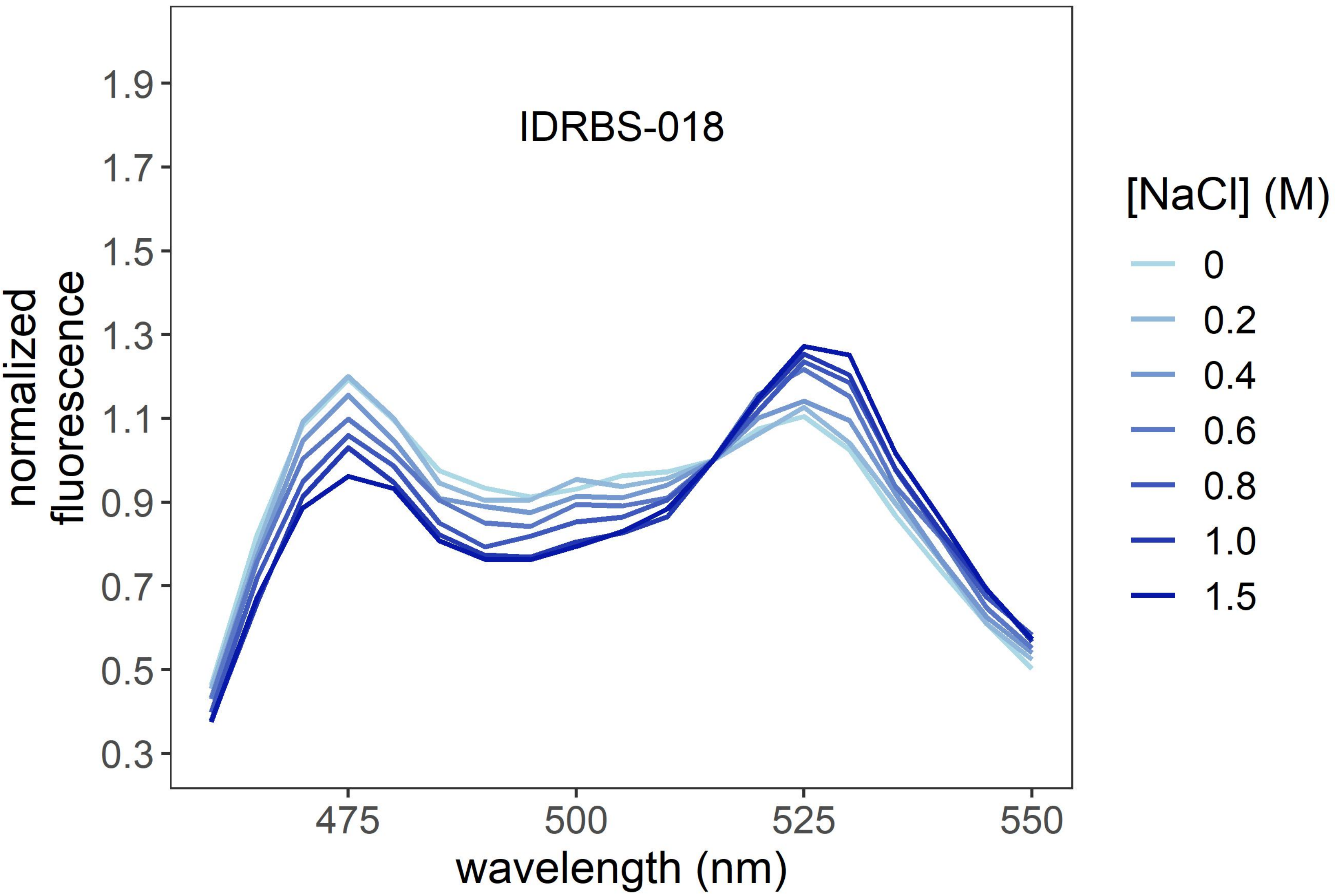

IDRBS-018
1.9
1.7
1.5
1.3
1.1
0.9
0.7
0.5
0.3
normalized
fluorescence
475
500
525
550
wavelength (nm)
[NaCl] (M)
0
0.2
0.4
0.6
0.8
1.0
1.5

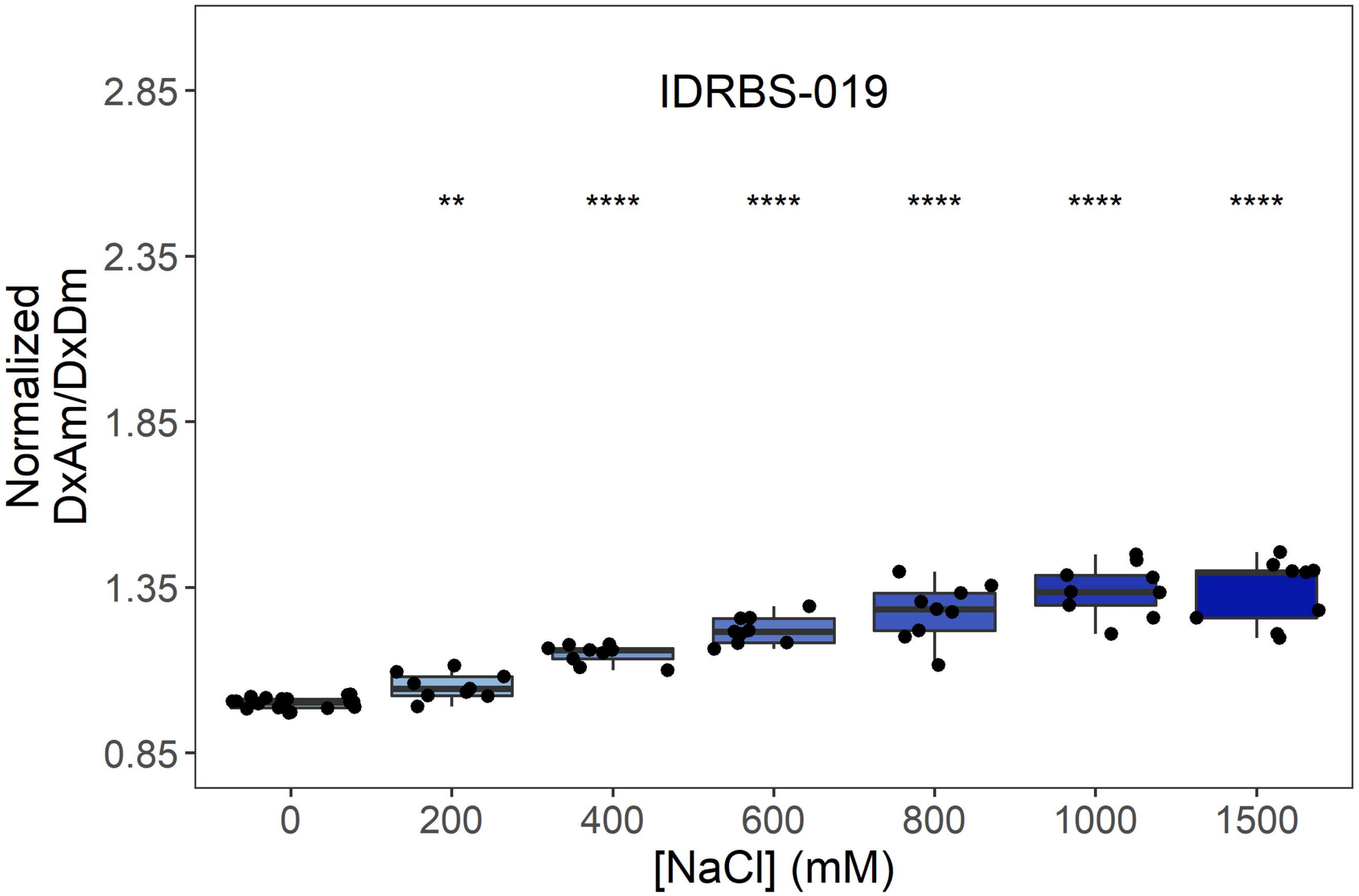
IDRBS-019
Normalized
DxAm/DxDm
2.85
2.35
1.85
1.35
0.85
**
****
****
****
****
****
0
200
400
600
800
1000
1500
[NaCl] (mM)

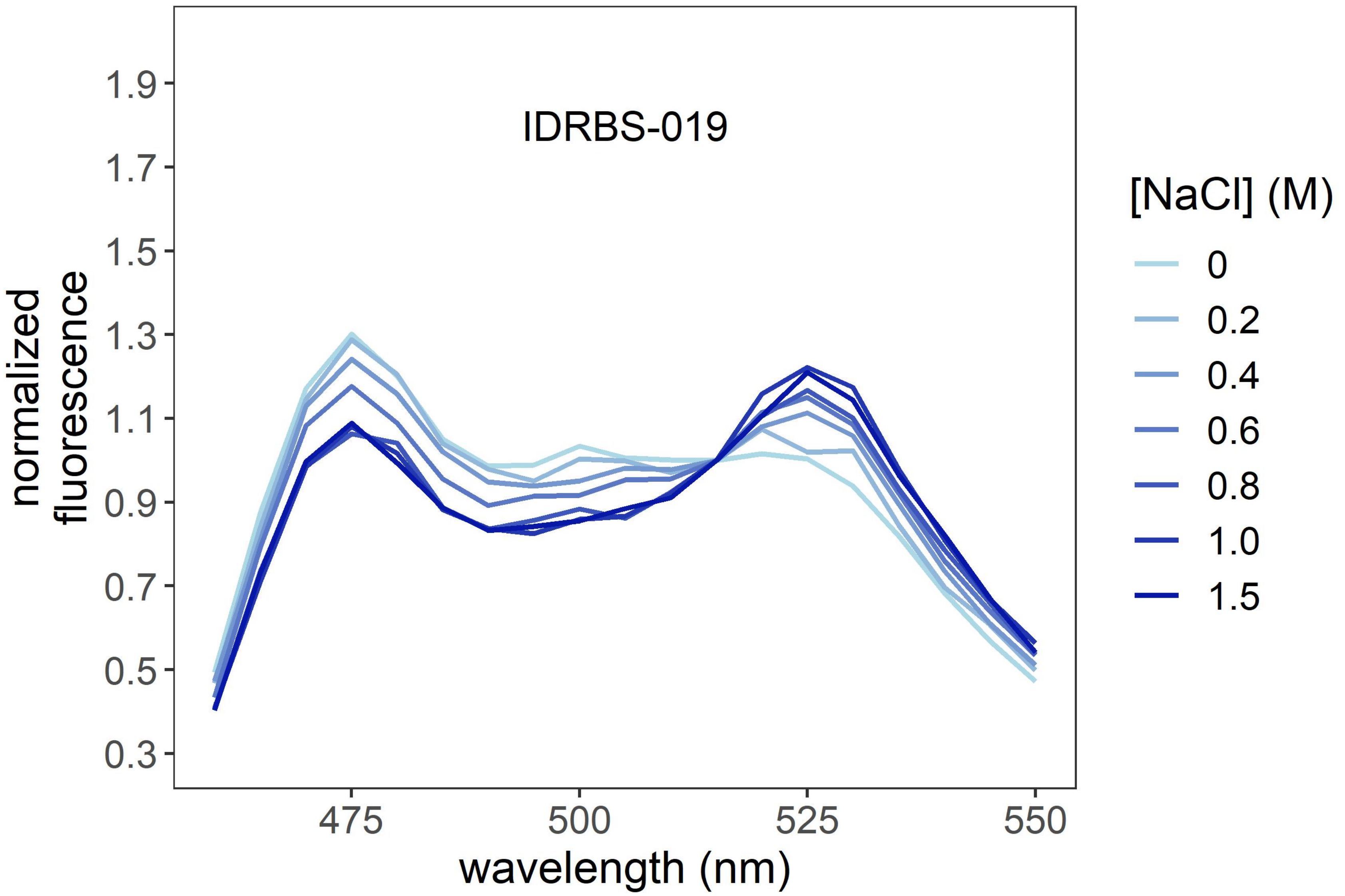
IDRBS-019
normalized
fluorescence
1.9
1.7
1.5
1.3
1.1
0.9
0.7
0.5
0.3
475
500
525
550
wavelength (nm)
[NaCl] (M)
0
0.2
0.4
0.6
0.8
1.0
1.5

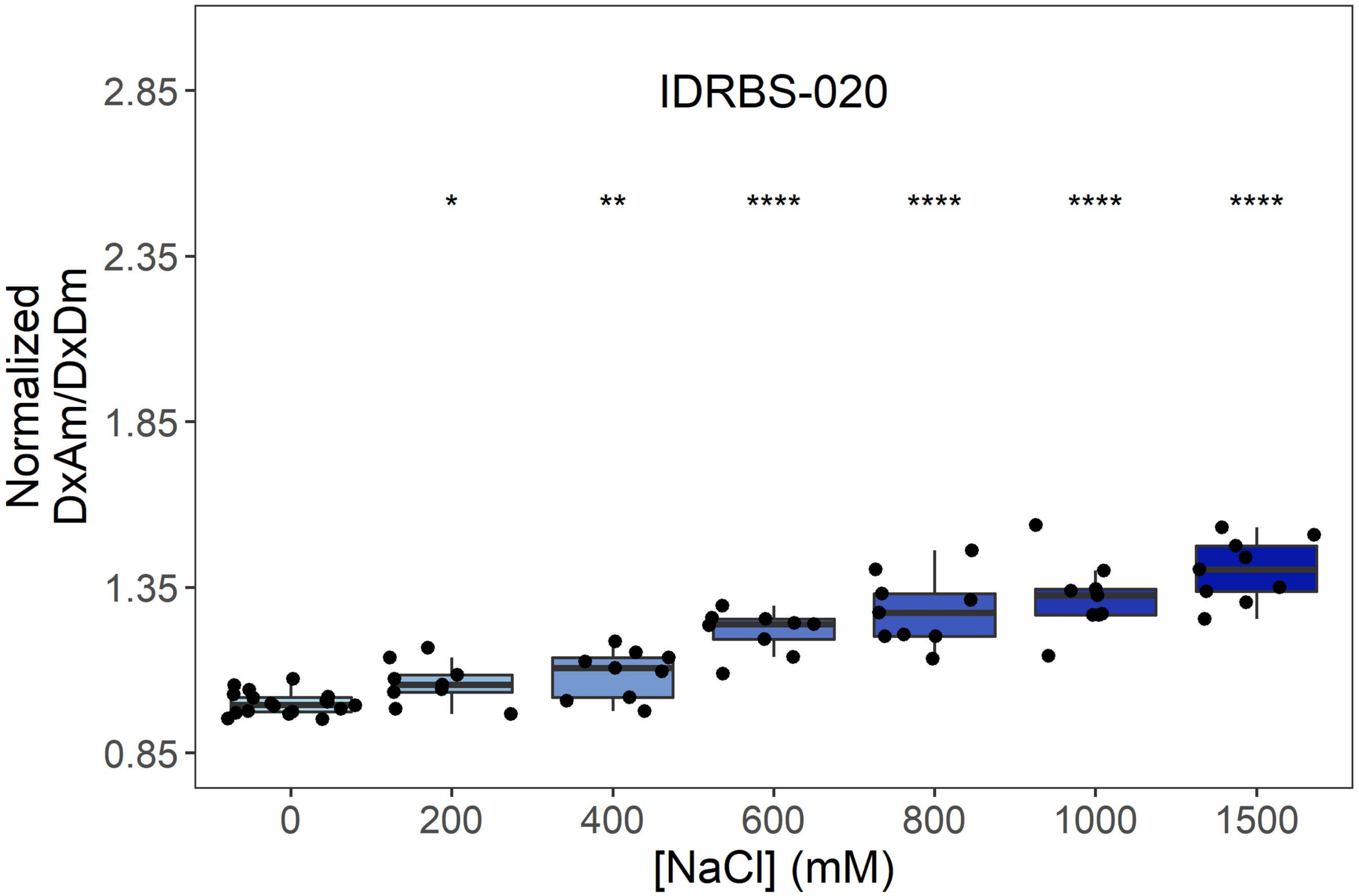
IDRBS-020
*
**
****
****
****
****
2.85
2.35
1.85
1.35
0.85
Normalized
DxAm/DxDm
0
200
400
600
800
1000
1500
[NaCl] (mM)

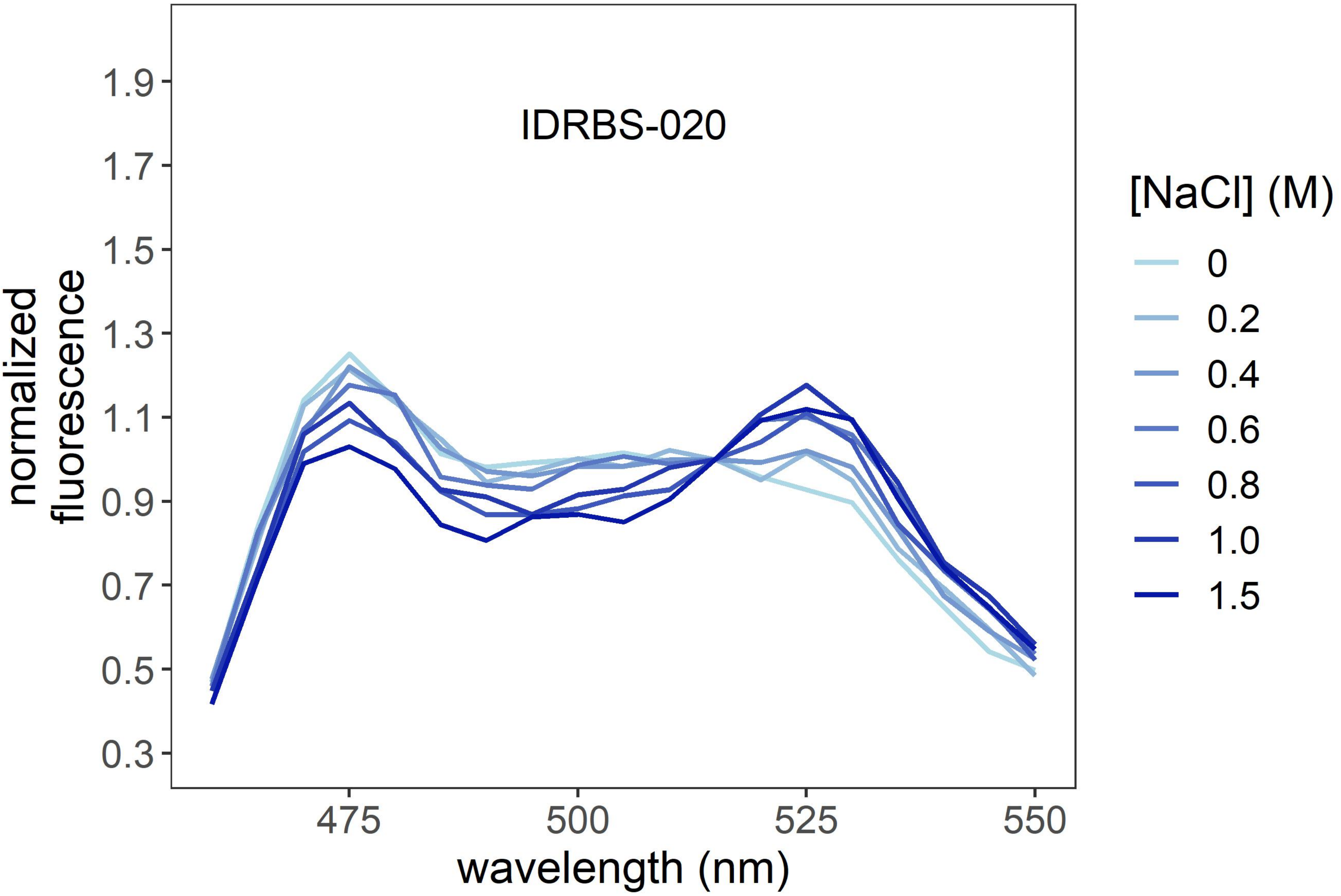
IDRBS-020
1.9
1.7
1.5
1.3
1.1
0.9
0.7
0.5
0.3
normalized
fluorescence
475
500
525
550
wavelength (nm)
[NaCl] (M)
0
0.2
0.4
0.6
0.8
1.0
1.5

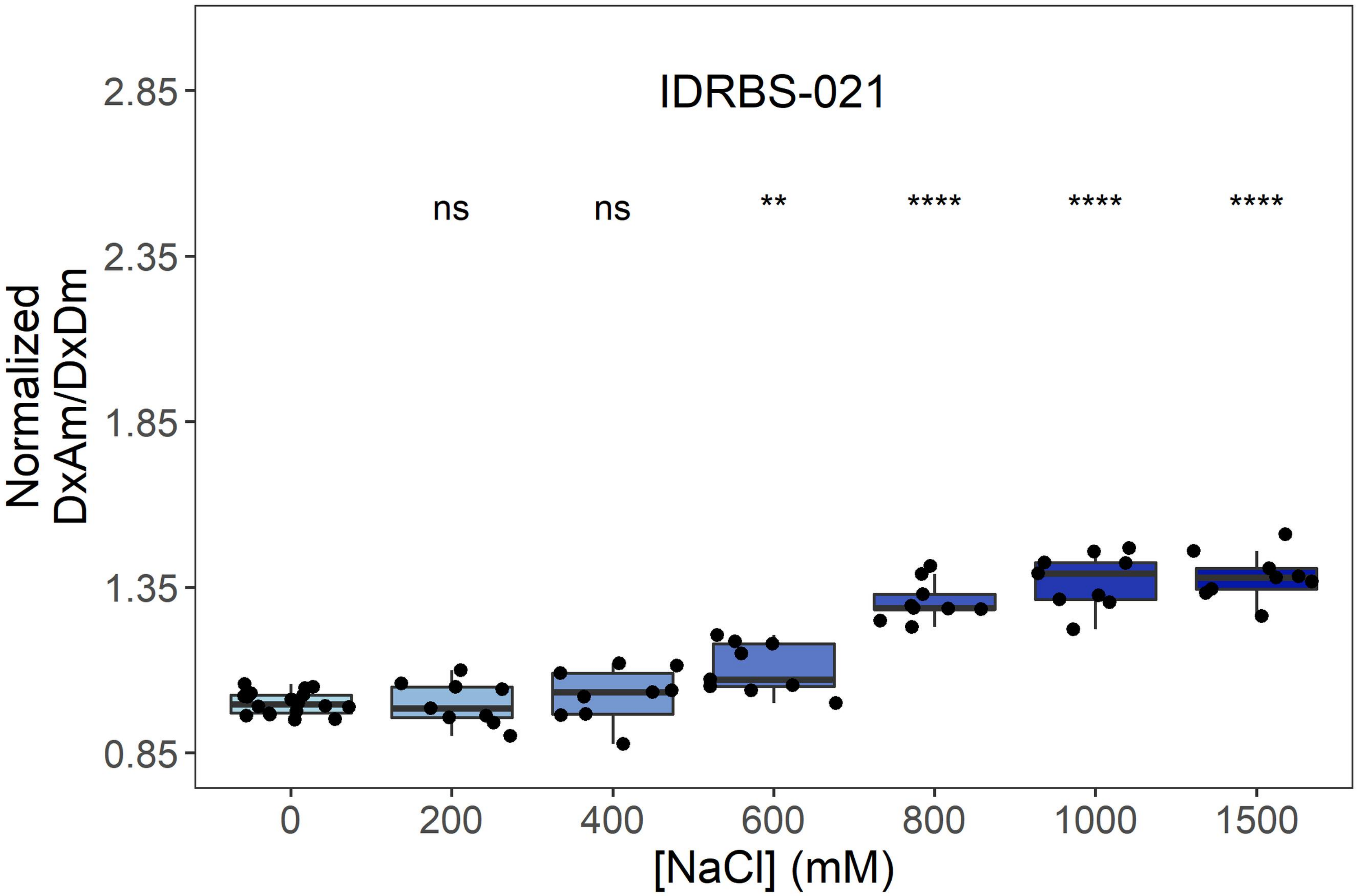
IDRBS-021
ns
ns
**
****
****
****
Normalized
DxAm/DxDm
2.85
2.35
1.85
1.35
0.85
0
200
400
600
800
1000
1500
[NaCl] (mM)

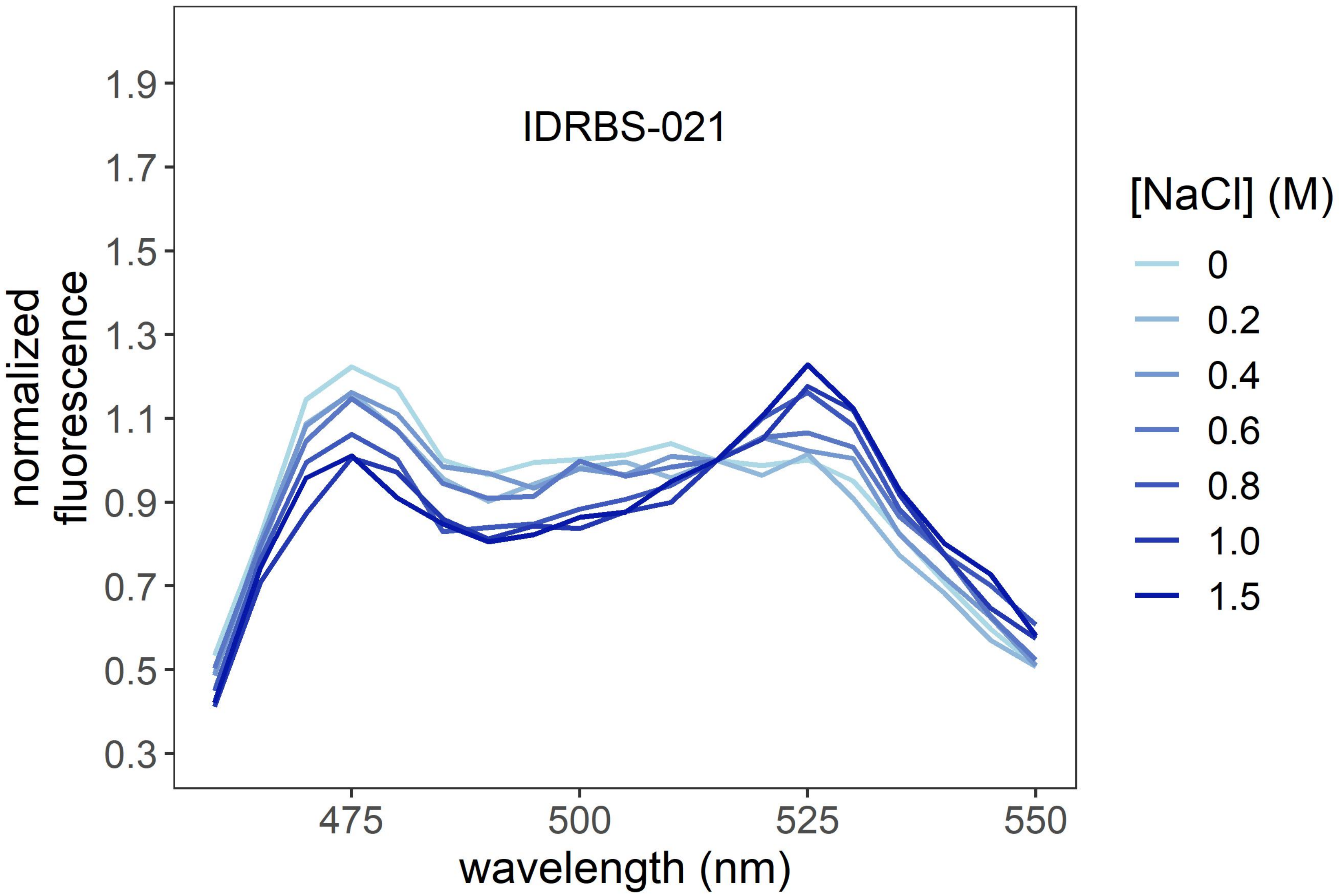
IDRBS-021
normalized
fluorescence
1.9
1.7
1.5
1.3
1.1
0.9
0.7
0.5
0.3
475
500
525
550
wavelength (nm)
[NaCl] (M)
0
0.2
0.4
0.6
0.8
1.0
1.5

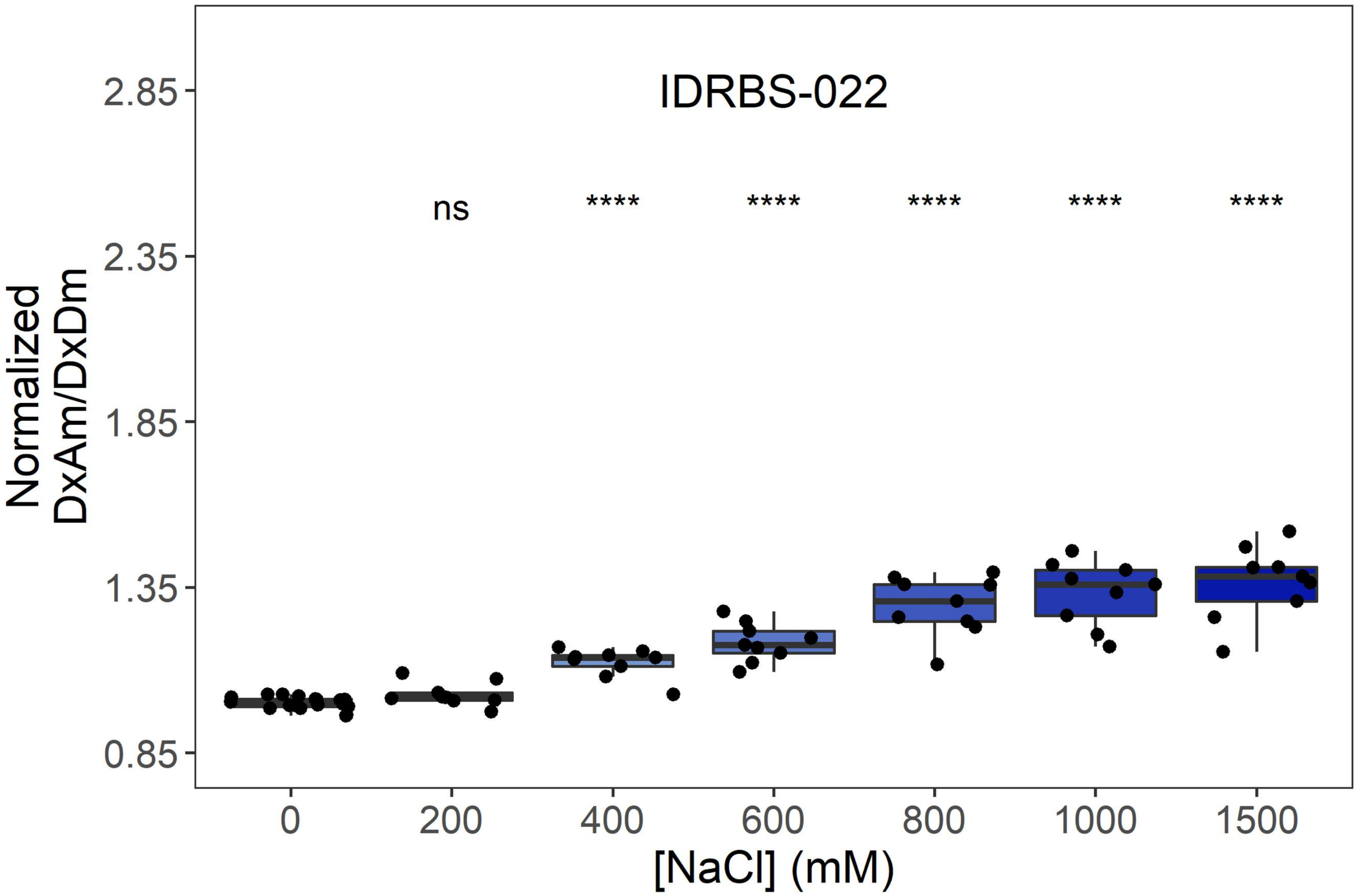

IDRBS-022
ns
****
****
****
****
****
Normalized DxAm/DxDm
2.85
2.35
1.85
1.35
0.85
0
200
400
600
800
1000
1500
[NaCl] (mM)


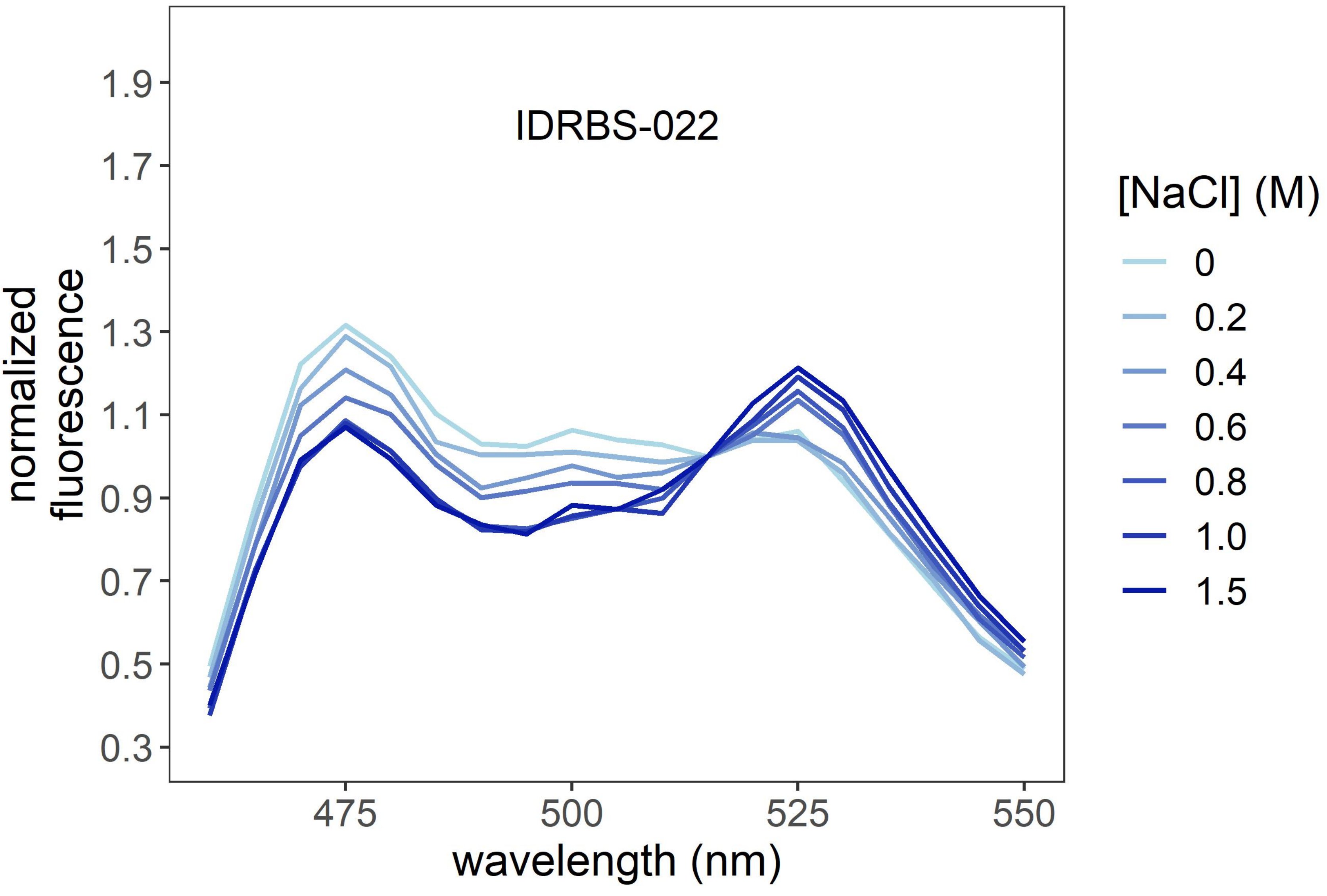

IDRBS-022
normalized fluorescence
1.9
1.7
1.5
1.3
1.1
0.9
0.7
0.5
0.3
475
500
525
550
wavelength (nm)
[NaCl] (M)
0
0.2
0.4
0.6
0.8
1.0
1.5

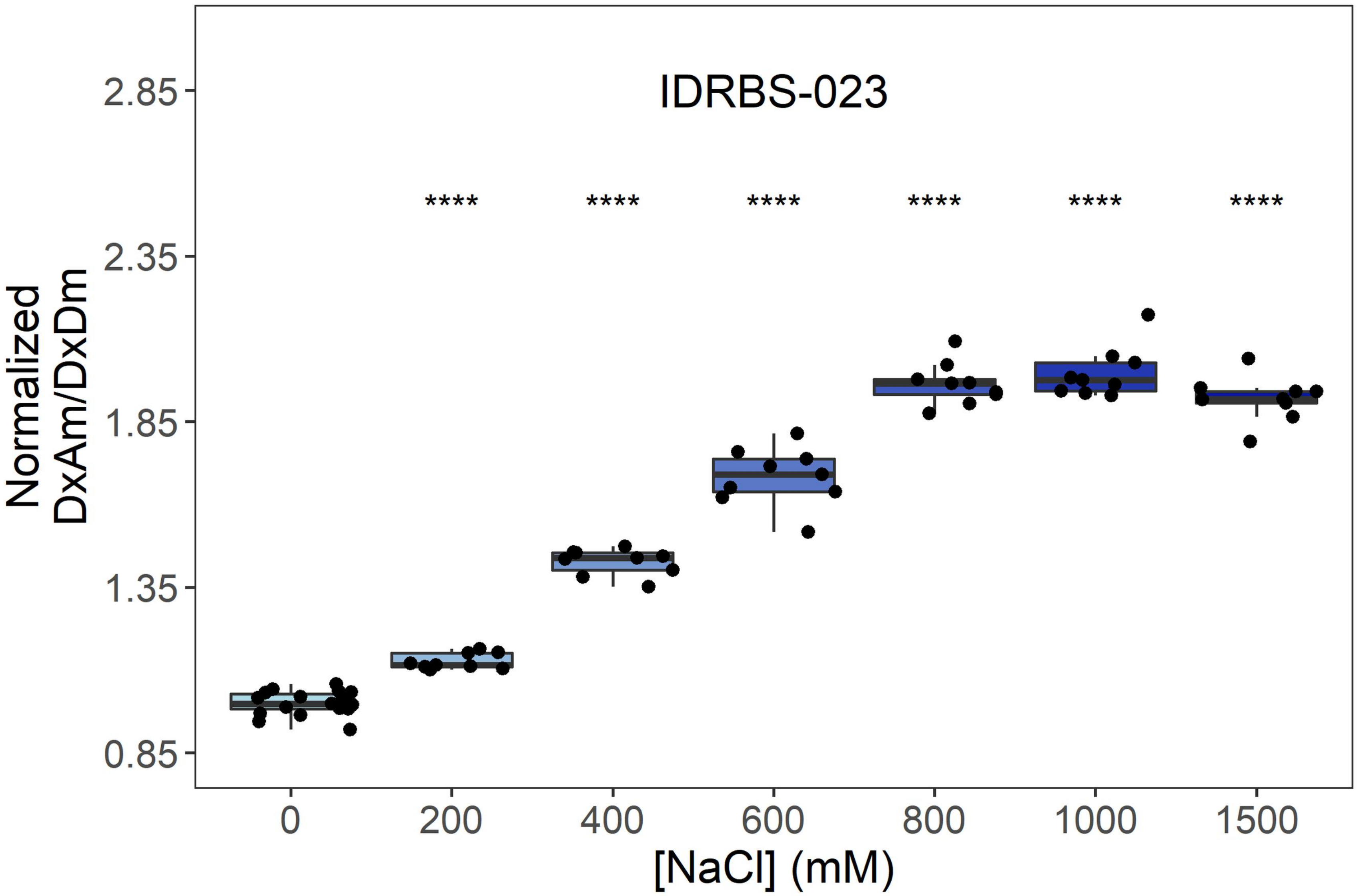

IDRBS-023
****
Normalized
DxAm/DxDm
[NaCl] (mM)
0.85
1.35
1.85
2.35
2.85
0
200
400
600
800
1000
1500


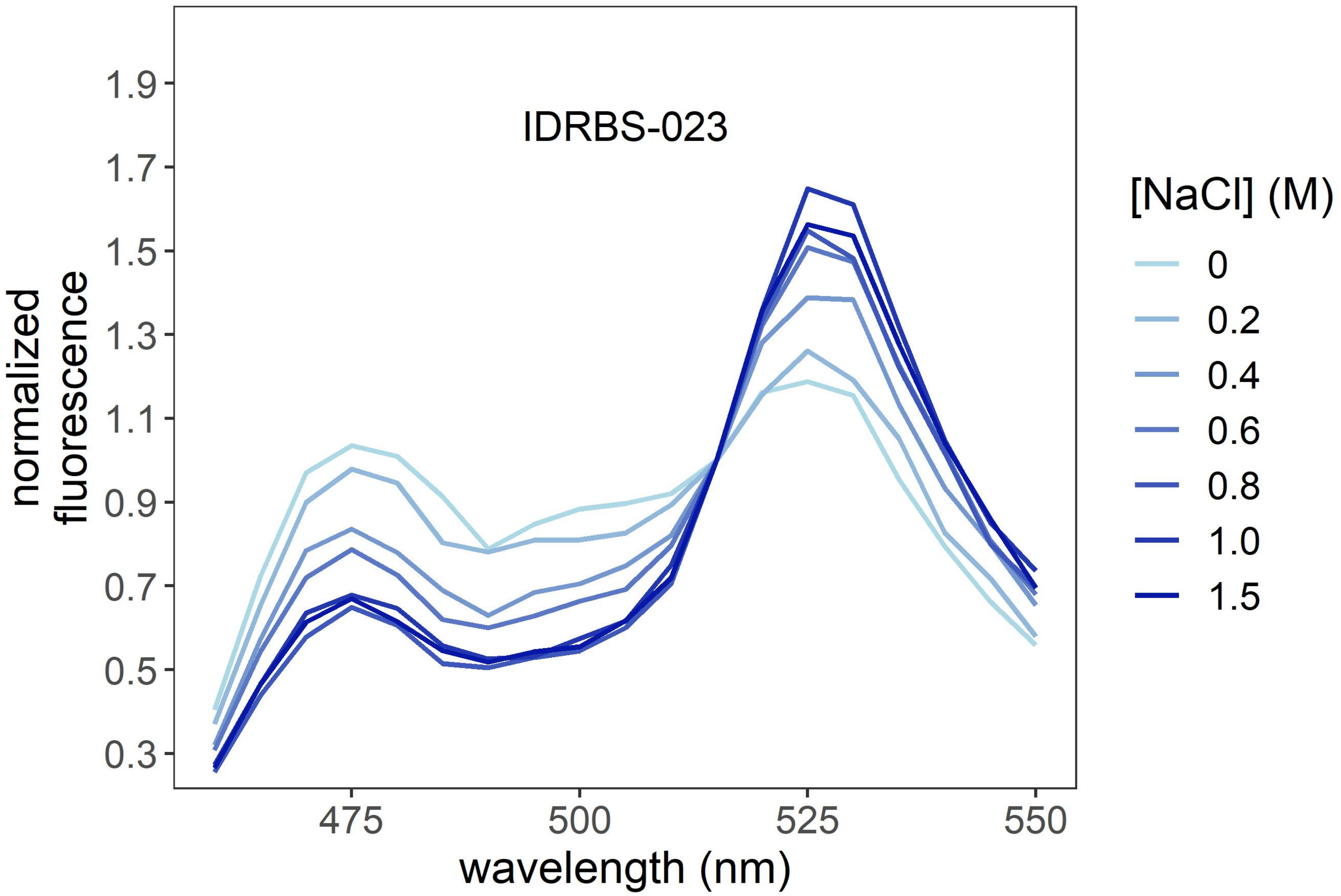

IDRBS-023
normalized
fluorescence
wavelength (nm)
[NaCl] (M)
0
0.2
0.4
0.6
0.8
1.0
1.5
0.3
0.5
0.7
0.9
1.1
1.3
1.5
1.7
1.9
475
500
525
550

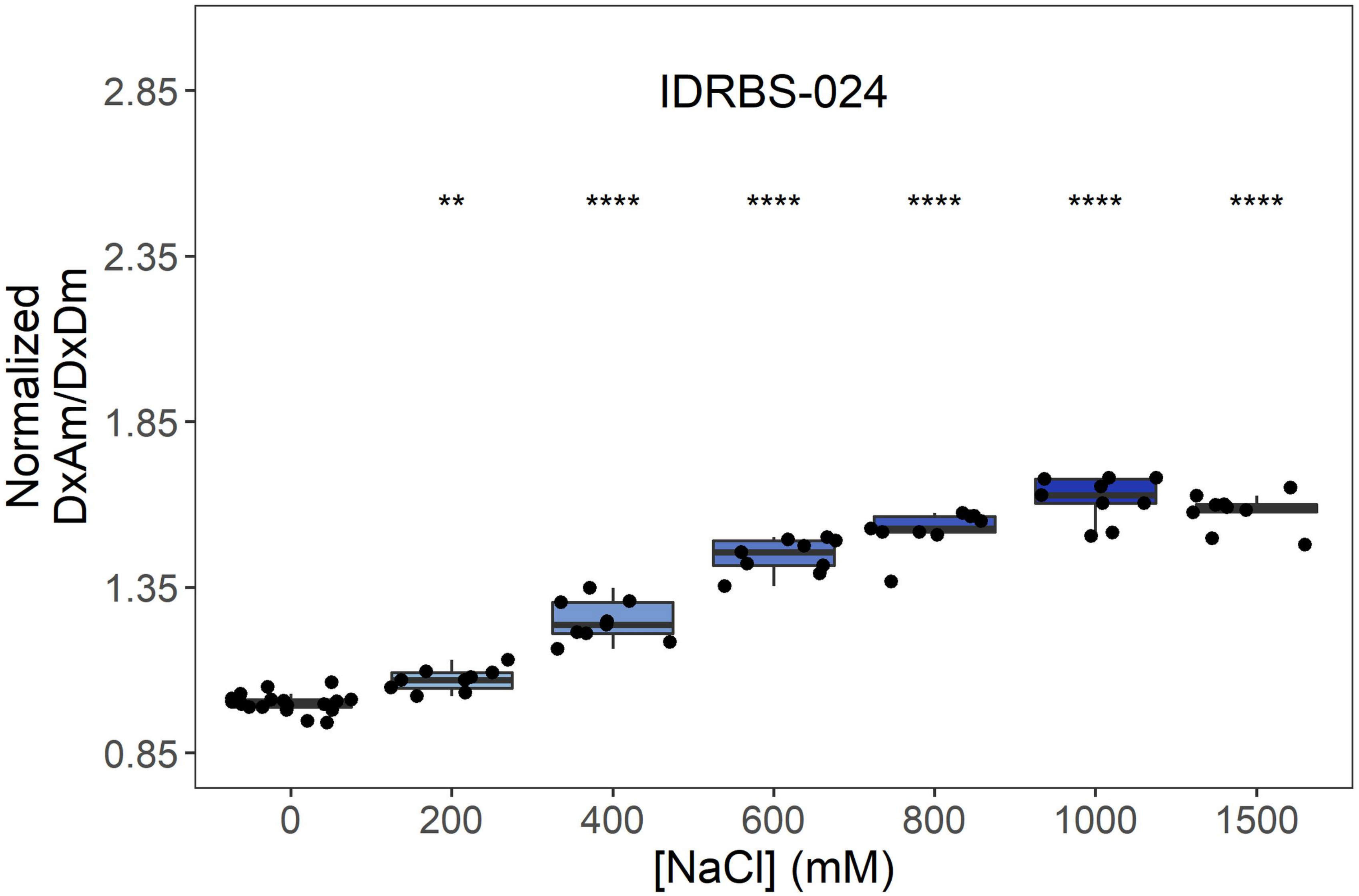
IDRBS-024
Normalized
DxAm/DxDm
2.85
2.35
1.85
1.35
0.85
**
****
****
****
****
****
0
200
400
600
800
1000
1500
[NaCl] (mM)

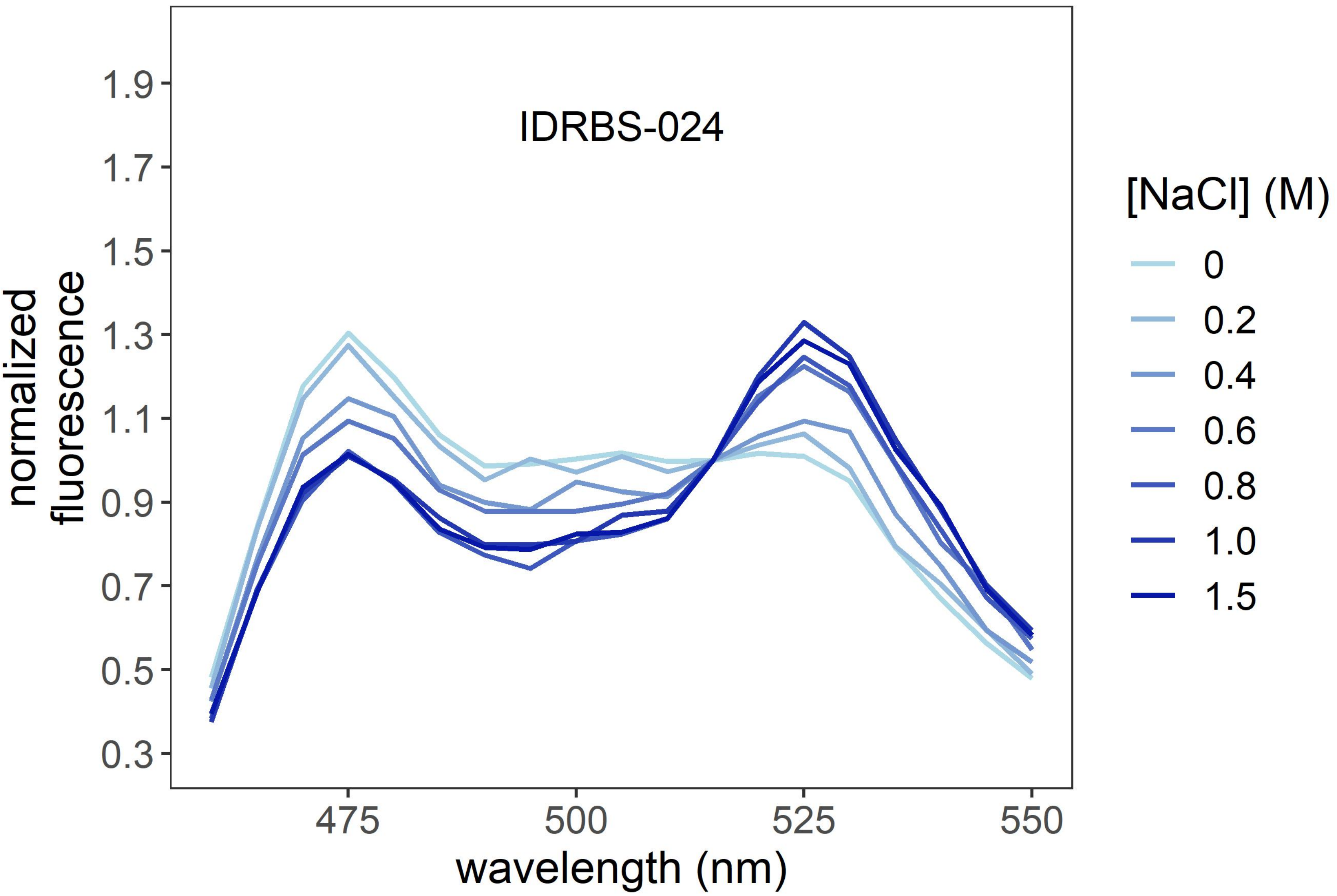
IDRBS-024
normalized
fluorescence
1.9
1.7
1.5
1.3
1.1
0.9
0.7
0.5
0.3
475
500
525
550
wavelength (nm)
[NaCl] (M)
0
0.2
0.4
0.6
0.8
1.0
1.5

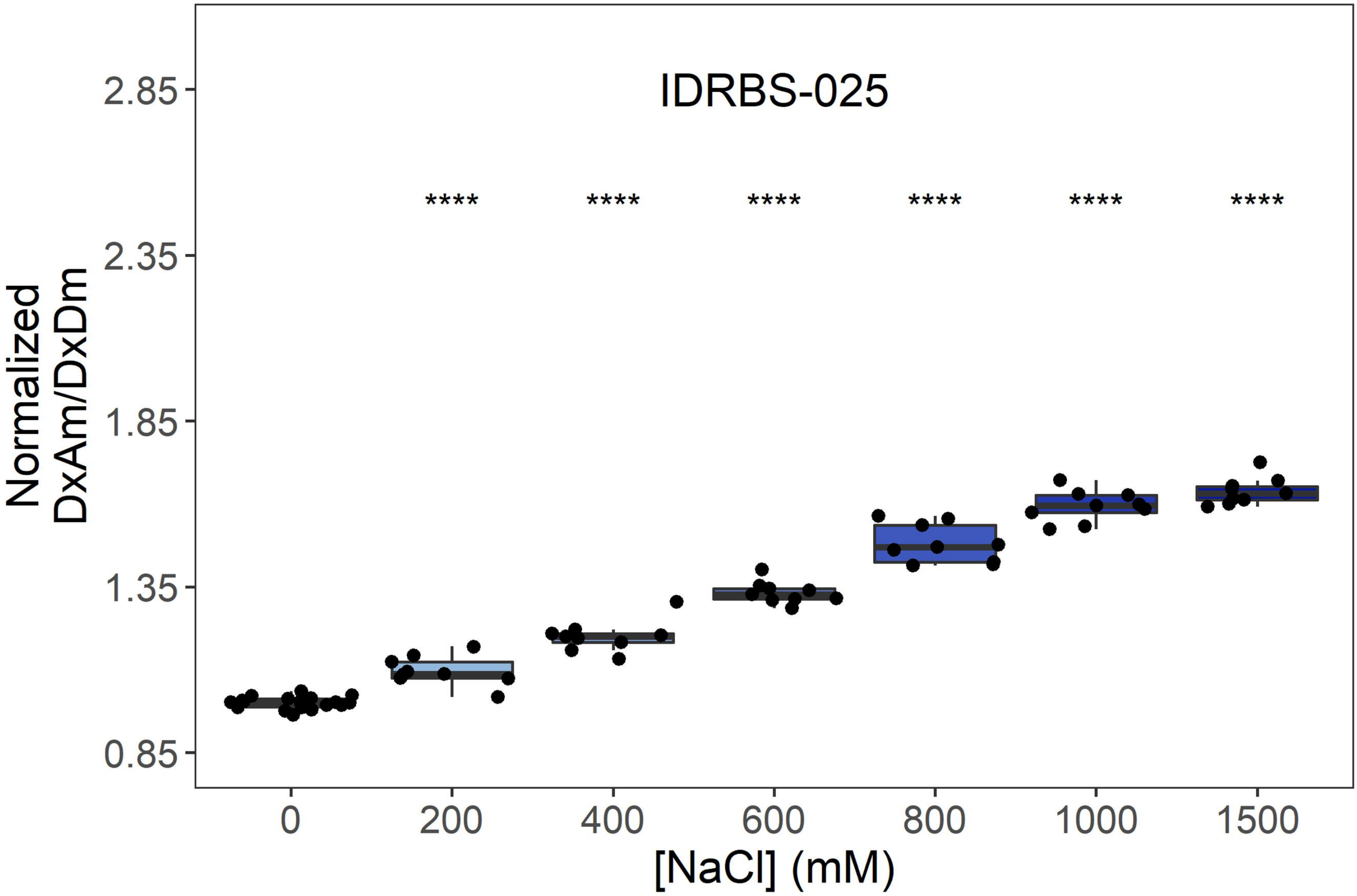
IDRBS-025
Normalized DxAm/DxDm
[NaCl] (mM)
0
200
400
600
800
1000
1500
2.85
2.35
1.85
1.35
0.85
**** **** **** **** **** ****

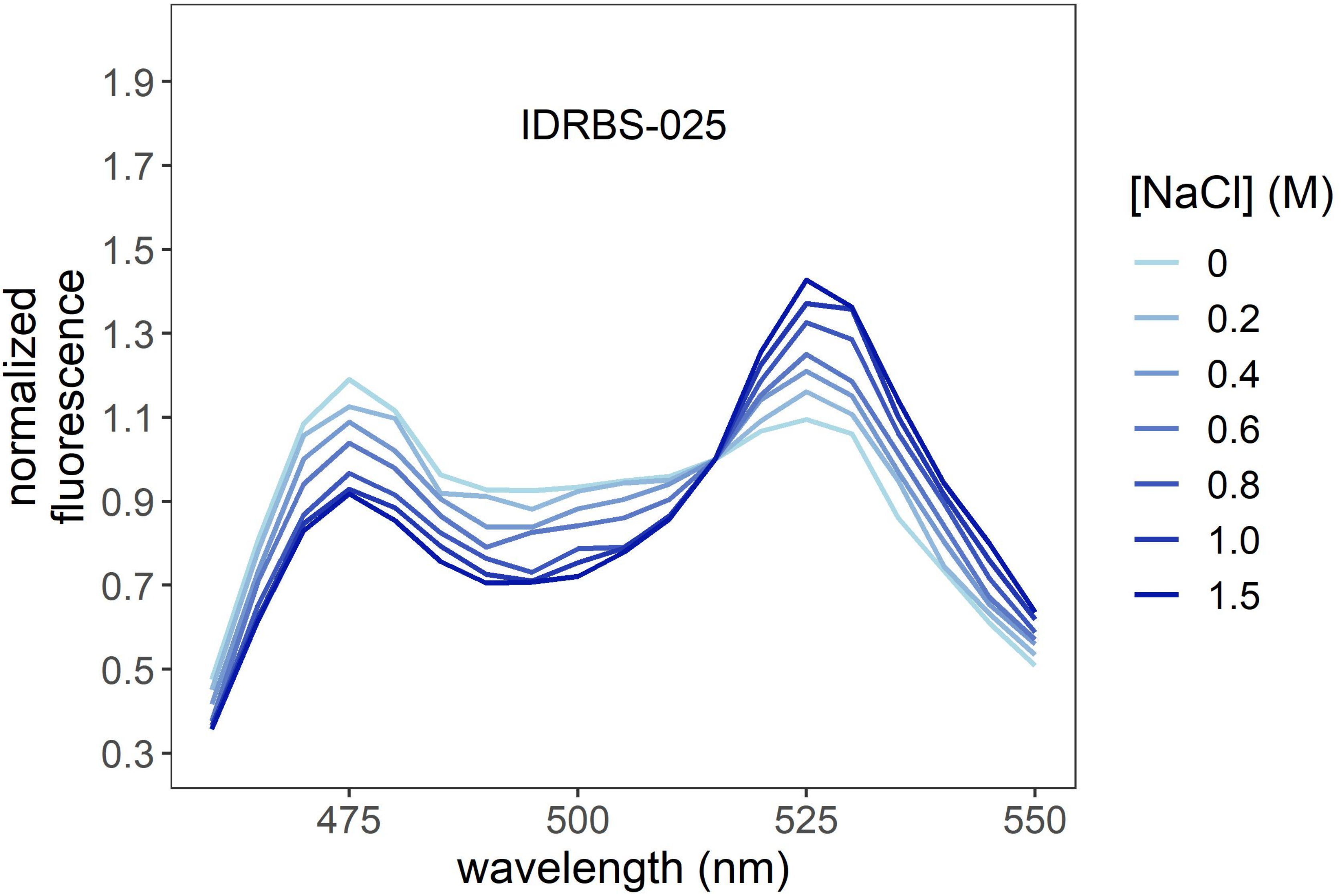
IDRBS-025
normalized fluorescence
wavelength (nm)
475
500
525
550
[NaCl] (M)
0
0.2
0.4
0.6
0.8
1.0
1.5

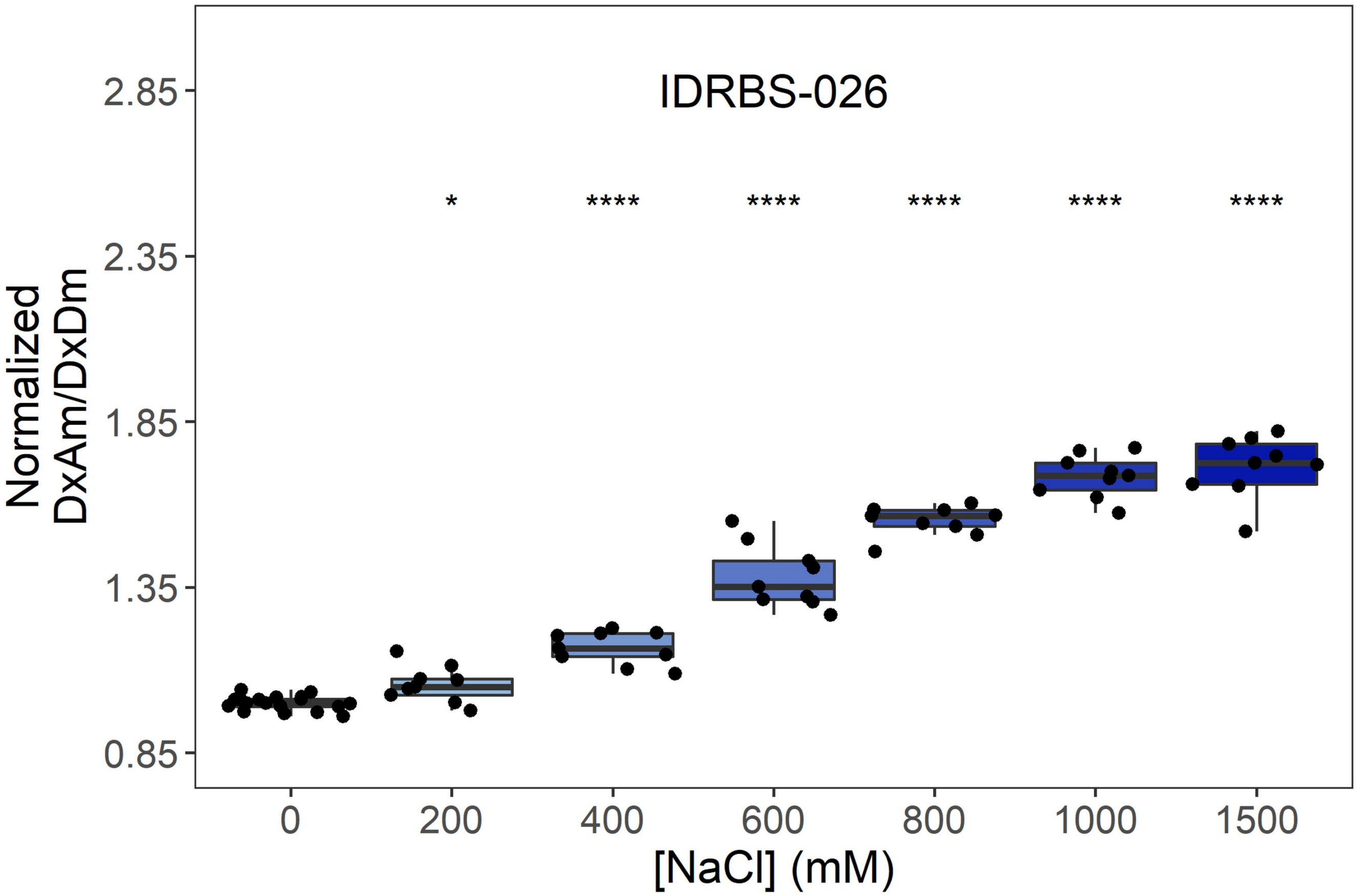
IDRBS-026
*
****
****
****
****
****
Normalized
DxAm/DxDm
2.85
2.35
1.85
1.35
0.85
0
200
400
600
800
1000
1500
[NaCl] (mM)

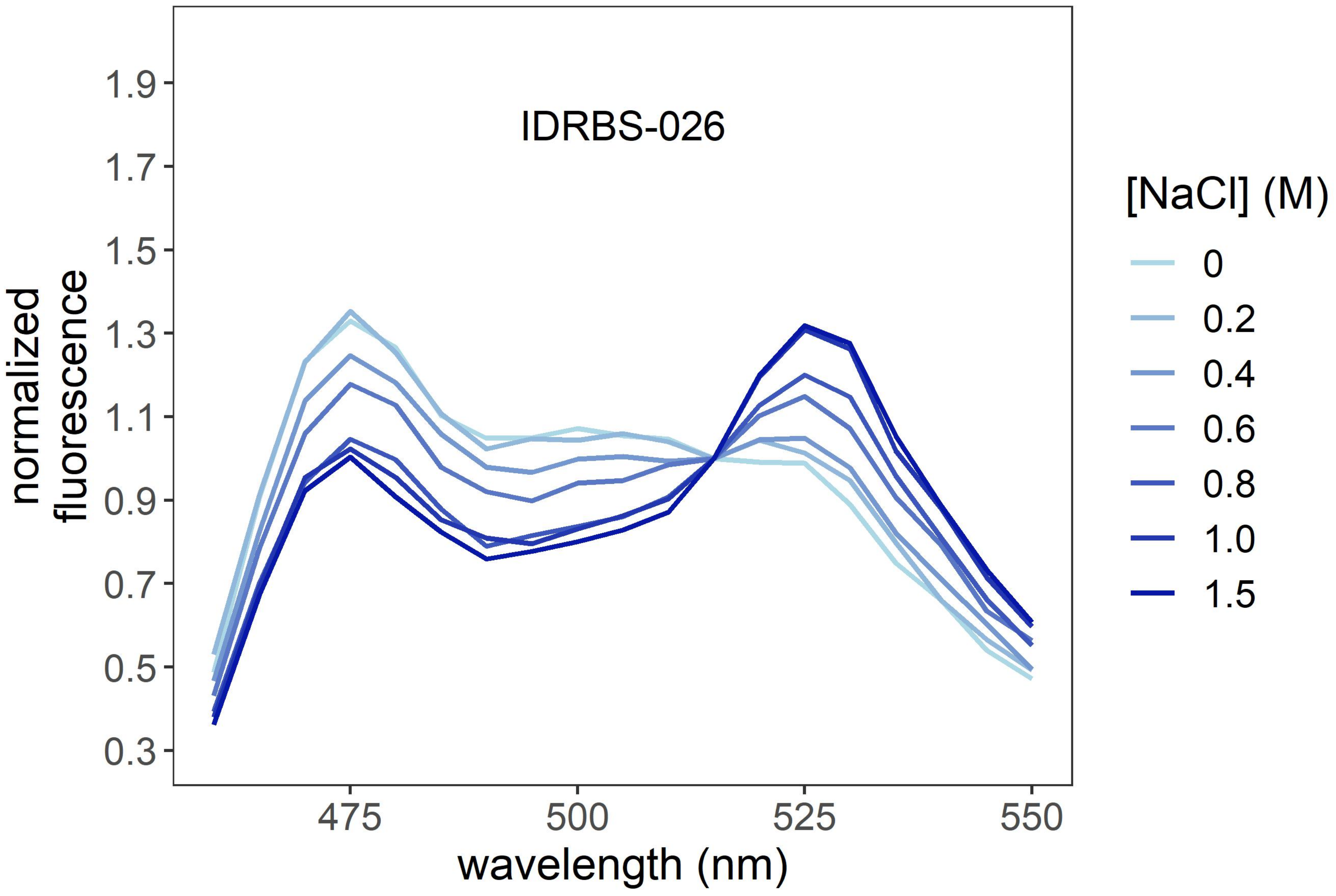
IDRBS-026
normalized
fluorescence
1.9
1.7
1.5
1.3
1.1
0.9
0.7
0.5
0.3
475
500
525
550
wavelength (nm)
[NaCl] (M)
0
0.2
0.4
0.6
0.8
1.0
1.5

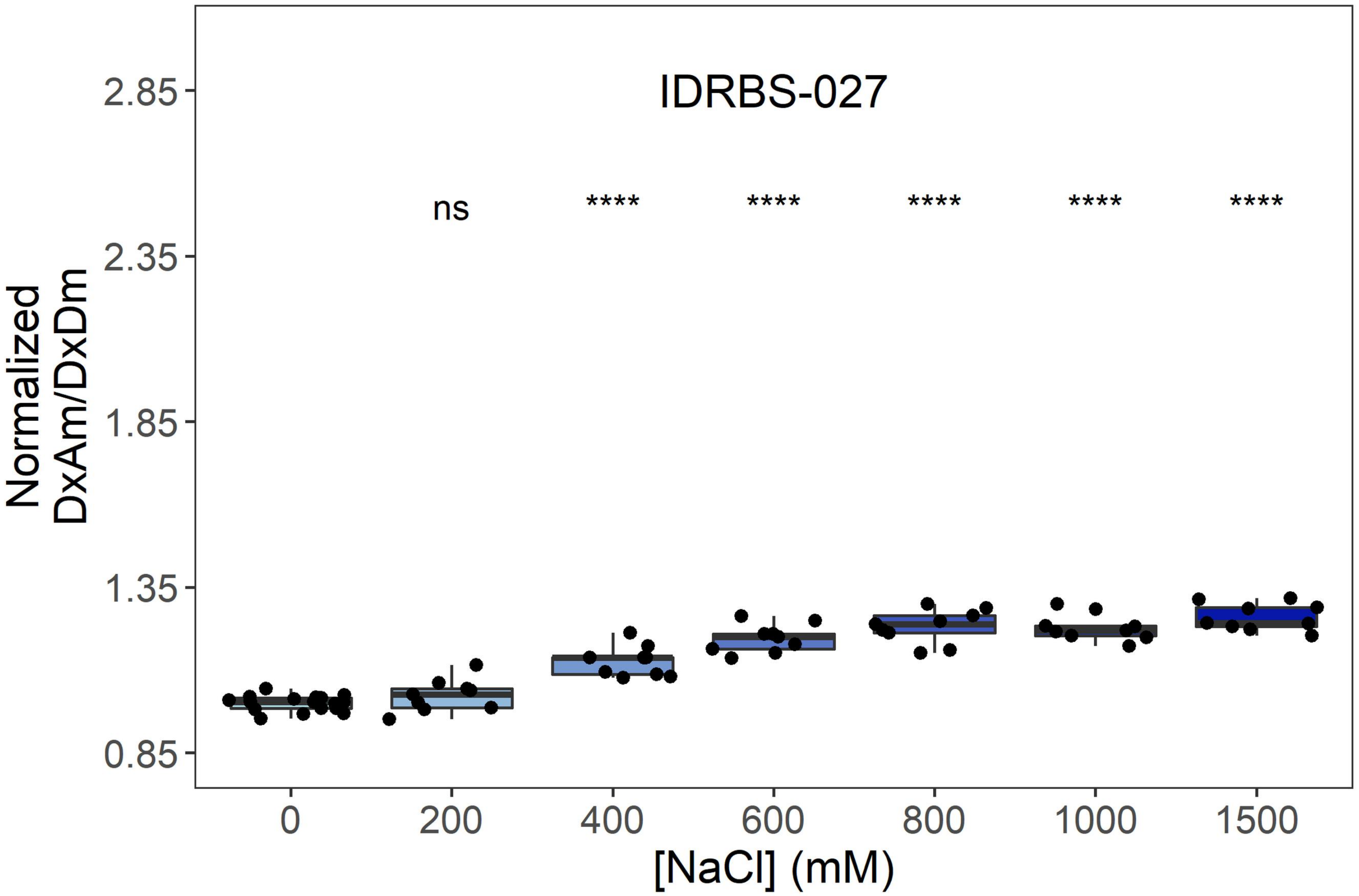
IDRBS-027
ns
****
****
****
****
****
Normalized DxAm/DxDm
2.85
2.35
1.85
1.35
0.85
0
200
400
600
800
1000
1500
[NaCl] (mM)

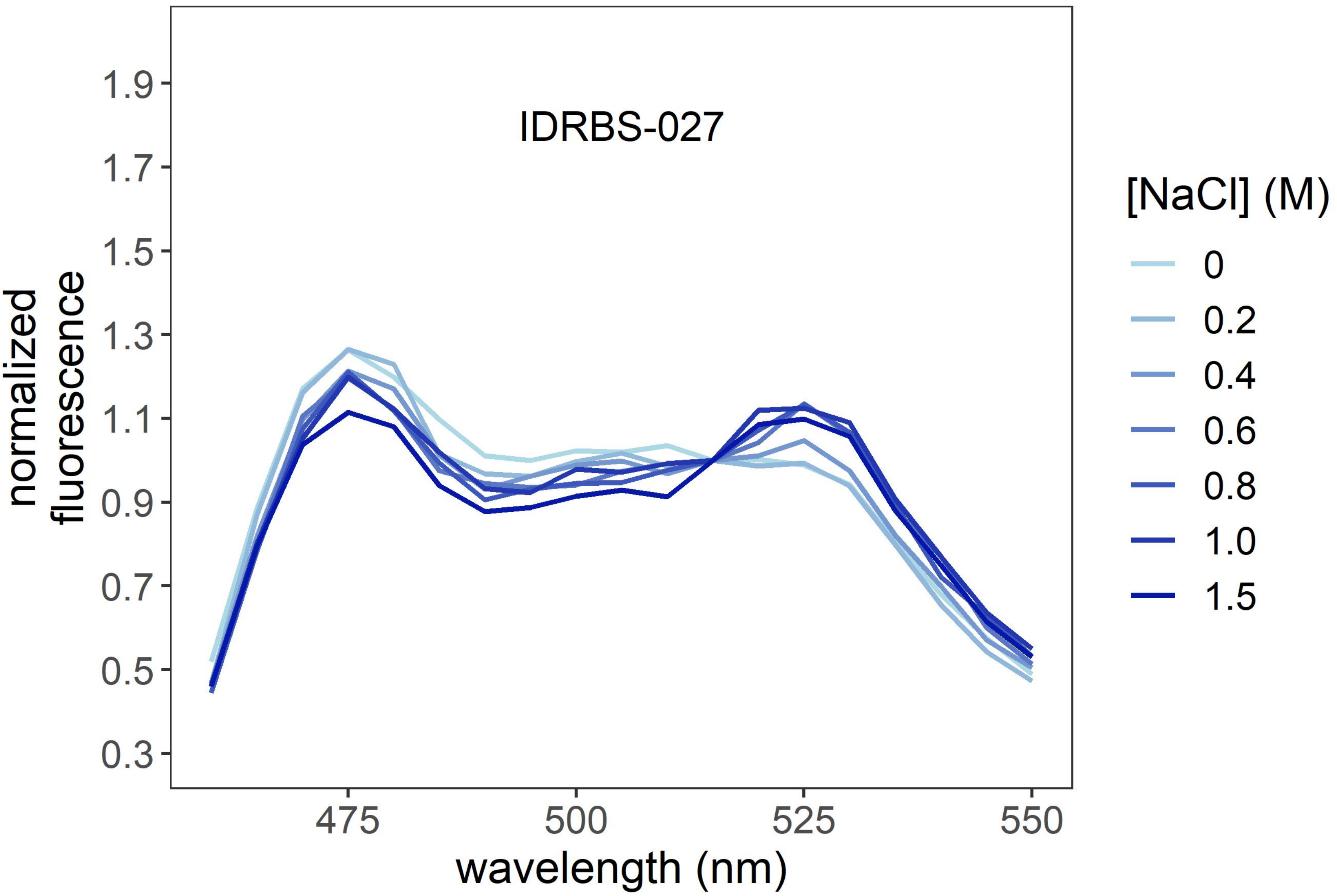
IDRBS-027
normalized fluorescence
1.9
1.7
1.5
1.3
1.1
0.9
0.7
0.5
0.3
475
500
525
550
wavelength (nm)
[NaCl] (M)
0
0.2
0.4
0.6
0.8
1.0
1.5

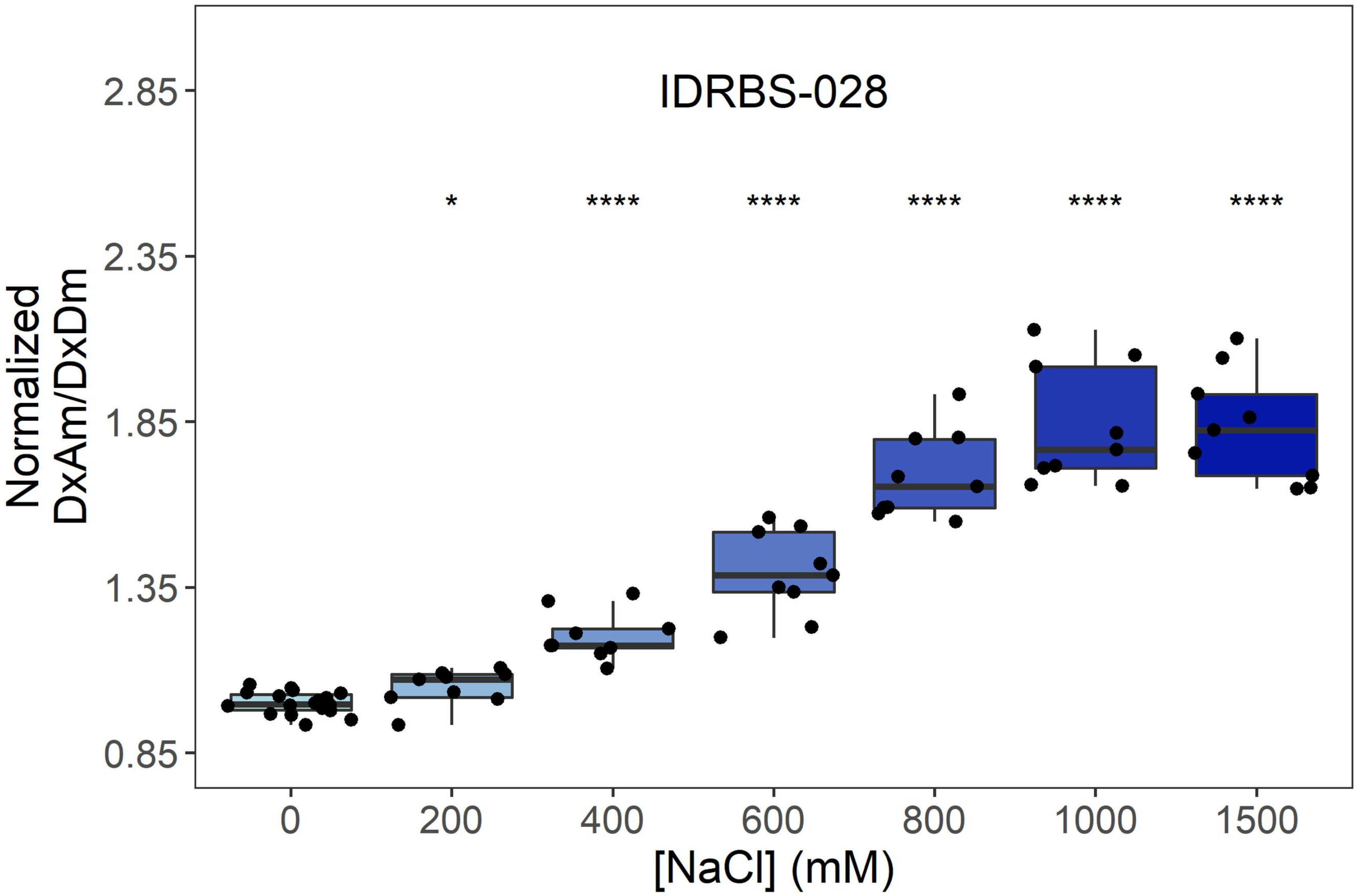
IDRBS-028
*
****
****
****
****
****
Normalized DxAm/DxDm
2.85
2.35
1.85
1.35
0.85
0
200
400
600
800
1000
1500
[NaCl] (mM)

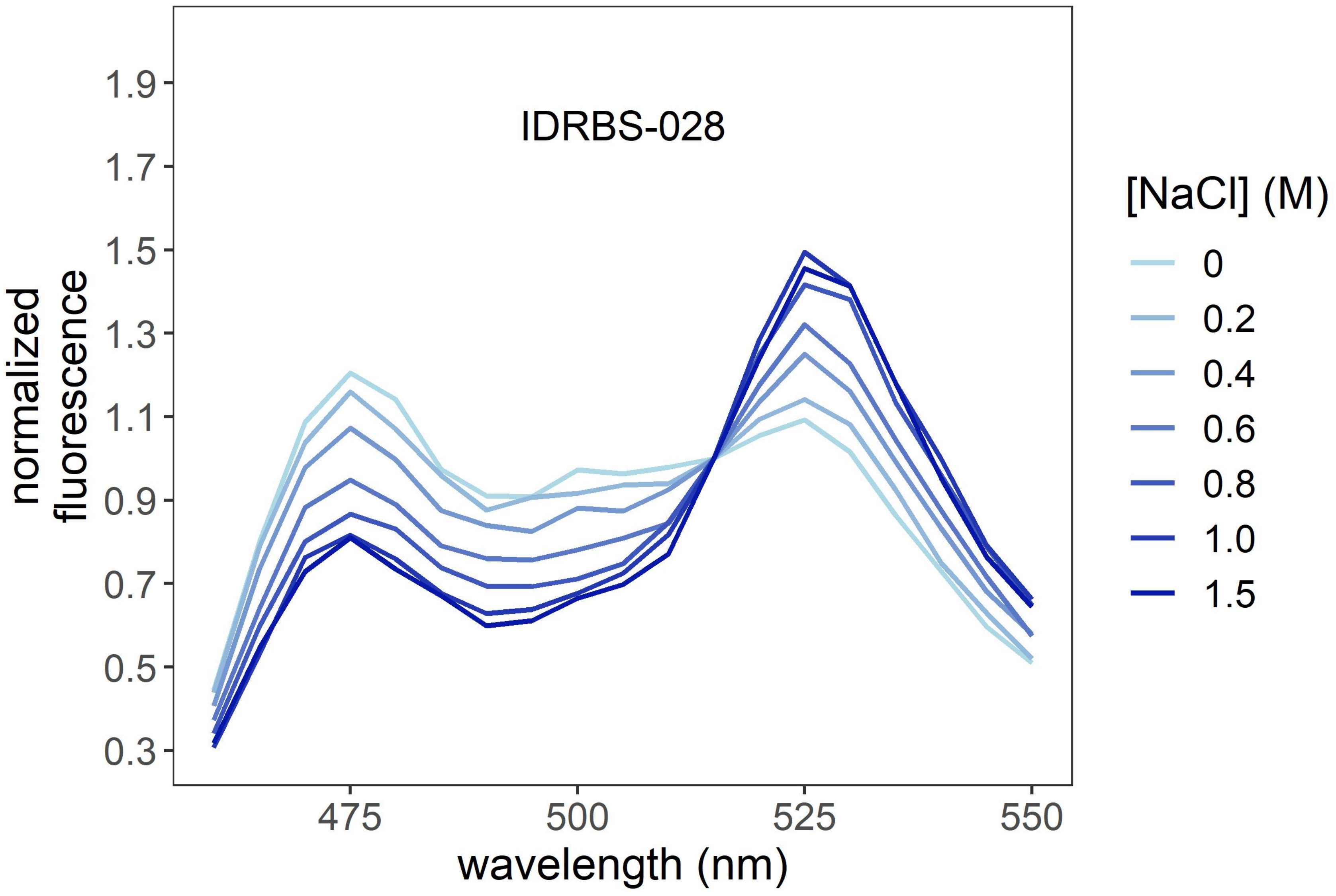
IDRBS-028
normalized fluorescence
1.9
1.7
1.5
1.3
1.1
0.9
0.7
0.5
0.3
475
500
525
550
wavelength (nm)
[NaCl] (M)
0
0.2
0.4
0.6
0.8
1.0
1.5

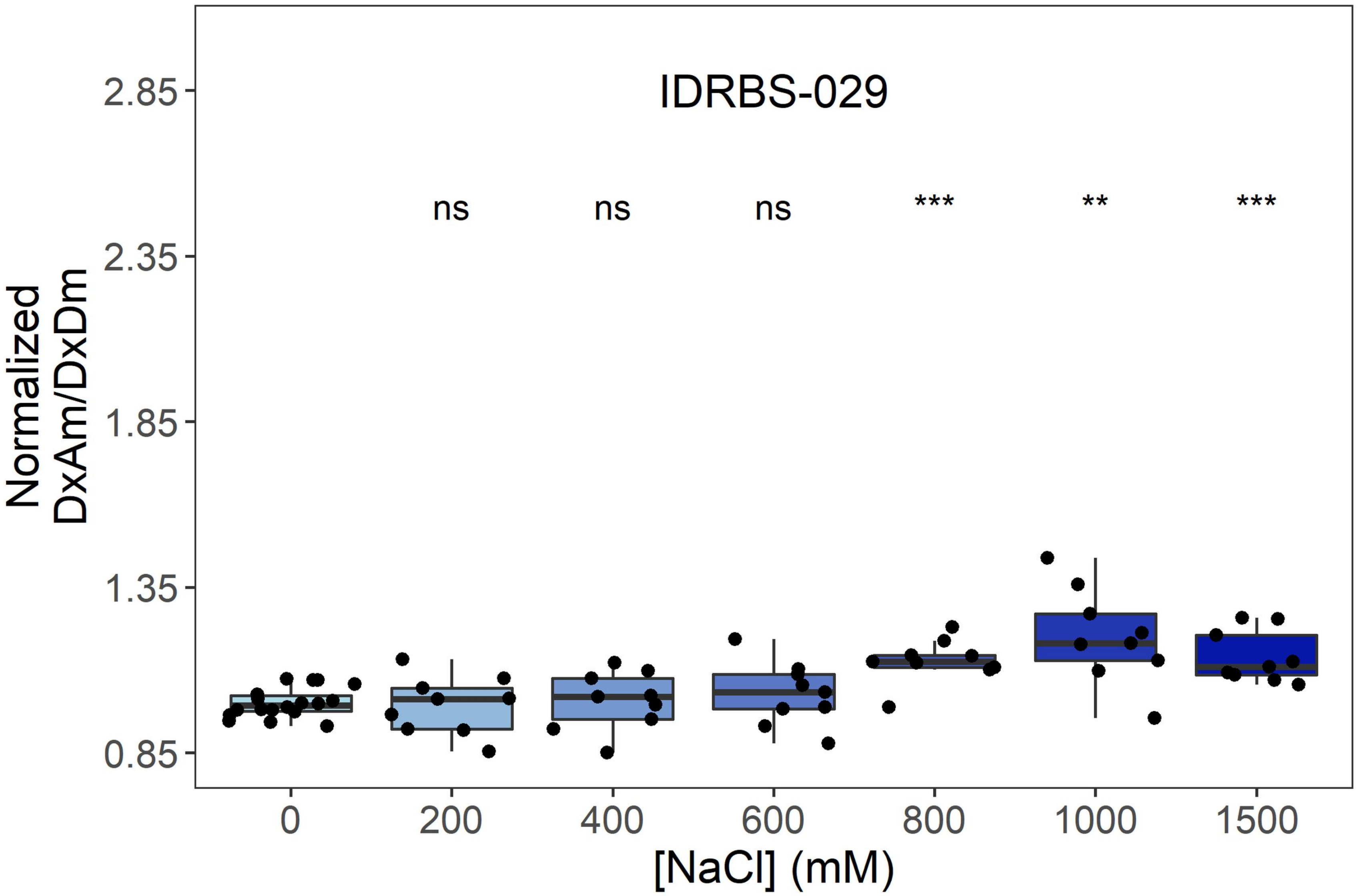

IDRBS-029
ns
ns
ns
***
**
***
Normalized
DxAm/DxDm
2.85
2.35
1.85
1.35
0.85
0
200
400
600
800
1000
1500
[NaCl] (mM)


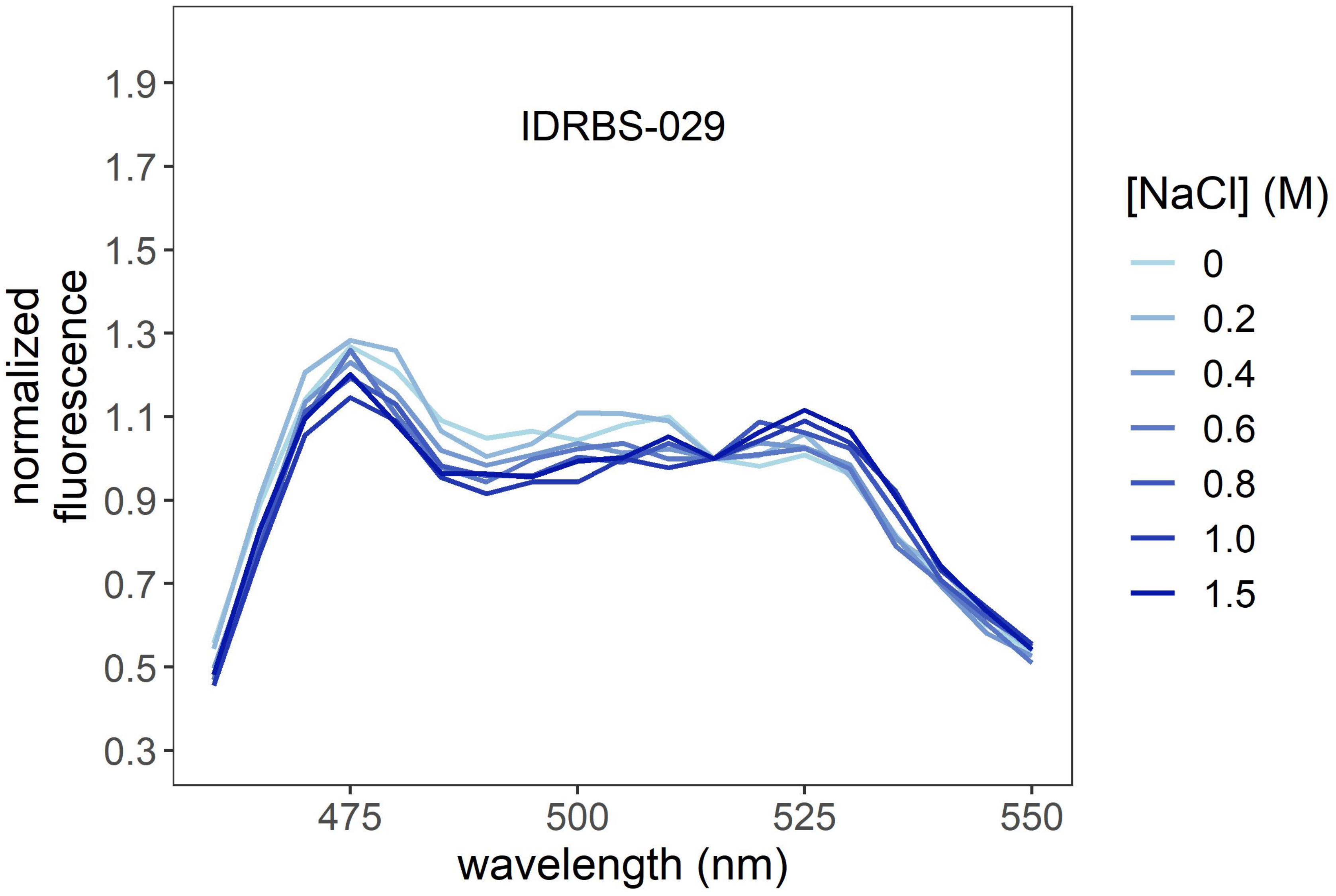

IDRBS-029
normalized
fluorescence
1.9
1.7
1.5
1.3
1.1
0.9
0.7
0.5
0.3
475
500
525
550
wavelength (nm)
[NaCl] (M)
0
0.2
0.4
0.6
0.8
1.0
1.5

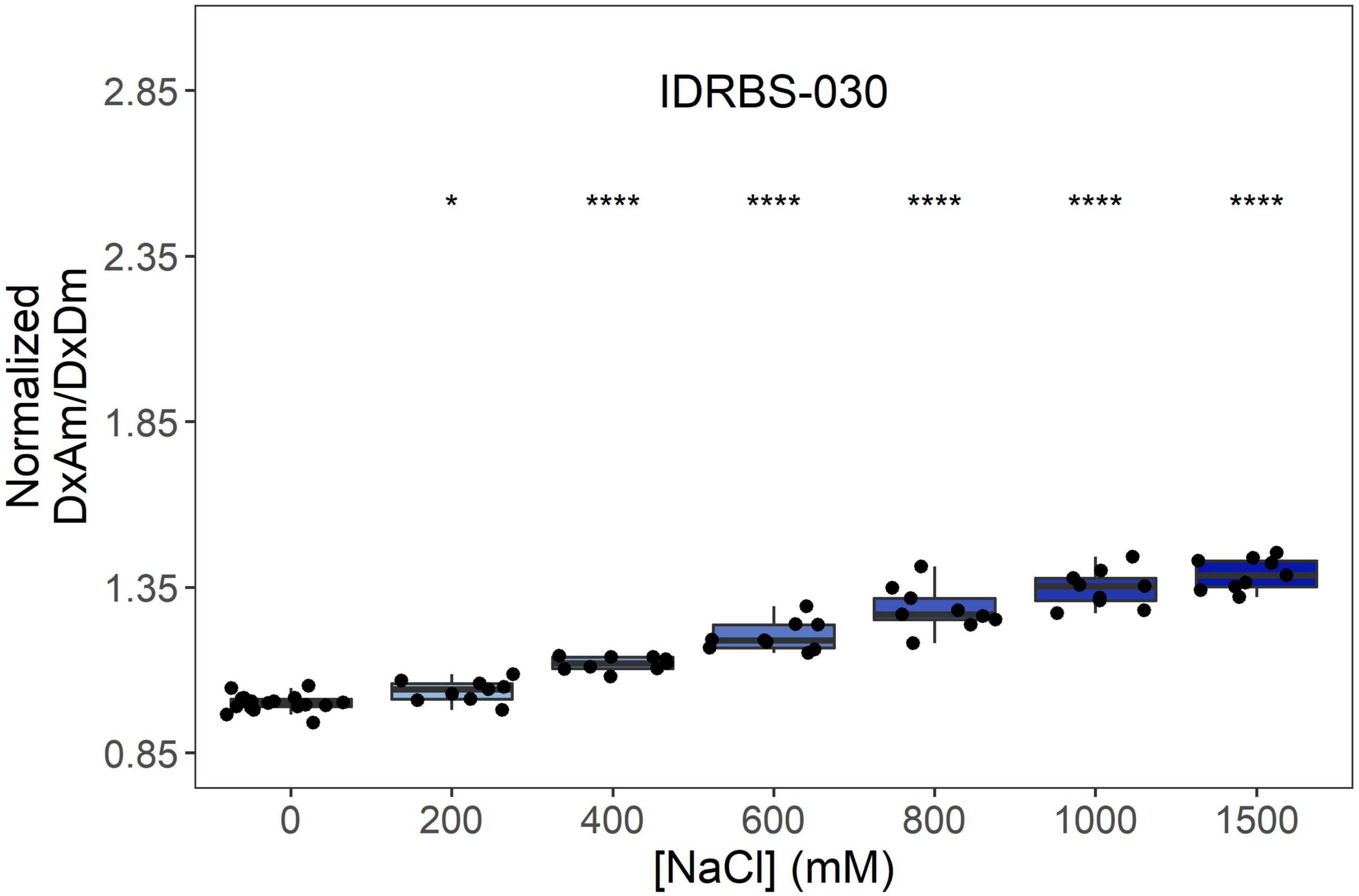

IDRBS-030
*
****
****
****
****
****
2.85
2.35
1.85
1.35
0.85
Normalized DxAm/DxDm
0
200
400
600
800
1000
1500
[NaCl] (mM)


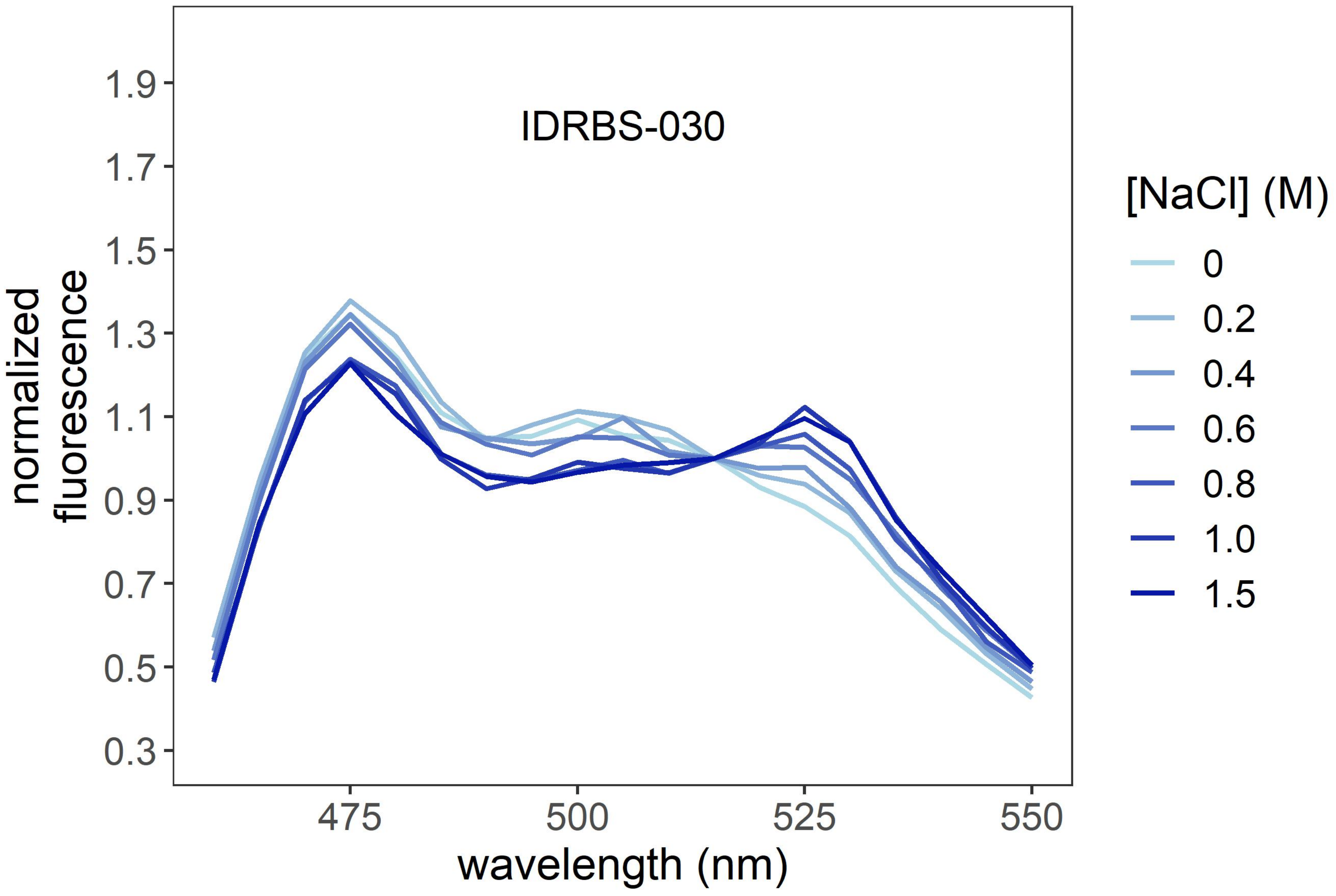

IDRBS-030
1.9
1.7
1.5
1.3
1.1
0.9
0.7
0.5
0.3
normalized fluorescence
475
500
525
550
wavelength (nm)
[NaCl] (M)
0
0.2
0.4
0.6
0.8
1.0
1.5

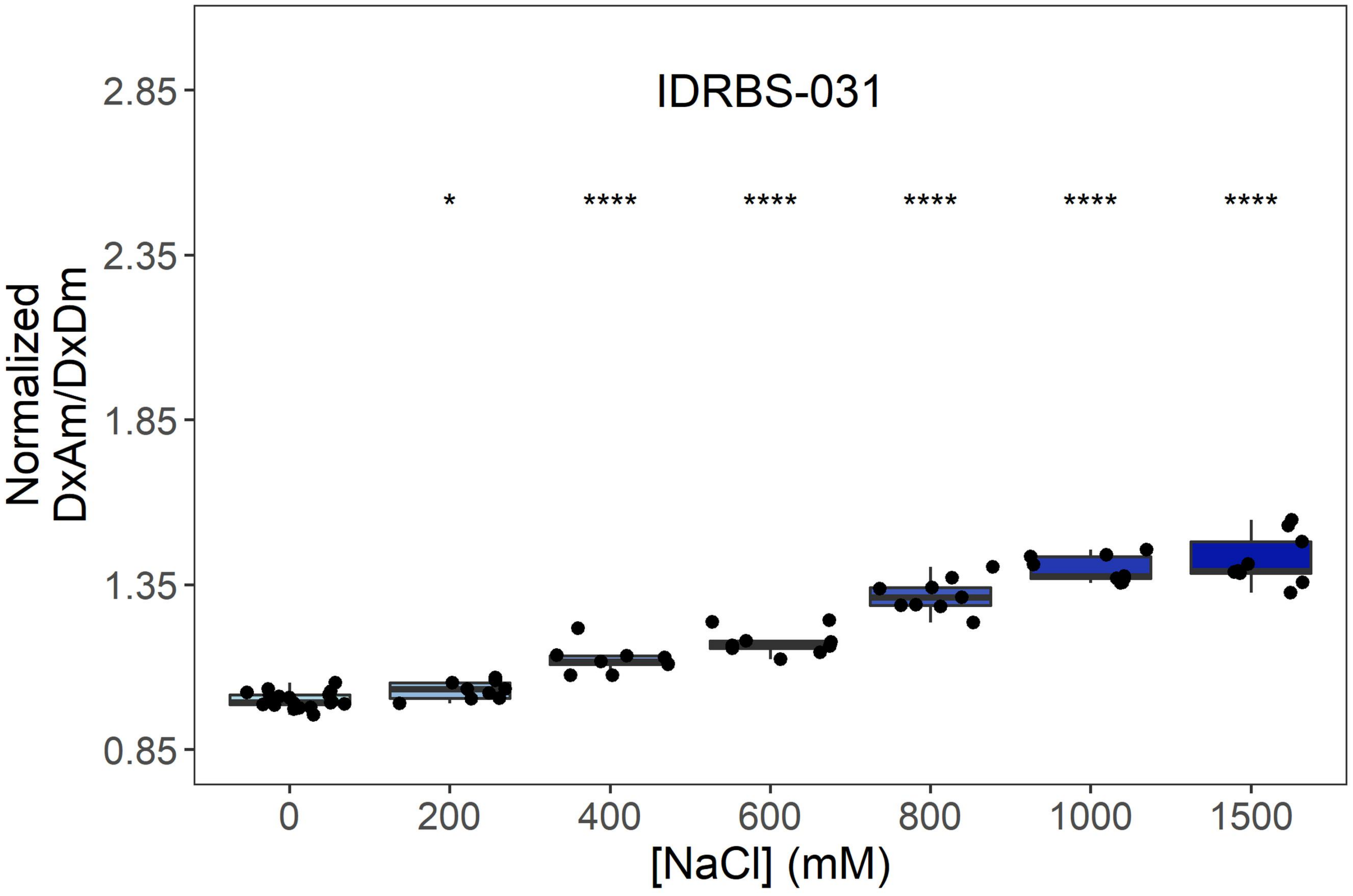
IDRBS-031
*
****
****
****
****
****
Normalized
DxAm/DxDm
2.85
2.35
1.85
1.35
0.85
0
200
400
600
800
1000
1500
[NaCl] (mM)

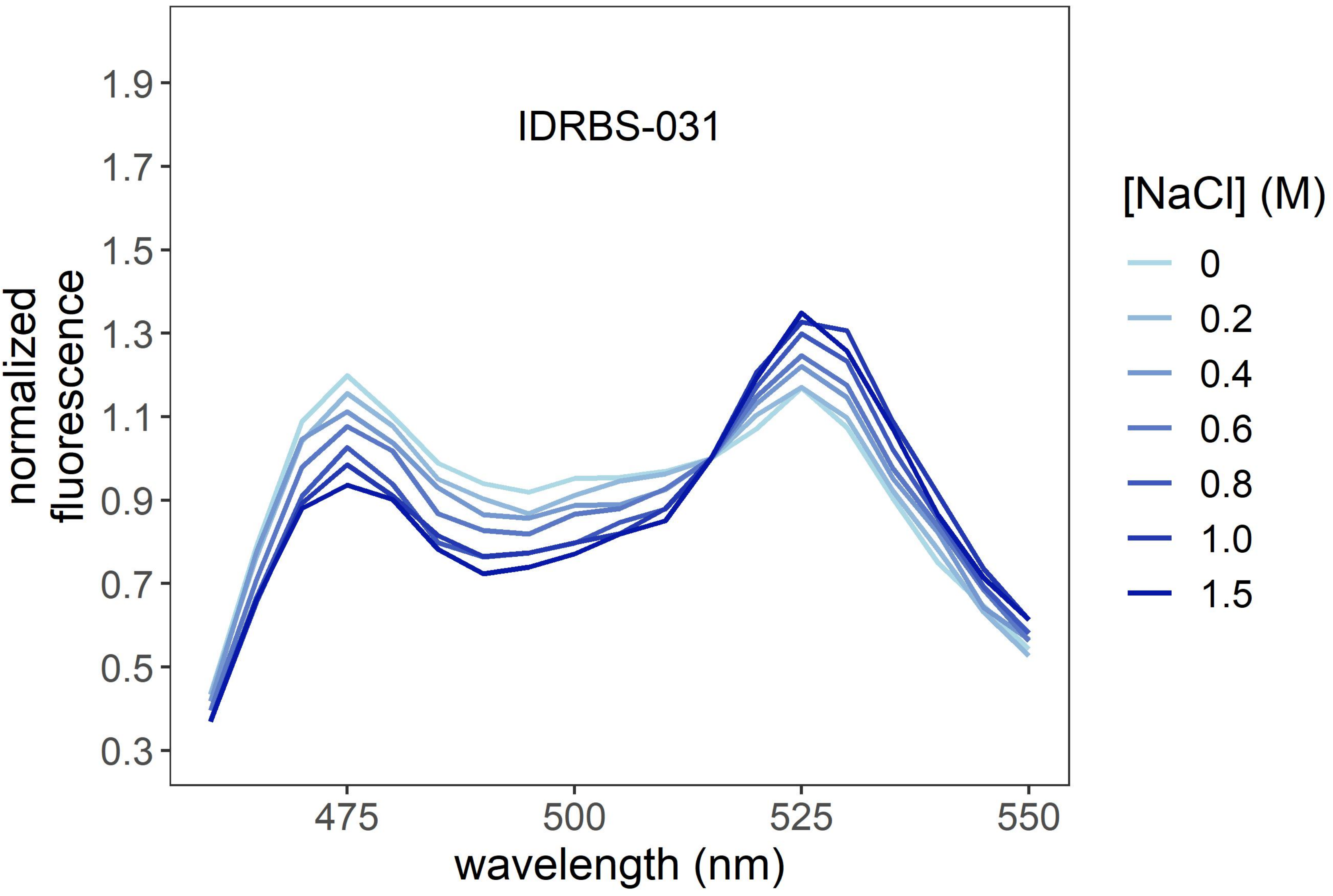
IDRBS-031
normalized
fluorescence
1.9
1.7
1.5
1.3
1.1
0.9
0.7
0.5
0.3
475
500
525
550
wavelength (nm)
[NaCl] (M)
0
0.2
0.4
0.6
0.8
1.0
1.5

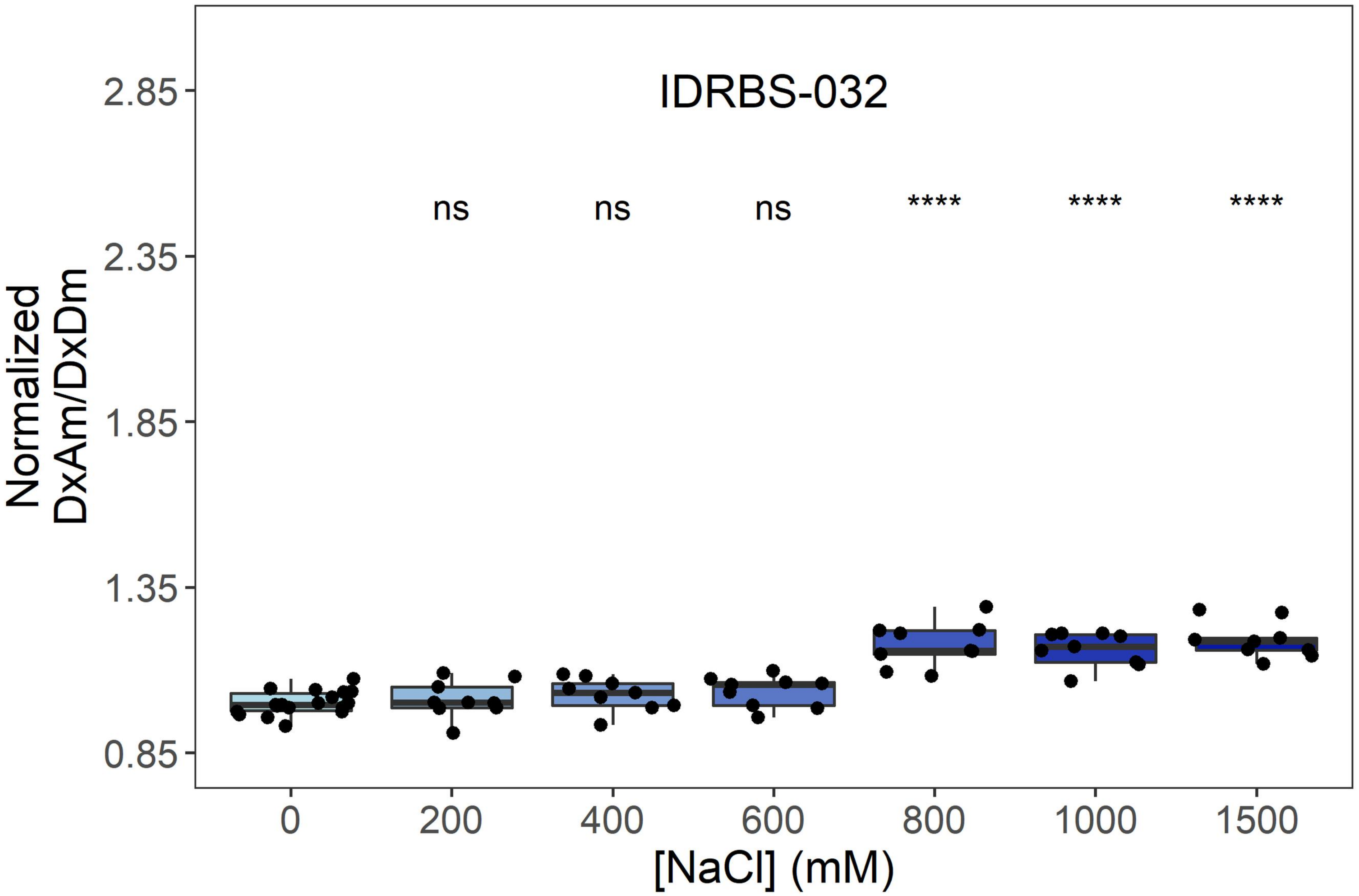
IDRBS-032
ns
ns
ns
****
****
****
Normalized DxAm/DxDm
2.85
2.35
1.85
1.35
0.85
0
200
400
600
800
1000
1500
[NaCl] (mM)

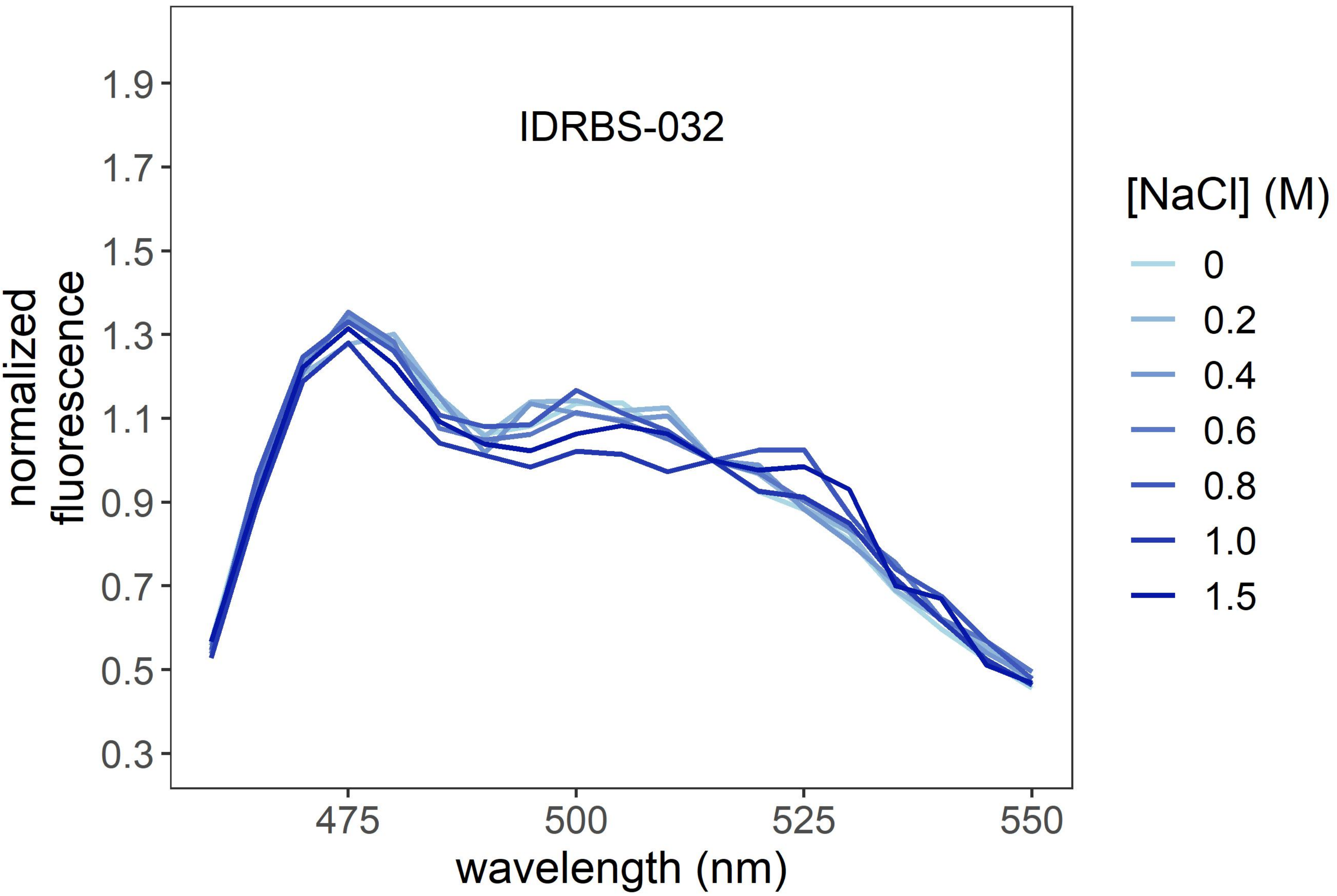
IDRBS-032
normalized fluorescence
1.9
1.7
1.5
1.3
1.1
0.9
0.7
0.5
0.3
475
500
525
550
wavelength (nm)
[NaCl] (M)
0
0.2
0.4
0.6
0.8
1.0
1.5

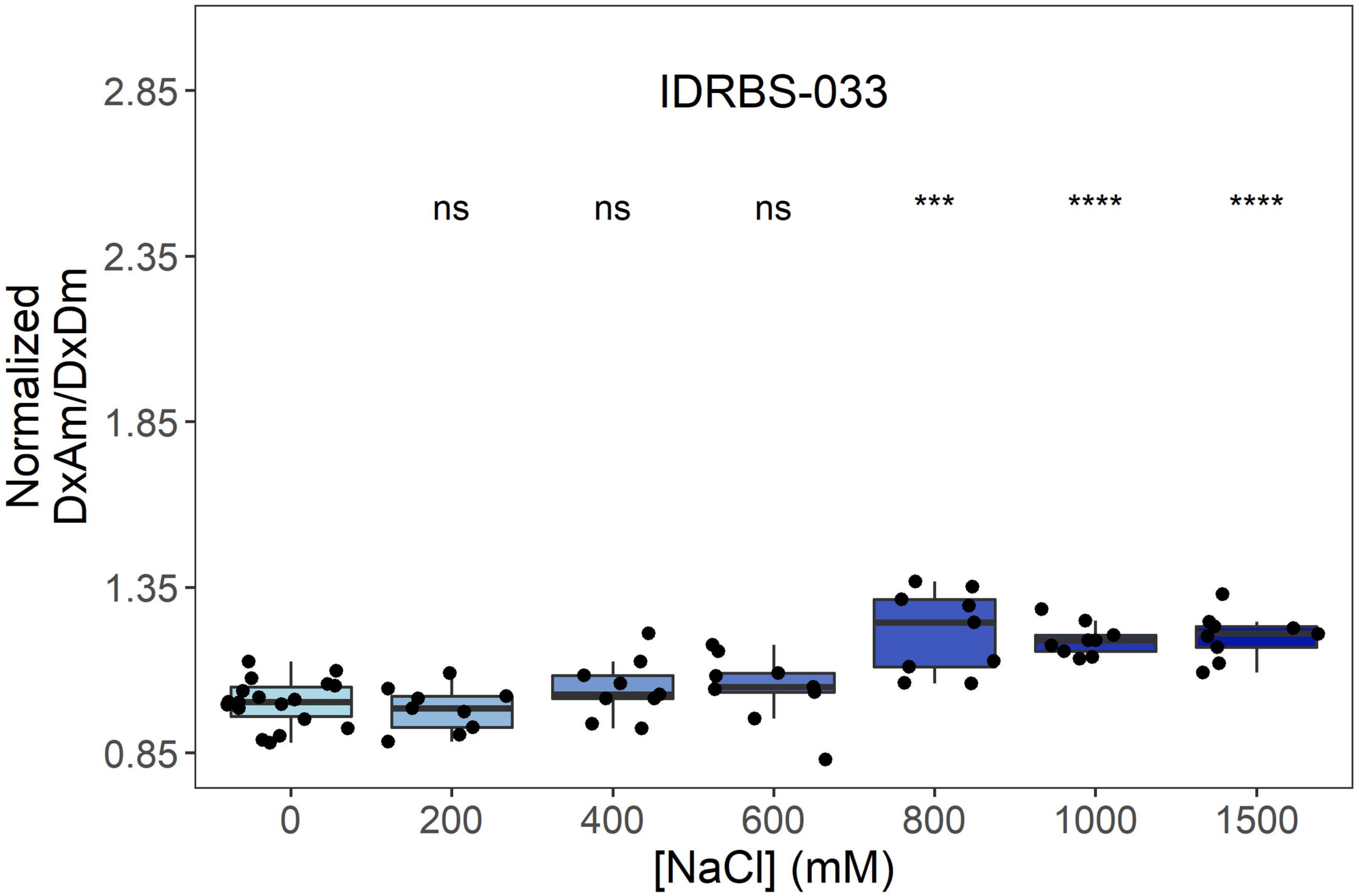
IDRBS-033
ns
ns
ns
***
****
****
2.85
2.35
1.85
1.35
0.85
Normalized DxAm/DxDm
0
200
400
600
800
1000
1500
[NaCl] (mM)

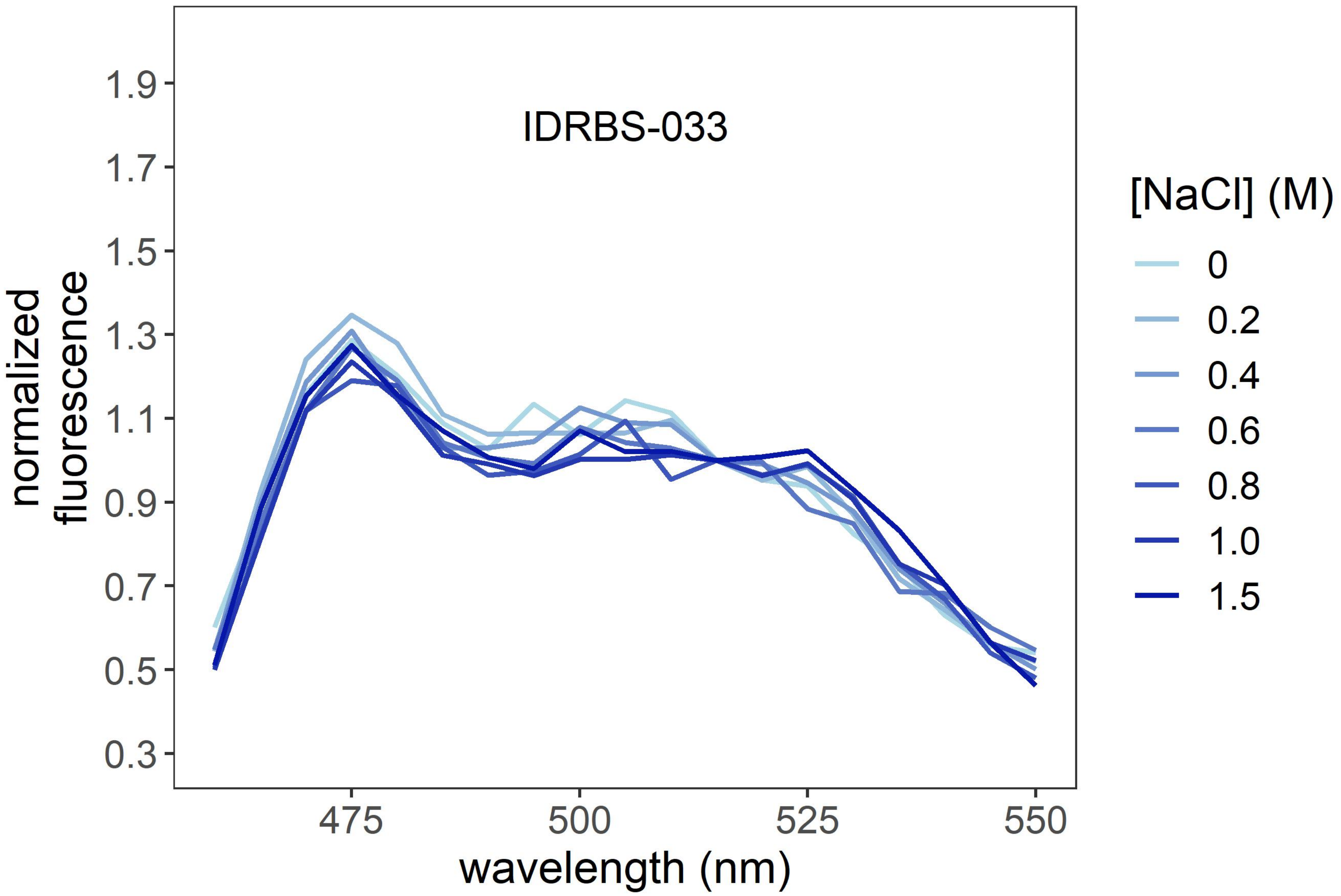
IDRBS-033
1.9
1.7
1.5
1.3
1.1
0.9
0.7
0.5
0.3
normalized fluorescence
475
500
525
550
wavelength (nm)
[NaCl] (M)
0
0.2
0.4
0.6
0.8
1.0
1.5

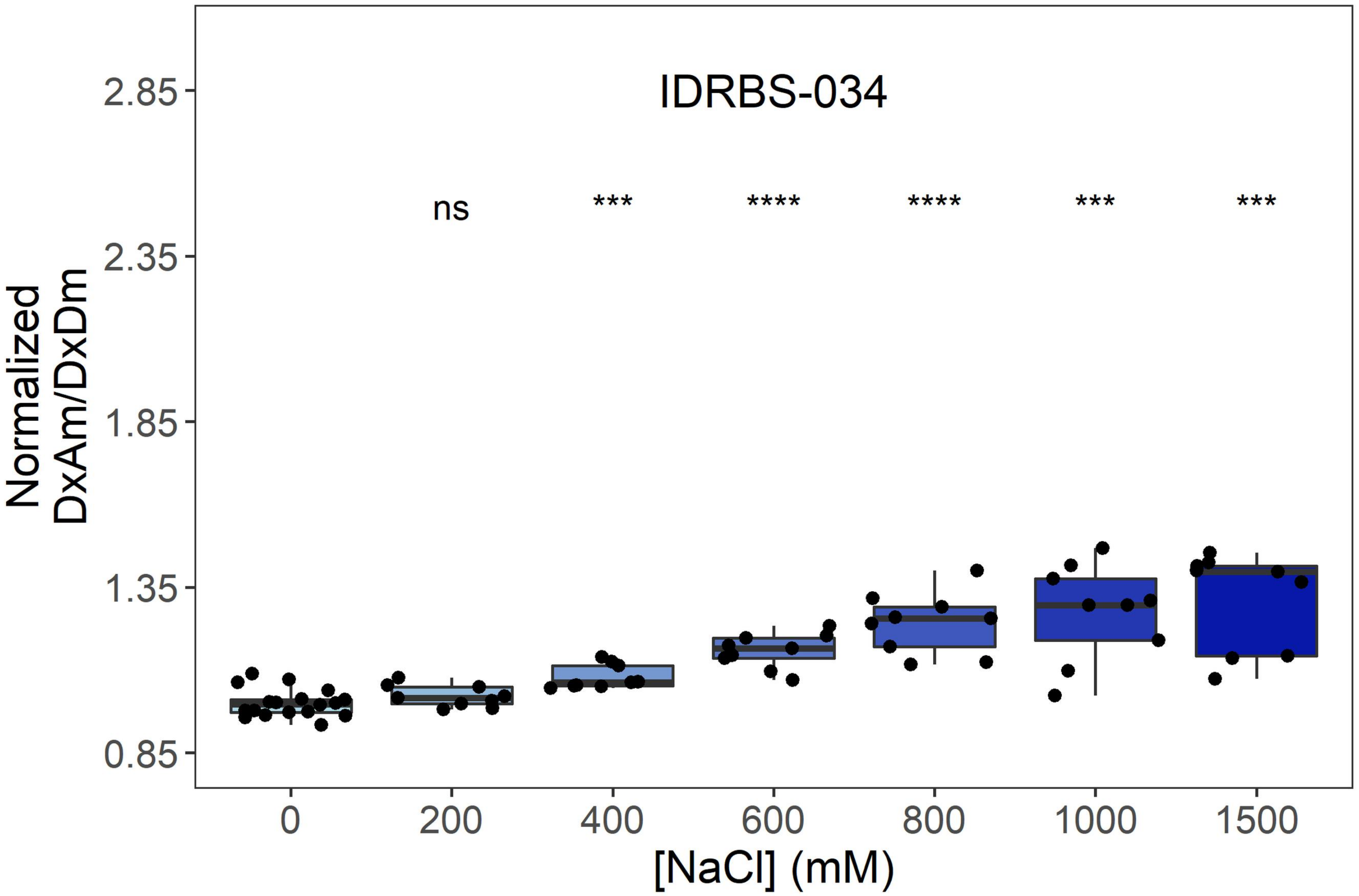

IDRBS-034
ns
***
****
****
***
***
2.85
2.35
1.85
1.35
0.85
Normalized
DxAm/DxDm
0
200
400
600
800
1000
1500
[NaCl] (mM)


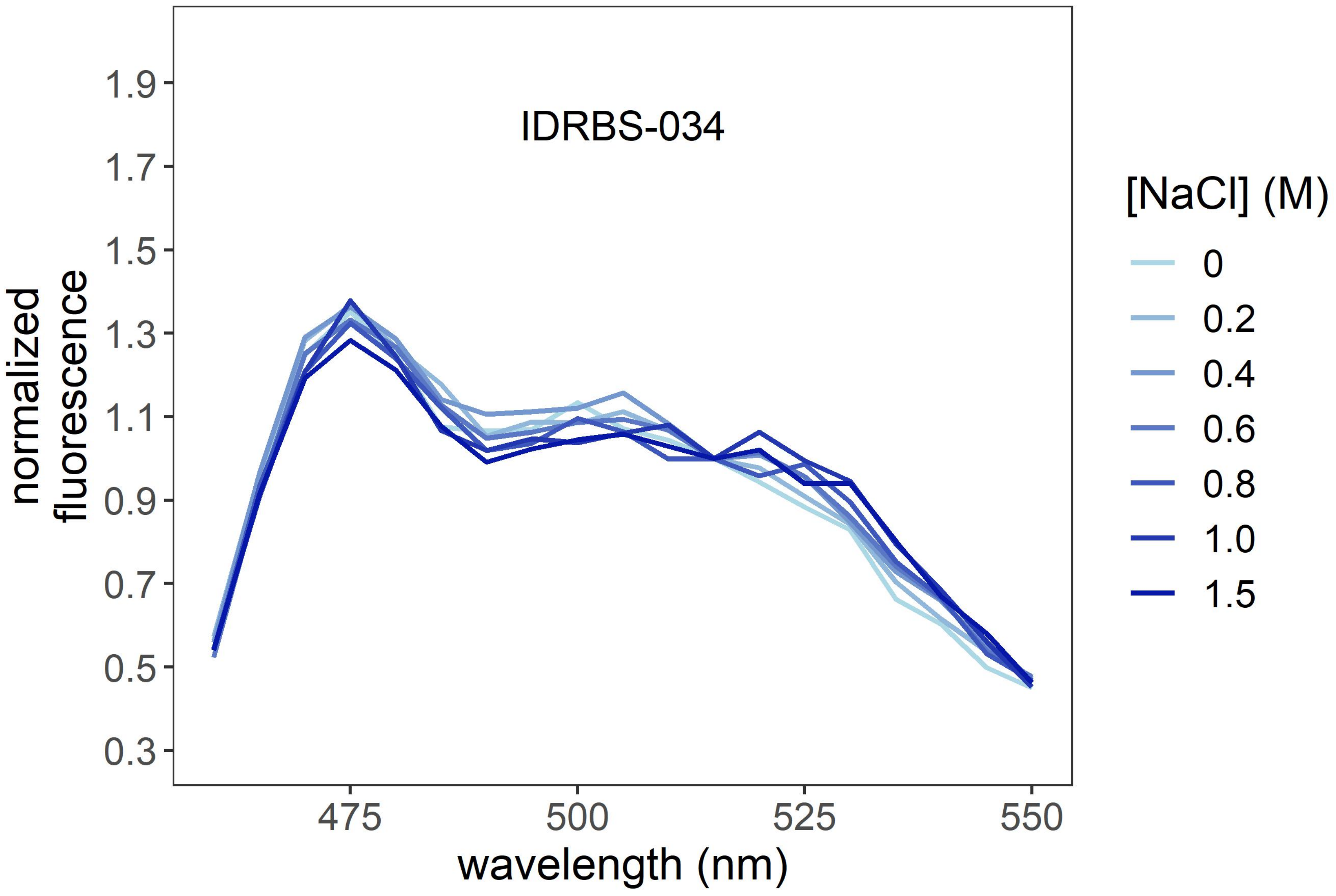

IDRBS-034
1.9
1.7
1.5
1.3
1.1
0.9
0.7
0.5
0.3
normalized
fluorescence
475
500
525
550
wavelength (nm)
[NaCl] (M)
0
0.2
0.4
0.6
0.8
1.0
1.5

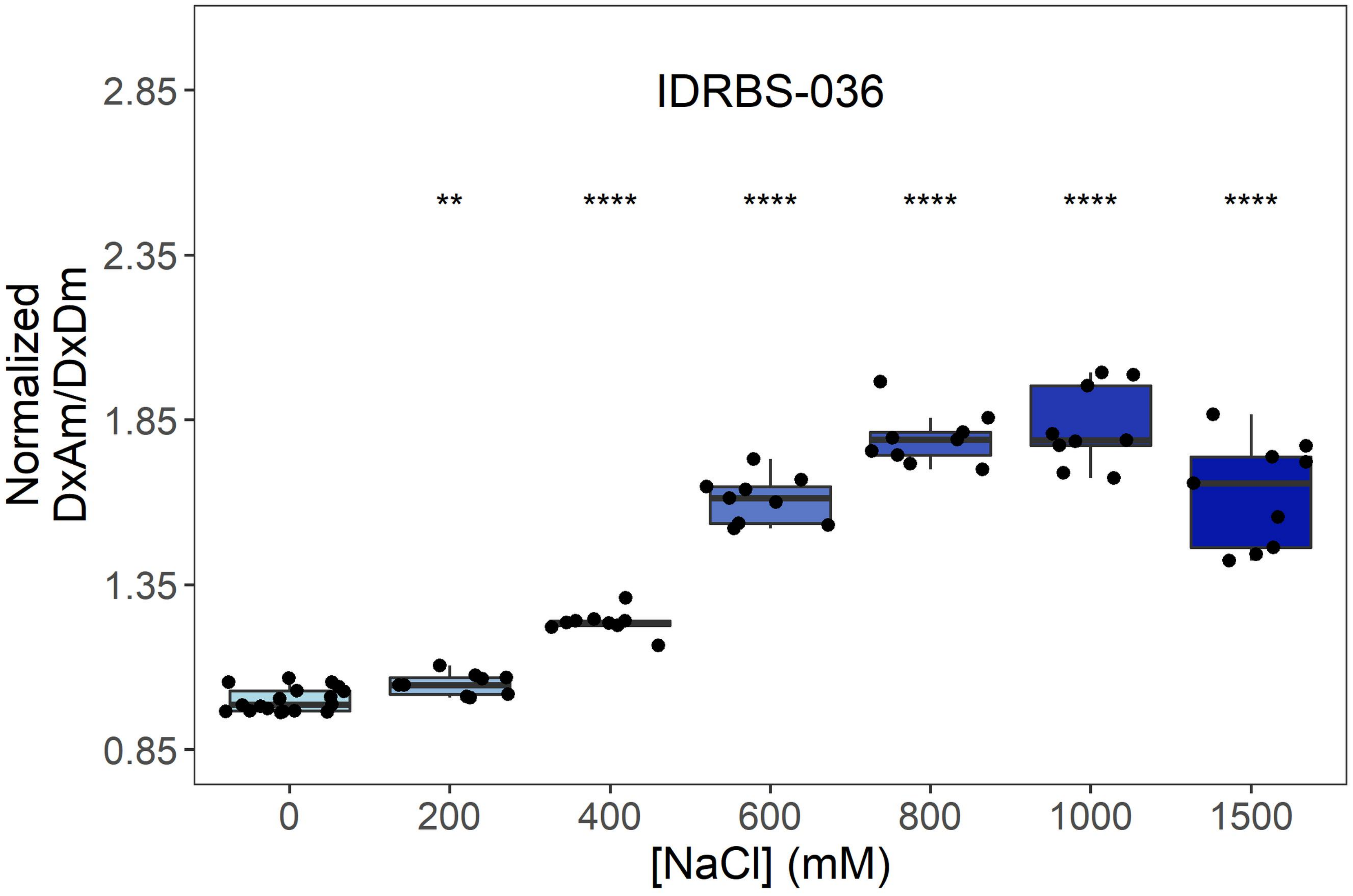

IDRBS-036
**
****
****
****
****
****
2.85
2.35
1.85
1.35
0.85
Normalized
DxAm/DxDm
0
200
400
600
800
1000
1500
[NaCl] (mM)


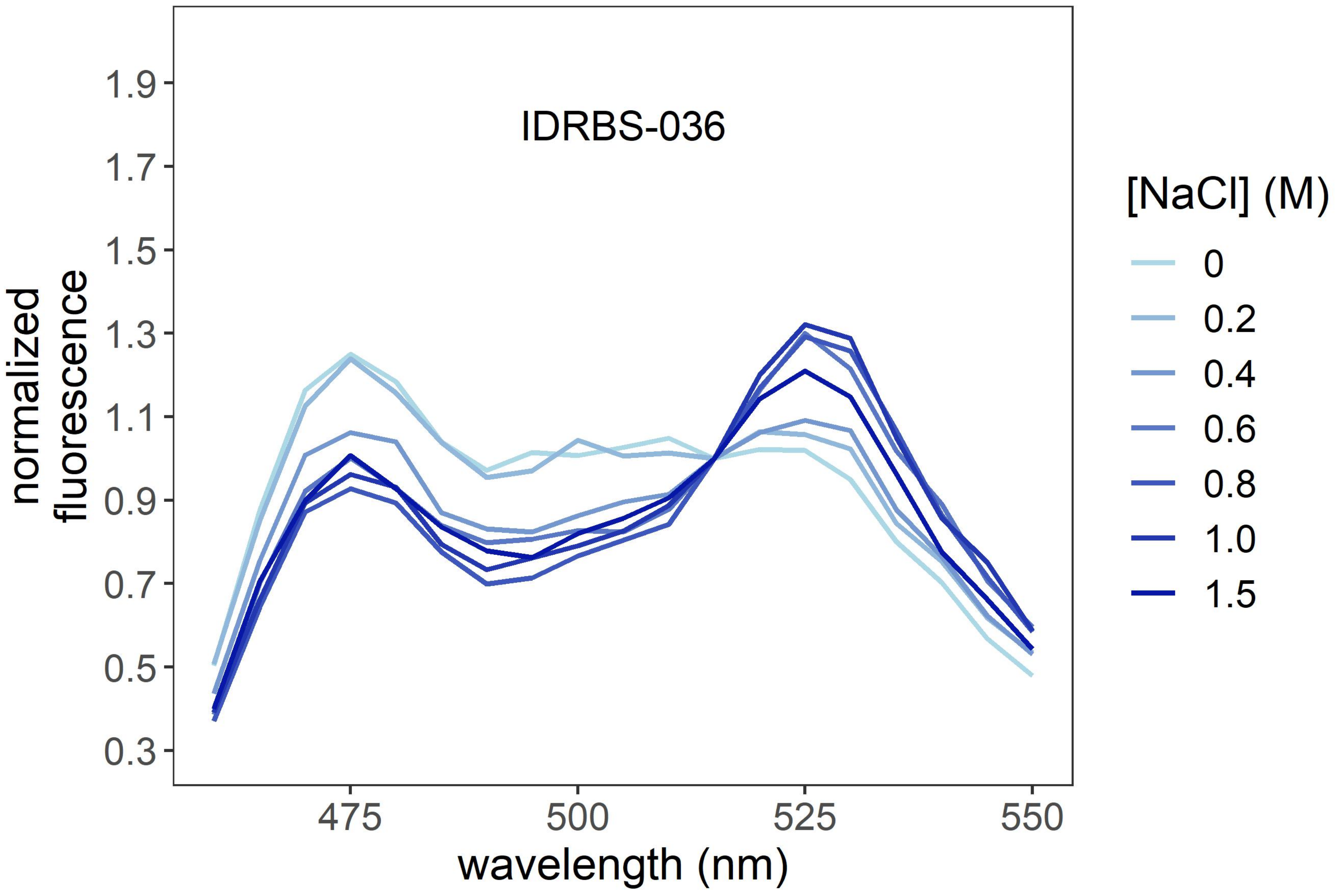

IDRBS-036
1.9
1.7
1.5
1.3
1.1
0.9
0.7
0.5
0.3
normalized
fluorescence
475
500
525
550
wavelength (nm)
[NaCl] (M)
0
0.2
0.4
0.6
0.8
1.0
1.5

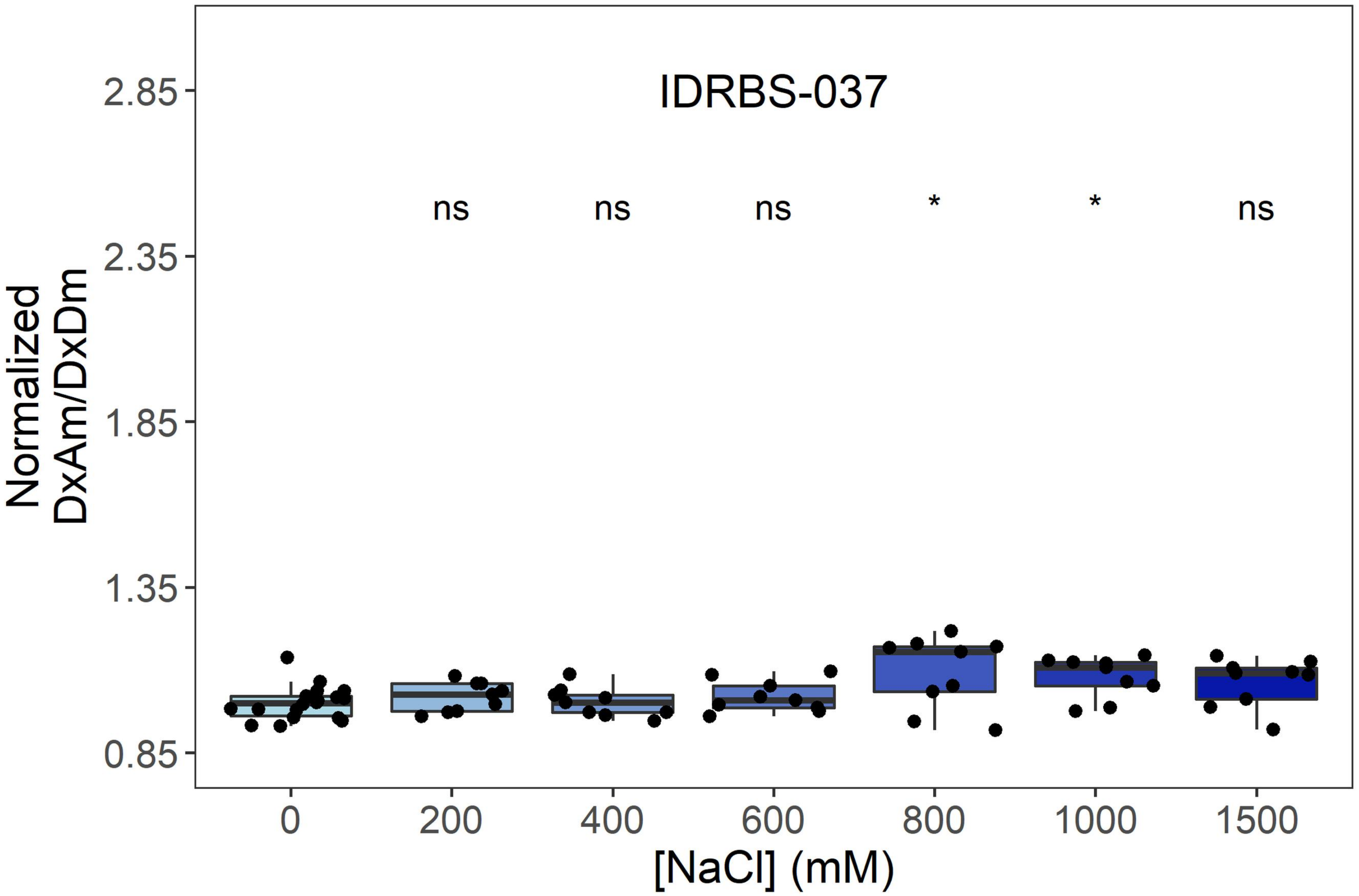
IDRBS-037
ns
ns
ns
*
*
ns
Normalized
DxAm/DxDm
2.85
2.35
1.85
1.35
0.85
0
200
400
600
800
1000
1500
[NaCl] (mM)

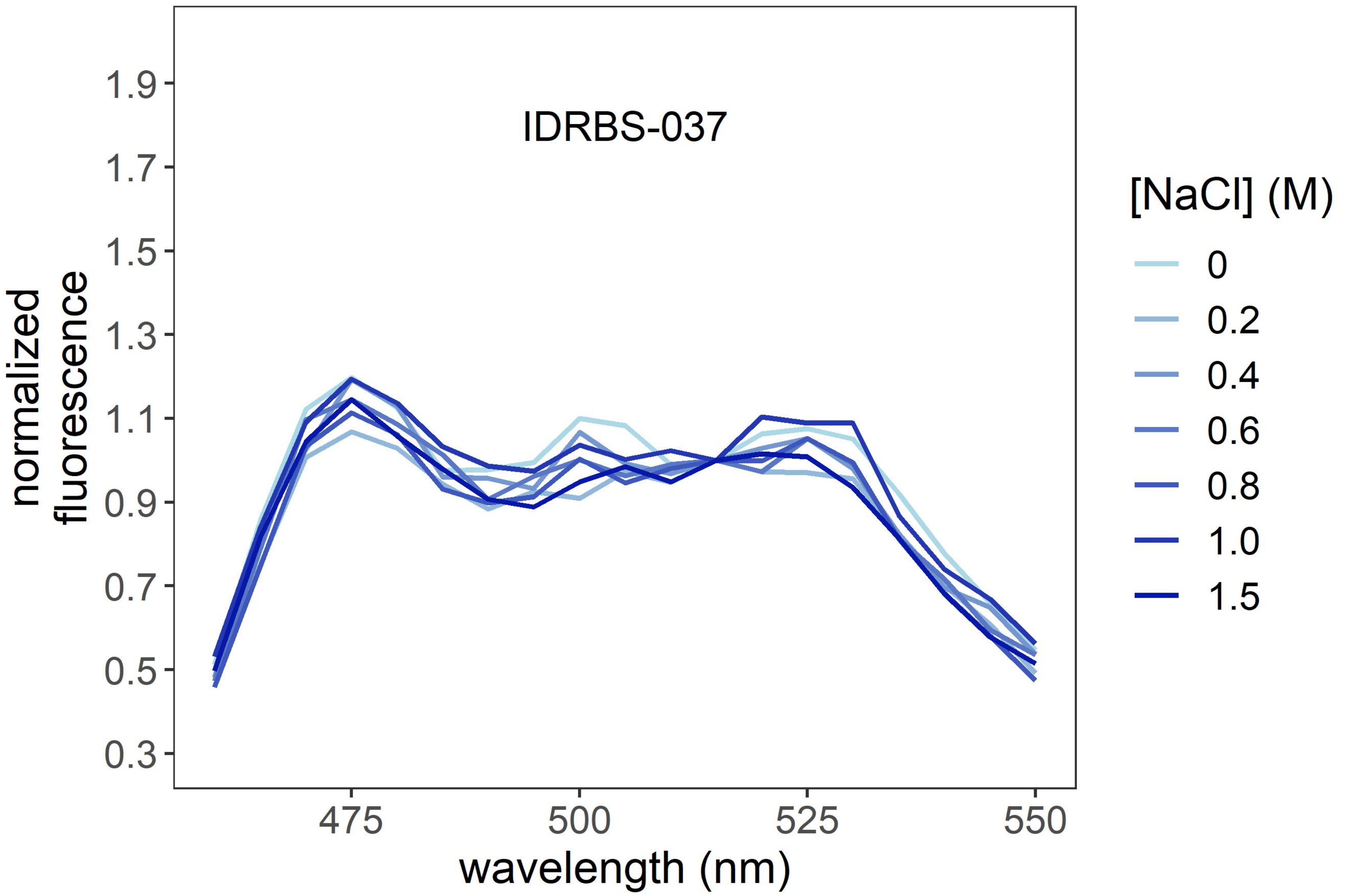
IDRBS-037
normalized
fluorescence
1.9
1.7
1.5
1.3
1.1
0.9
0.7
0.5
0.3
475
500
525
550
wavelength (nm)
[NaCl] (M)
0
0.2
0.4
0.6
0.8
1.0
1.5

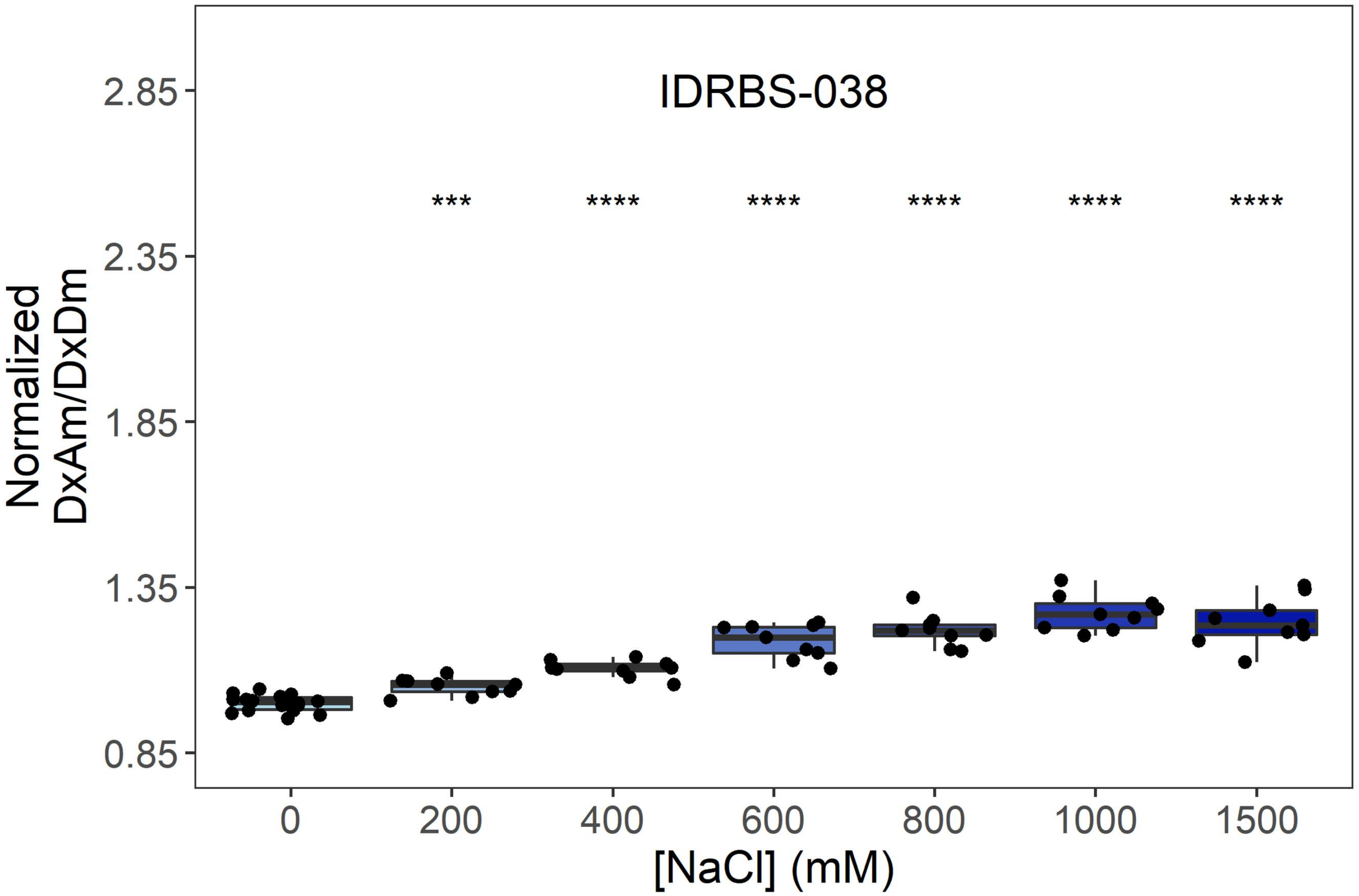

IDRBS-038
Normalized
DxAm/DxDm
2.85
2.35
1.85
1.35
0.85
***
****
****
****
****
****
0
200
400
600
800
1000
1500
[NaCl] (mM)


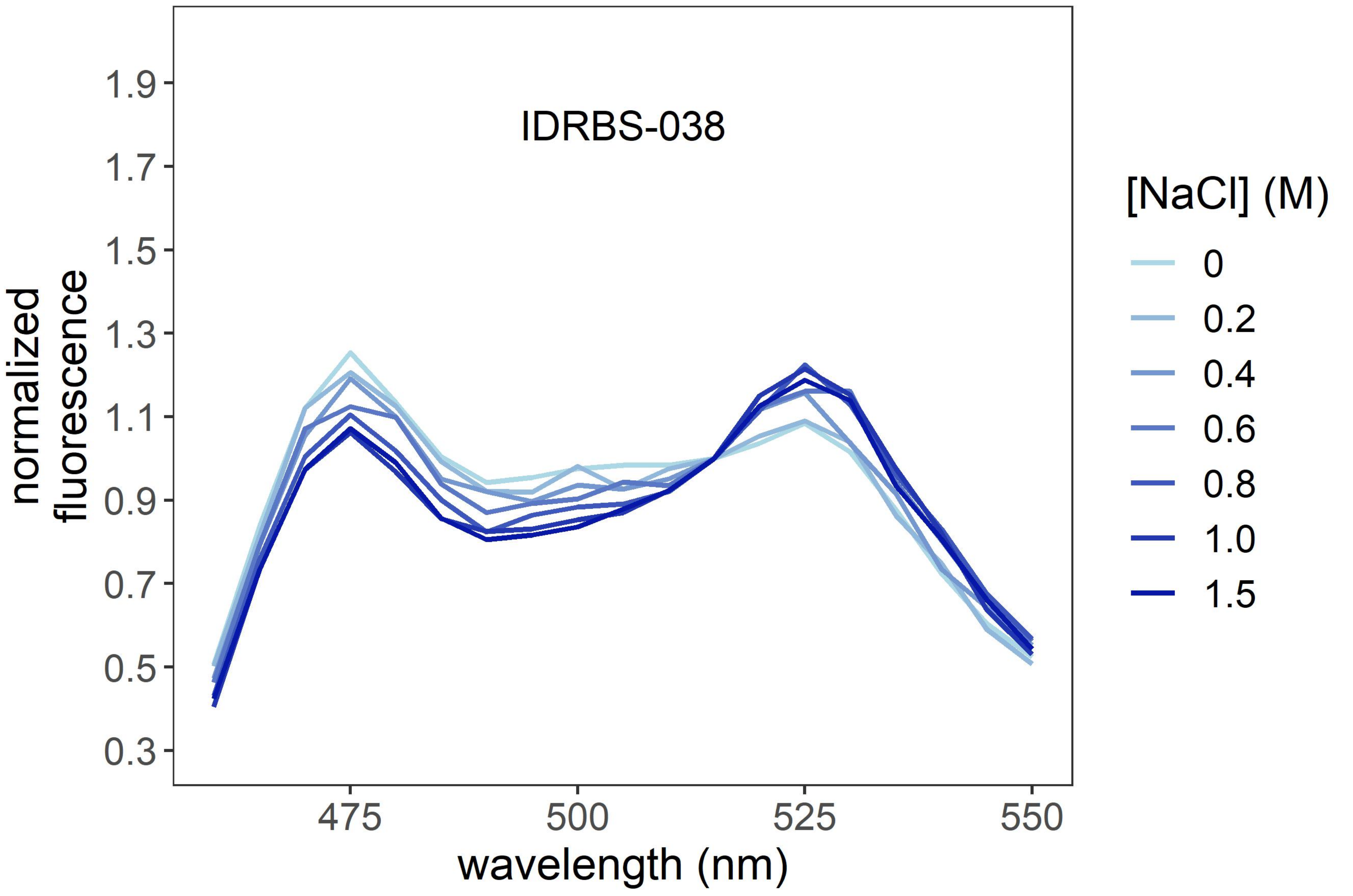

IDRBS-038
normalized
fluorescence
1.9
1.7
1.5
1.3
1.1
0.9
0.7
0.5
0.3
475
500
525
550
wavelength (nm)
[NaCl] (M)
0
0.2
0.4
0.6
0.8
1.0
1.5

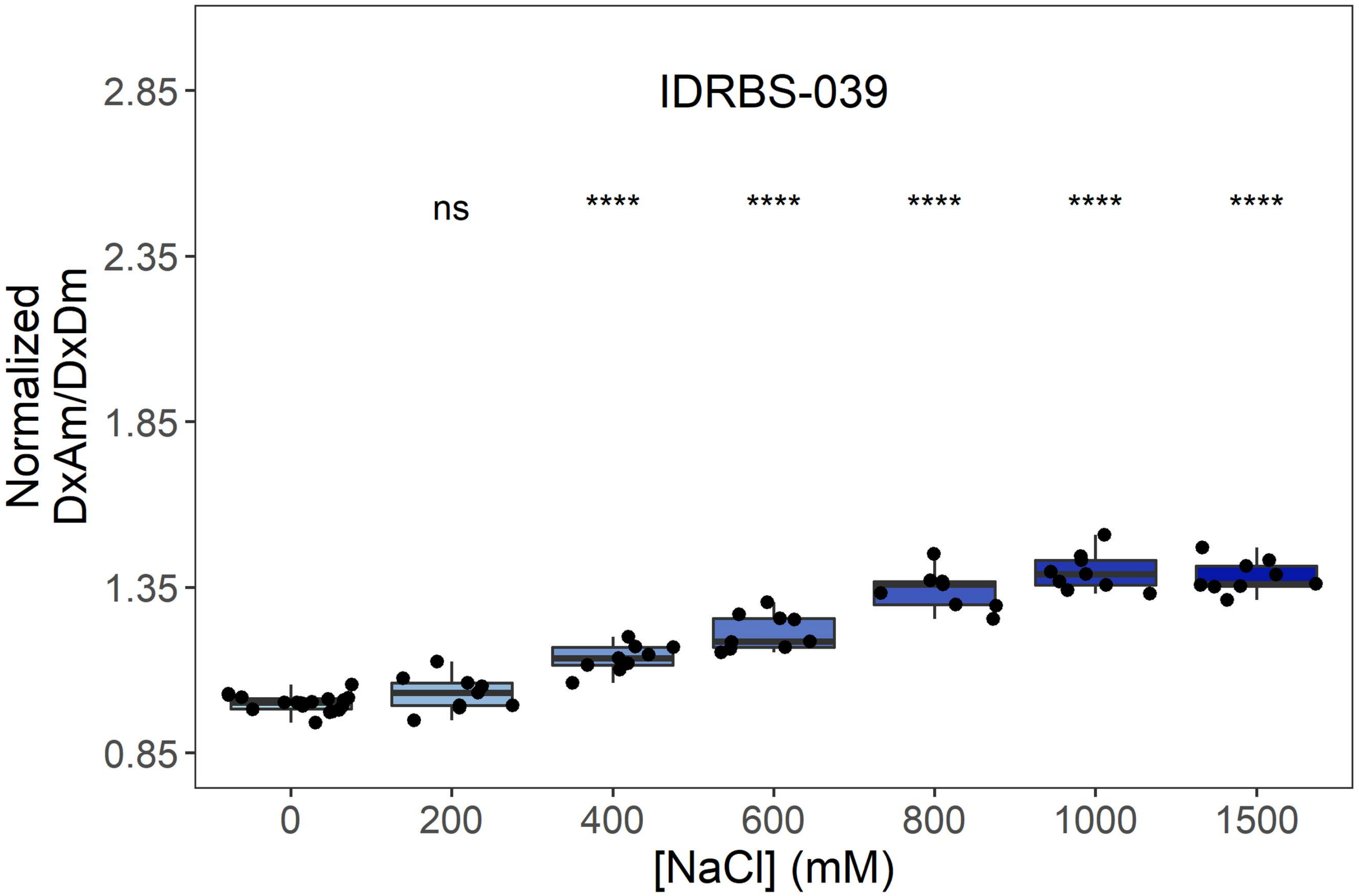
IDRBS-039
ns
****
****
****
****
****
Normalized
DxAm/DxDm
2.85
2.35
1.85
1.35
0.85
0
200
400
600
800
1000
1500
[NaCl] (mM)

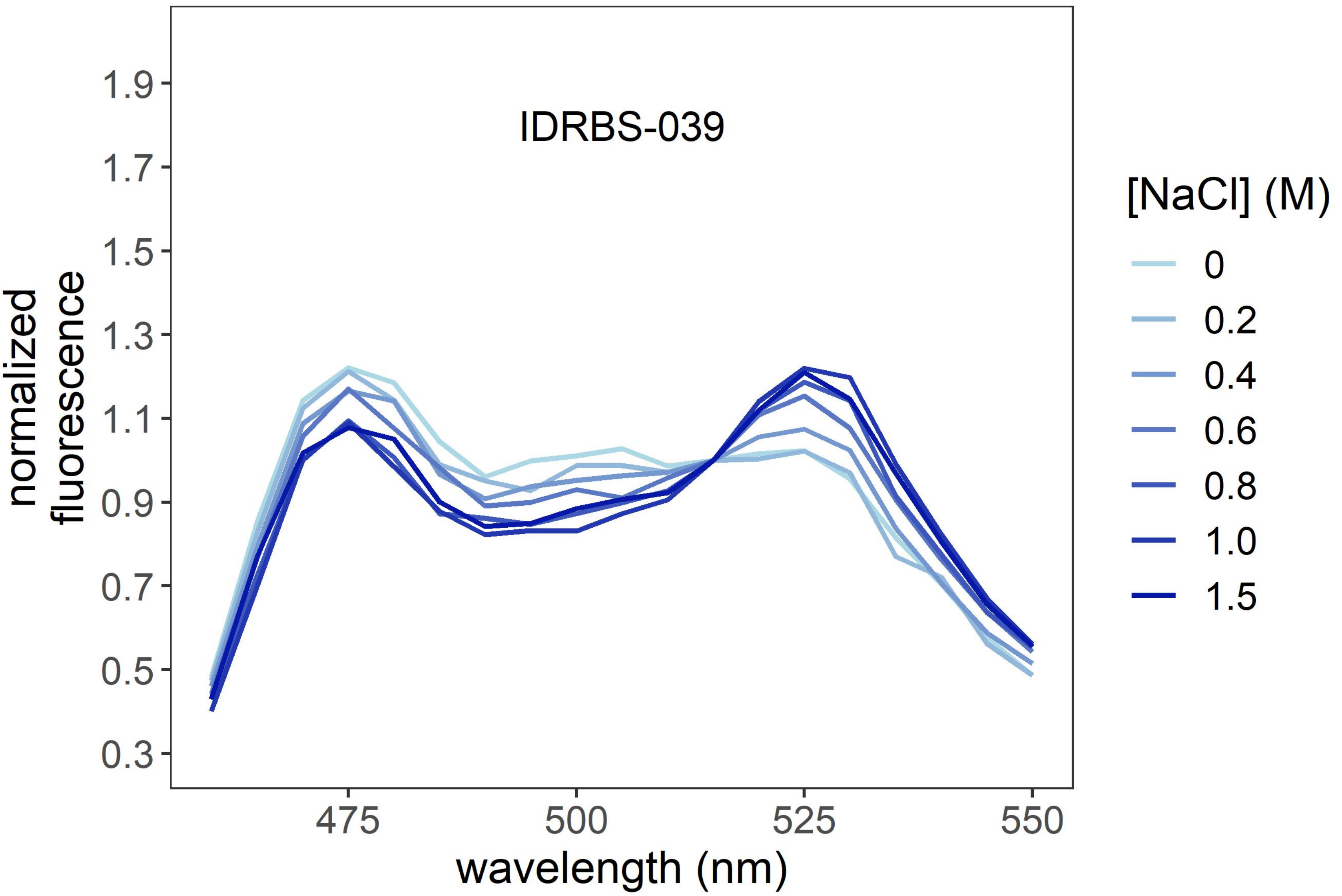
IDRBS-039
normalized
fluorescence
1.9
1.7
1.5
1.3
1.1
0.9
0.7
0.5
0.3
475
500
525
550
wavelength (nm)
[NaCl] (M)
0
0.2
0.4
0.6
0.8
1.0
1.5

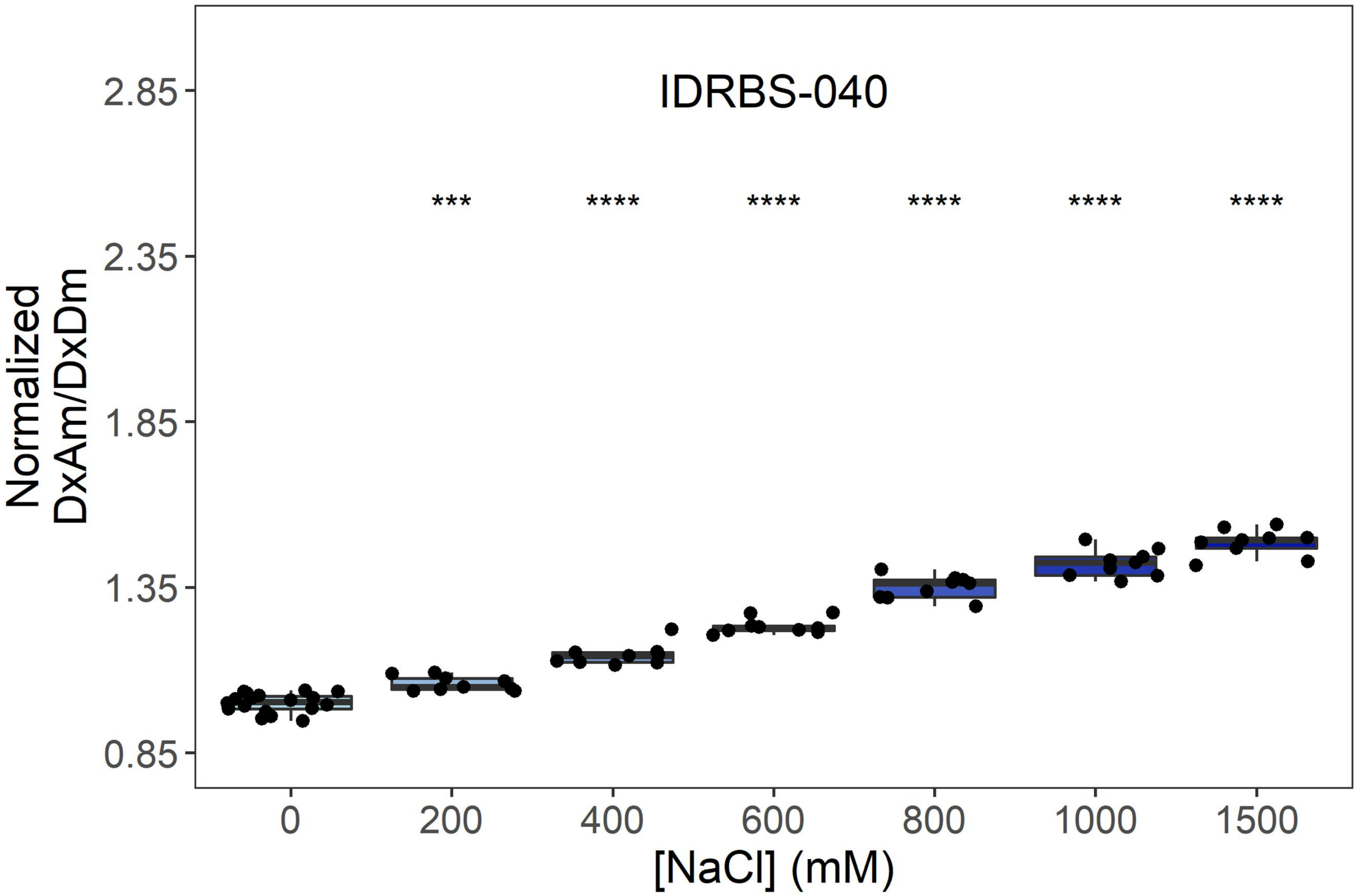
IDRBS-040
***
****
****
****
****
****
Normalized DxAm/DxDm
2.85
2.35
1.85
1.35
0.85
0
200
400
600
800
1000
1500
[NaCl] (mM)

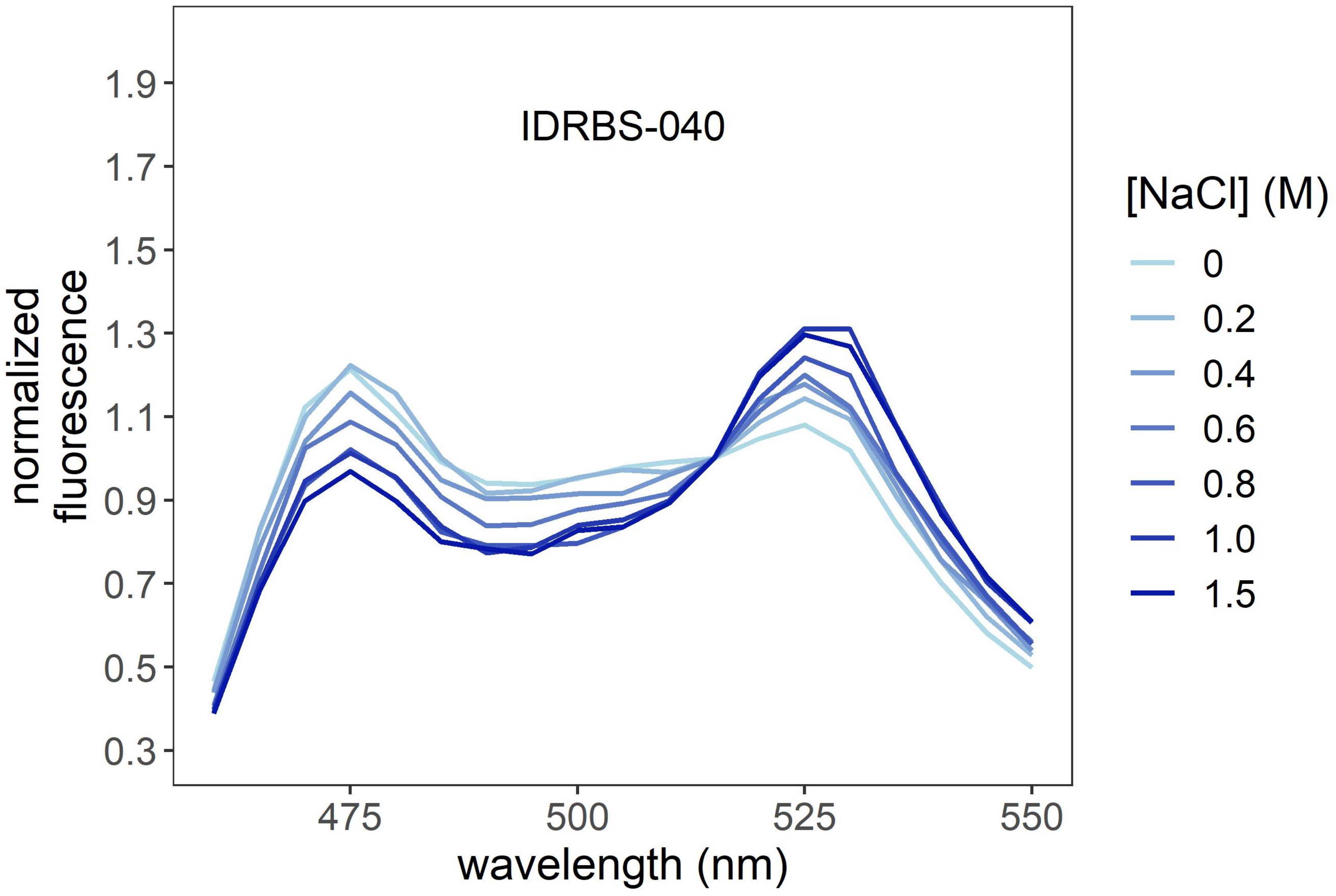
IDRBS-040
normalized fluorescence
1.9
1.7
1.5
1.3
1.1
0.9
0.7
0.5
0.3
475
500
525
550
wavelength (nm)
[NaCl] (M)
0
0.2
0.4
0.6
0.8
1.0
1.5

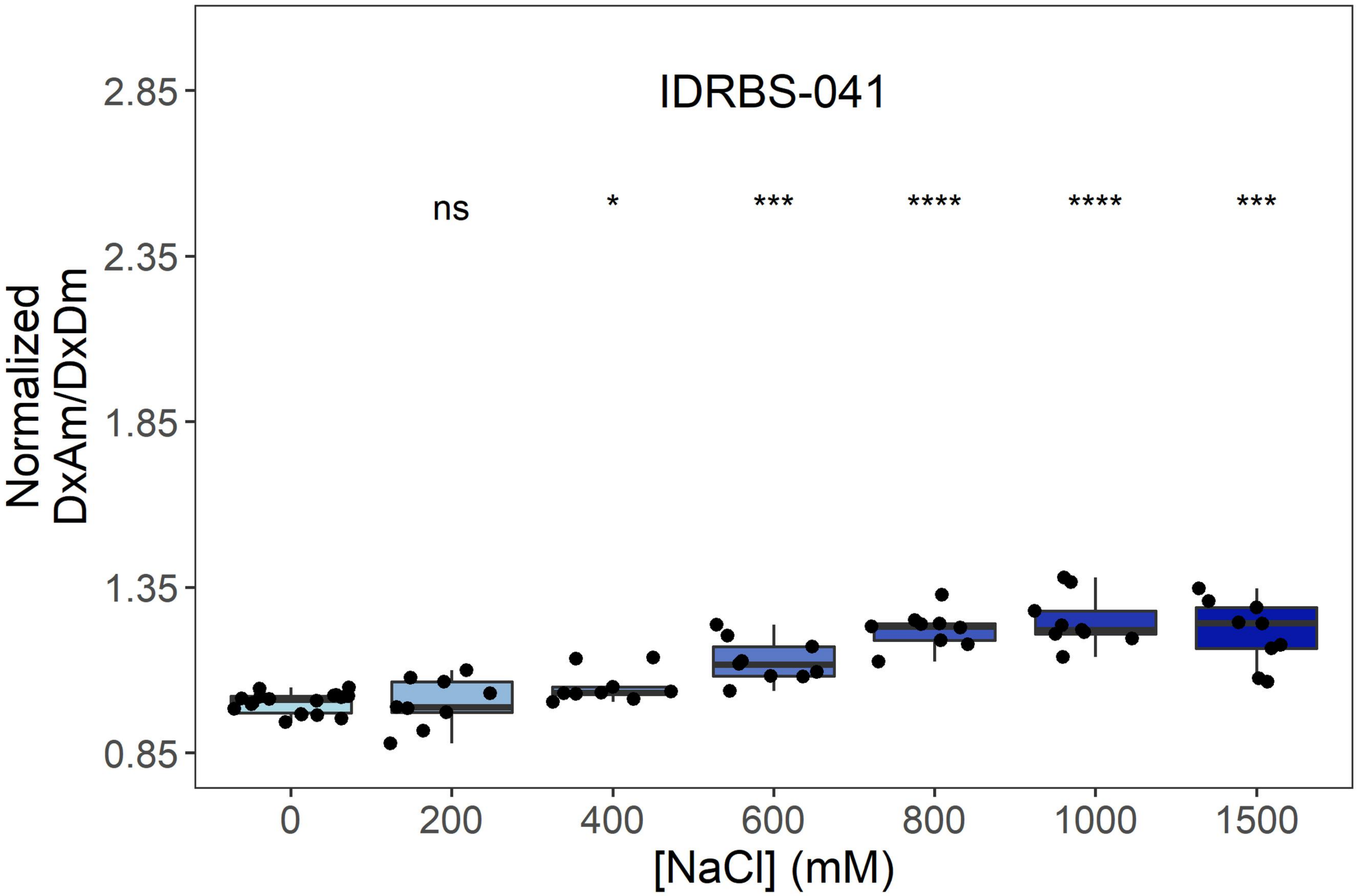
IDRBS-041
ns
*
***
****
****
***
Normalized
DxAm/DxDm
2.85
2.35
1.85
1.35
0.85
0
200
400
600
800
1000
1500
[NaCl] (mM)

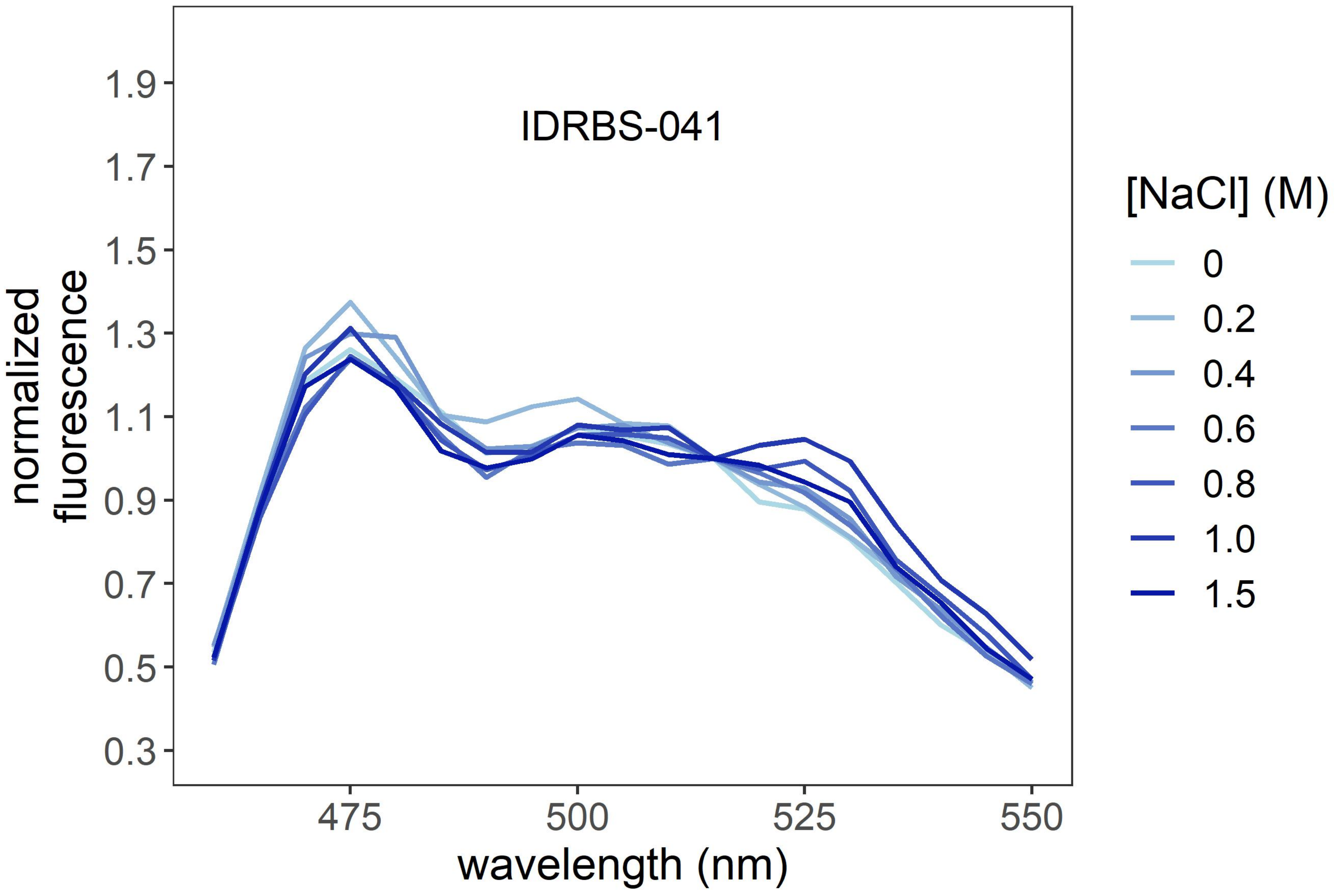
IDRBS-041
normalized
fluorescence
1.9
1.7
1.5
1.3
1.1
0.9
0.7
0.5
0.3
475
500
525
550
wavelength (nm)
[NaCl] (M)
0
0.2
0.4
0.6
0.8
1.0
1.5

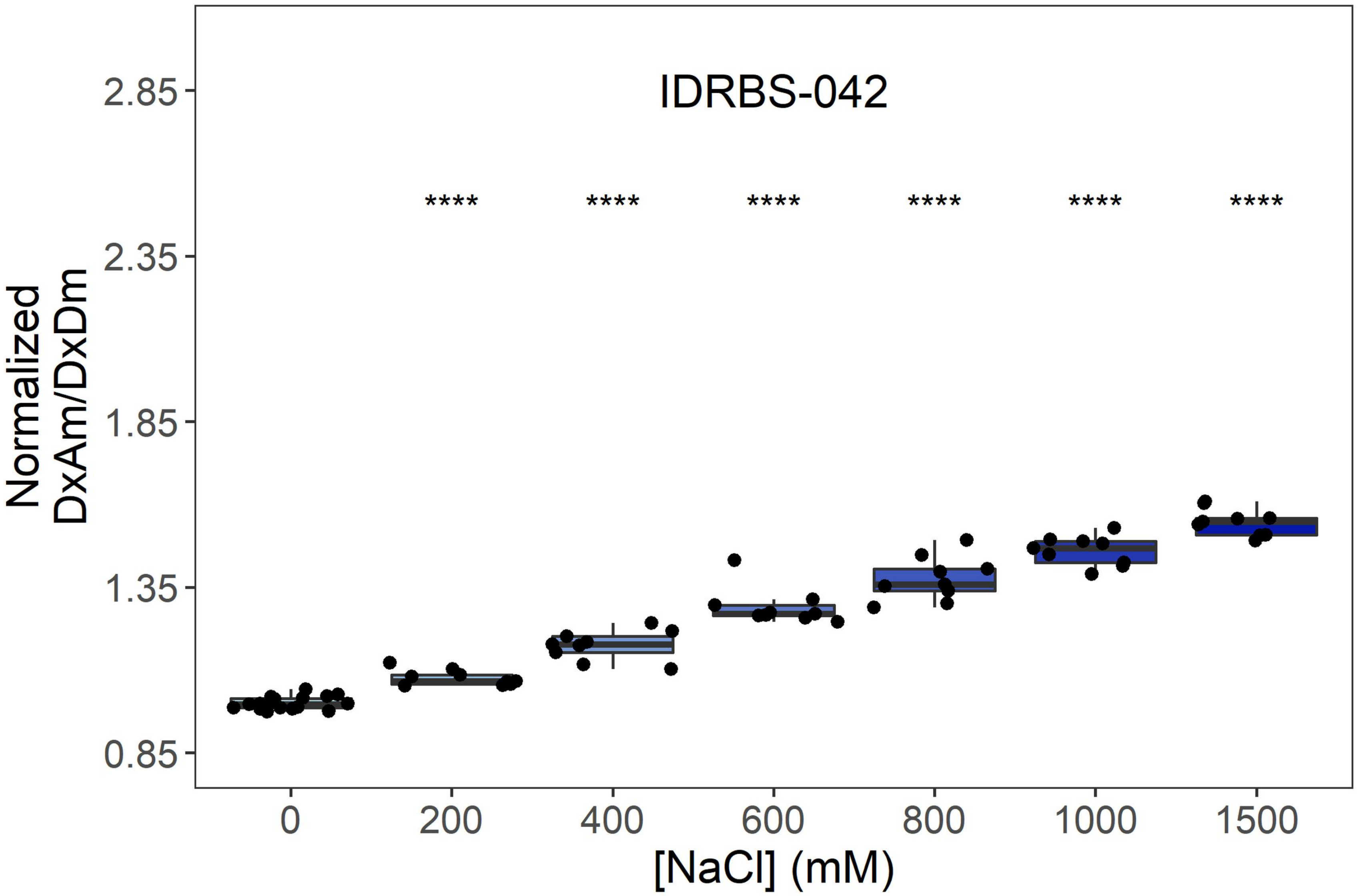
IDRBS-042
****
2.85
2.35
1.85
1.35
0.85
Normalized
DxAm/DxDm
0
200
400
600
800
1000
1500
[NaCl] (mM)

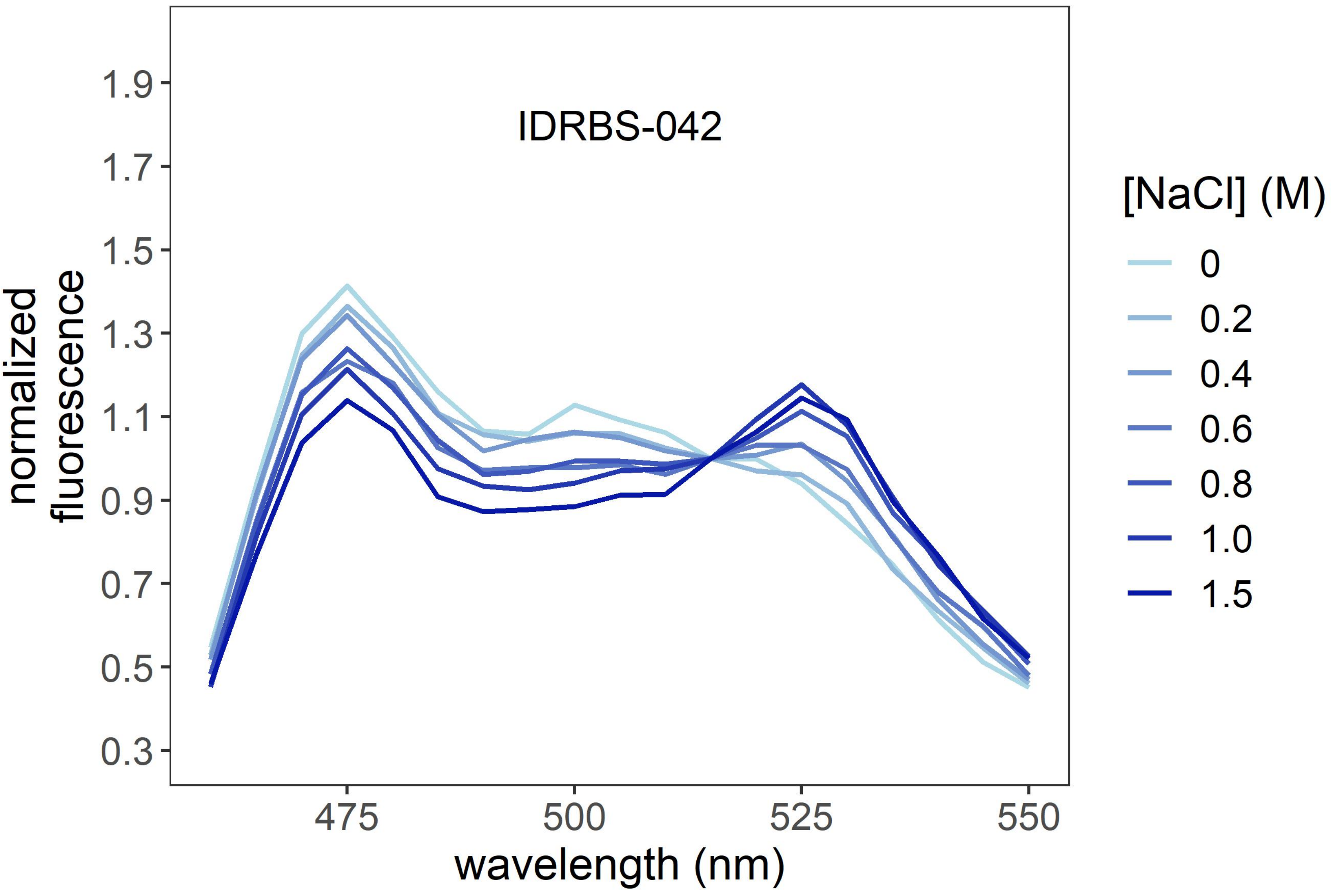
IDRBS-042
1.9
1.7
1.5
1.3
1.1
0.9
0.7
0.5
0.3
normalized
fluorescence
475
500
525
550
wavelength (nm)
[NaCl] (M)
0
0.2
0.4
0.6
0.8
1.0
1.5

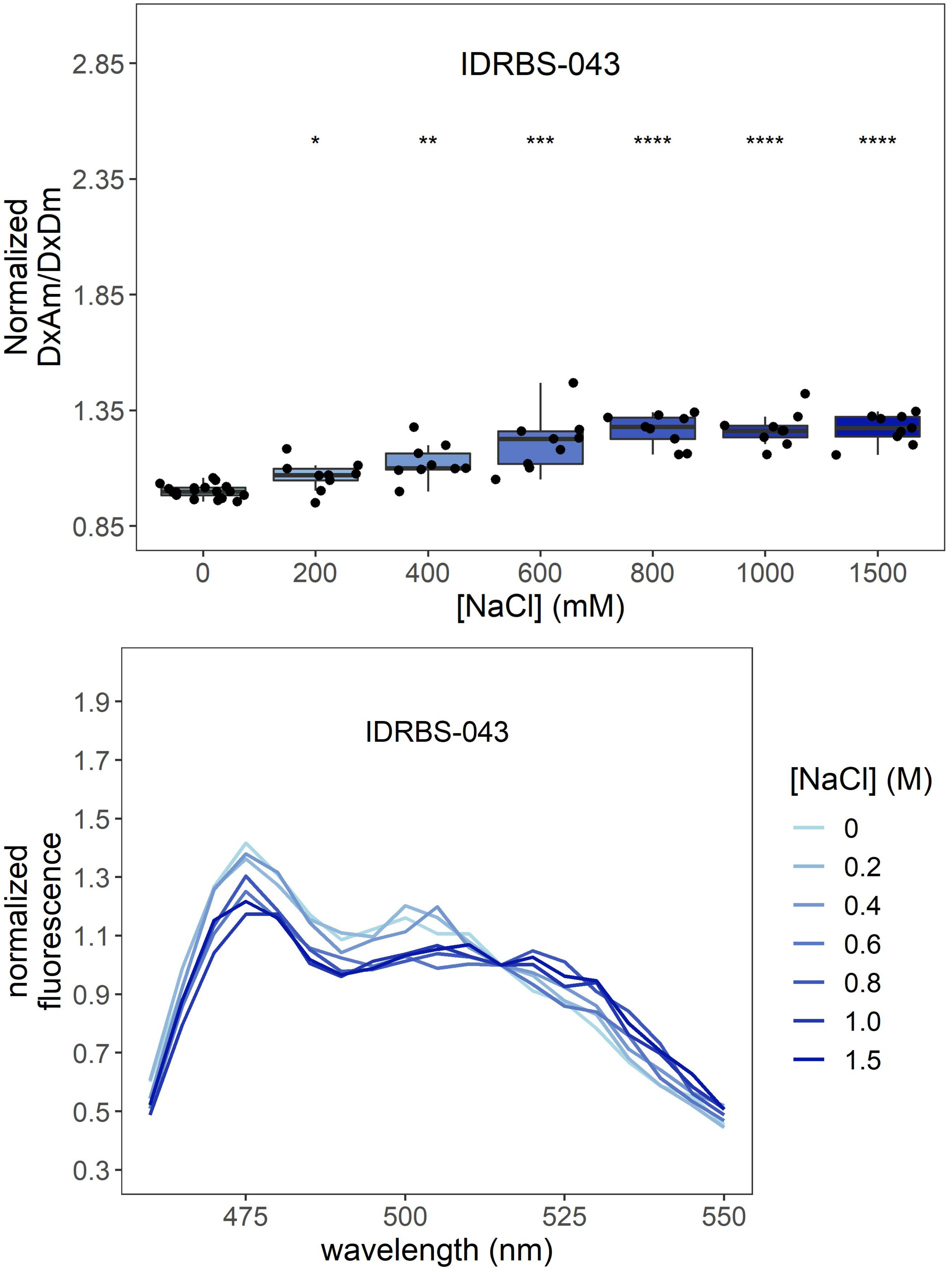
IDRBS-043
Normalized
DxAm/DxDm
2.85
2.35
1.85
1.35
0.85
*
**
***
****
****
****
0
200
400
600
800
1000
1500
[NaCl] (mM)
IDRBS-043
normalized
fluorescence
1.9
1.7
1.5
1.3
1.1
0.9
0.7
0.5
0.3
475
500
525
550
wavelength (nm)
[NaCl] (M)
0
0.2
0.4
0.6
0.8
1.0
1.5

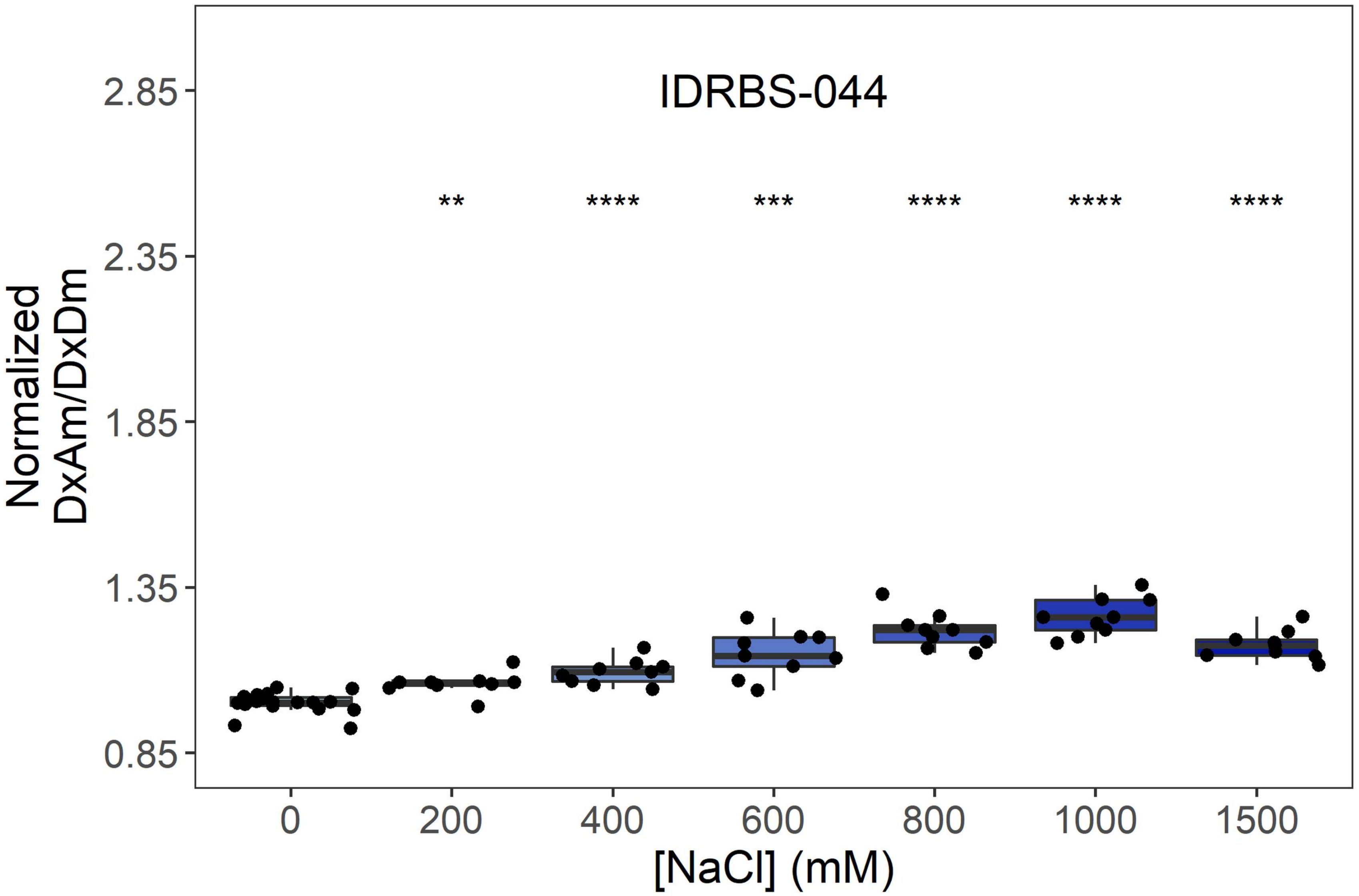
IDRBS-044
Normalized DxAm/DxDm
2.85
2.35
1.85
1.35
0.85
**
****
***
****
****
****
0
200
400
600
800
1000
1500
[NaCl] (mM)

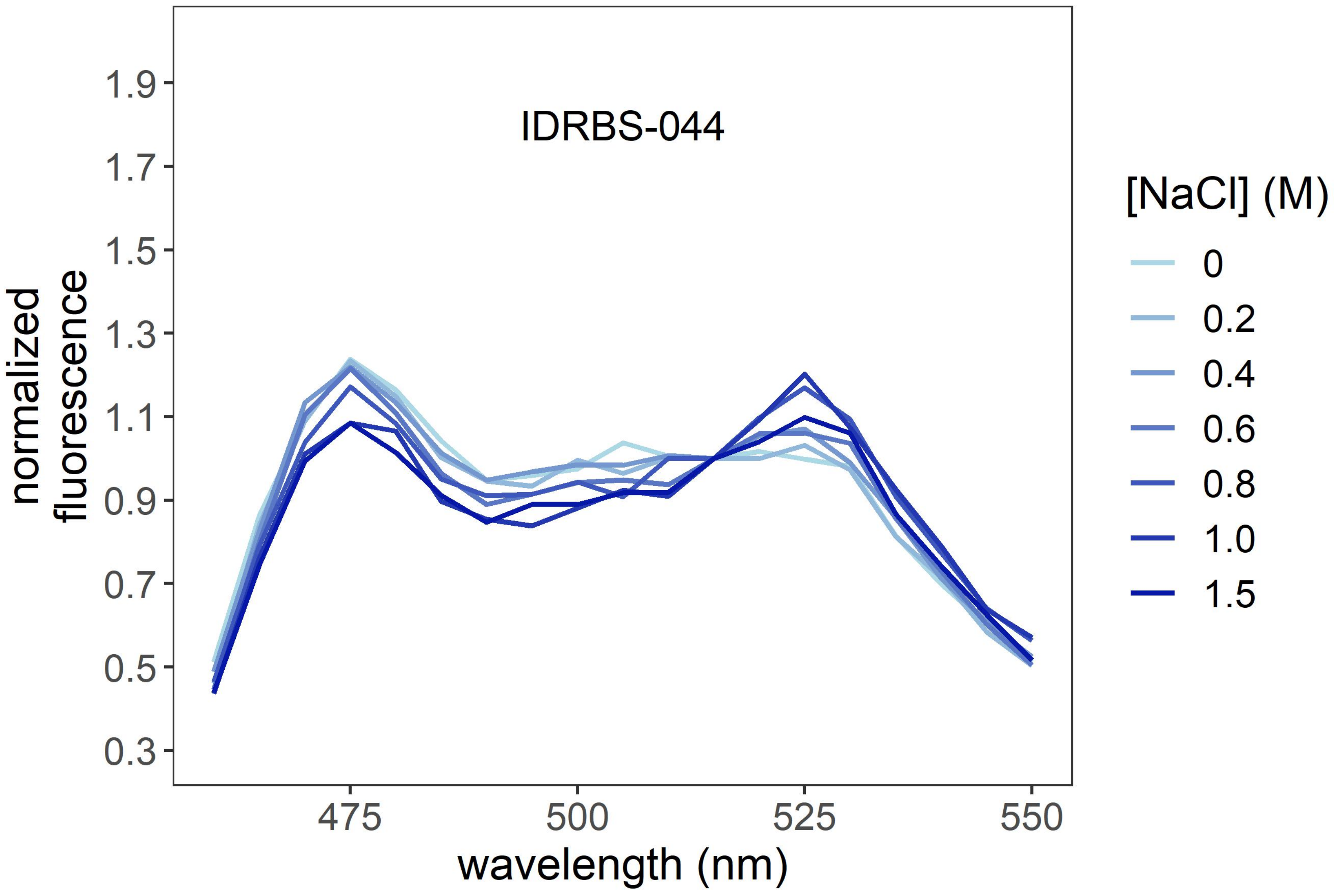
IDRBS-044
normalized fluorescence
1.9
1.7
1.5
1.3
1.1
0.9
0.7
0.5
0.3
475
500
525
550
wavelength (nm)
[NaCl] (M)
0
0.2
0.4
0.6
0.8
1.0
1.5

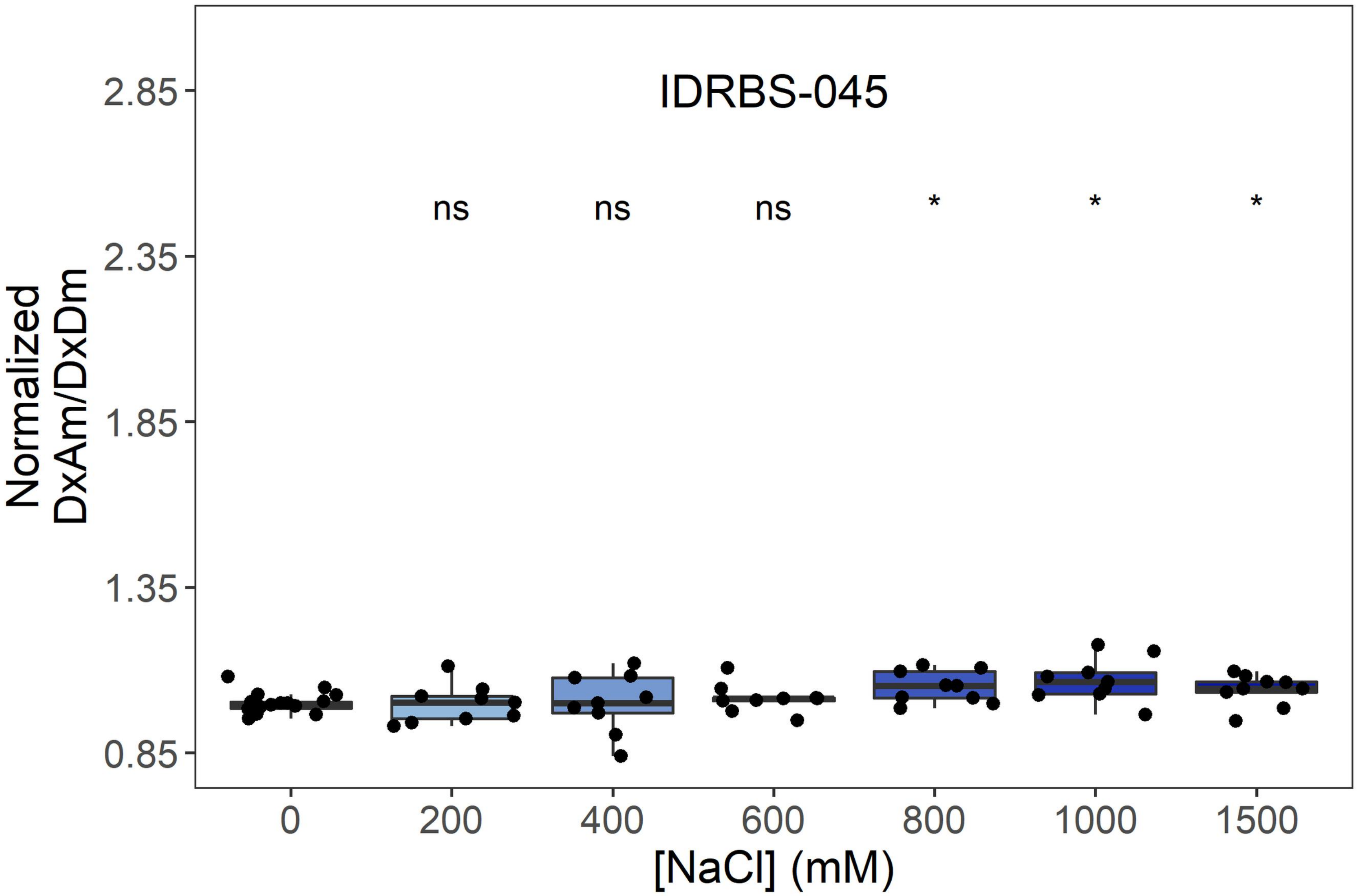
IDRBS-045
ns
ns
ns
*
*
*
Normalized DxAm/DxDm
2.85
2.35
1.85
1.35
0.85
0
200
400
600
800
1000
1500
[NaCl] (mM)

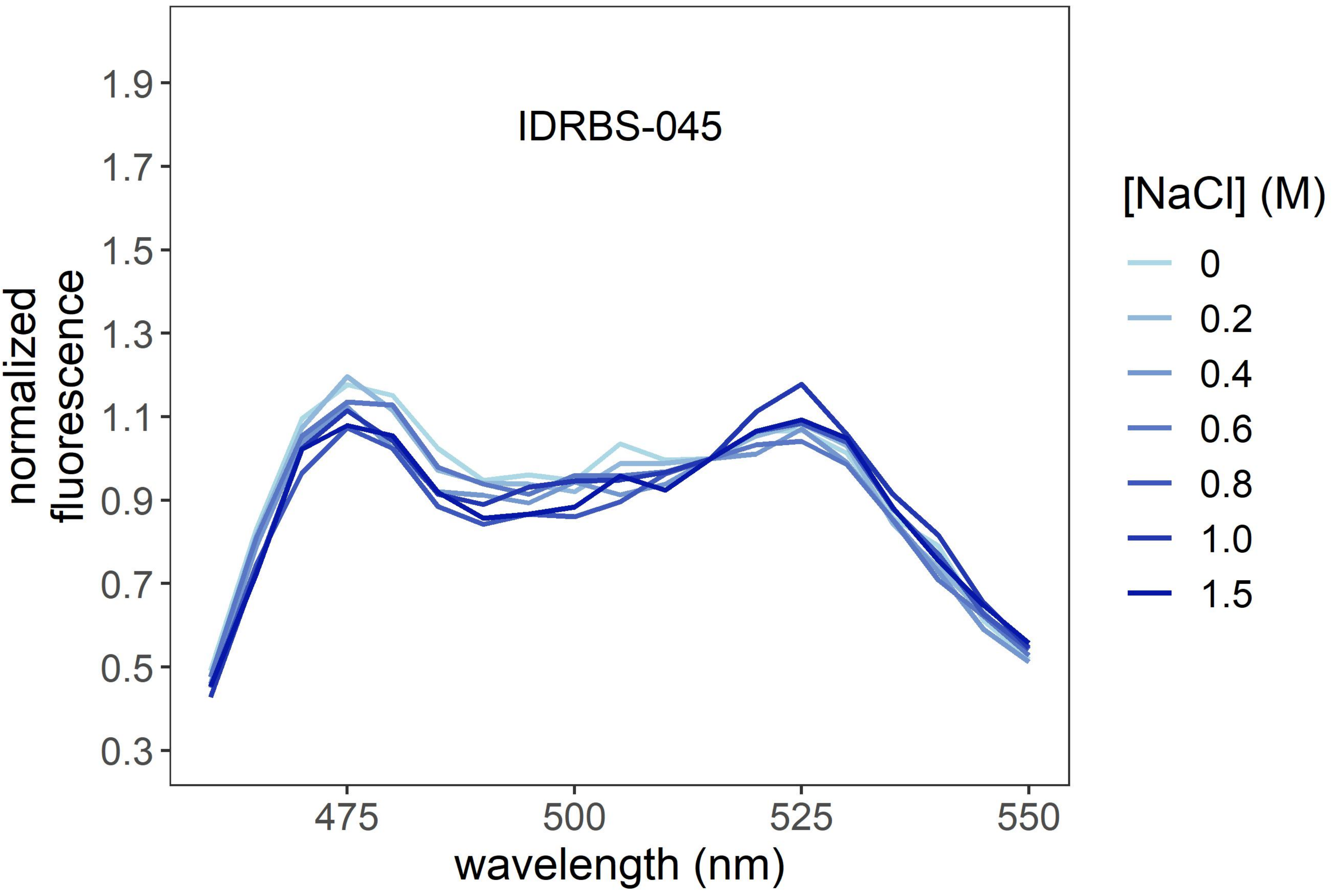
IDRBS-045
normalized fluorescence
1.9
1.7
1.5
1.3
1.1
0.9
0.7
0.5
0.3
475
500
525
550
wavelength (nm)
[NaCl] (M)
0
0.2
0.4
0.6
0.8
1.0
1.5

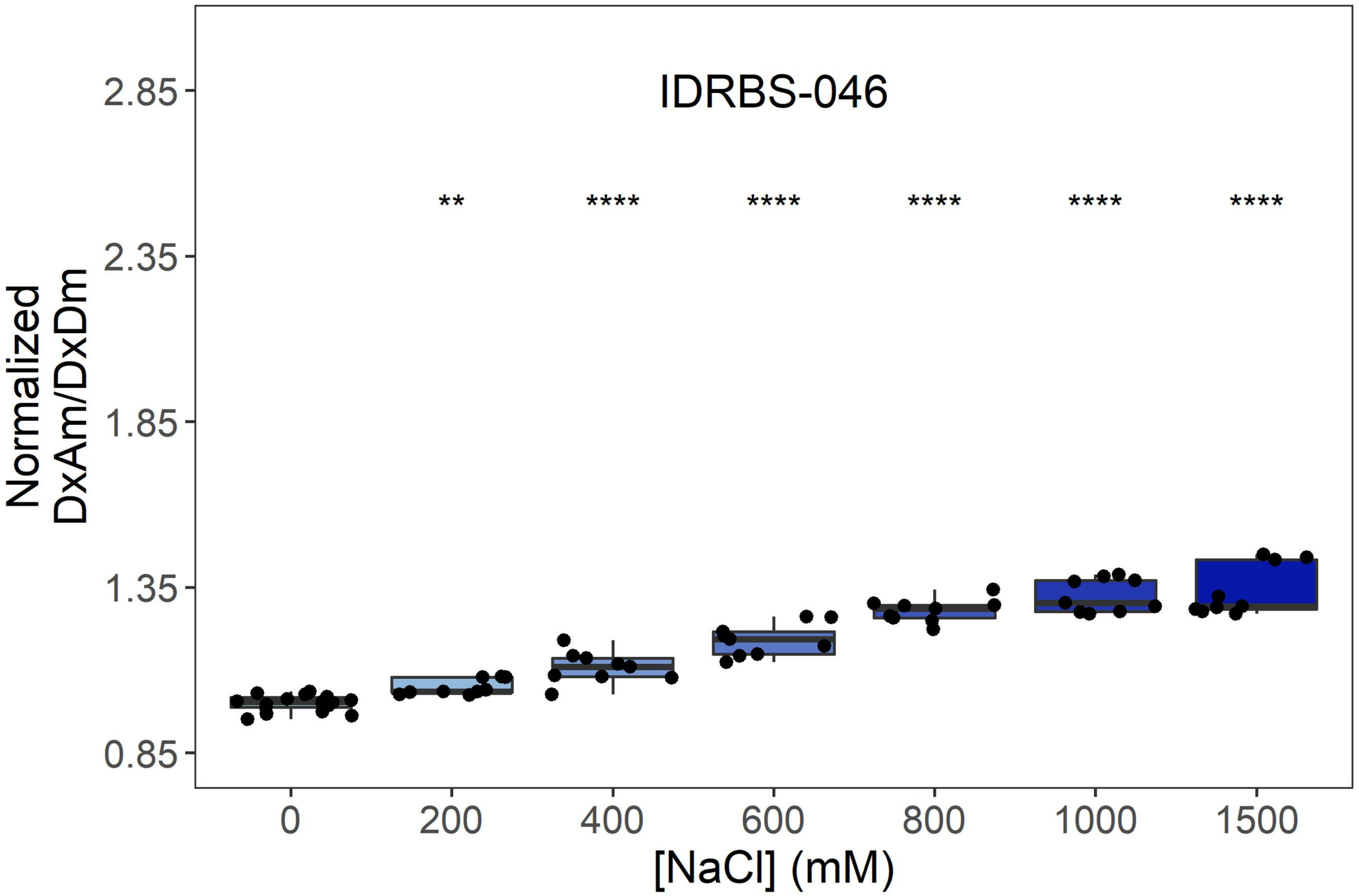
IDRBS-046
Normalized DxAm/DxDm
2.85
2.35
1.85
1.35
0.85
**
****
****
****
****
****
0
200
400
600
800
1000
1500
[NaCl] (mM)

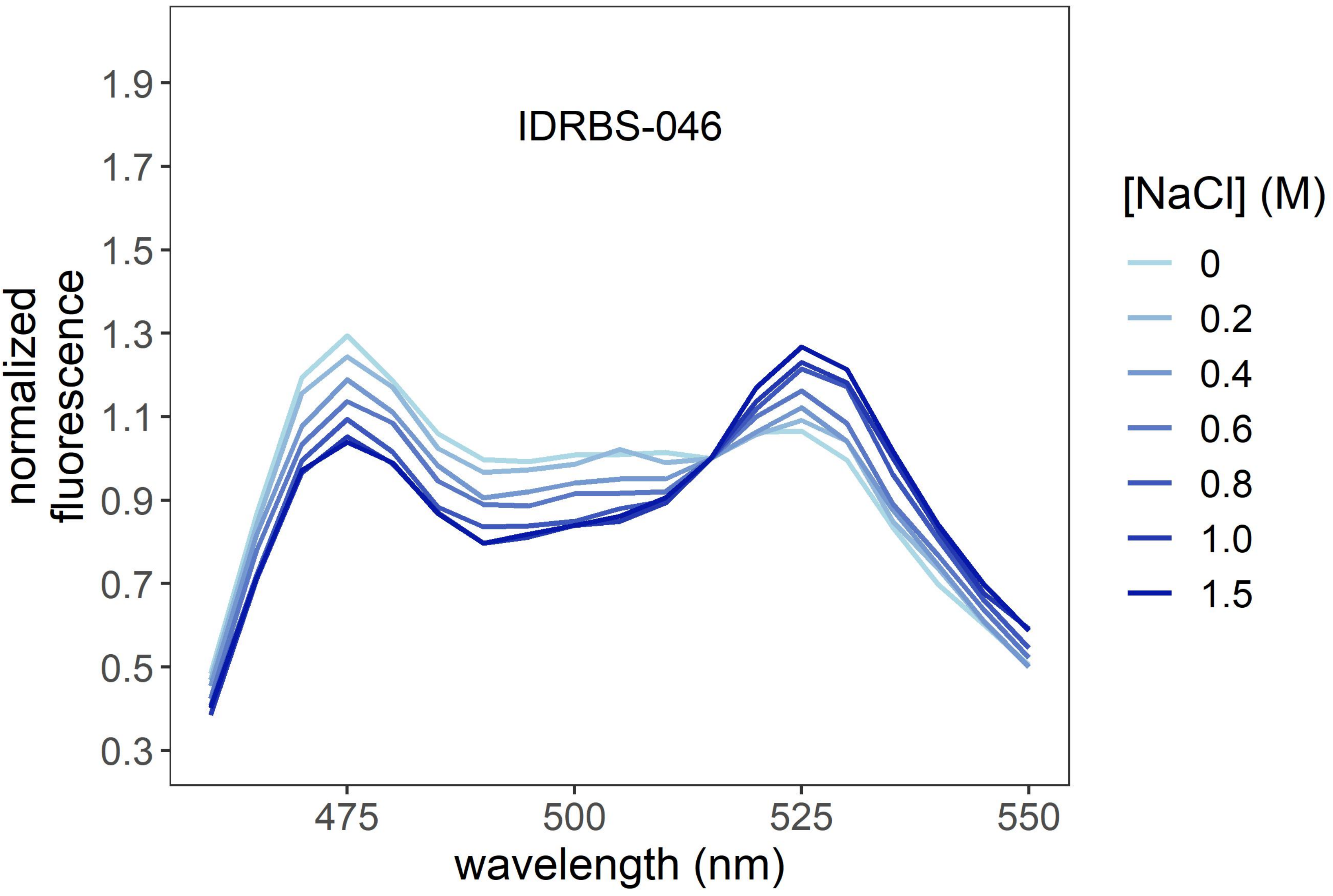
IDRBS-046
normalized fluorescence
1.9
1.7
1.5
1.3
1.1
0.9
0.7
0.5
0.3
475
500
525
550
wavelength (nm)
[NaCl] (M)
0
0.2
0.4
0.6
0.8
1.0
1.5

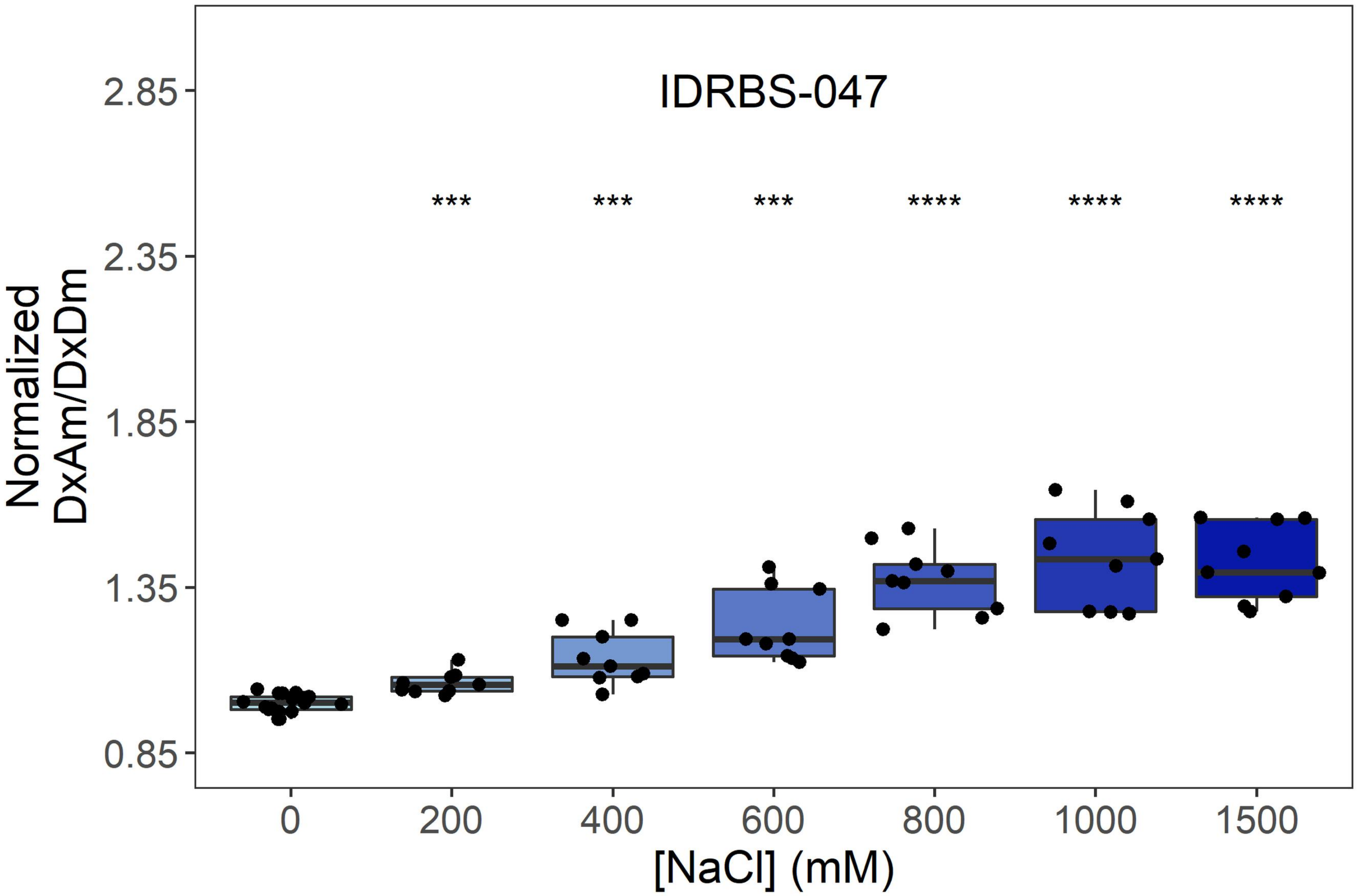
IDRBS-047
***
***
***
****
****
****
Normalized DxAm/DxDm
2.85
2.35
1.85
1.35
0.85
0
200
400
600
800
1000
1500
[NaCl] (mM)

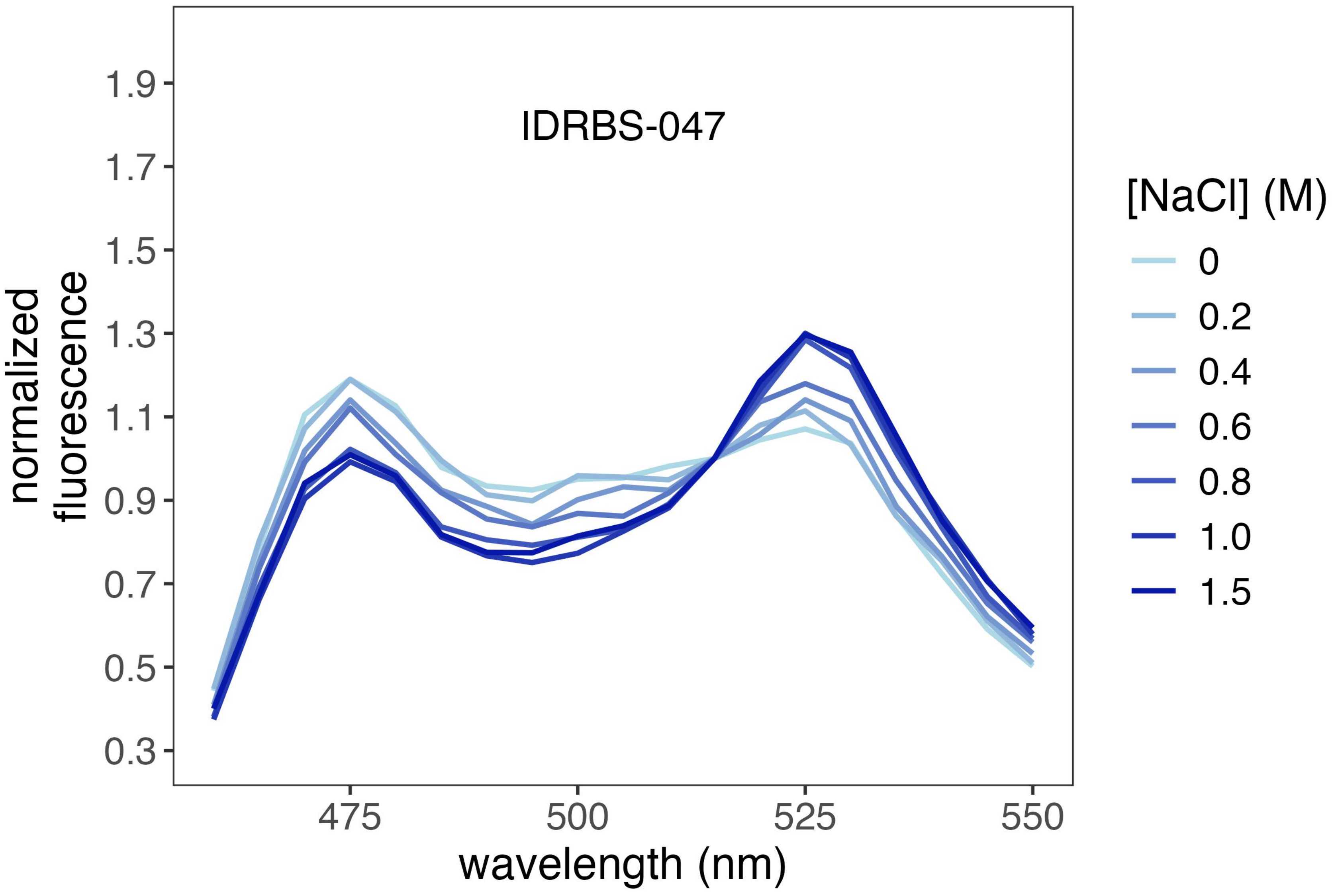
IDRBS-047
normalized fluorescence
1.9
1.7
1.5
1.3
1.1
0.9
0.7
0.5
0.3
475
500
525
550
wavelength (nm)
[NaCl] (M)
0
0.2
0.4
0.6
0.8
1.0
1.5

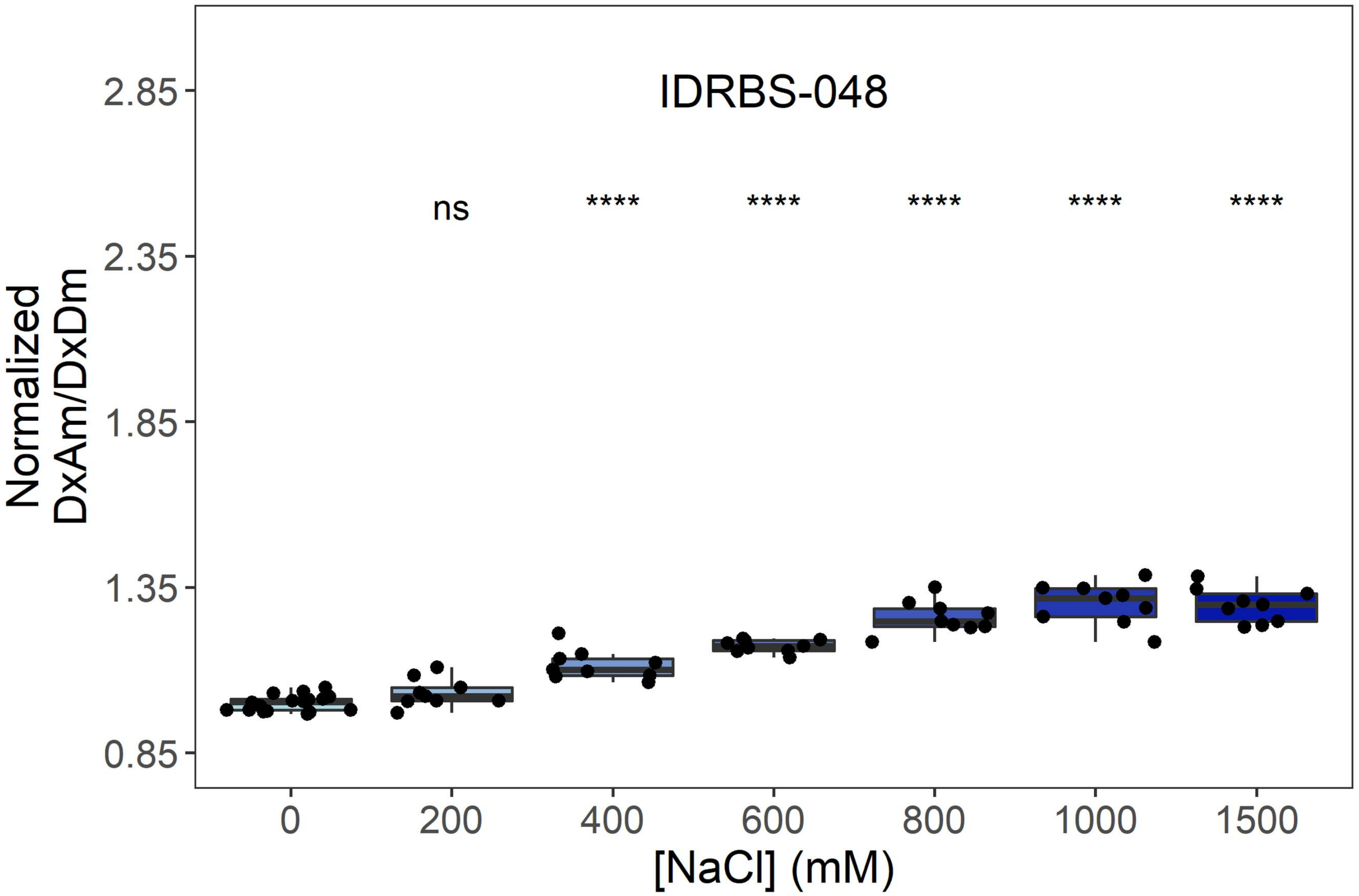
IDRBS-048
ns
****
****
****
****
****
Normalized DxAm/DxDm
2.85
2.35
1.85
1.35
0.85
0
200
400
600
800
1000
1500
[NaCl] (mM)

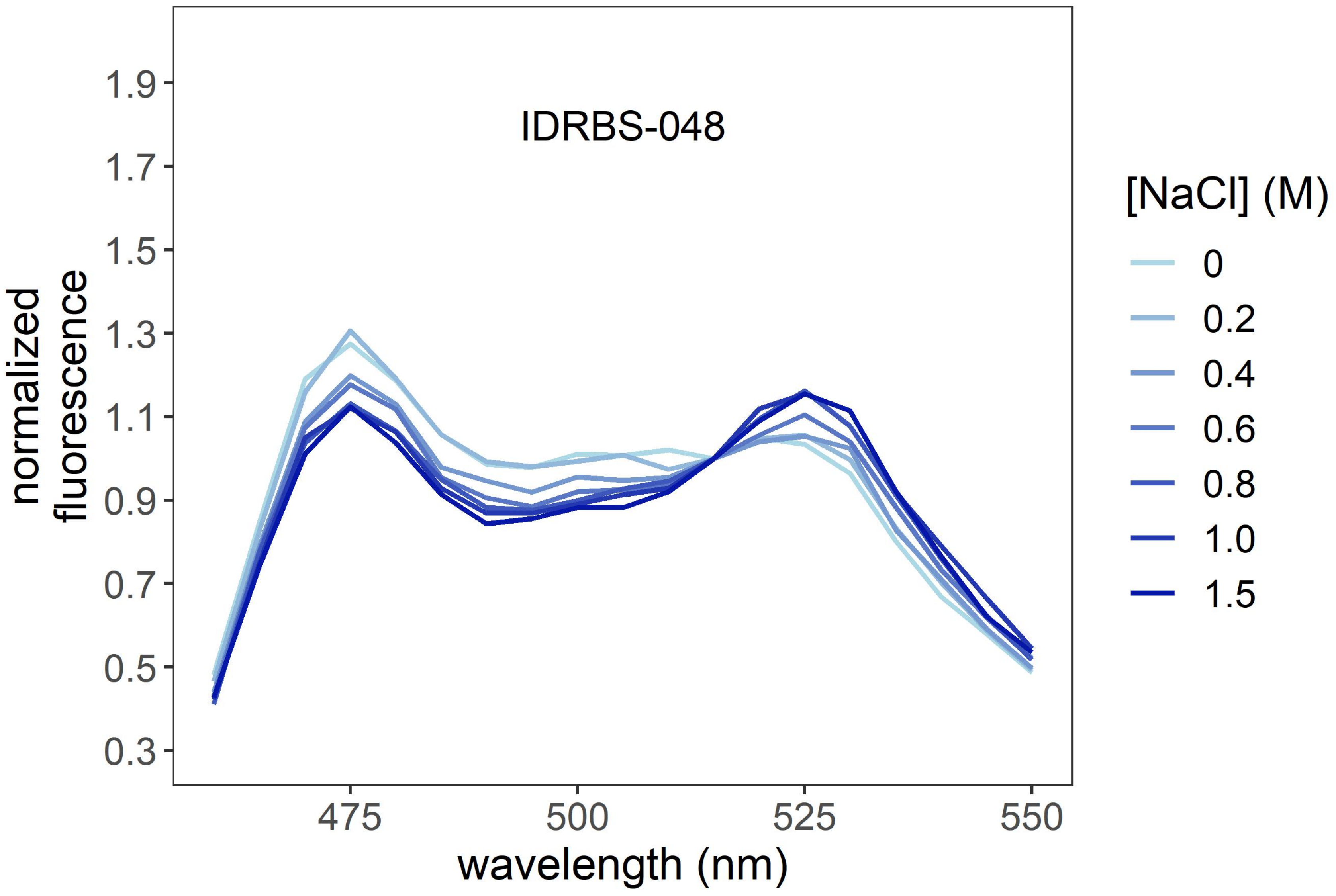
IDRBS-048
normalized fluorescence
1.9
1.7
1.5
1.3
1.1
0.9
0.7
0.5
0.3
475
500
525
550
wavelength (nm)
[NaCl] (M)
0
0.2
0.4
0.6
0.8
1.0
1.5

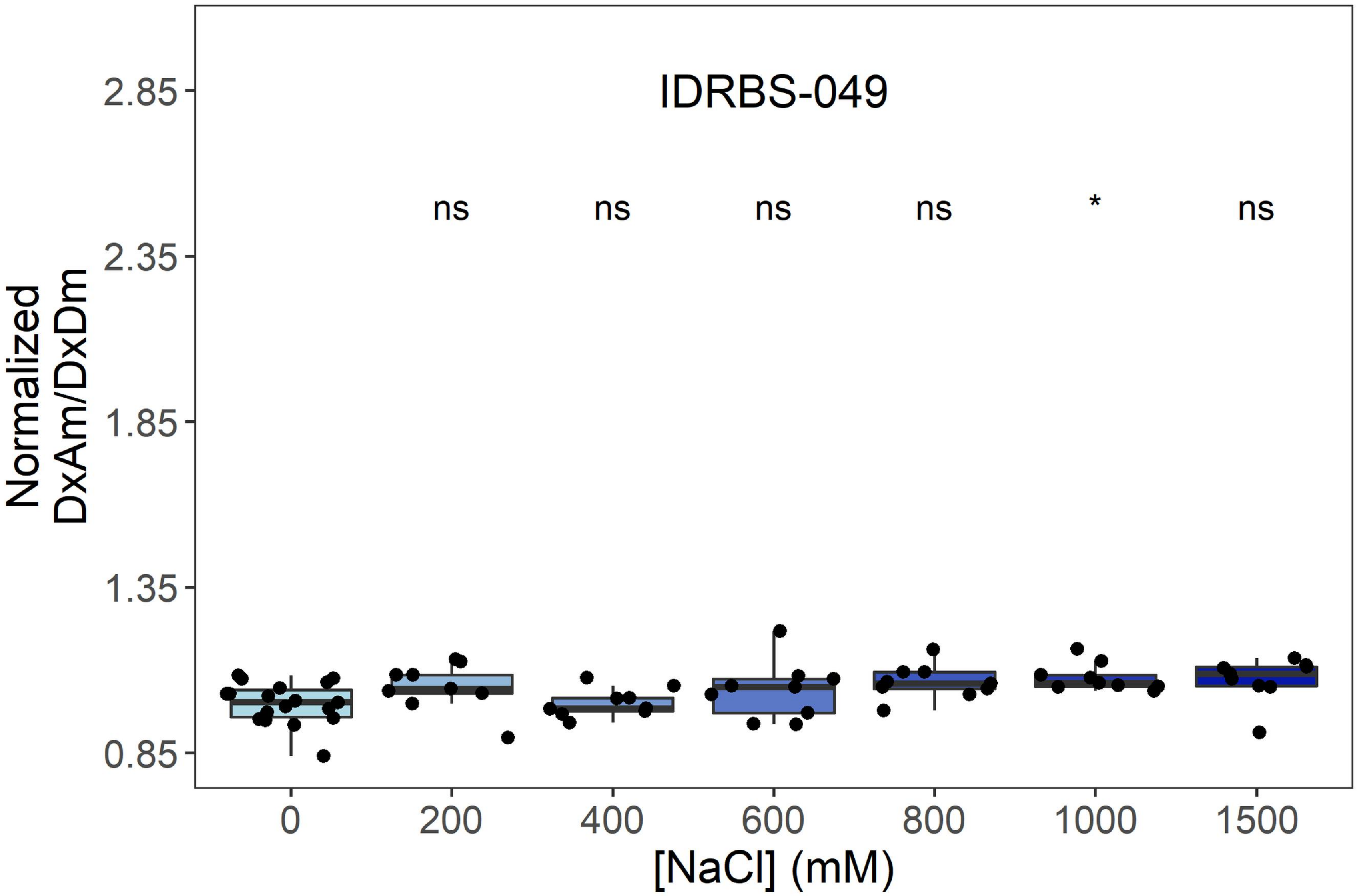
IDRBS-049
ns
ns
ns
ns
*
ns
Normalized DxAm/DxDm
2.85
2.35
1.85
1.35
0.85
0
200
400
600
800
1000
1500
[NaCl] (mM)

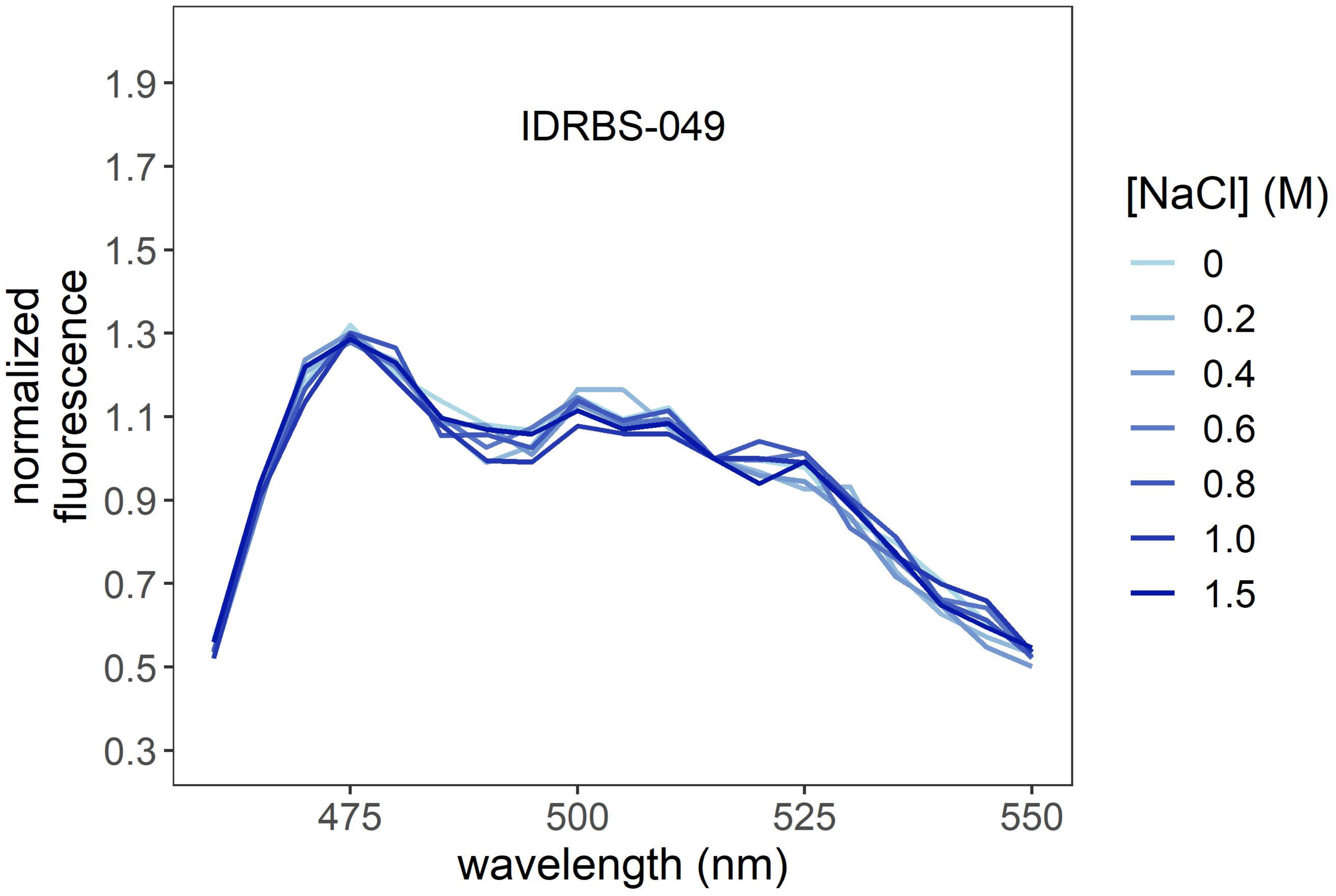
IDRBS-049
normalized fluorescence
1.9
1.7
1.5
1.3
1.1
0.9
0.7
0.5
0.3
475
500
525
550
wavelength (nm)
[NaCl] (M)
0
0.2
0.4
0.6
0.8
1.0
1.5

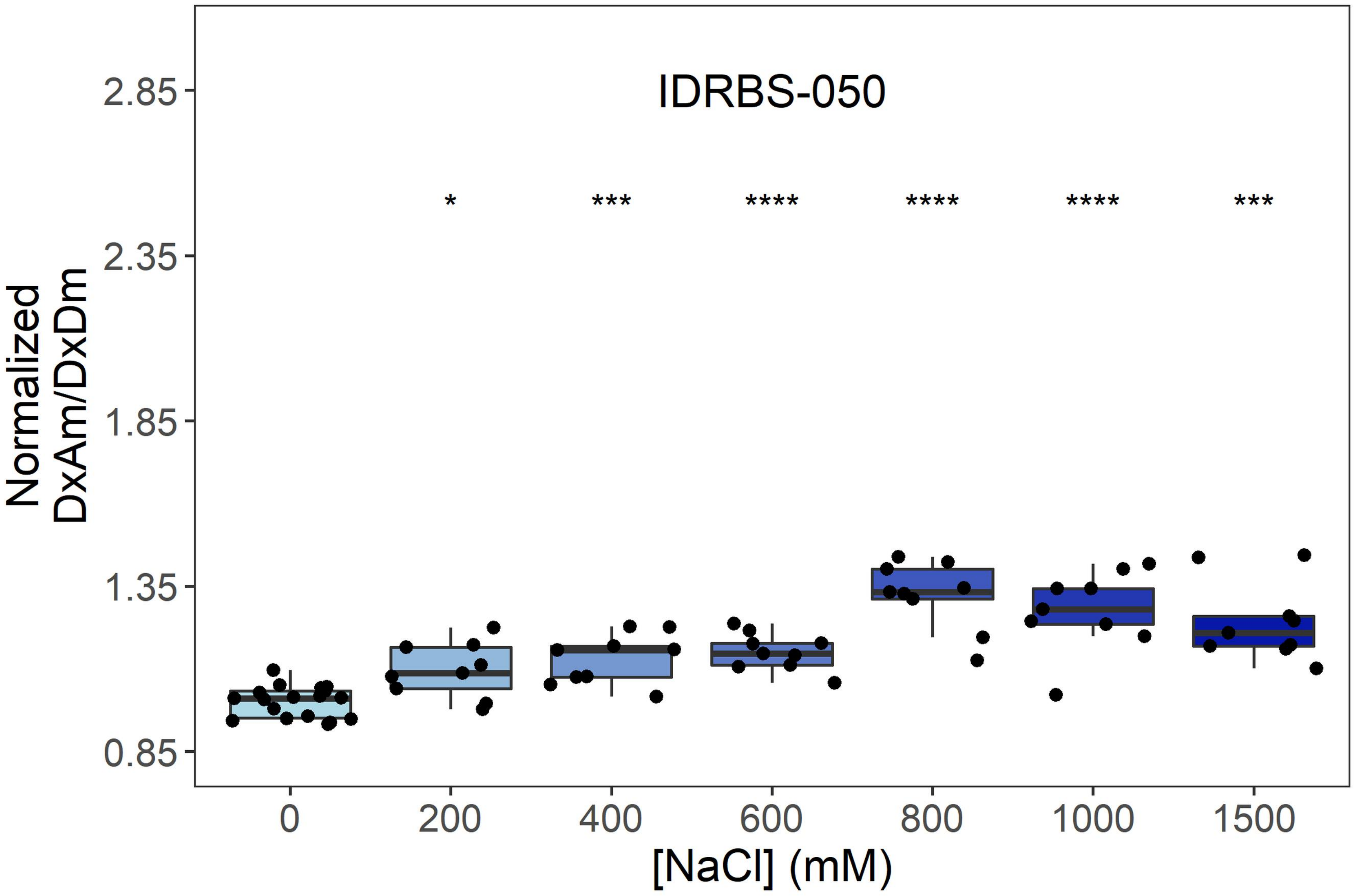
IDRBS-050
Normalized
DxAm/DxDm
2.85
2.35
1.85
1.35
0.85
*
***
****
****
****
***
0
200
400
600
800
1000
1500
[NaCl] (mM)

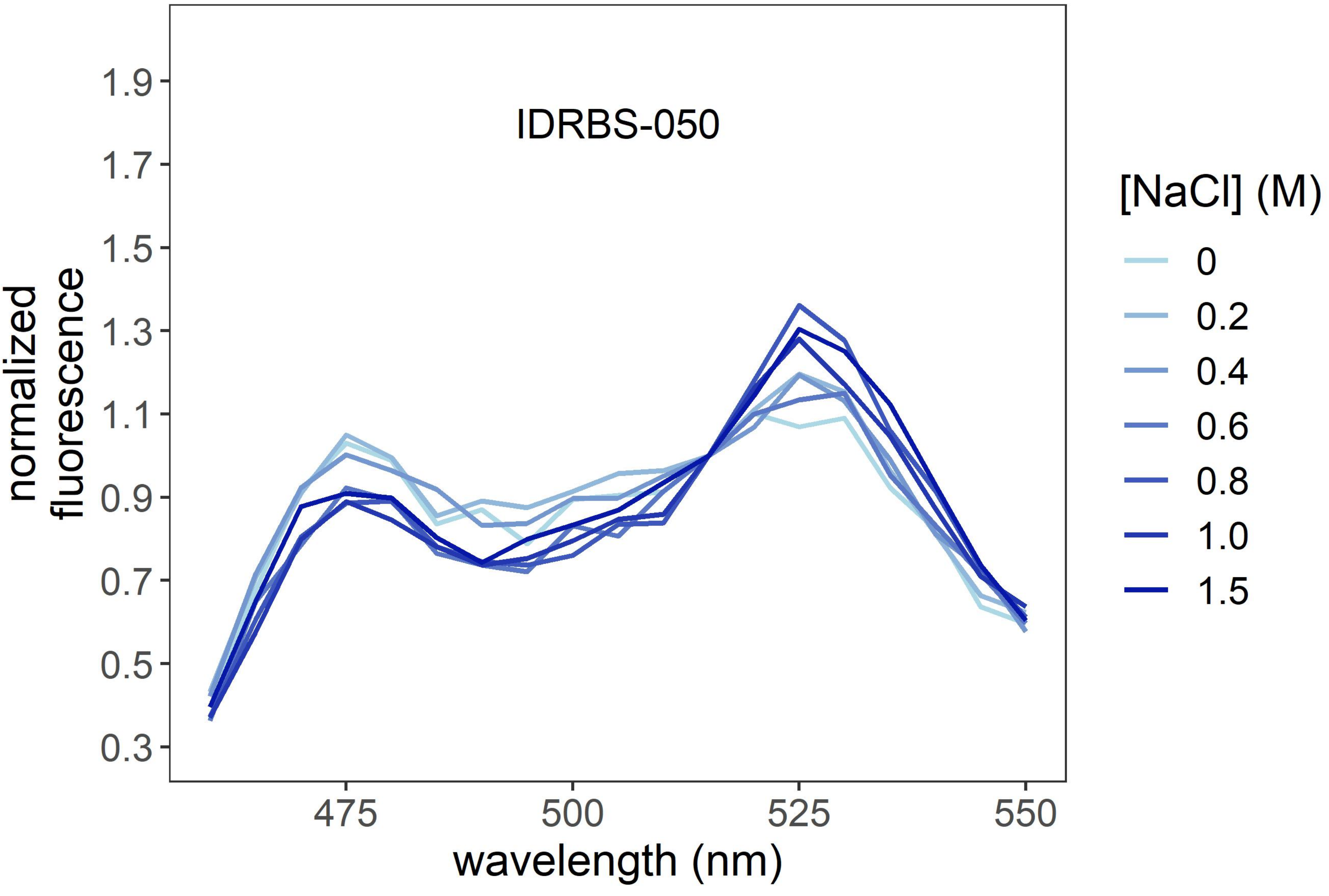
IDRBS-050
normalized
fluorescence
1.9
1.7
1.5
1.3
1.1
0.9
0.7
0.5
0.3
475
500
525
550
wavelength (nm)
[NaCl] (M)
0
0.2
0.4
0.6
0.8
1.0
1.5

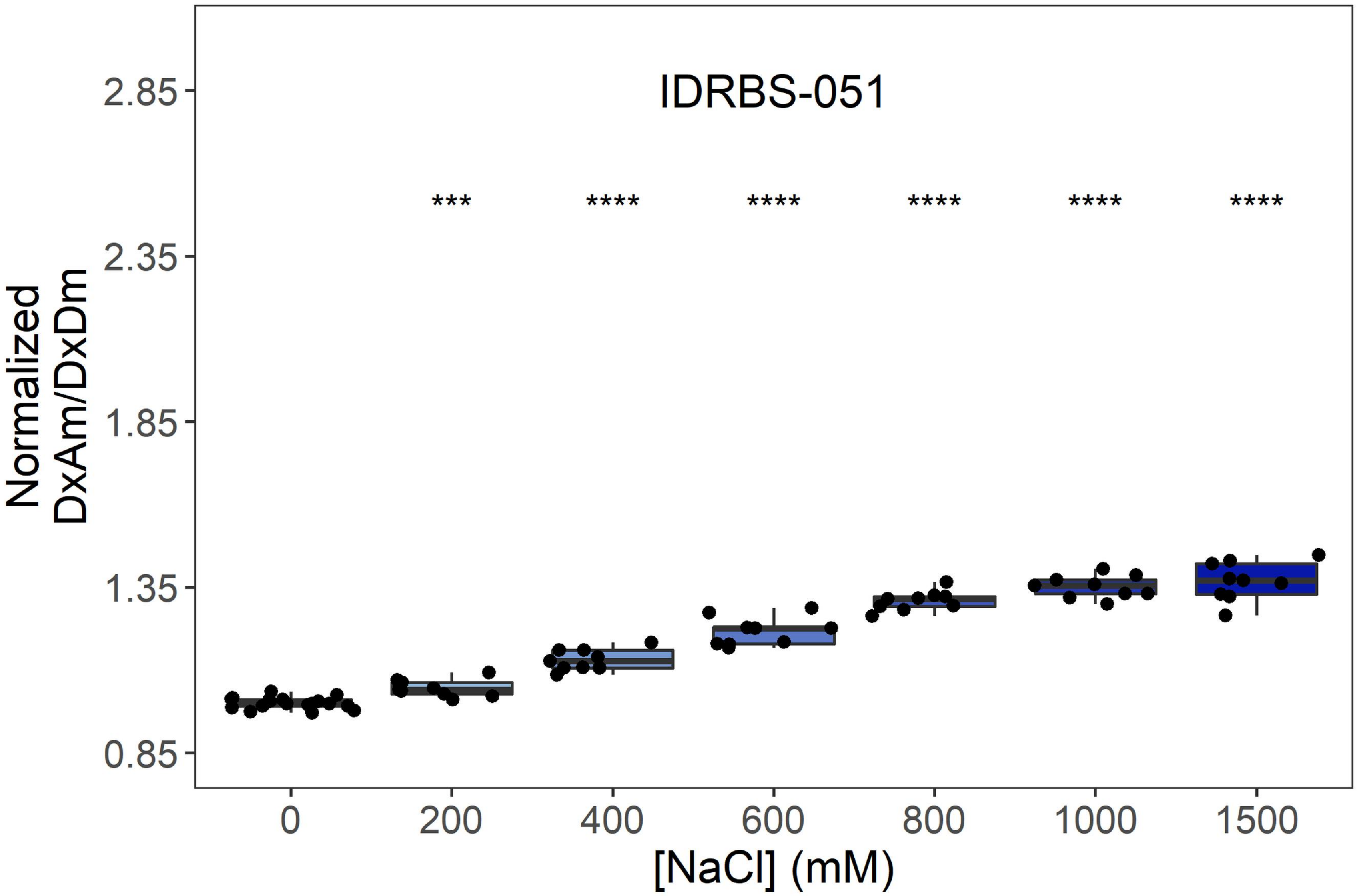

IDRBS-051
***
****
****
****
****
****
2.85
2.35
1.85
1.35
0.85
Normalized DxAm/DxDm
0
200
400
600
800
1000
1500
[NaCl] (mM)


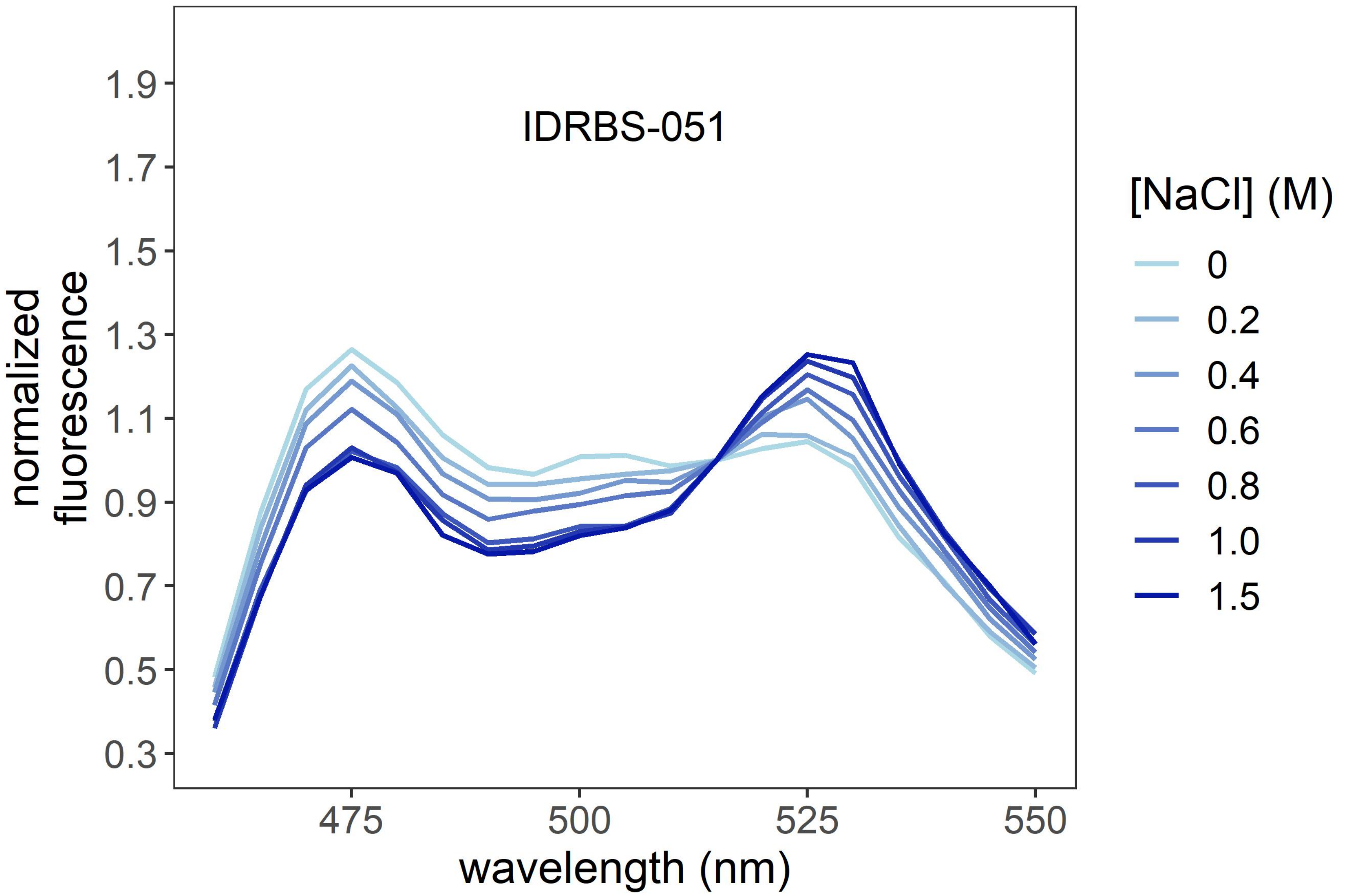

IDRBS-051
1.9
1.7
1.5
1.3
1.1
0.9
0.7
0.5
0.3
normalized fluorescence
475
500
525
550
wavelength (nm)
[NaCl] (M)
0
0.2
0.4
0.6
0.8
1.0
1.5

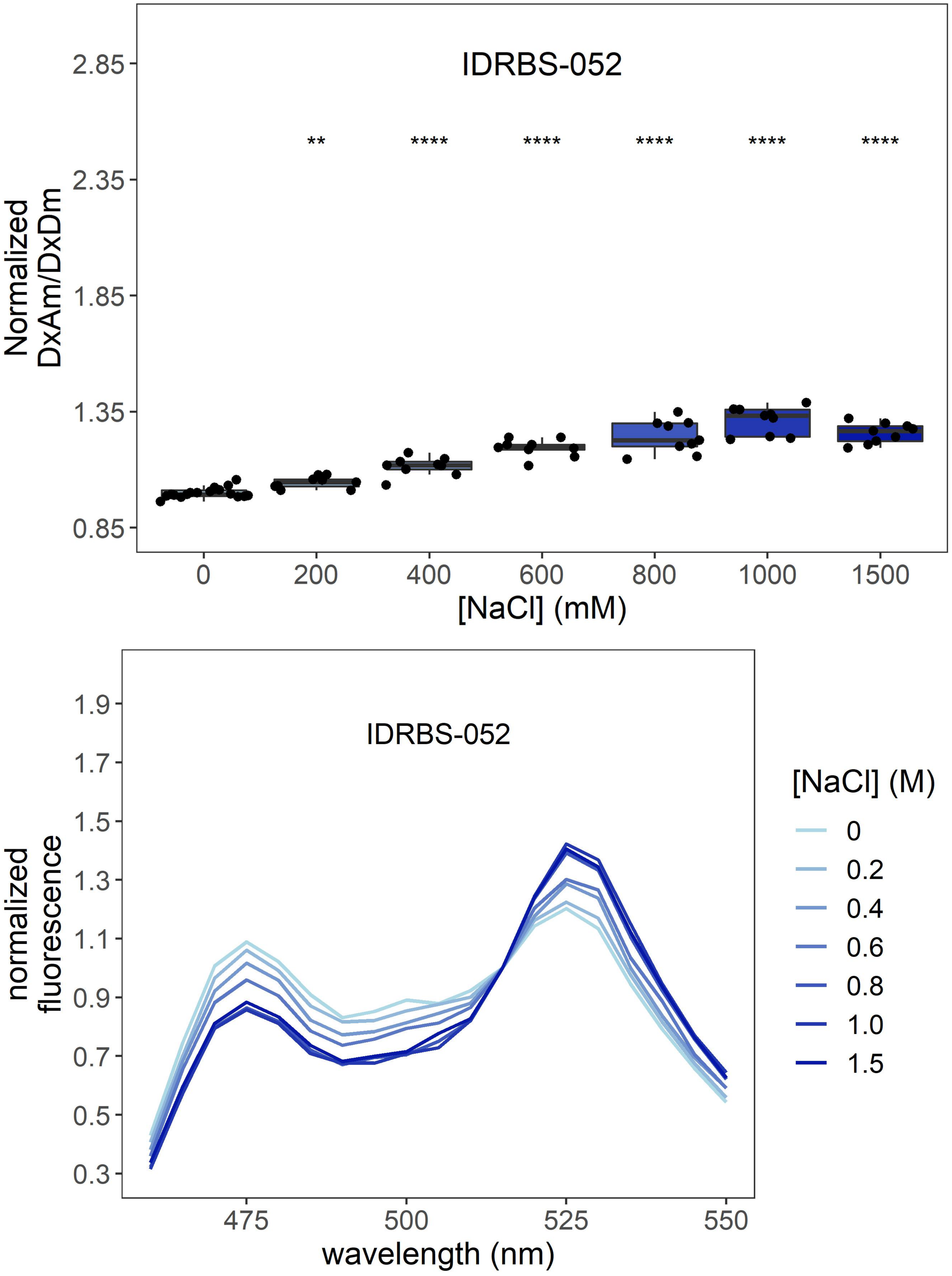

IDRBS-052
**
****
****
****
****
****
Normalized DxAm/DxDm
2.85
2.35
1.85
1.35
0.85
0
200
400
600
800
1000
1500
[NaCl] (mM)
IDRBS-052
normalized fluorescence
1.9
1.7
1.5
1.3
1.1
0.9
0.7
0.5
0.3
475
500
525
550
wavelength (nm)
[NaCl] (M)
0
0.2
0.4
0.6
0.8
1.0
1.5

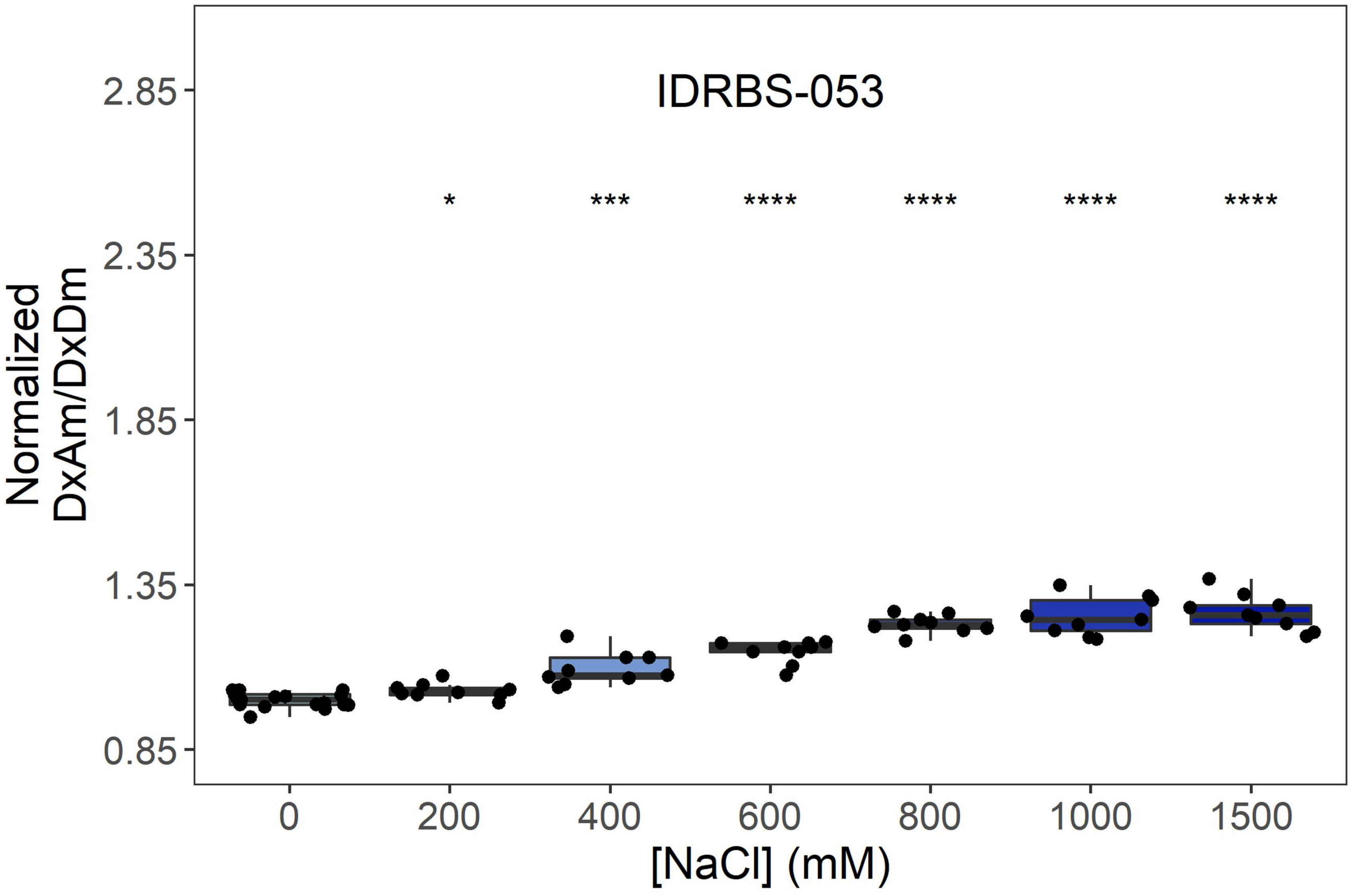
IDRBS-053
Normalized DxAm/DxDm
*
***
****
****
****
****
2.85
2.35
1.85
1.35
0.85
0
200
400
600
800
1000
1500
[NaCl] (mM)

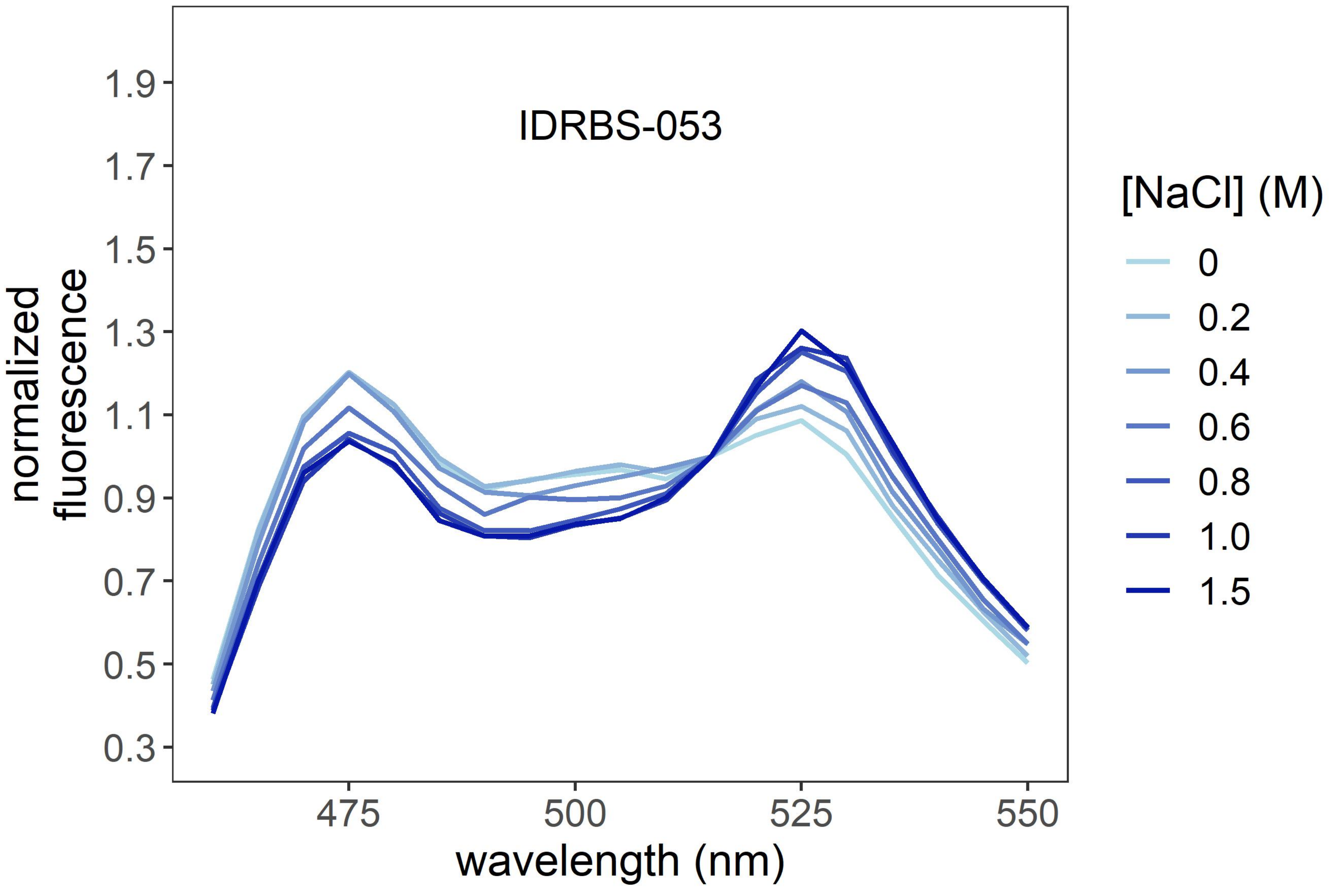
IDRBS-053
normalized fluorescence
1.9
1.7
1.5
1.3
1.1
0.9
0.7
0.5
0.3
475
500
525
550
wavelength (nm)
[NaCl] (M)
0
0.2
0.4
0.6
0.8
1.0
1.5

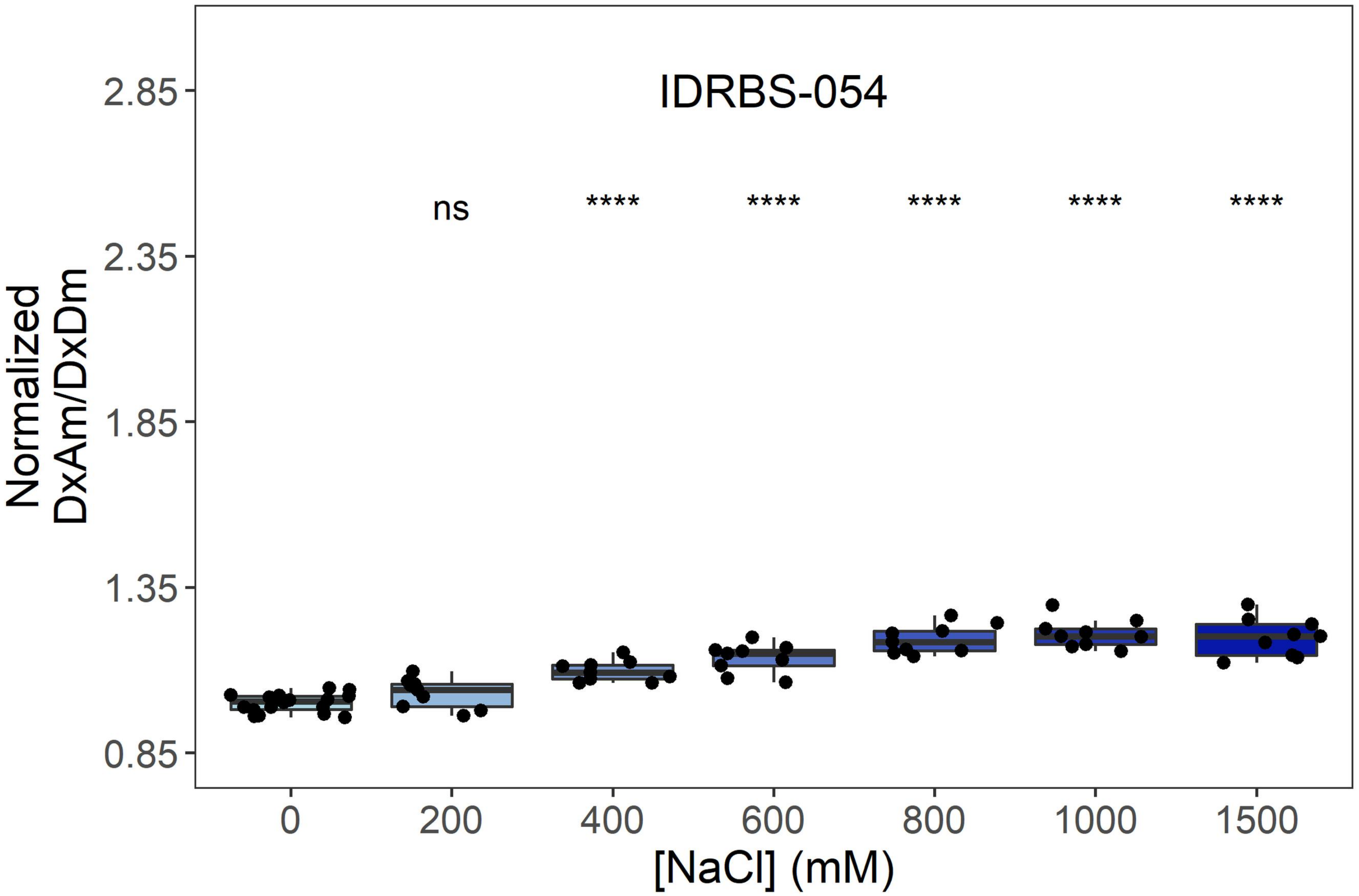

IDRBS-054
Normalized DxAm/DxDm
2.85
2.35
1.85
1.35
0.85
ns
****
****
****
****
****
0
200
400
600
800
1000
1500
[NaCl] (mM)


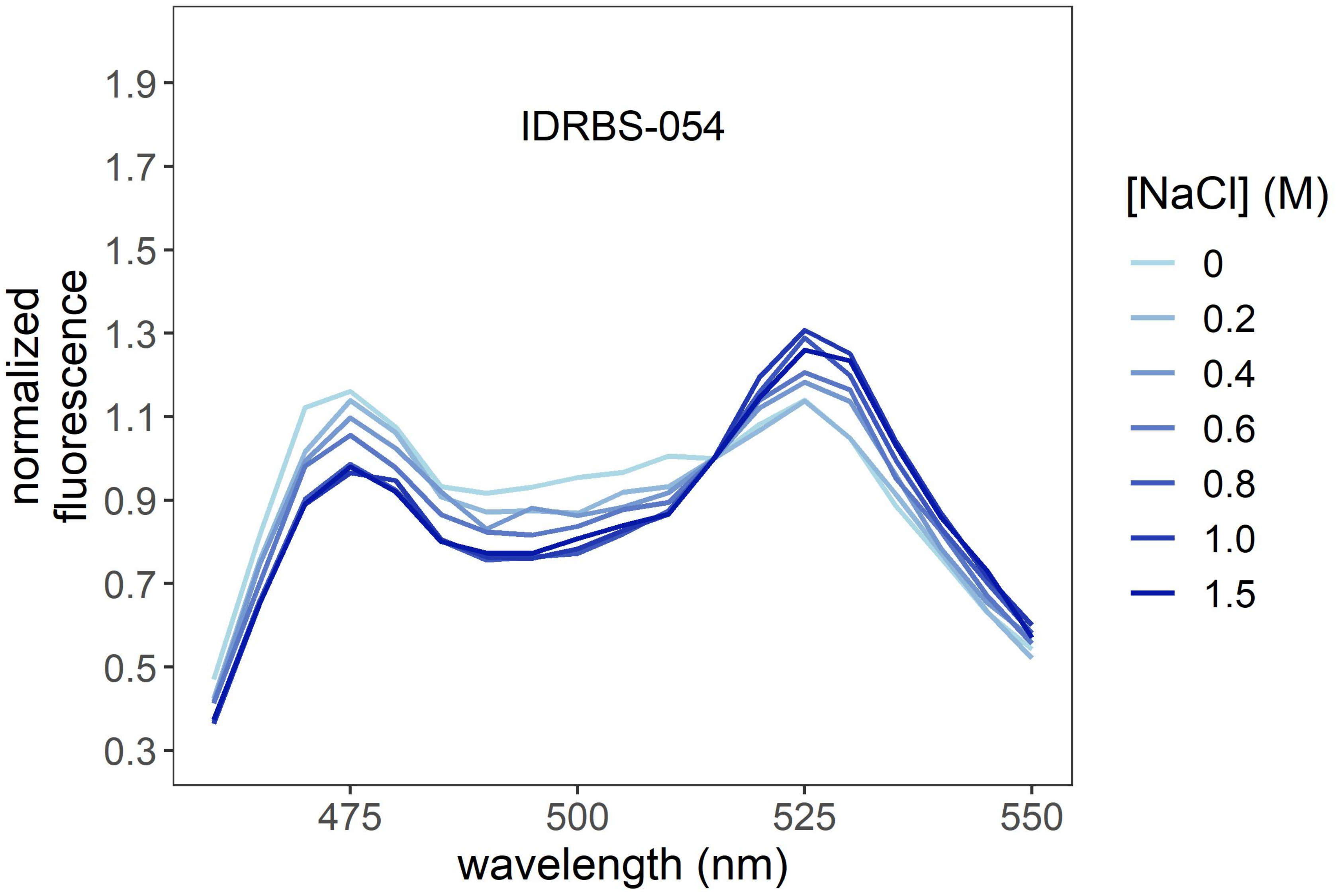

IDRBS-054
normalized fluorescence
1.9
1.7
1.5
1.3
1.1
0.9
0.7
0.5
0.3
475
500
525
550
wavelength (nm)
[NaCl] (M)
0
0.2
0.4
0.6
0.8
1.0
1.5

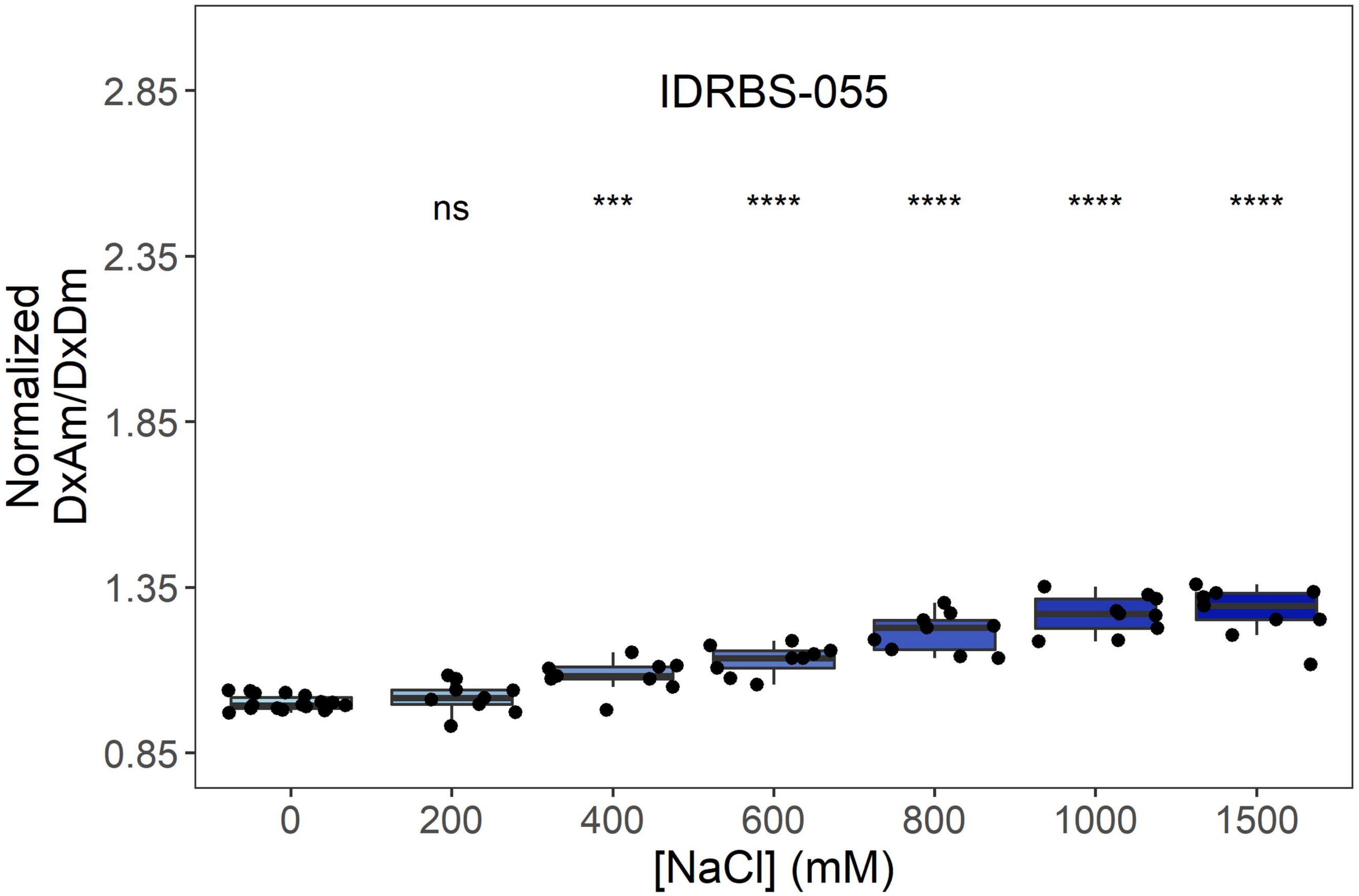

IDRBS-055
ns
***
****
****
****
****
Normalized DxAm/DxDm
2.85
2.35
1.85
1.35
0.85
0
200
400
600
800
1000
1500
[NaCl] (mM)


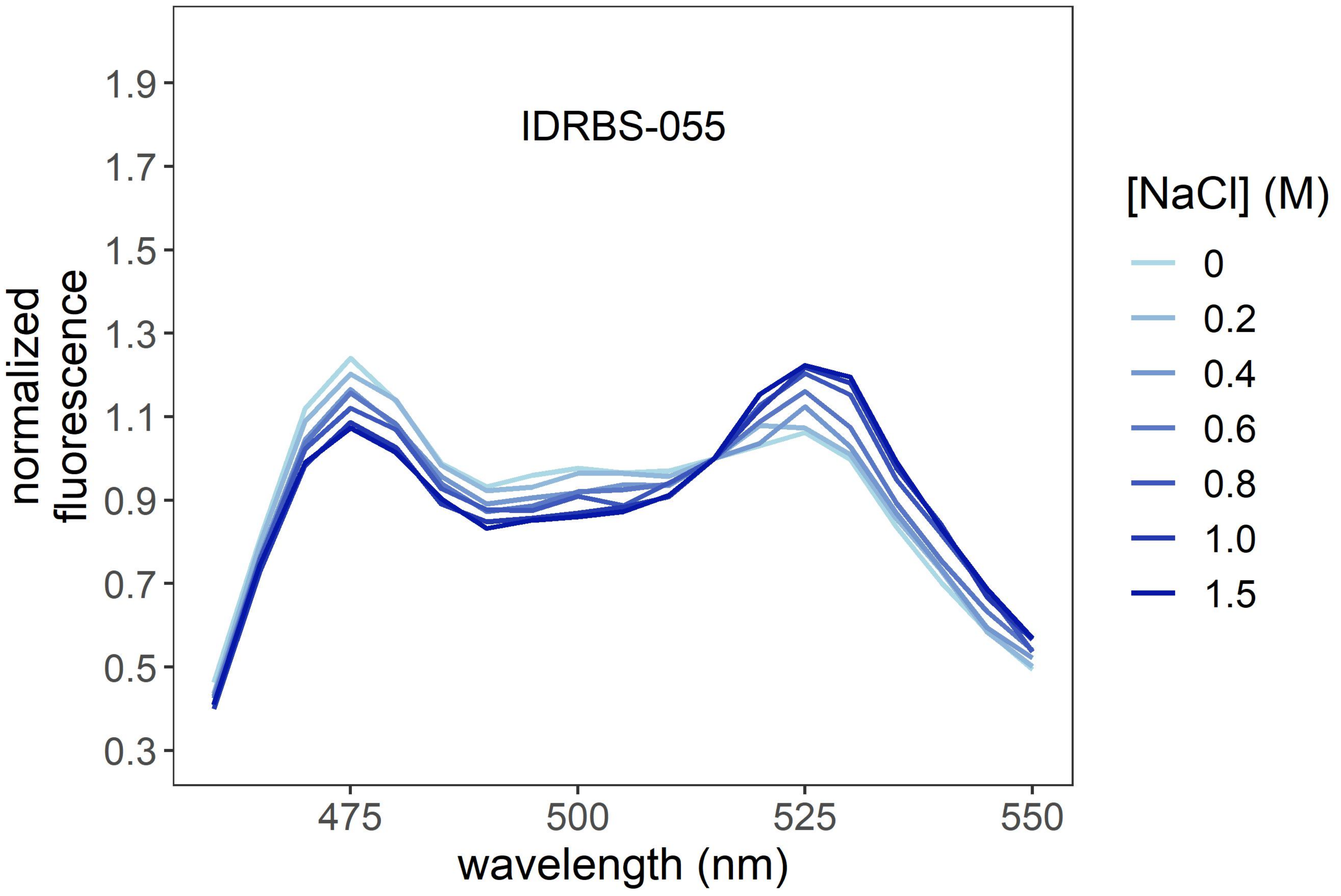

IDRBS-055
normalized fluorescence
1.9
1.7
1.5
1.3
1.1
0.9
0.7
0.5
0.3
475
500
525
550
wavelength (nm)
[NaCl] (M)
0
0.2
0.4
0.6
0.8
1.0
1.5

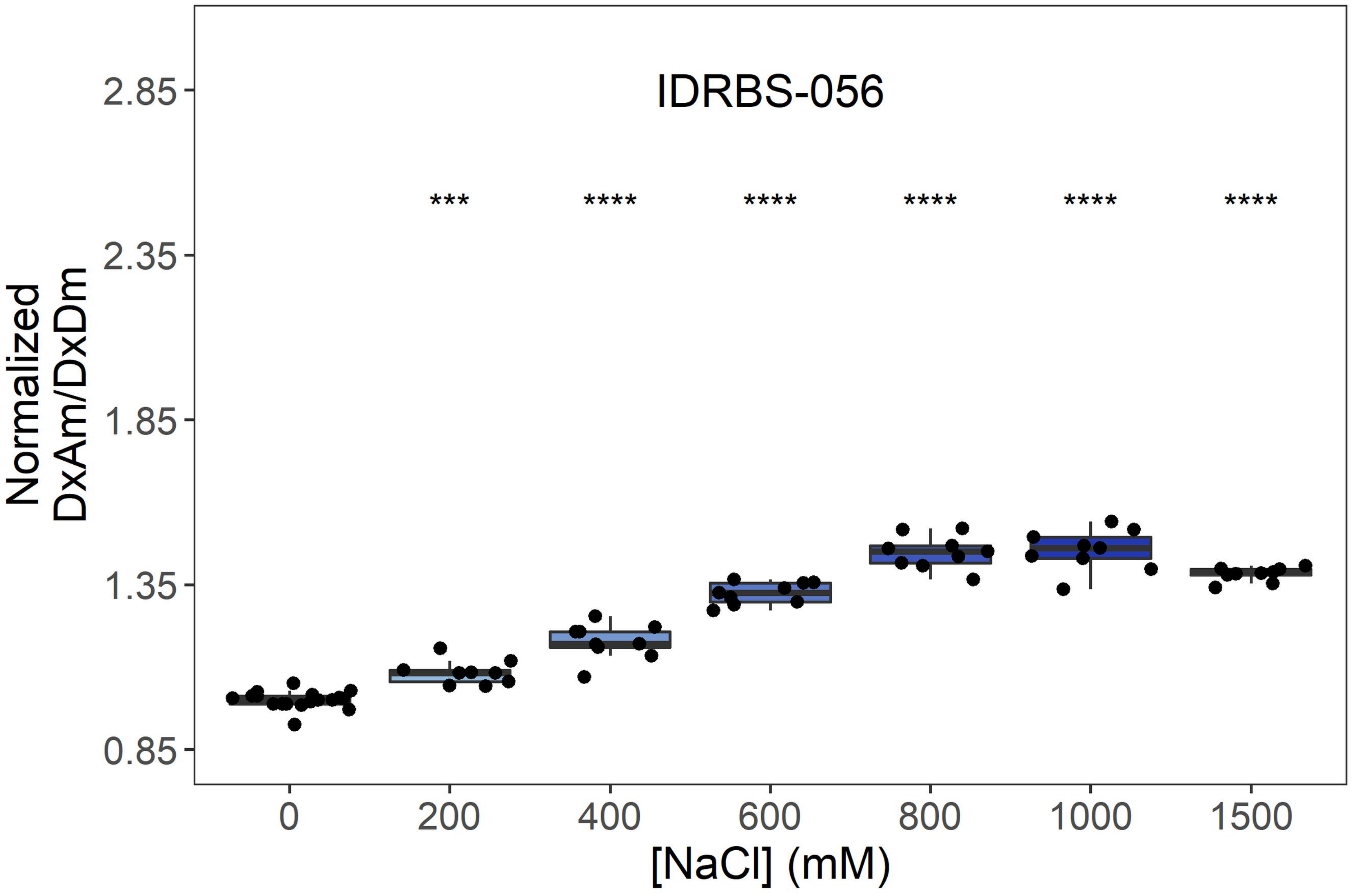
IDRBS-056
Normalized DxAm/DxDm
2.85
2.35
1.85
1.35
0.85
***
****
****
****
****
****
0
200
400
600
800
1000
1500
[NaCl] (mM)

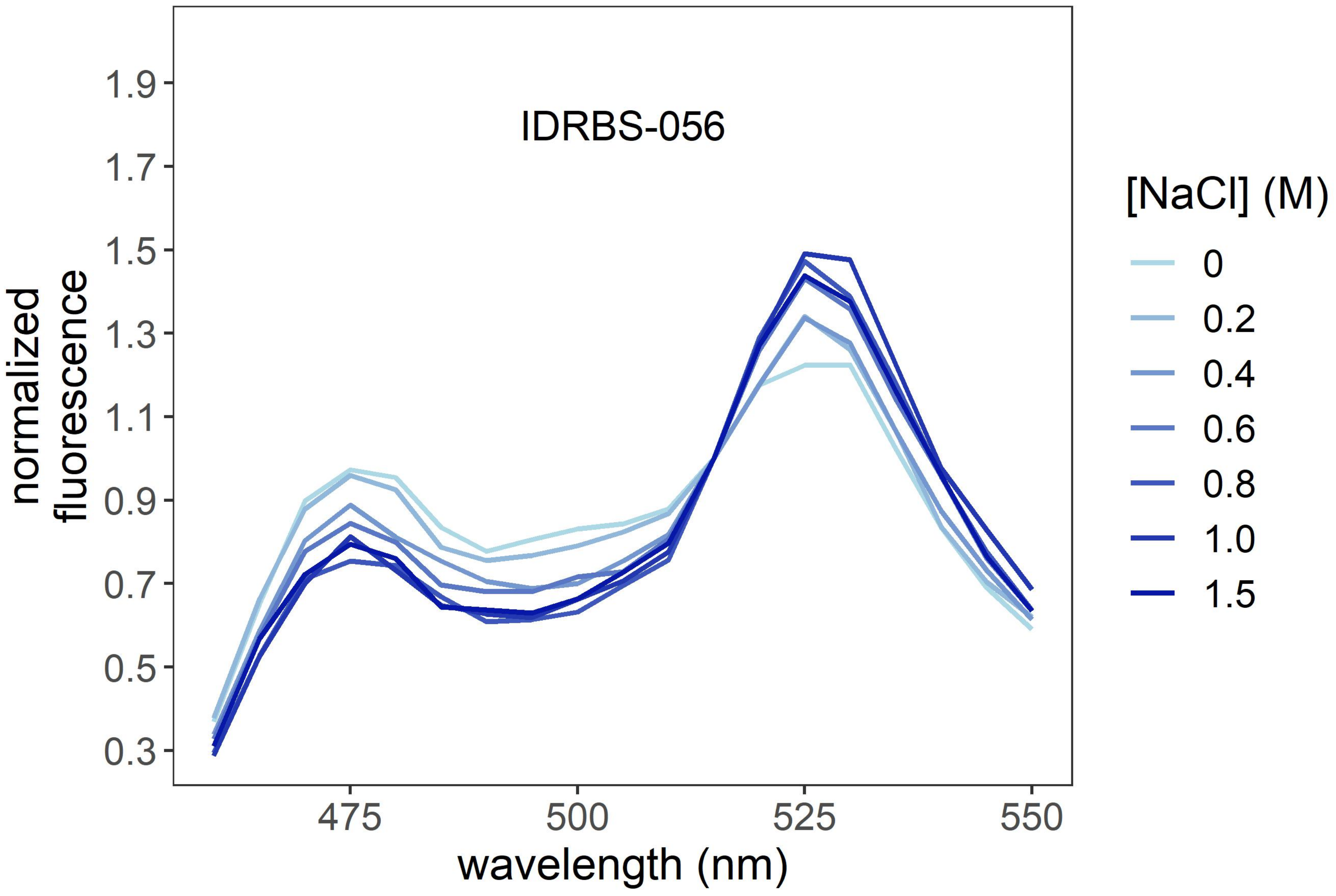
IDRBS-056
normalized fluorescence
1.9
1.7
1.5
1.3
1.1
0.9
0.7
0.5
0.3
475
500
525
550
wavelength (nm)
[NaCl] (M)
0
0.2
0.4
0.6
0.8
1.0
1.5

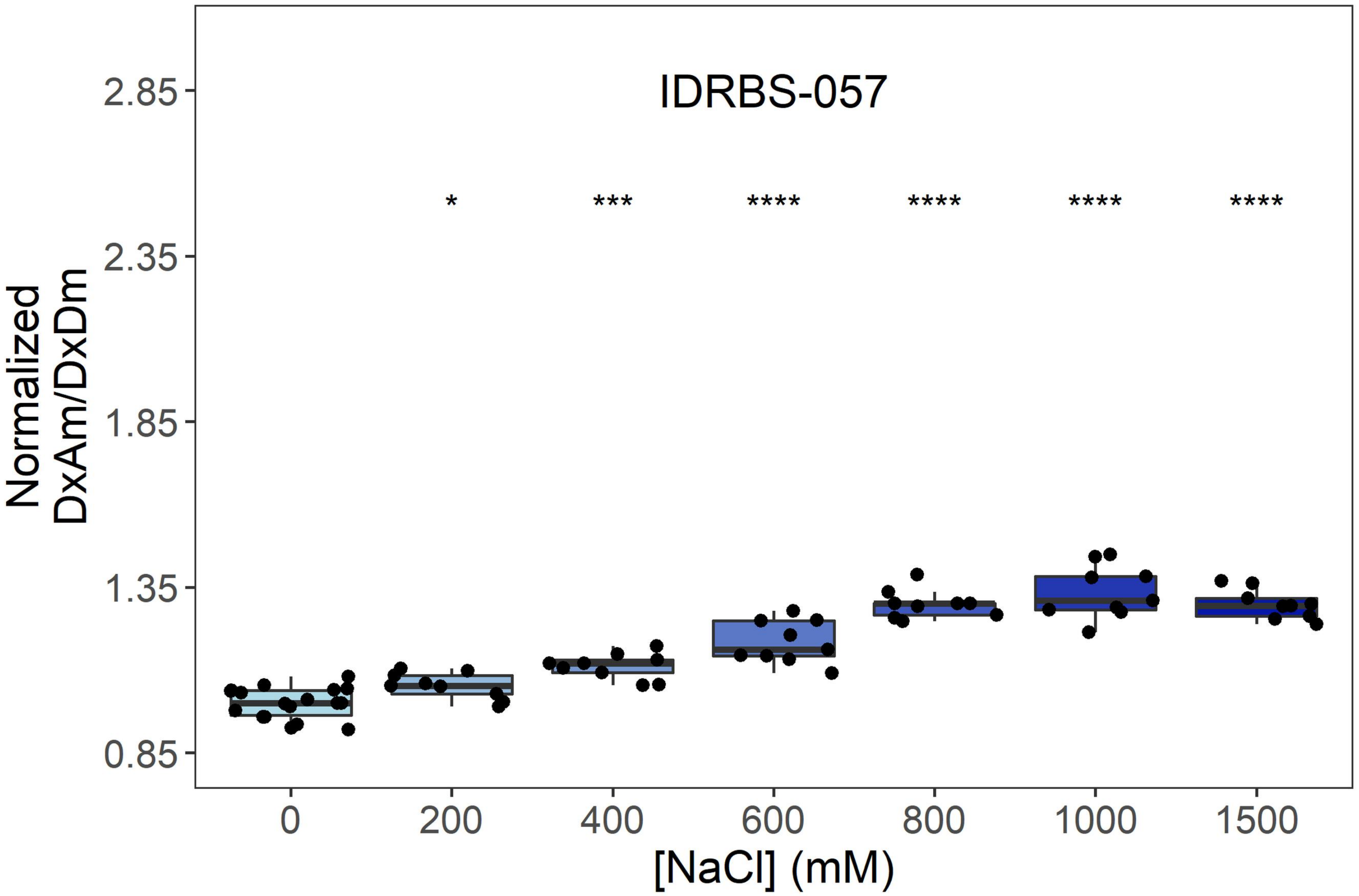
IDRBS-057
Normalized DxAm/DxDm
2.85
2.35
1.85
1.35
0.85
* *** **** **** **** ****
0
200
400
600
800
1000
1500
[NaCl] (mM)

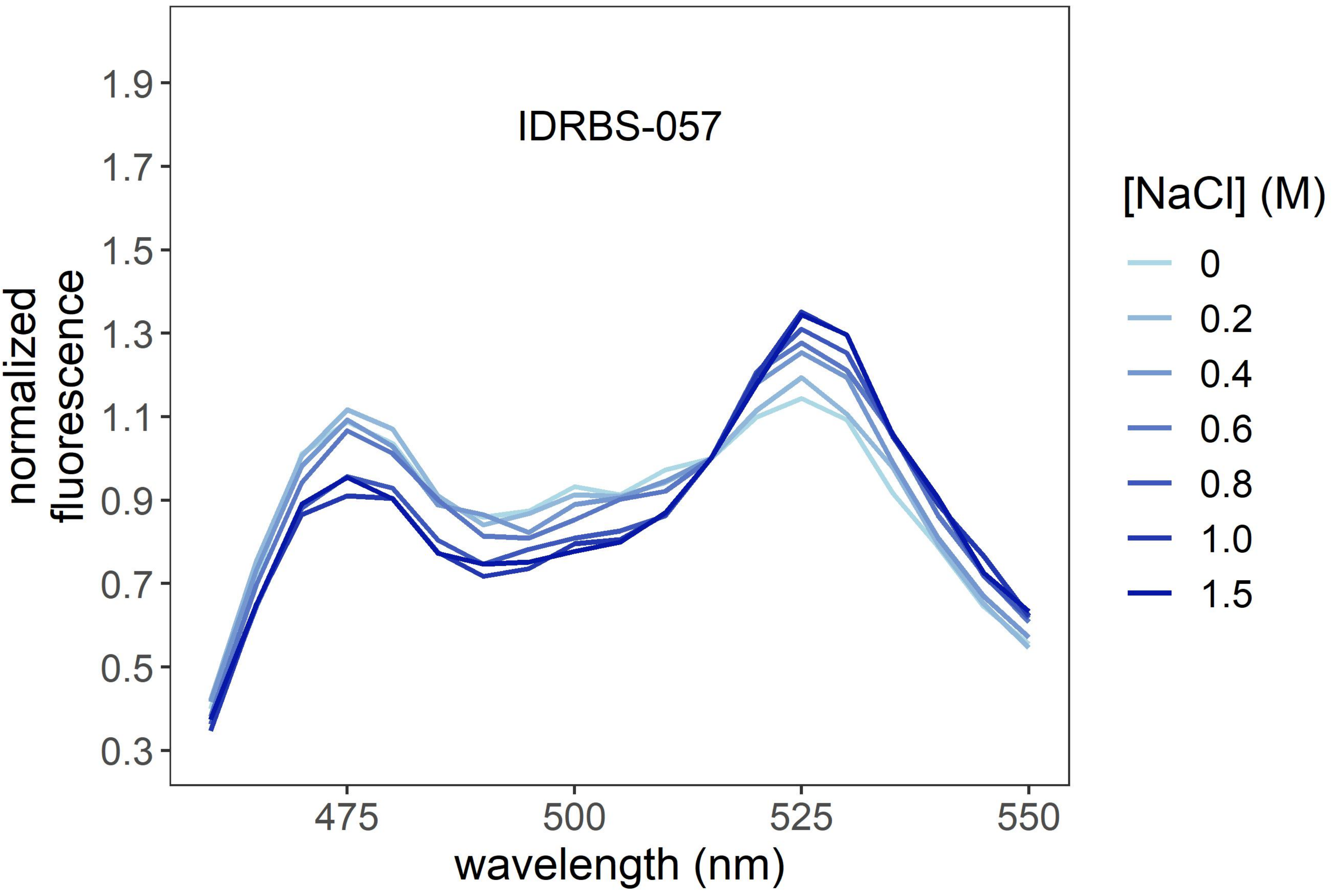
IDRBS-057
normalized fluorescence
1.9
1.7
1.5
1.3
1.1
0.9
0.7
0.5
0.3
475
500
525
550
wavelength (nm)
[NaCl] (M)
0
0.2
0.4
0.6
0.8
1.0
1.5

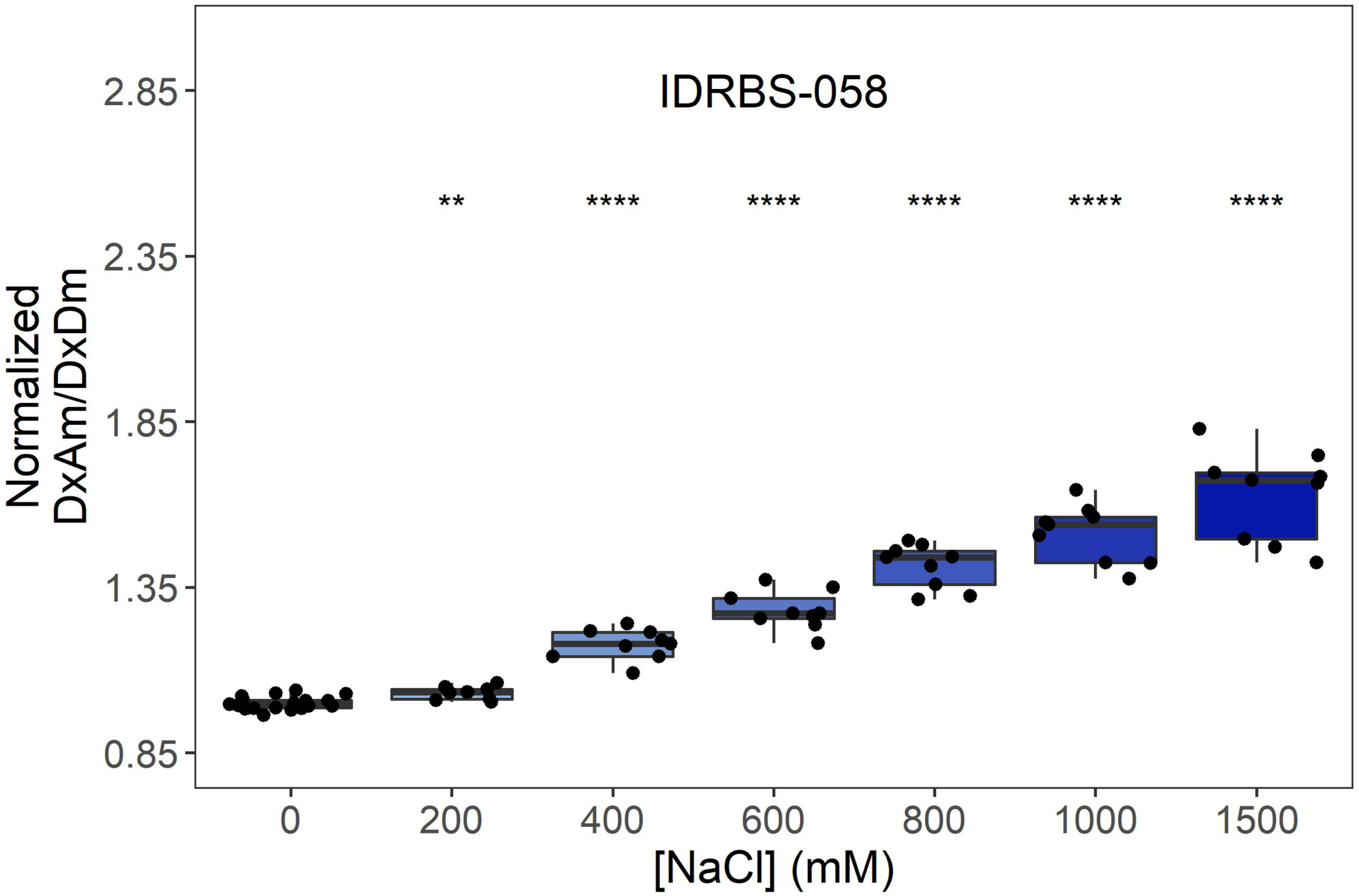
IDRBS-058
**
****
****
****
****
****
Normalized DxAm/DxDm
2.85
2.35
1.85
1.35
0.85
0
200
400
600
800
1000
1500
[NaCl] (mM)

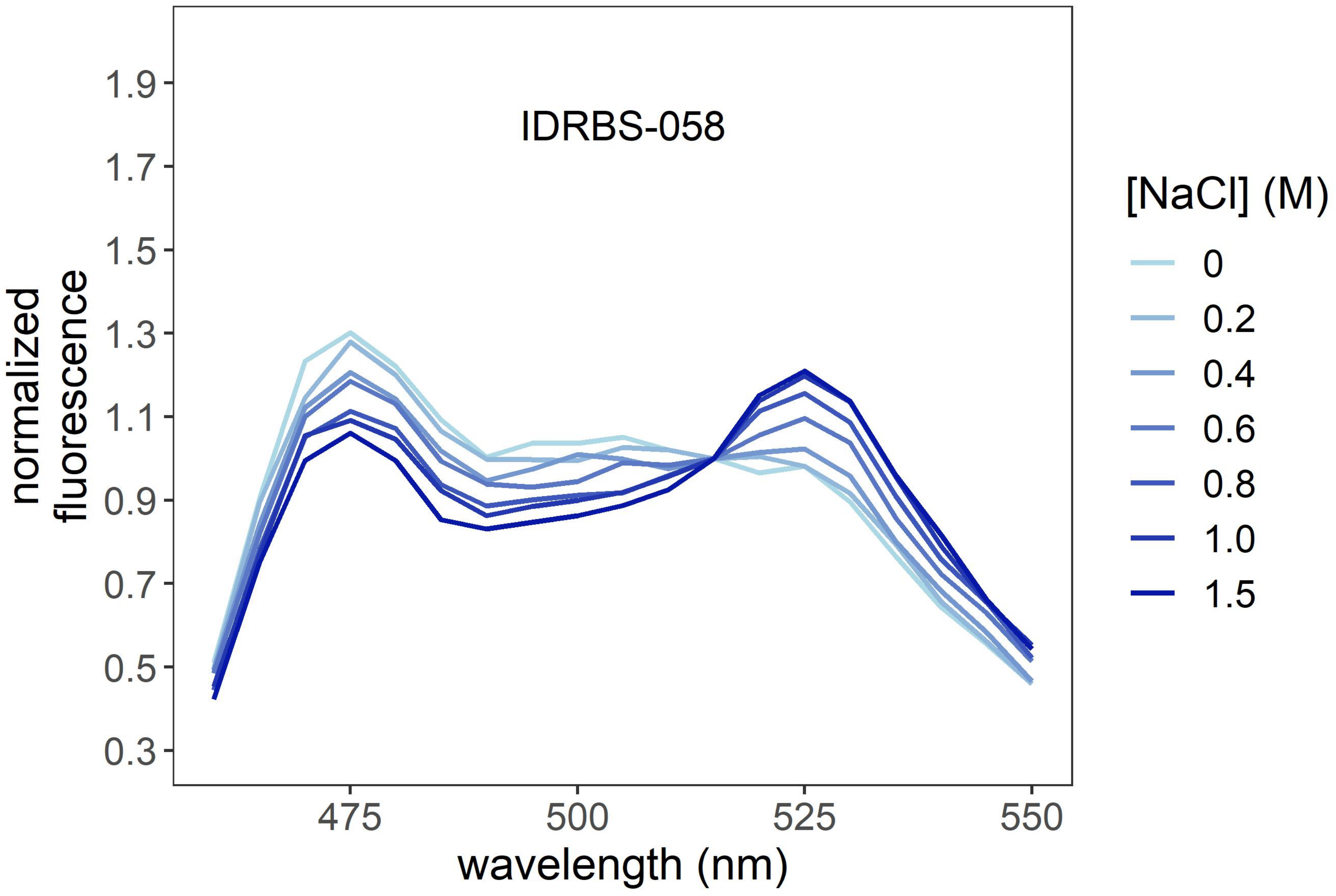
IDRBS-058
normalized fluorescence
1.9
1.7
1.5
1.3
1.1
0.9
0.7
0.5
0.3
475
500
525
550
wavelength (nm)
[NaCl] (M)
0
0.2
0.4
0.6
0.8
1.0
1.5

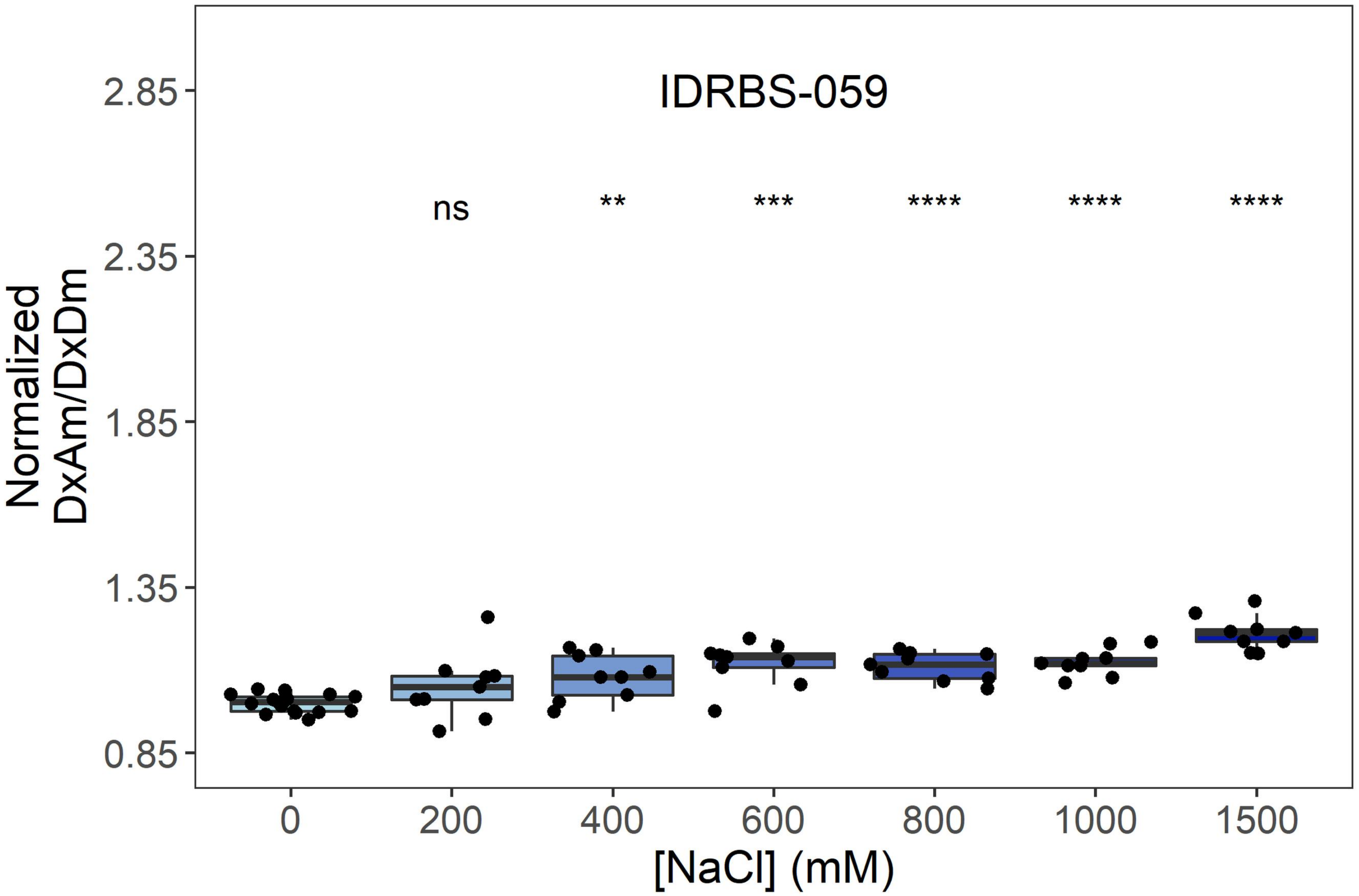
IDRBS-059
ns
**
***
****
****
****
Normalized DxAm/DxDm
2.85
2.35
1.85
1.35
0.85
0
200
400
600
800
1000
1500
[NaCl] (mM)

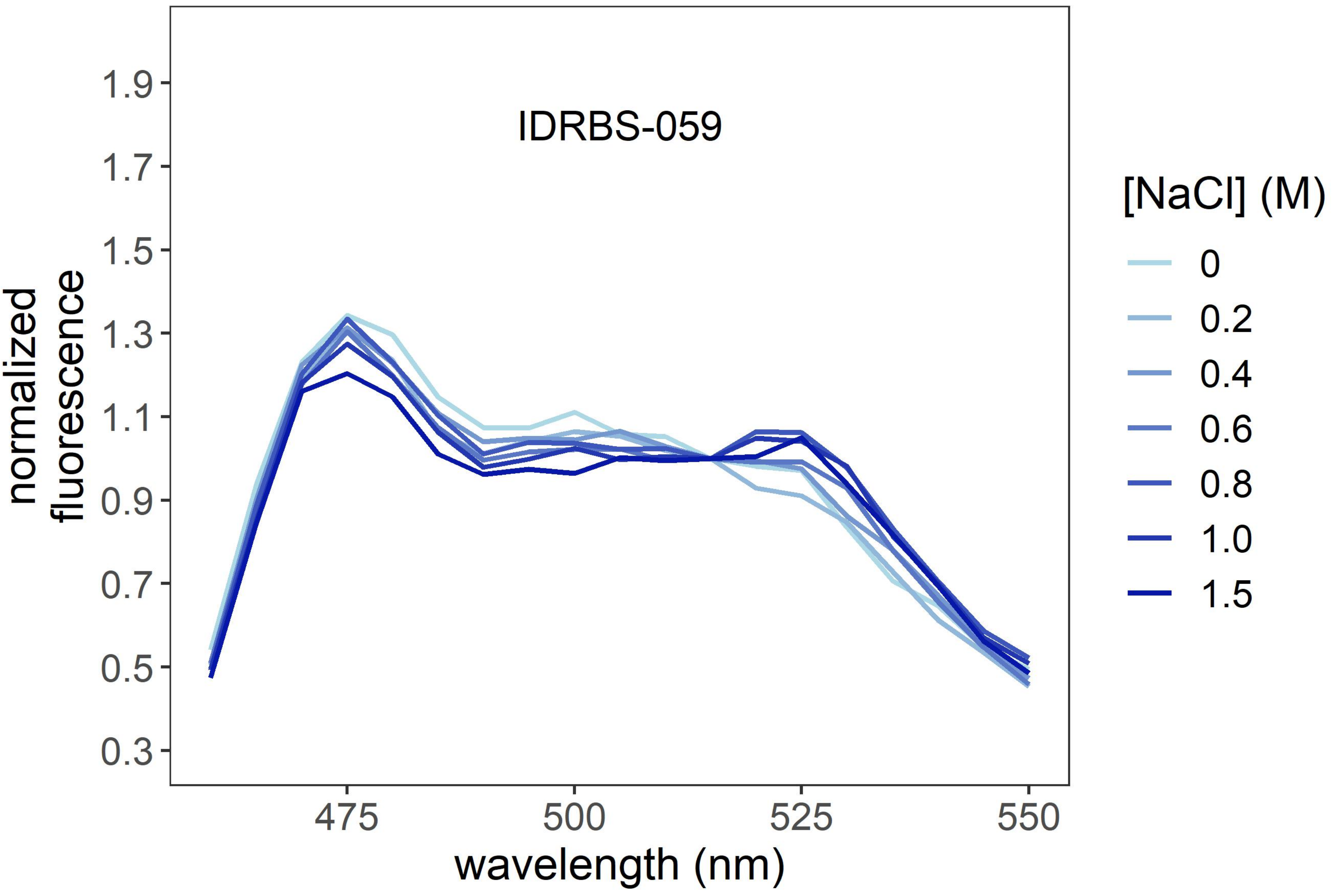
IDRBS-059
normalized fluorescence
1.9
1.7
1.5
1.3
1.1
0.9
0.7
0.5
0.3
475
500
525
550
wavelength (nm)
[NaCl] (M)
0
0.2
0.4
0.6
0.8
1.0
1.5

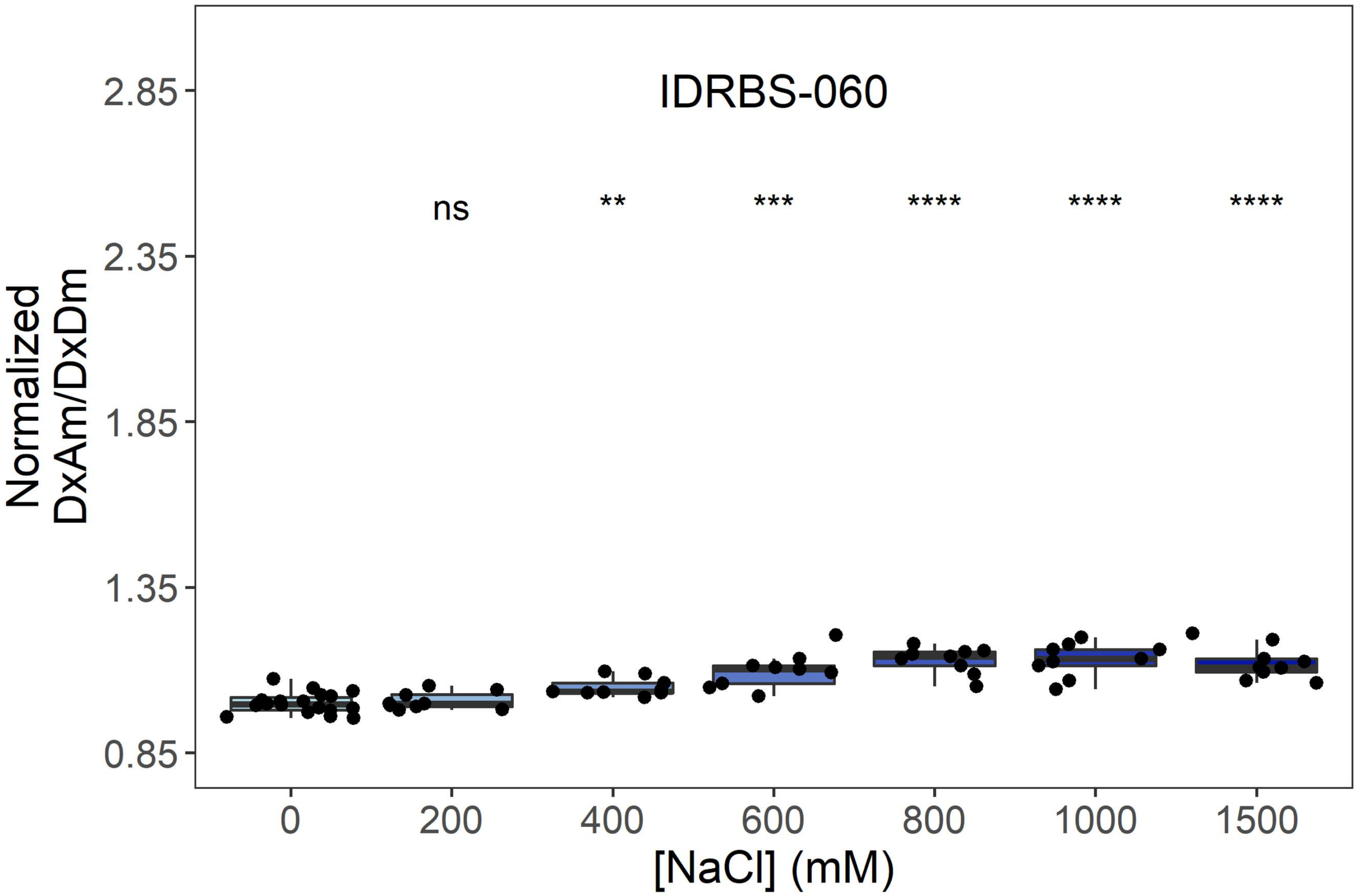
IDRBS-060
ns
**
***
****
****
****
Normalized DxAm/DxDm
2.85
2.35
1.85
1.35
0.85
0
200
400
600
800
1000
1500
[NaCl] (mM)

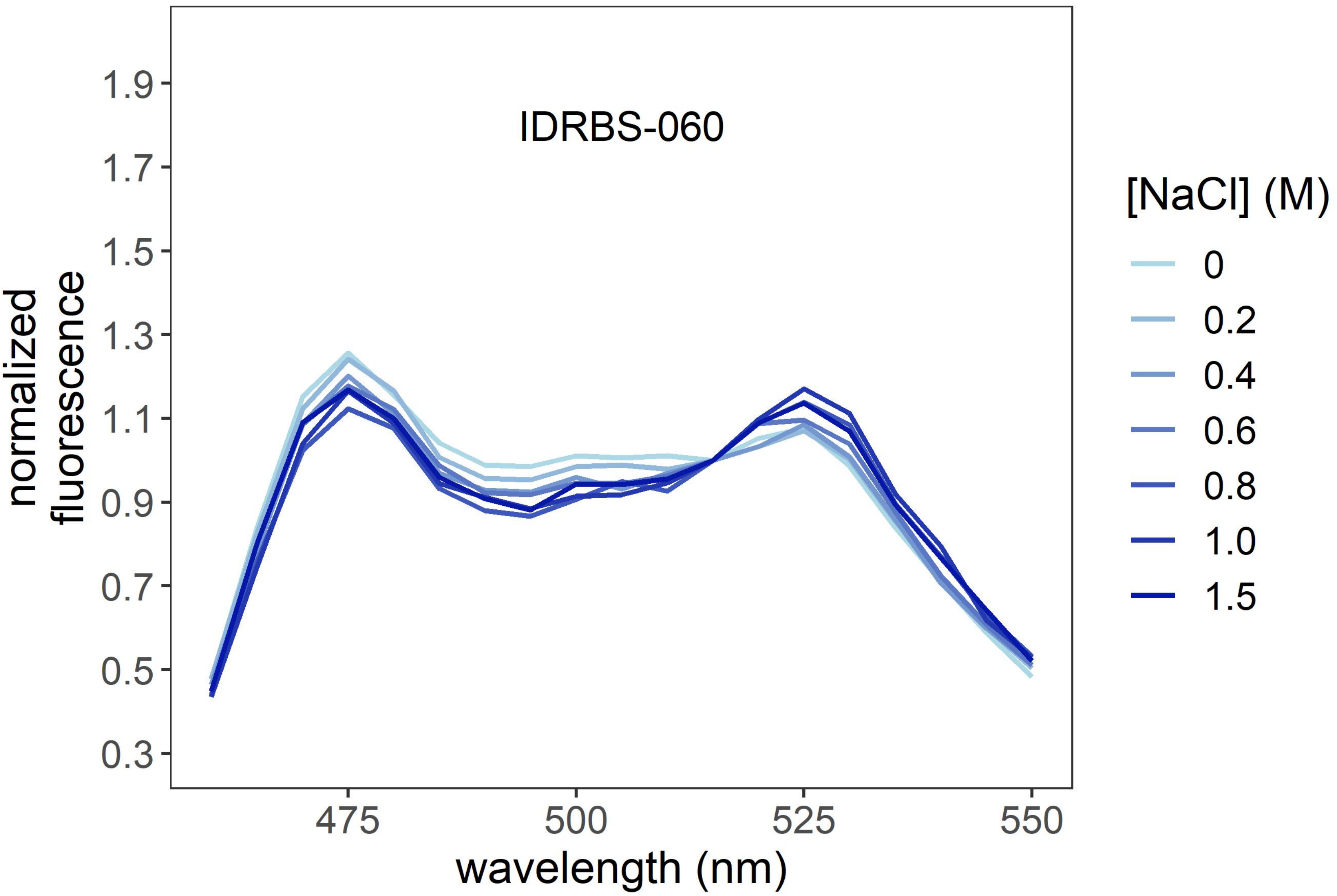
IDRBS-060
normalized fluorescence
1.9
1.7
1.5
1.3
1.1
0.9
0.7
0.5
0.3
475
500
525
550
wavelength (nm)
[NaCl] (M)
0
0.2
0.4
0.6
0.8
1.0
1.5

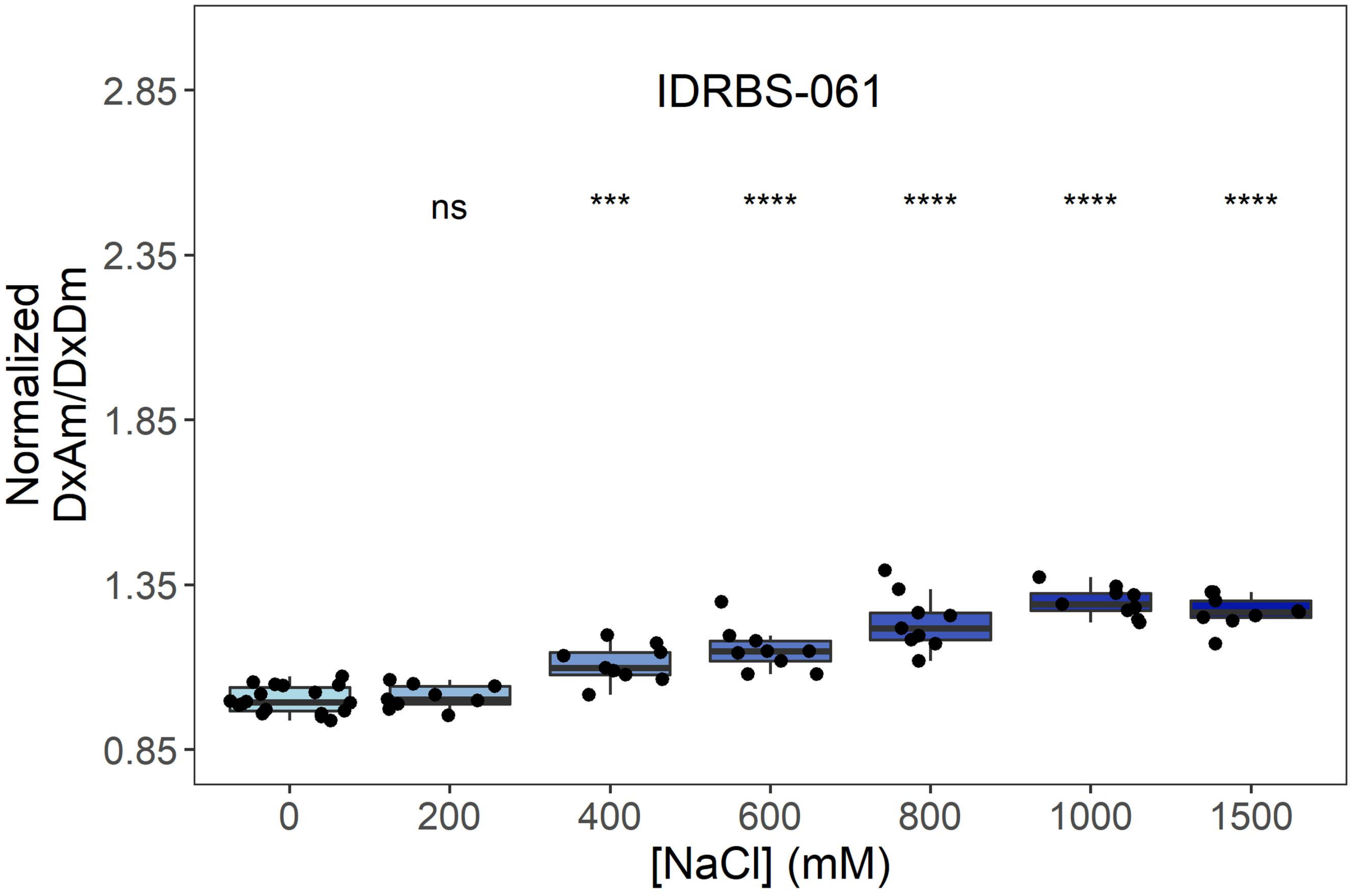
IDRBS-061
ns
***
****
****
****
****
Normalized
DxAm/DxDm
2.85
2.35
1.85
1.35
0.85
0
200
400
600
800
1000
1500
[NaCl] (mM)

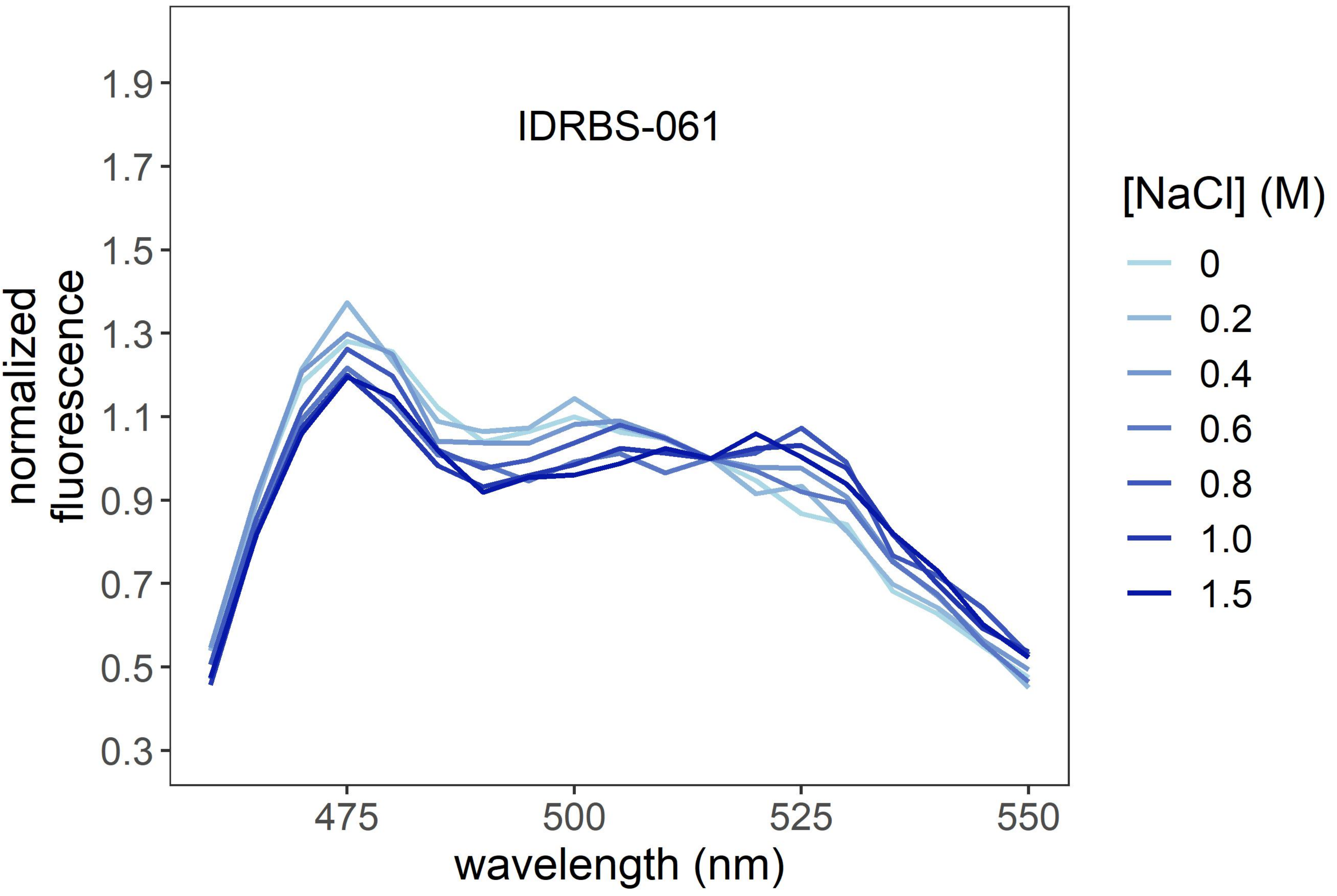
IDRBS-061
normalized
fluorescence
1.9
1.7
1.5
1.3
1.1
0.9
0.7
0.5
0.3
475
500
525
550
wavelength (nm)
[NaCl] (M)
0
0.2
0.4
0.6
0.8
1.0
1.5

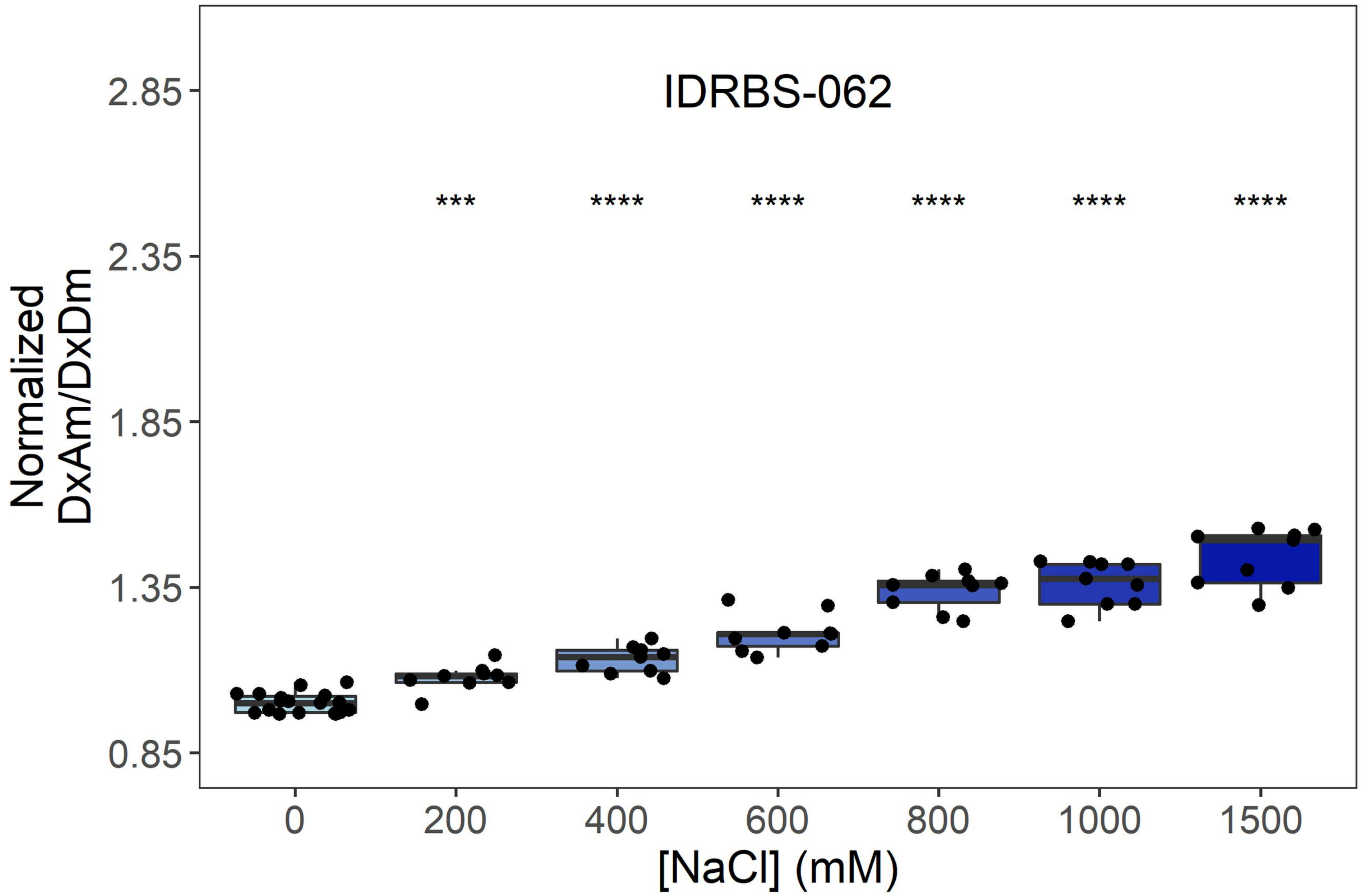
IDRBS-062
Normalized DxAm/DxDm
2.85
2.35
1.85
1.35
0.85
***
****
****
****
****
****
0
200
400
600
800
1000
1500
[NaCl] (mM)

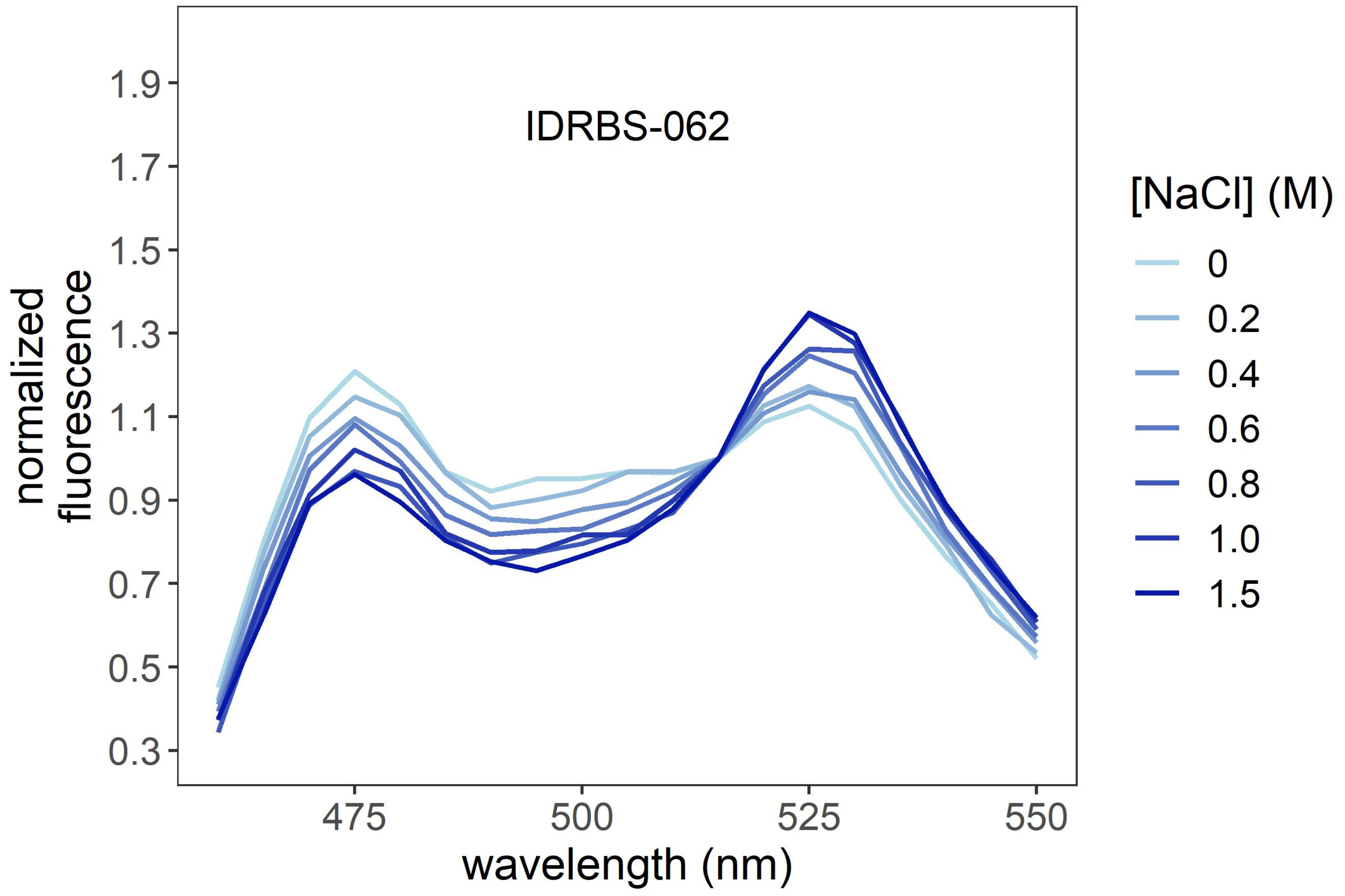
IDRBS-062
normalized fluorescence
1.9
1.7
1.5
1.3
1.1
0.9
0.7
0.5
0.3
475
500
525
550
wavelength (nm)
[NaCl] (M)
0
0.2
0.4
0.6
0.8
1.0
1.5

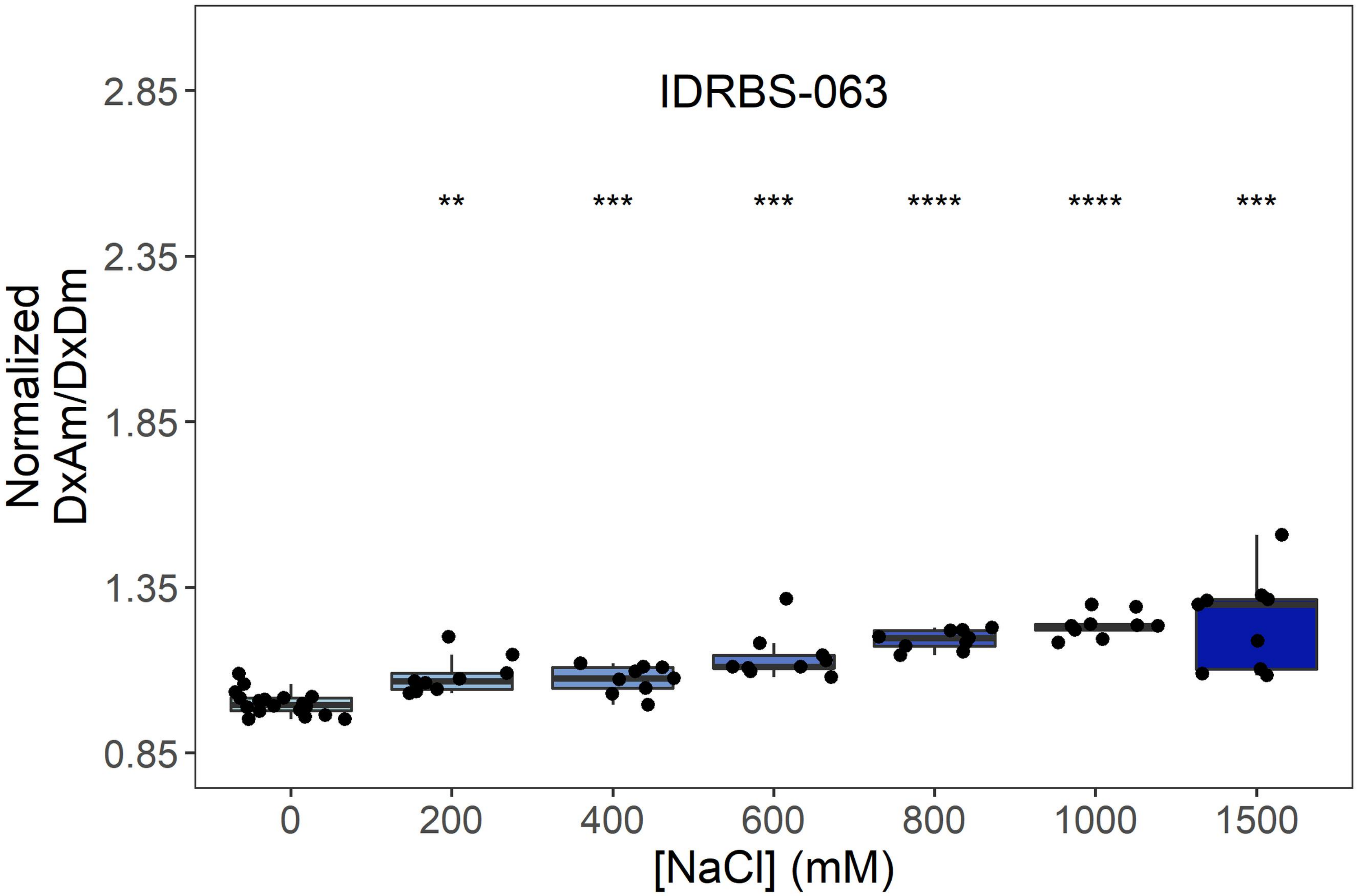
IDRBS-063
**
***
***
****
****
***
Normalized
DxAm/DxDm
2.85
2.35
1.85
1.35
0.85
0
200
400
600
800
1000
1500
[NaCl] (mM)

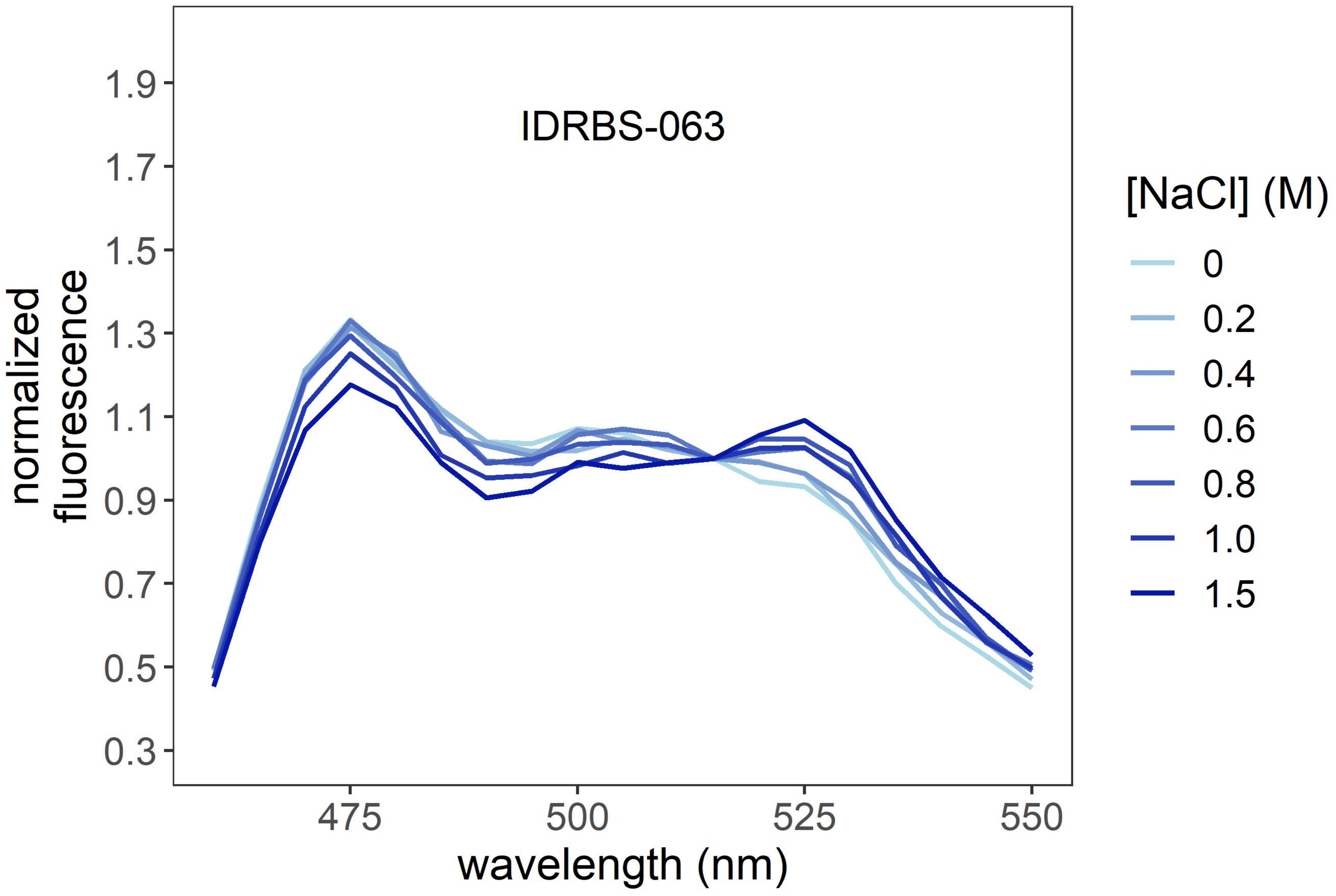
IDRBS-063
normalized
fluorescence
1.9
1.7
1.5
1.3
1.1
0.9
0.7
0.5
0.3
475
500
525
550
wavelength (nm)
[NaCl] (M)
0
0.2
0.4
0.6
0.8
1.0
1.5

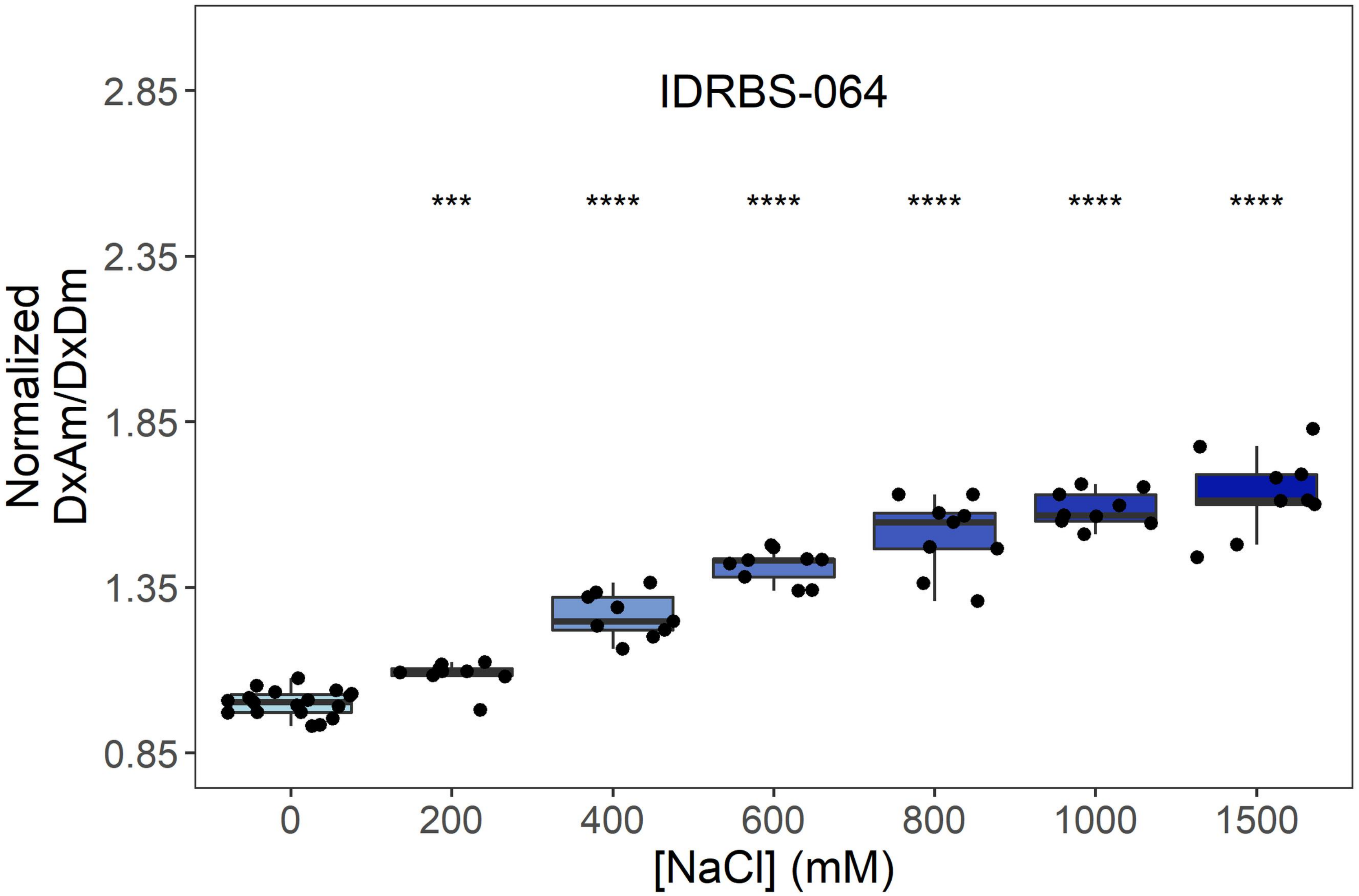
IDRBS-064
***
****
****
****
****
****
Normalized
DxAm/DxDm
2.85
2.35
1.85
1.35
0.85
0
200
400
600
800
1000
1500
[NaCl] (mM)

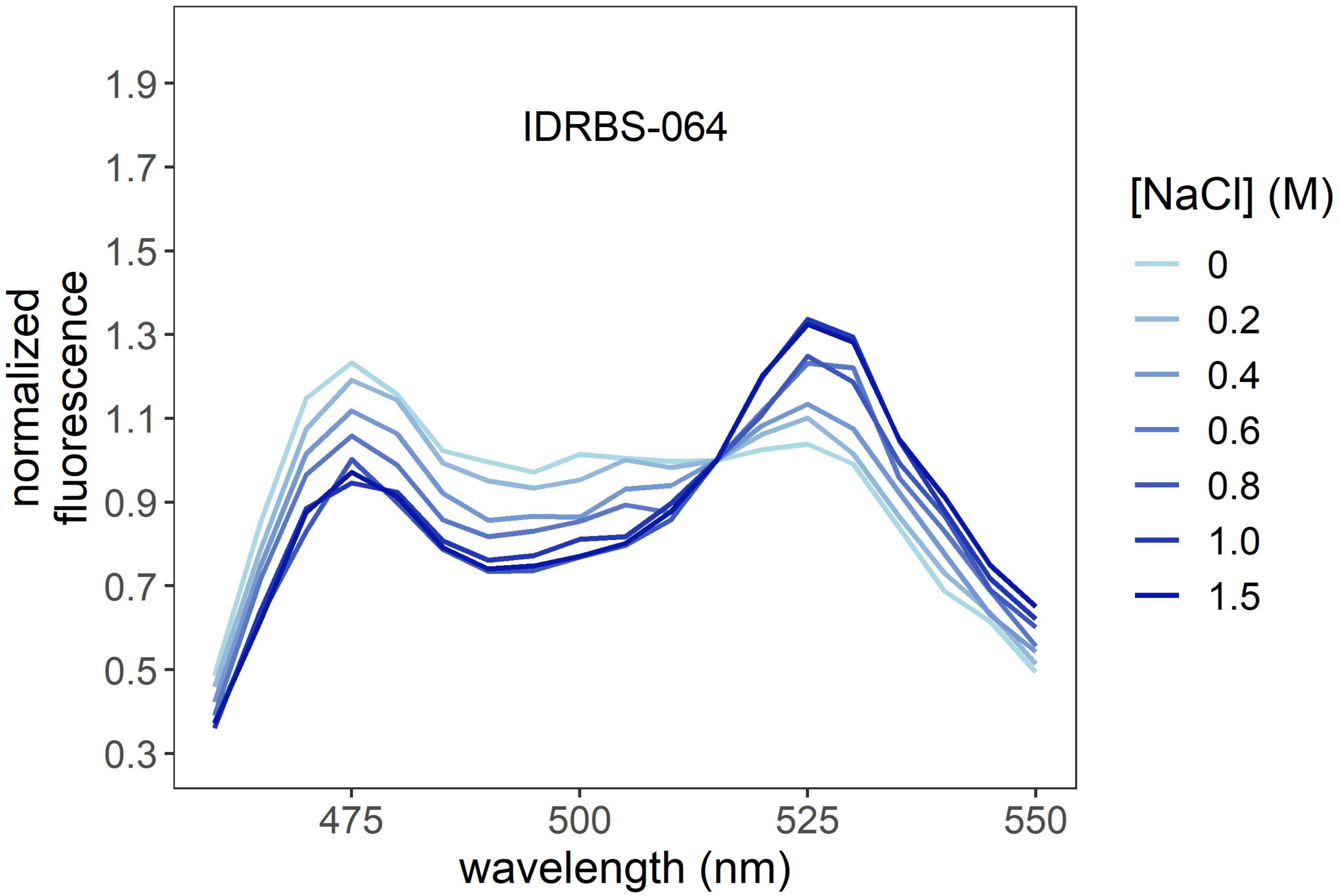
IDRBS-064
normalized
fluorescence
1.9
1.7
1.5
1.3
1.1
0.9
0.7
0.5
0.3
475
500
525
550
wavelength (nm)
[NaCl] (M)
0
0.2
0.4
0.6
0.8
1.0
1.5

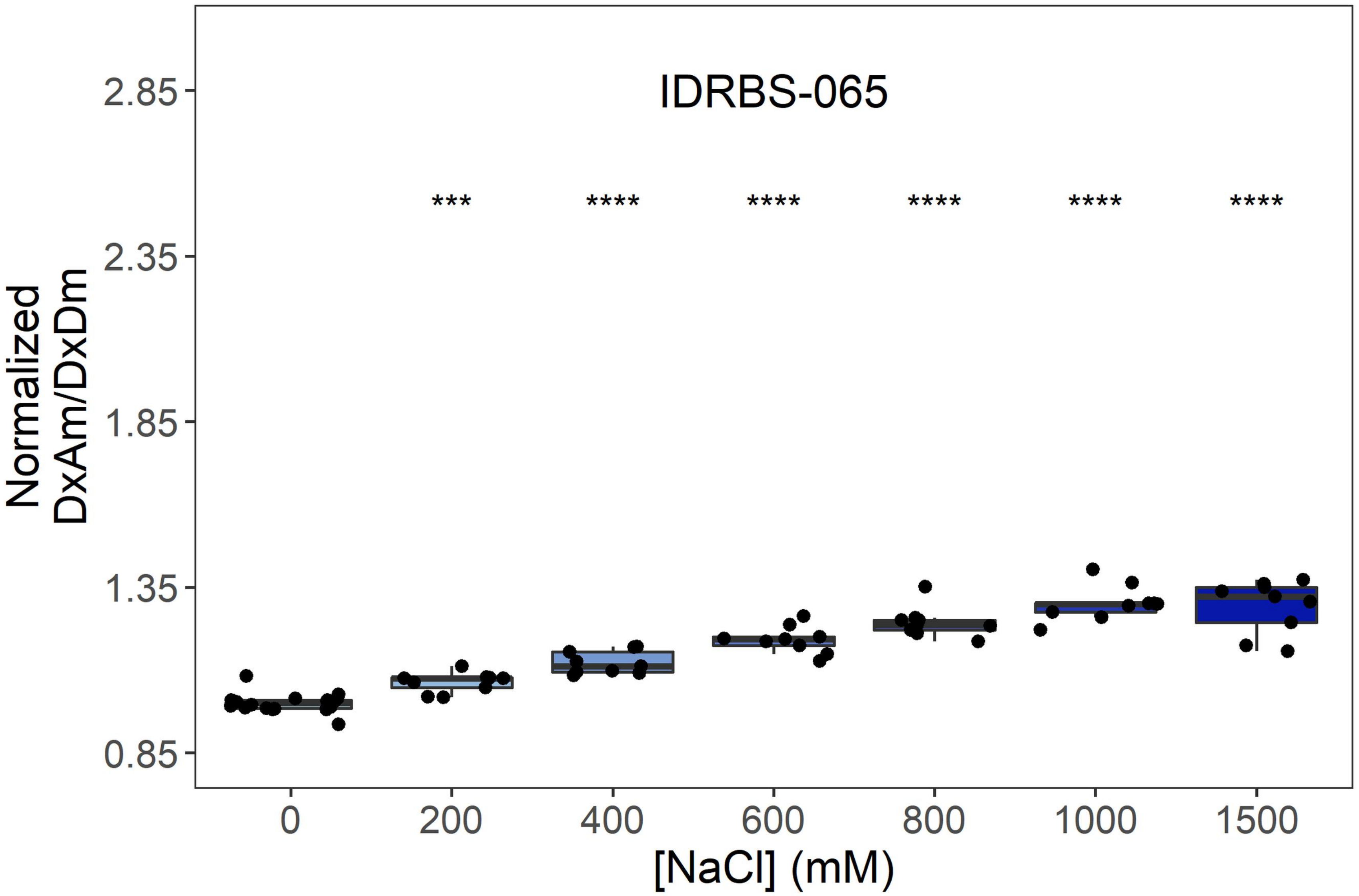
IDRBS-065
Normalized DxAm/DxDm
2.85
2.35
1.85
1.35
0.85
***
****
****
****
****
****
0
200
400
600
800
1000
1500
[NaCl] (mM)

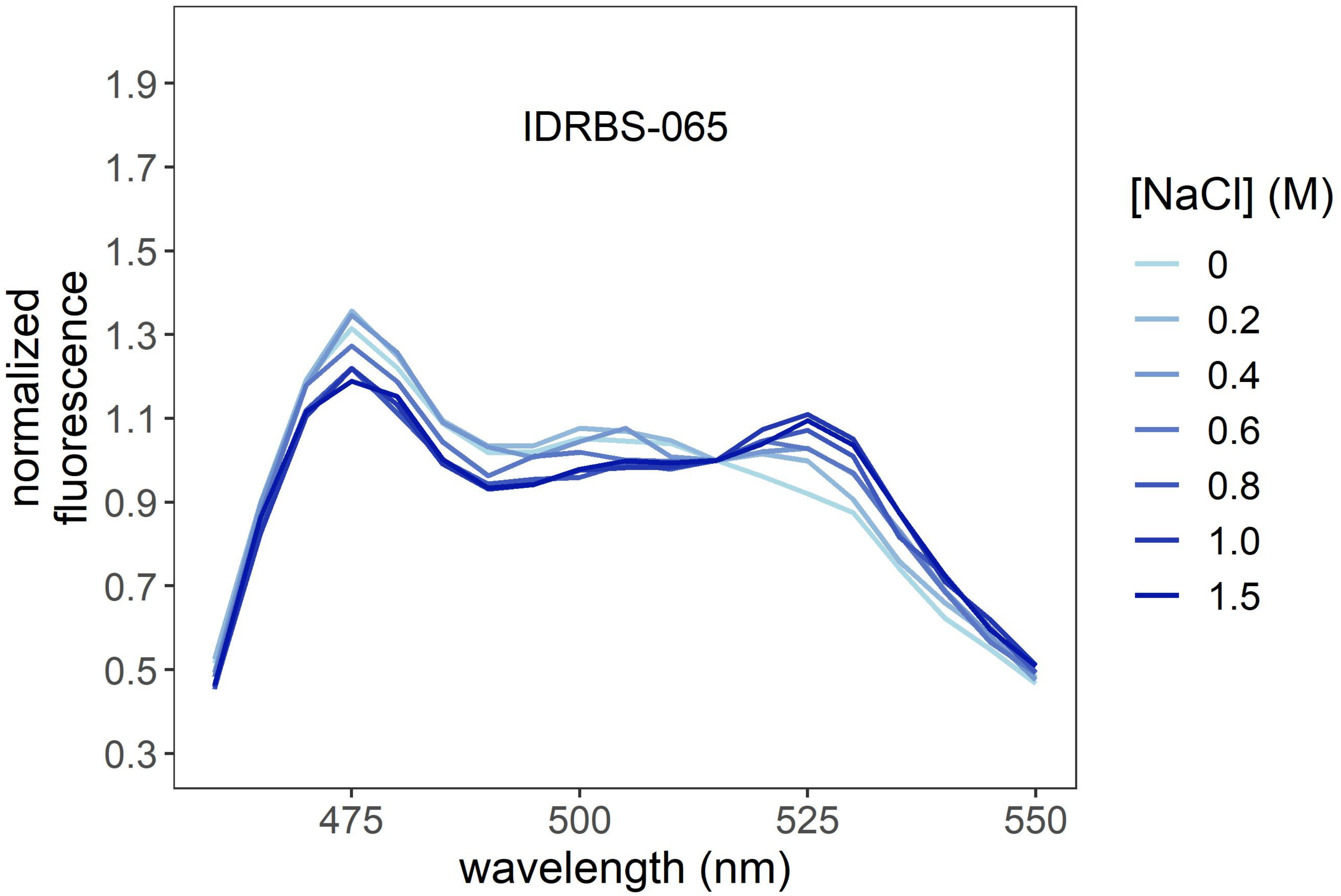
IDRBS-065
normalized fluorescence
1.9
1.7
1.5
1.3
1.1
0.9
0.7
0.5
0.3
475
500
525
550
wavelength (nm)
[NaCl] (M)
0
0.2
0.4
0.6
0.8
1.0
1.5

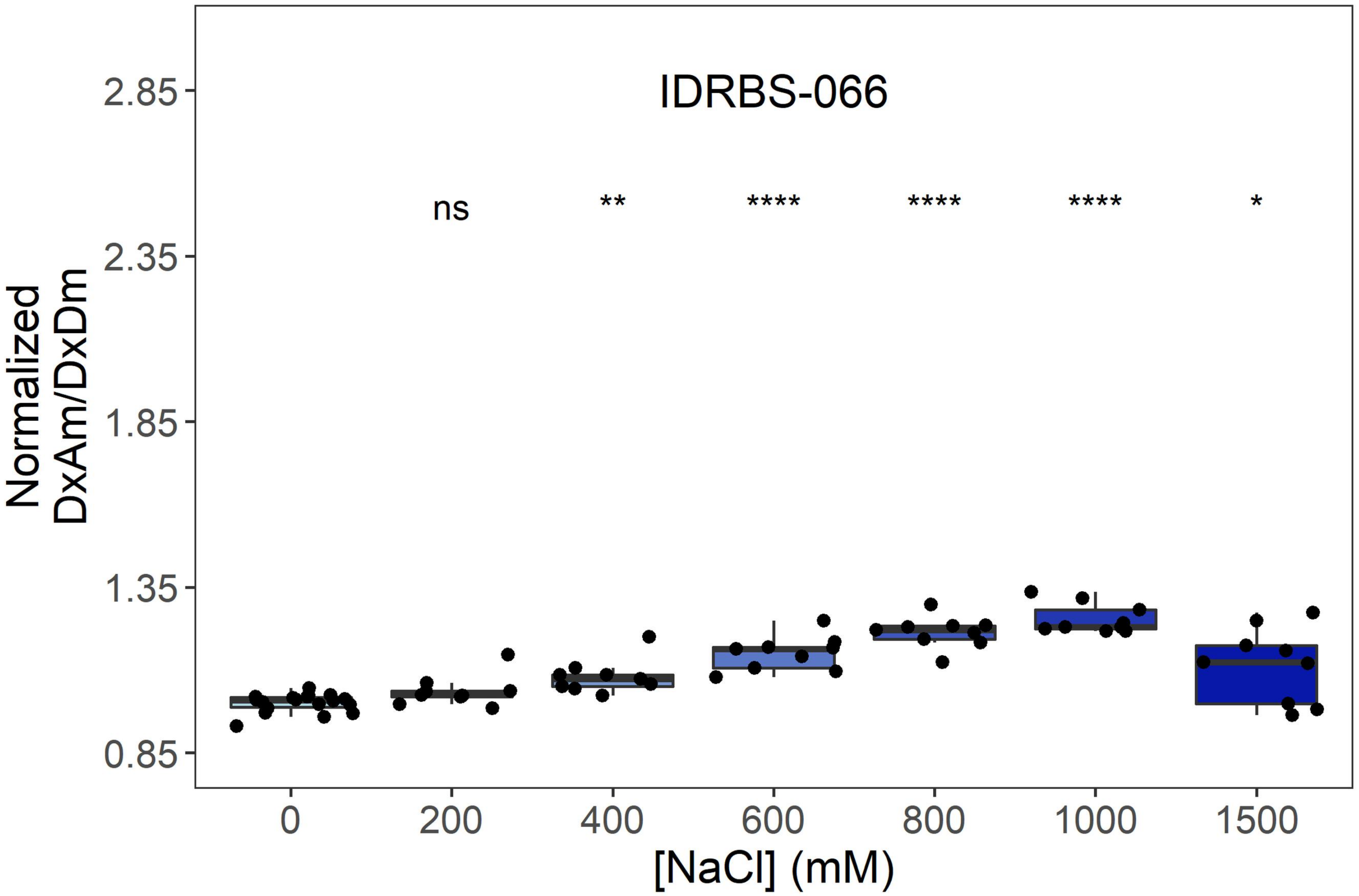
IDRBS-066
ns
**
****
****
****
*
2.85
2.35
1.85
1.35
0.85
Normalized
DxAm/DxDm
0
200
400
600
800
1000
1500
[NaCl] (mM)

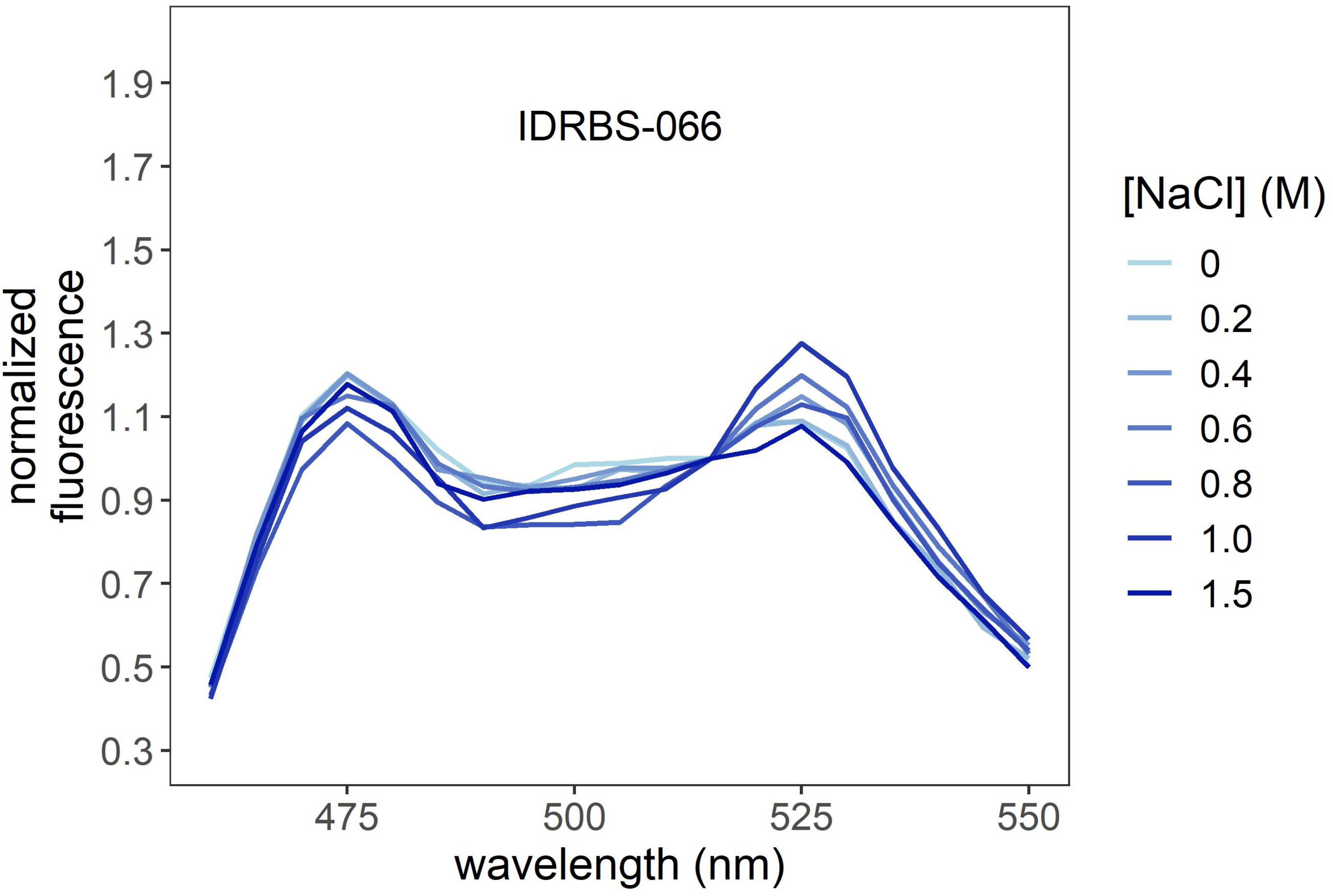
IDRBS-066
1.9
1.7
1.5
1.3
1.1
0.9
0.7
0.5
0.3
normalized
fluorescence
475
500
525
550
wavelength (nm)
[NaCl] (M)
0
0.2
0.4
0.6
0.8
1.0
1.5

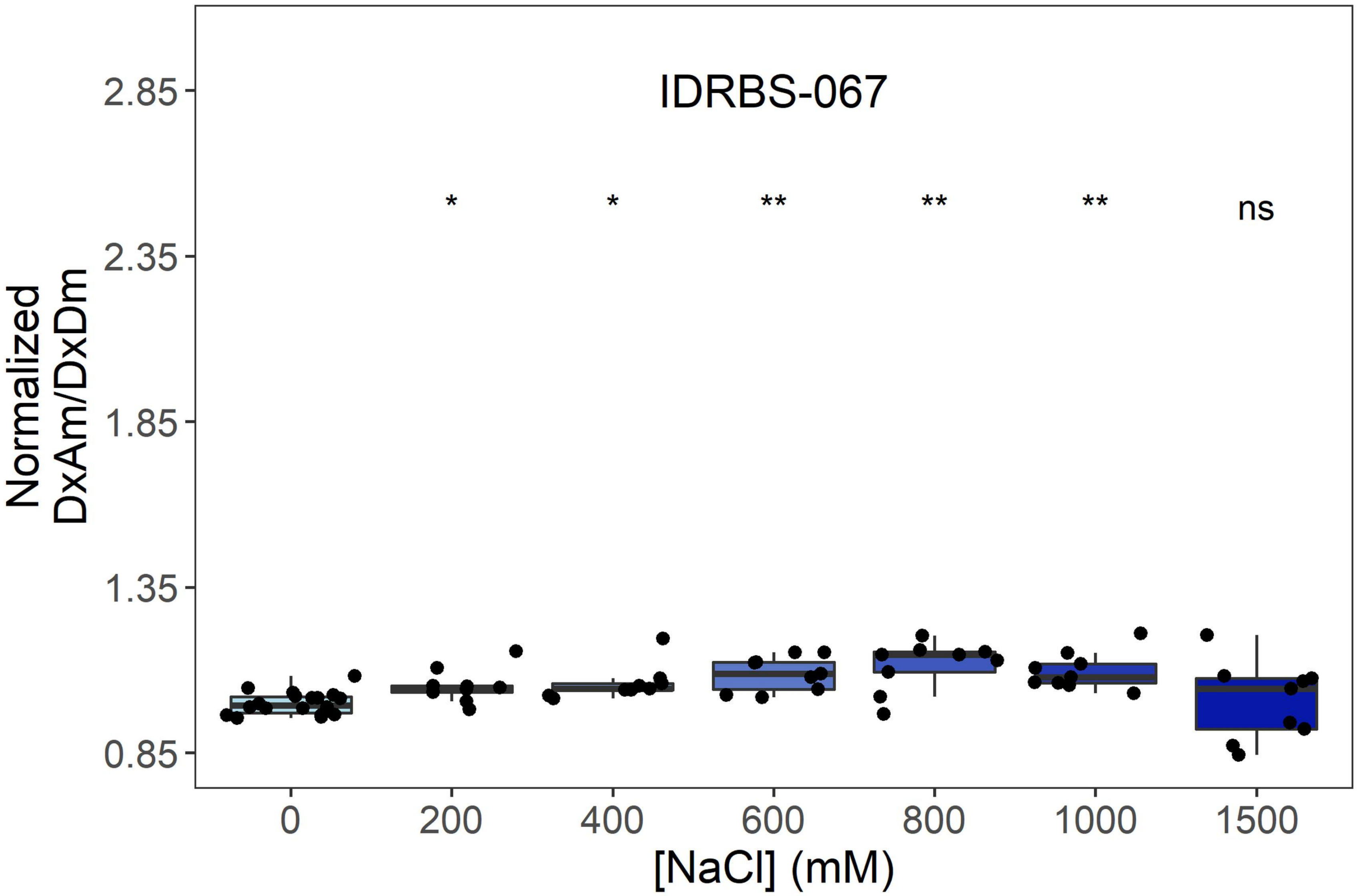
IDRBS-067
Normalized DxAm/DxDm
2.85
2.35
1.85
1.35
0.85
*
*
**
**
**
ns
0
200
400
600
800
1000
1500
[NaCl] (mM)

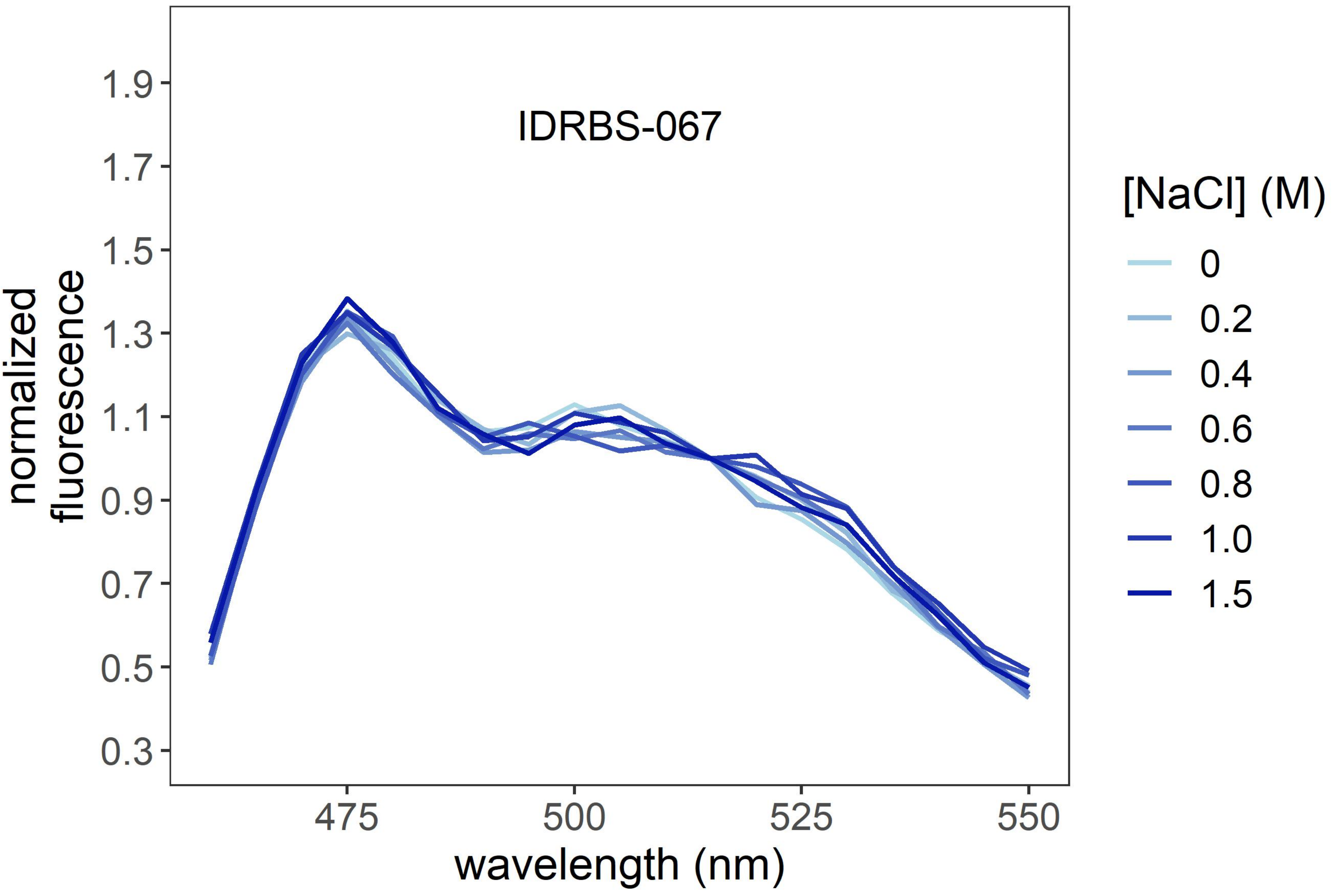
IDRBS-067
normalized fluorescence
1.9
1.7
1.5
1.3
1.1
0.9
0.7
0.5
0.3
475
500
525
550
wavelength (nm)
[NaCl] (M)
0
0.2
0.4
0.6
0.8
1.0
1.5

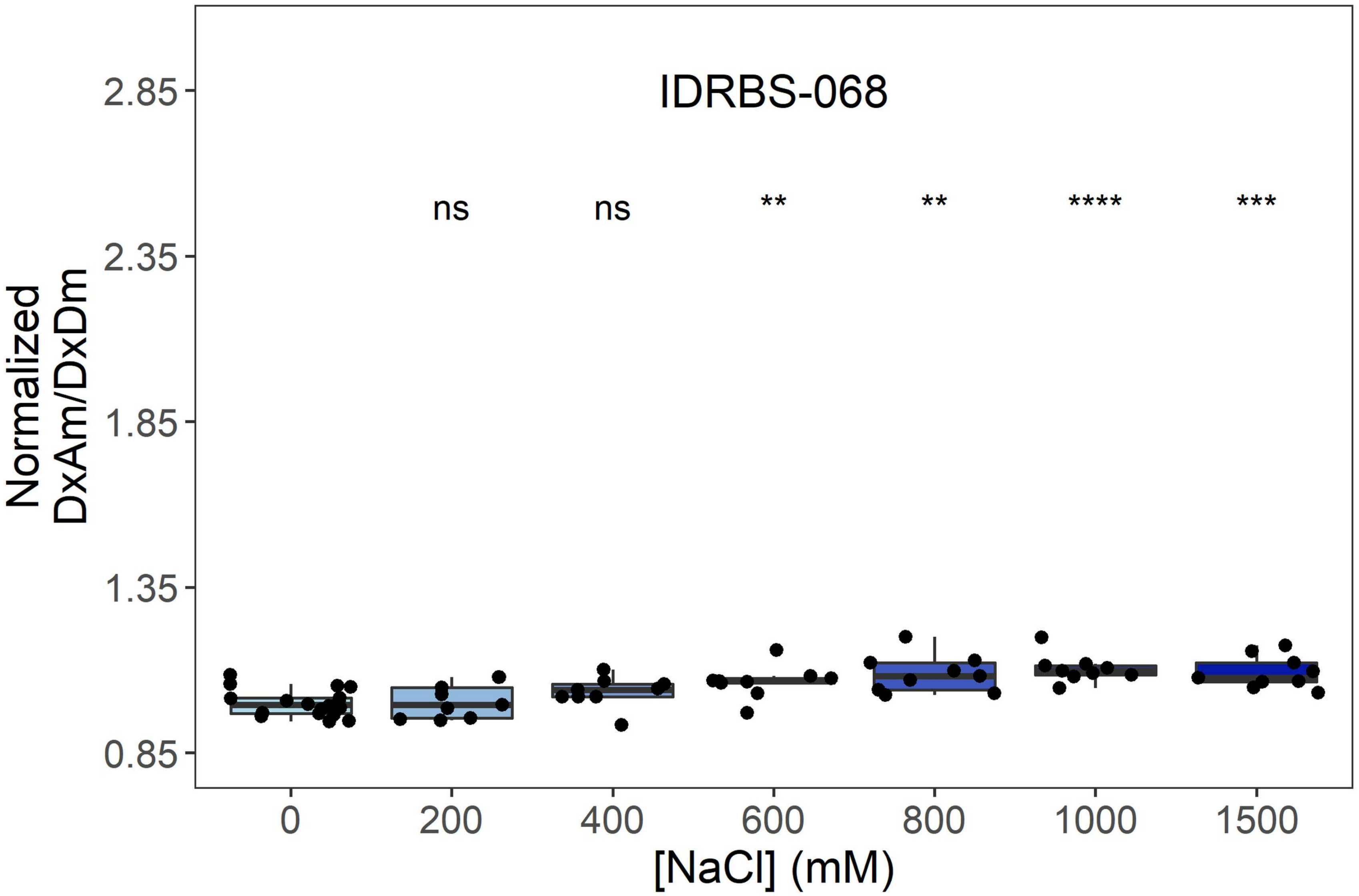
IDRBS-068
ns
ns
**
**
****
***
2.85
2.35
1.85
1.35
0.85
Normalized DxAm/DxDm
0
200
400
600
800
1000
1500
[NaCl] (mM)

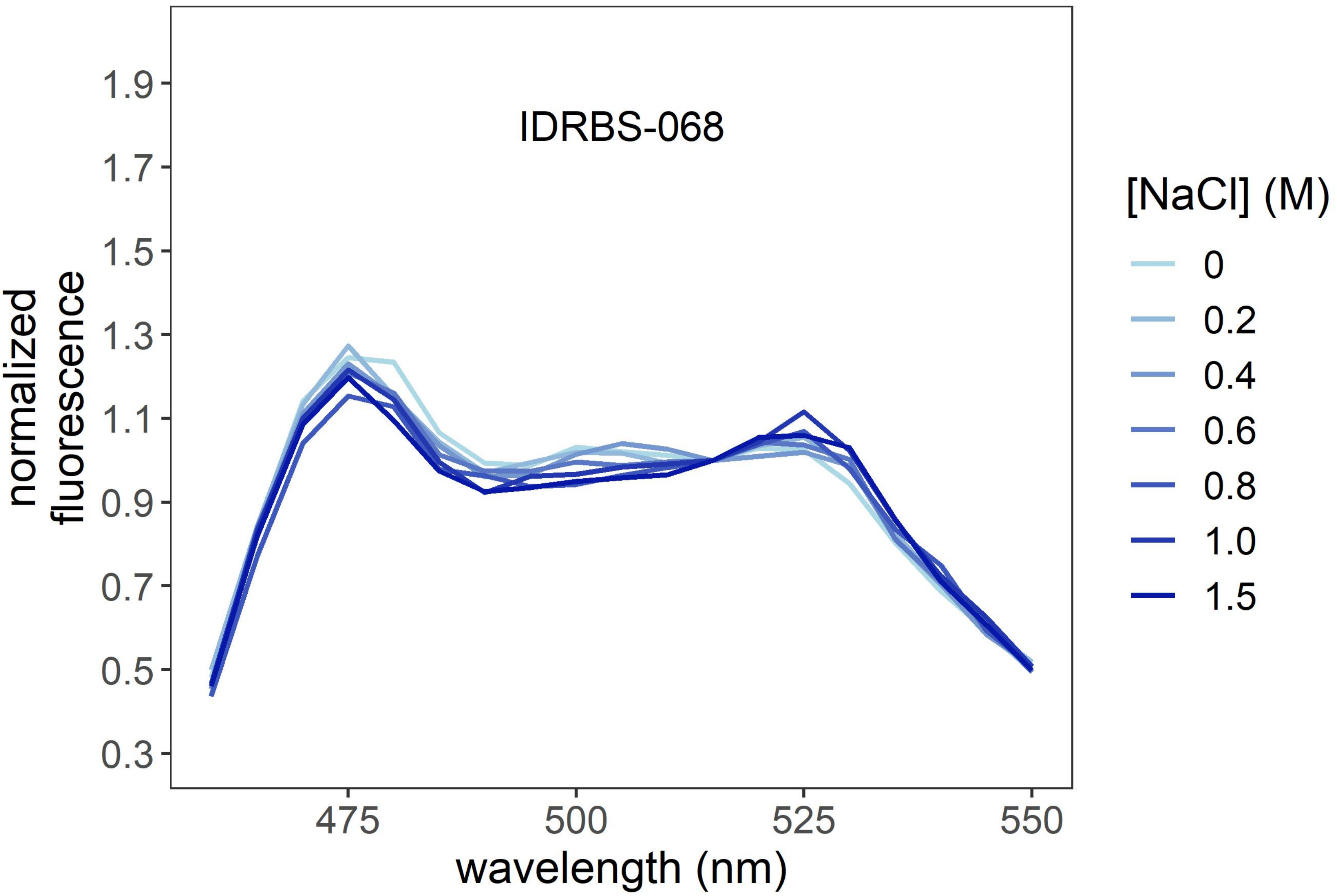
IDRBS-068
1.9
1.7
1.5
1.3
1.1
0.9
0.7
0.5
0.3
normalized fluorescence
475
500
525
550
wavelength (nm)
[NaCl] (M)
0
0.2
0.4
0.6
0.8
1.0
1.5

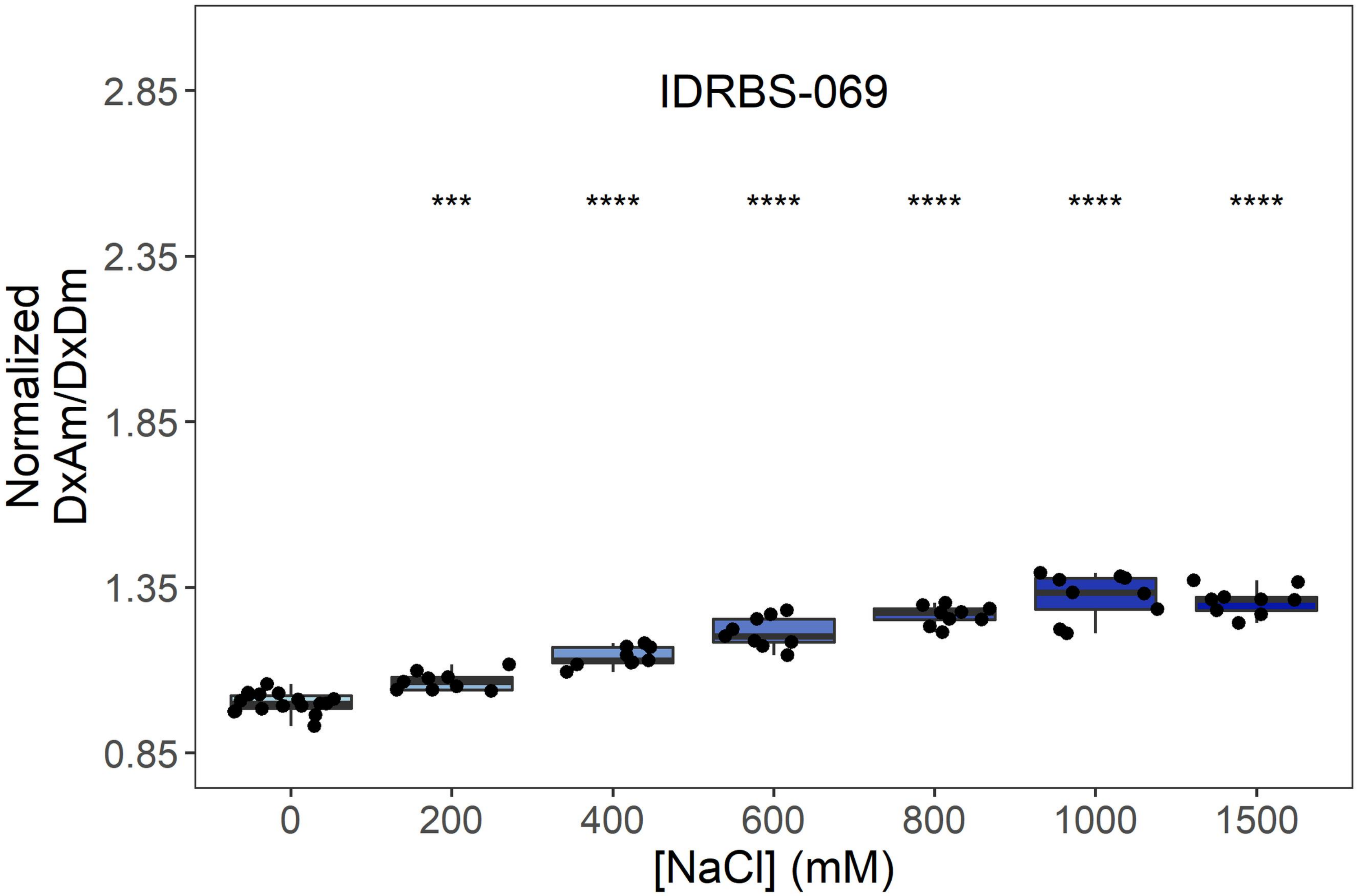

IDRBS-069
***
****
****
****
****
****
2.85
2.35
1.85
1.35
0.85
Normalized
DxAm/DxDm
0
200
400
600
800
1000
1500
[NaCl] (mM)


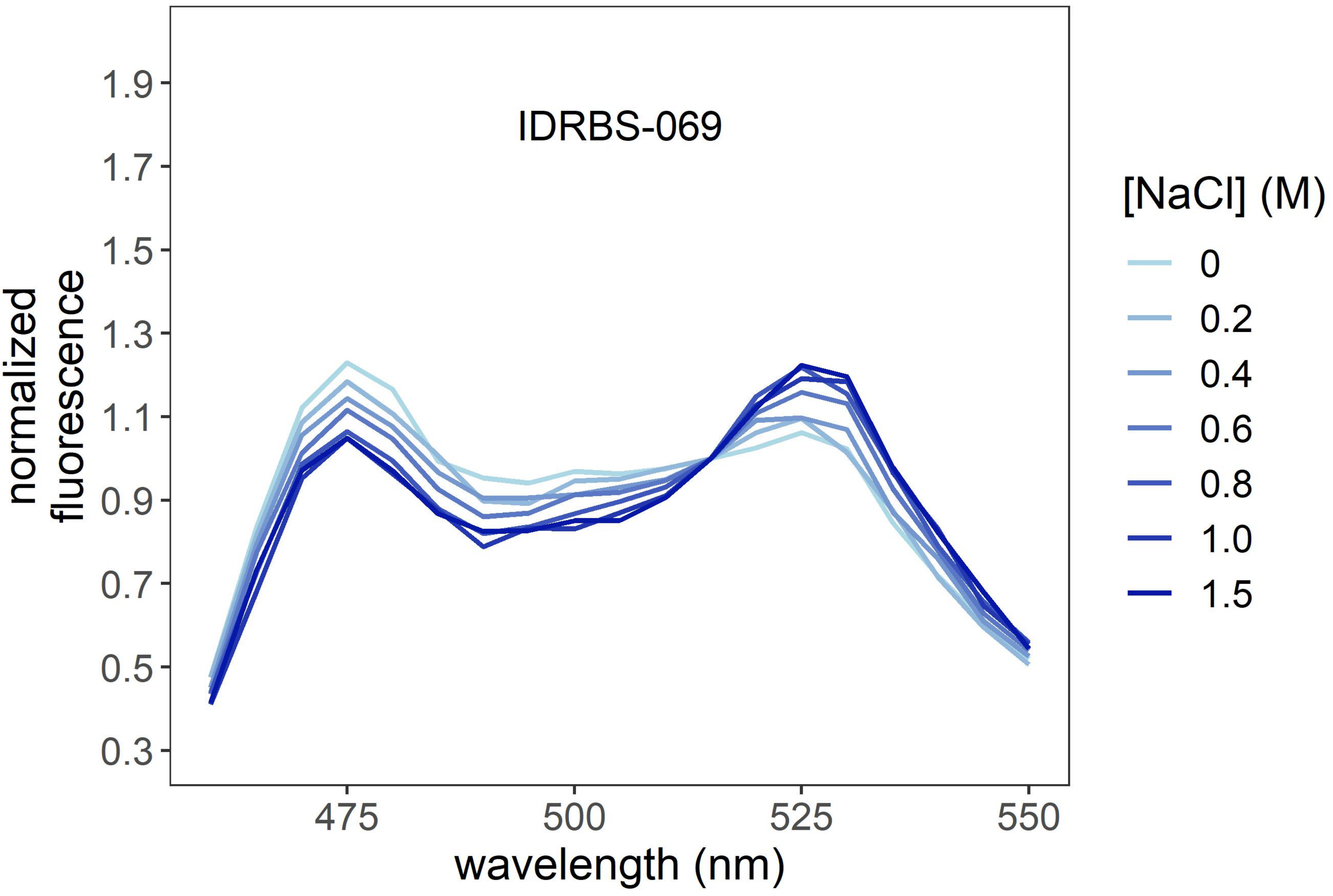

IDRBS-069
1.9
1.7
1.5
1.3
1.1
0.9
0.7
0.5
0.3
normalized
fluorescence
475
500
525
550
wavelength (nm)
[NaCl] (M)
0
0.2
0.4
0.6
0.8
1.0
1.5

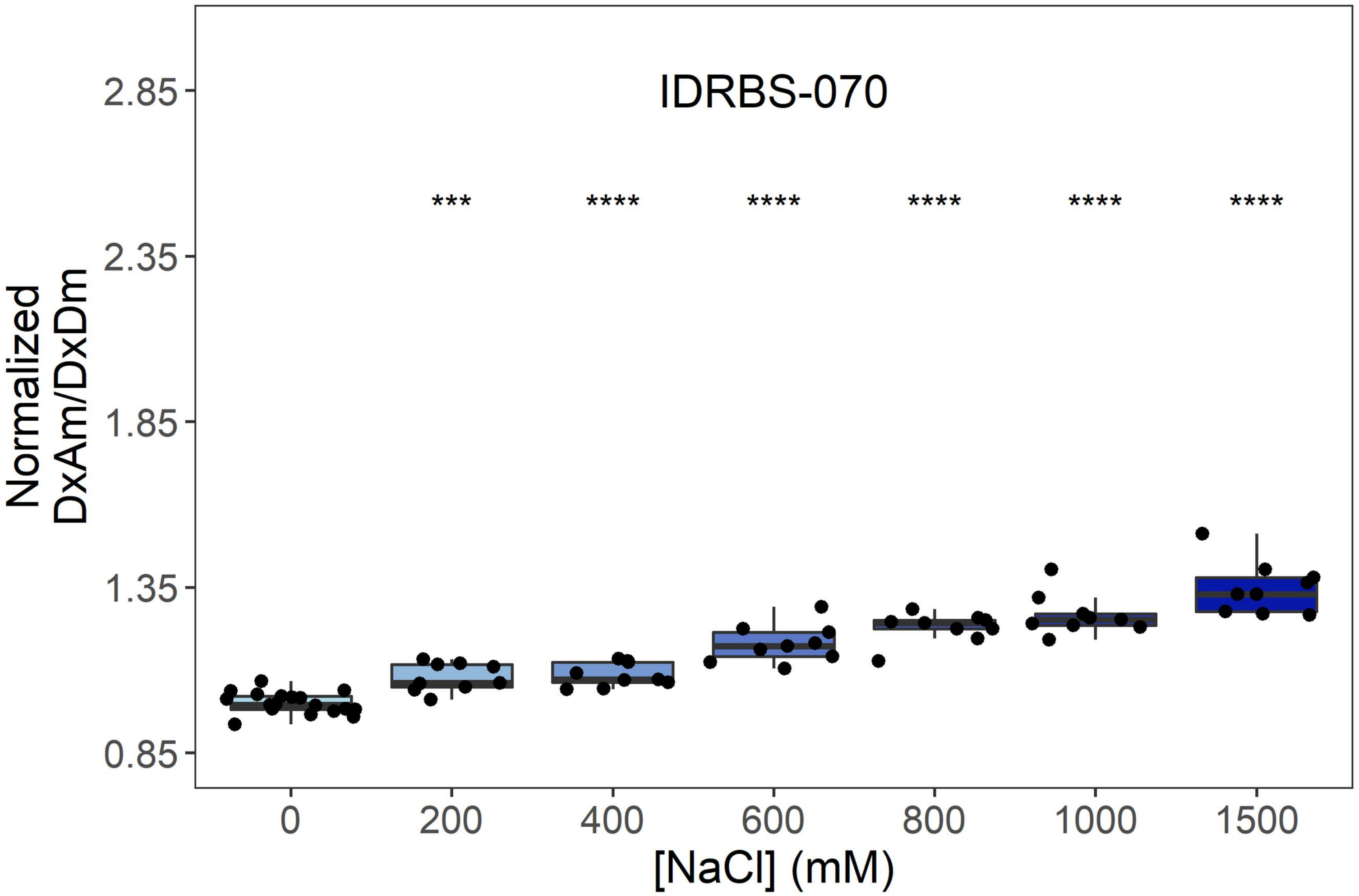

IDRBS-070
***
****
****
****
****
****
Normalized DxAm/DxDm
2.85
2.35
1.85
1.35
0.85
0
200
400
600
800
1000
1500
[NaCl] (mM)


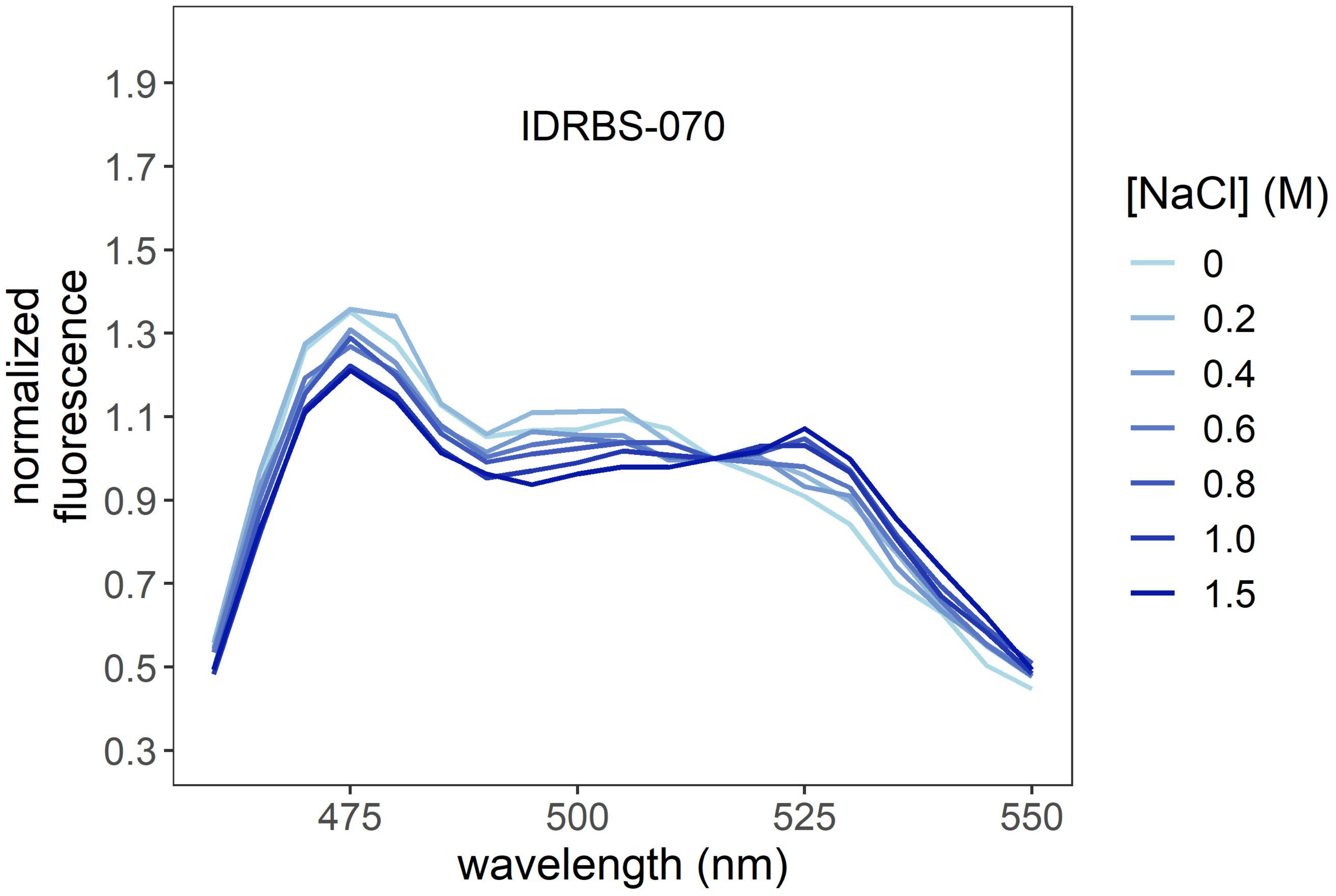

IDRBS-070
normalized fluorescence
1.9
1.7
1.5
1.3
1.1
0.9
0.7
0.5
0.3
475
500
525
550
wavelength (nm)
[NaCl] (M)
0
0.2
0.4
0.6
0.8
1.0
1.5

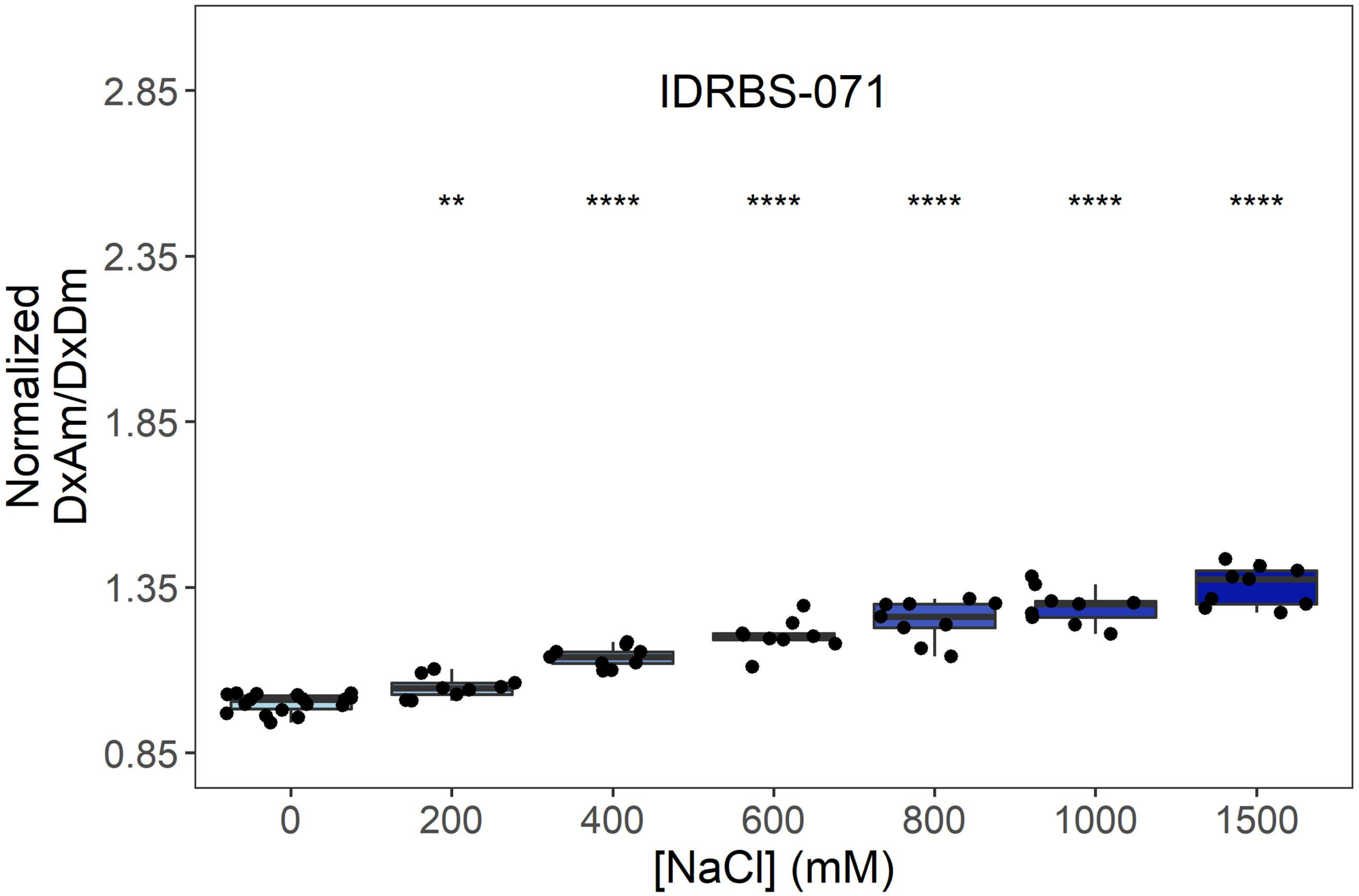
IDRBS-071
Normalized
DxAm/DxDm
2.85
2.35
1.85
1.35
0.85
**
****
****
****
****
****
0
200
400
600
800
1000
1500
[NaCl] (mM)

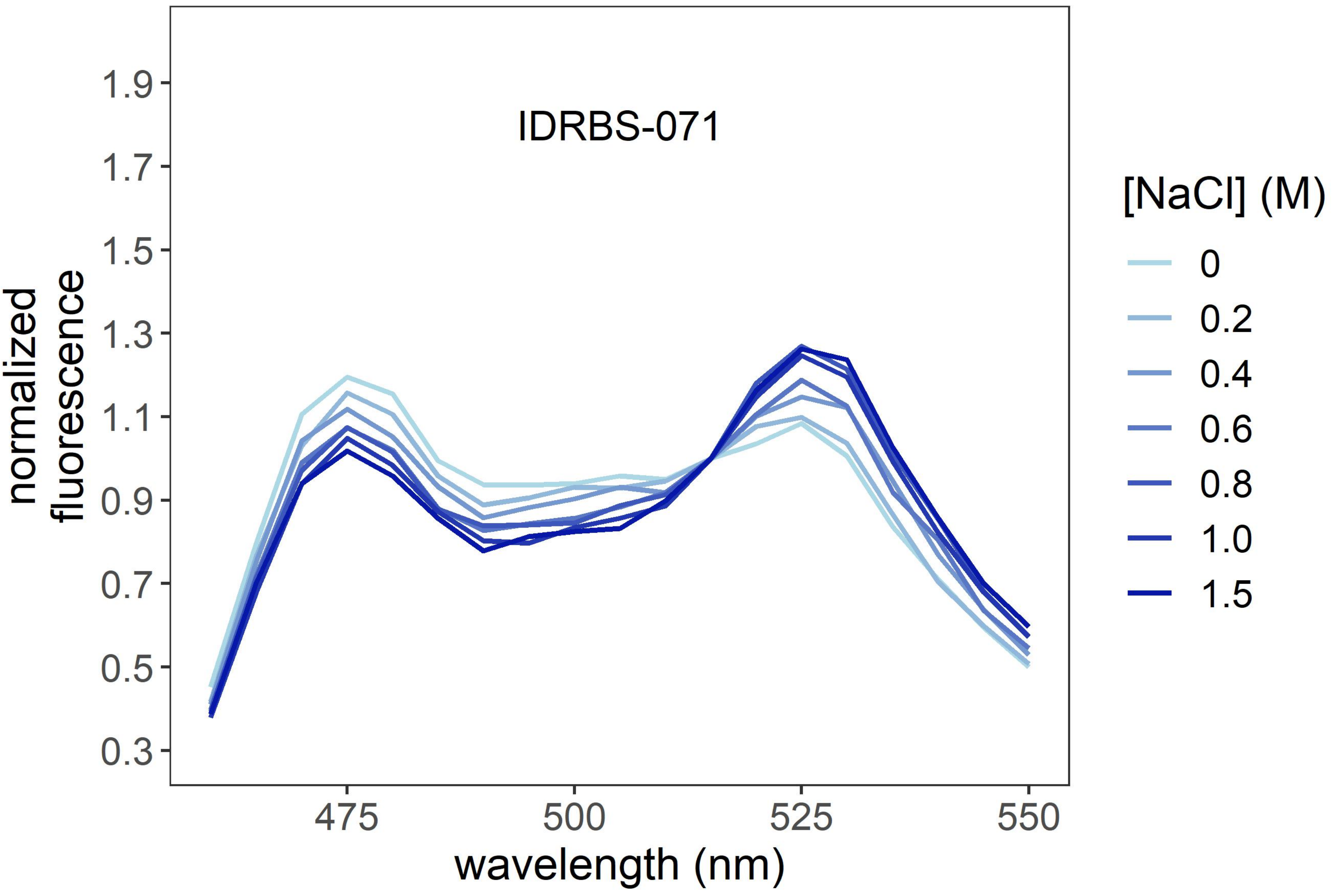
IDRBS-071
normalized
fluorescence
1.9
1.7
1.5
1.3
1.1
0.9
0.7
0.5
0.3
475
500
525
550
wavelength (nm)
[NaCl] (M)
0
0.2
0.4
0.6
0.8
1.0
1.5

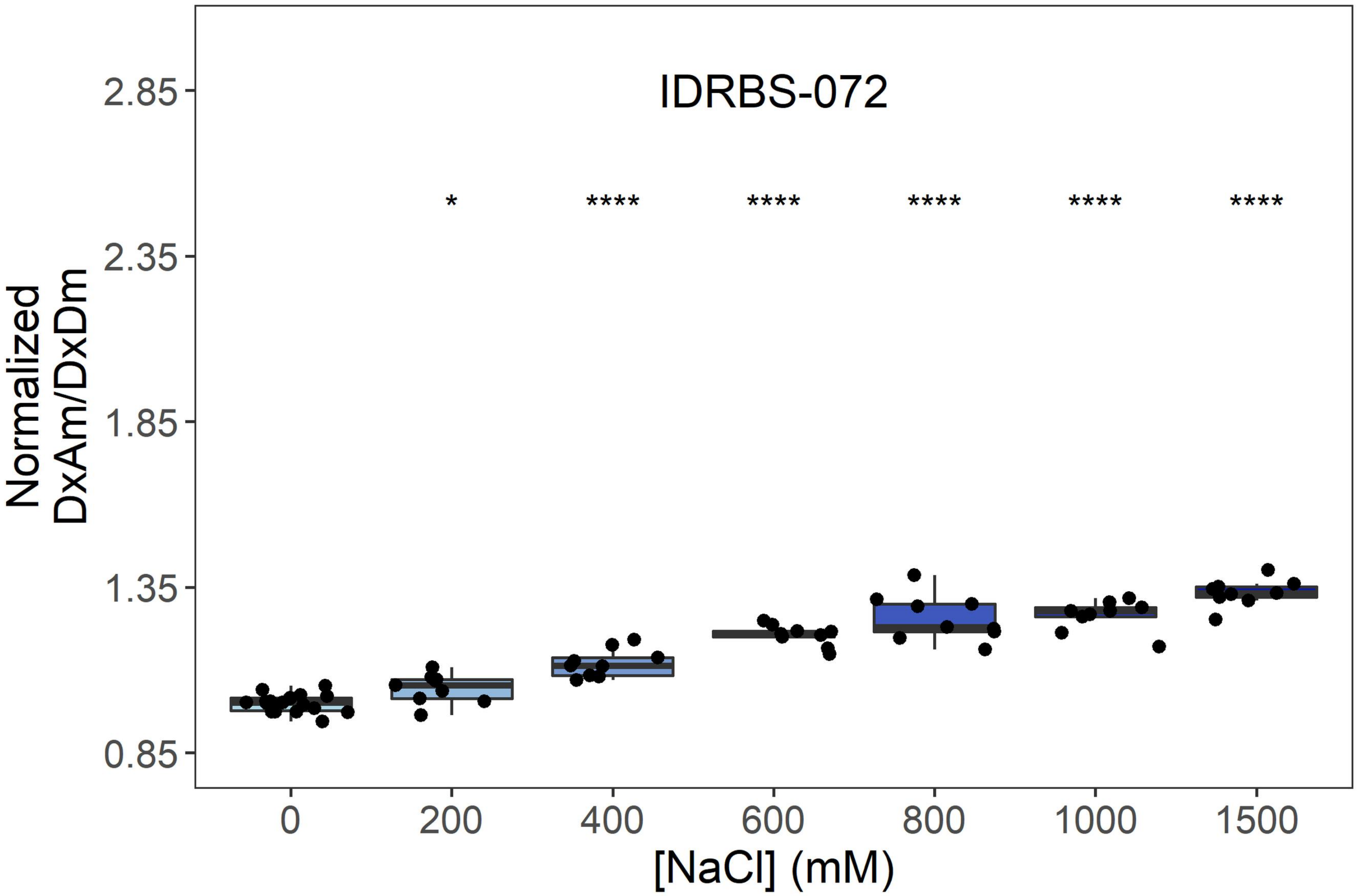
IDRBS-072
Normalized
DxAm/DxDm
2.85
2.35
1.85
1.35
0.85
*
****
****
****
****
****
0
200
400
600
800
1000
1500
[NaCl] (mM)

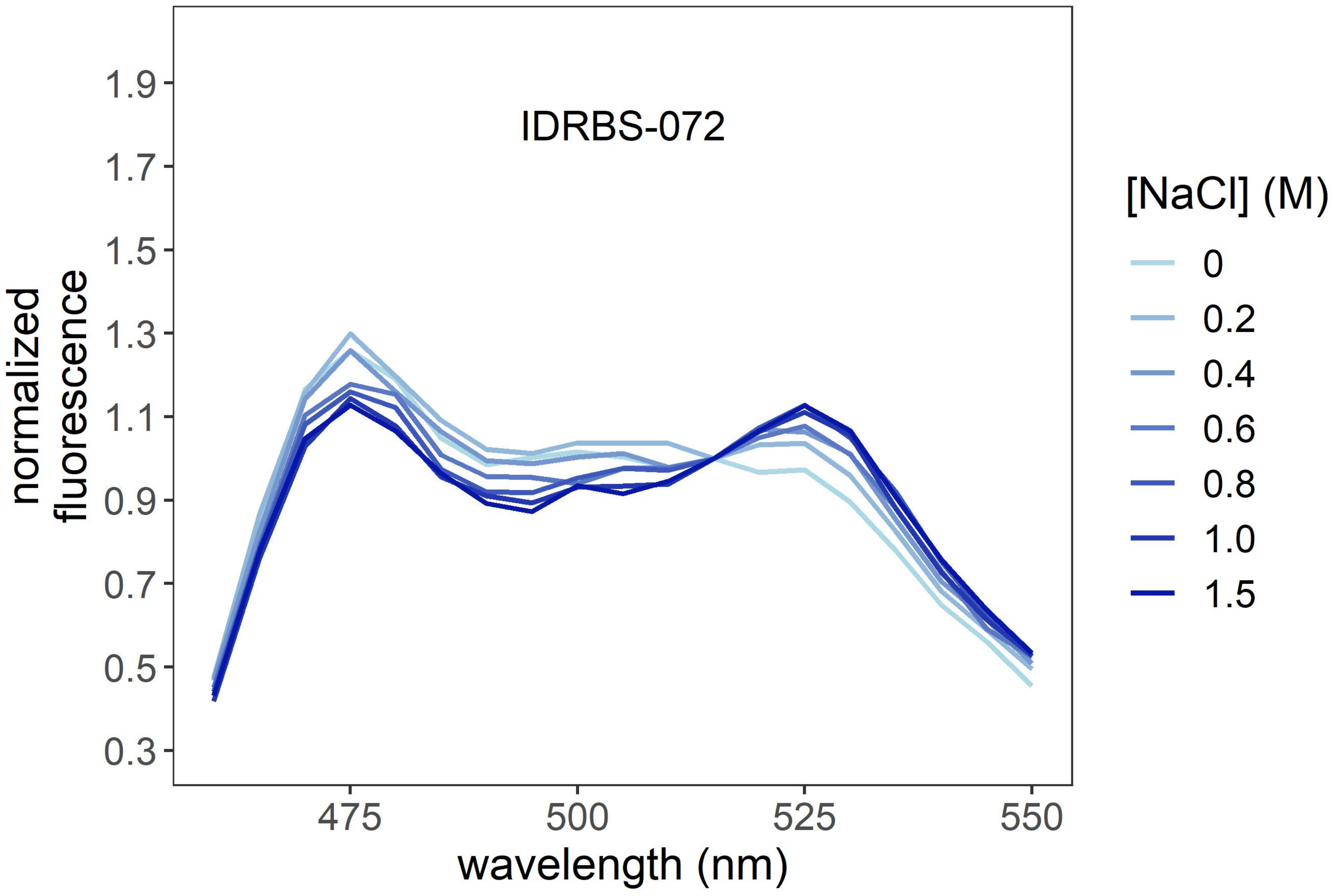
IDRBS-072
normalized
fluorescence
1.9
1.7
1.5
1.3
1.1
0.9
0.7
0.5
0.3
475
500
525
550
wavelength (nm)
[NaCl] (M)
0
0.2
0.4
0.6
0.8
1.0
1.5

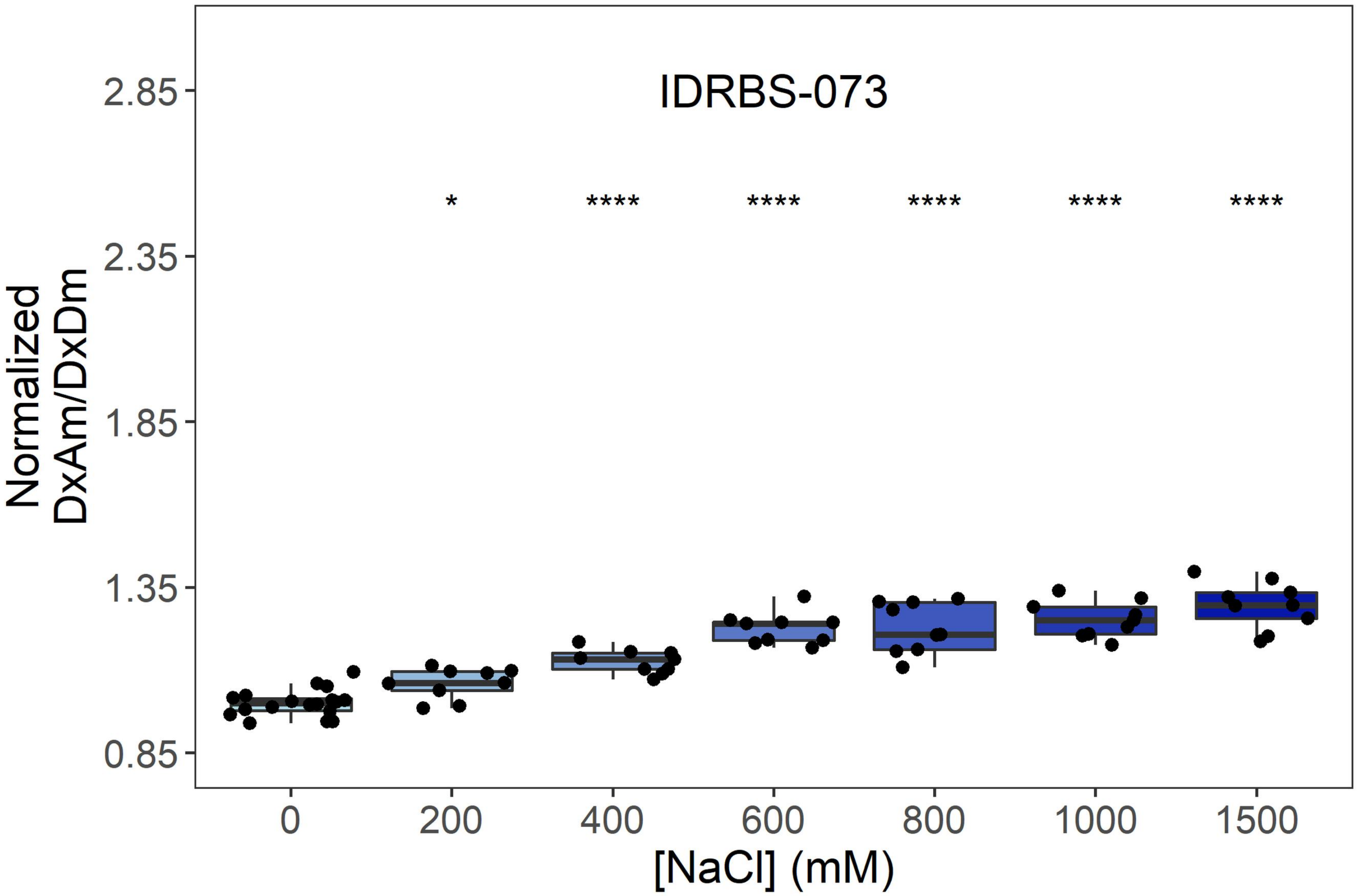
IDRBS-073
Normalized DxAm/DxDm
2.85
2.35
1.85
1.35
0.85
*
****
****
****
****
****
0
200
400
600
800
1000
1500
[NaCl] (mM)

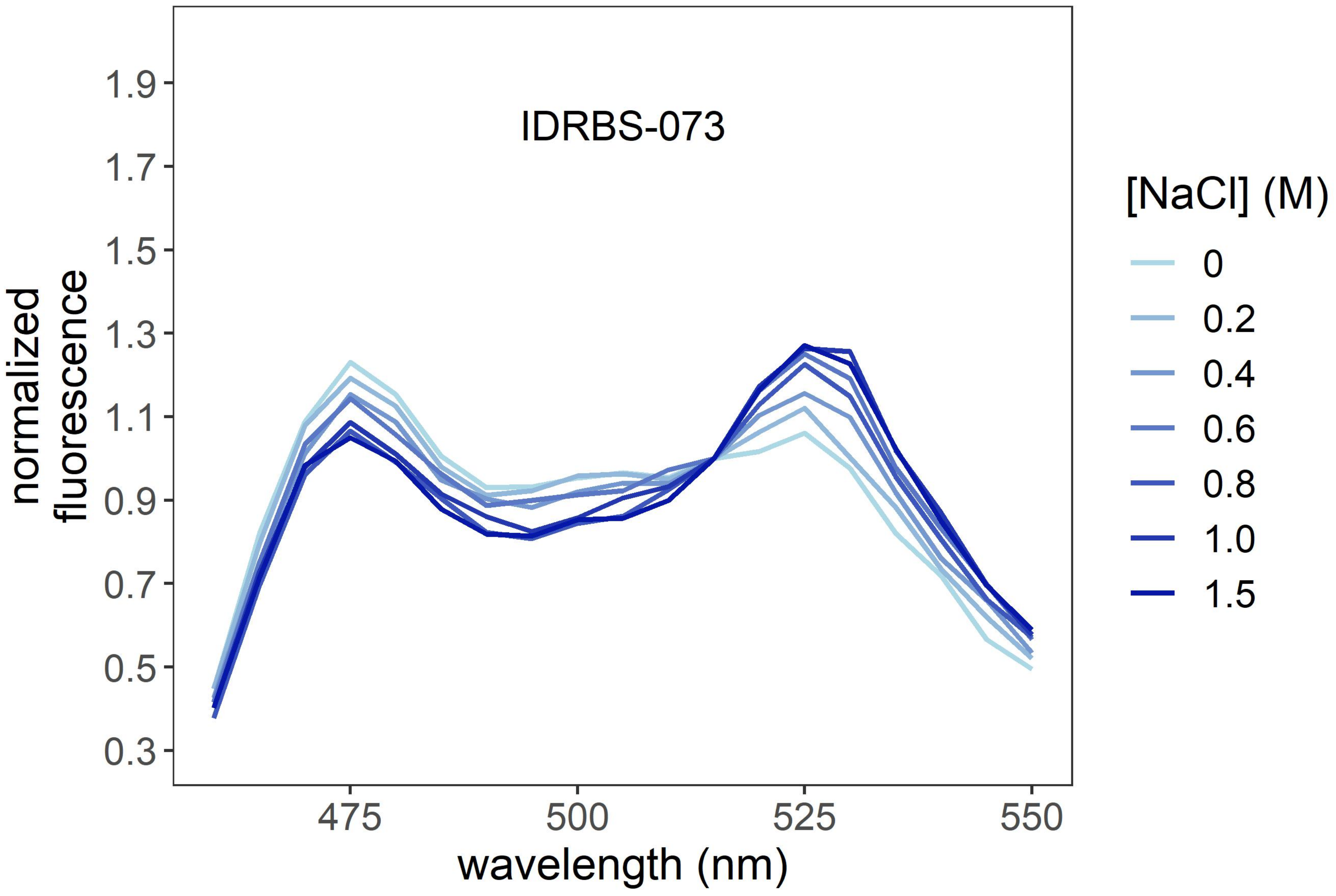
IDRBS-073
normalized fluorescence
1.9
1.7
1.5
1.3
1.1
0.9
0.7
0.5
0.3
475
500
525
550
wavelength (nm)
[NaCl] (M)
0
0.2
0.4
0.6
0.8
1.0
1.5

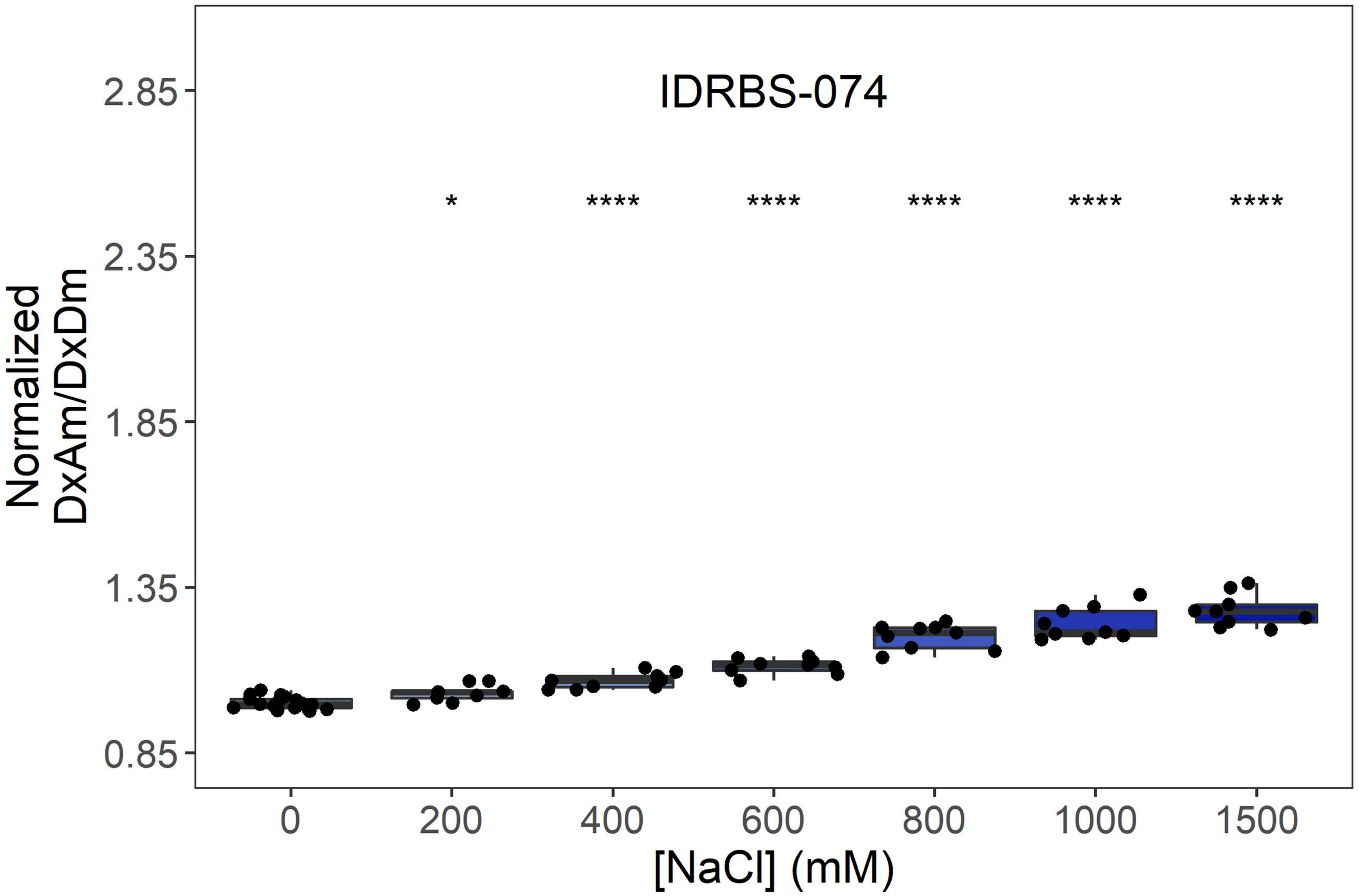
IDRBS-074
*
****
****
****
****
****
2.85
2.35
1.85
1.35
0.85
Normalized
DxAm/DxDm
0
200
400
600
800
1000
1500
[NaCl] (mM)

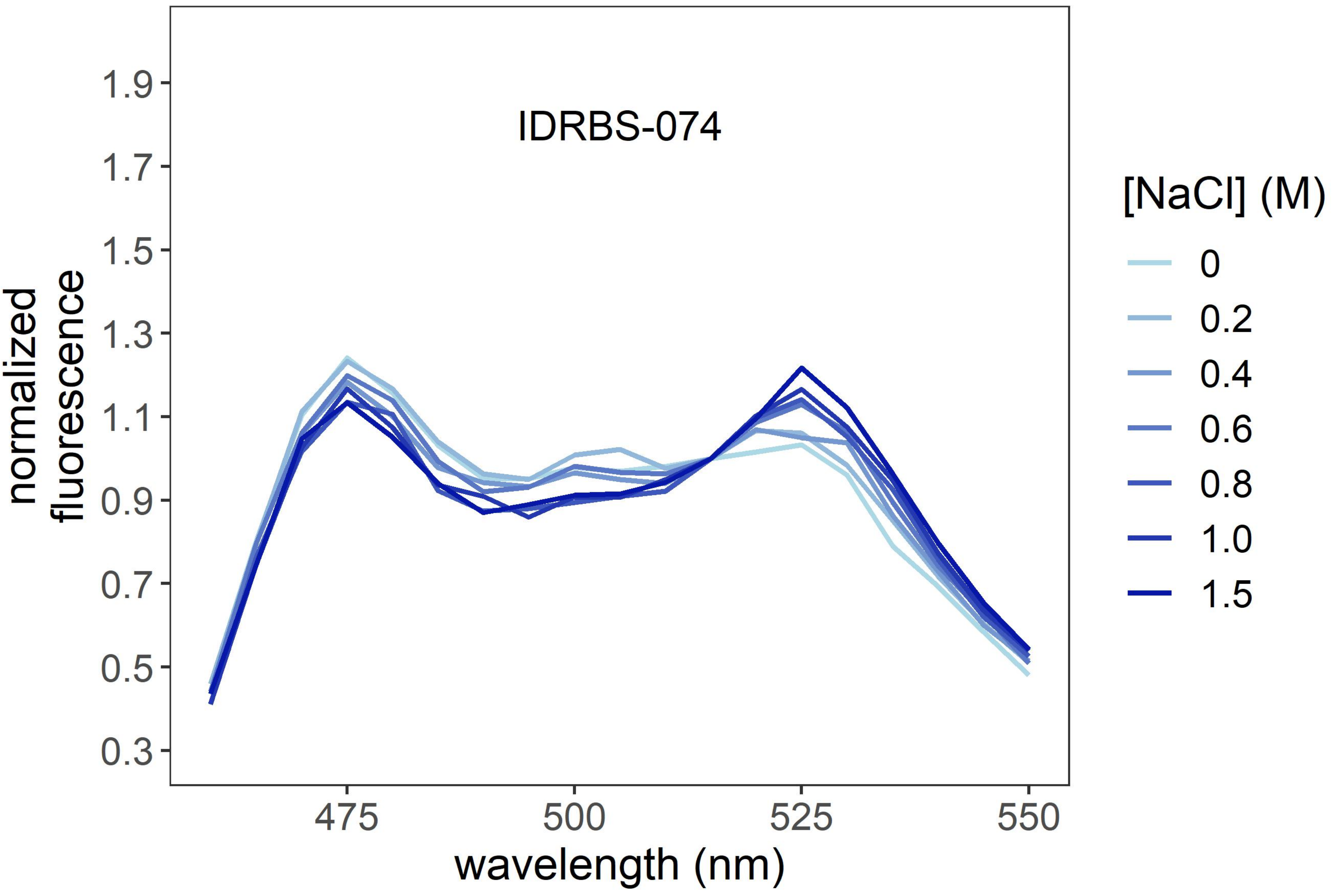
IDRBS-074
1.9
1.7
1.5
1.3
1.1
0.9
0.7
0.5
0.3
normalized
fluorescence
475
500
525
550
wavelength (nm)
[NaCl] (M)
0
0.2
0.4
0.6
0.8
1.0
1.5

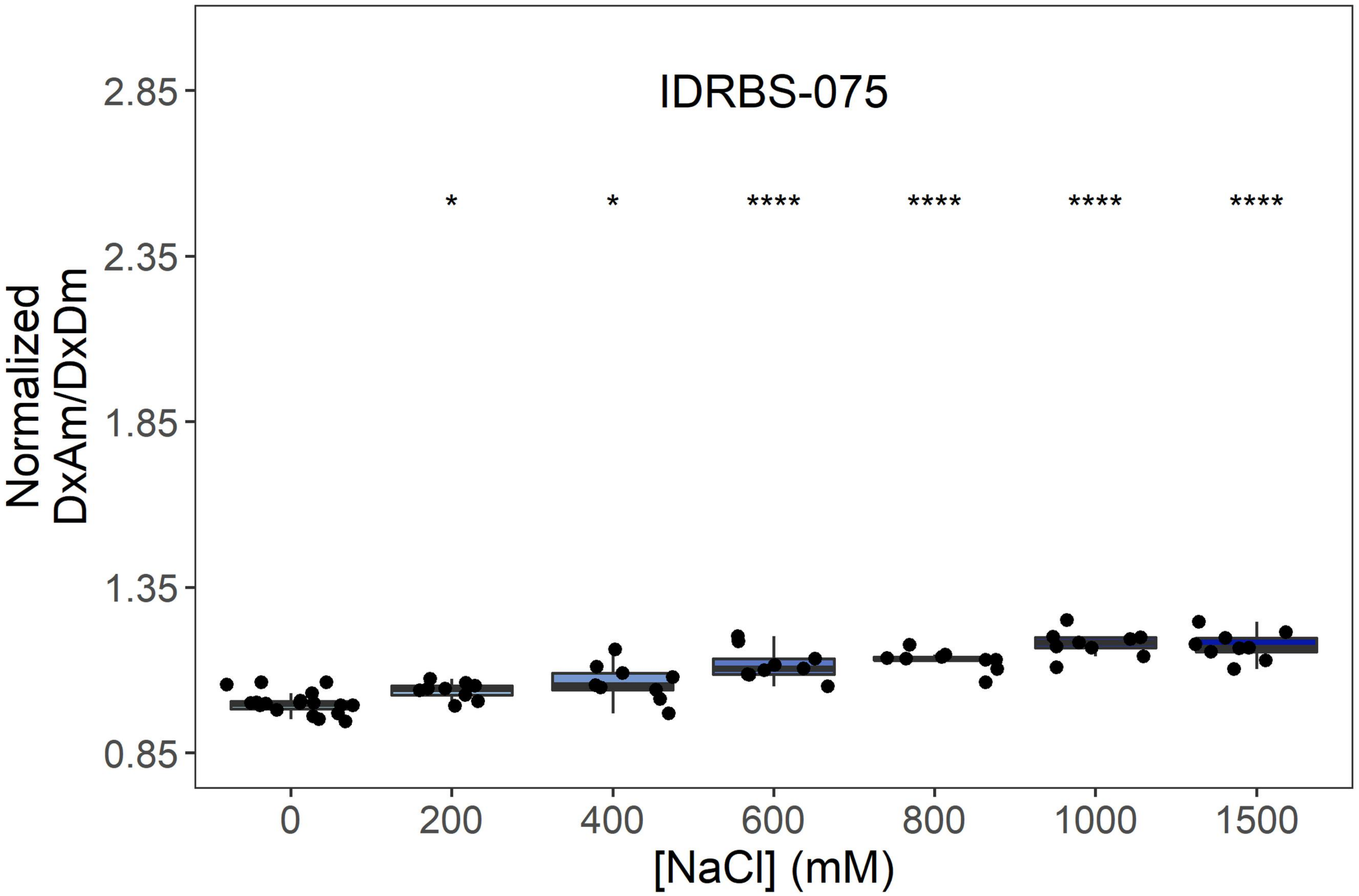
IDRBS-075
Normalized DxAm/DxDm
2.85
2.35
1.85
1.35
0.85
* * **** **** **** ****
0
200
400
600
800
1000
1500
[NaCl] (mM)

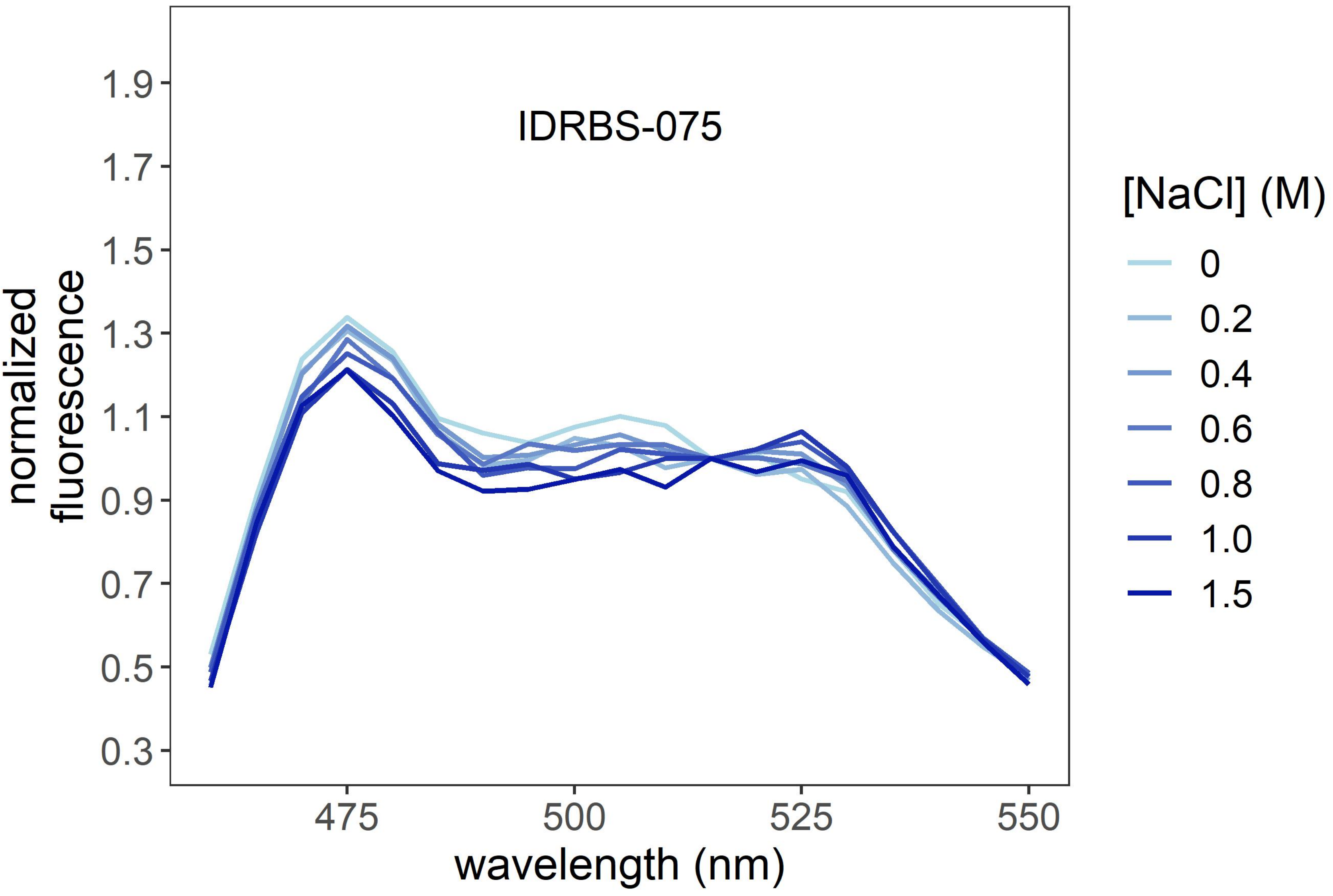
IDRBS-075
normalized fluorescence
1.9
1.7
1.5
1.3
1.1
0.9
0.7
0.5
0.3
475
500
525
550
wavelength (nm)
[NaCl] (M)
0
0.2
0.4
0.6
0.8
1.0
1.5

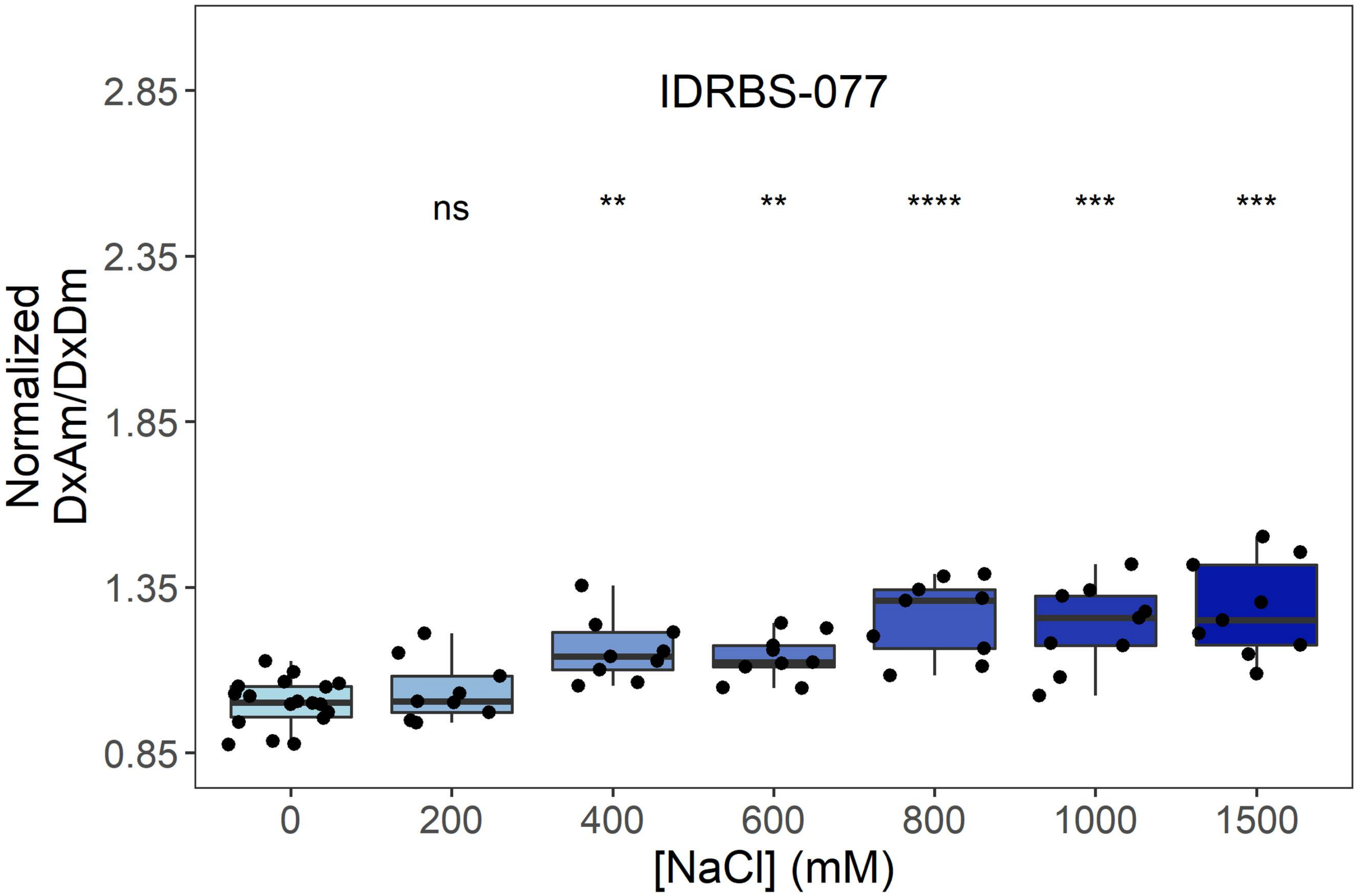
IDRBS-077
ns
**
**
****
***
***
Normalized DxAm/DxDm
2.85
2.35
1.85
1.35
0.85
0
200
400
600
800
1000
1500
[NaCl] (mM)

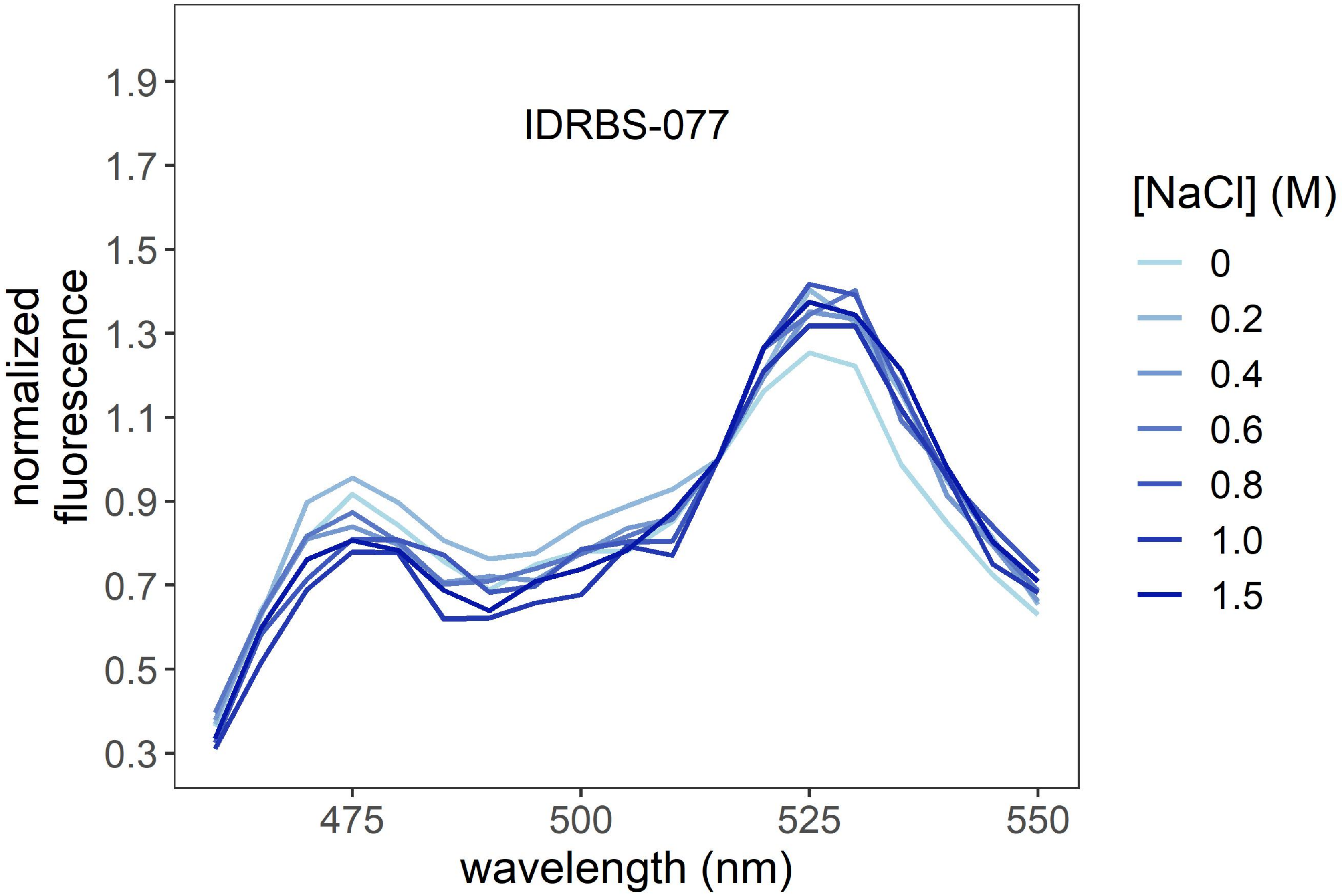
IDRBS-077
normalized fluorescence
1.9
1.7
1.5
1.3
1.1
0.9
0.7
0.5
0.3
475
500
525
550
wavelength (nm)
[NaCl] (M)
0
0.2
0.4
0.6
0.8
1.0
1.5

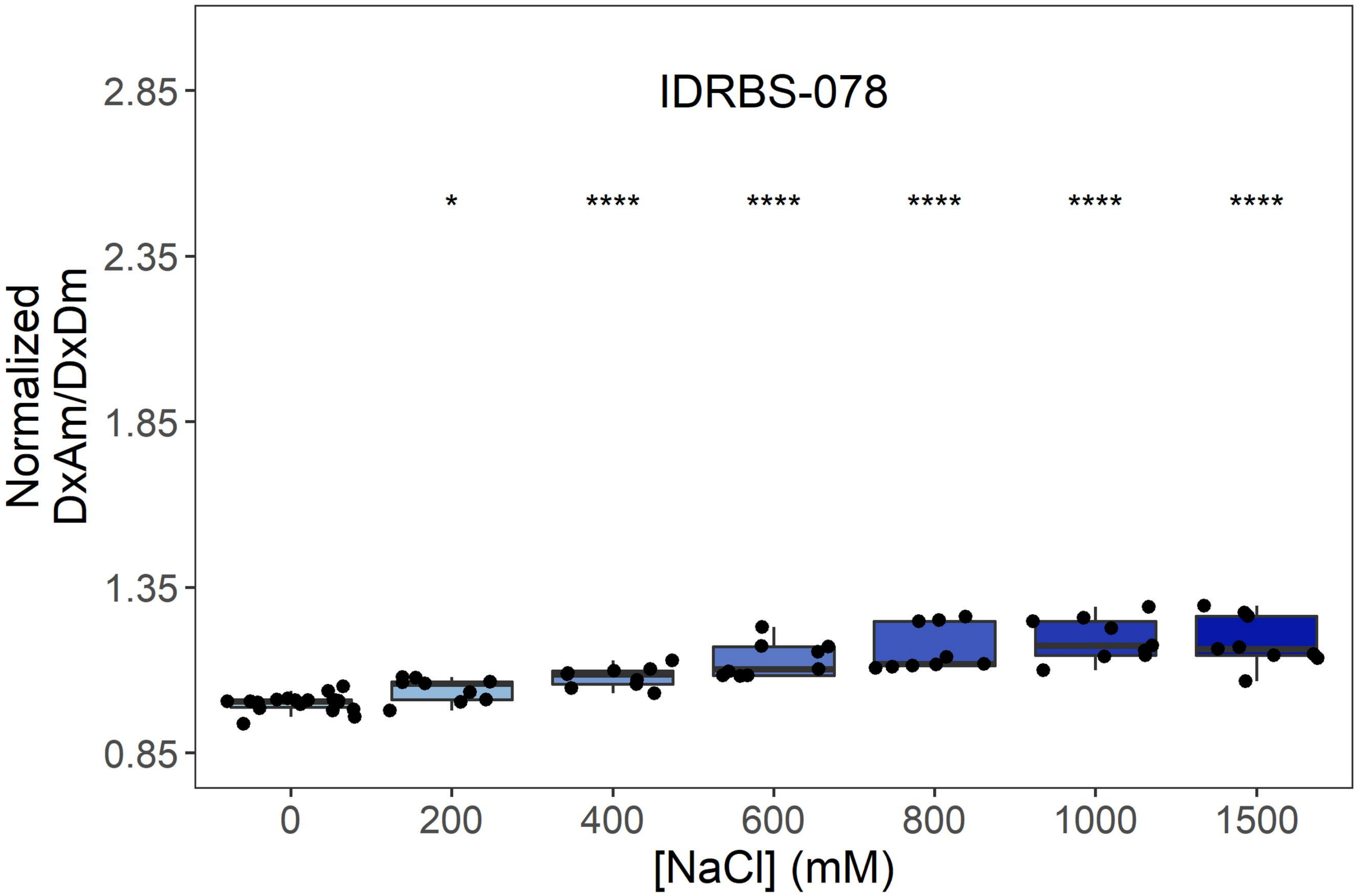
IDRBS-078
Normalized DxAm/DxDm
2.85
2.35
1.85
1.35
0.85
*
****
****
****
****
****
0
200
400
600
800
1000
1500
[NaCl] (mM)

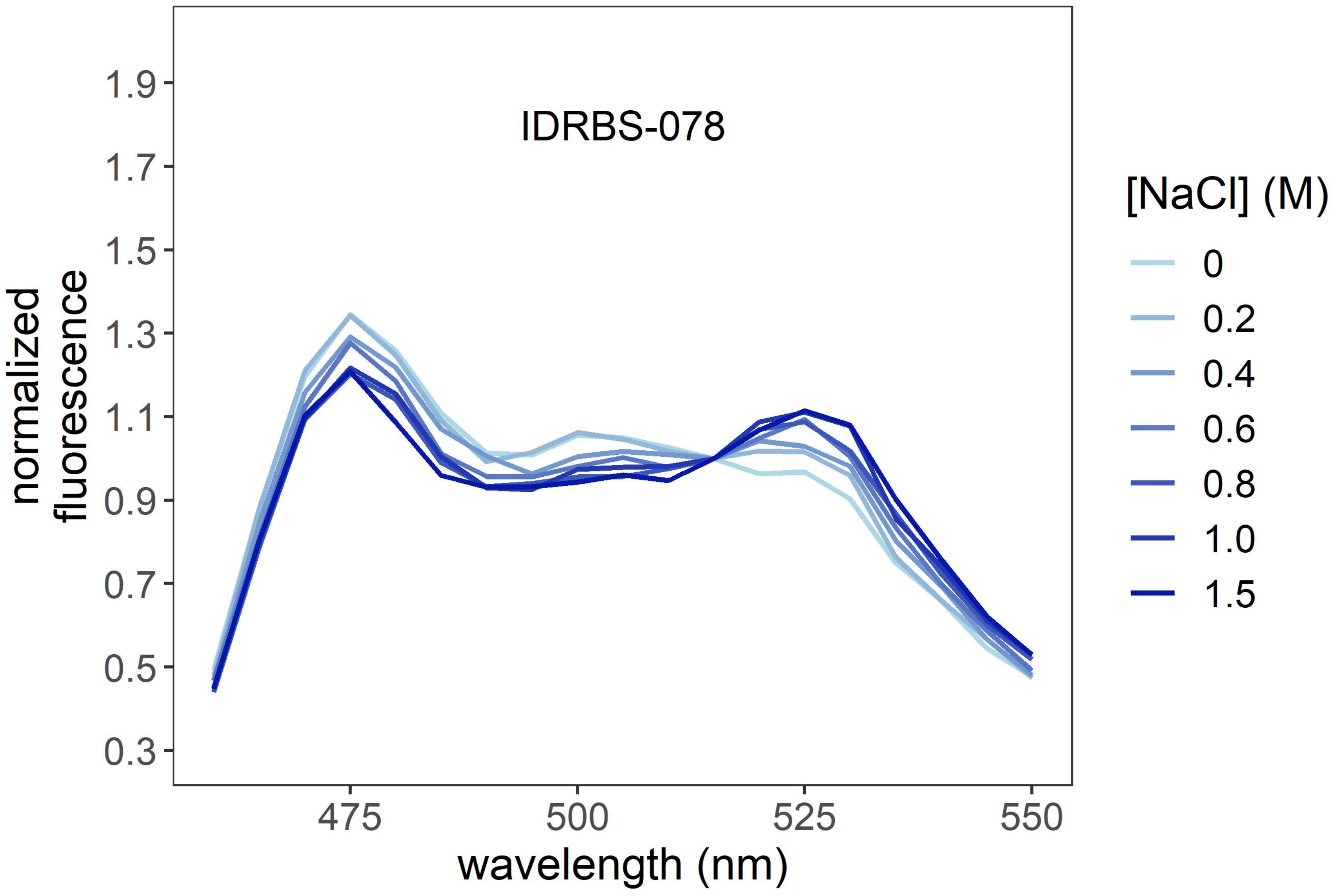
IDRBS-078
normalized fluorescence
1.9
1.7
1.5
1.3
1.1
0.9
0.7
0.5
0.3
475
500
525
550
wavelength (nm)
[NaCl] (M)
0
0.2
0.4
0.6
0.8
1.0
1.5

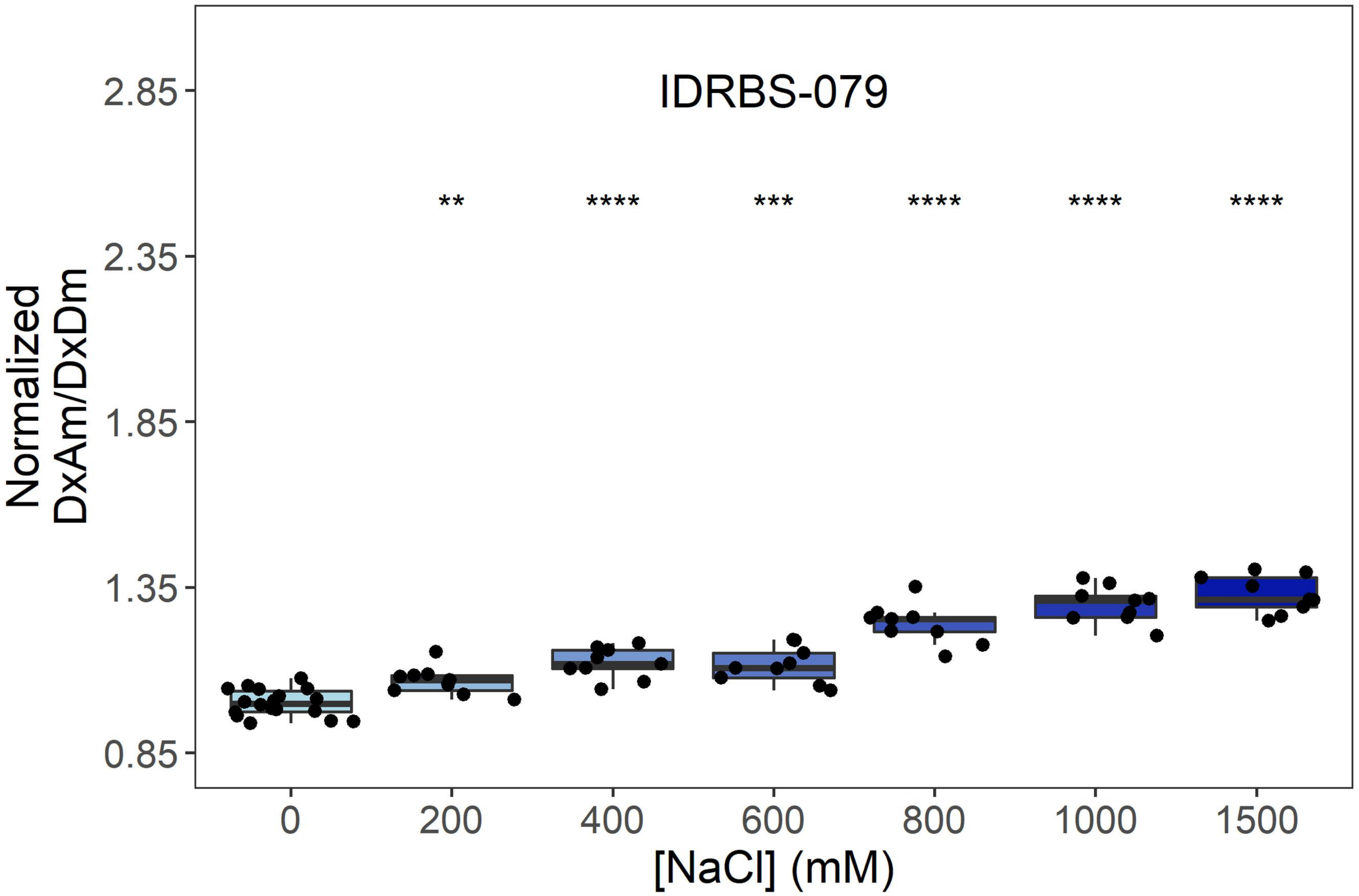
IDRBS-079
Normalized DxAm/DxDm
2.85
2.35
1.85
1.35
0.85
**
****
***
****
****
****
0
200
400
600
800
1000
1500
[NaCl] (mM)

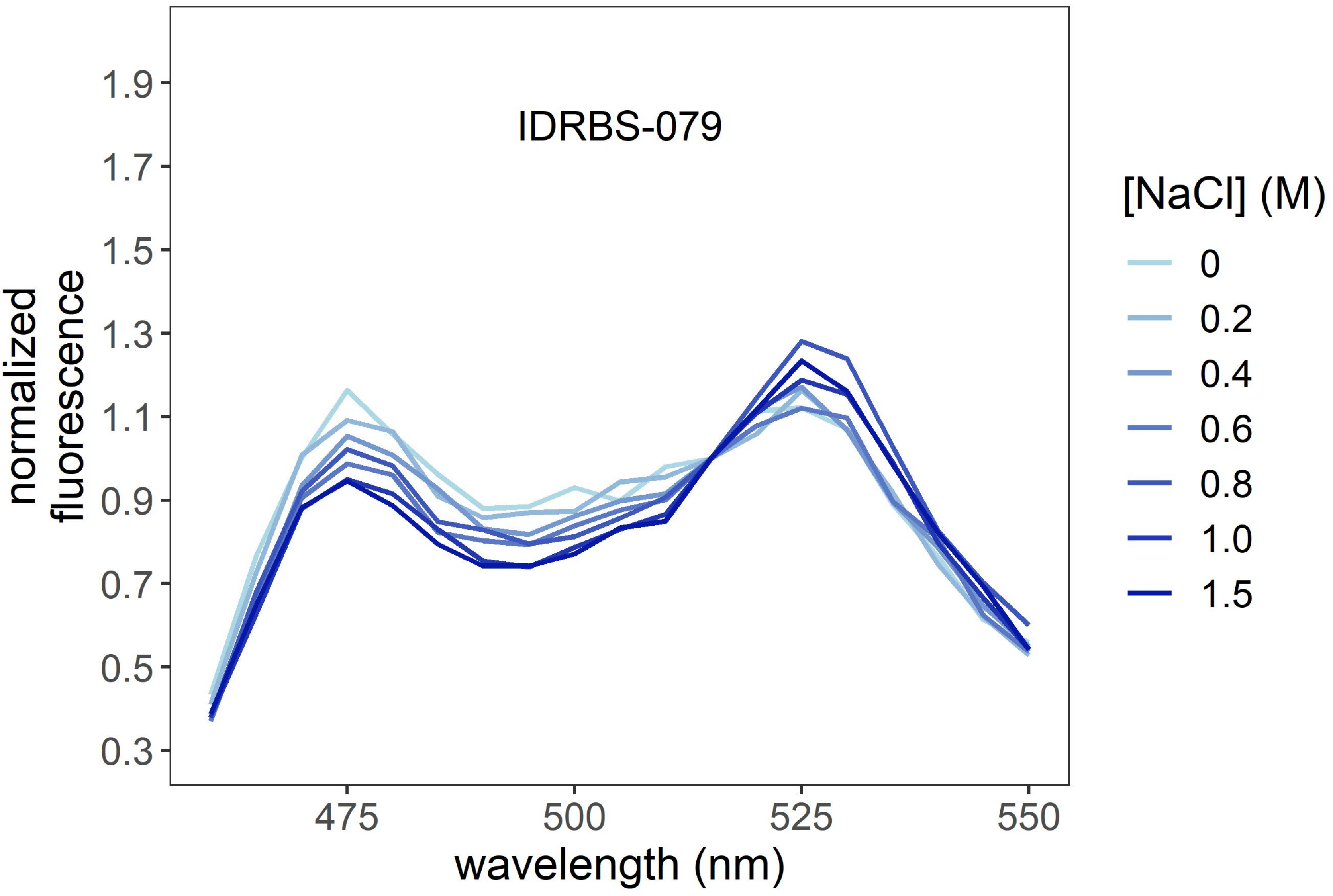
IDRBS-079
normalized fluorescence
1.9
1.7
1.5
1.3
1.1
0.9
0.7
0.5
0.3
475
500
525
550
wavelength (nm)
[NaCl] (M)
0
0.2
0.4
0.6
0.8
1.0
1.5

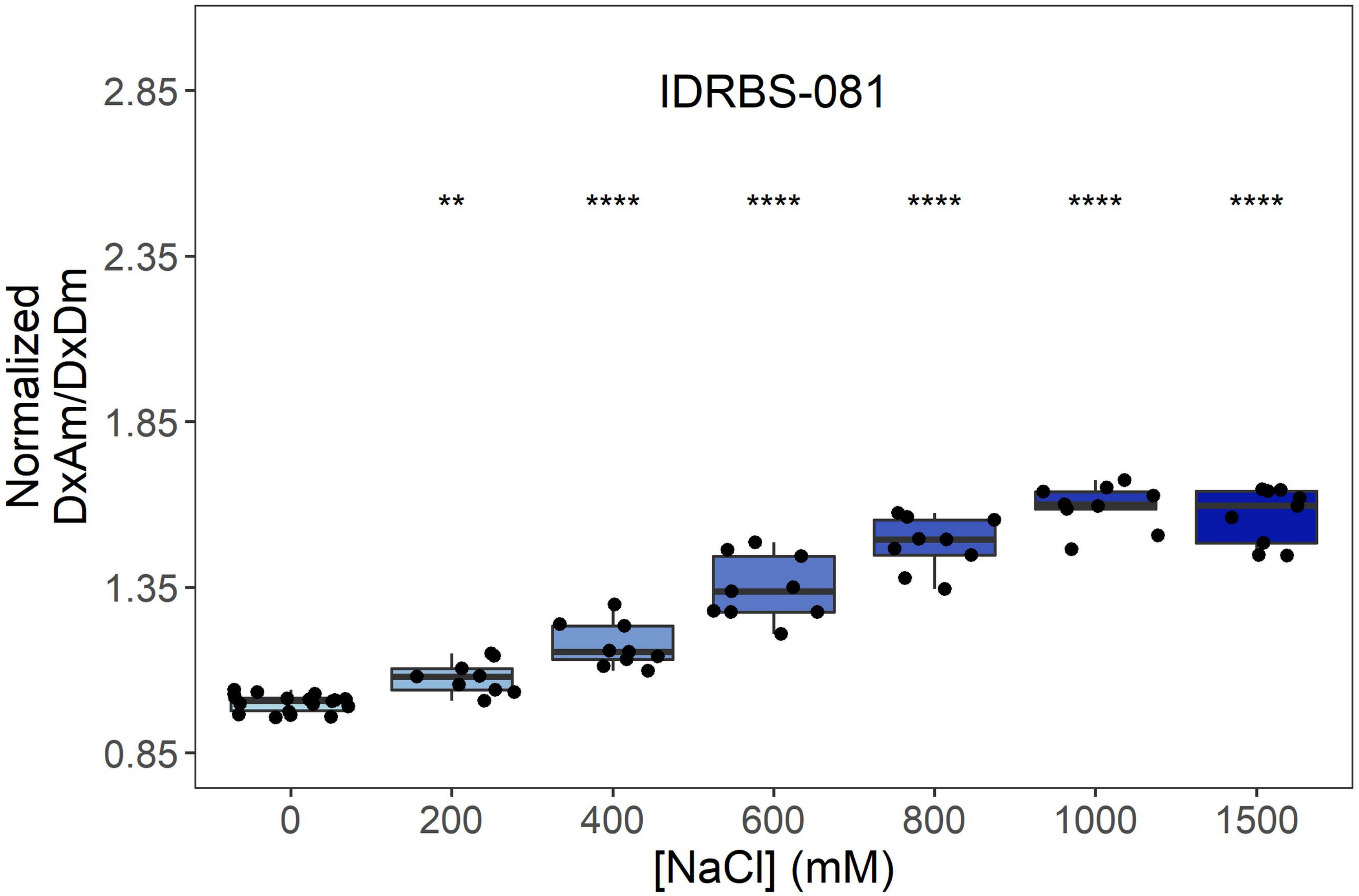
IDRBS-081
Normalized DxAm/DxDm
2.85
2.35
1.85
1.35
0.85
**
****
****
****
****
****
0
200
400
600
800
1000
1500
[NaCl] (mM)

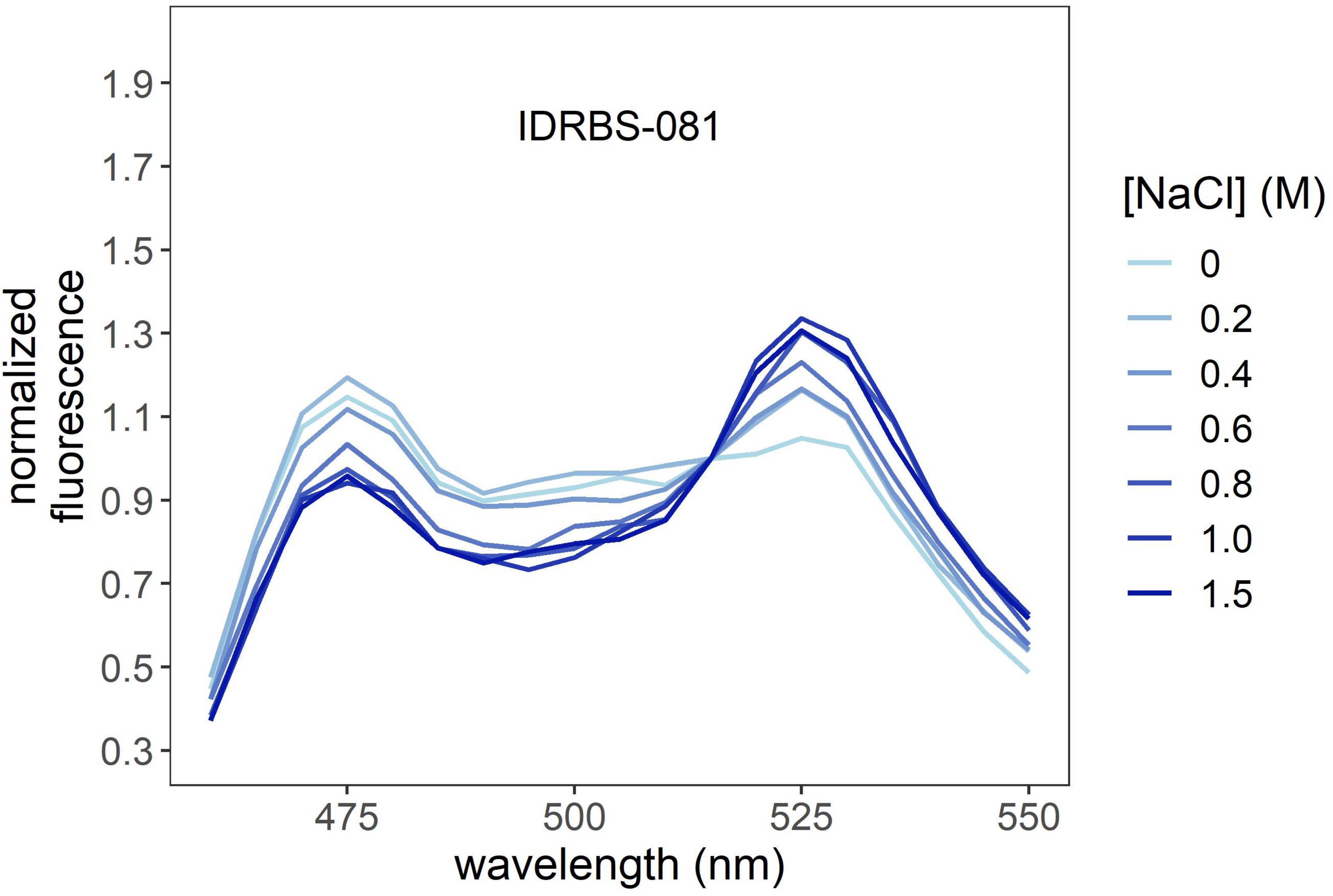
IDRBS-081
normalized fluorescence
1.9
1.7
1.5
1.3
1.1
0.9
0.7
0.5
0.3
475
500
525
550
wavelength (nm)
[NaCl] (M)
0
0.2
0.4
0.6
0.8
1.0
1.5

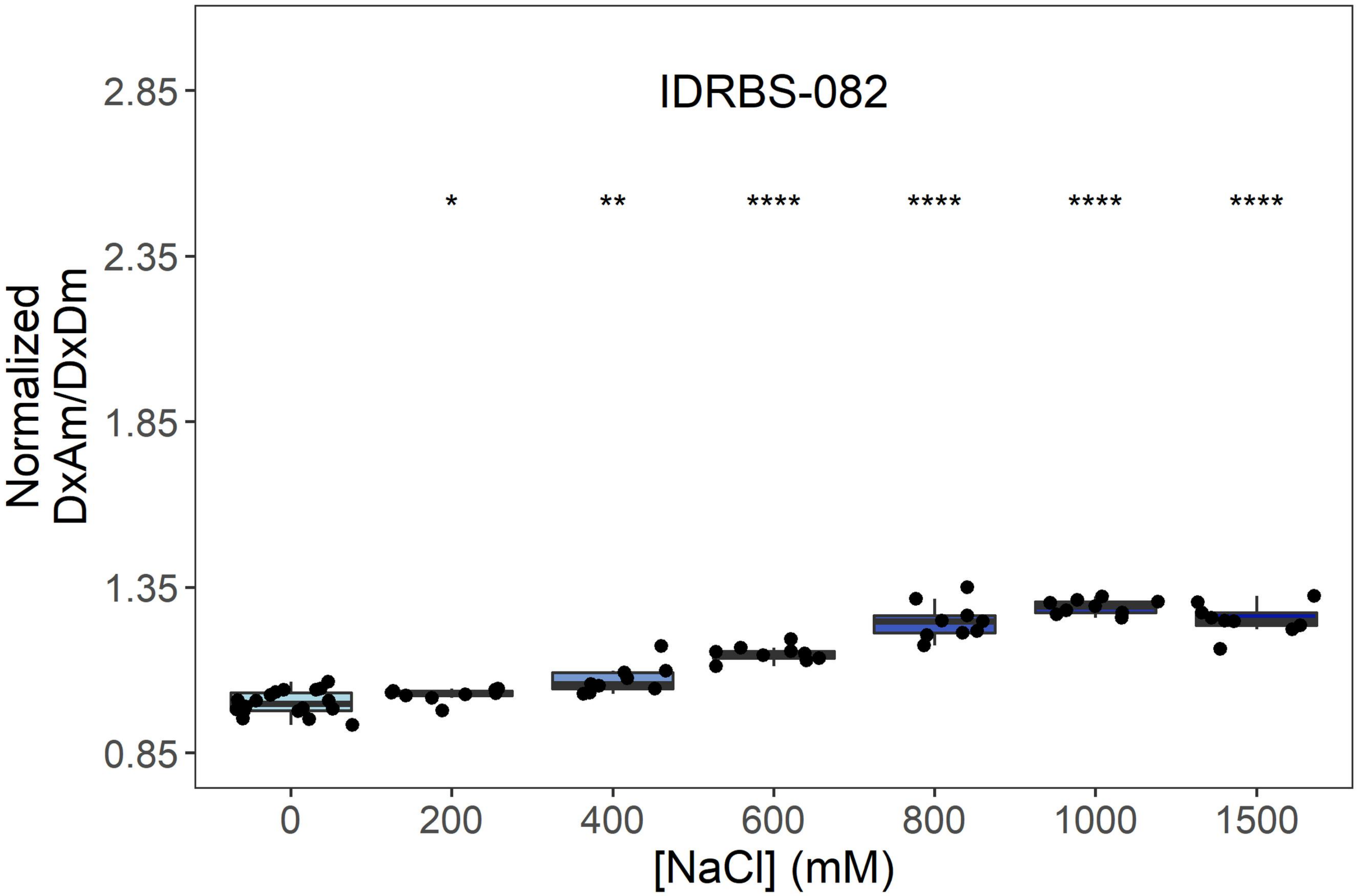
IDRBS-082
*
**
****
****
****
****
Normalized
DxAm/DxDm
2.85
2.35
1.85
1.35
0.85
0
200
400
600
800
1000
1500
[NaCl] (mM)

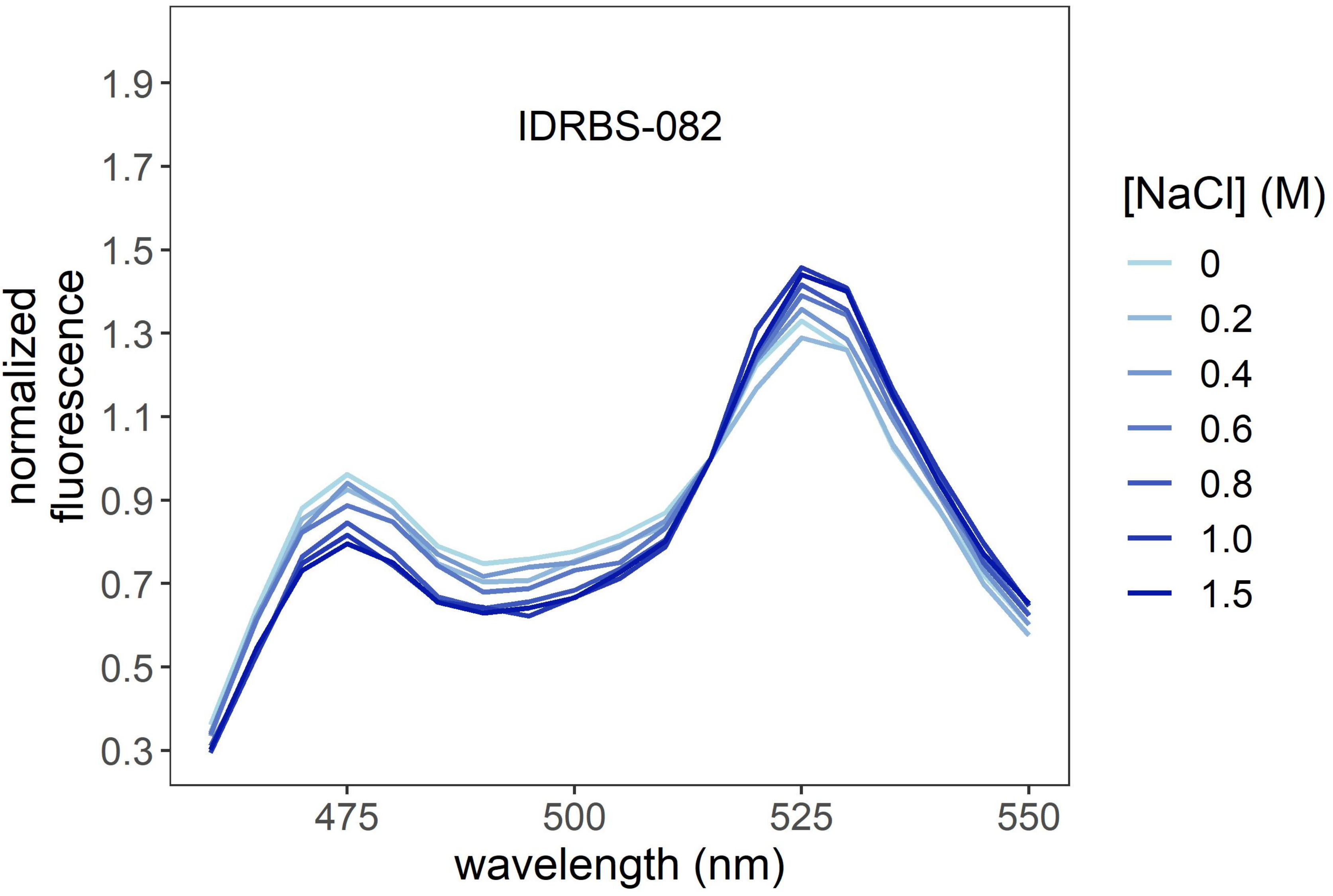
IDRBS-082
normalized
fluorescence
1.9
1.7
1.5
1.3
1.1
0.9
0.7
0.5
0.3
475
500
525
550
wavelength (nm)
[NaCl] (M)
0
0.2
0.4
0.6
0.8
1.0
1.5

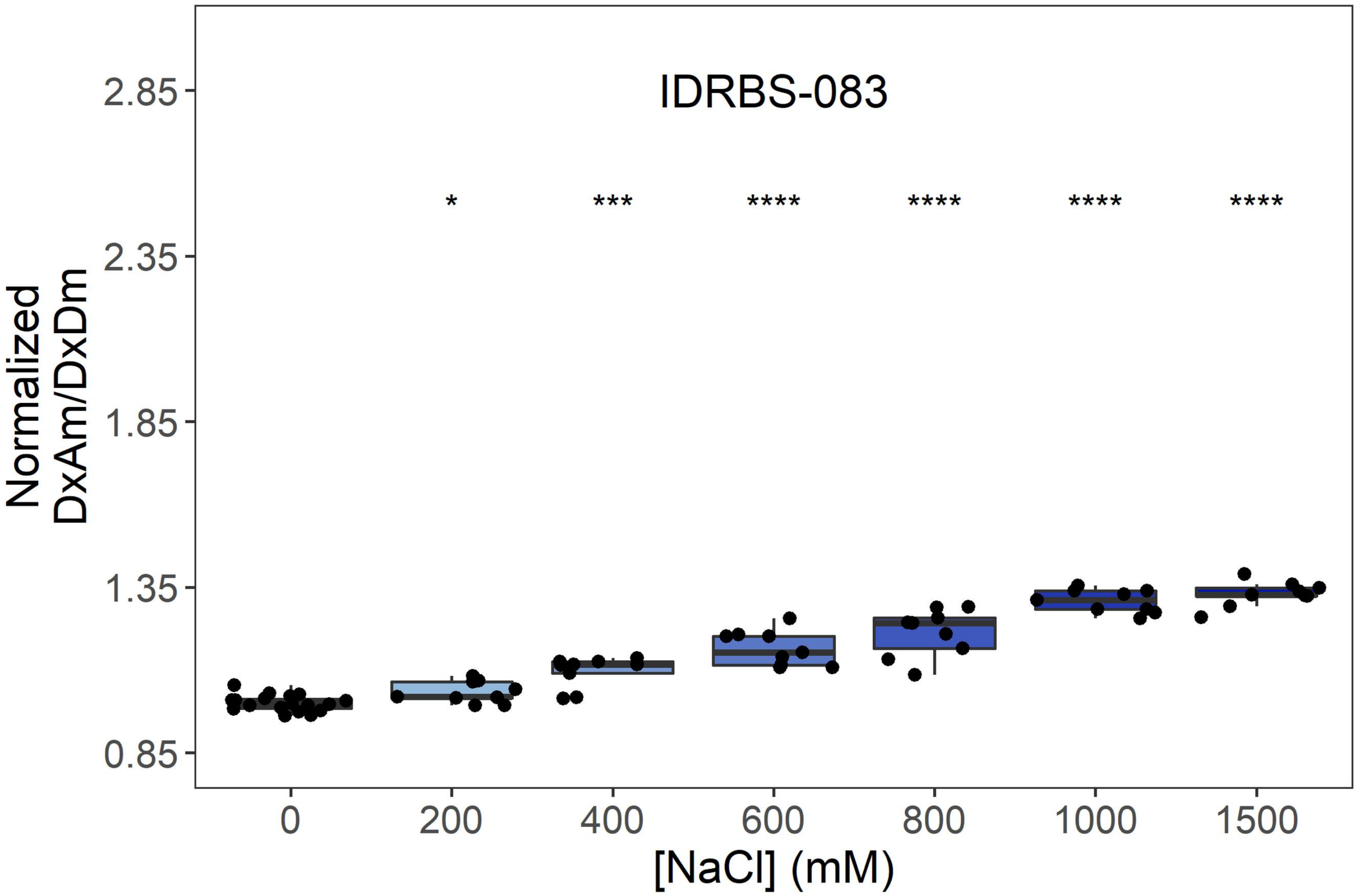
IDRBS-083
Normalized DxAm/DxDm
2.85
2.35
1.85
1.35
0.85
*
***
****
****
****
****
0
200
400
600
800
1000
1500
[NaCl] (mM)

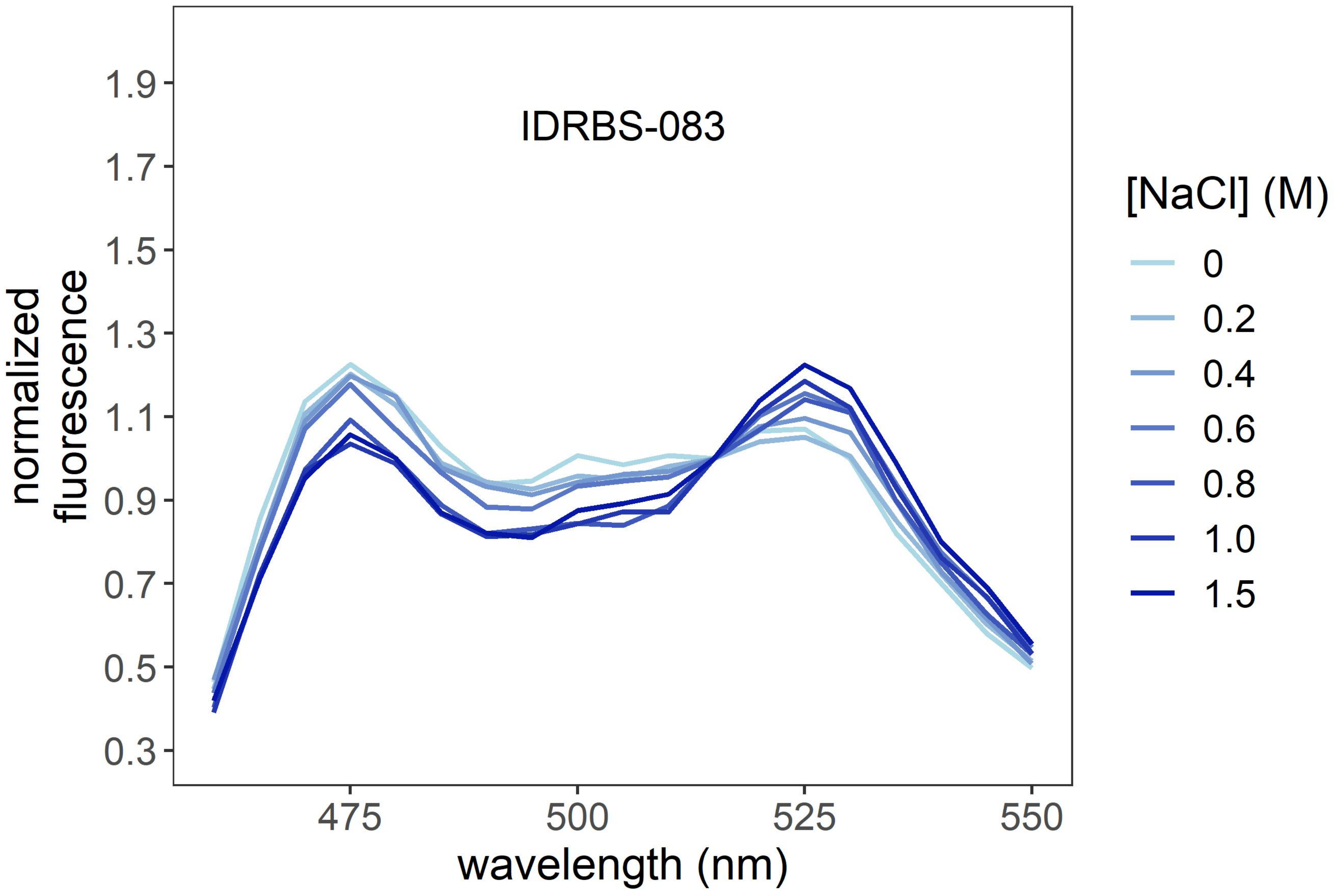
IDRBS-083
normalized fluorescence
1.9
1.7
1.5
1.3
1.1
0.9
0.7
0.5
0.3
475
500
525
550
wavelength (nm)
[NaCl] (M)
0
0.2
0.4
0.6
0.8
1.0
1.5

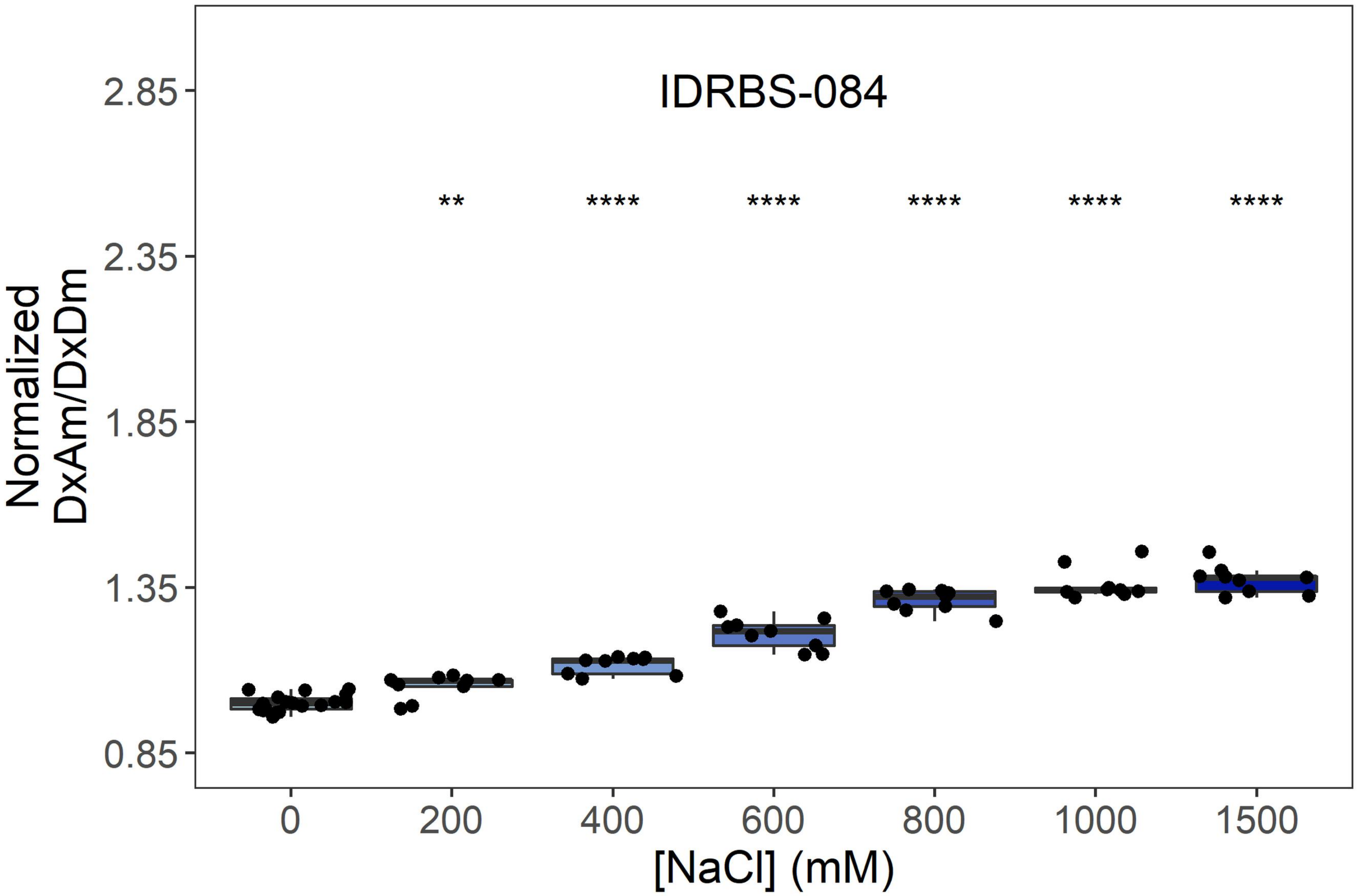

IDRBS-084
**
****
****
****
****
****
2.85
2.35
1.85
1.35
0.85
Normalized DxAm/DxDm
0
200
400
600
800
1000
1500
[NaCl] (mM)


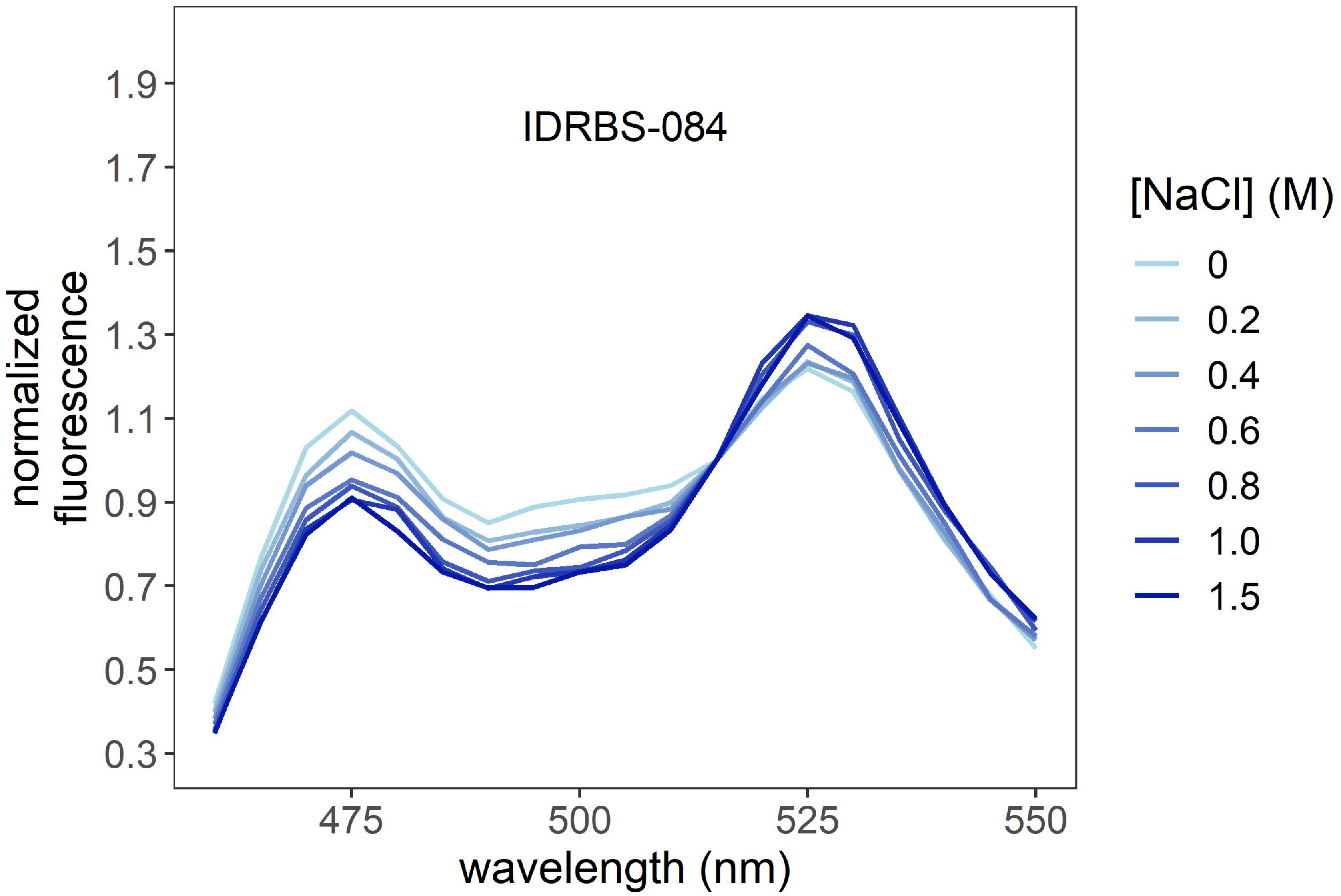

IDRBS-084
1.9
1.7
1.5
1.3
1.1
0.9
0.7
0.5
0.3
normalized fluorescence
475
500
525
550
wavelength (nm)
[NaCl] (M)
0
0.2
0.4
0.6
0.8
1.0
1.5

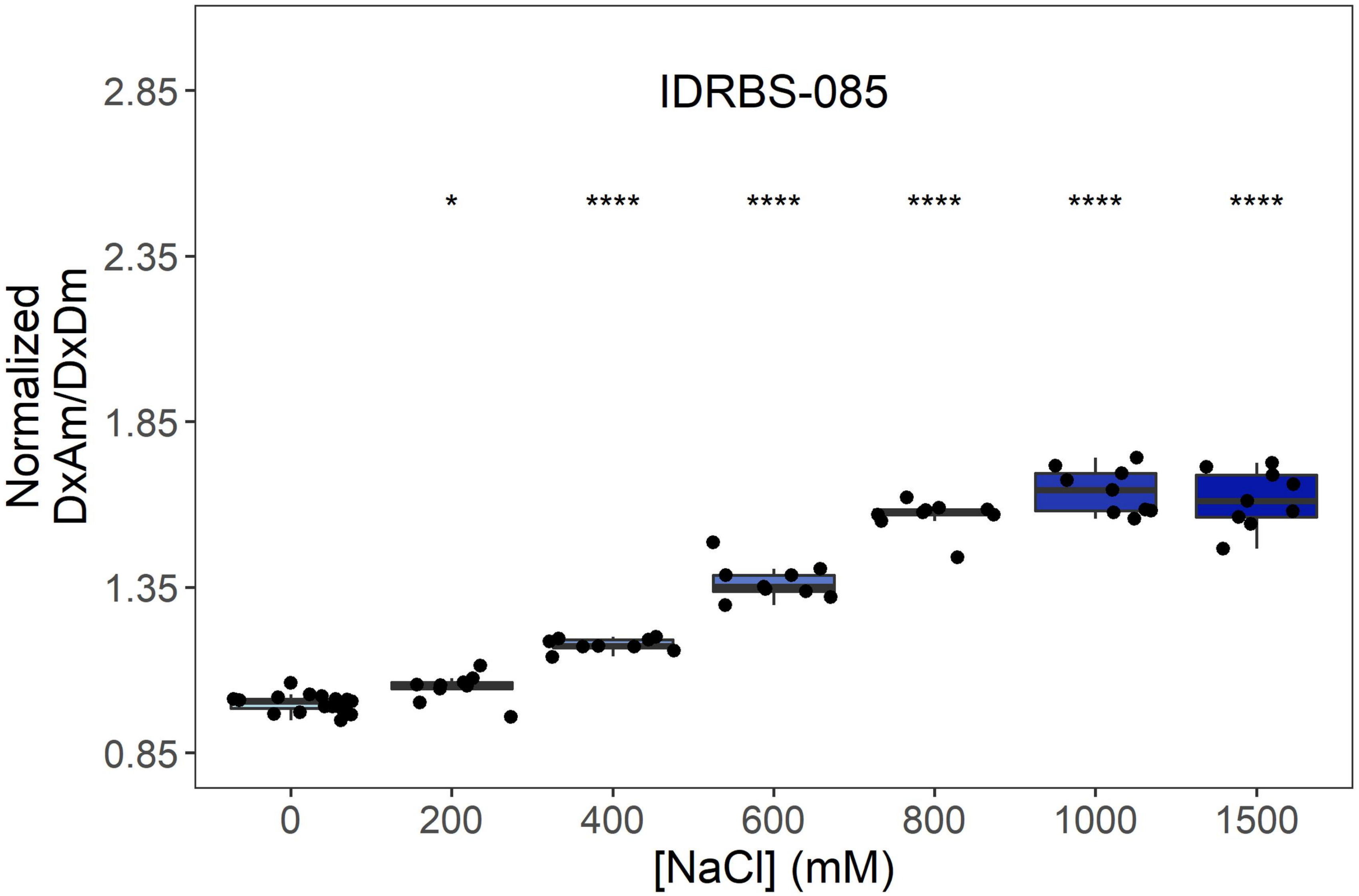
IDRBS-085
Normalized
DxAm/DxDm
2.85
2.35
1.85
1.35
0.85
*
****
****
****
****
****
0
200
400
600
800
1000
1500
[NaCl] (mM)

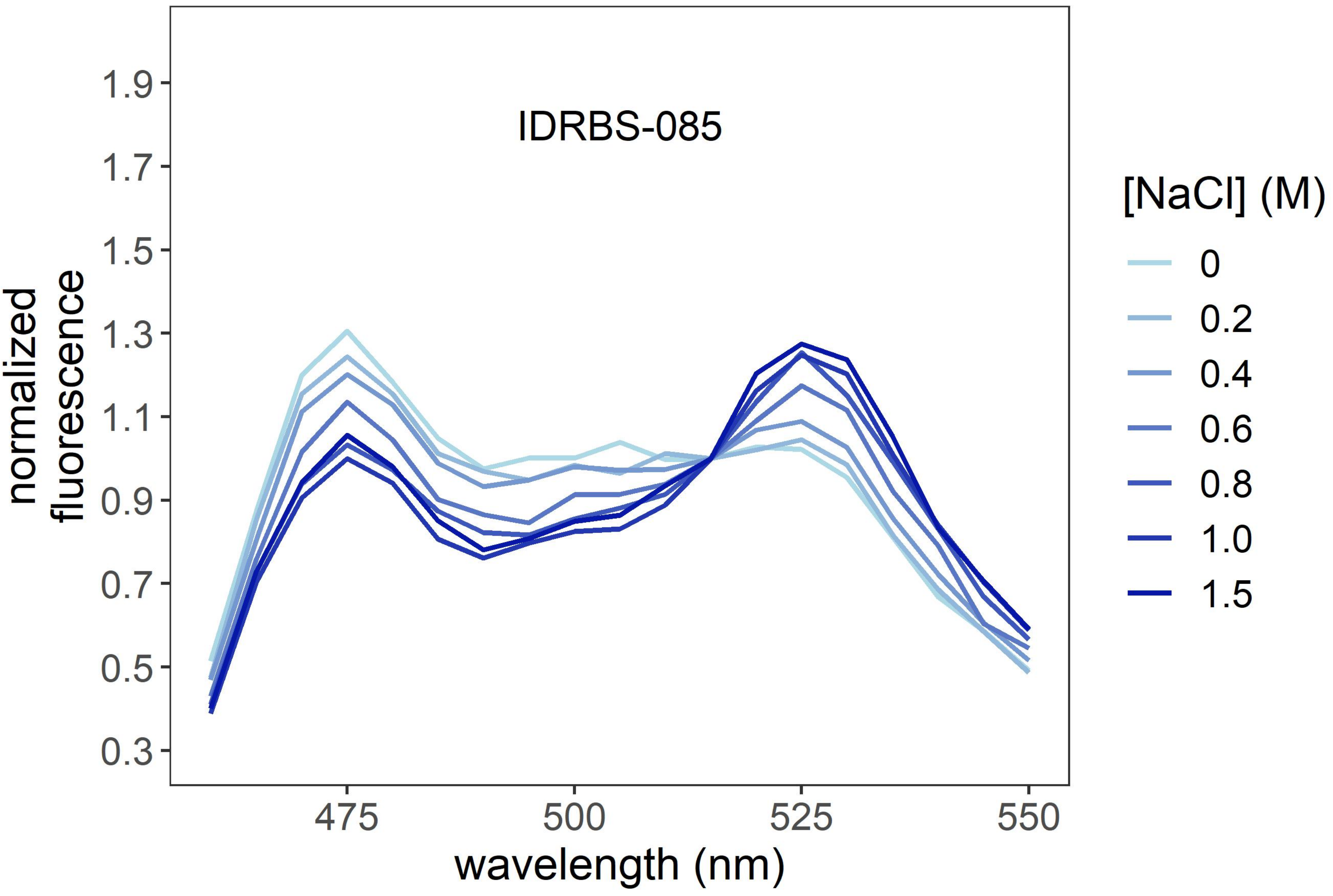
IDRBS-085
normalized
fluorescence
1.9
1.7
1.5
1.3
1.1
0.9
0.7
0.5
0.3
475
500
525
550
wavelength (nm)
[NaCl] (M)
0
0.2
0.4
0.6
0.8
1.0
1.5

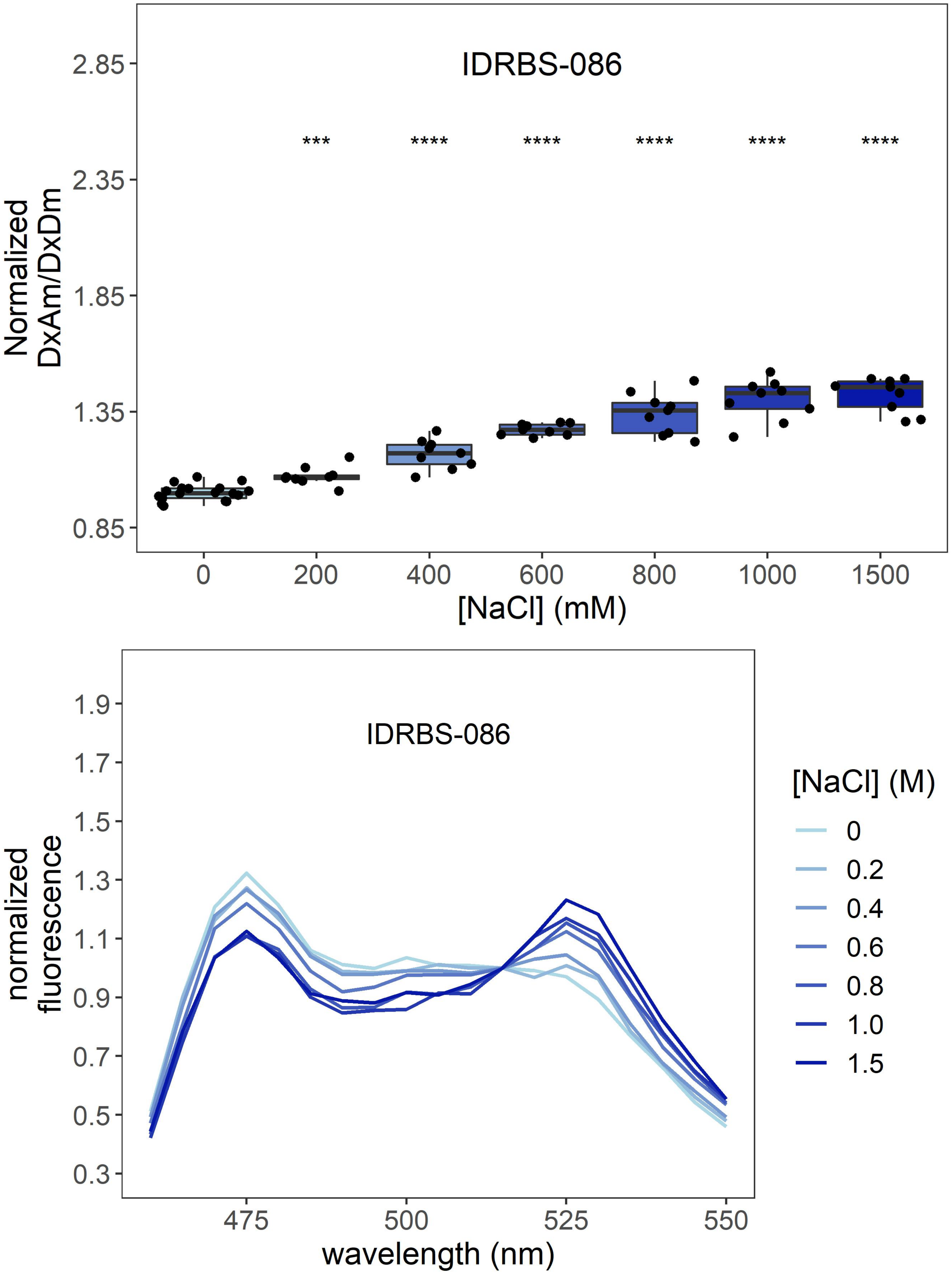
IDRBS-086
2.85
2.35
1.85
1.35
0.85
Normalized
DxAm/DxDm
***
****
****
****
****
****
0
200
400
600
800
1000
1500
[NaCl] (mM)
IDRBS-086
1.9
1.7
1.5
1.3
1.1
0.9
0.7
0.5
0.3
normalized
fluorescence
475
500
525
550
wavelength (nm)
[NaCl] (M)
0
0.2
0.4
0.6
0.8
1.0
1.5

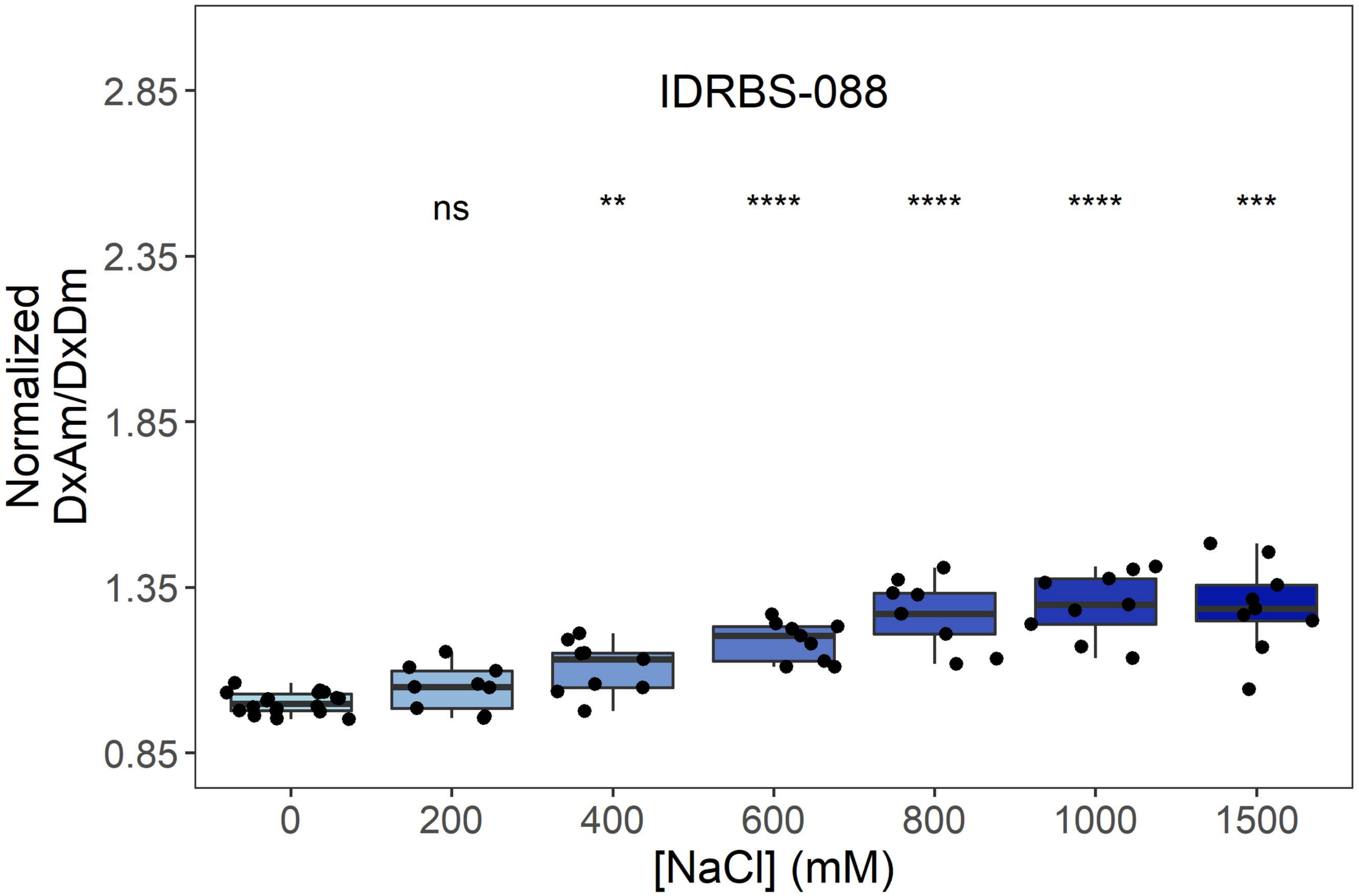
IDRBS-088
ns
**
****
****
****
***
Normalized DxAm/DxDm
2.85
2.35
1.85
1.35
0.85
0
200
400
600
800
1000
1500
[NaCl] (mM)

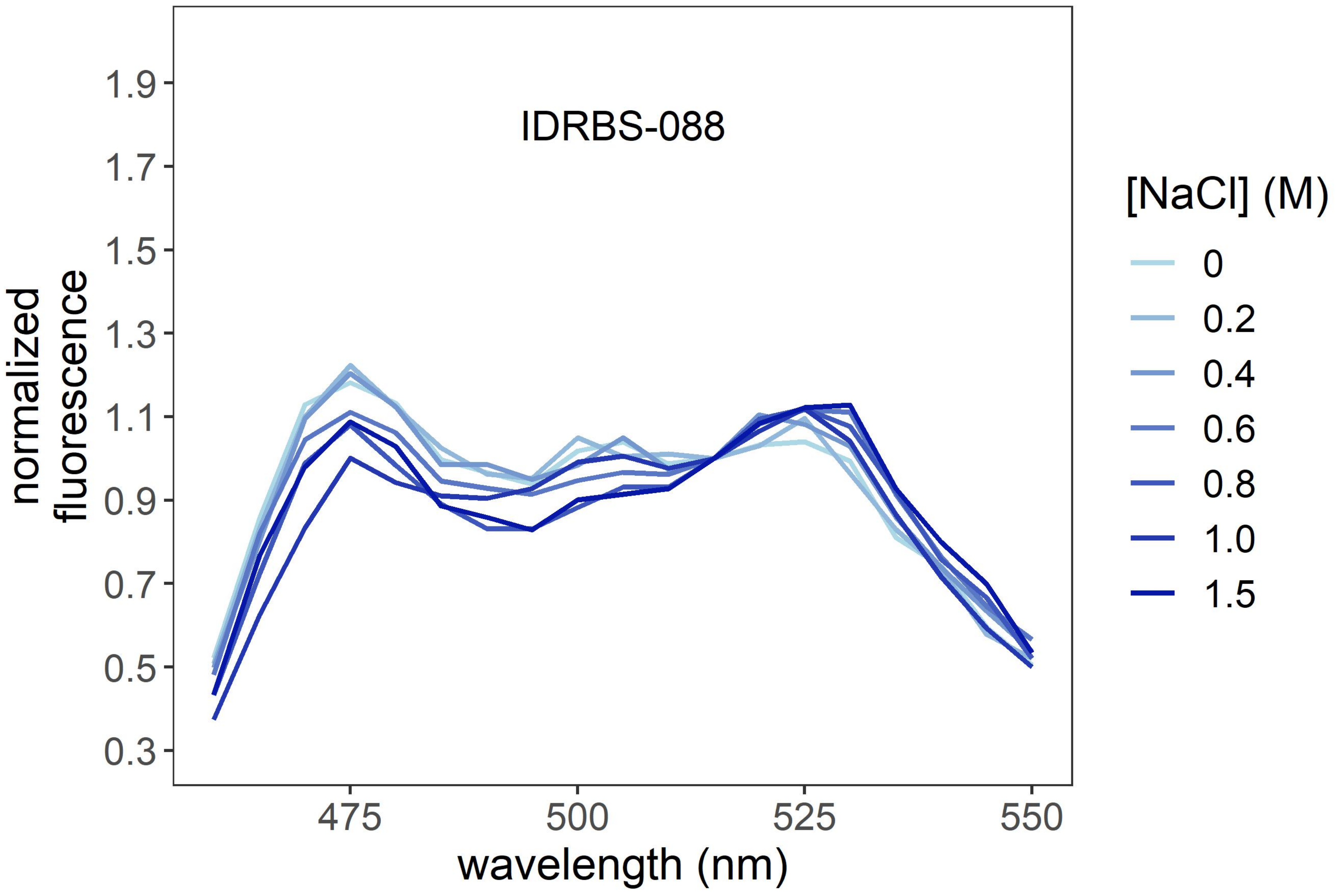
IDRBS-088
normalized fluorescence
1.9
1.7
1.5
1.3
1.1
0.9
0.7
0.5
0.3
475
500
525
550
wavelength (nm)
[NaCl] (M)
0
0.2
0.4
0.6
0.8
1.0
1.5

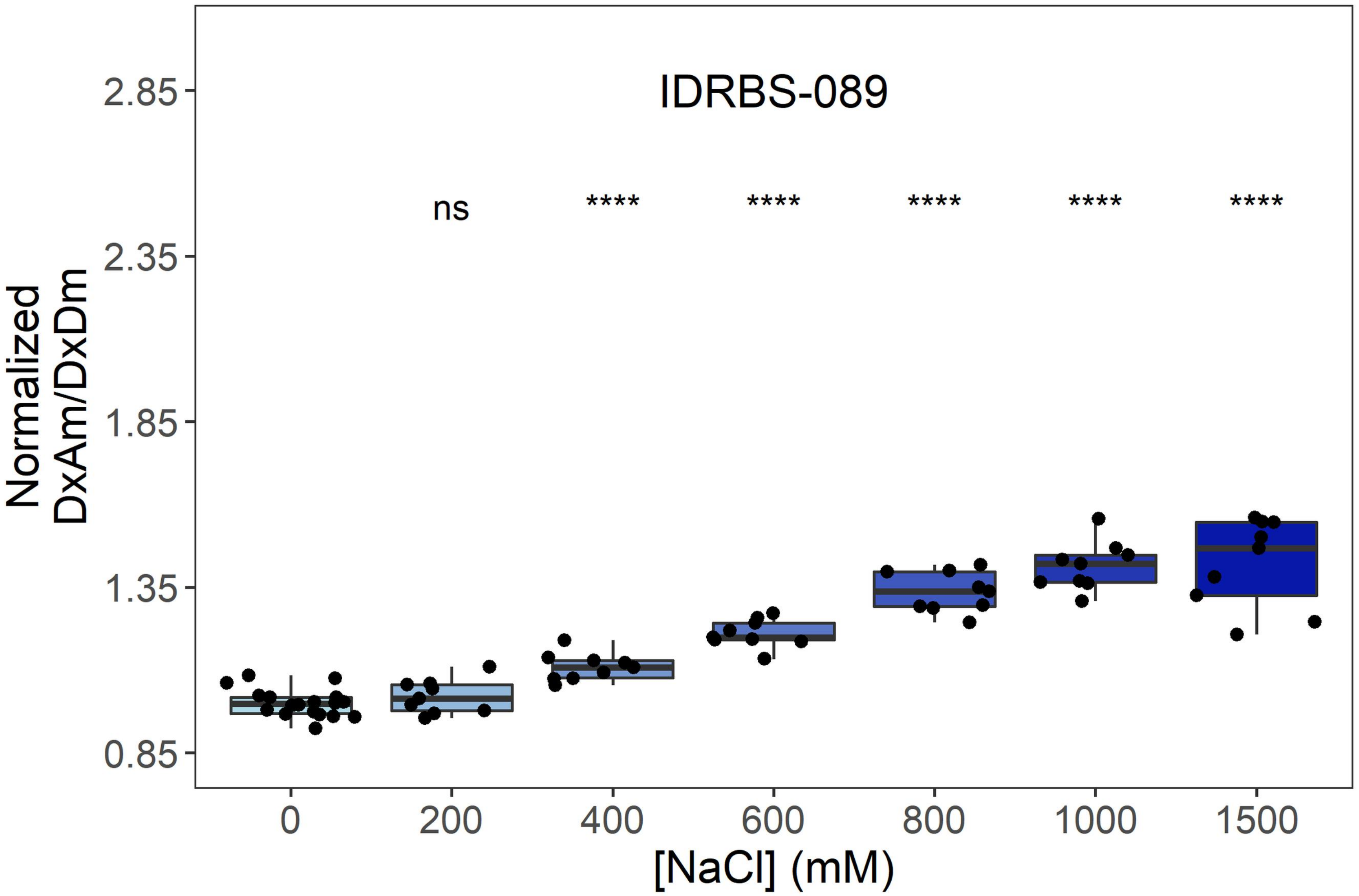
IDRBS-089
ns
****
****
****
****
****
2.85
2.35
1.85
1.35
0.85
Normalized
DxAm/DxDm
0
200
400
600
800
1000
1500
[NaCl] (mM)

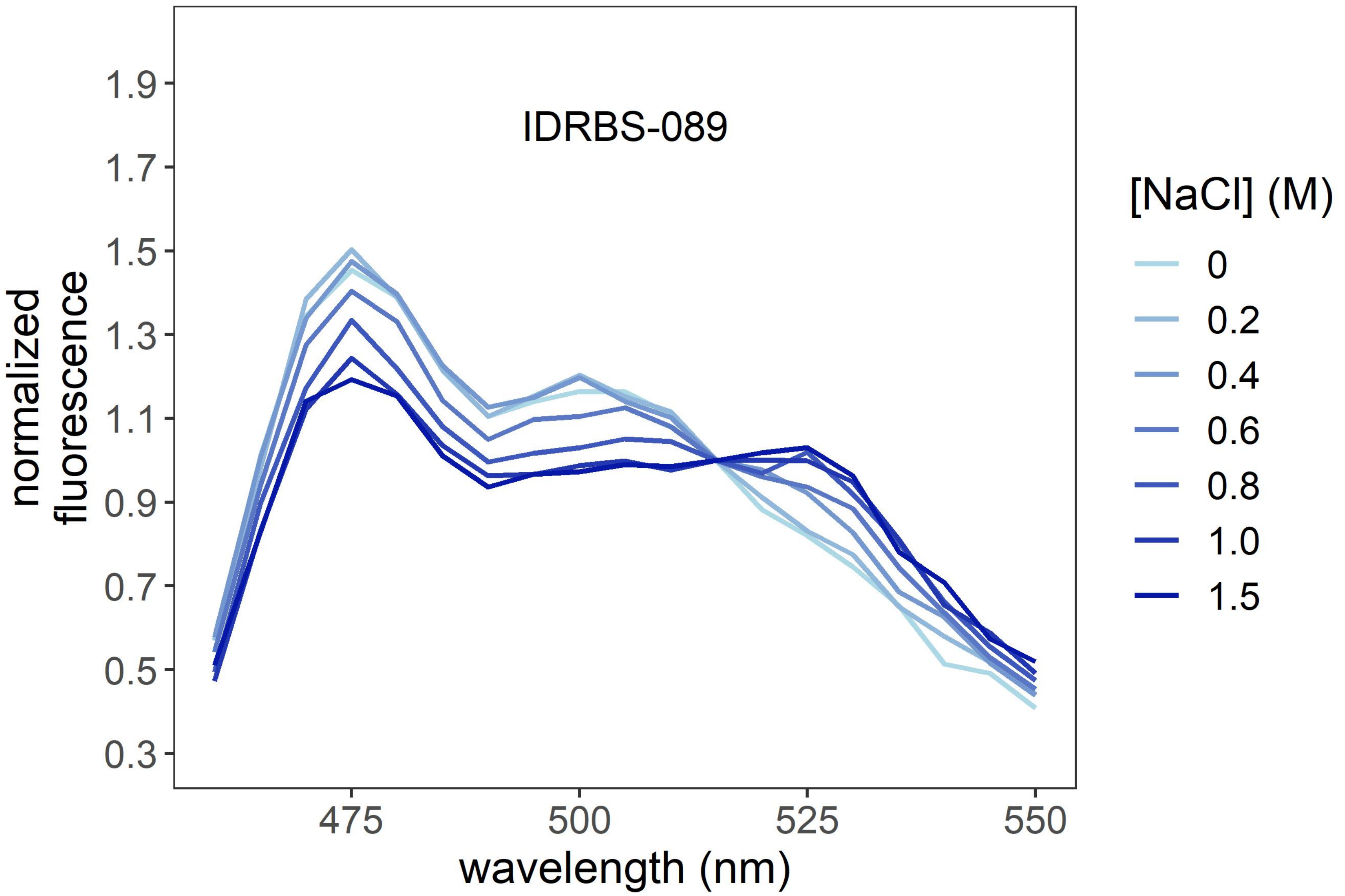
IDRBS-089
1.9
1.7
1.5
1.3
1.1
0.9
0.7
0.5
0.3
normalized
fluorescence
475
500
525
550
wavelength (nm)
[NaCl] (M)
0
0.2
0.4
0.6
0.8
1.0
1.5

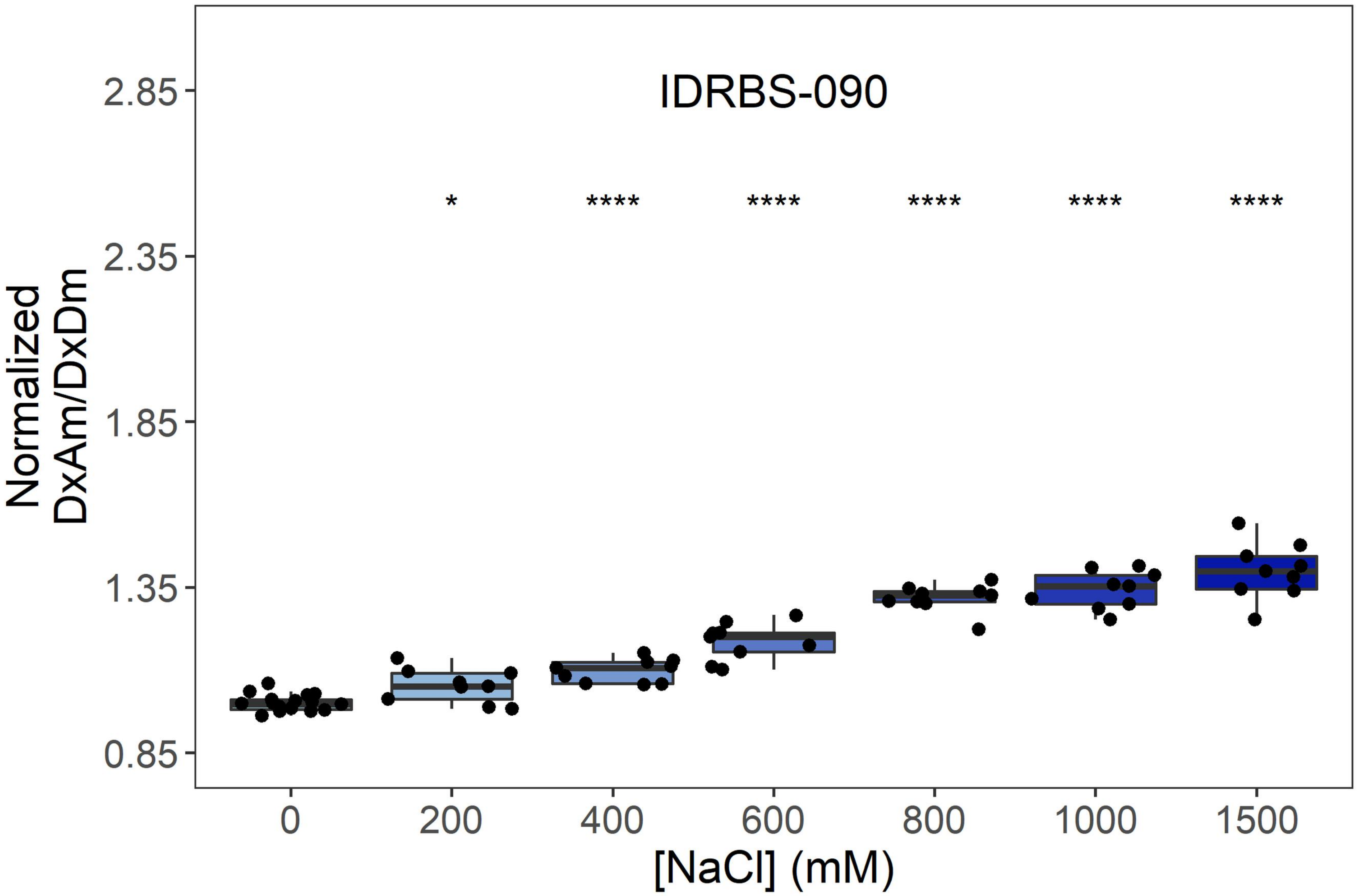
IDRBS-090
Normalized DxAm/DxDm
2.85
2.35
1.85
1.35
0.85
*
****
****
****
****
****
0
200
400
600
800
1000
1500
[NaCl] (mM)

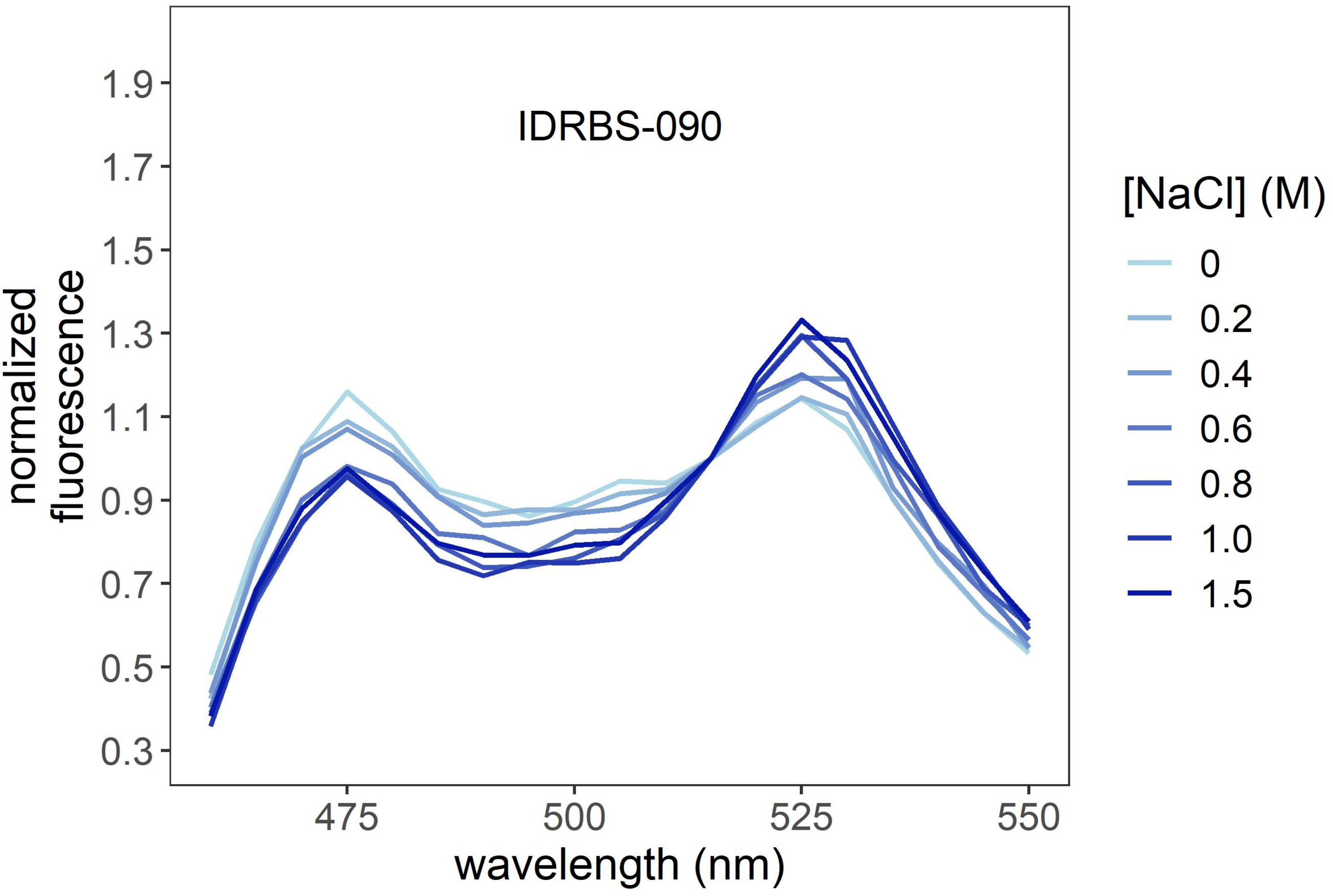
IDRBS-090
normalized fluorescence
1.9
1.7
1.5
1.3
1.1
0.9
0.7
0.5
0.3
475
500
525
550
wavelength (nm)
[NaCl] (M)
0
0.2
0.4
0.6
0.8
1.0
1.5

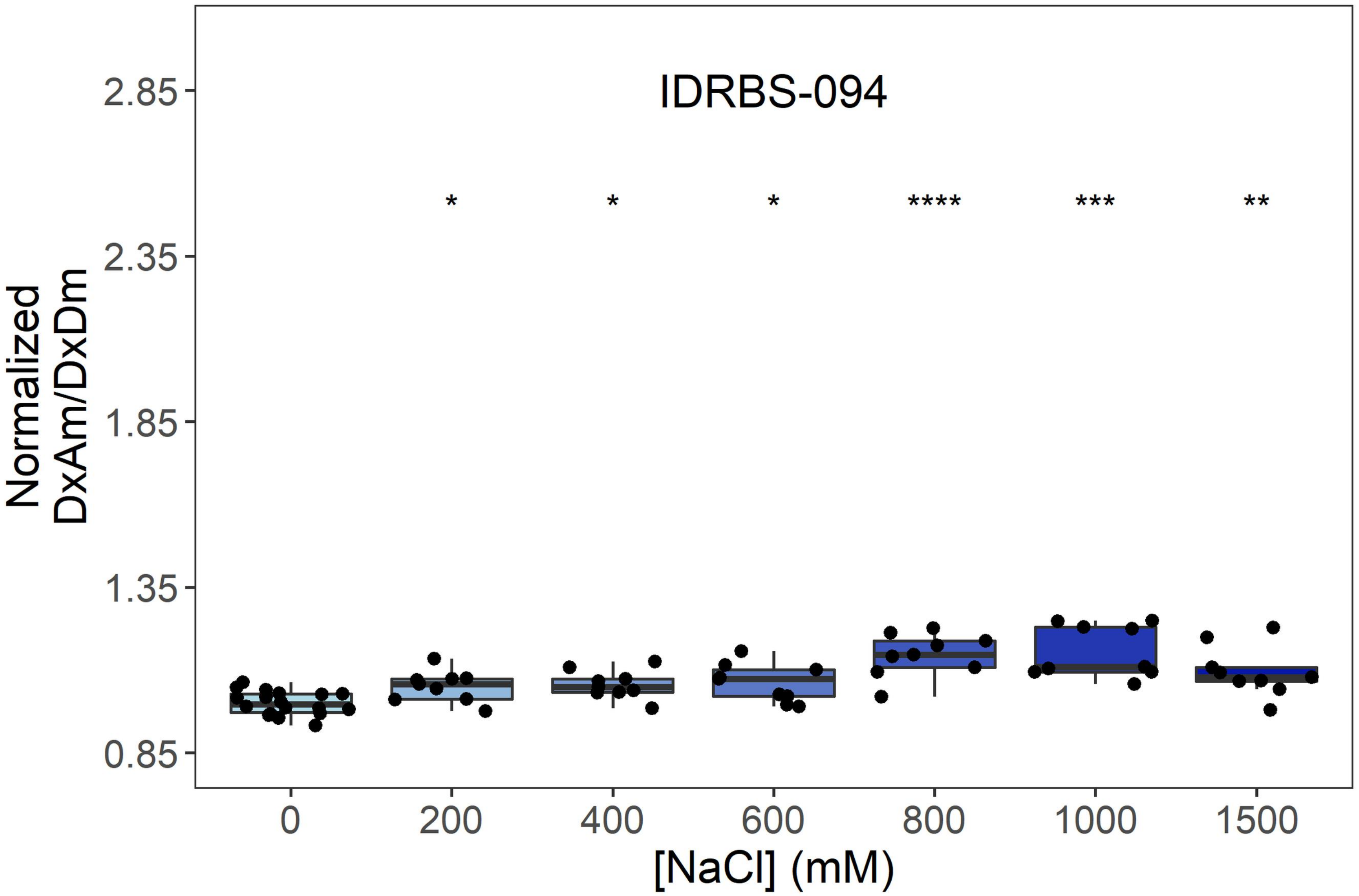
IDRBS-094
Normalized DxAm/DxDm
2.85
2.35
1.85
1.35
0.85
*
*
*
****
***
**
0
200
400
600
800
1000
1500
[NaCl] (mM)

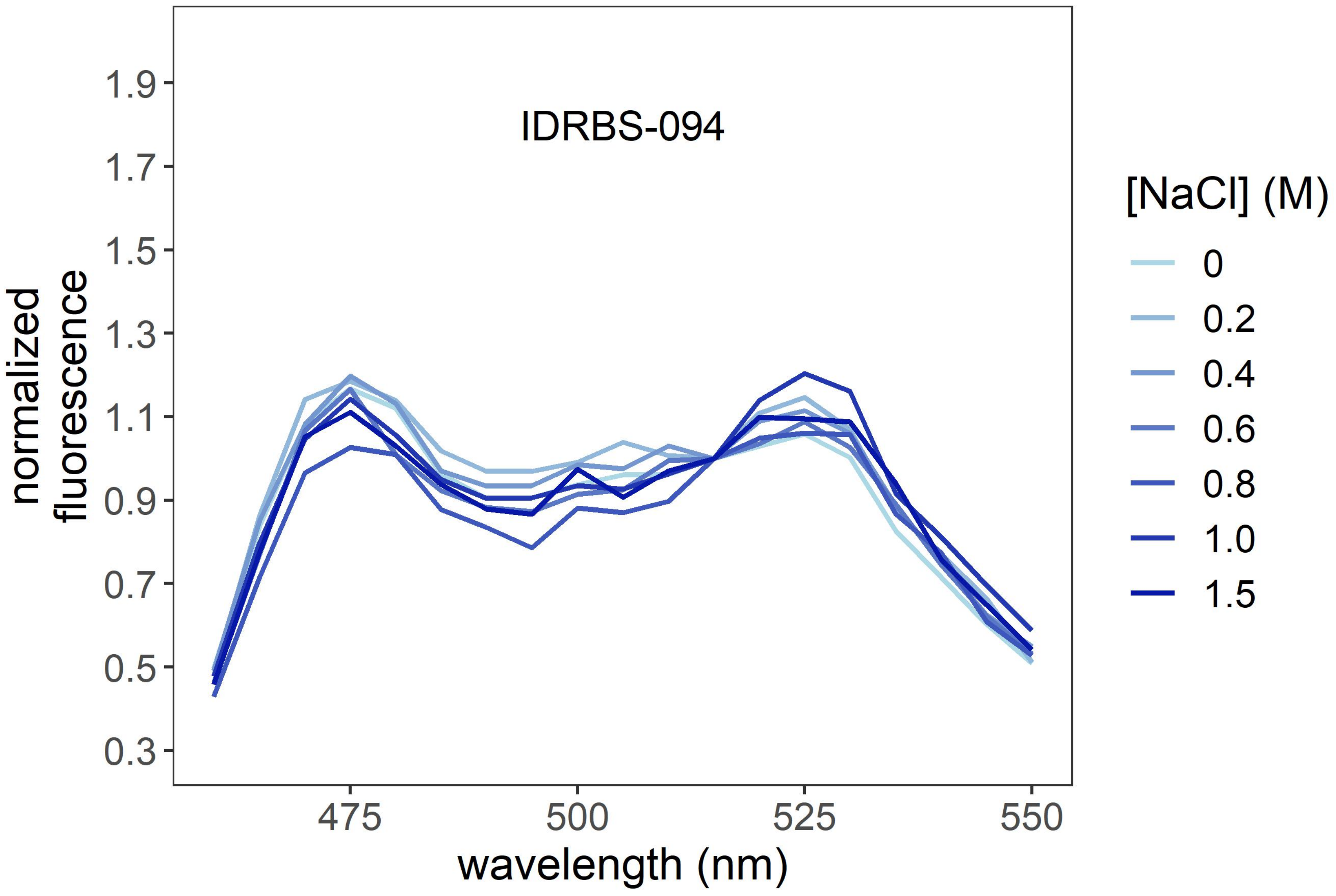
IDRBS-094
normalized fluorescence
1.9
1.7
1.5
1.3
1.1
0.9
0.7
0.5
0.3
475
500
525
550
wavelength (nm)
[NaCl] (M)
0
0.2
0.4
0.6
0.8
1.0
1.5

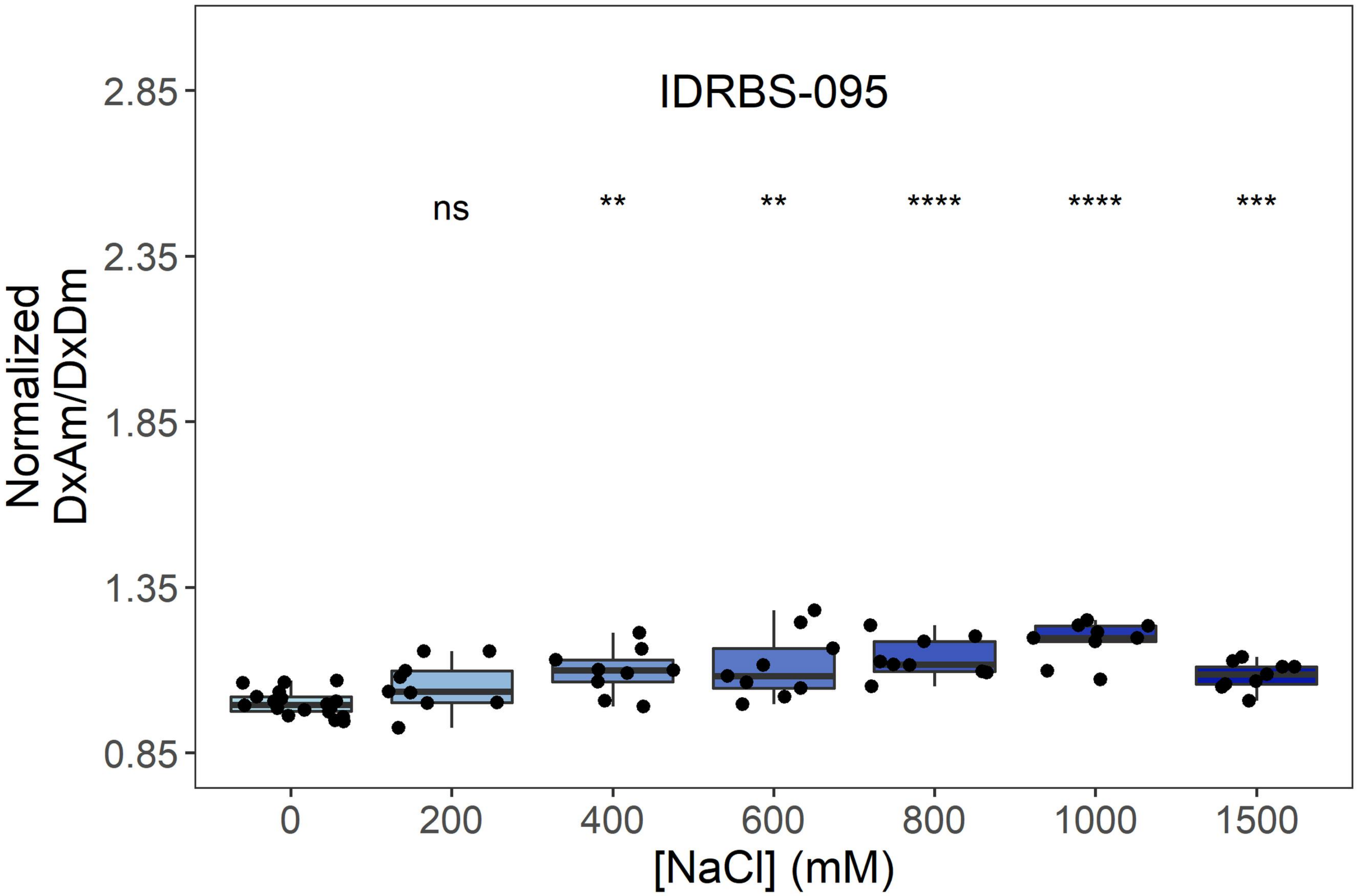

IDRBS-095
ns
**
**
****
****
***
Normalized
DxAm/DxDm
2.85
2.35
1.85
1.35
0.85
0
200
400
600
800
1000
1500
[NaCl] (mM)


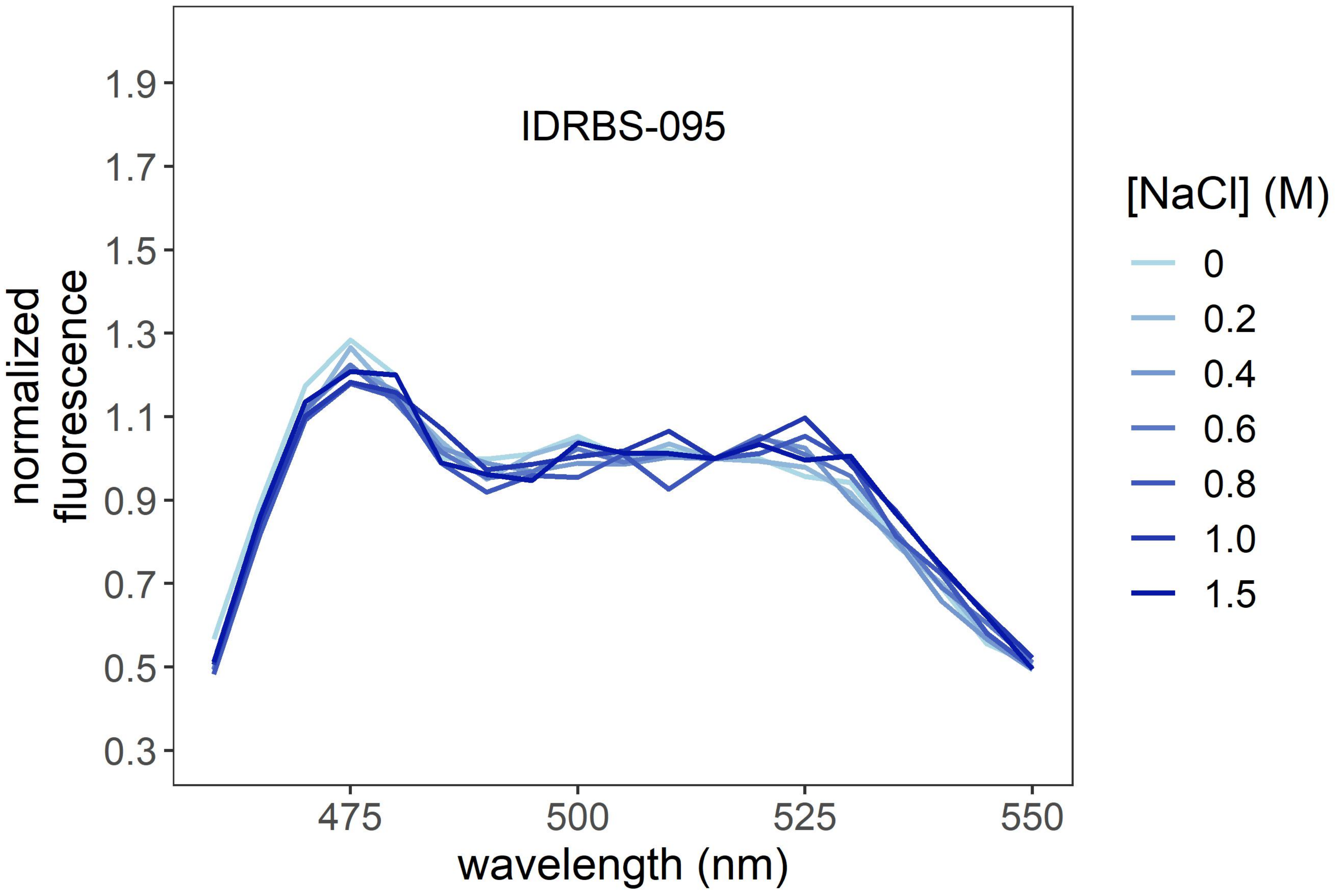

IDRBS-095
normalized
fluorescence
1.9
1.7
1.5
1.3
1.1
0.9
0.7
0.5
0.3
475
500
525
550
wavelength (nm)
[NaCl] (M)
0
0.2
0.4
0.6
0.8
1.0
1.5

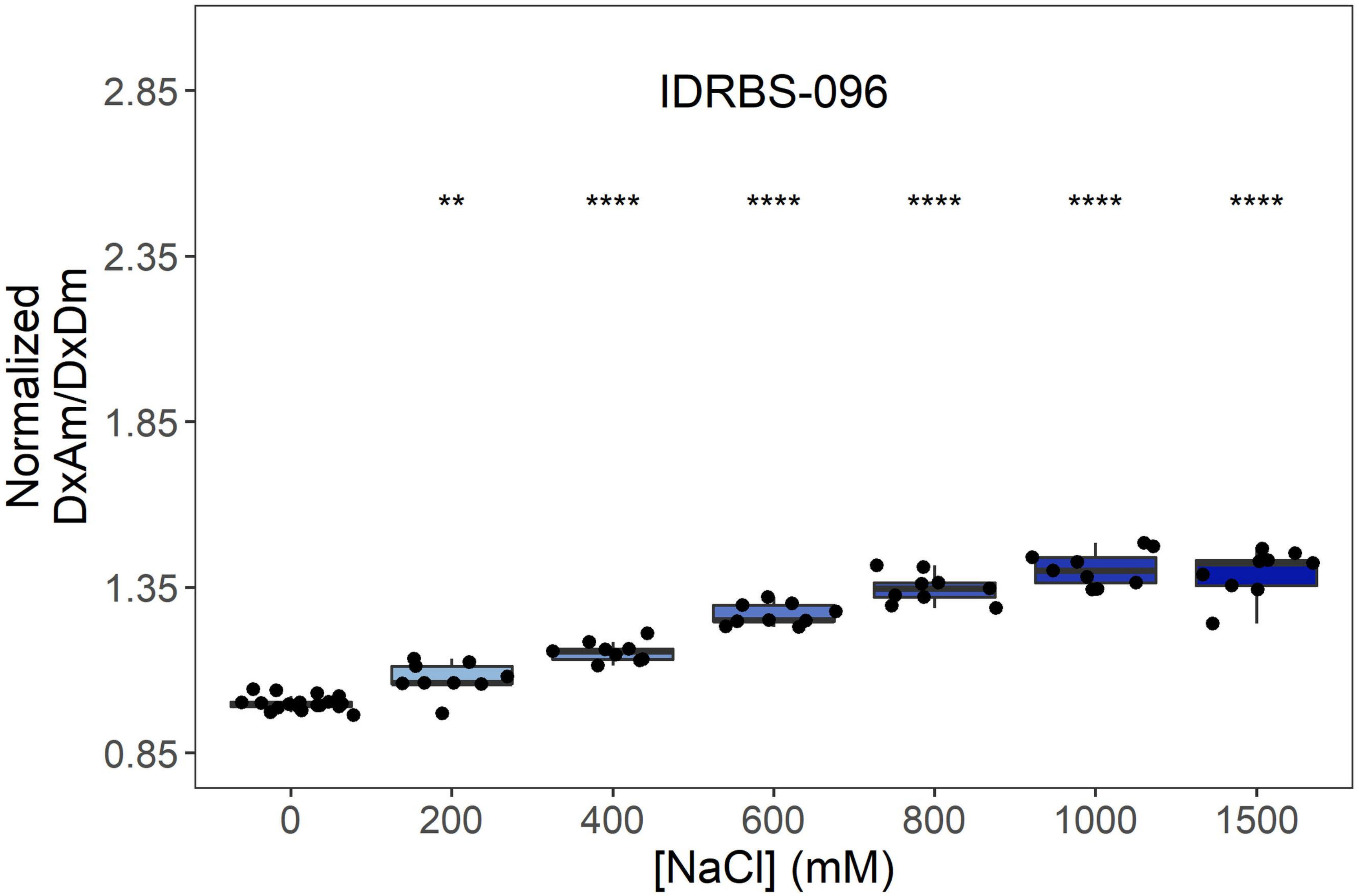
IDRBS-096
**
****
****
****
****
****
Normalized
DxAm/DxDm
2.85
2.35
1.85
1.35
0.85
0
200
400
600
800
1000
1500
[NaCl] (mM)

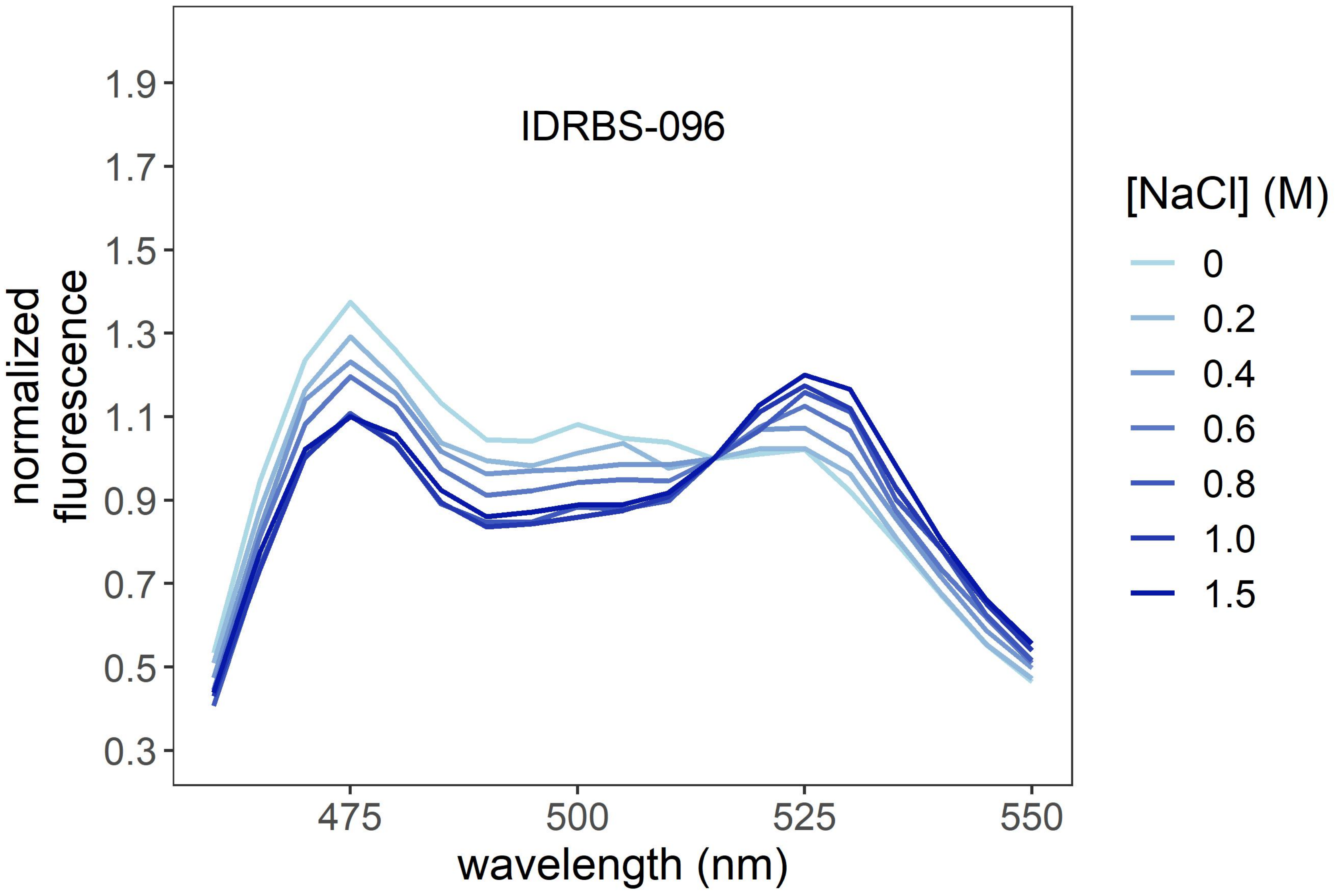
IDRBS-096
normalized
fluorescence
1.9
1.7
1.5
1.3
1.1
0.9
0.7
0.5
0.3
475
500
525
550
wavelength (nm)
[NaCl] (M)
0
0.2
0.4
0.6
0.8
1.0
1.5

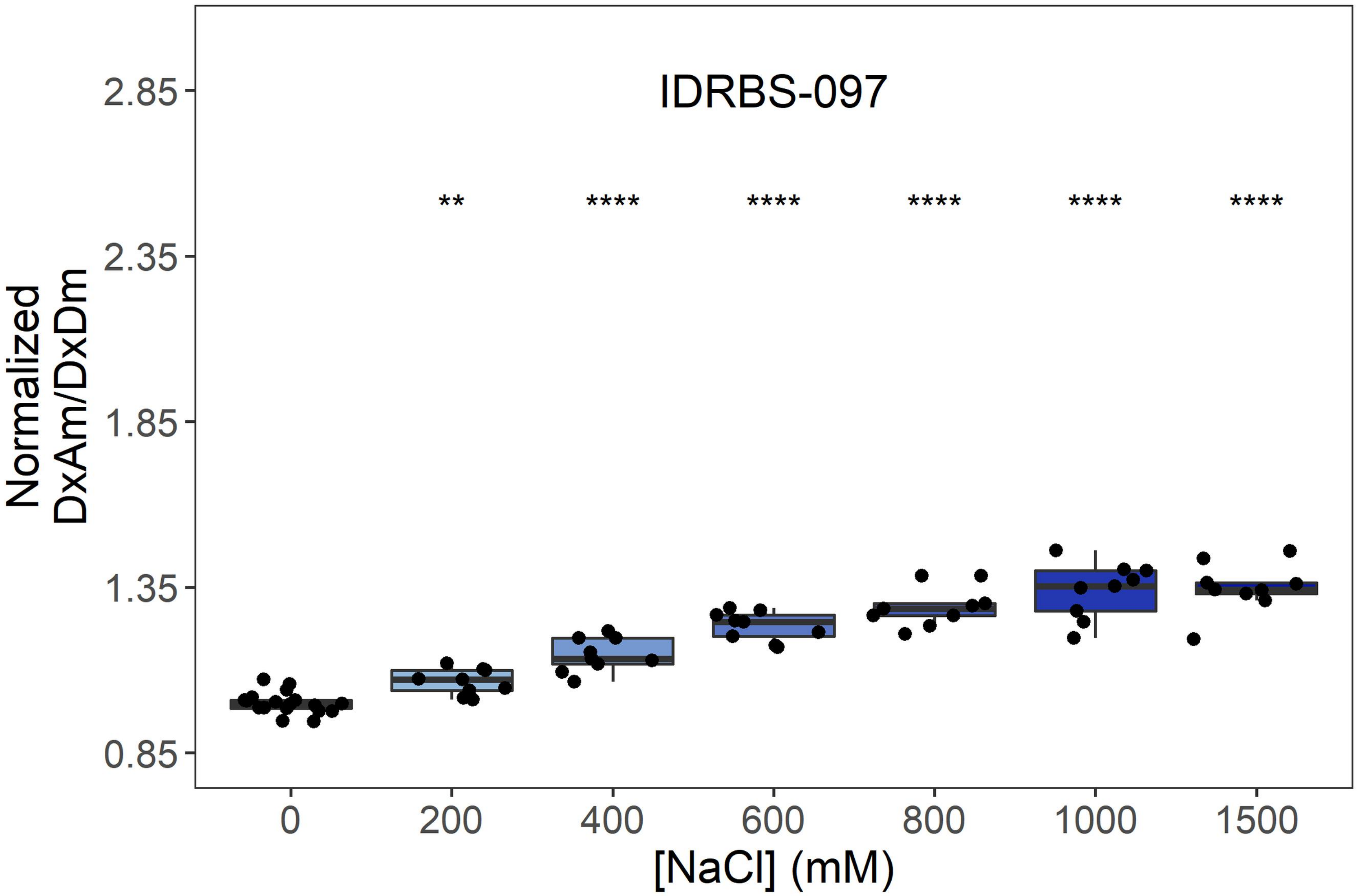
IDRBS-097
**
****
****
****
****
****
Normalized
DxAm/DxDm
2.85
2.35
1.85
1.35
0.85
0
200
400
600
800
1000
1500
[NaCl] (mM)

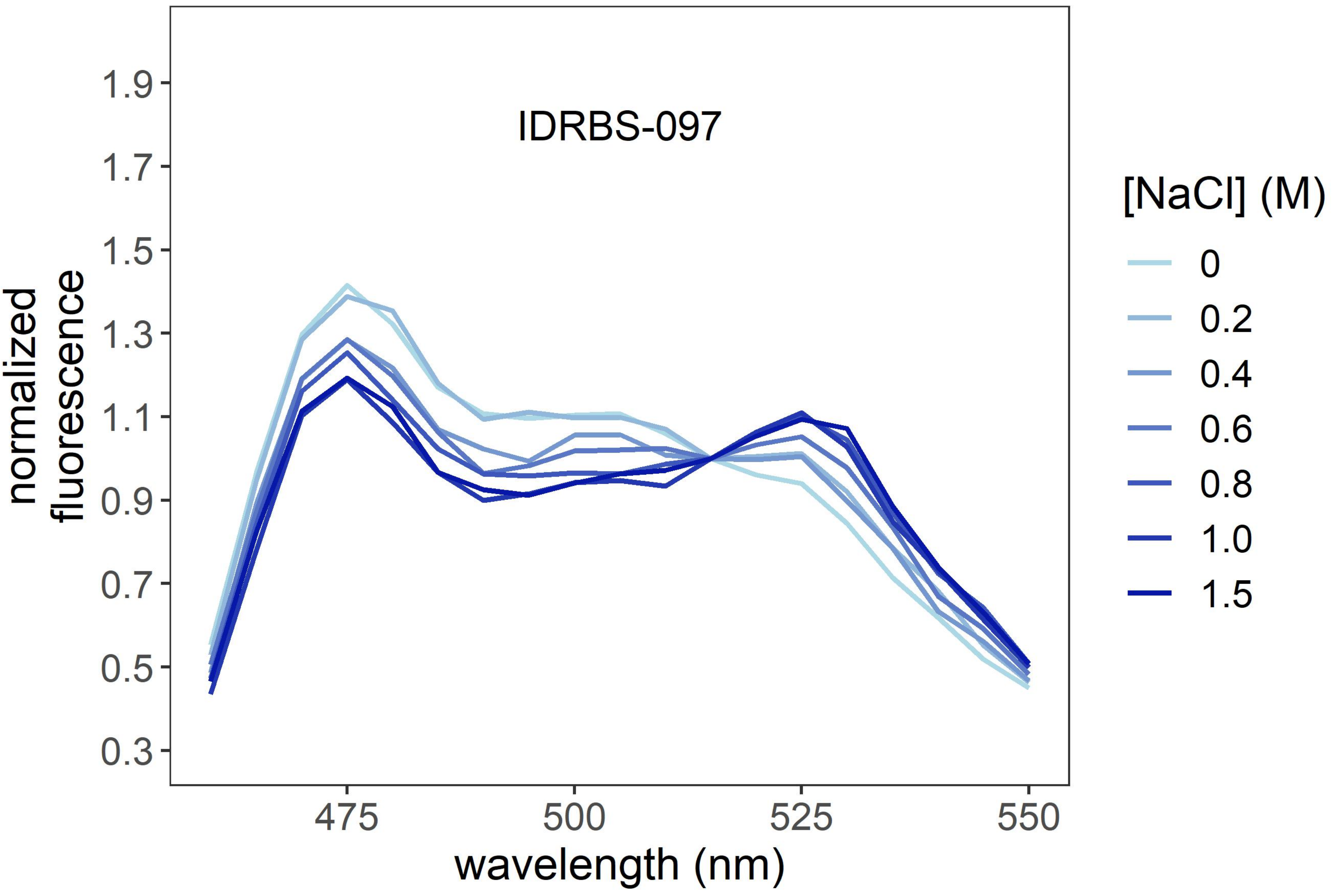
IDRBS-097
normalized
fluorescence
1.9
1.7
1.5
1.3
1.1
0.9
0.7
0.5
0.3
475
500
525
550
wavelength (nm)
[NaCl] (M)
0
0.2
0.4
0.6
0.8
1.0
1.5

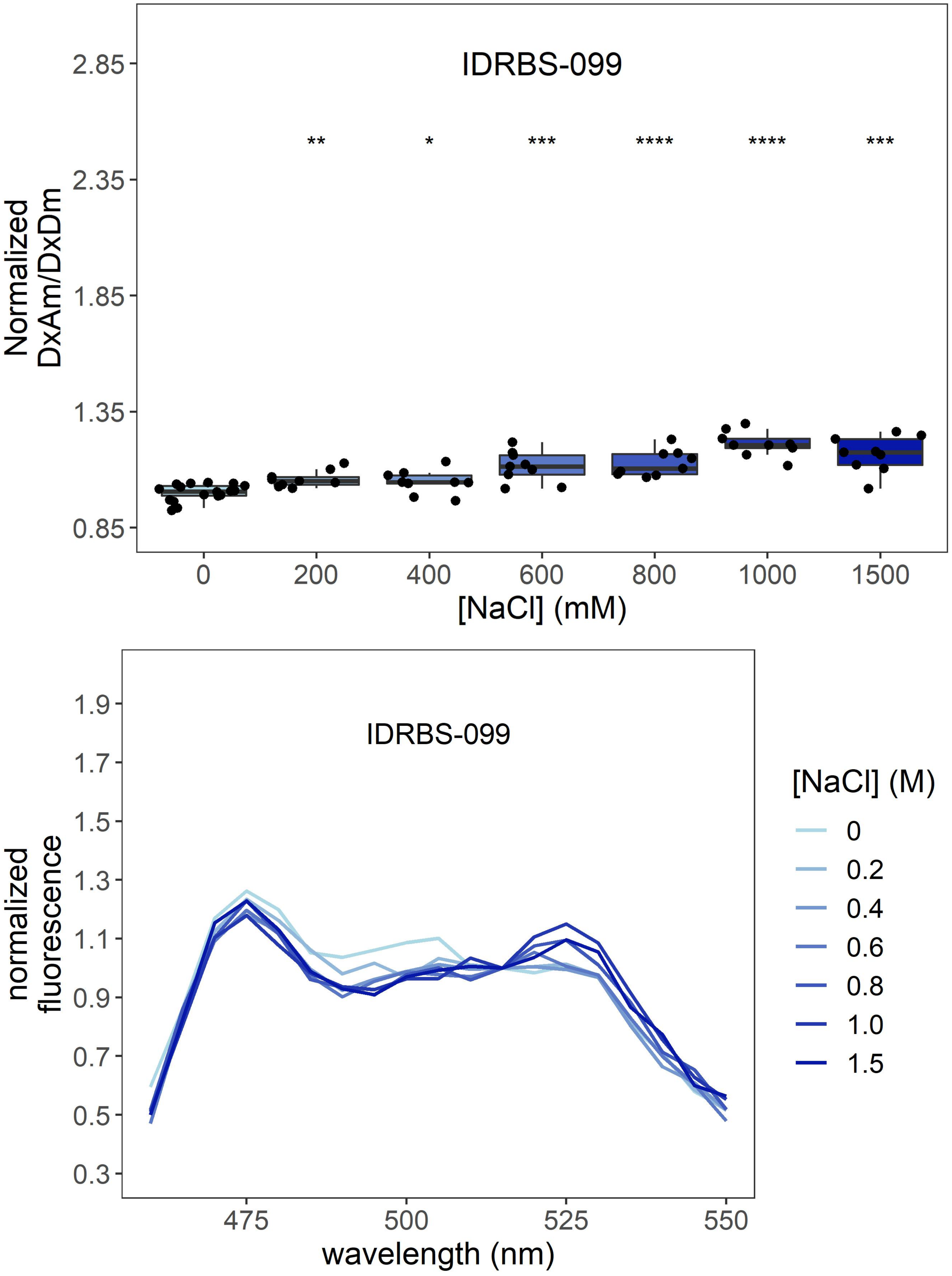
IDRBS-099
Normalized DxAm/DxDm
[NaCl] (mM)
0
200
400
600
800
1000
1500
**
*
***
****
****
***
0.85
1.35
1.85
2.35
2.85
IDRBS-099
normalized fluorescence
wavelength (nm)
475
500
525
550
0.3
0.5
0.7
0.9
1.1
1.3
1.5
1.7
1.9
[NaCl] (M)
0
0.2
0.4
0.6
0.8
1.0
1.5

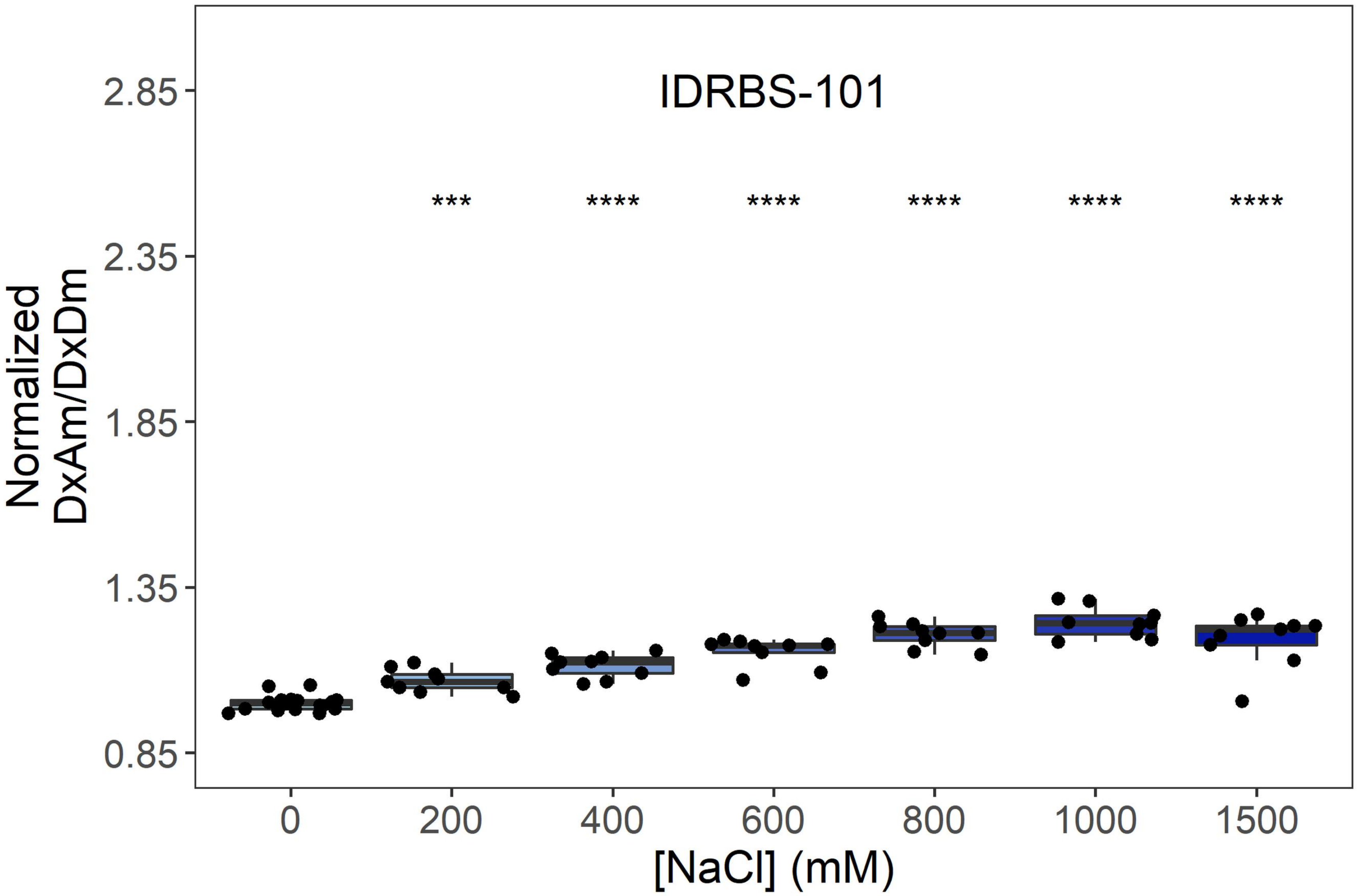
IDRBS-101
***
****
****
****
****
****
Normalized DxAm/DxDm
2.85
2.35
1.85
1.35
0.85
0
200
400
600
800
1000
1500
[NaCl] (mM)

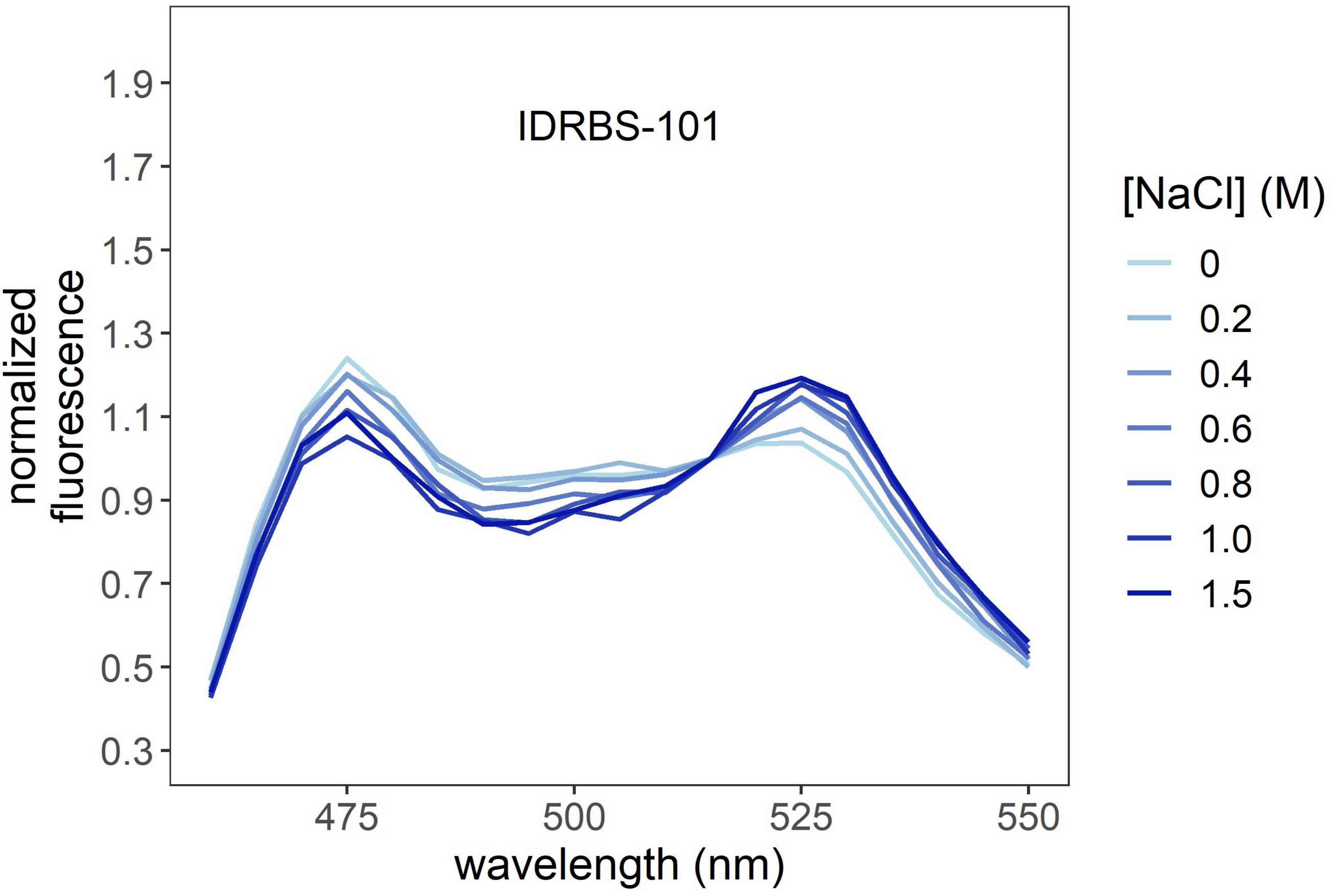
IDRBS-101
normalized fluorescence
1.9
1.7
1.5
1.3
1.1
0.9
0.7
0.5
0.3
475
500
525
550
wavelength (nm)
[NaCl] (M)
0
0.2
0.4
0.6
0.8
1.0
1.5

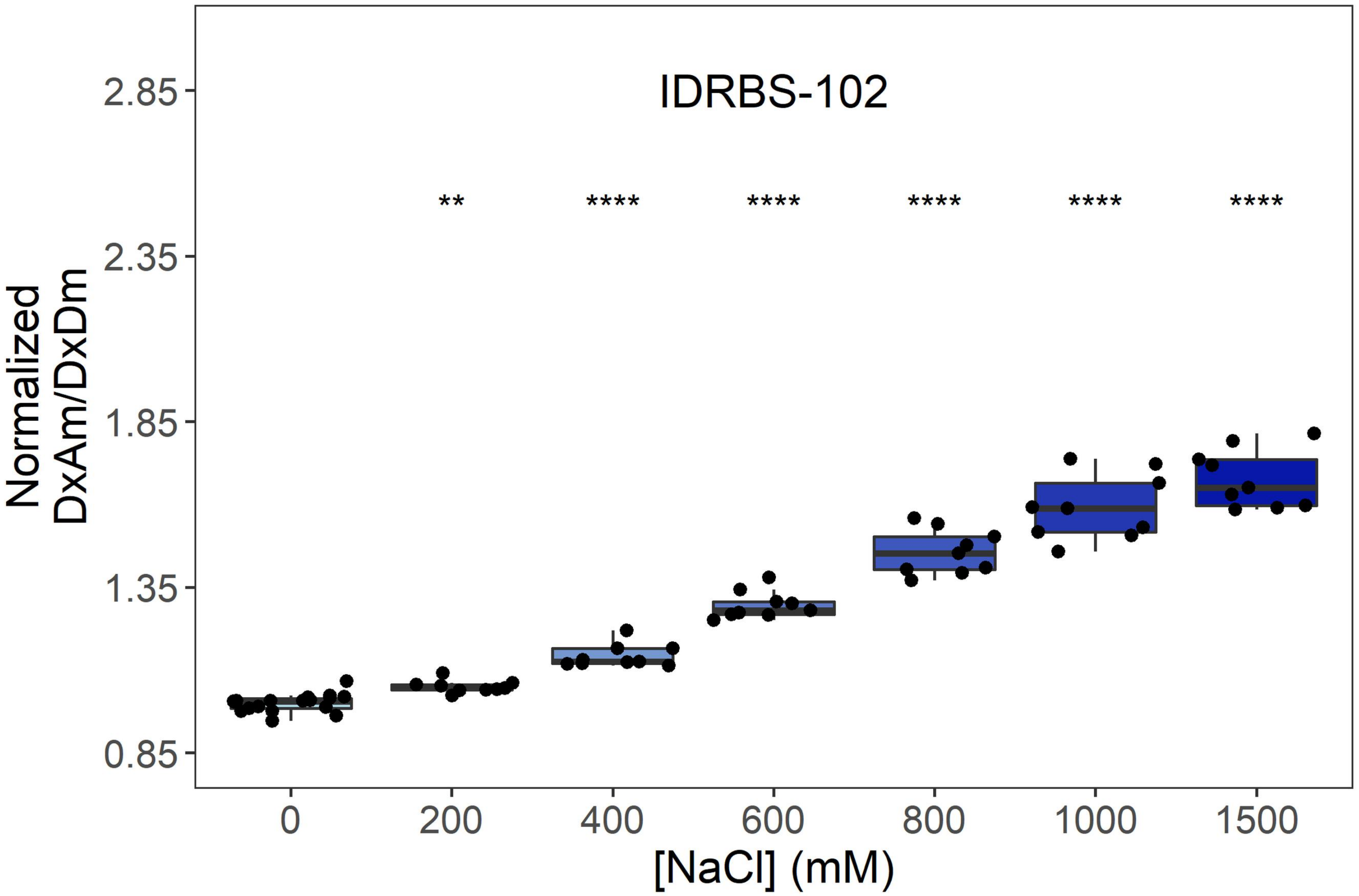
IDRBS-102
**
****
****
****
****
****
Normalized
DxAm/DxDm
2.85
2.35
1.85
1.35
0.85
0
200
400
600
800
1000
1500
[NaCl] (mM)

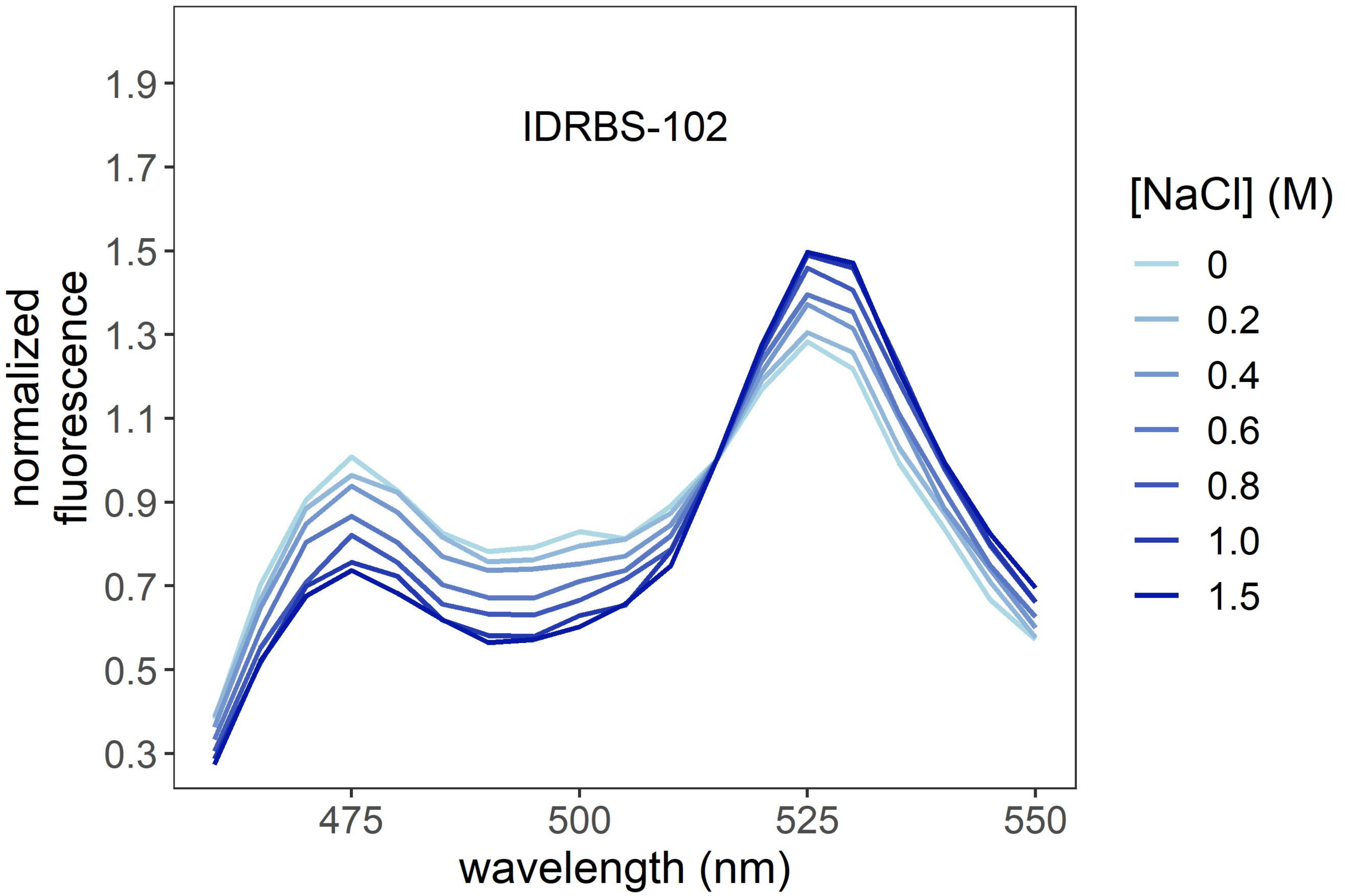
IDRBS-102
normalized
fluorescence
1.9
1.7
1.5
1.3
1.1
0.9
0.7
0.5
0.3
475
500
525
550
wavelength (nm)
[NaCl] (M)
0
0.2
0.4
0.6
0.8
1.0
1.5

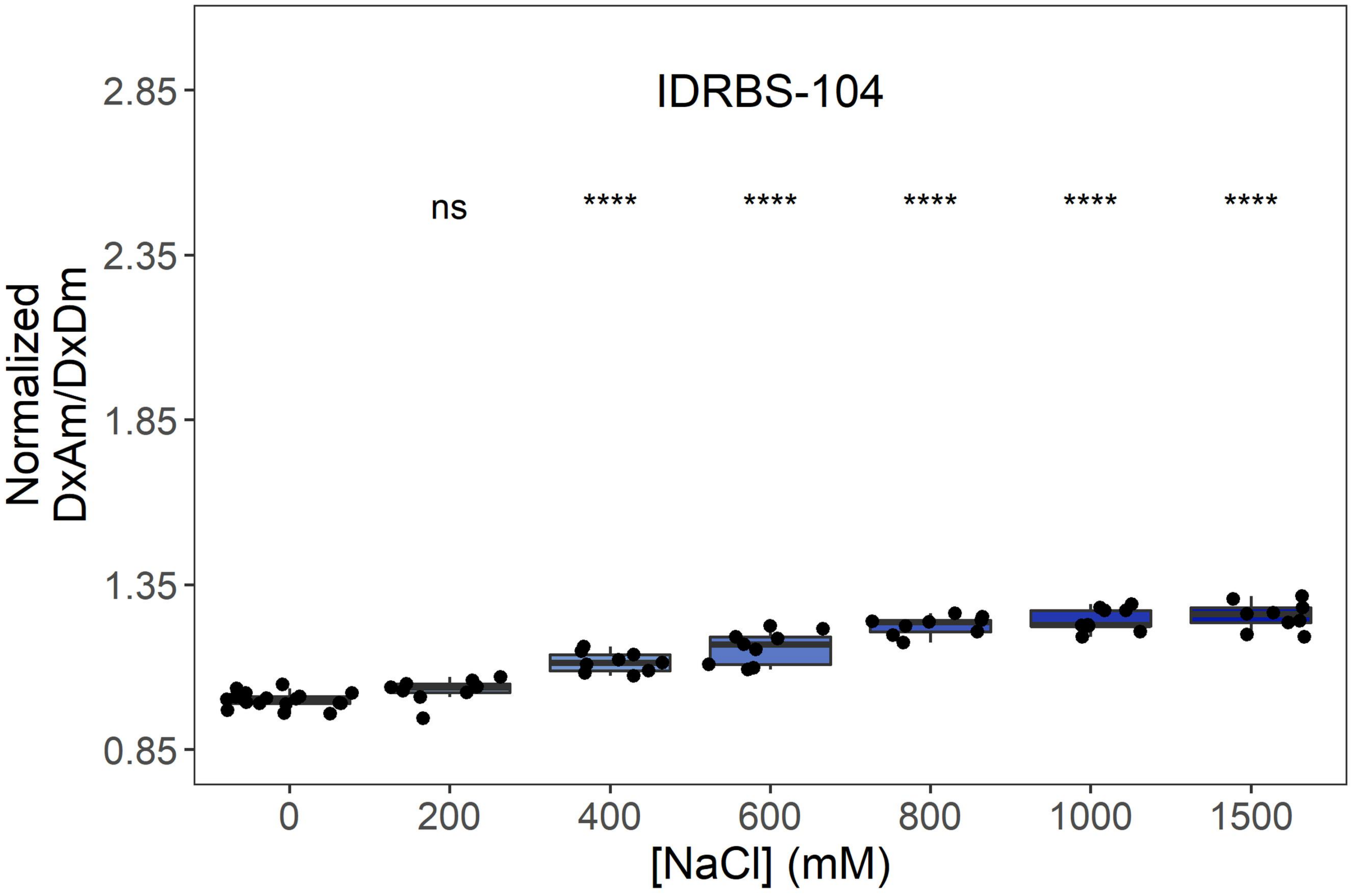
IDRBS-104
ns
****
****
****
****
****
2.85
2.35
1.85
1.35
0.85
Normalized
DxAm/DxDm
0
200
400
600
800
1000
1500
[NaCl] (mM)

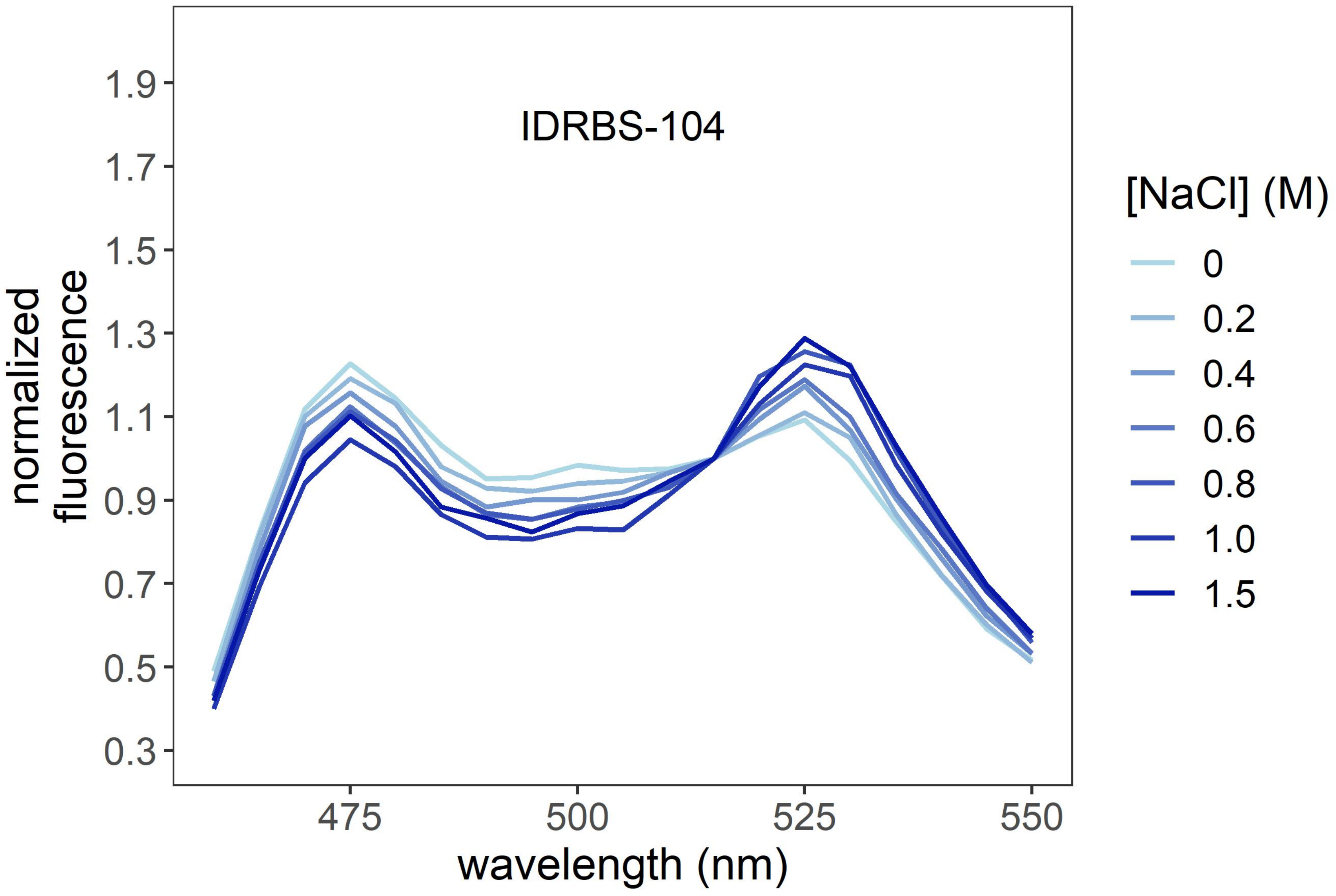
IDRBS-104
1.9
1.7
1.5
1.3
1.1
0.9
0.7
0.5
0.3
normalized
fluorescence
475
500
525
550
wavelength (nm)
[NaCl] (M)
0
0.2
0.4
0.6
0.8
1.0
1.5

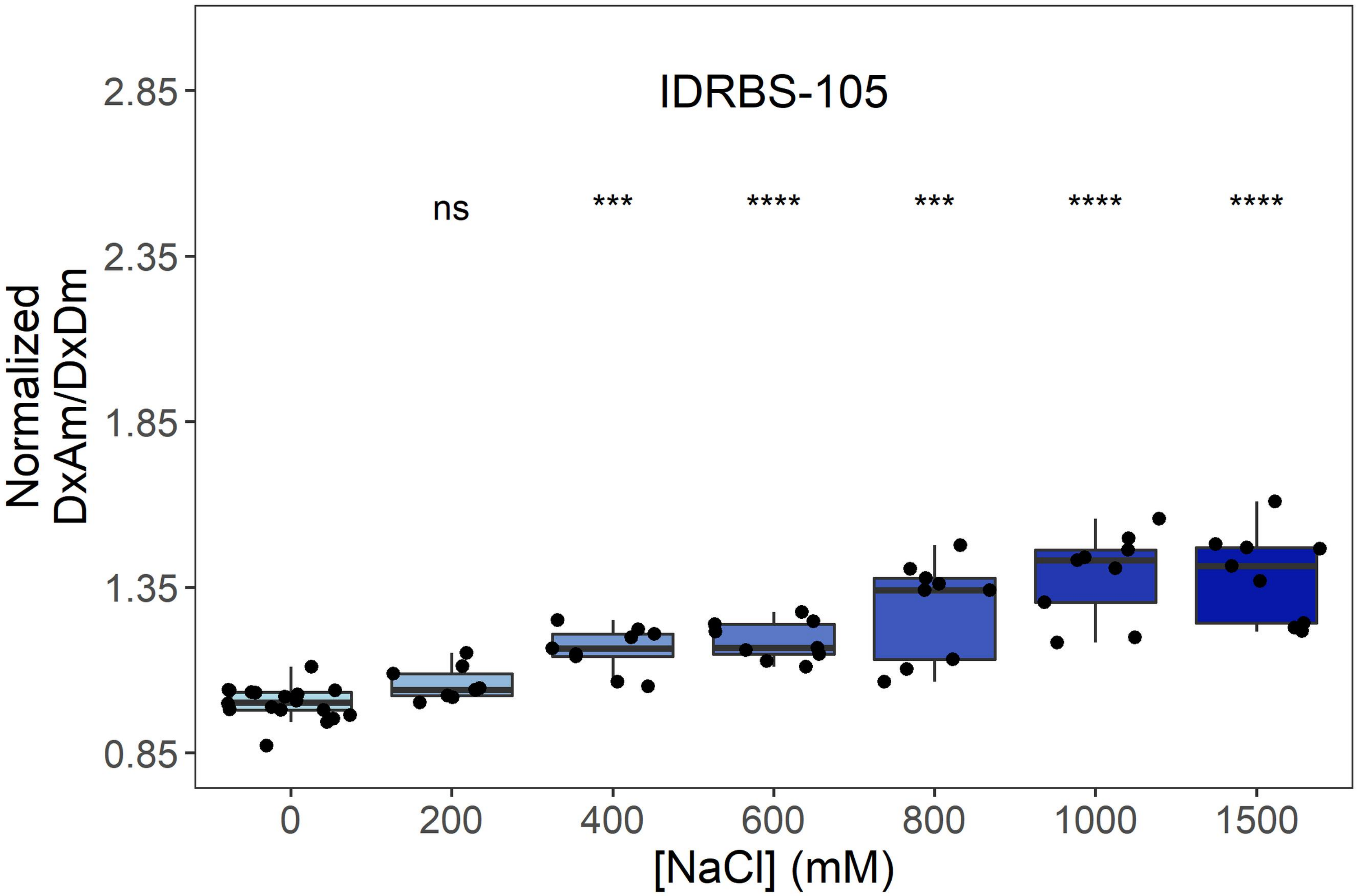
IDRBS-105
ns
***
****
***
****
****
Normalized
DxAm/DxDm
2.85
2.35
1.85
1.35
0.85
0
200
400
600
800
1000
1500
[NaCl] (mM)

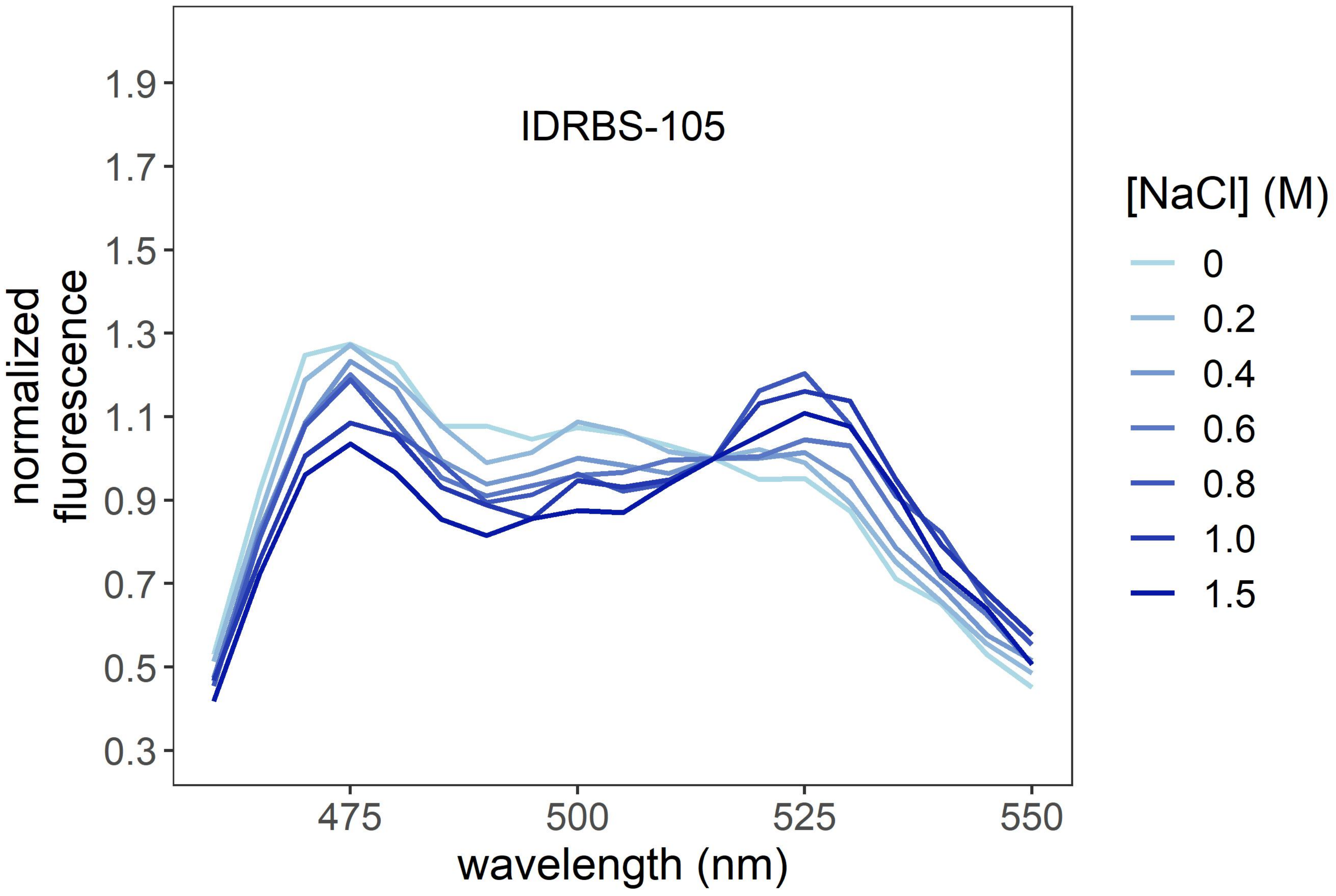
IDRBS-105
normalized
fluorescence
1.9
1.7
1.5
1.3
1.1
0.9
0.7
0.5
0.3
475
500
525
550
wavelength (nm)
[NaCl] (M)
0
0.2
0.4
0.6
0.8
1.0
1.5

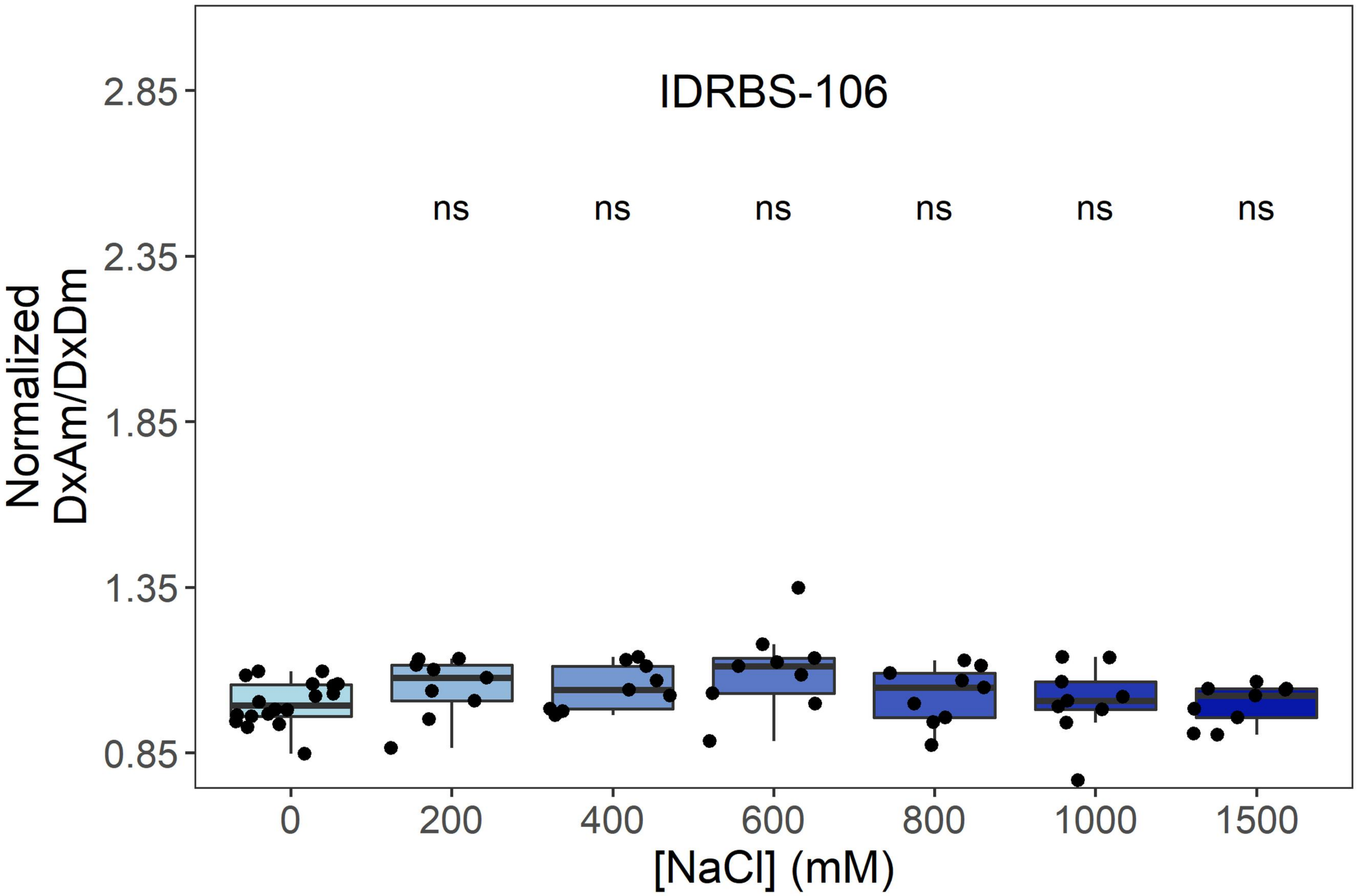
IDRBS-106
ns
ns
ns
ns
ns
ns
Normalized DxAm/DxDm
2.85
2.35
1.85
1.35
0.85
0
200
400
600
800
1000
1500
[NaCl] (mM)

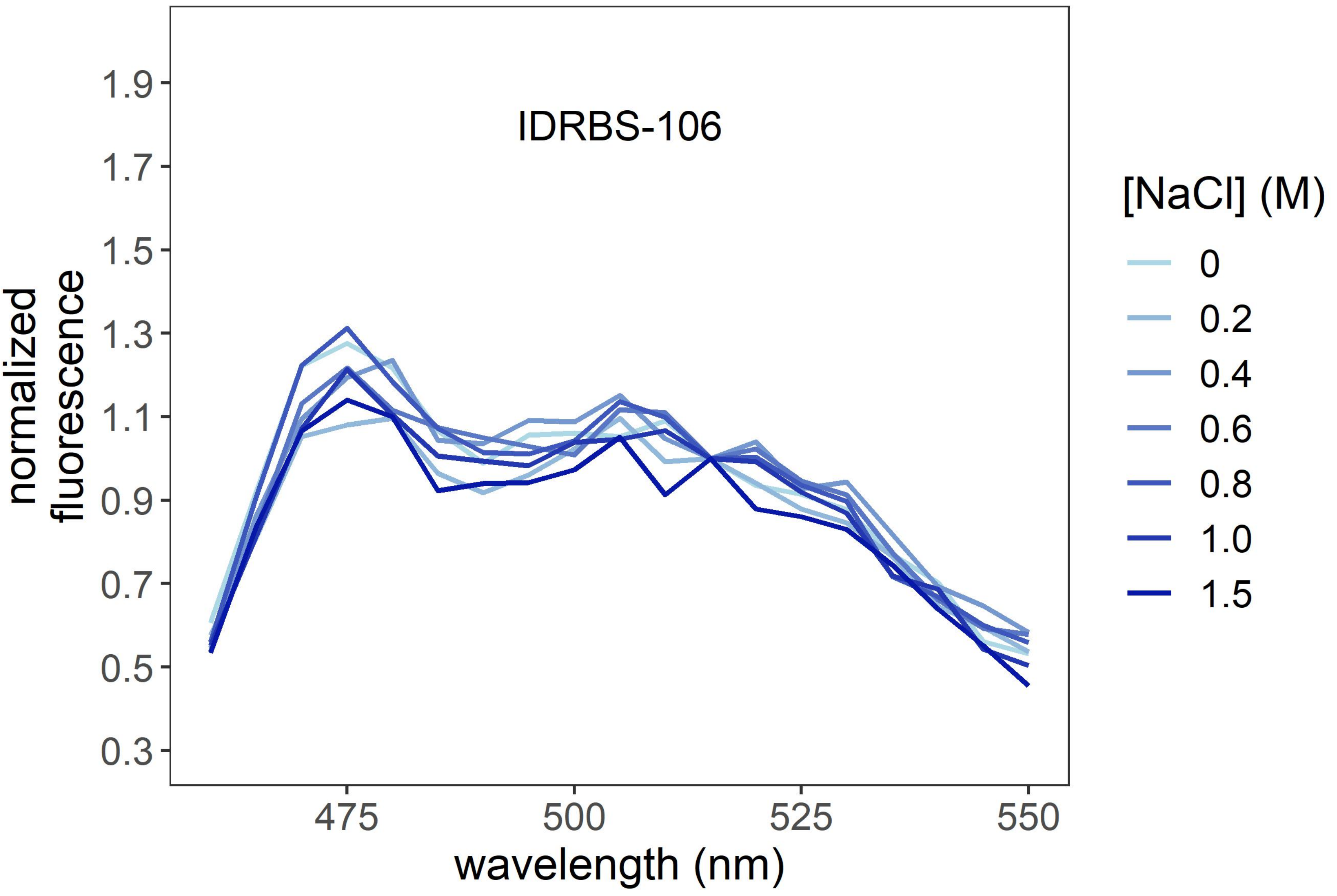
IDRBS-106
normalized fluorescence
1.9
1.7
1.5
1.3
1.1
0.9
0.7
0.5
0.3
475
500
525
550
wavelength (nm)
[NaCl] (M)
0
0.2
0.4
0.6
0.8
1.0
1.5

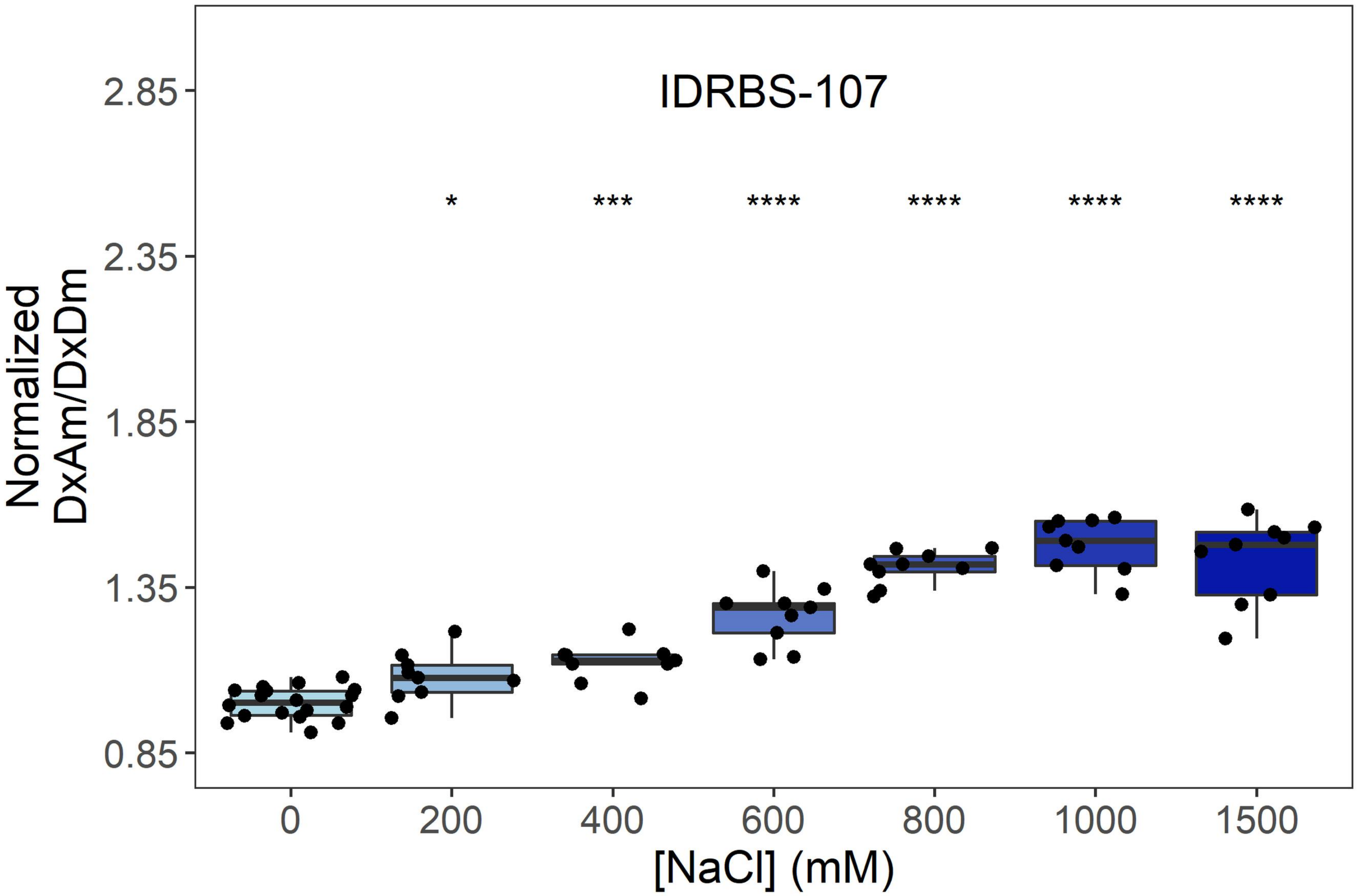
IDRBS-107
*
***
****
****
****
****
Normalized DxAm/DxDm
2.85
2.35
1.85
1.35
0.85
0
200
400
600
800
1000
1500
[NaCl] (mM)

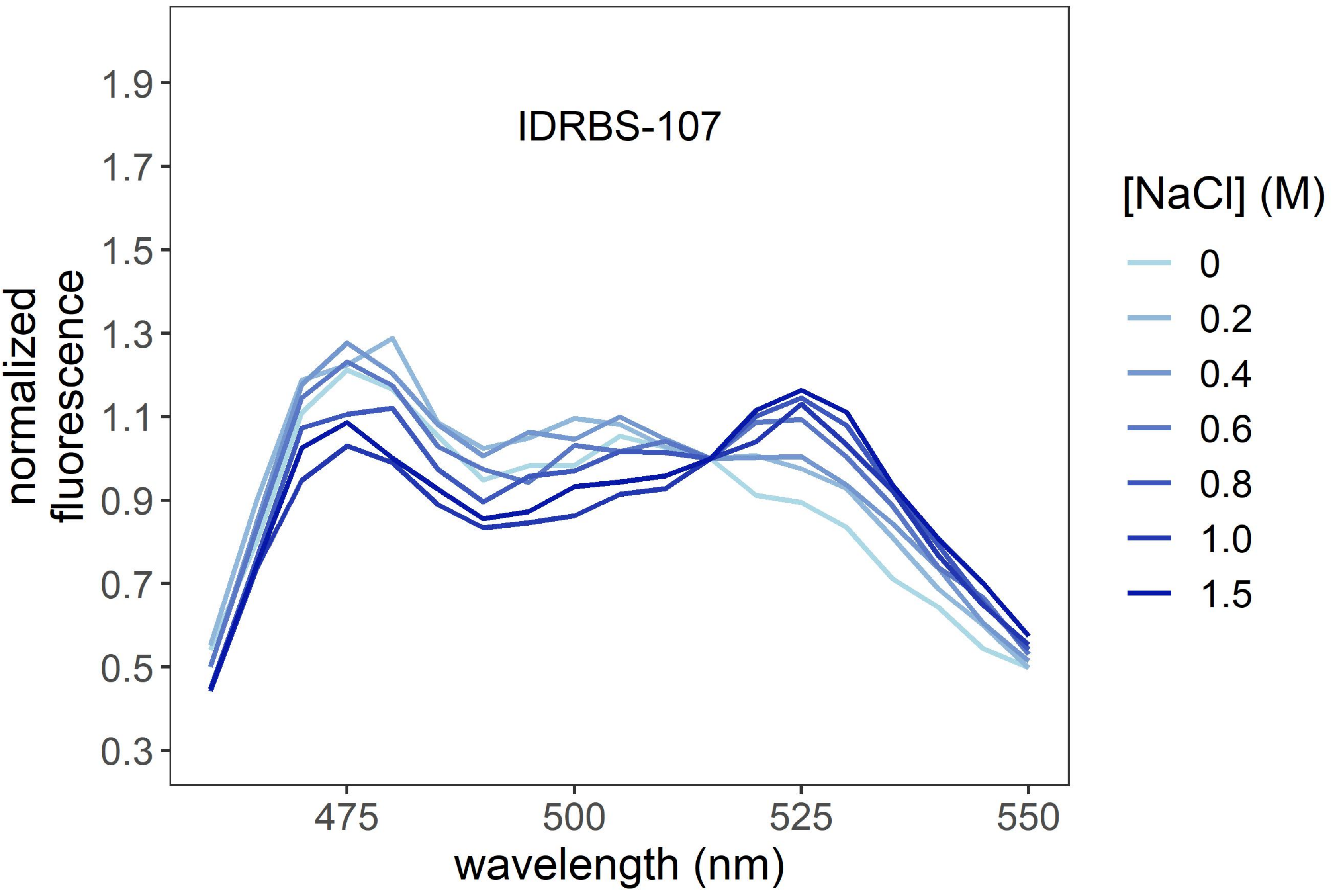
IDRBS-107
normalized fluorescence
1.9
1.7
1.5
1.3
1.1
0.9
0.7
0.5
0.3
475
500
525
550
wavelength (nm)
[NaCl] (M)
0
0.2
0.4
0.6
0.8
1.0
1.5

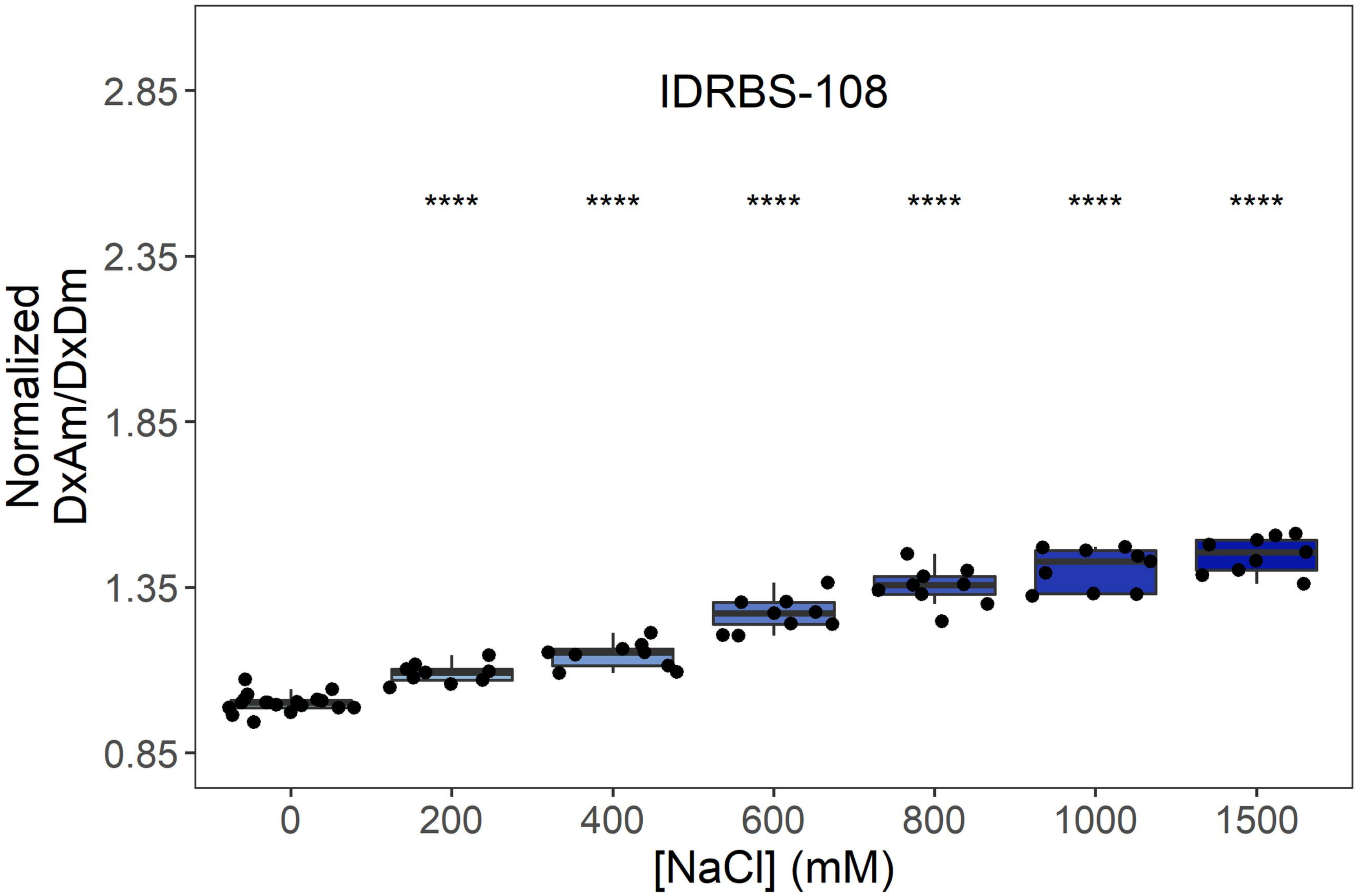

IDRBS-108
Normalized DxAm/DxDm
2.85
2.35
1.85
1.35
0.85
****
0
200
400
600
800
1000
1500
[NaCl] (mM)


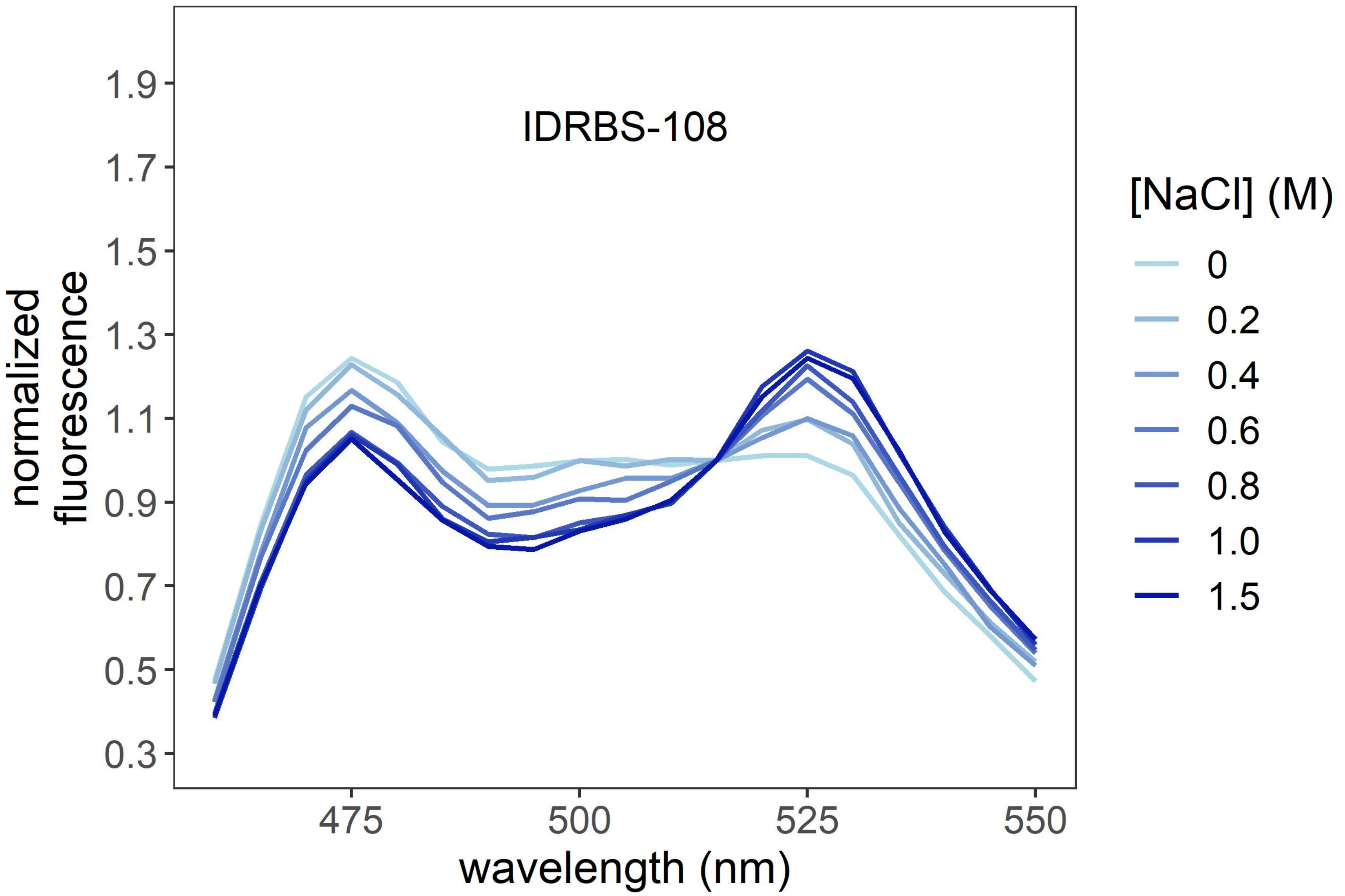

IDRBS-108
normalized fluorescence
1.9
1.7
1.5
1.3
1.1
0.9
0.7
0.5
0.3
475
500
525
550
wavelength (nm)
[NaCl] (M)
0
0.2
0.4
0.6
0.8
1.0
1.5

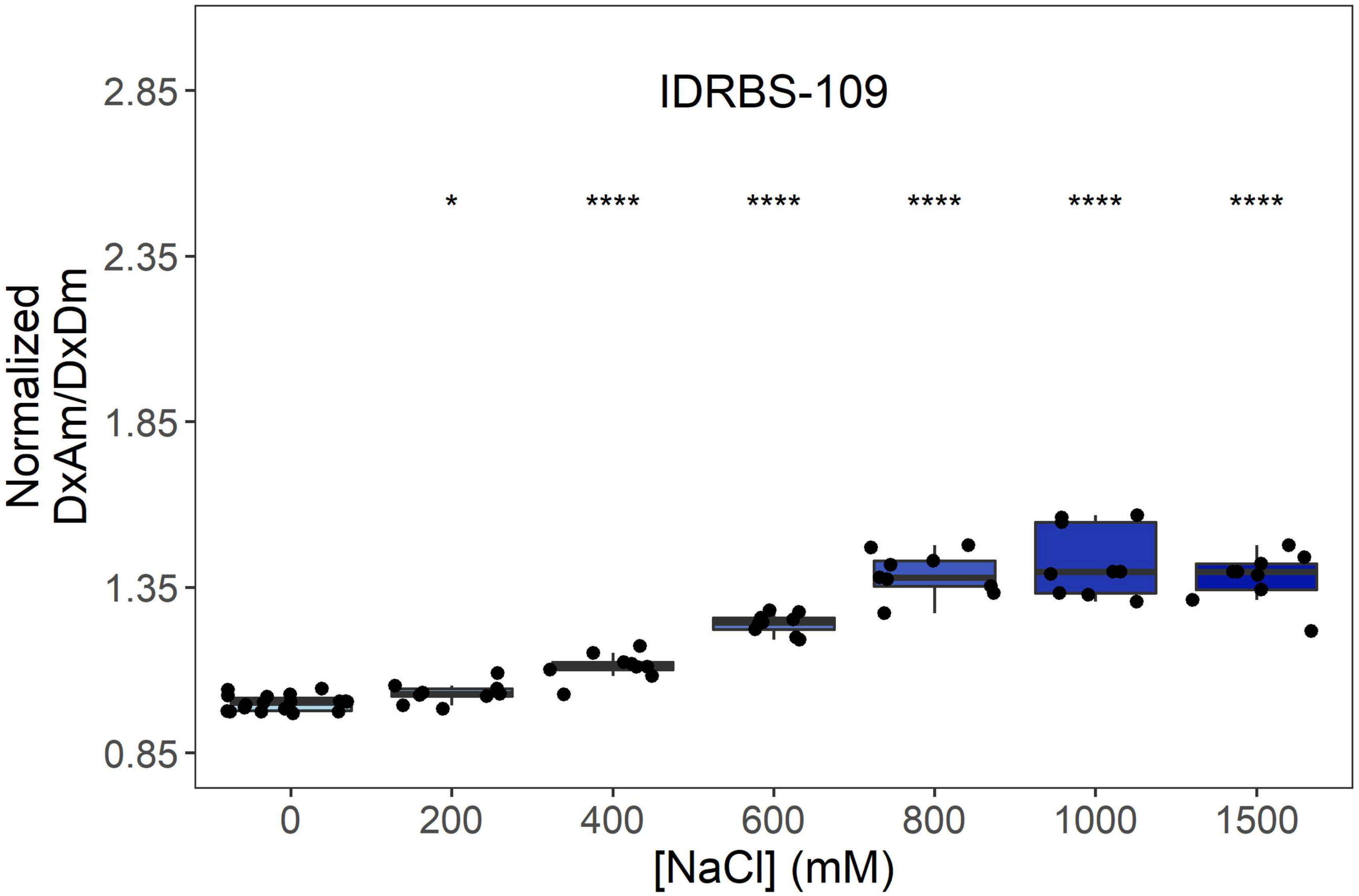
IDRBS-109
Normalized DxAm/DxDm
2.85
2.35
1.85
1.35
0.85
*
****
****
****
****
****
0
200
400
600
800
1000
1500
[NaCl] (mM)
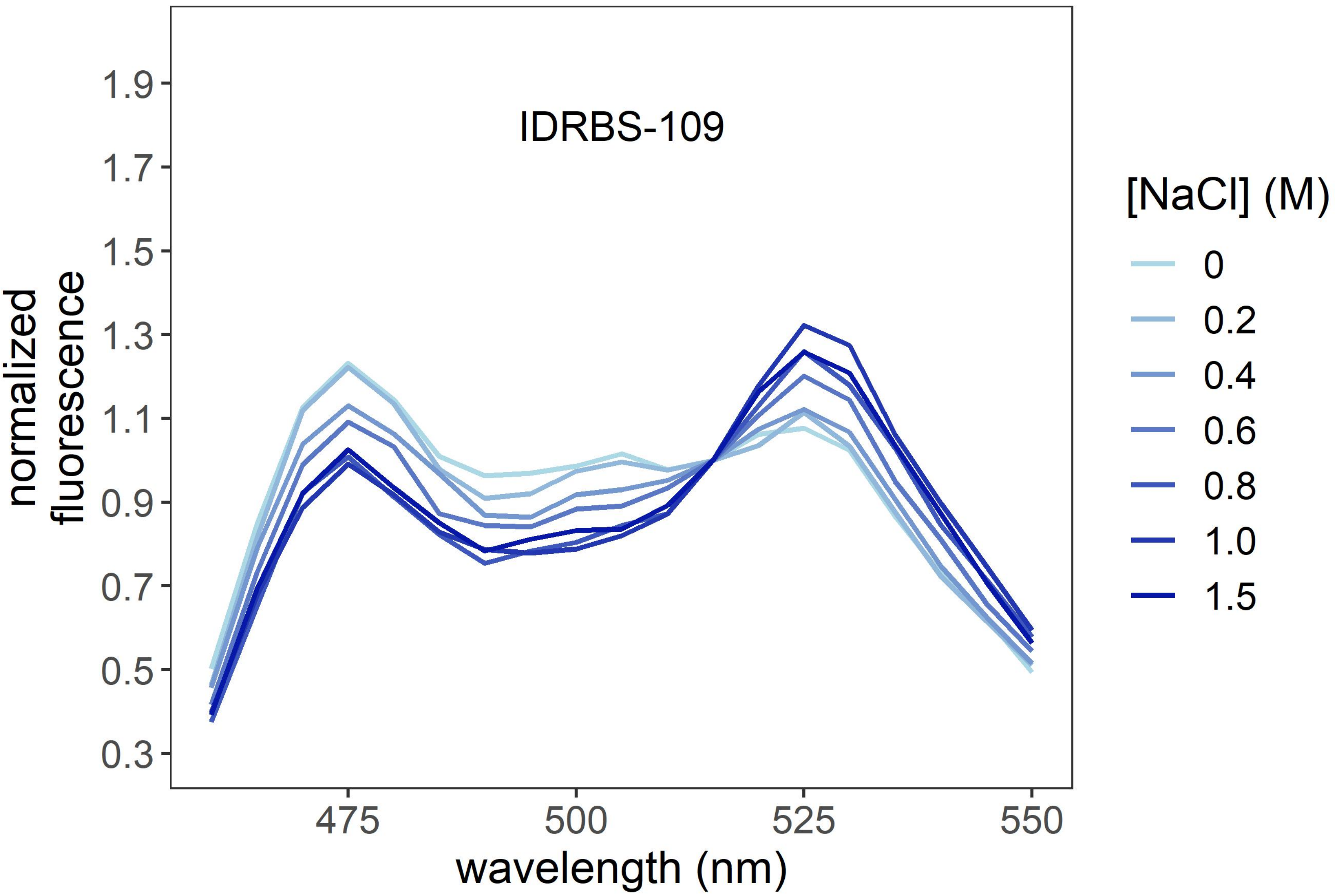
IDRBS-109
normalized fluorescence
1.9
1.7
1.5
1.3
1.1
0.9
0.7
0.5
0.3
475
500
525
550
wavelength (nm)
[NaCl] (M)
0
0.2
0.4
0.6
0.8
1.0
1.5

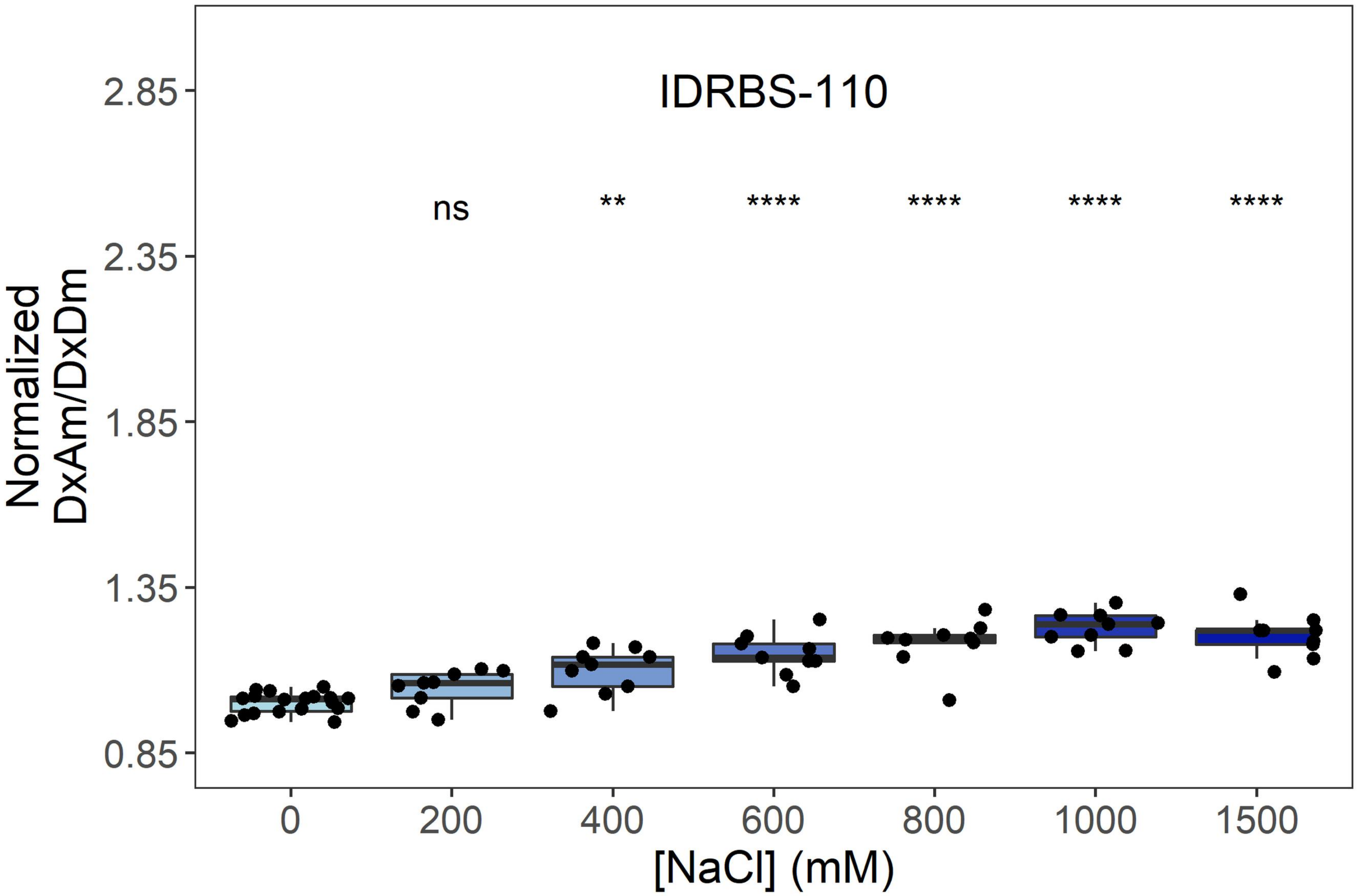
IDRBS-110
ns
**
****
****
****
****
2.85
2.35
1.85
1.35
0.85
Normalized
DxAm/DxDm
0
200
400
600
800
1000
1500
[NaCl] (mM)

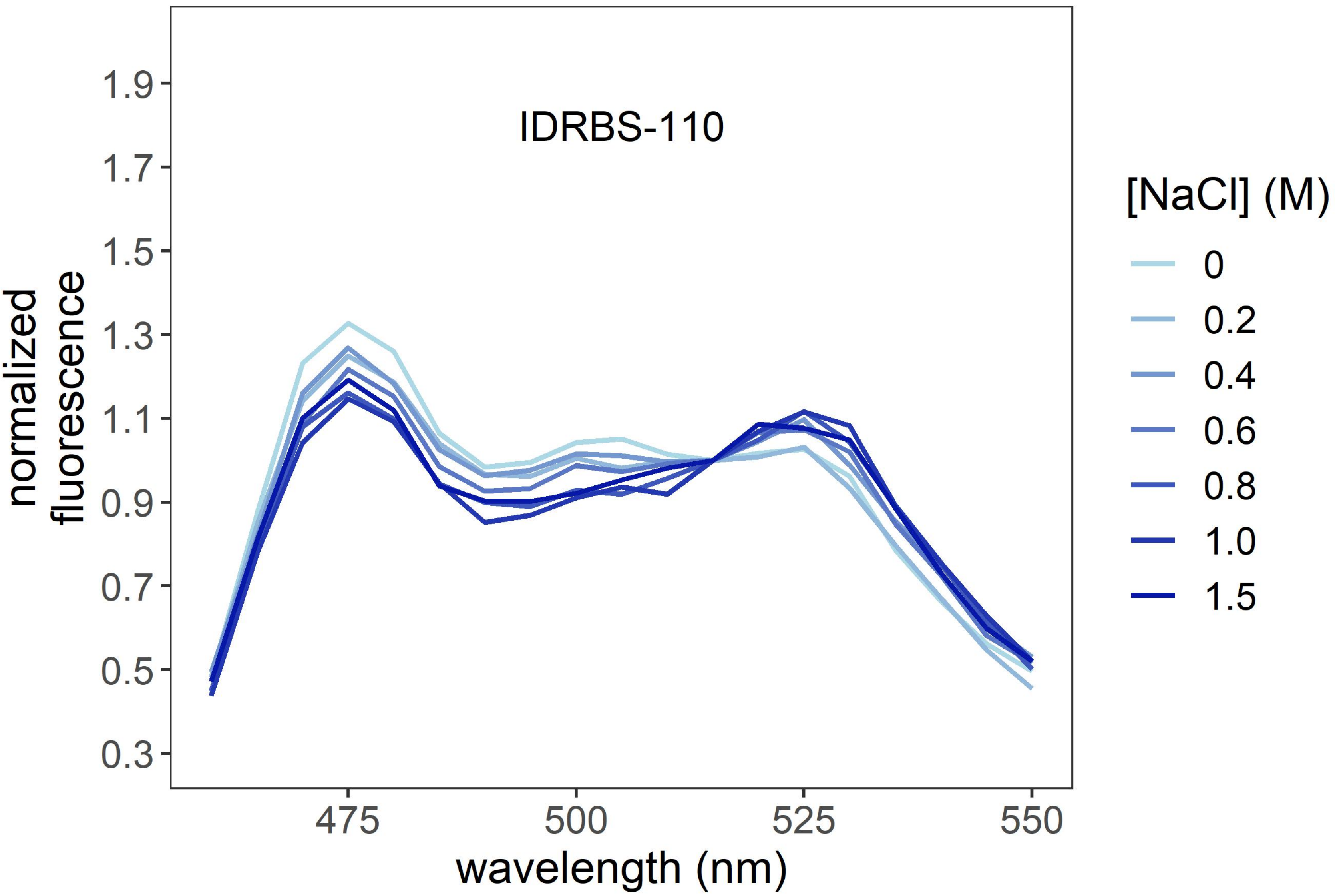
IDRBS-110
1.9
1.7
1.5
1.3
1.1
0.9
0.7
0.5
0.3
normalized
fluorescence
475
500
525
550
wavelength (nm)
[NaCl] (M)
0
0.2
0.4
0.6
0.8
1.0
1.5

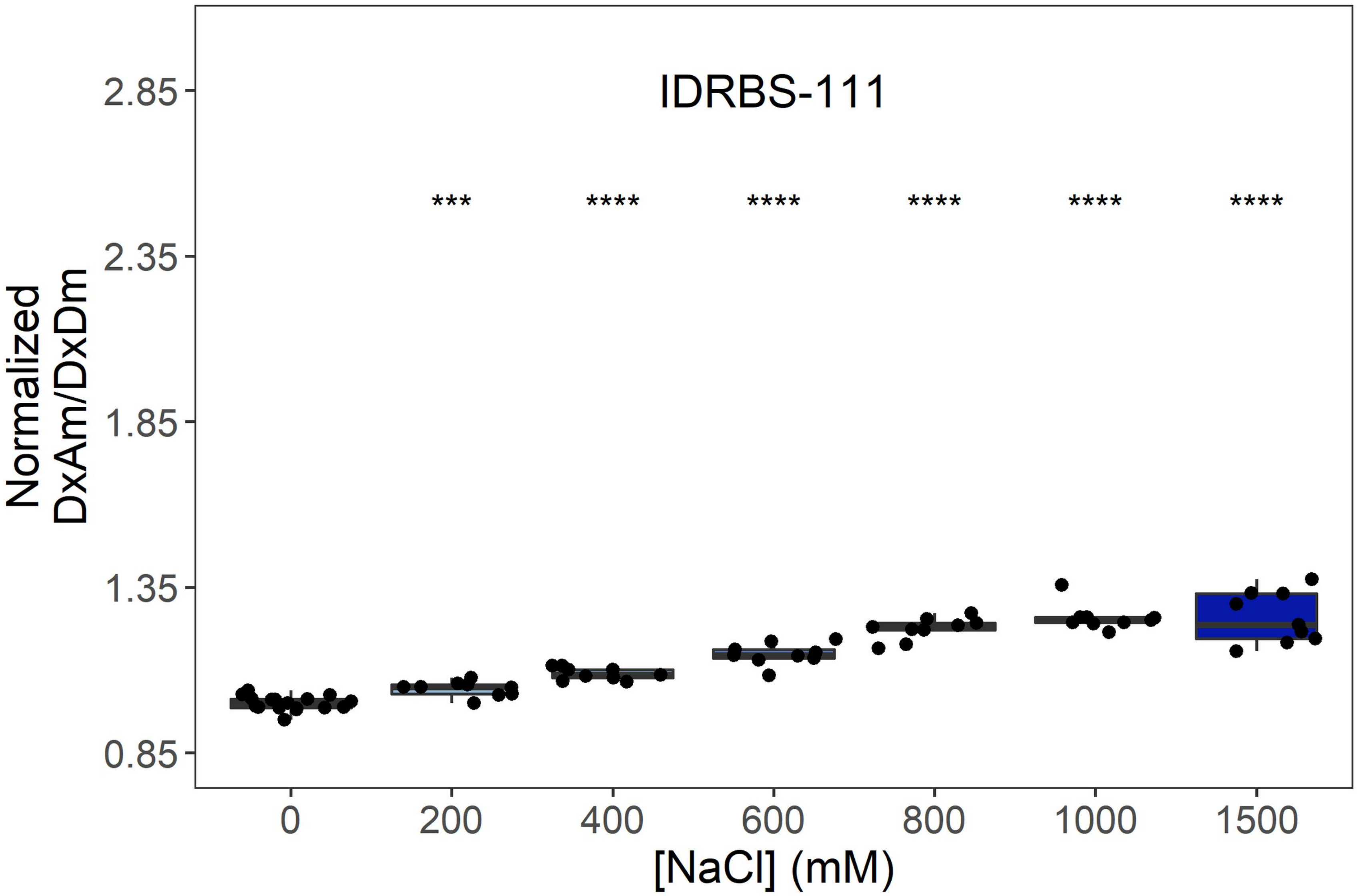
IDRBS-111
***
****
****
****
****
****
Normalized DxAm/DxDm
2.85
2.35
1.85
1.35
0.85
0
200
400
600
800
1000
1500
[NaCl] (mM)

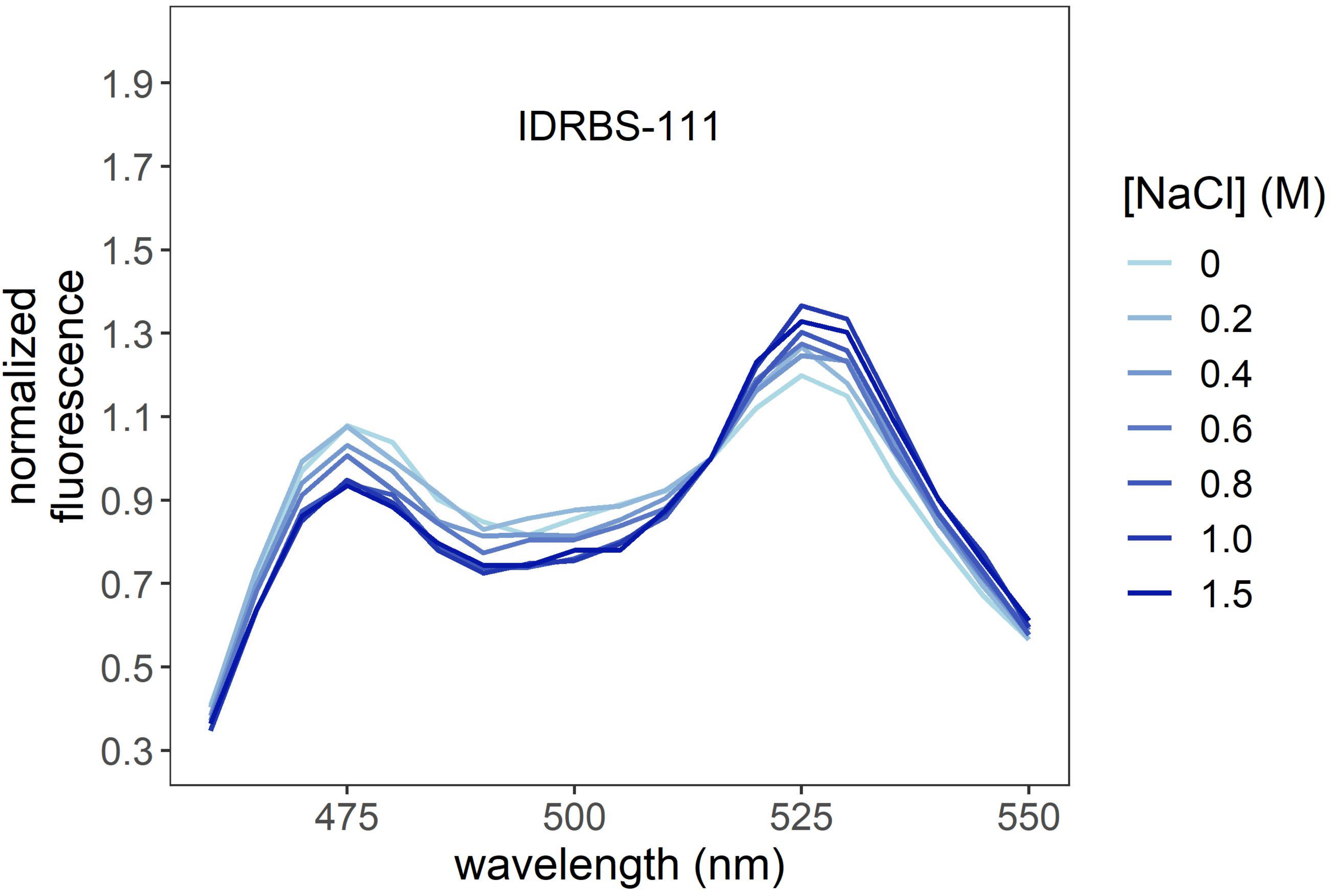
IDRBS-111
normalized fluorescence
1.9
1.7
1.5
1.3
1.1
0.9
0.7
0.5
0.3
475
500
525
550
wavelength (nm)
[NaCl] (M)
0
0.2
0.4
0.6
0.8
1.0
1.5

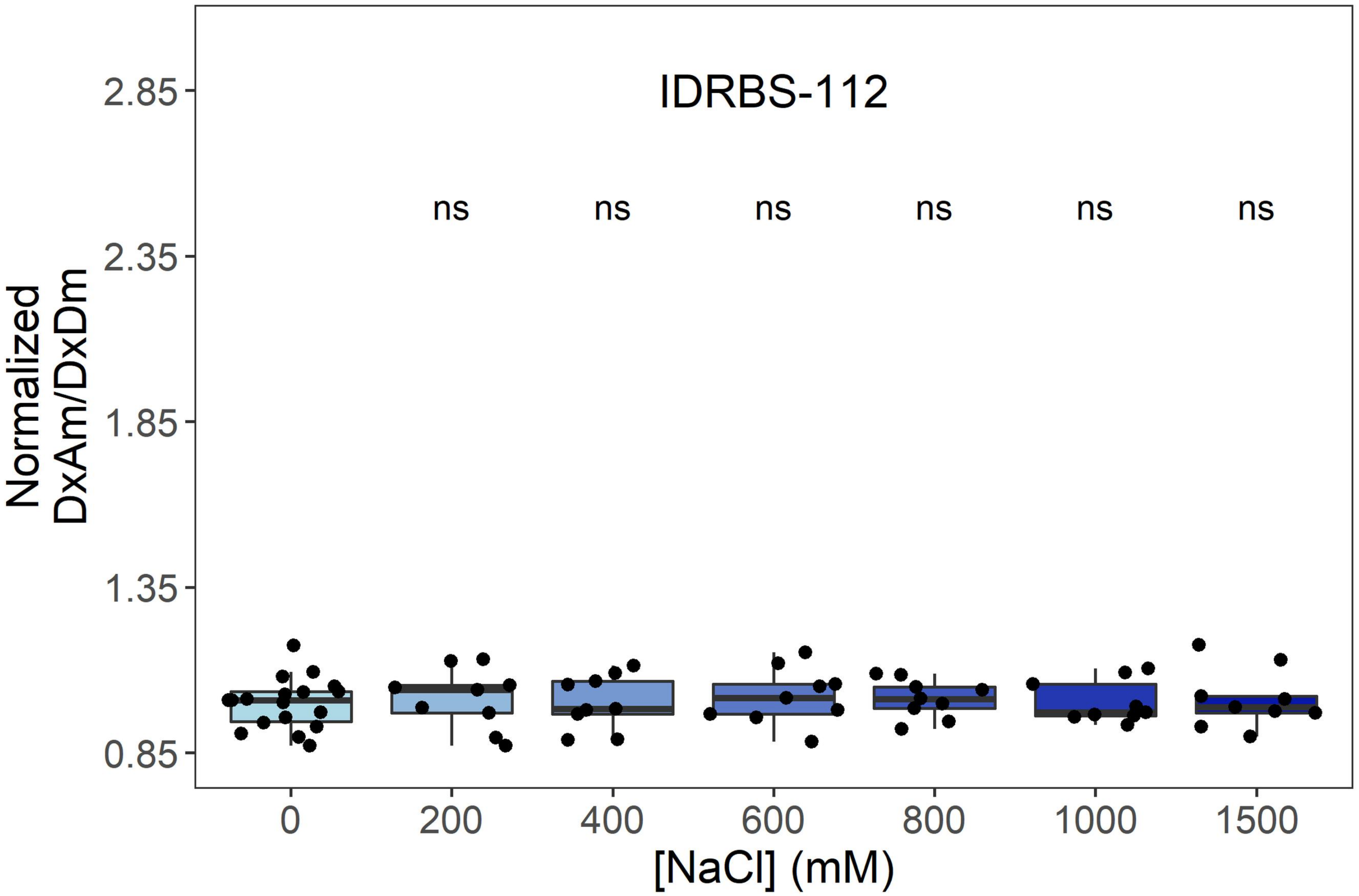
IDRBS-112
ns
ns
ns
ns
ns
ns
Normalized
DxAm/DxDm
2.85
2.35
1.85
1.35
0.85
0
200
400
600
800
1000
1500
[NaCl] (mM)

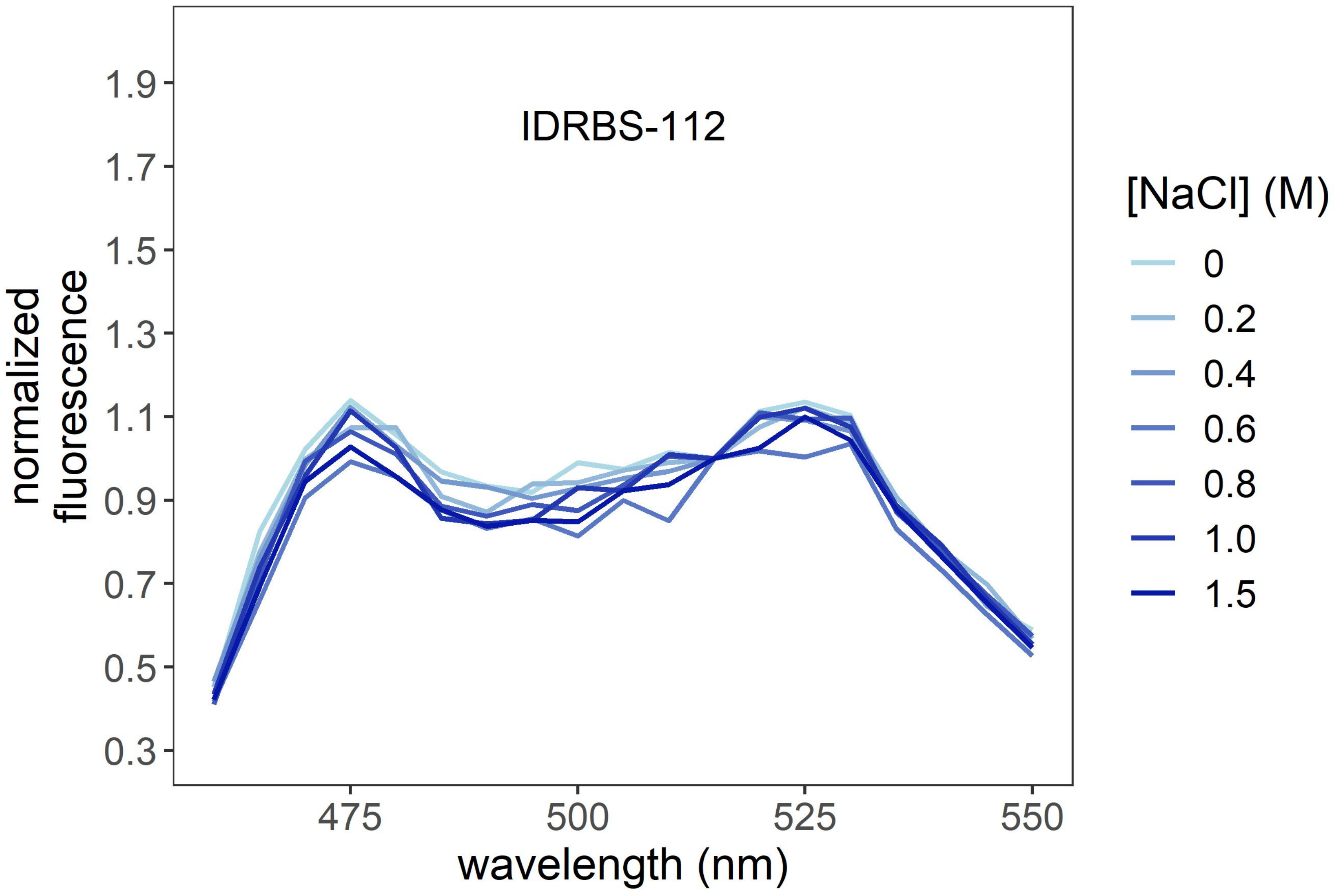
IDRBS-112
normalized
fluorescence
1.9
1.7
1.5
1.3
1.1
0.9
0.7
0.5
0.3
475
500
525
550
wavelength (nm)
[NaCl] (M)
0
0.2
0.4
0.6
0.8
1.0
1.5

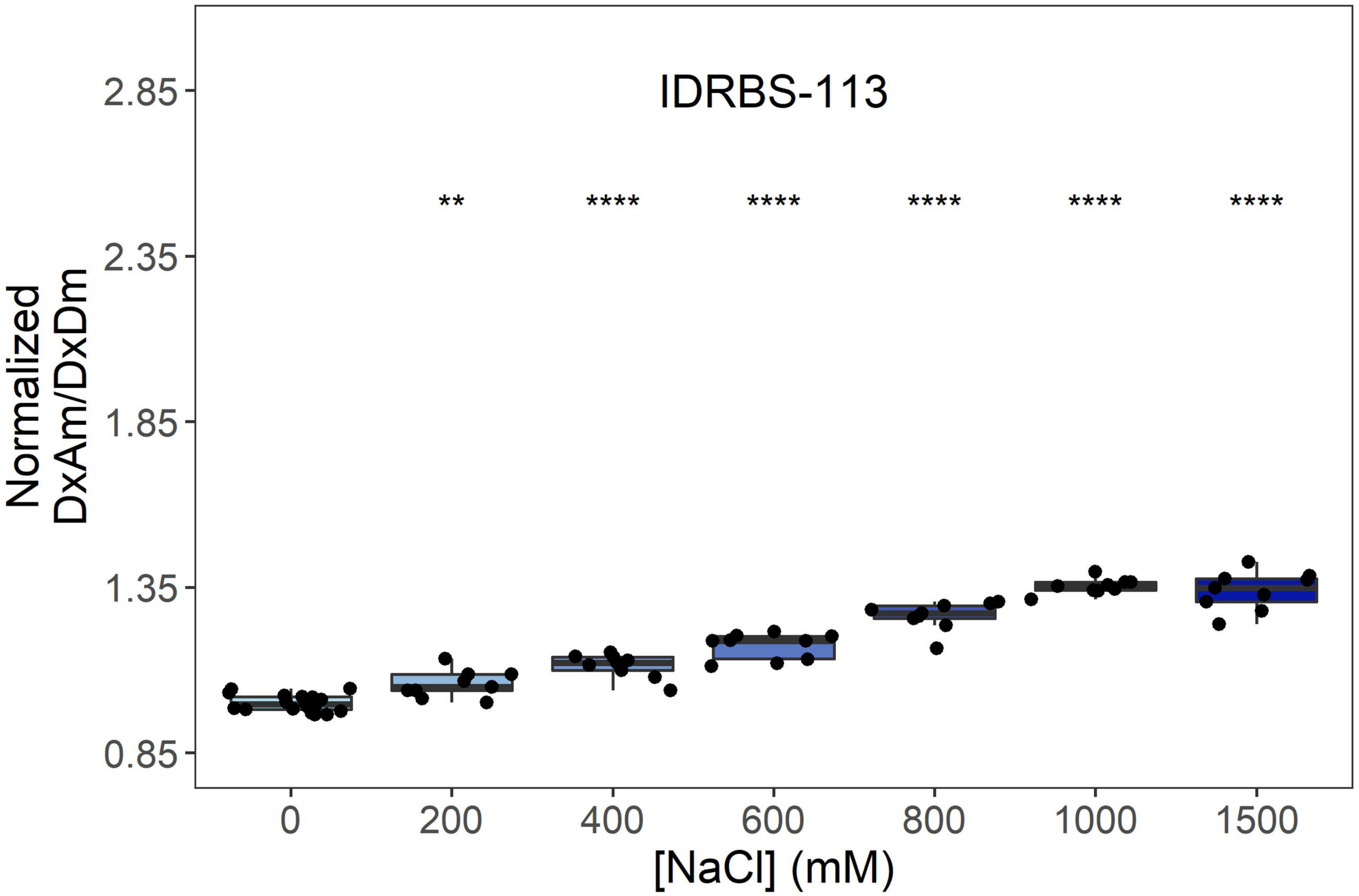
IDRBS-113
Normalized DxAm/DxDm
2.85
2.35
1.85
1.35
0.85
**
****
****
****
****
****
0
200
400
600
800
1000
1500
[NaCl] (mM)

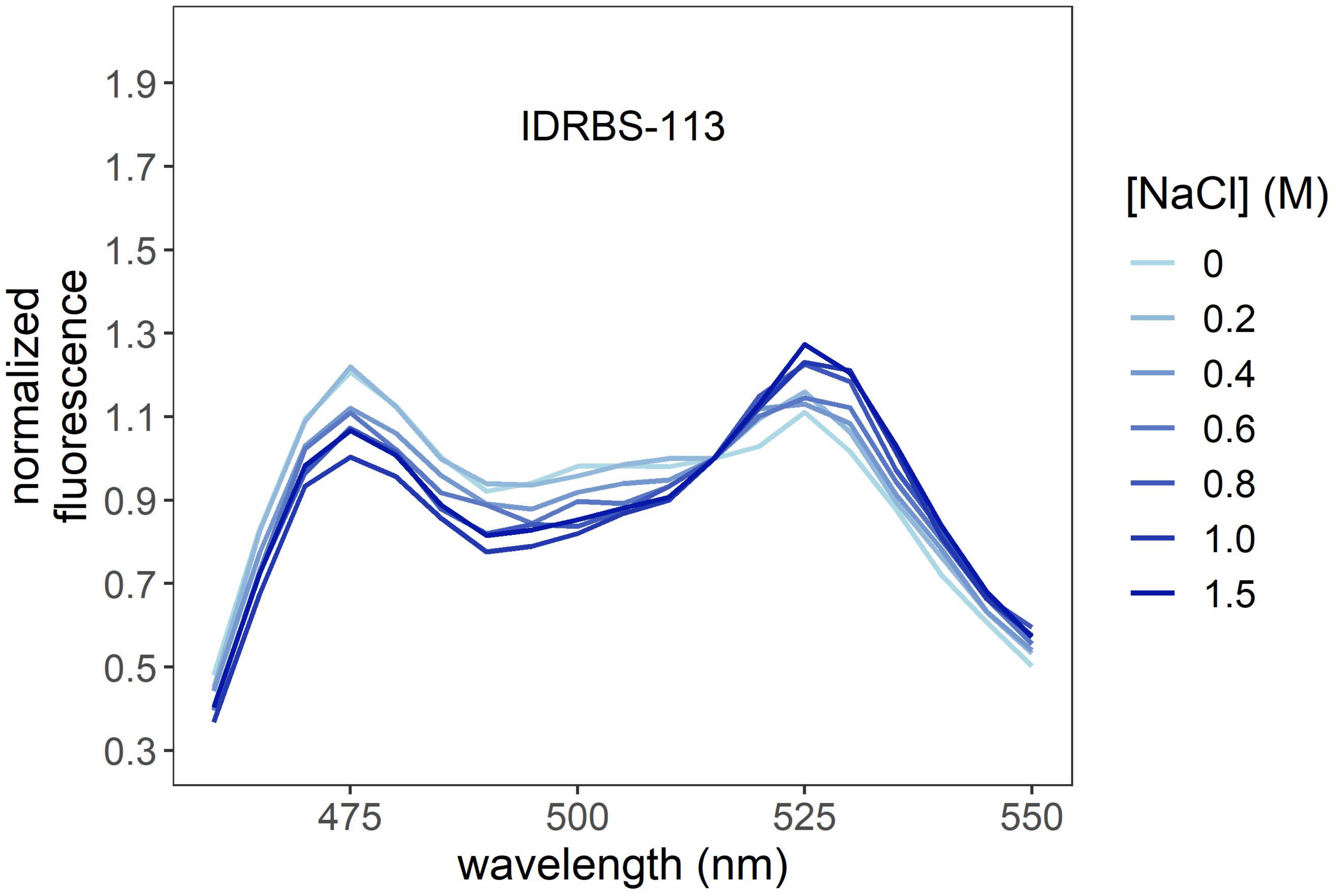
IDRBS-113
normalized fluorescence
1.9
1.7
1.5
1.3
1.1
0.9
0.7
0.5
0.3
475
500
525
550
wavelength (nm)
[NaCl] (M)
0
0.2
0.4
0.6
0.8
1.0
1.5

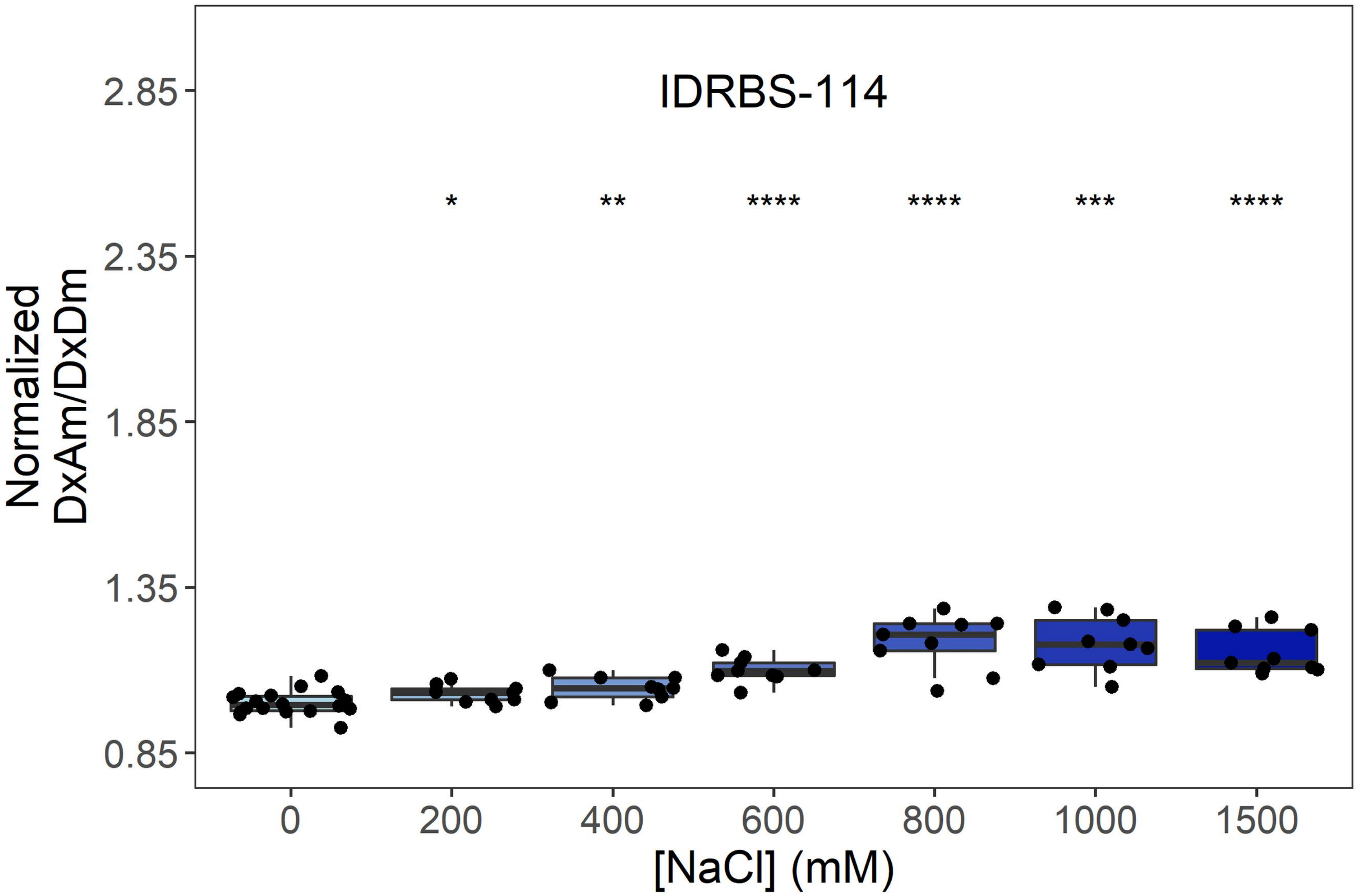

IDRBS-114
*
**
****
****
***
****
Normalized
DxAm/DxDm
2.85
2.35
1.85
1.35
0.85
0
200
400
600
800
1000
1500
[NaCl] (mM)


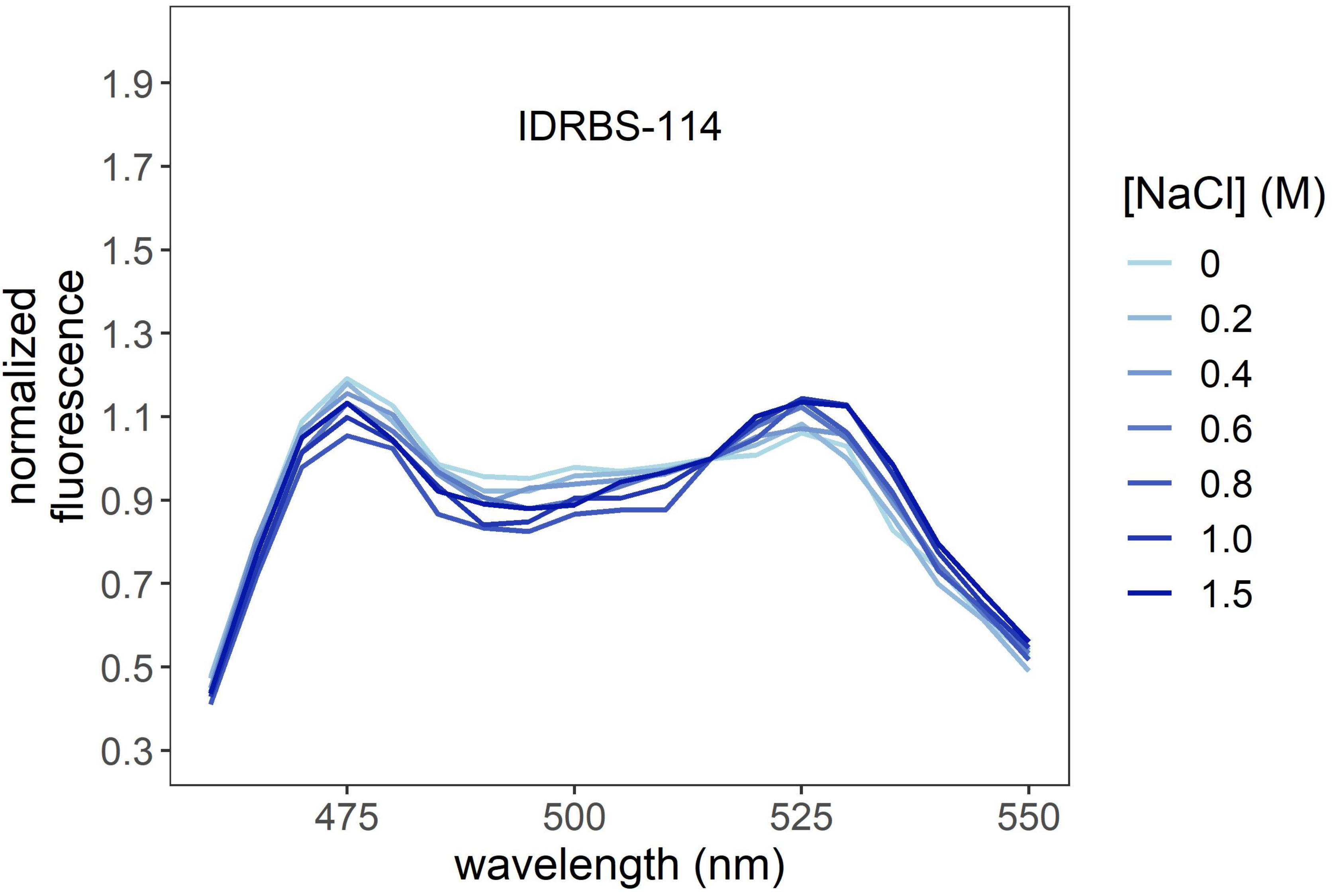

IDRBS-114
normalized
fluorescence
1.9
1.7
1.5
1.3
1.1
0.9
0.7
0.5
0.3
475
500
525
550
wavelength (nm)
[NaCl] (M)
0
0.2
0.4
0.6
0.8
1.0
1.5

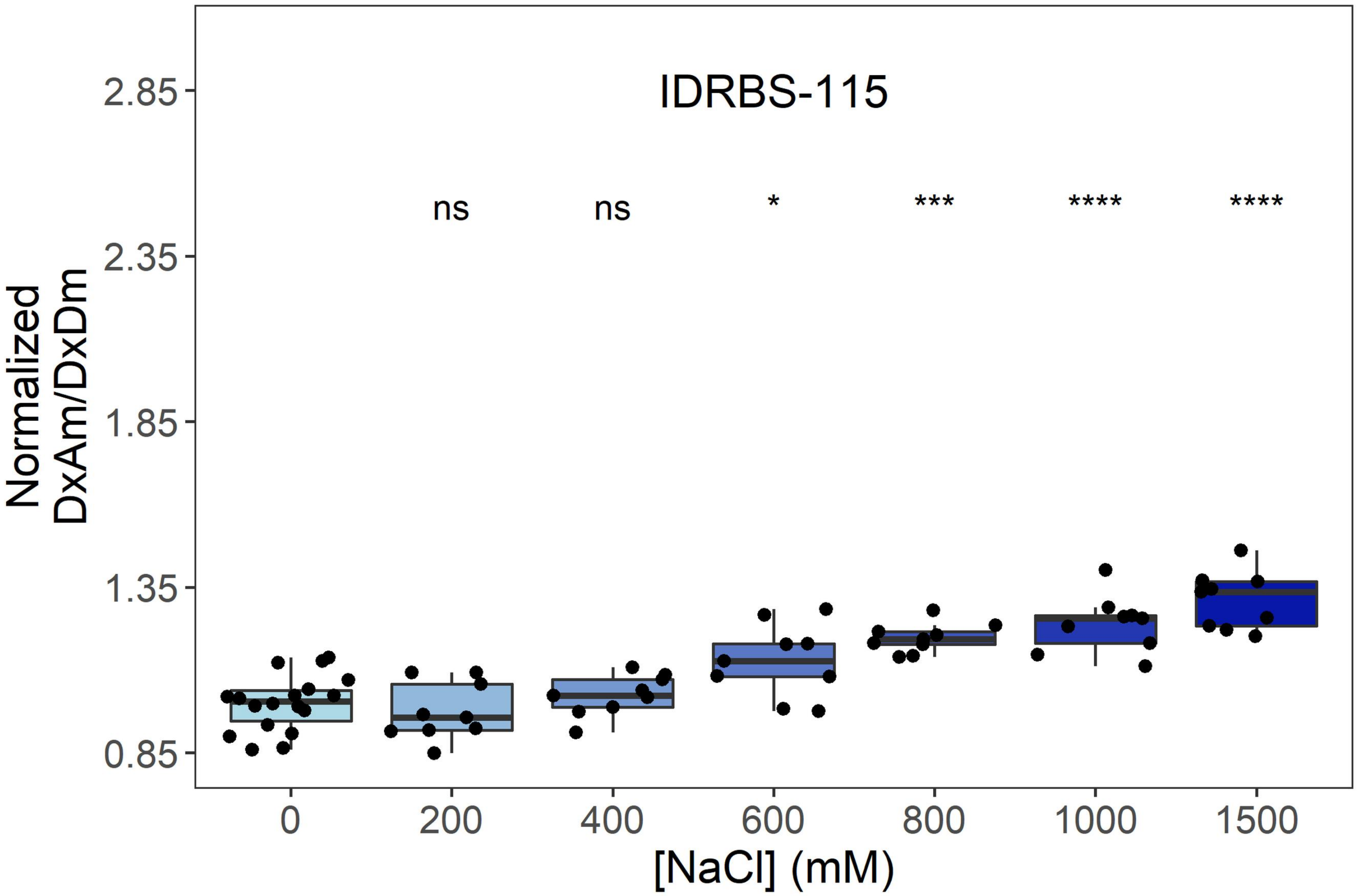
IDRBS-115
ns
ns
*
***
****
****
Normalized DxAm/DxDm
2.85
2.35
1.85
1.35
0.85
0
200
400
600
800
1000
1500
[NaCl] (mM)

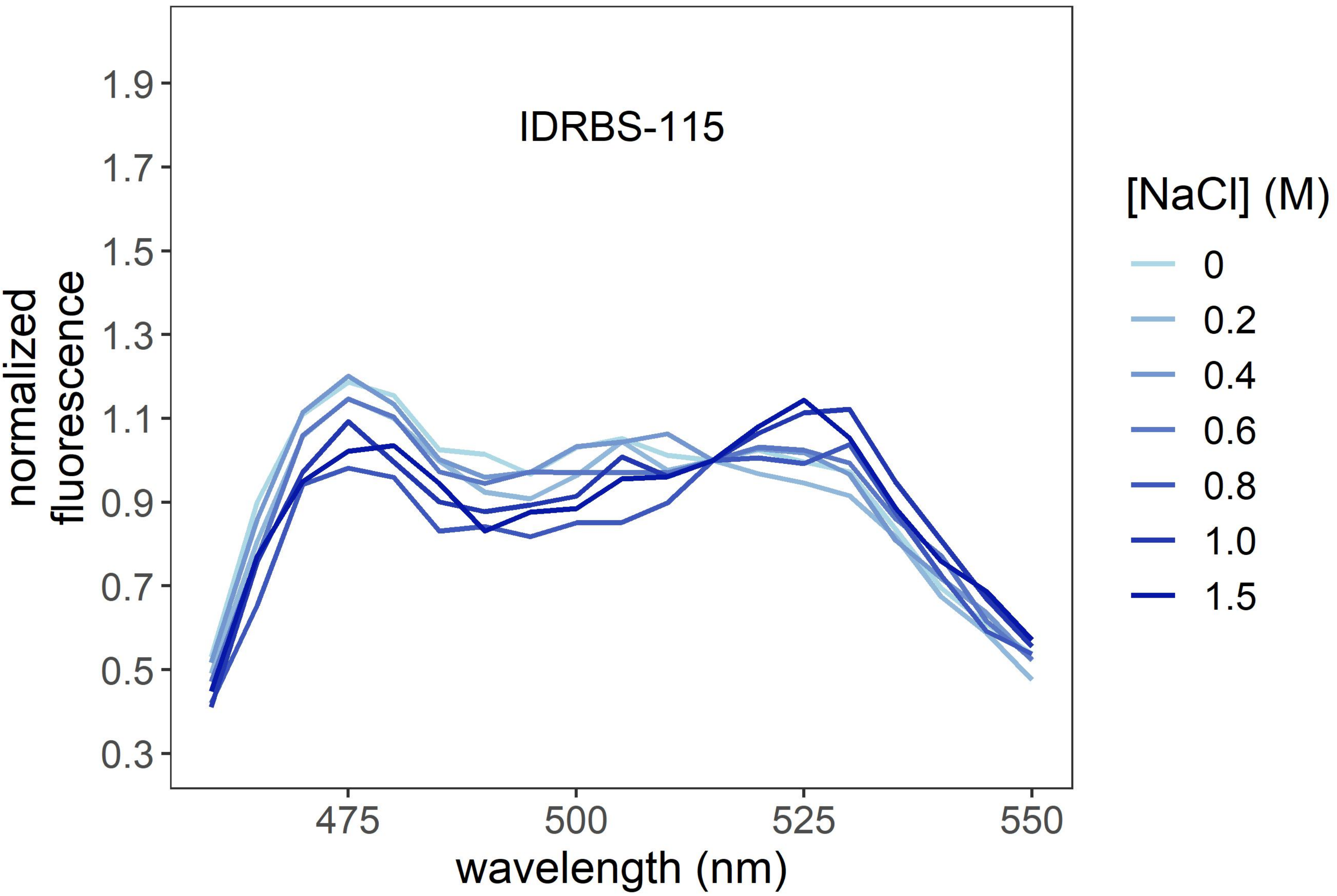
IDRBS-115
normalized fluorescence
1.9
1.7
1.5
1.3
1.1
0.9
0.7
0.5
0.3
475
500
525
550
wavelength (nm)
[NaCl] (M)
0
0.2
0.4
0.6
0.8
1.0
1.5

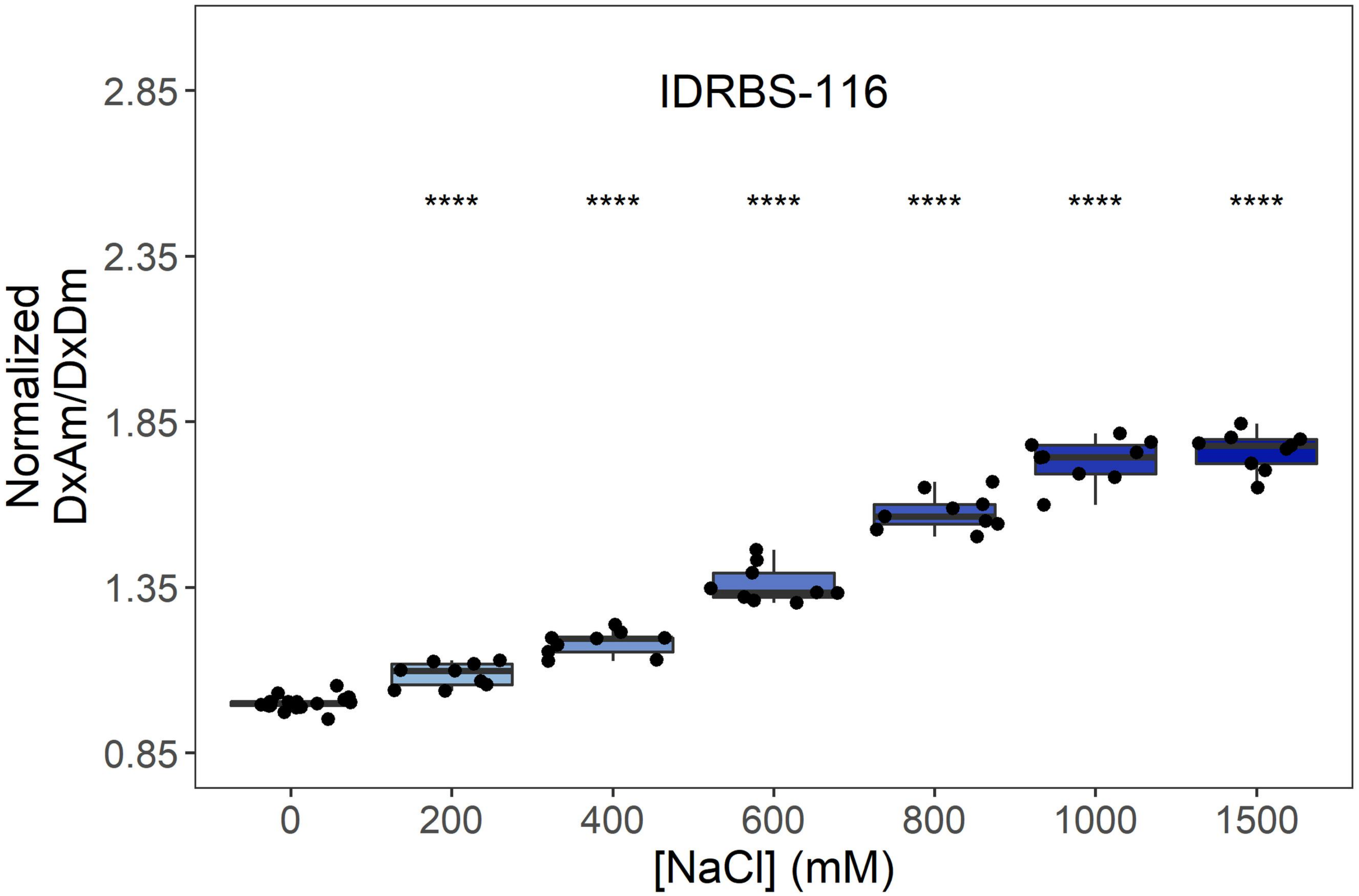
IDRBS-116
****
Normalized DxAm/DxDm
2.85
2.35
1.85
1.35
0.85
0
200
400
600
800
1000
1500
[NaCl] (mM)

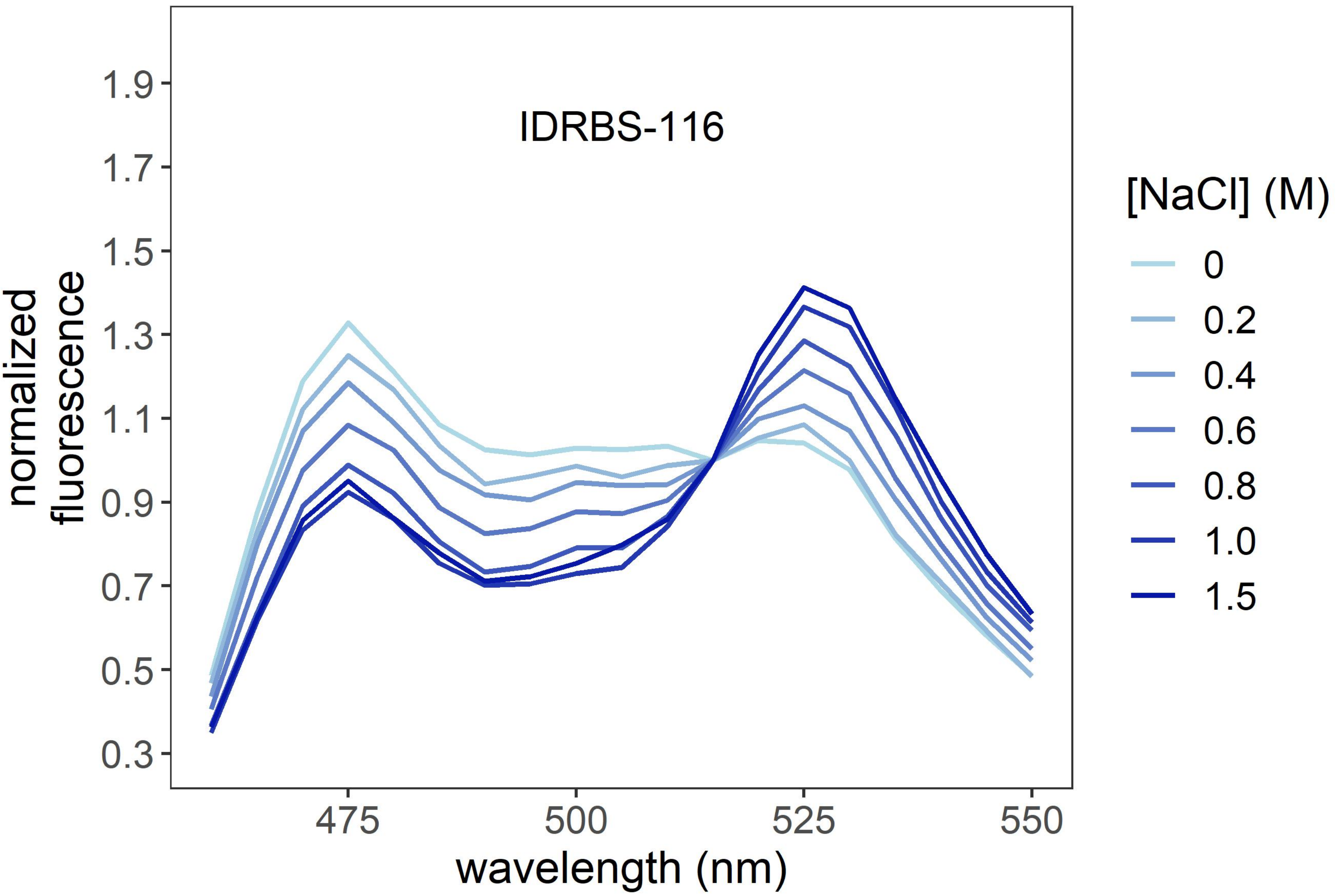
IDRBS-116
normalized fluorescence
1.9
1.7
1.5
1.3
1.1
0.9
0.7
0.5
0.3
475
500
525
550
wavelength (nm)
[NaCl] (M)
0
0.2
0.4
0.6
0.8
1.0
1.5

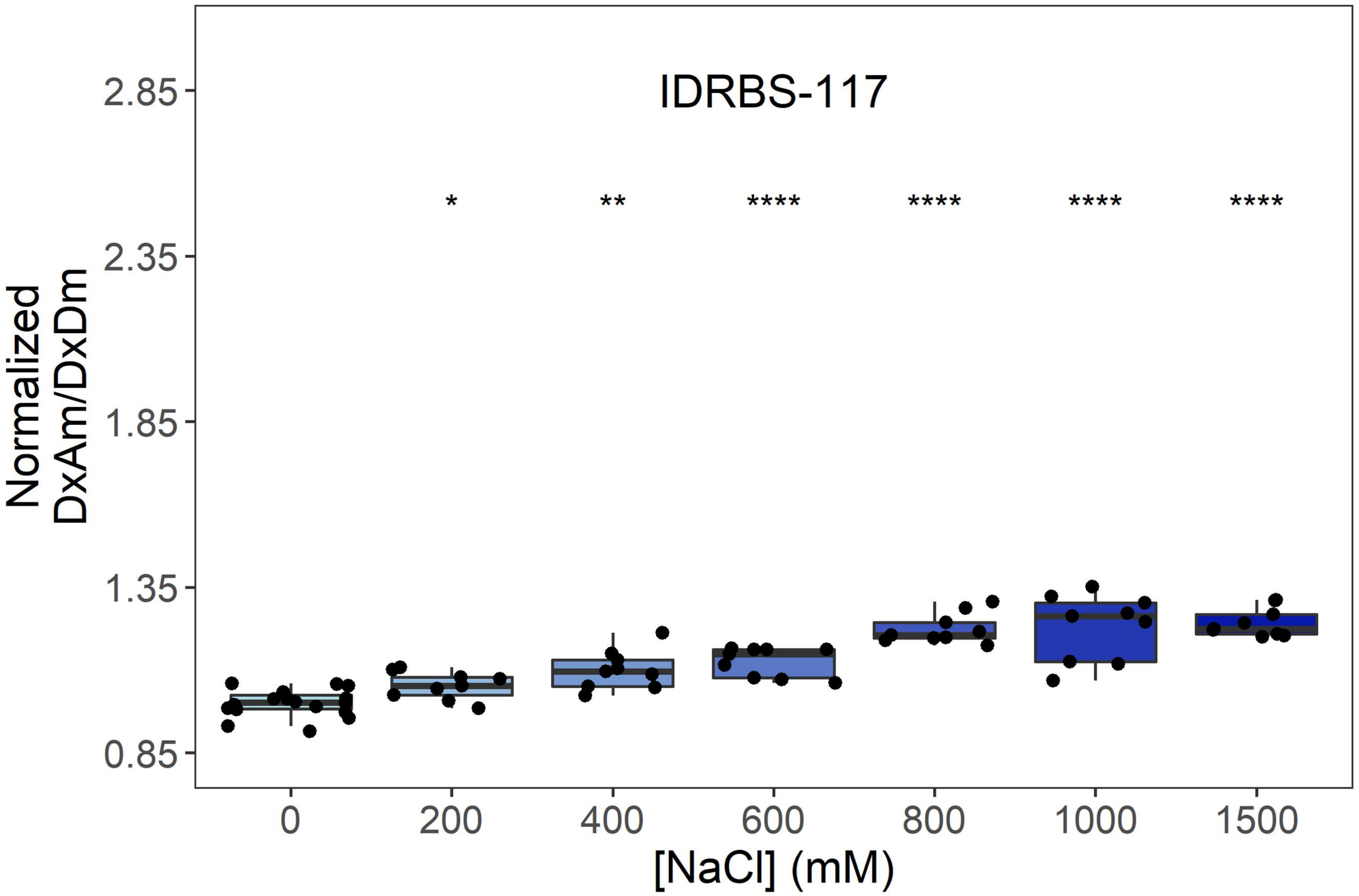

IDRBS-117
Normalized DxAm/DxDm
2.85
2.35
1.85
1.35
0.85
*
**
****
****
****
****
0
200
400
600
800
1000
1500
[NaCl] (mM)


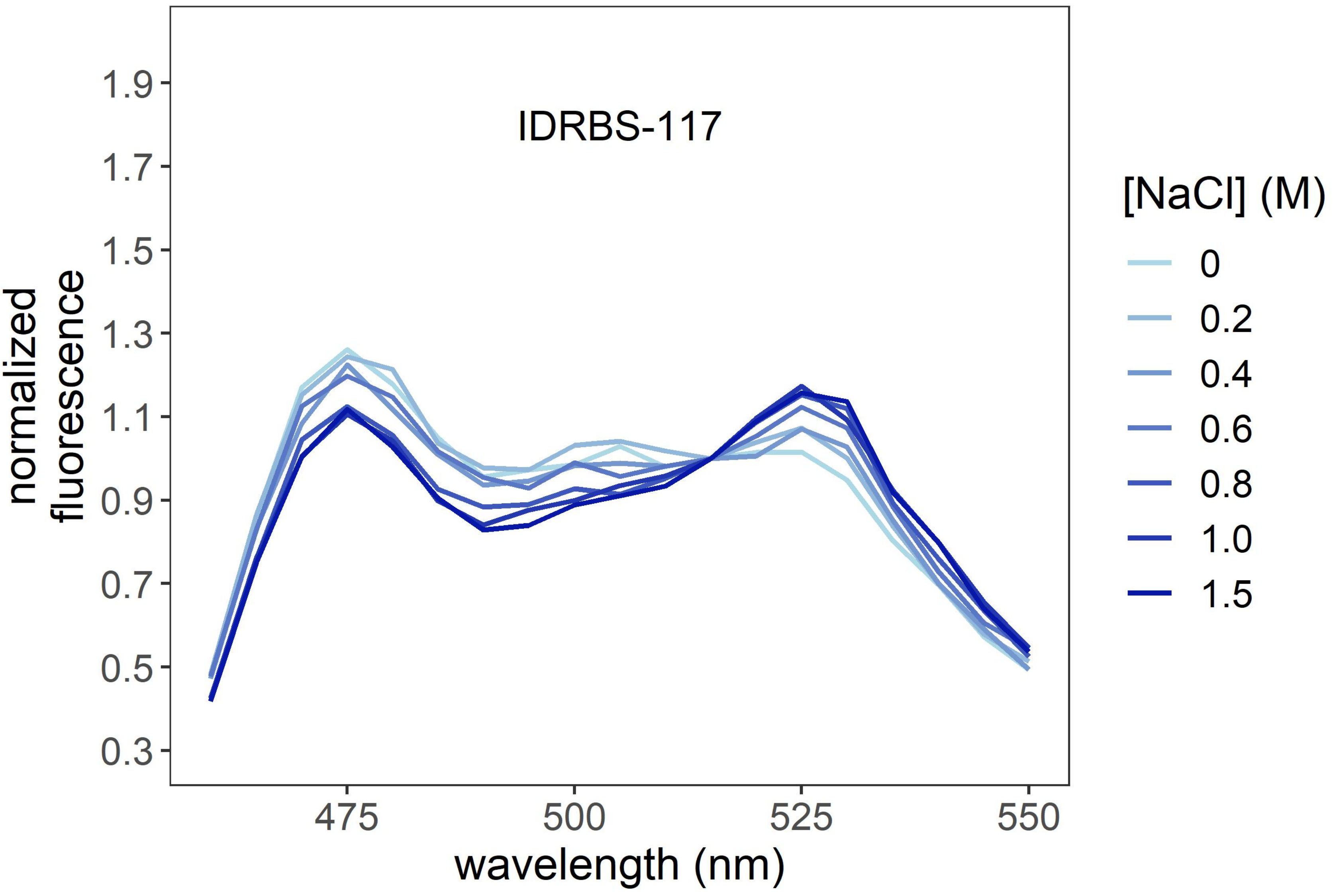

IDRBS-117
normalized fluorescence
1.9
1.7
1.5
1.3
1.1
0.9
0.7
0.5
0.3
475
500
525
550
wavelength (nm)
[NaCl] (M)
0
0.2
0.4
0.6
0.8
1.0
1.5

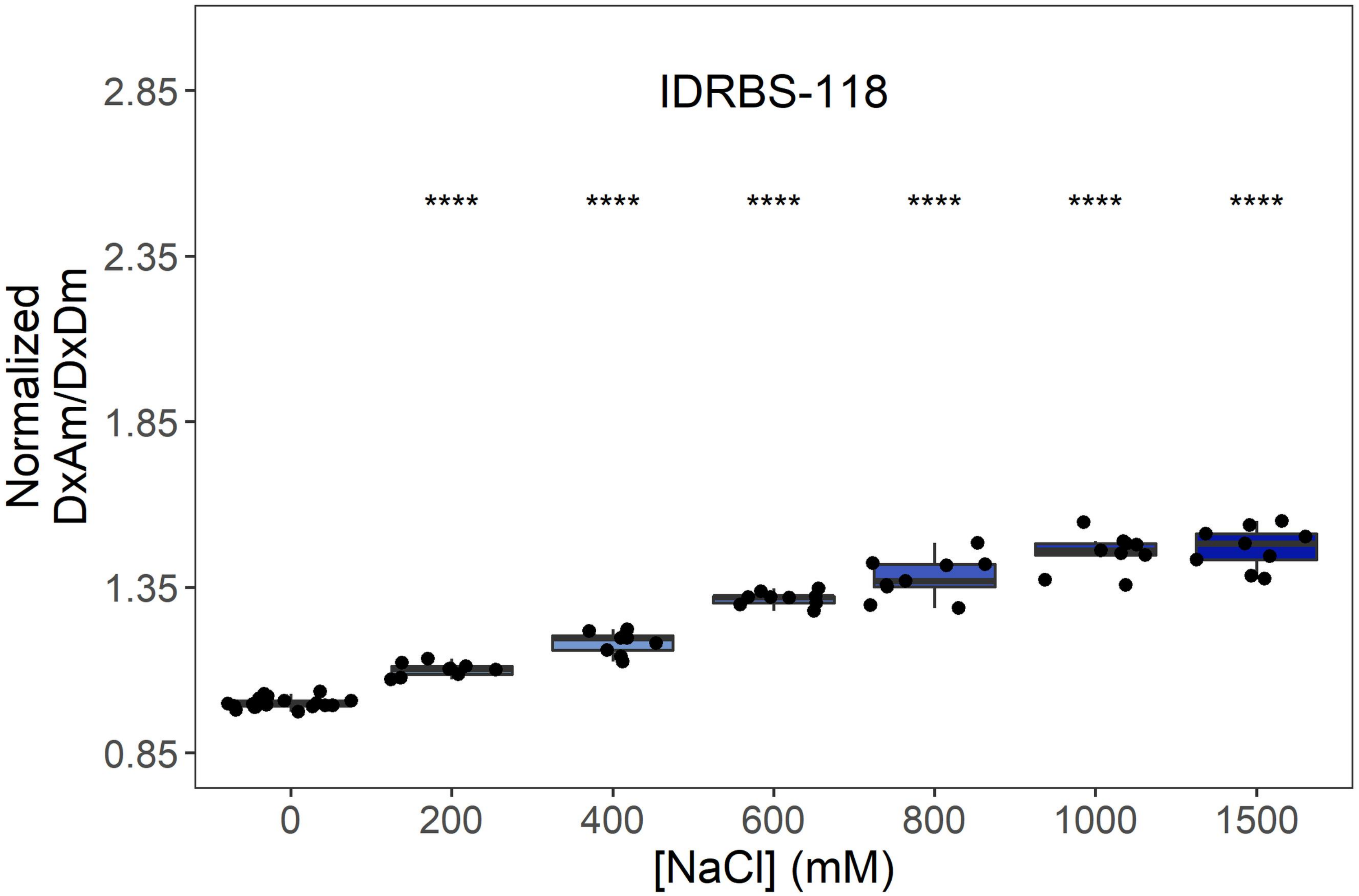
IDRBS-118
Normalized DxAm/DxDm
2.85
2.35
1.85
1.35
0.85
****
0
200
400
600
800
1000
1500
[NaCl] (mM)

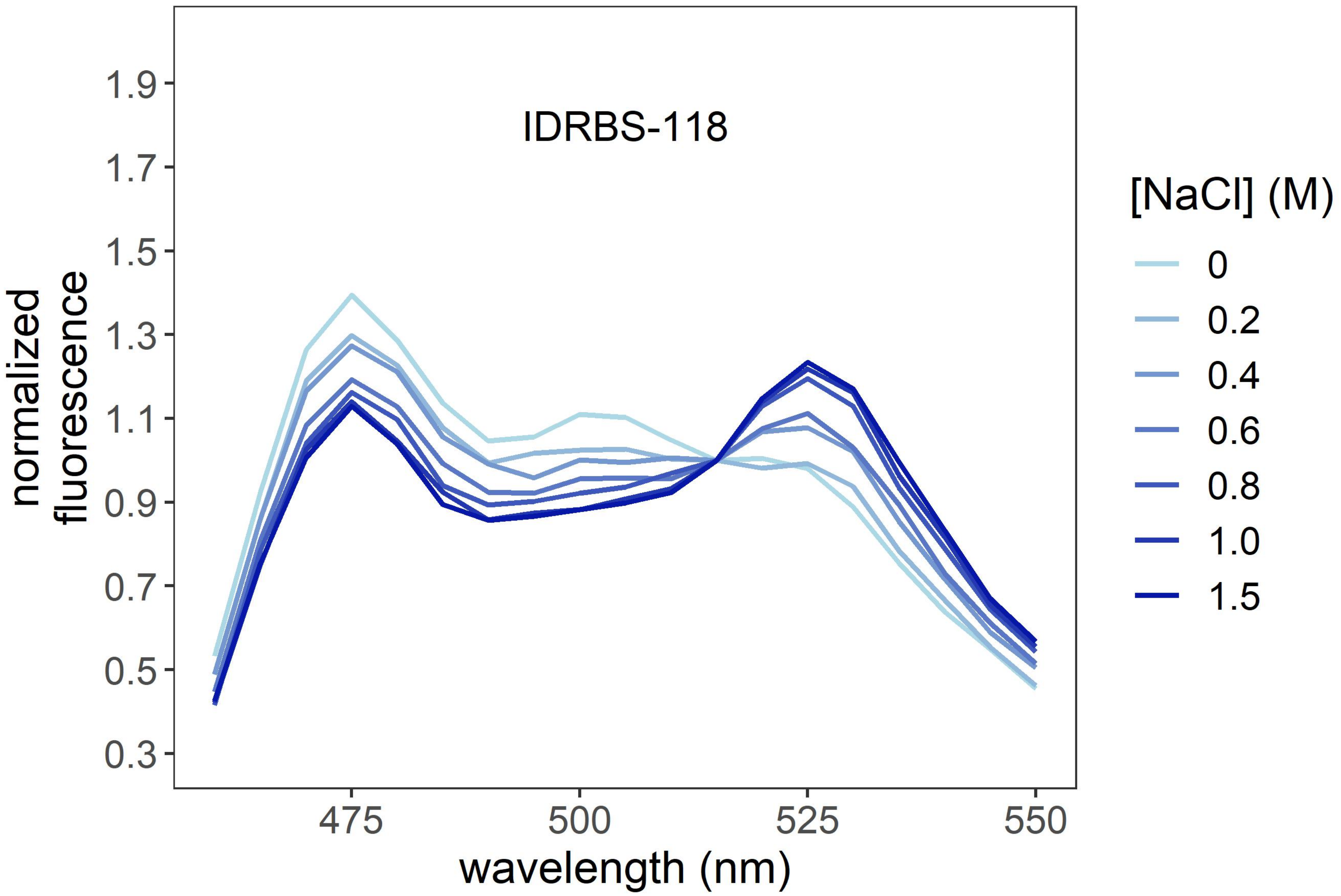
IDRBS-118
normalized fluorescence
1.9
1.7
1.5
1.3
1.1
0.9
0.7
0.5
0.3
475
500
525
550
wavelength (nm)
[NaCl] (M)
0
0.2
0.4
0.6
0.8
1.0
1.5

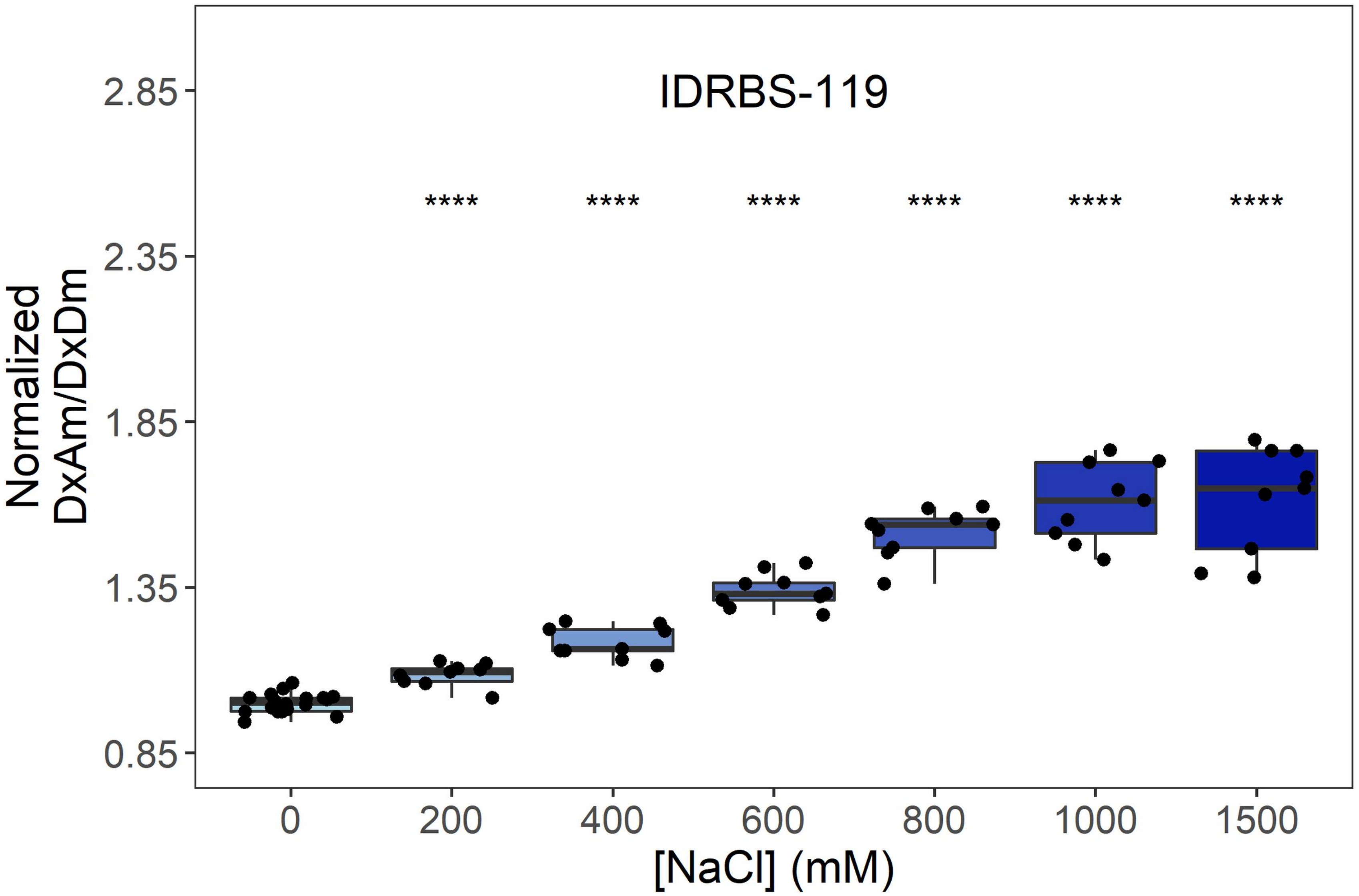
IDRBS-119
Normalized
DxAm/DxDm
2.85
2.35
1.85
1.35
0.85
****
****
****
****
****
****
0
200
400
600
800
1000
1500
[NaCl] (mM)

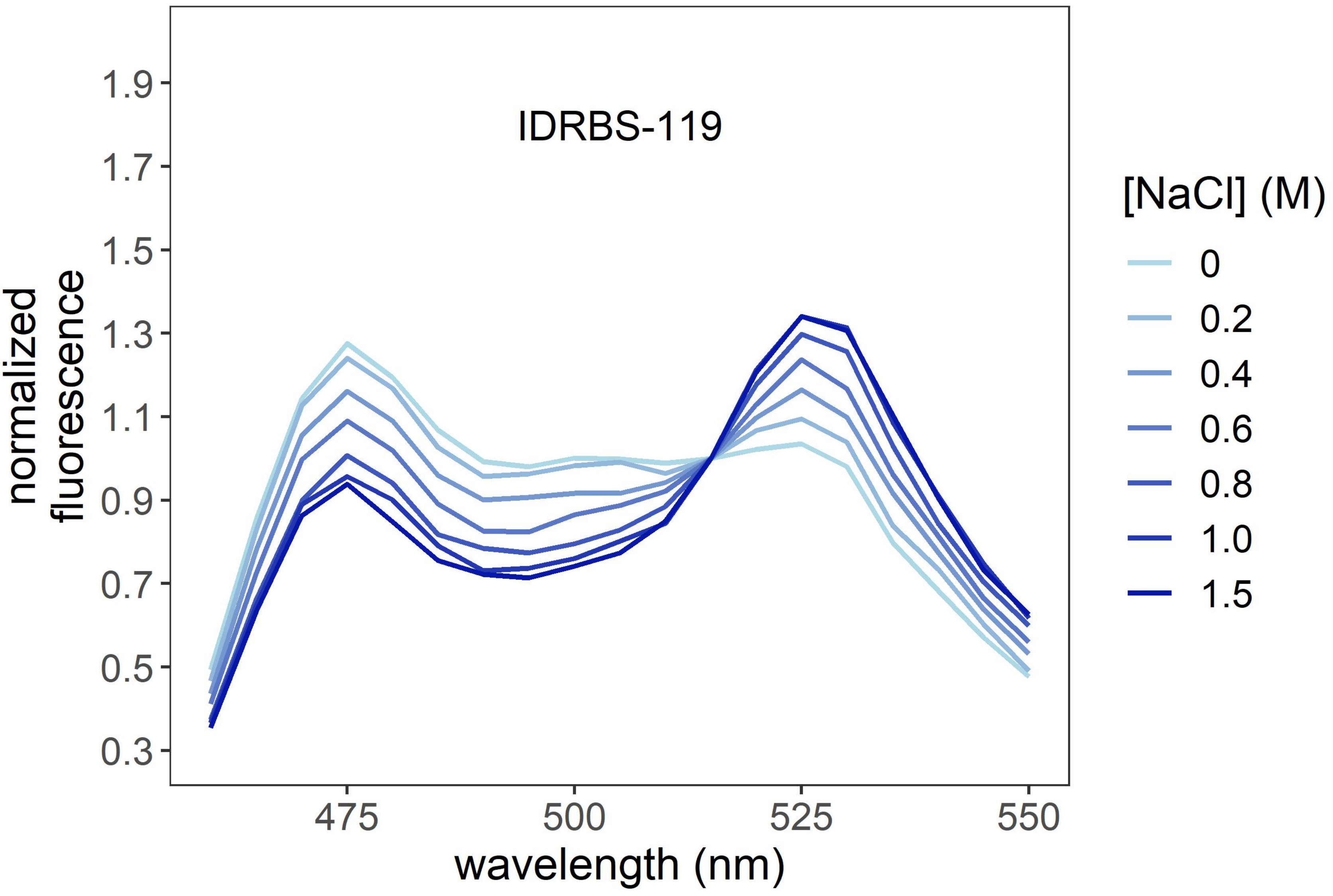
IDRBS-119
normalized
fluorescence
1.9
1.7
1.5
1.3
1.1
0.9
0.7
0.5
0.3
475
500
525
550
wavelength (nm)
[NaCl] (M)
0
0.2
0.4
0.6
0.8
1.0
1.5

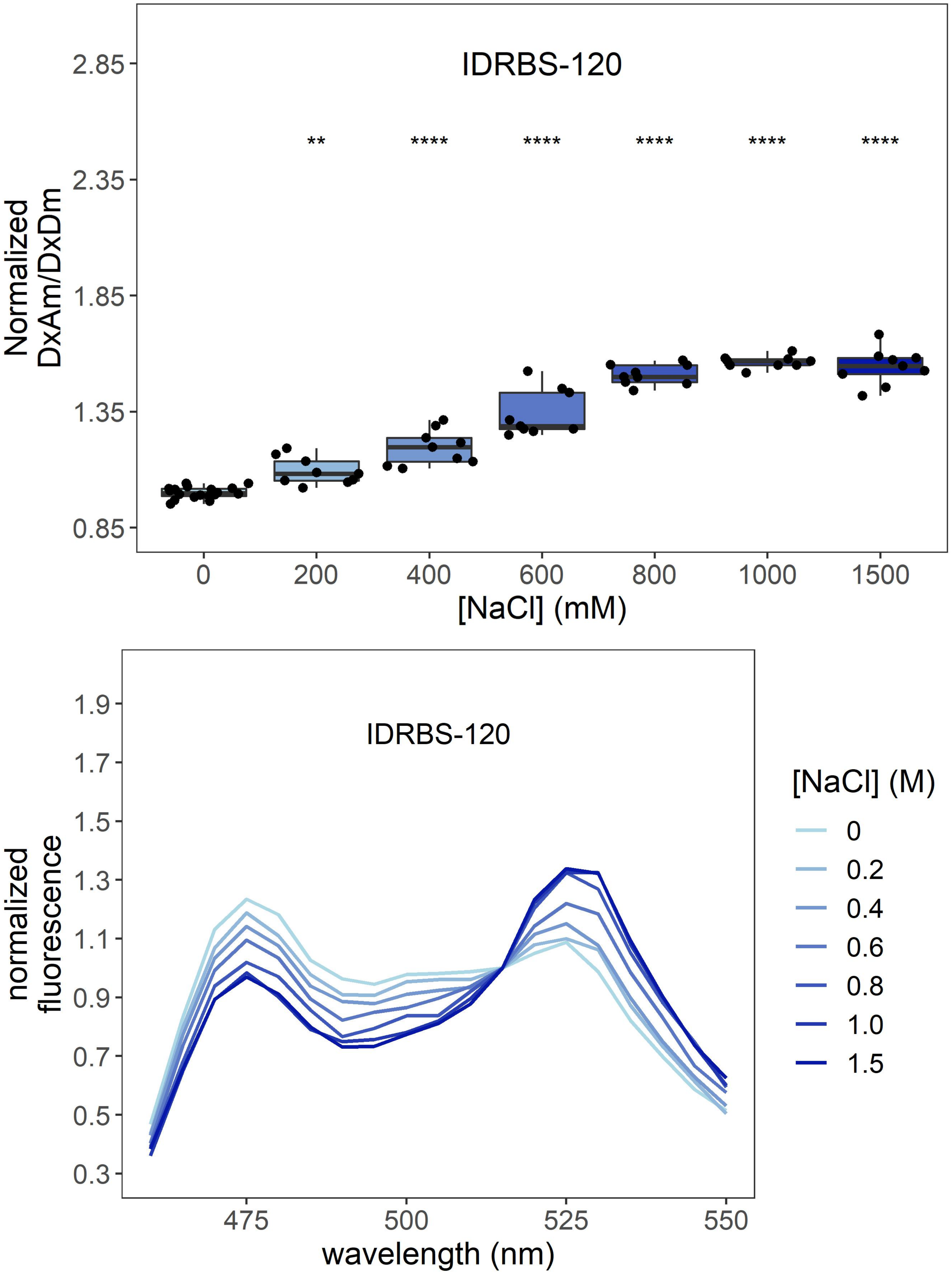

IDRBS-120
Normalized
DxAm/DxDm
2.85
2.35
1.85
1.35
0.85
**
****
****
****
****
****
0
200
400
600
800
1000
1500
[NaCl] (mM)
IDRBS-120
normalized
fluorescence
1.9
1.7
1.5
1.3
1.1
0.9
0.7
0.5
0.3
475
500
525
550
wavelength (nm)
[NaCl] (M)
0
0.2
0.4
0.6
0.8
1.0
1.5

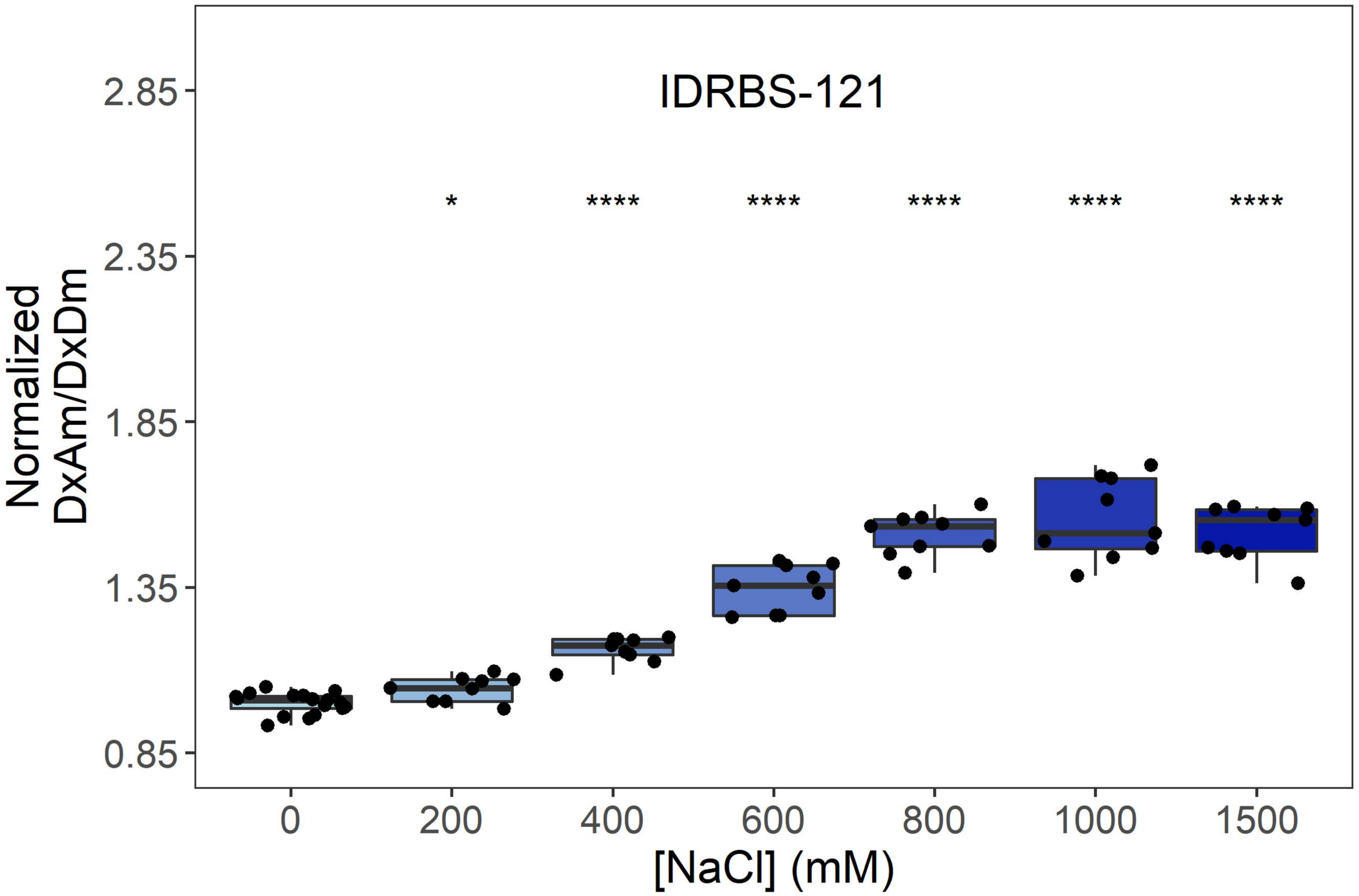

IDRBS-121
Normalized DxAm/DxDm
2.85
2.35
1.85
1.35
0.85
*
****
****
****
****
****
0
200
400
600
800
1000
1500
[NaCl] (mM)


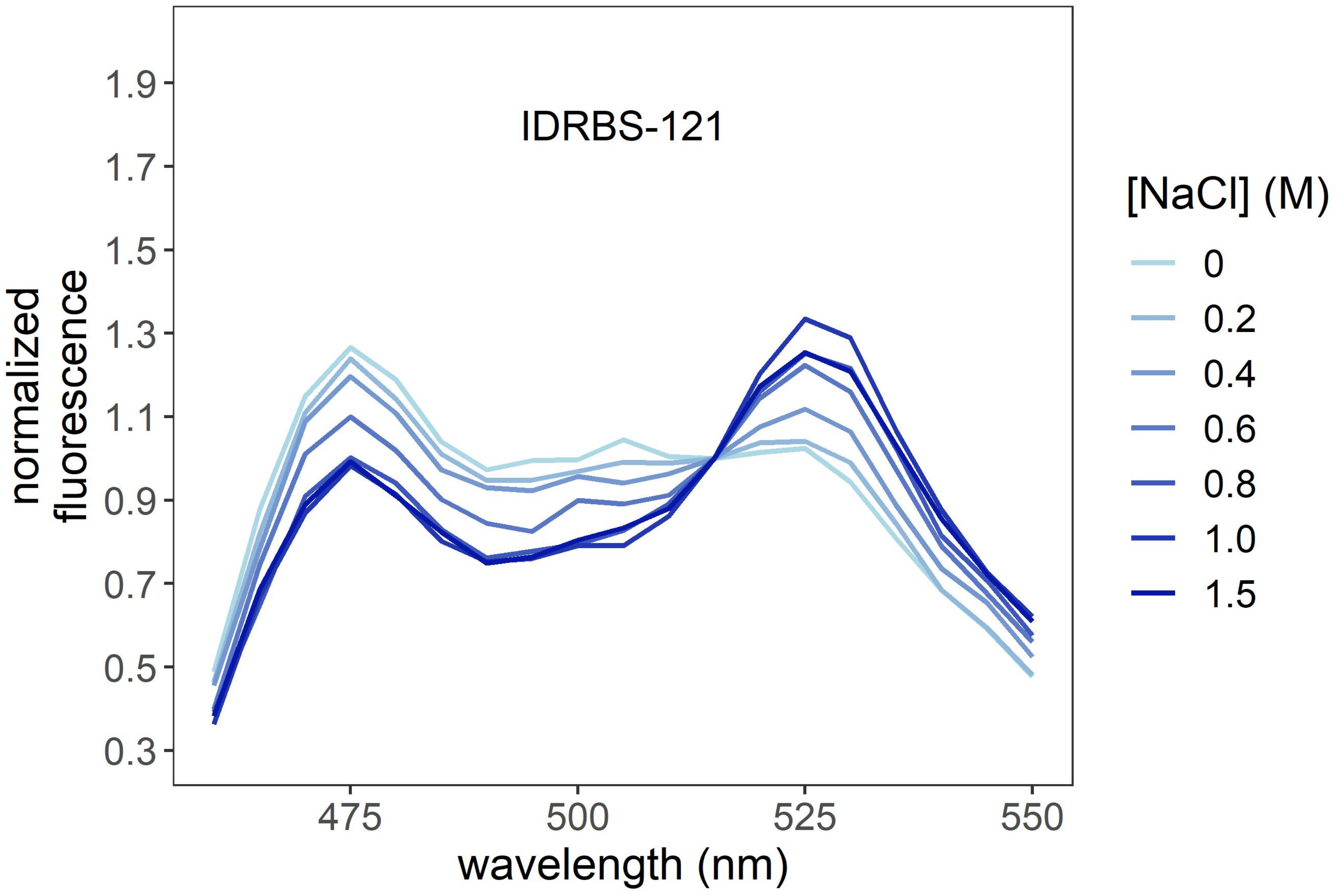

IDRBS-121
normalized fluorescence
1.9
1.7
1.5
1.3
1.1
0.9
0.7
0.5
0.3
475
500
525
550
wavelength (nm)
[NaCl] (M)
0
0.2
0.4
0.6
0.8
1.0
1.5

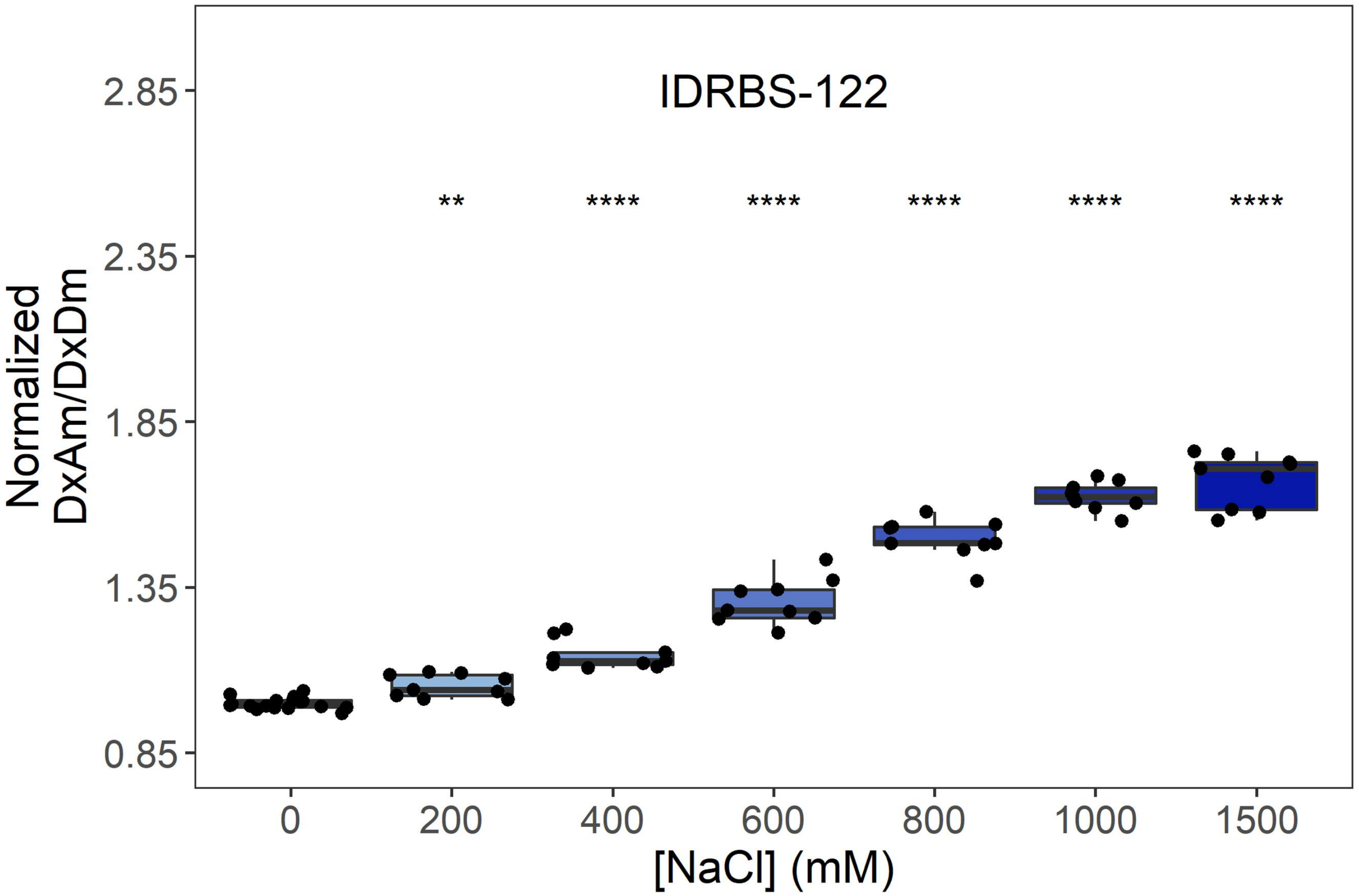
IDRBS-122
Normalized DxAm/DxDm
2.85
2.35
1.85
1.35
0.85
**
****
****
****
****
****
0
200
400
600
800
1000
1500
[NaCl] (mM)

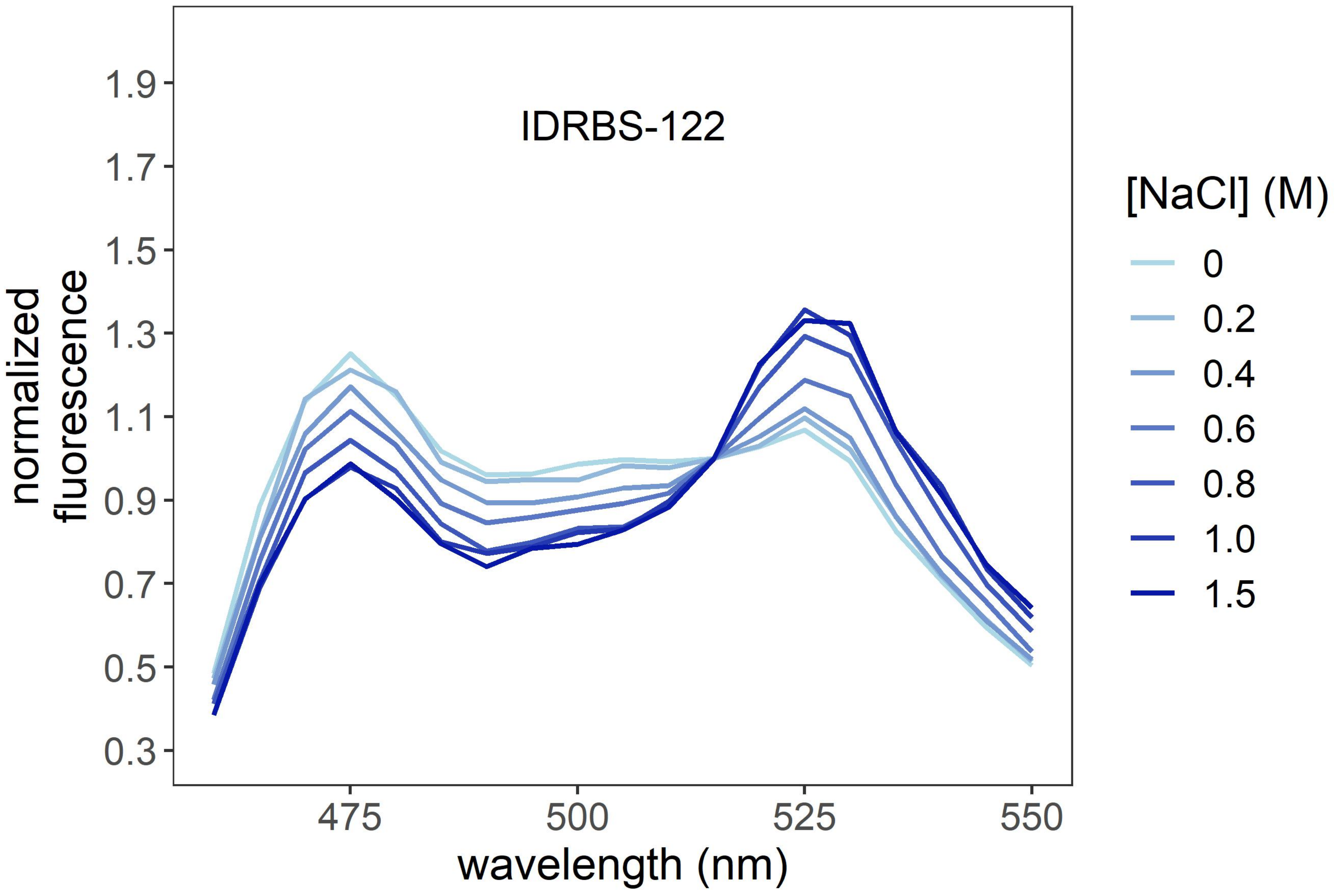
IDRBS-122
normalized fluorescence
1.9
1.7
1.5
1.3
1.1
0.9
0.7
0.5
0.3
475
500
525
550
wavelength (nm)
[NaCl] (M)
0
0.2
0.4
0.6
0.8
1.0
1.5

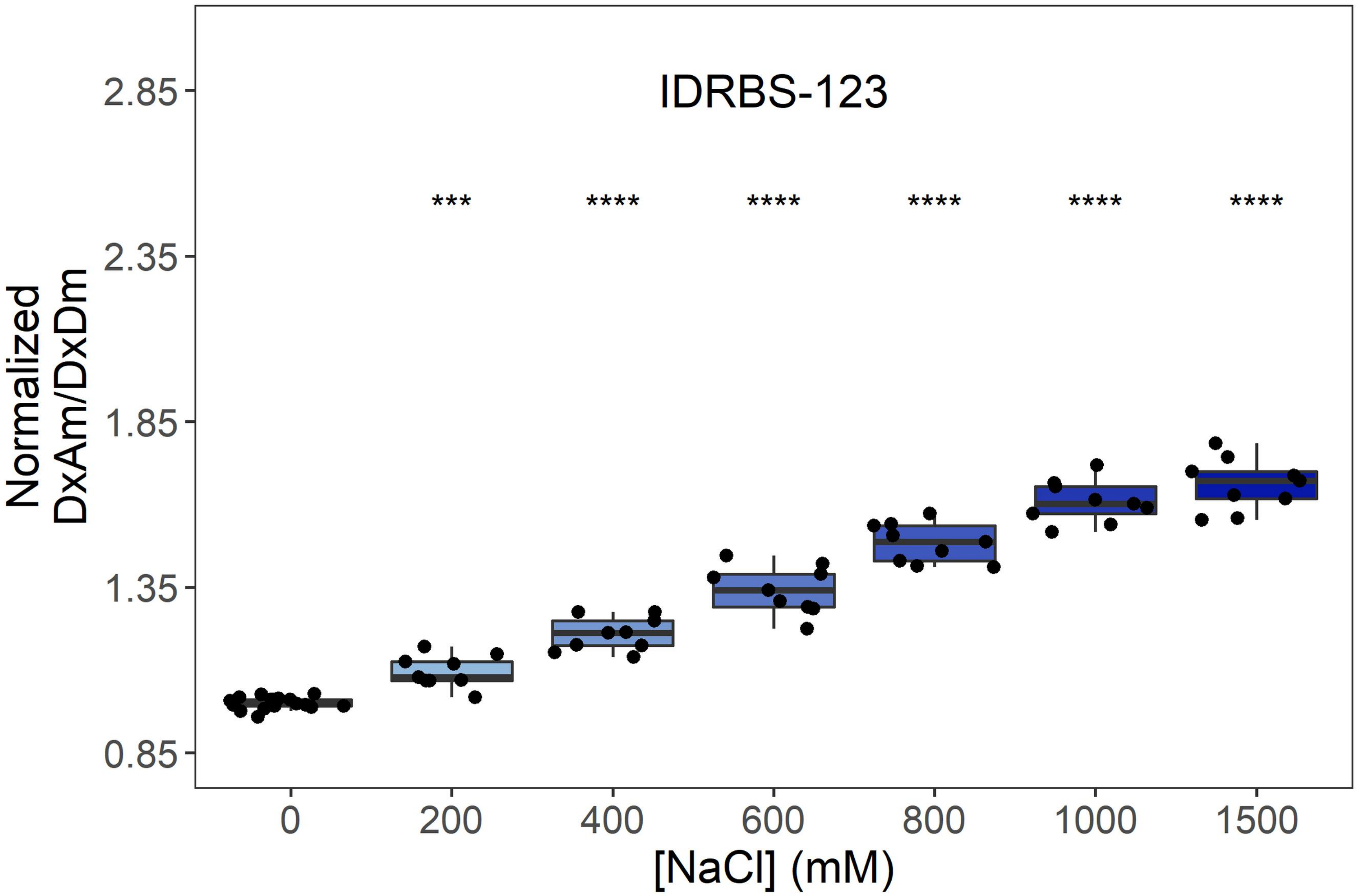
IDRBS-123
Normalized DxAm/DxDm
2.85
2.35
1.85
1.35
0.85
***
****
****
****
****
****
0
200
400
600
800
1000
1500
[NaCl] (mM)

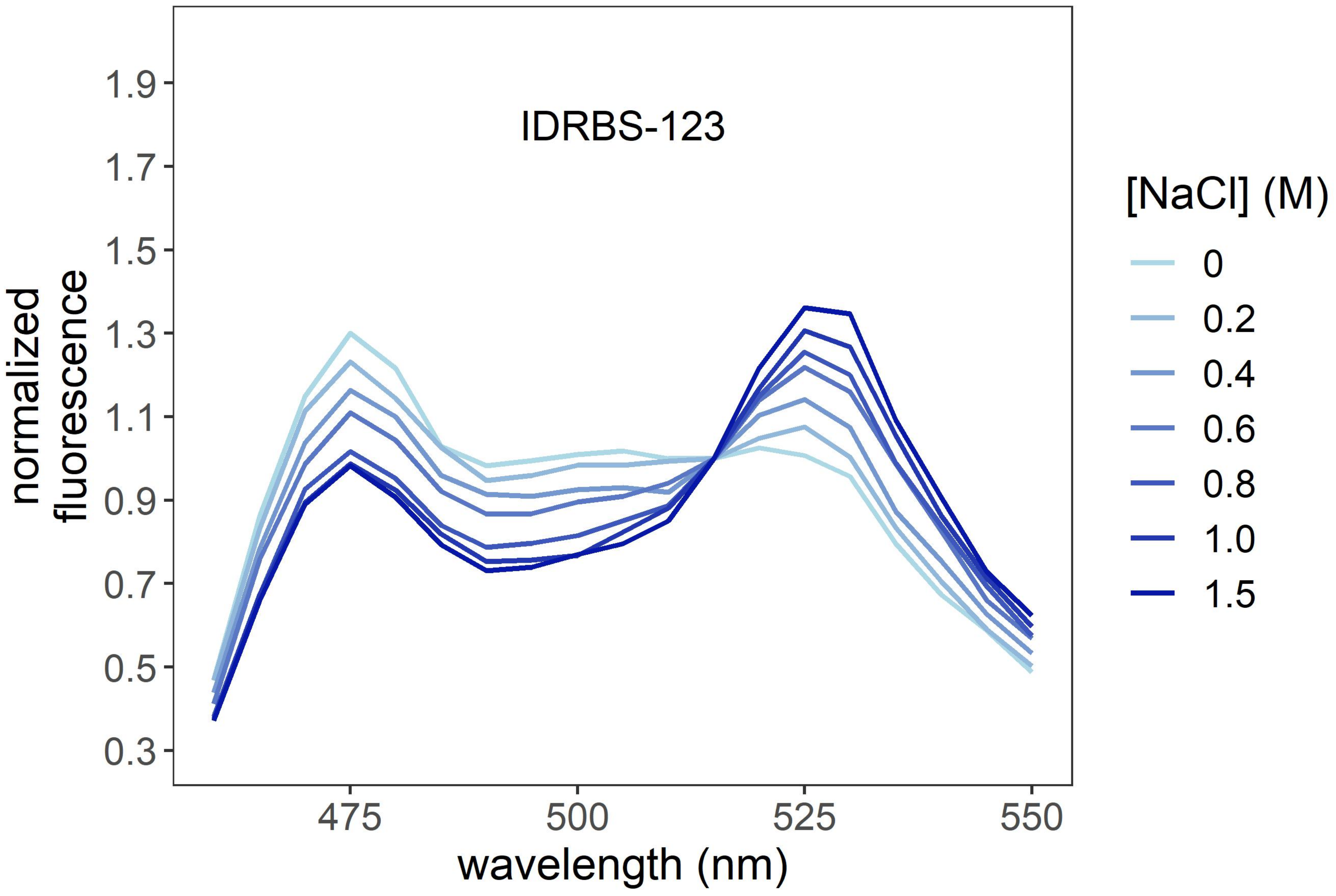
IDRBS-123
normalized fluorescence
1.9
1.7
1.5
1.3
1.1
0.9
0.7
0.5
0.3
475
500
525
550
wavelength (nm)
[NaCl] (M)
0
0.2
0.4
0.6
0.8
1.0
1.5

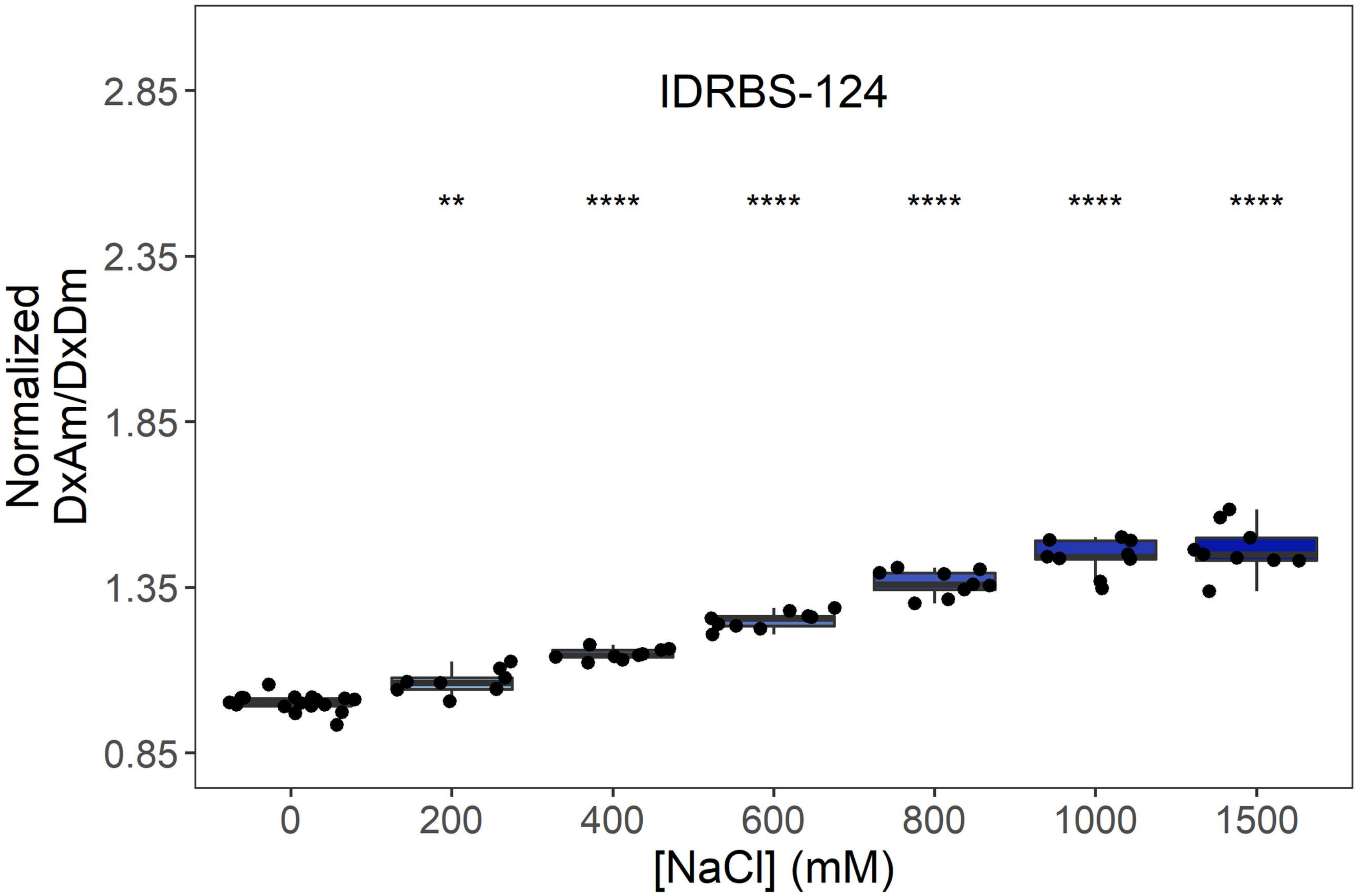
IDRBS-124
Normalized DxAm/DxDm
2.85
2.35
1.85
1.35
0.85
**
****
****
****
****
****
0
200
400
600
800
1000
1500
[NaCl] (mM)

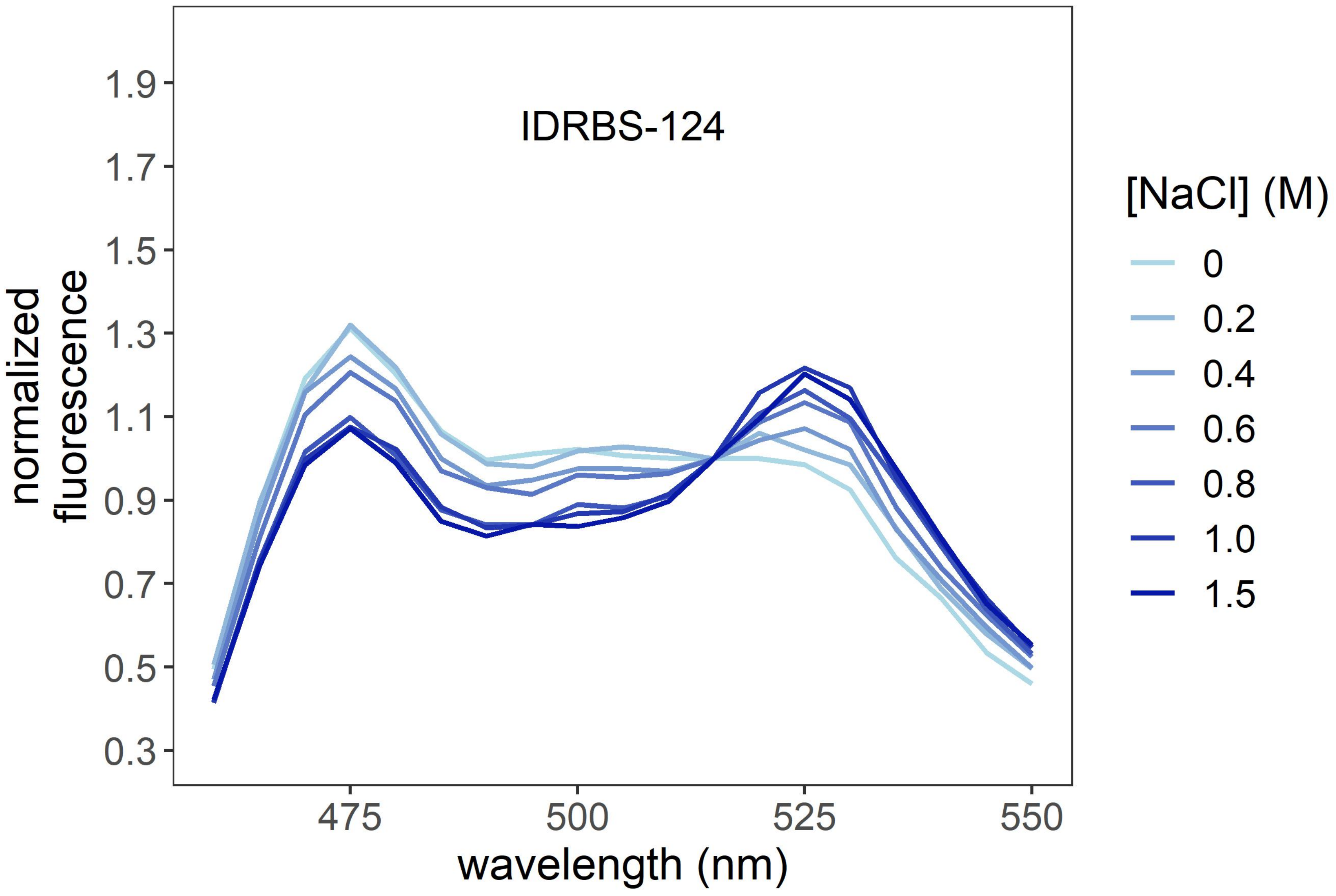
IDRBS-124
normalized fluorescence
1.9
1.7
1.5
1.3
1.1
0.9
0.7
0.5
0.3
475
500
525
550
wavelength (nm)
[NaCl] (M)
0
0.2
0.4
0.6
0.8
1.0
1.5

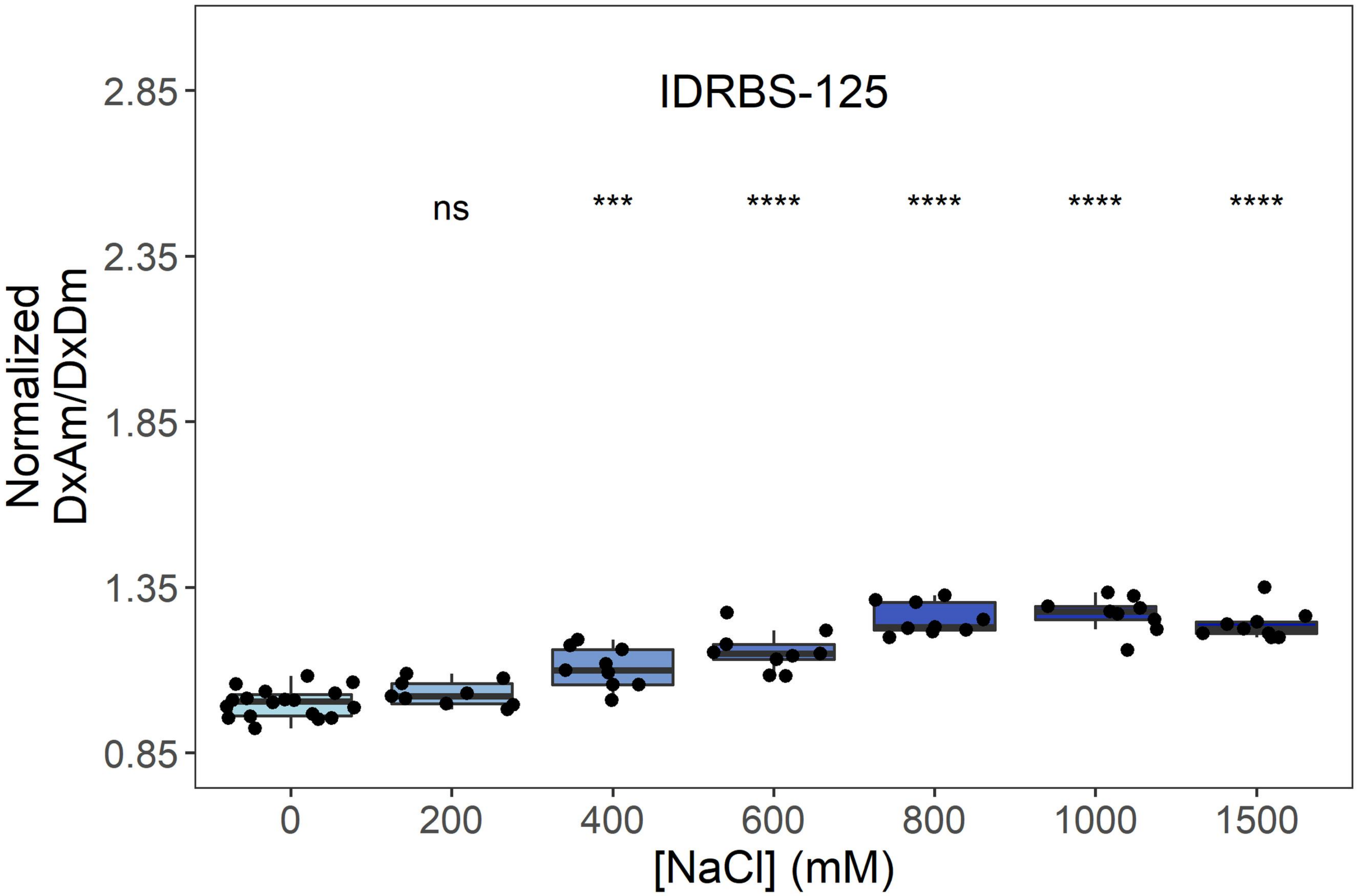
IDRBS-125
ns
***
****
****
****
****
Normalized DxAm/DxDm
2.85
2.35
1.85
1.35
0.85
0
200
400
600
800
1000
1500
[NaCl] (mM)

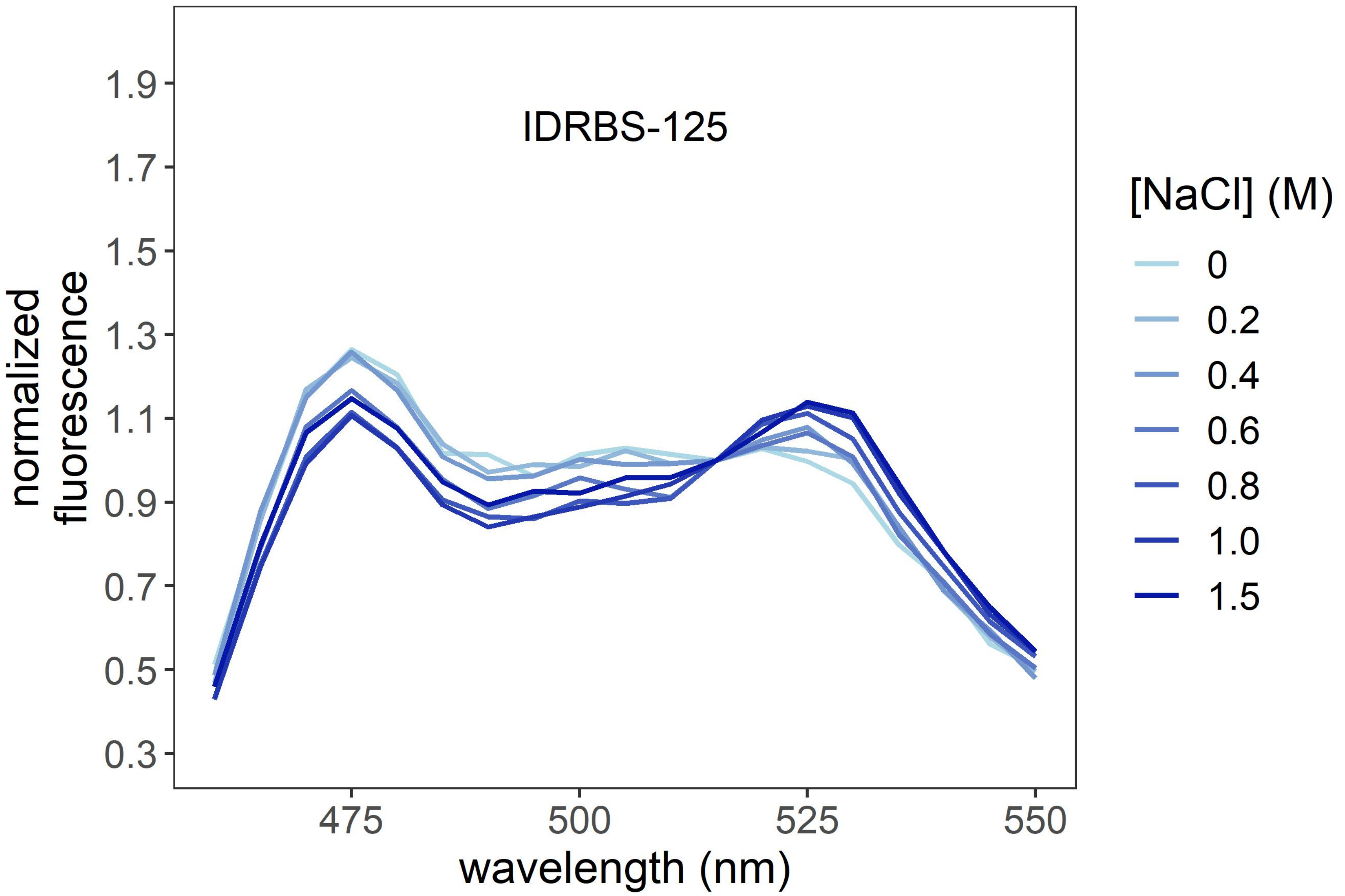
IDRBS-125
normalized fluorescence
1.9
1.7
1.5
1.3
1.1
0.9
0.7
0.5
0.3
475
500
525
550
wavelength (nm)
[NaCl] (M)
0
0.2
0.4
0.6
0.8
1.0
1.5

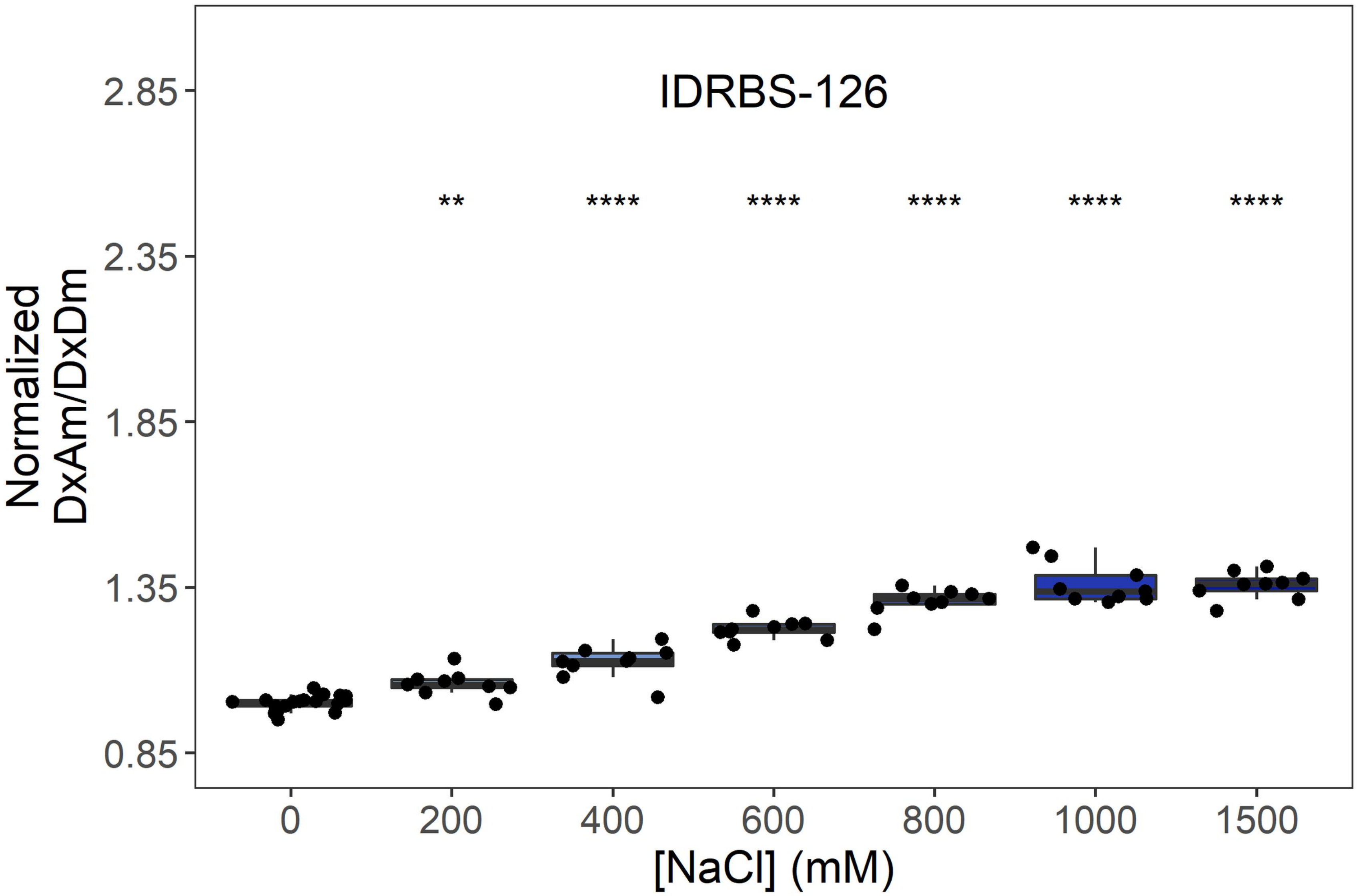

IDRBS-126
Normalized
DxAm/DxDm
2.85
2.35
1.85
1.35
0.85
**
****
****
****
****
****
0
200
400
600
800
1000
1500
[NaCl] (mM)


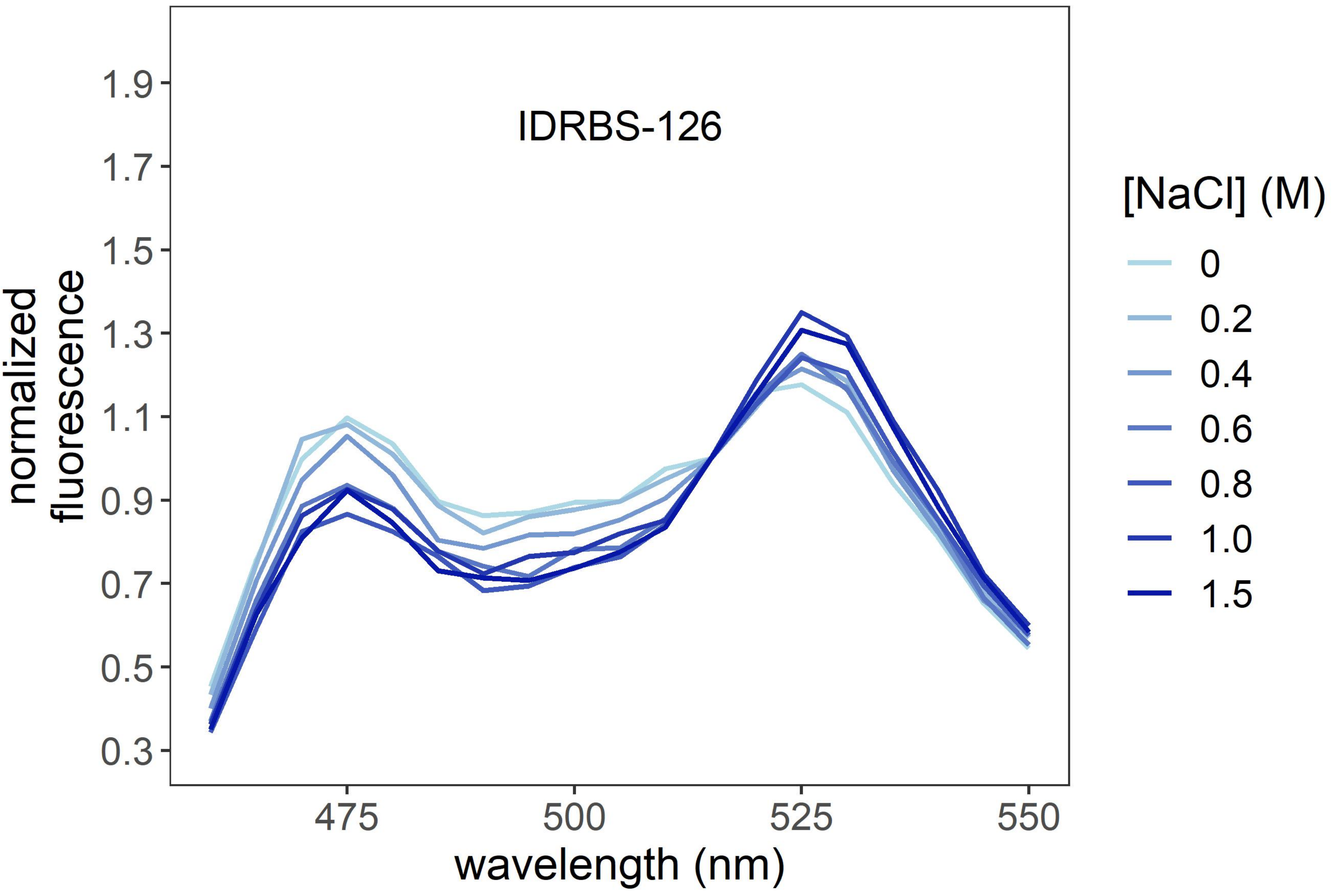

IDRBS-126
normalized
fluorescence
1.9
1.7
1.5
1.3
1.1
0.9
0.7
0.5
0.3
475
500
525
550
wavelength (nm)
[NaCl] (M)
0
0.2
0.4
0.6
0.8
1.0
1.5

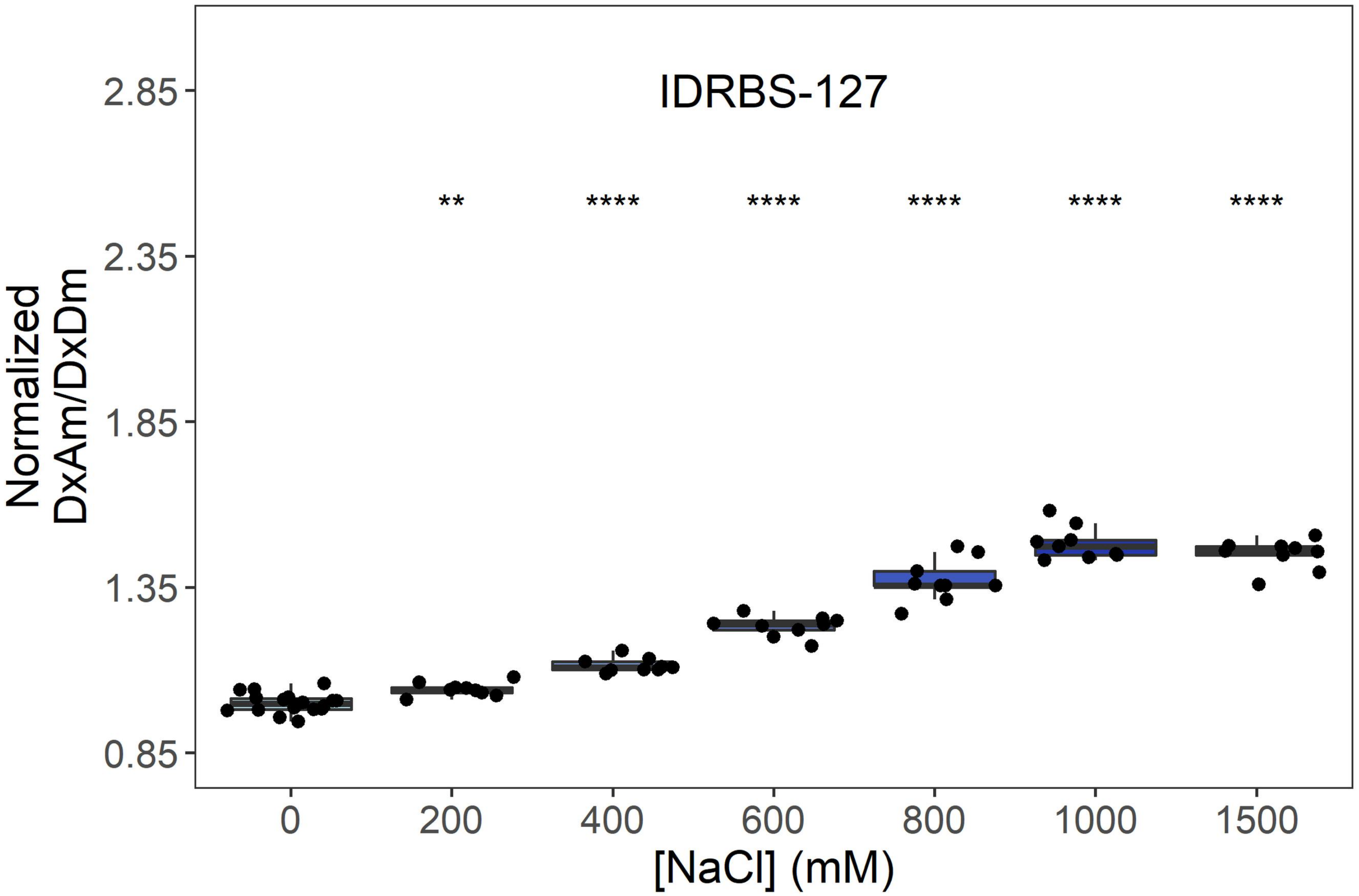
IDRBS-127
**
****
****
****
****
****
Normalized DxAm/DxDm
2.85
2.35
1.85
1.35
0.85
0
200
400
600
800
1000
1500
[NaCl] (mM)

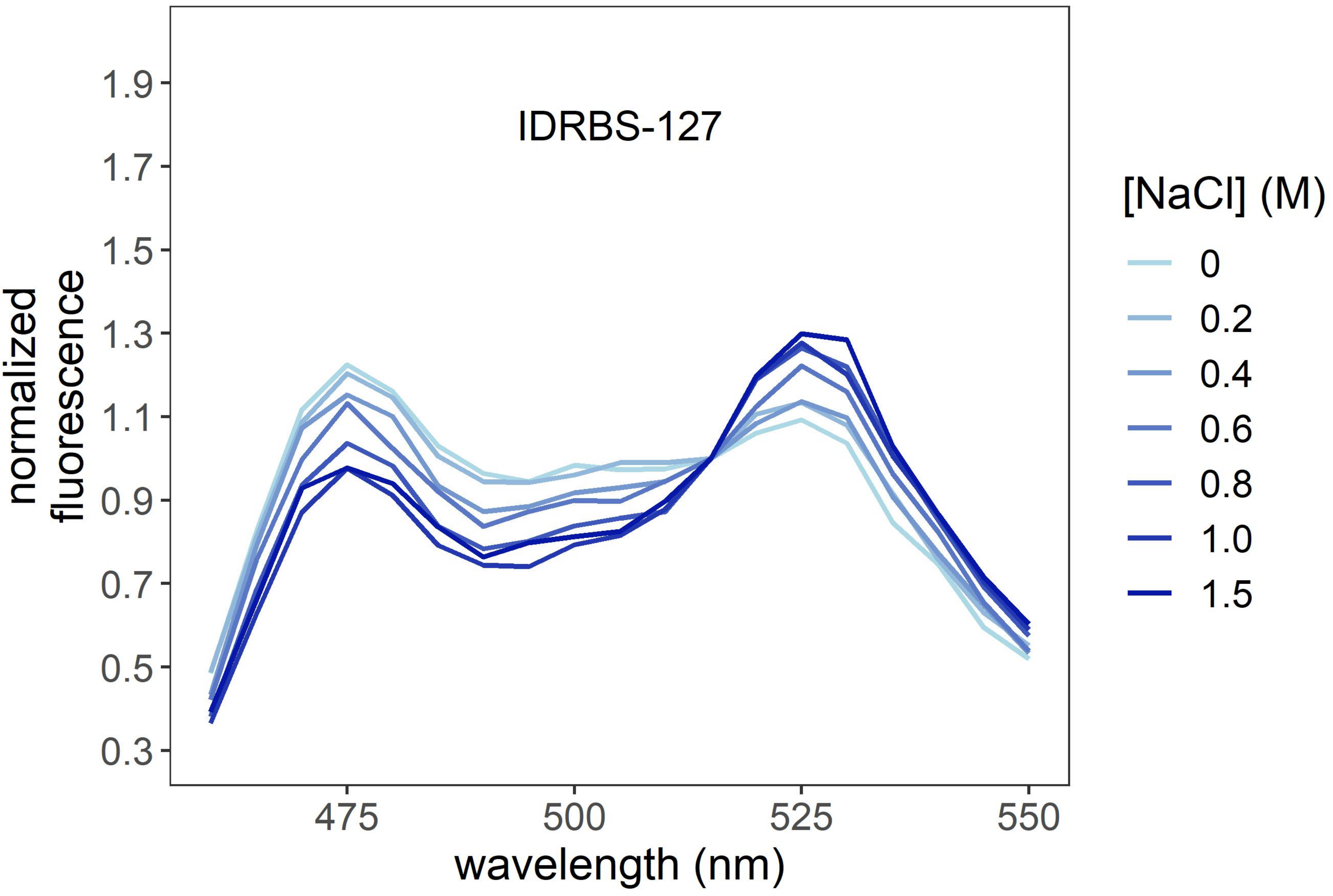
IDRBS-127
normalized fluorescence
1.9
1.7
1.5
1.3
1.1
0.9
0.7
0.5
0.3
475
500
525
550
wavelength (nm)
[NaCl] (M)
0
0.2
0.4
0.6
0.8
1.0
1.5

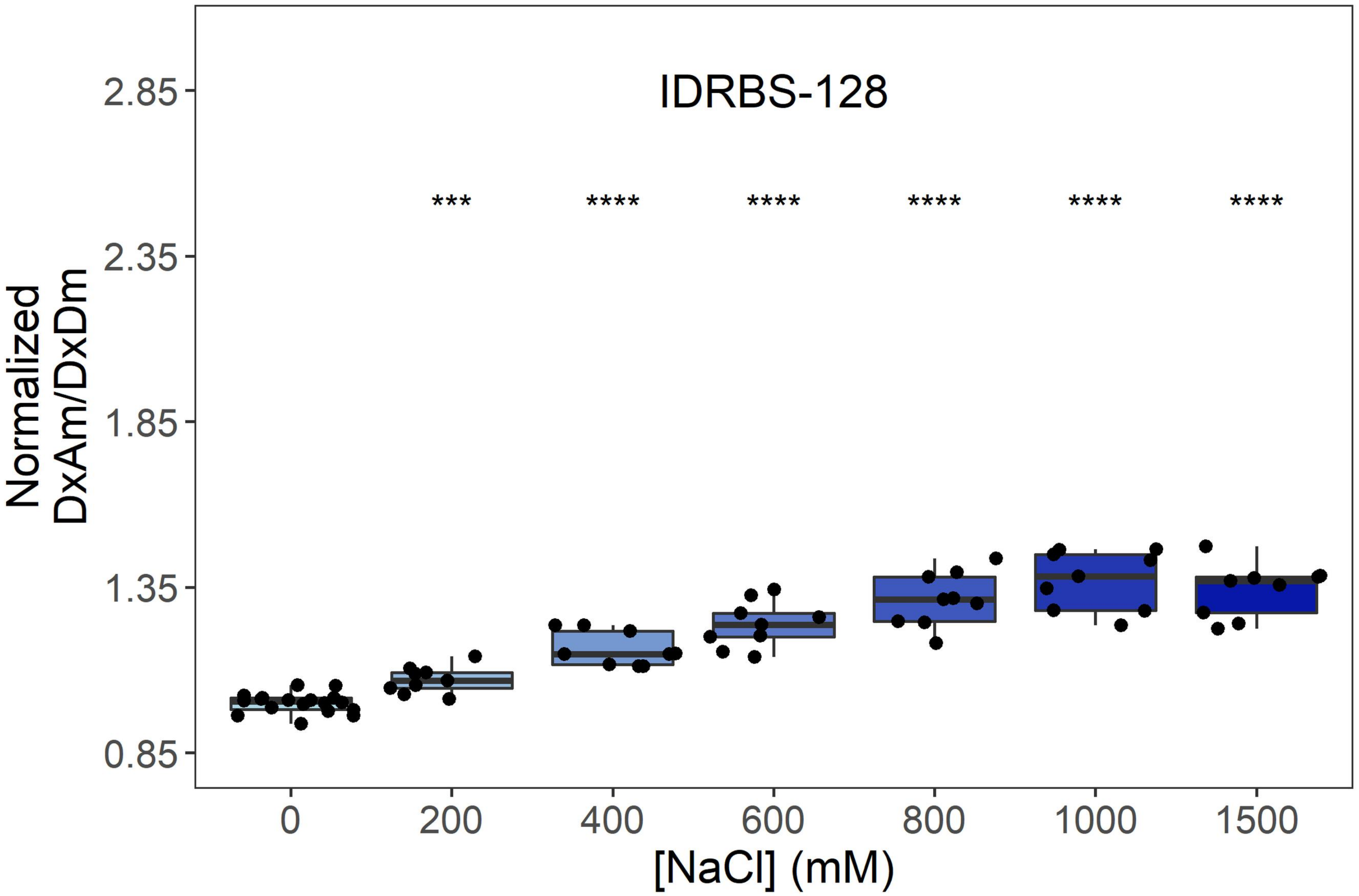
IDRBS-128
***
****
****
****
****
****
Normalized DxAm/DxDm
2.85
2.35
1.85
1.35
0.85
0
200
400
600
800
1000
1500
[NaCl] (mM)

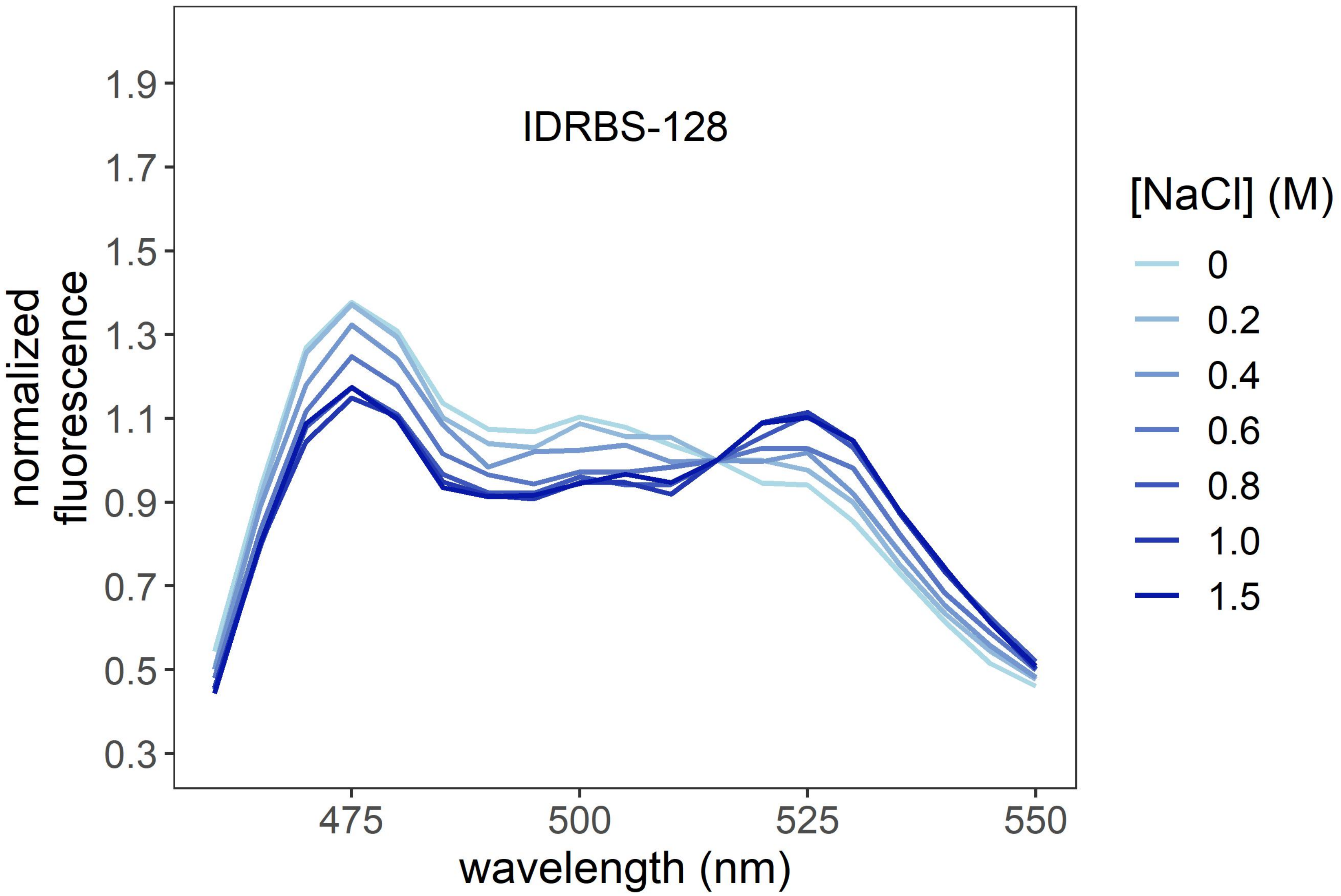
IDRBS-128
normalized fluorescence
1.9
1.7
1.5
1.3
1.1
0.9
0.7
0.5
0.3
475
500
525
550
wavelength (nm)
[NaCl] (M)
0
0.2
0.4
0.6
0.8
1.0
1.5

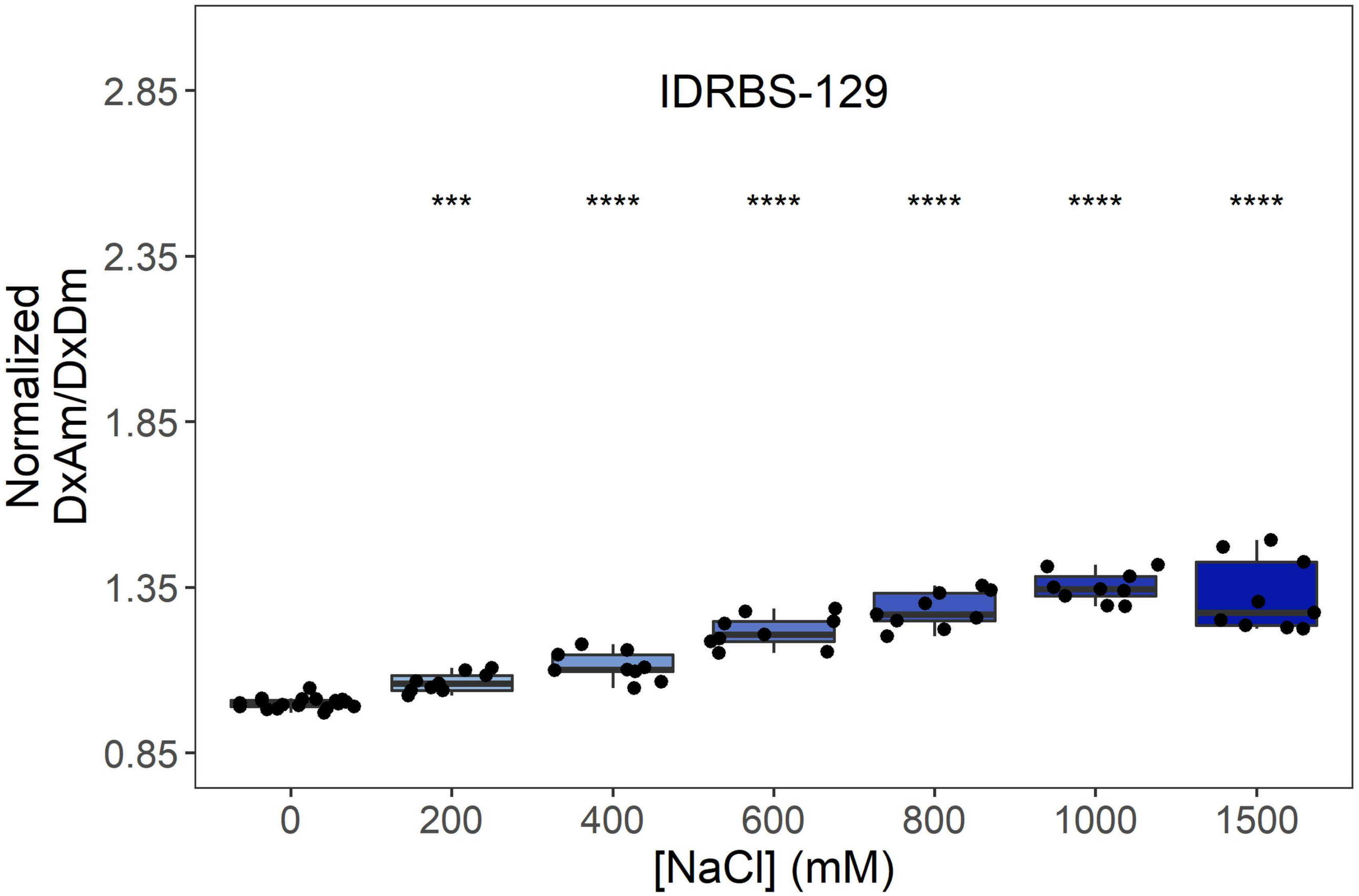
IDRBS-129
Normalized DxAm/DxDm
2.85
2.35
1.85
1.35
0.85
***
****
****
****
****
****
0
200
400
600
800
1000
1500
[NaCl] (mM)

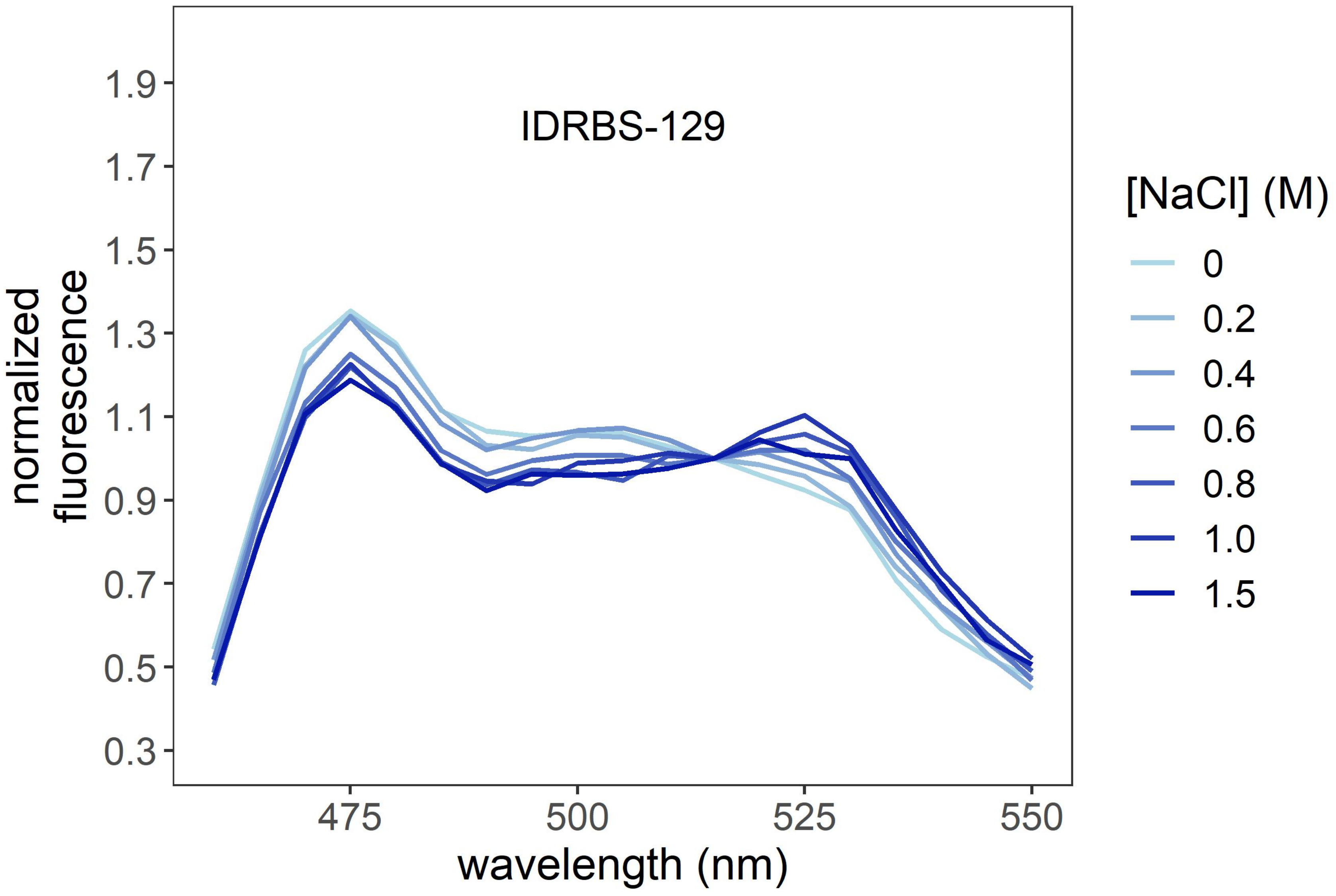
IDRBS-129
normalized fluorescence
1.9
1.7
1.5
1.3
1.1
0.9
0.7
0.5
0.3
475
500
525
550
wavelength (nm)
[NaCl] (M)
0
0.2
0.4
0.6
0.8
1.0
1.5

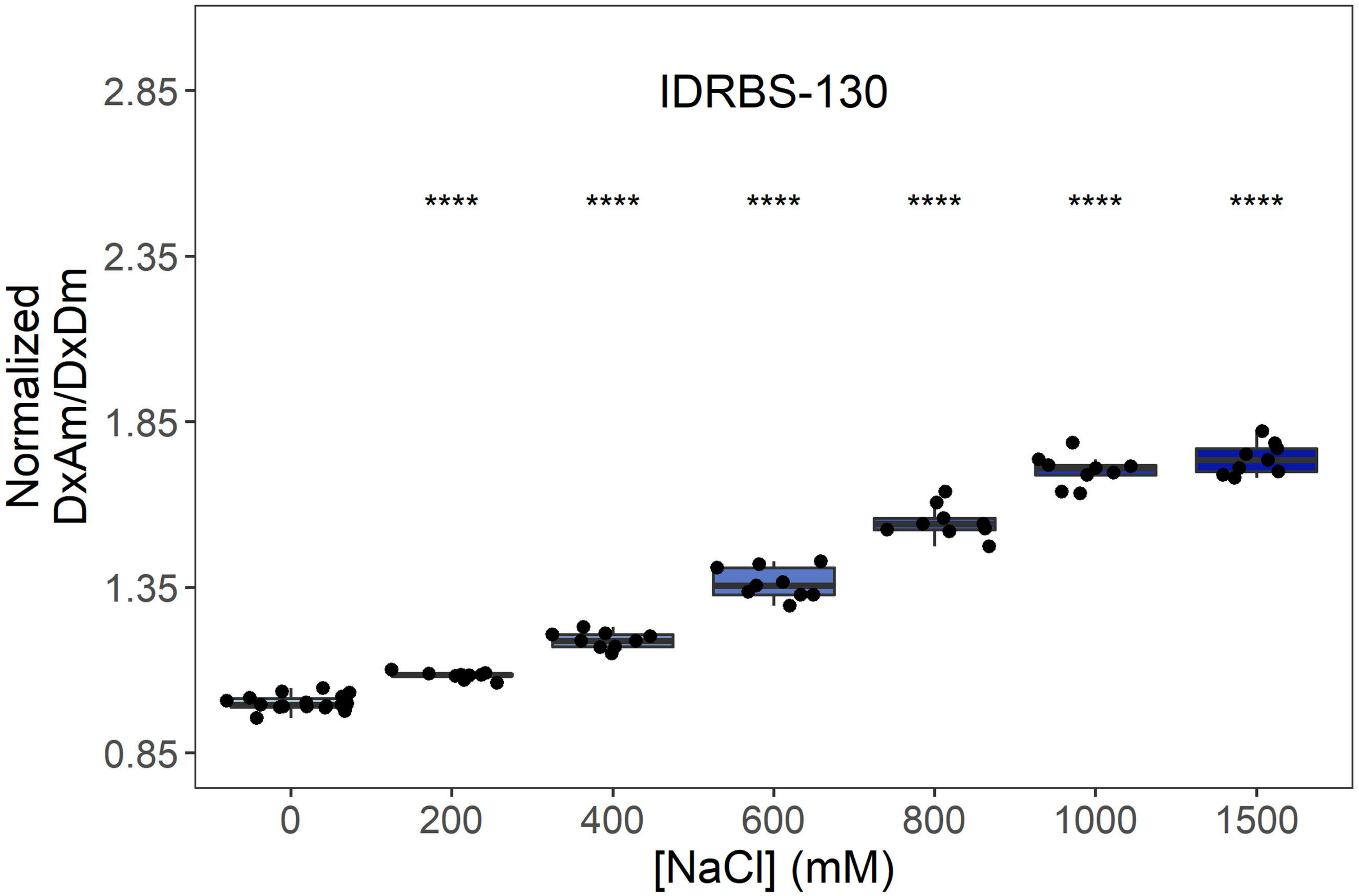
IDRBS-130
****
****
****
****
****
****
2.85
2.35
1.85
1.35
0.85
Normalized
DxAm/DxDm
0
200
400
600
800
1000
1500
[NaCl] (mM)

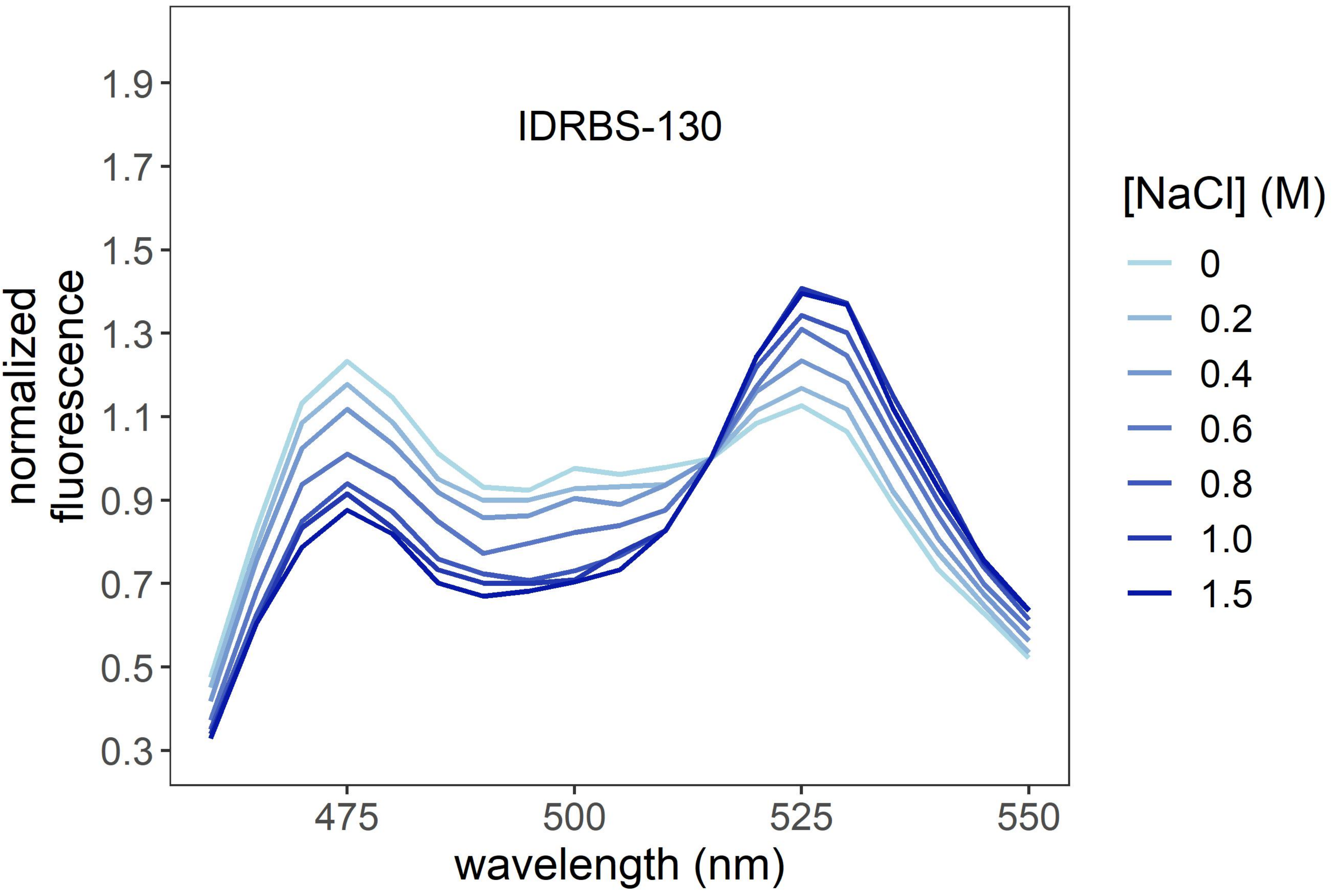
IDRBS-130
1.9
1.7
1.5
1.3
1.1
0.9
0.7
0.5
0.3
normalized
fluorescence
475
500
525
550
wavelength (nm)
[NaCl] (M)
0
0.2
0.4
0.6
0.8
1.0
1.5

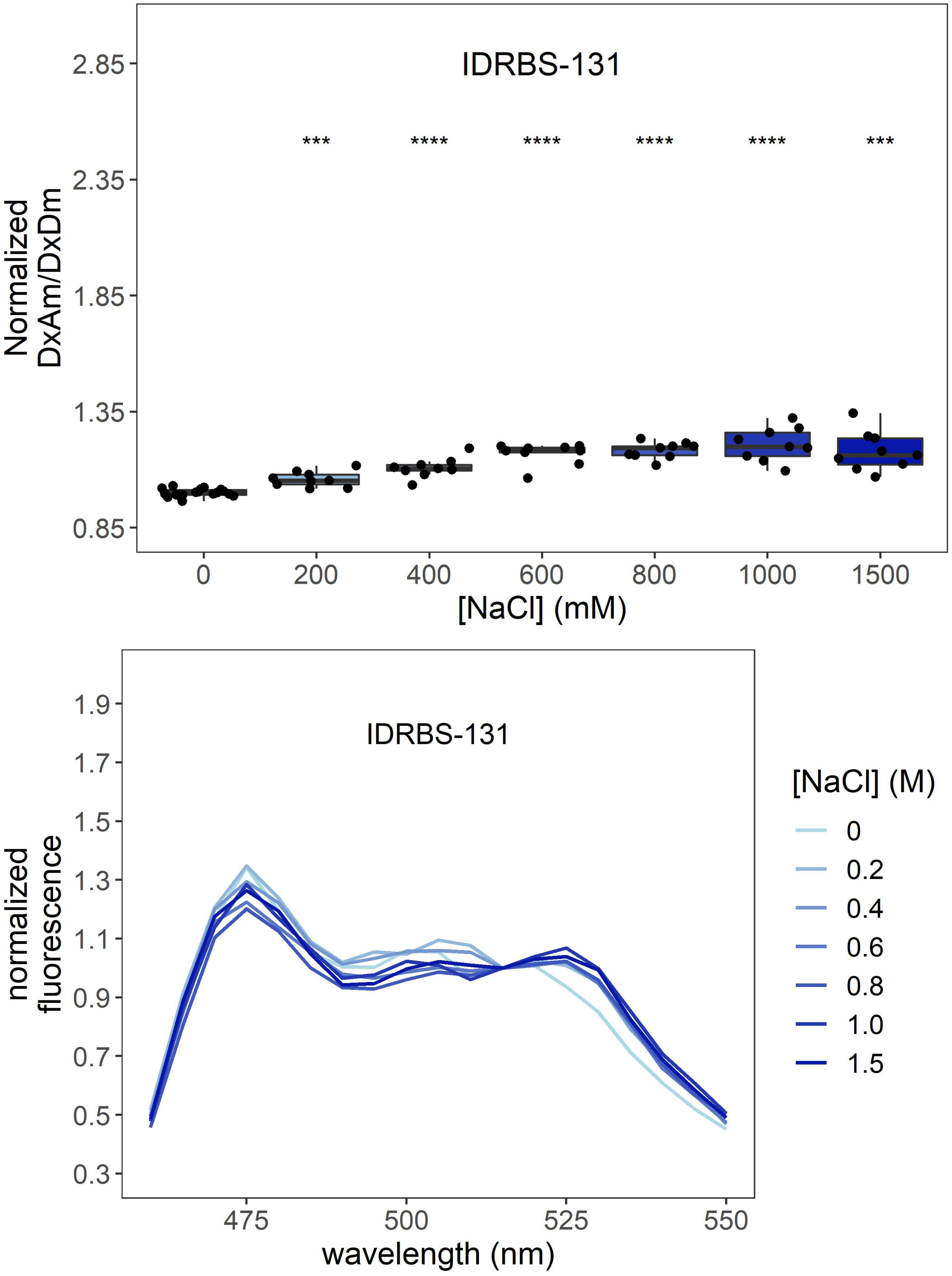
IDRBS-131
***
****
****
****
****
***
Normalized
DxAm/DxDm
2.85
2.35
1.85
1.35
0.85
0
200
400
600
800
1000
1500
[NaCl] (mM)
IDRBS-131
normalized
fluorescence
1.9
1.7
1.5
1.3
1.1
0.9
0.7
0.5
0.3
475
500
525
550
wavelength (nm)
[NaCl] (M)
0
0.2
0.4
0.6
0.8
1.0
1.5

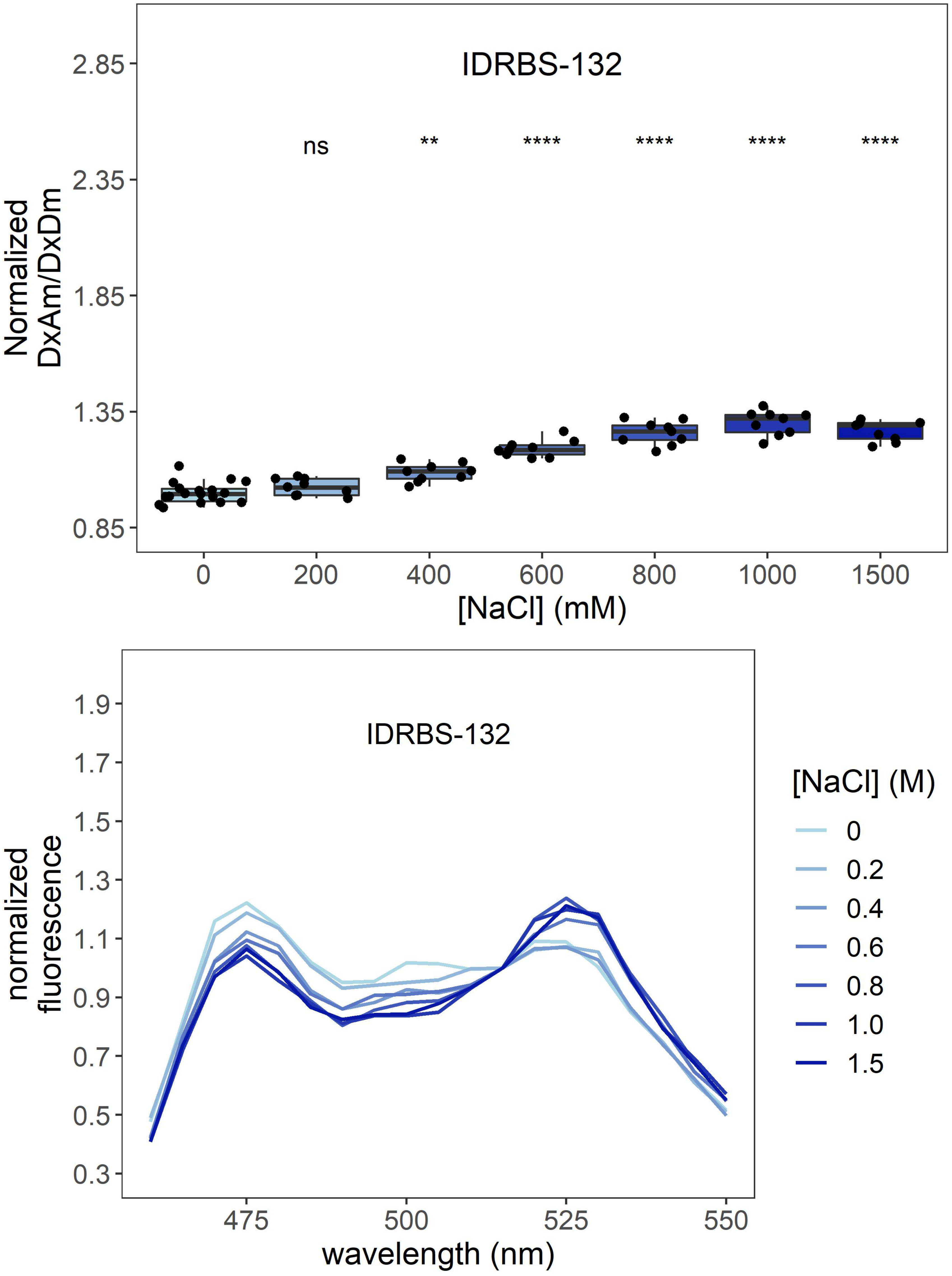

IDRBS-132
ns
**
****
****
****
****
Normalized DxAm/DxDm
2.85
2.35
1.85
1.35
0.85
0
200
400
600
800
1000
1500
[NaCl] (mM)
IDRBS-132
normalized fluorescence
1.9
1.7
1.5
1.3
1.1
0.9
0.7
0.5
0.3
475
500
525
550
wavelength (nm)
[NaCl] (M)
0
0.2
0.4
0.6
0.8
1.0
1.5

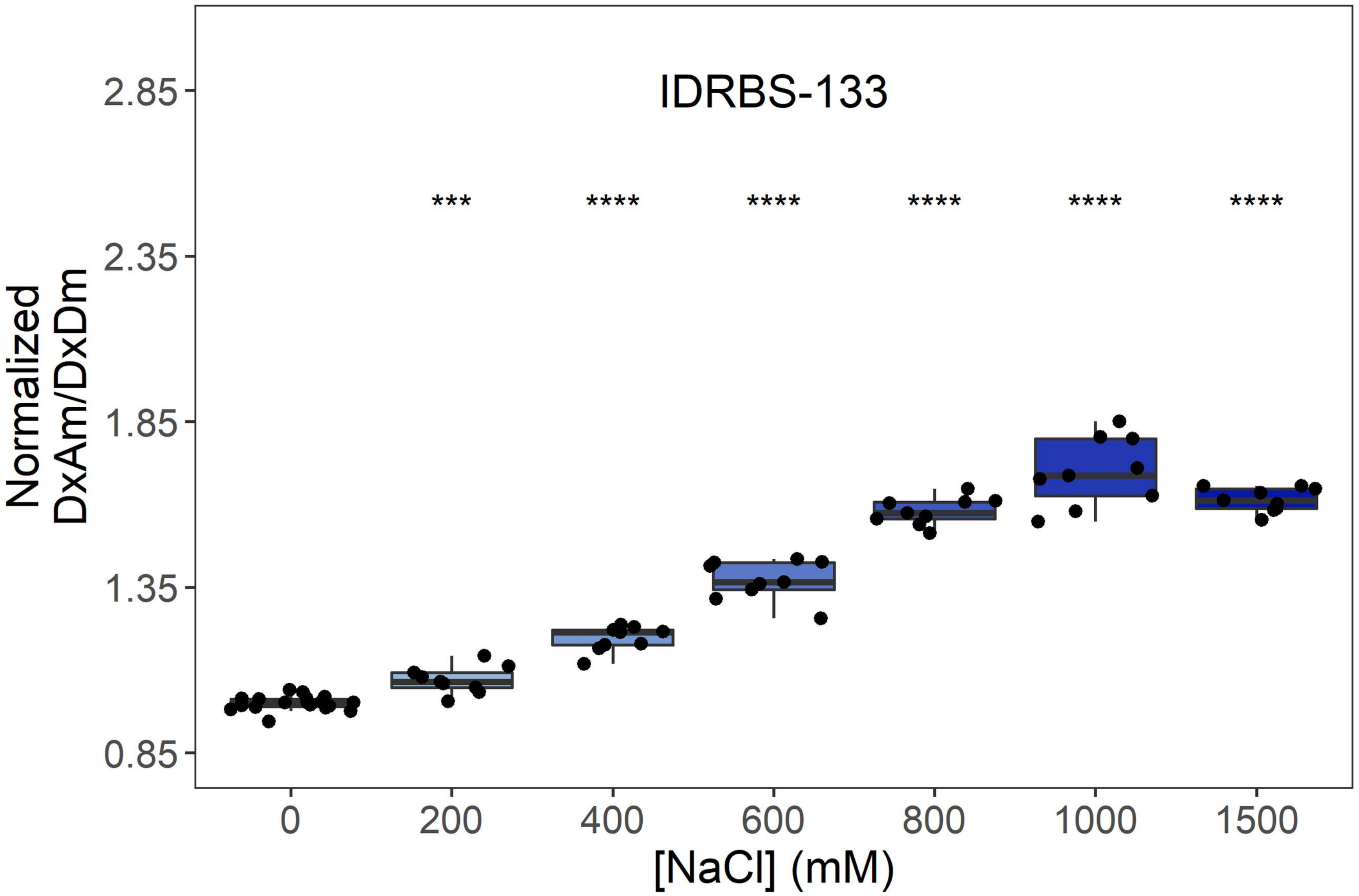
IDRBS-133
Normalized DxAm/DxDm
2.85
2.35
1.85
1.35
0.85
***
****
****
****
****
****
0
200
400
600
800
1000
1500
[NaCl] (mM)

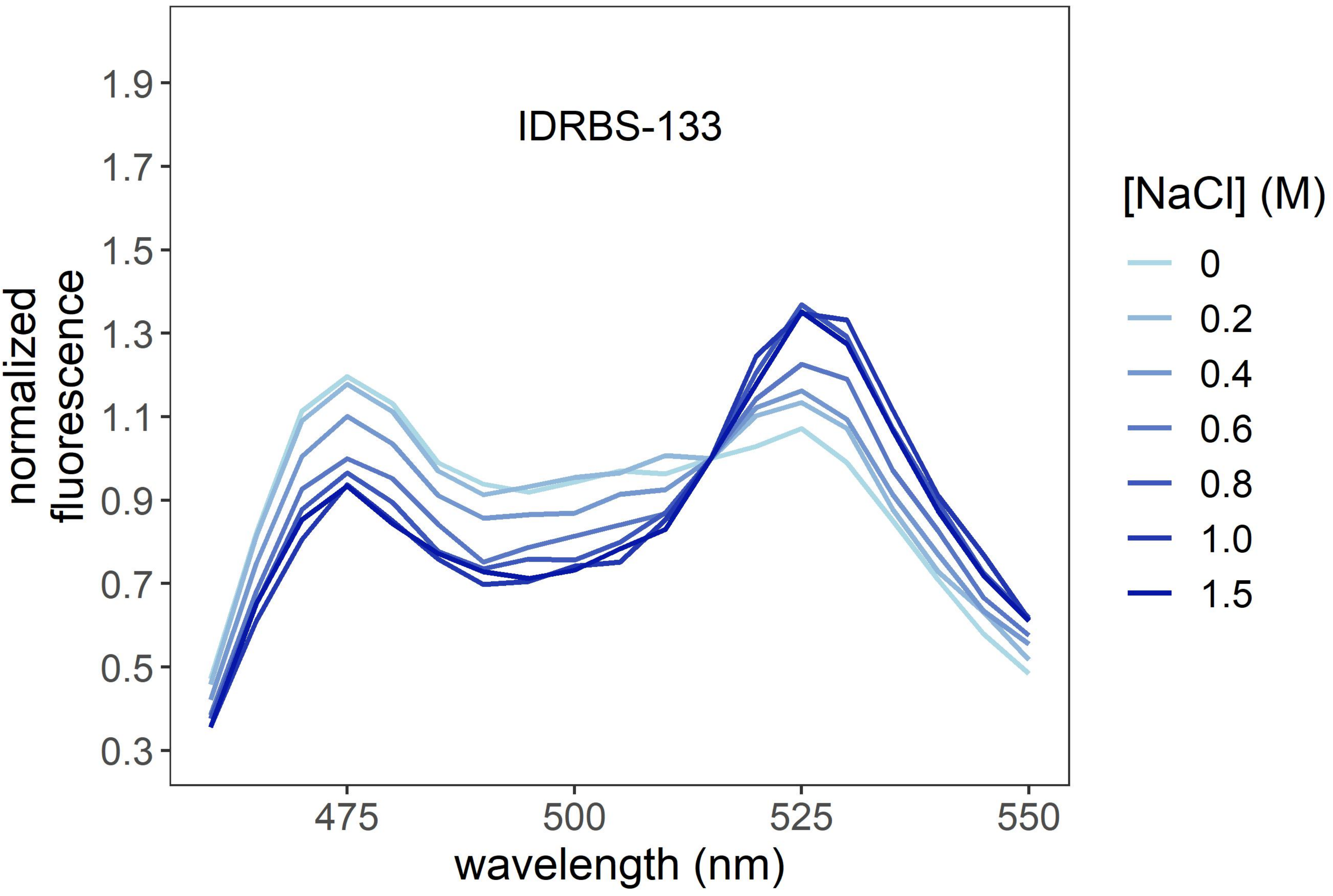
IDRBS-133
normalized fluorescence
1.9
1.7
1.5
1.3
1.1
0.9
0.7
0.5
0.3
475
500
525
550
wavelength (nm)
[NaCl] (M)
0
0.2
0.4
0.6
0.8
1.0
1.5

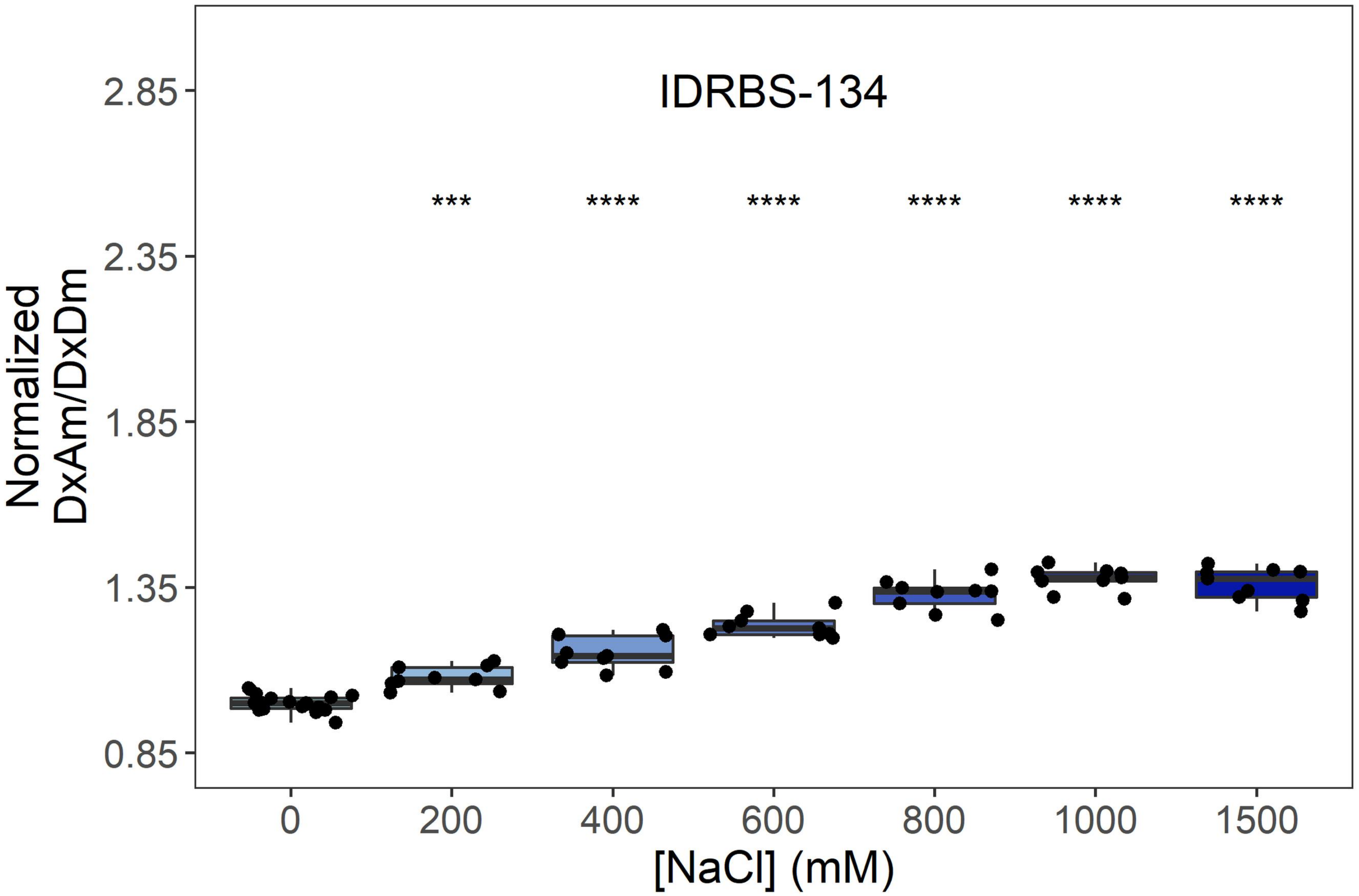

IDRBS-134
***
****
****
****
****
****
2.85
2.35
1.85
1.35
0.85
Normalized DxAm/DxDm
0
200
400
600
800
1000
1500
[NaCl] (mM)


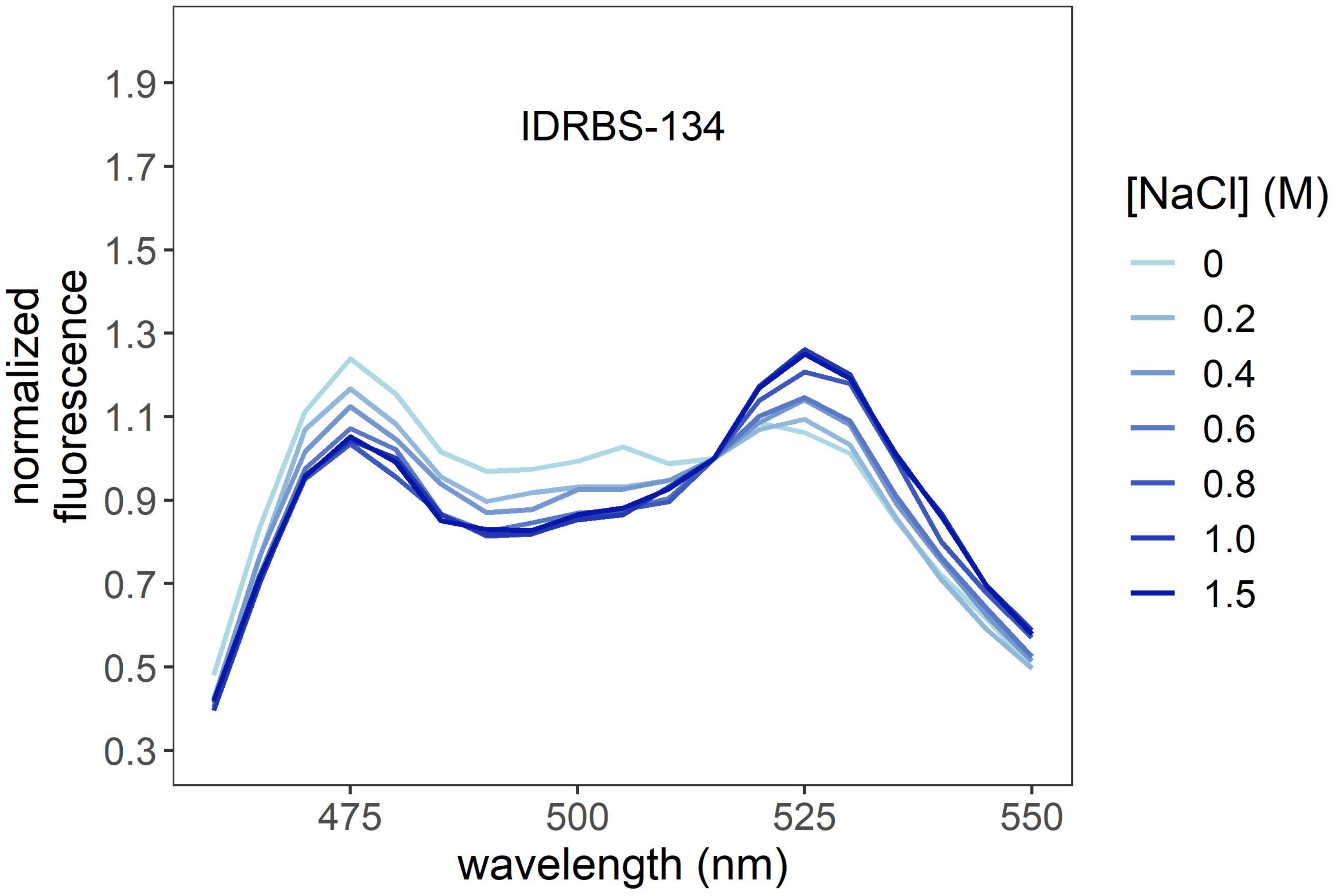

IDRBS-134
1.9
1.7
1.5
1.3
1.1
0.9
0.7
0.5
0.3
normalized fluorescence
475
500
525
550
wavelength (nm)
[NaCl] (M)
0
0.2
0.4
0.6
0.8
1.0
1.5

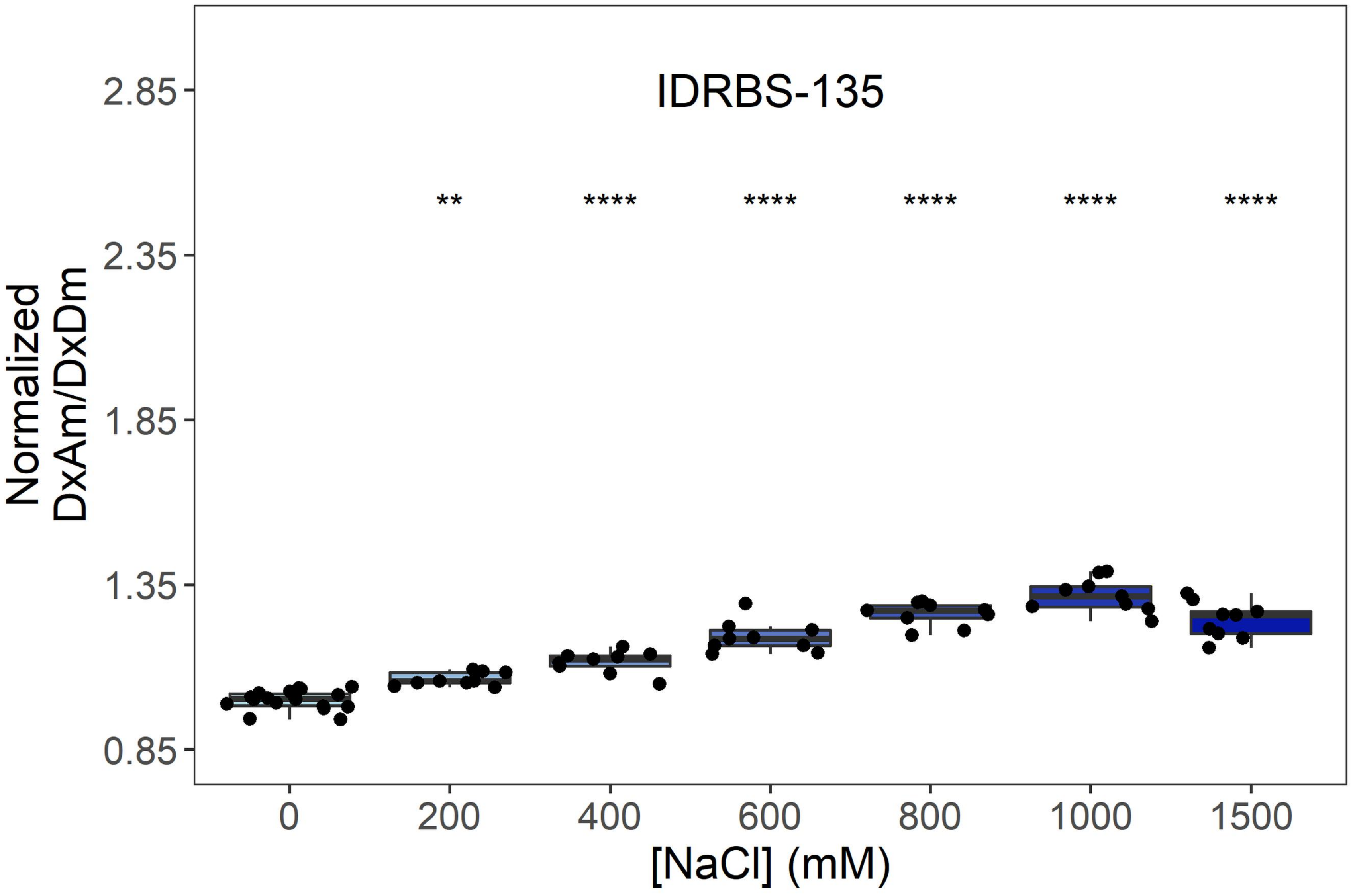

IDRBS-135
**
****
****
****
****
****
2.85
2.35
1.85
1.35
0.85
Normalized DxAm/DxDm
0
200
400
600
800
1000
1500
[NaCl] (mM)


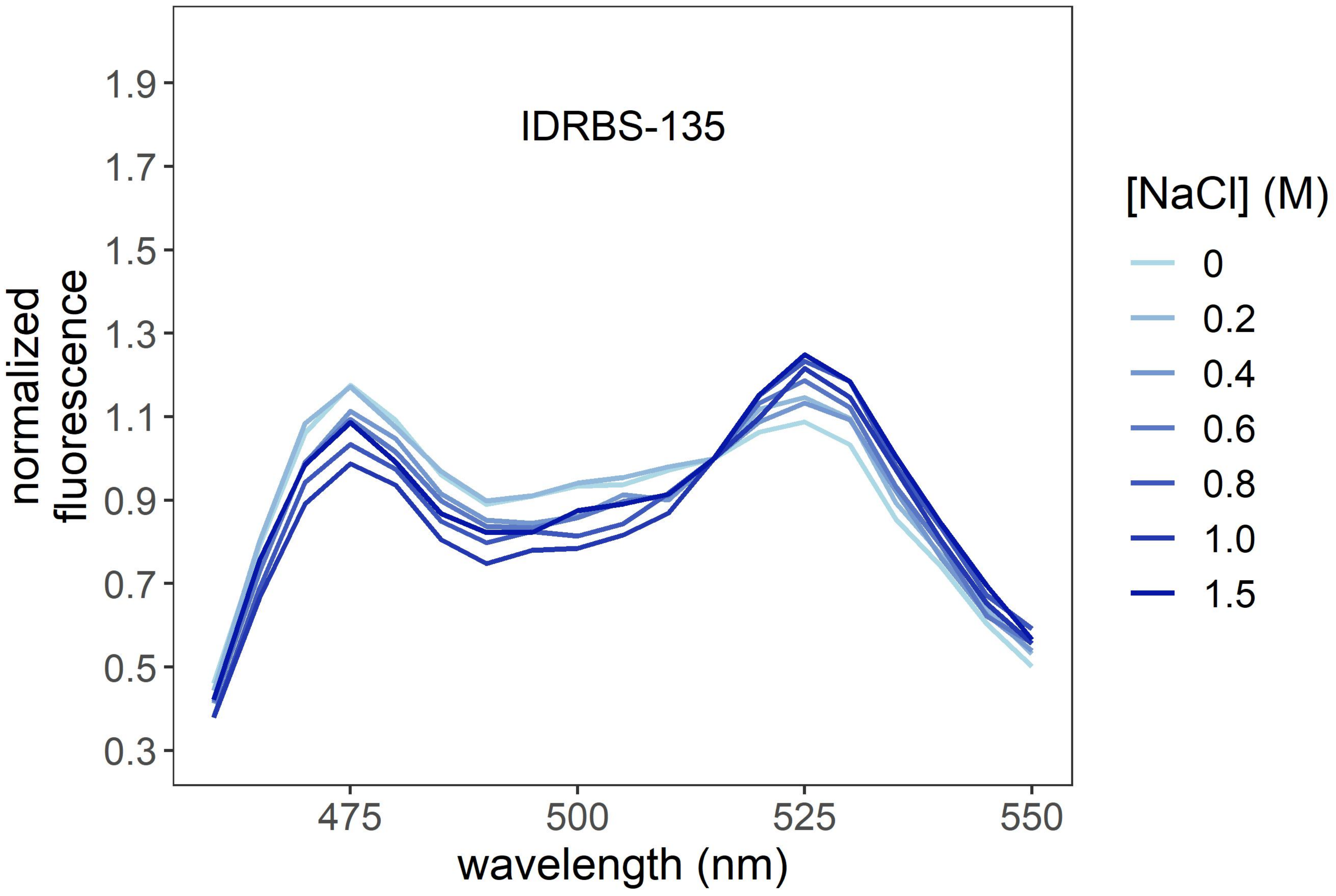

IDRBS-135
1.9
1.7
1.5
1.3
1.1
0.9
0.7
0.5
0.3
normalized fluorescence
475
500
525
550
wavelength (nm)
[NaCl] (M)
0
0.2
0.4
0.6
0.8
1.0
1.5

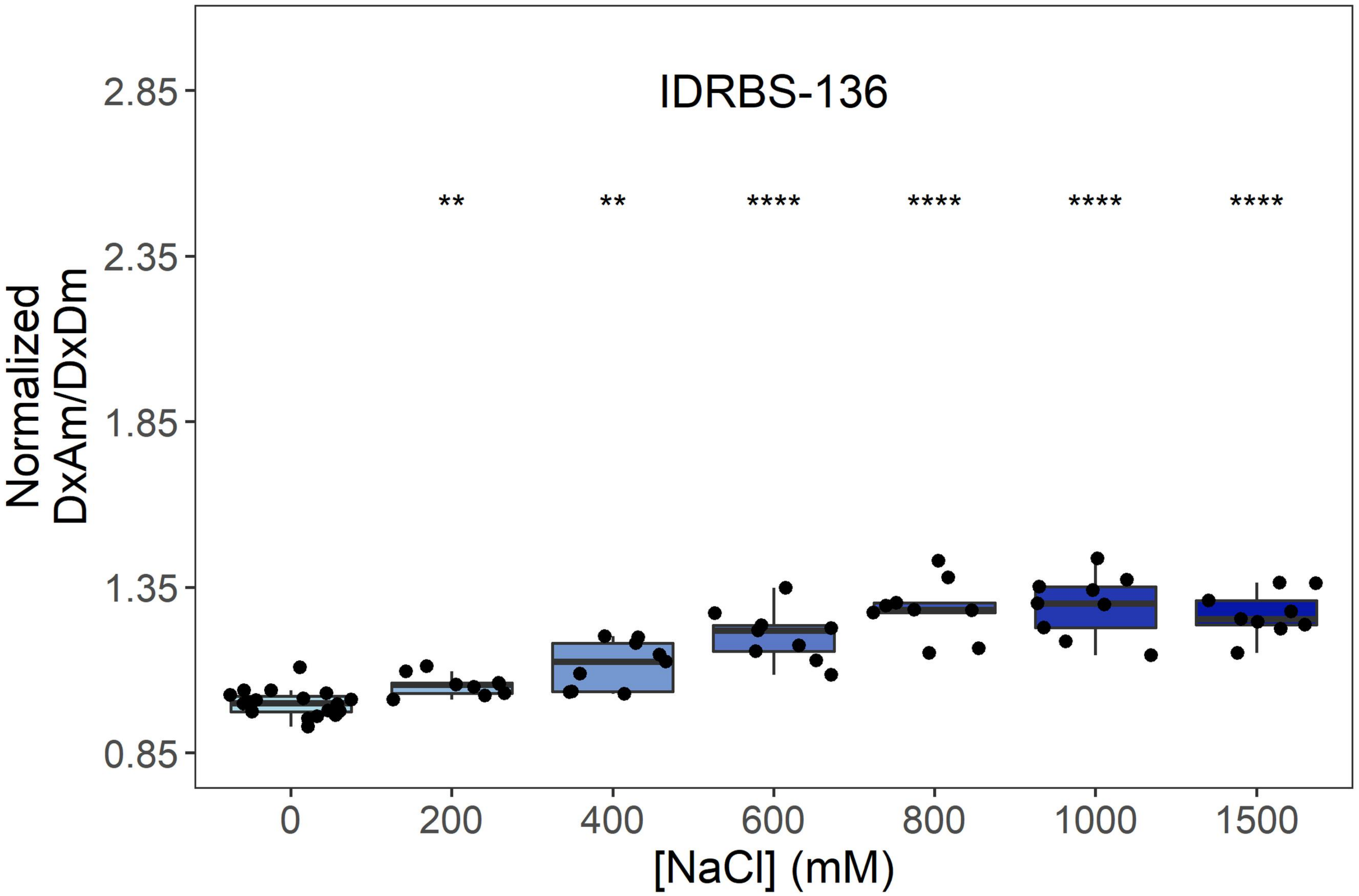

IDRBS-136
Normalized DxAm/DxDm
2.85
2.35
1.85
1.35
0.85
**
**
****
****
****
****
0
200
400
600
800
1000
1500
[NaCl] (mM)


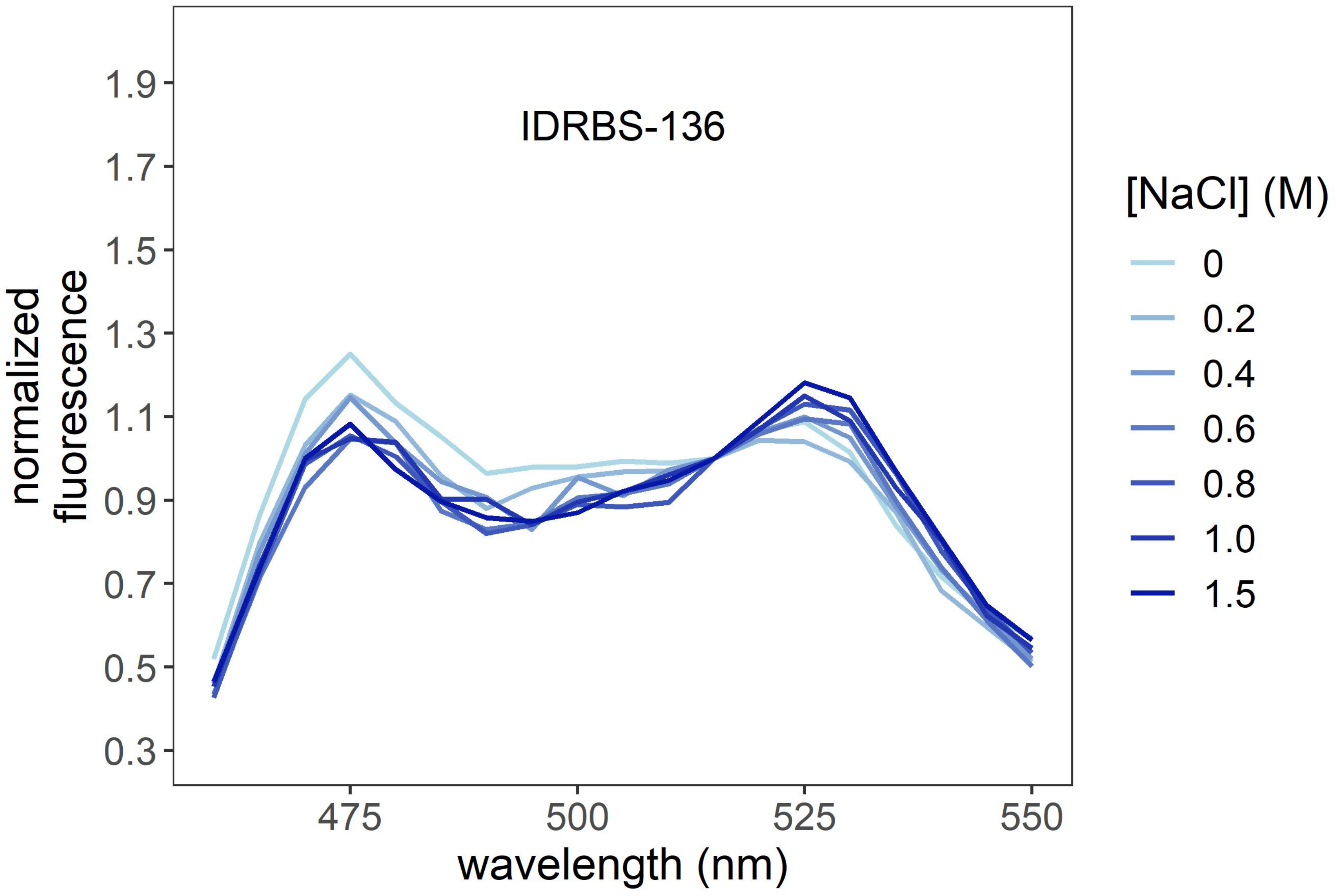

IDRBS-136
normalized fluorescence
1.9
1.7
1.5
1.3
1.1
0.9
0.7
0.5
0.3
475
500
525
550
wavelength (nm)
[NaCl] (M)
0
0.2
0.4
0.6
0.8
1.0
1.5

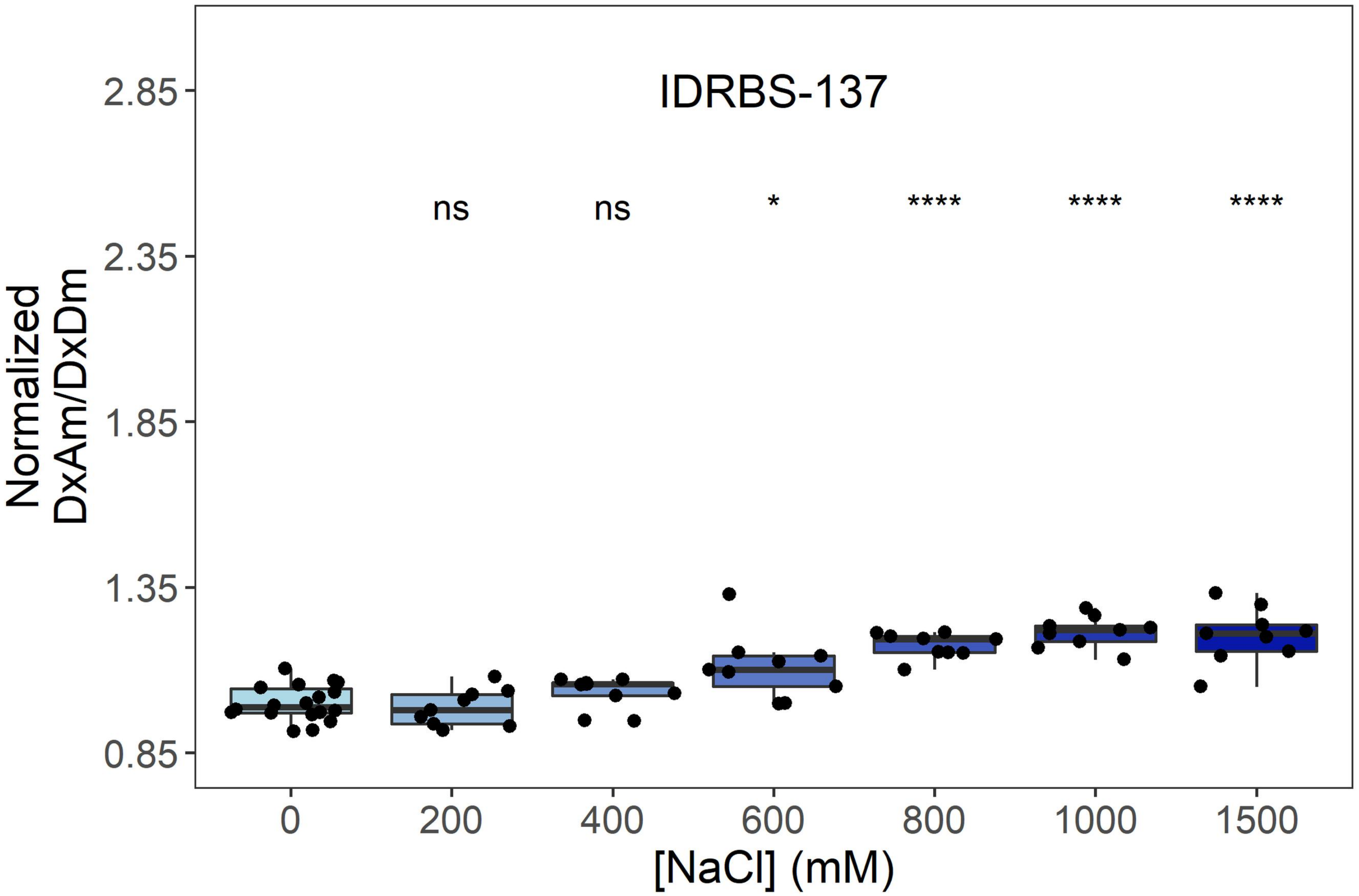
IDRBS-137
ns
ns
*
****
****
****
Normalized DxAm/DxDm
2.85
2.35
1.85
1.35
0.85
0
200
400
600
800
1000
1500
[NaCl] (mM)

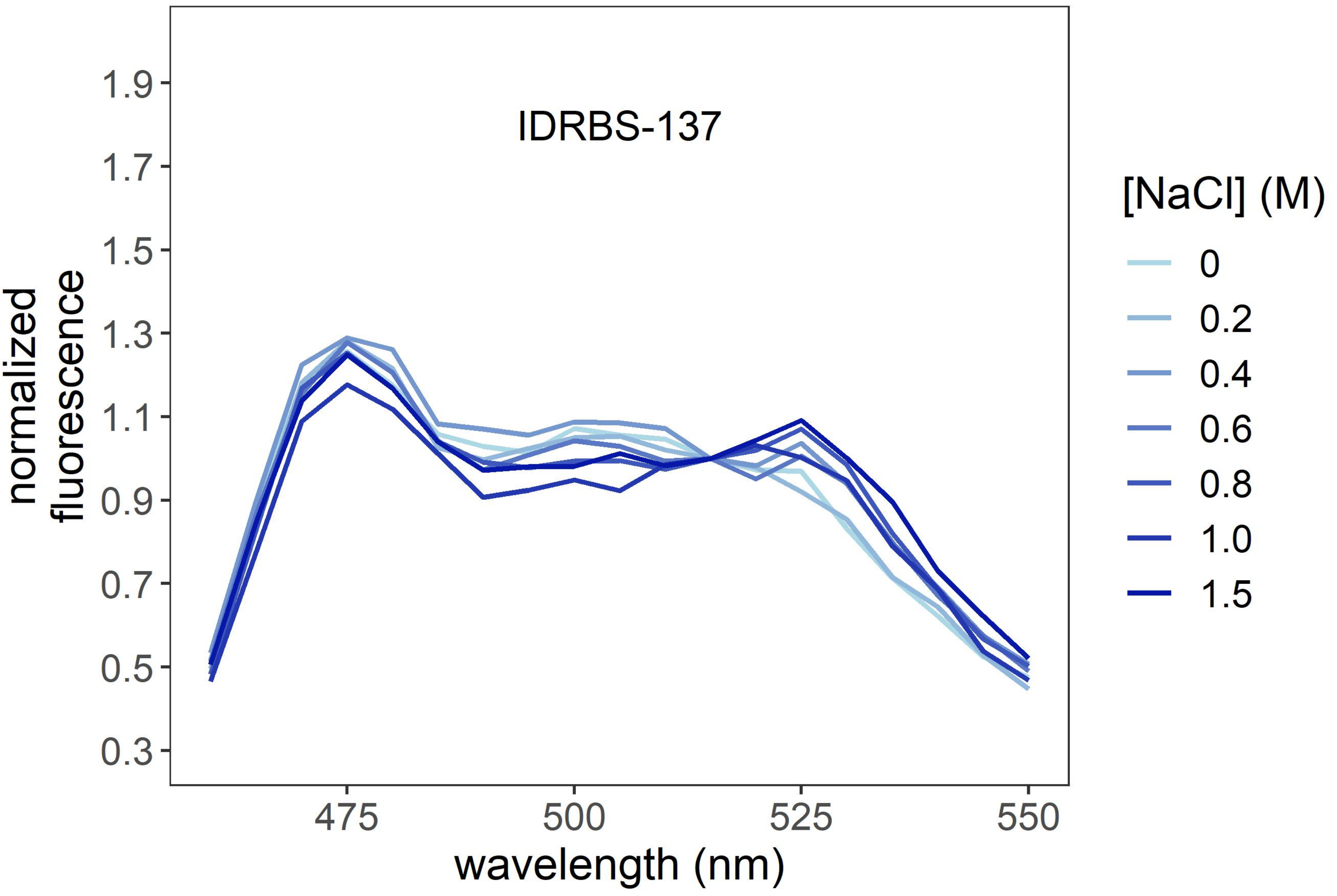
IDRBS-137
normalized fluorescence
1.9
1.7
1.5
1.3
1.1
0.9
0.7
0.5
0.3
475
500
525
550
wavelength (nm)
[NaCl] (M)
0
0.2
0.4
0.6
0.8
1.0
1.5

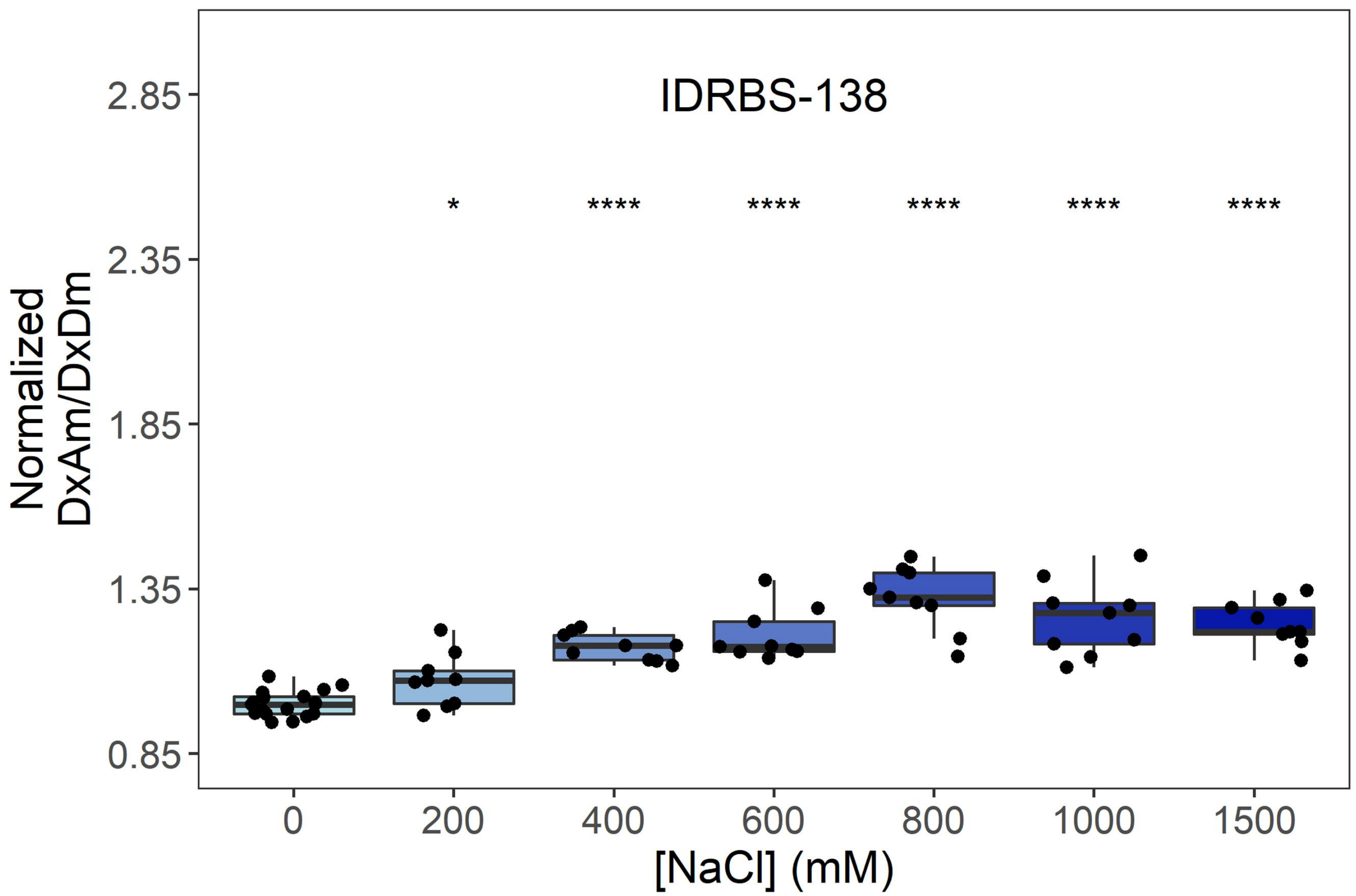

IDRBS-138
Normalized DxAm/DxDm
2.85
2.35
1.85
1.35
0.85
*
****
****
****
****
****
0
200
400
600
800
1000
1500
[NaCl] (mM)


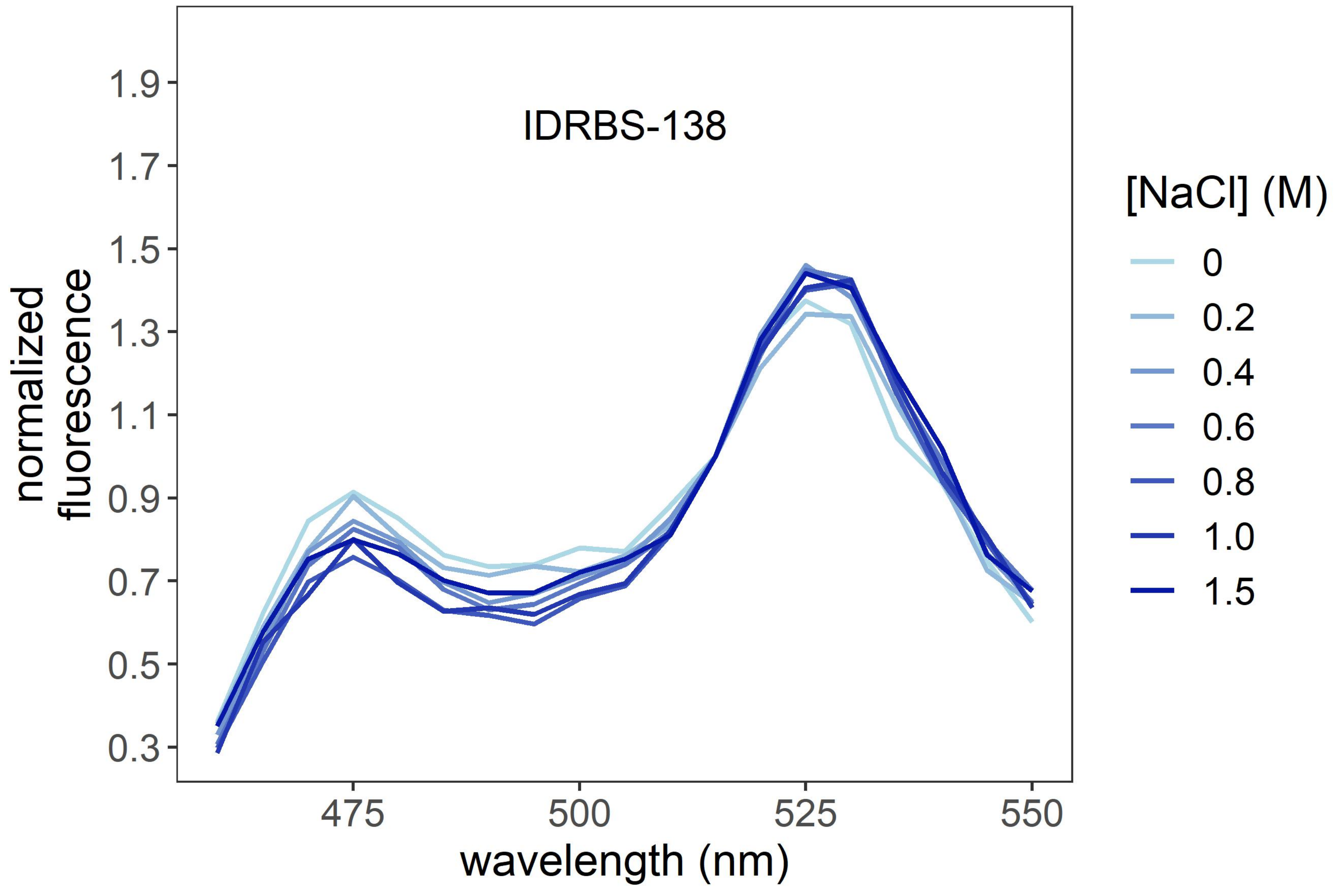

IDRBS-138
normalized fluorescence
1.9
1.7
1.5
1.3
1.1
0.9
0.7
0.5
0.3
475
500
525
550
wavelength (nm)
[NaCl] (M)
0
0.2
0.4
0.6
0.8
1.0
1.5

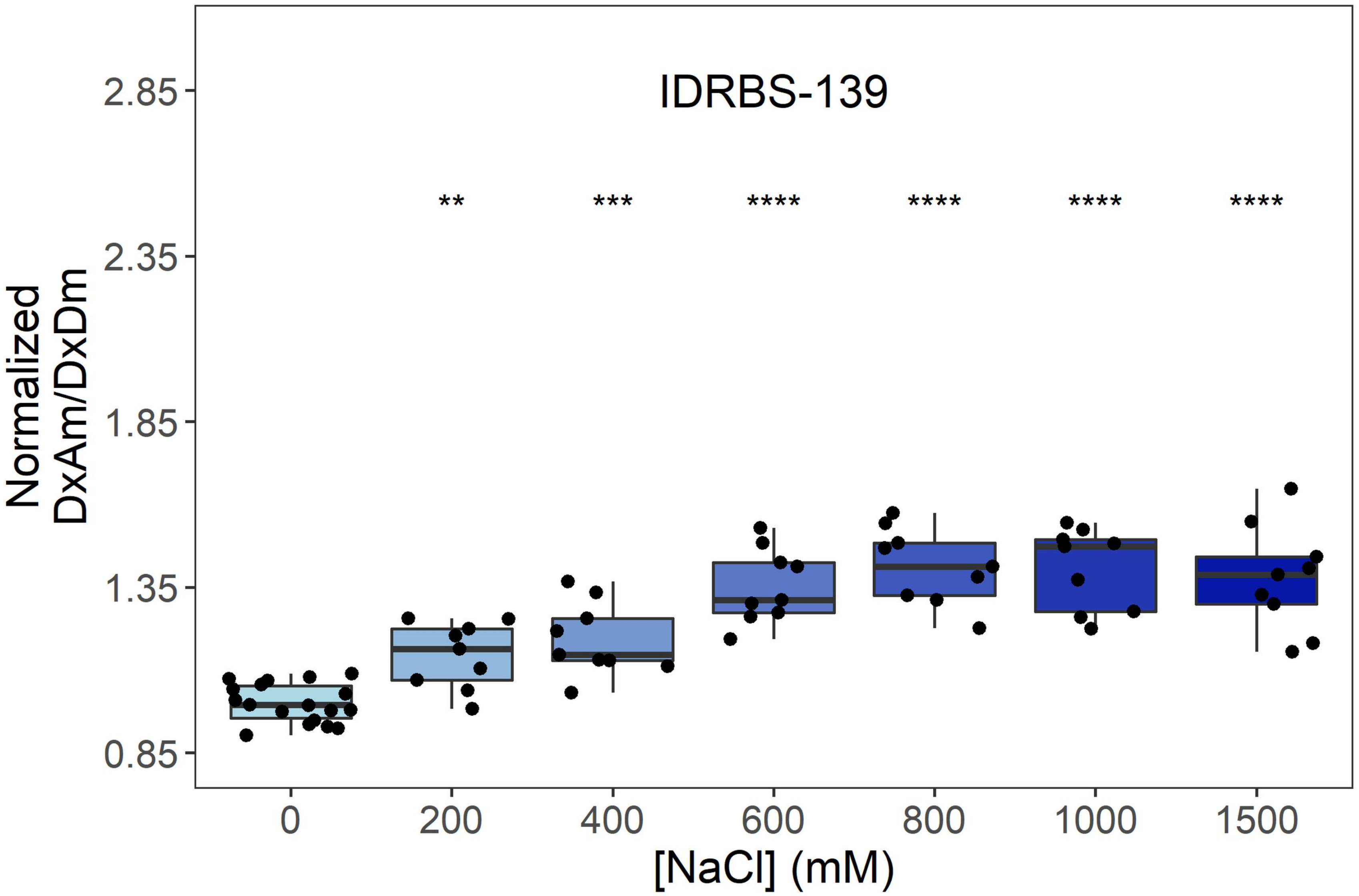
IDRBS-139
Normalized DxAm/DxDm
2.85
2.35
1.85
1.35
0.85
** *** **** **** **** ****
0
200
400
600
800
1000
1500
[NaCl] (mM)

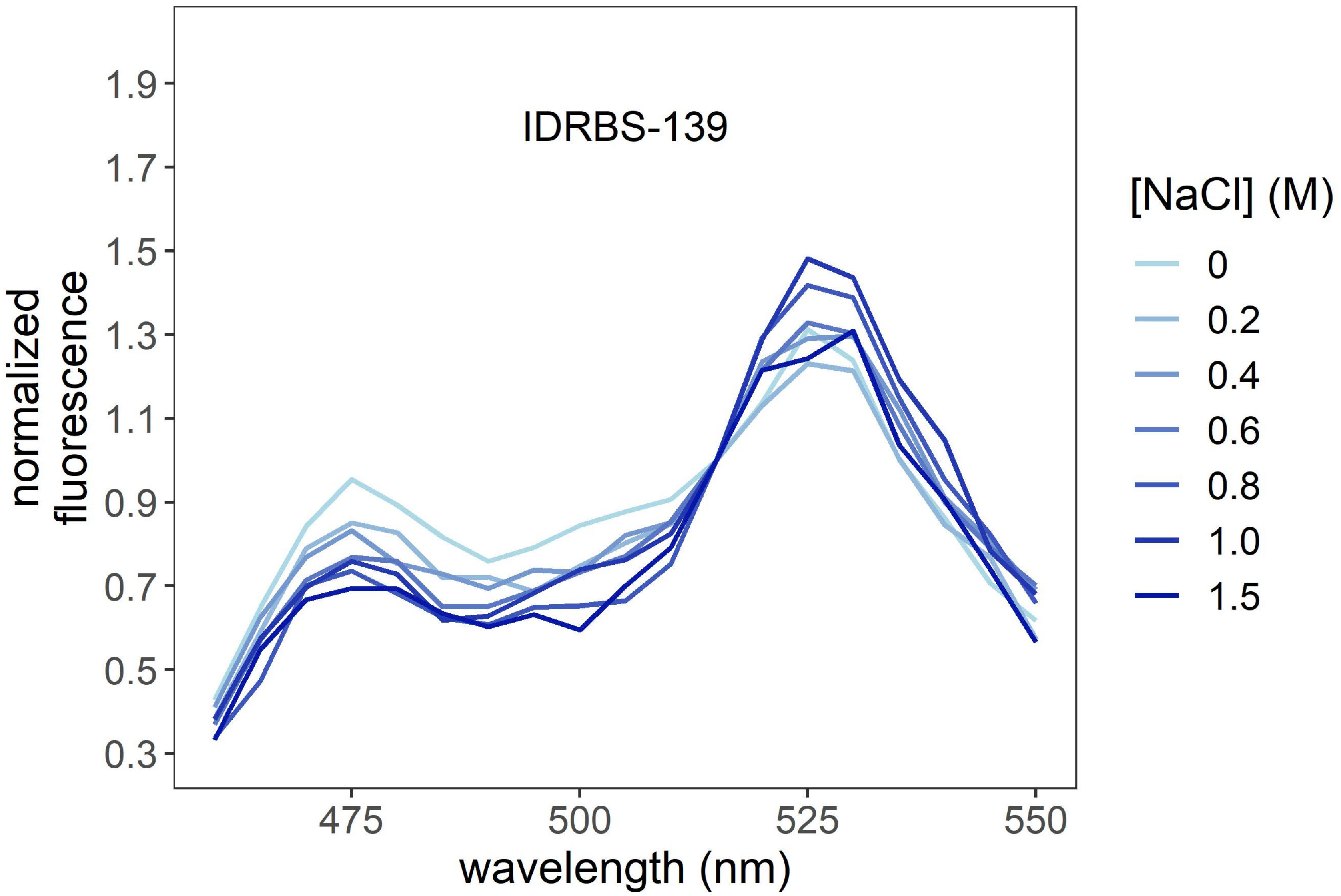
IDRBS-139
normalized fluorescence
1.9
1.7
1.5
1.3
1.1
0.9
0.7
0.5
0.3
475
500
525
550
wavelength (nm)
[NaCl] (M)
0
0.2
0.4
0.6
0.8
1.0
1.5

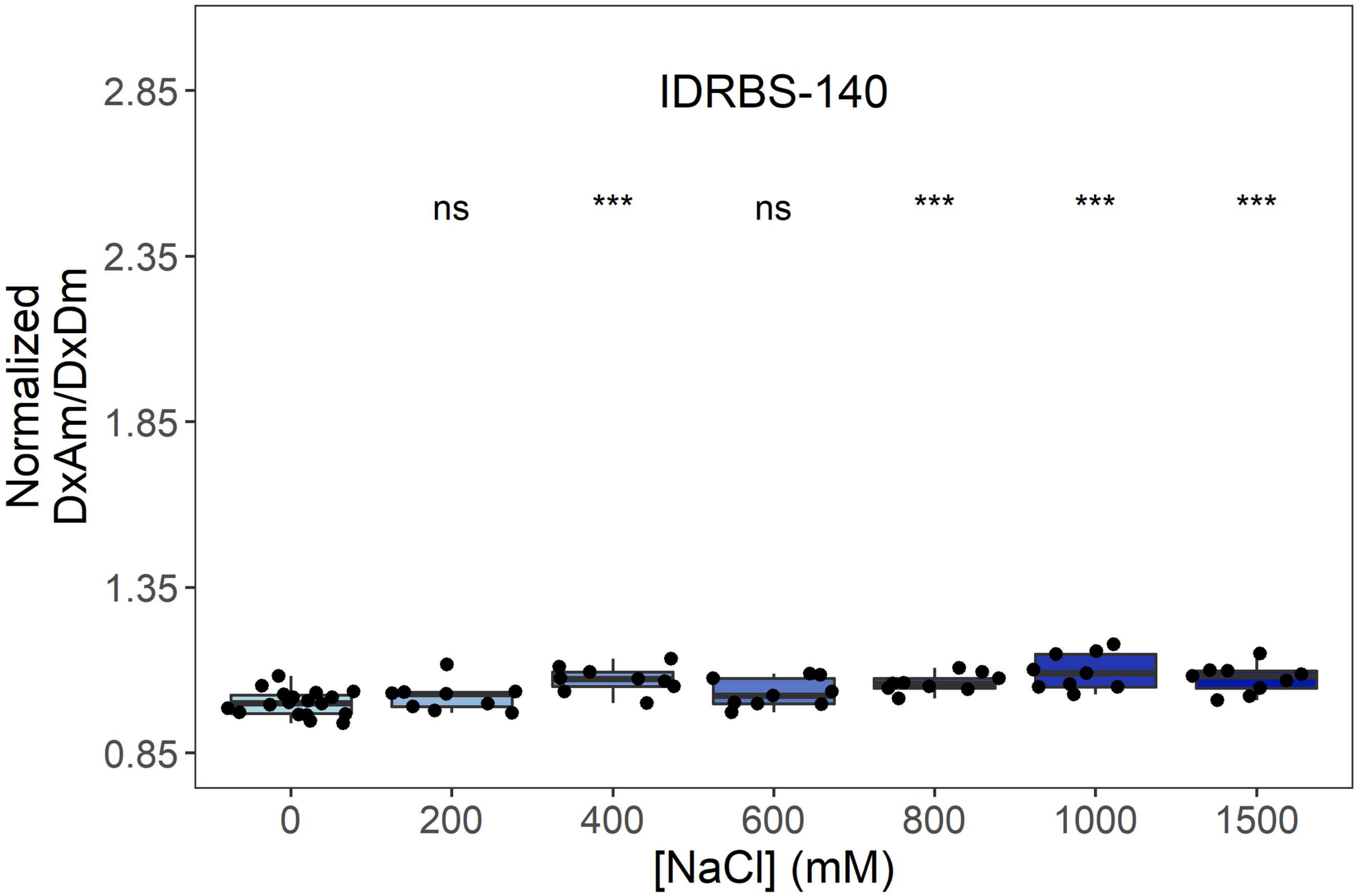
IDRBS-140
ns
***
ns
***
***
***
2.85
2.35
1.85
1.35
0.85
Normalized DxAm/DxDm
0
200
400
600
800
1000
1500
[NaCl] (mM)

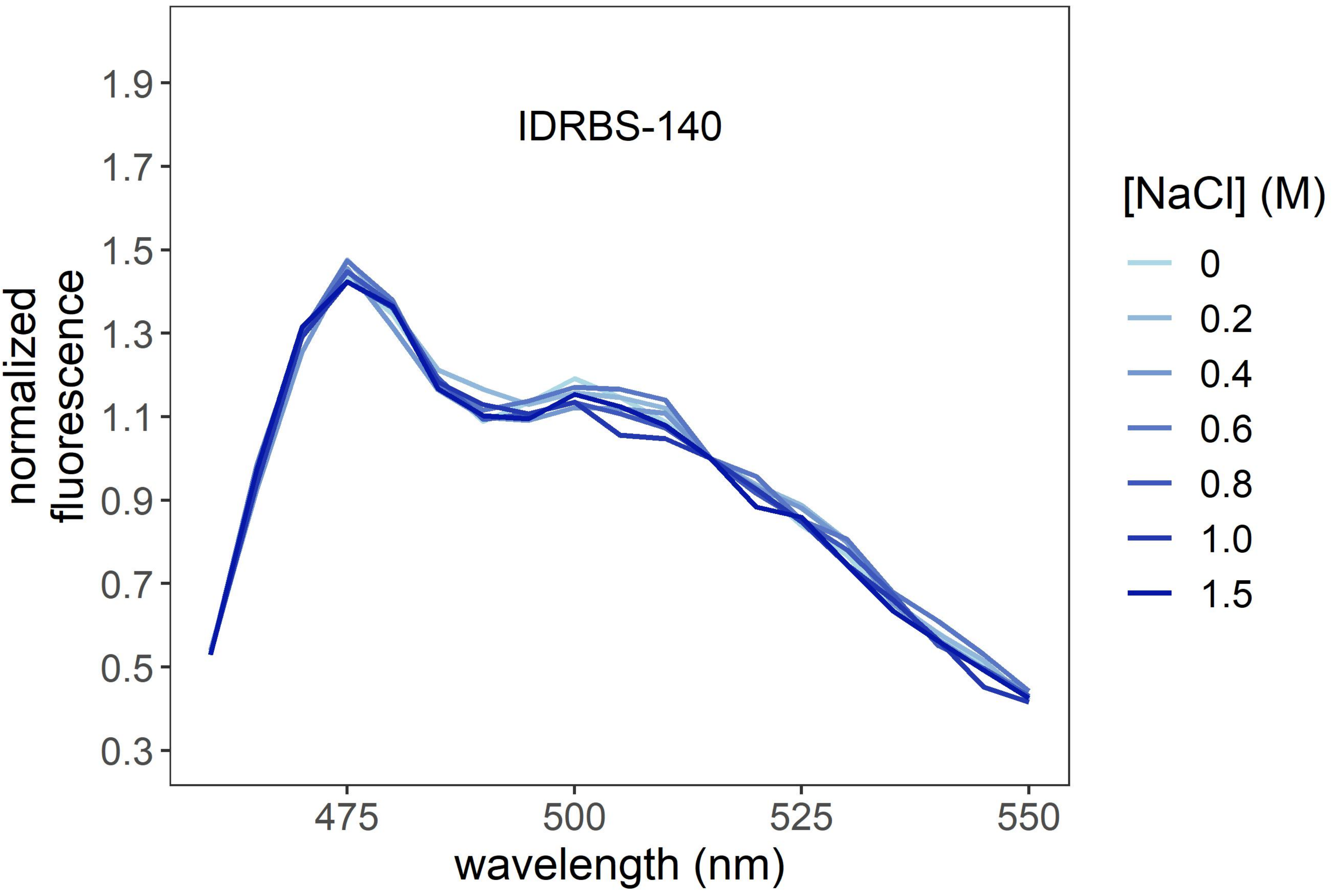
IDRBS-140
1.9
1.7
1.5
1.3
1.1
0.9
0.7
0.5
0.3
normalized fluorescence
475
500
525
550
wavelength (nm)
[NaCl] (M)
0
0.2
0.4
0.6
0.8
1.0
1.5

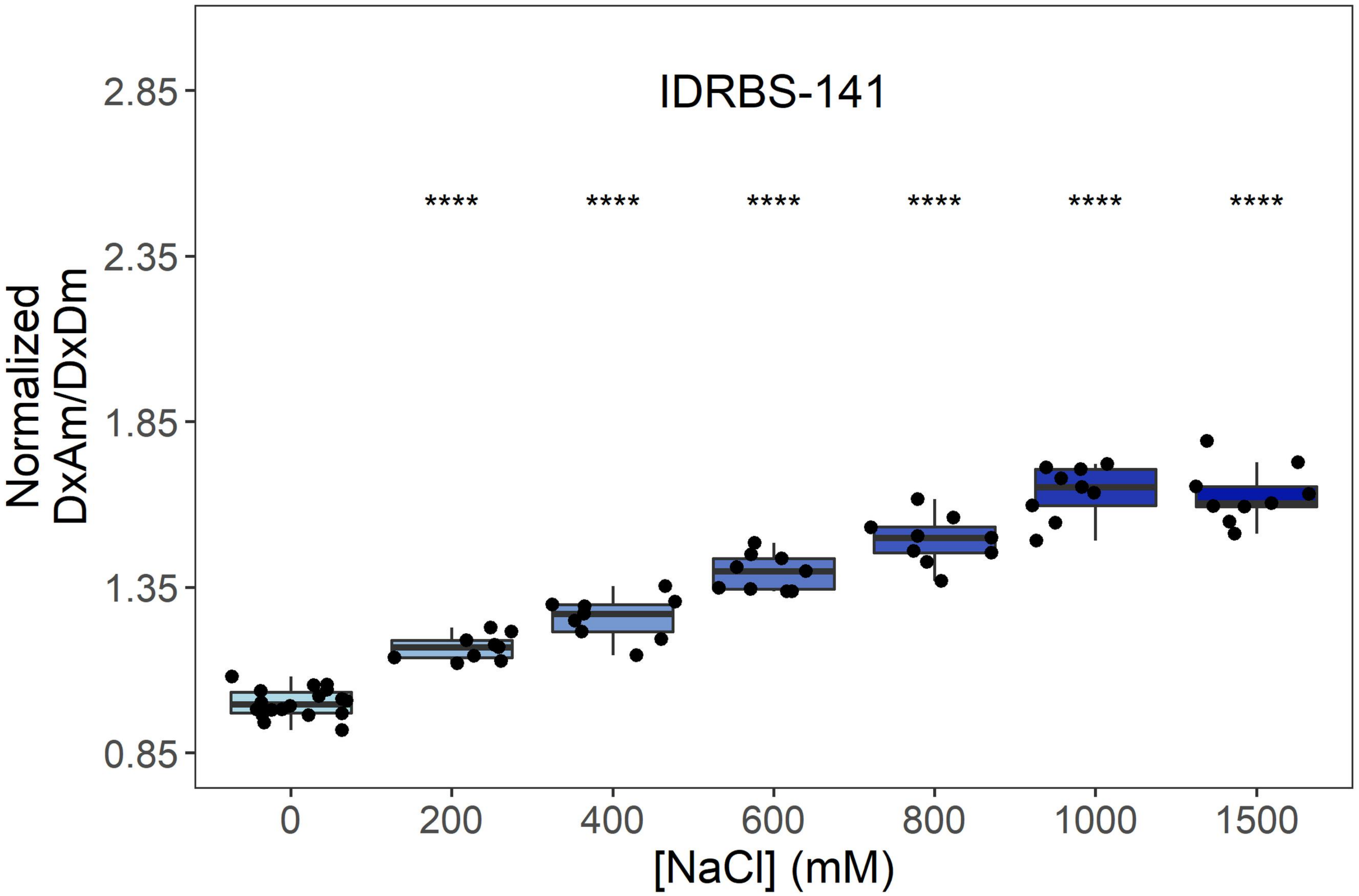
IDRBS-141
****
2.85
2.35
1.85
1.35
0.85
Normalized DxAm/DxDm
0
200
400
600
800
1000
1500
[NaCl] (mM)

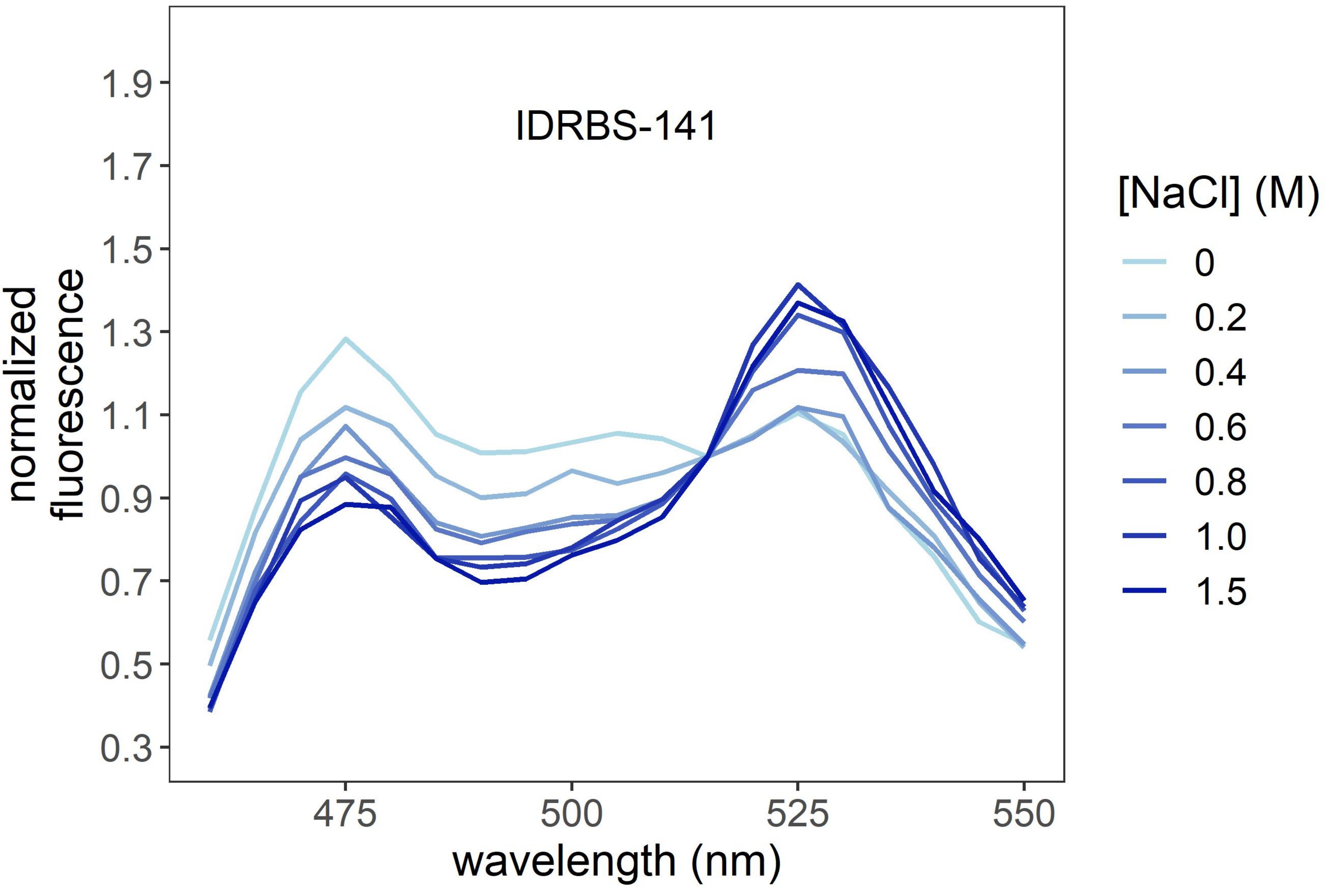
IDRBS-141
1.9
1.7
1.5
1.3
1.1
0.9
0.7
0.5
0.3
normalized fluorescence
475
500
525
550
wavelength (nm)
[NaCl] (M)
0
0.2
0.4
0.6
0.8
1.0
1.5

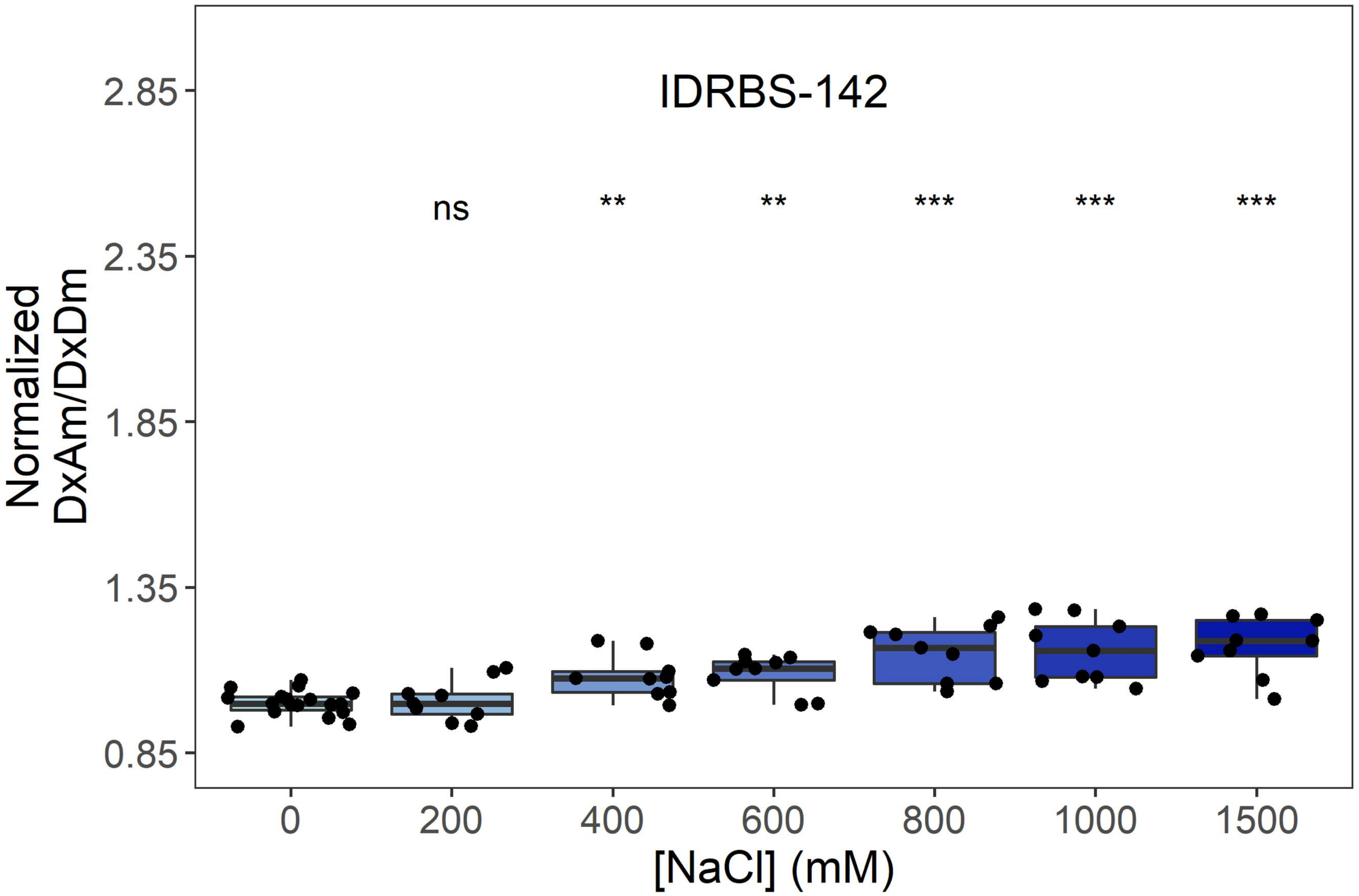
IDRBS-142
ns
**
**
***
***
***
2.85
2.35
1.85
1.35
0.85
Normalized
DxAm/DxDm
0
200
400
600
800
1000
1500
[NaCl] (mM)

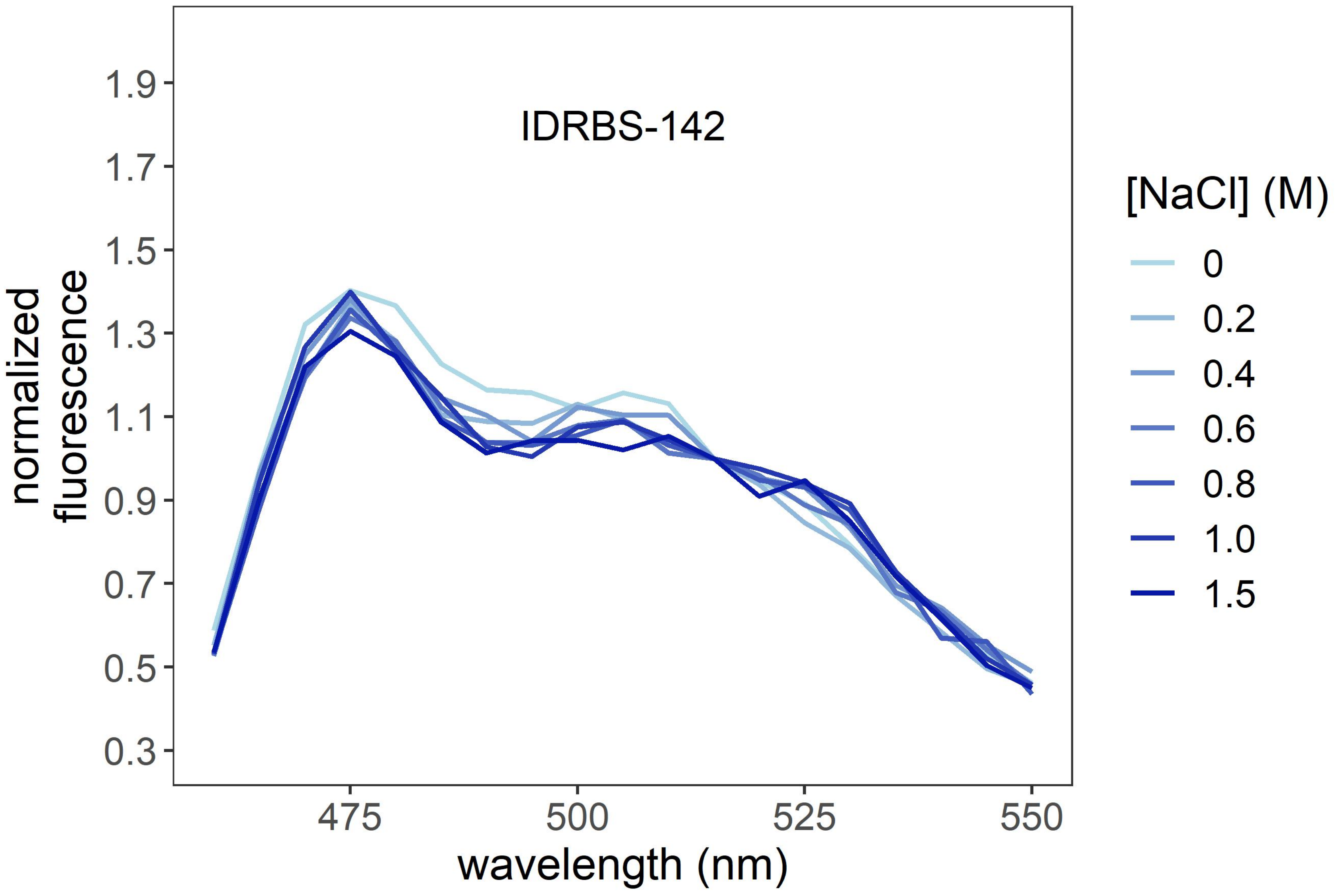
IDRBS-142
1.9
1.7
1.5
1.3
1.1
0.9
0.7
0.5
0.3
normalized
fluorescence
475
500
525
550
wavelength (nm)
[NaCl] (M)
0
0.2
0.4
0.6
0.8
1.0
1.5

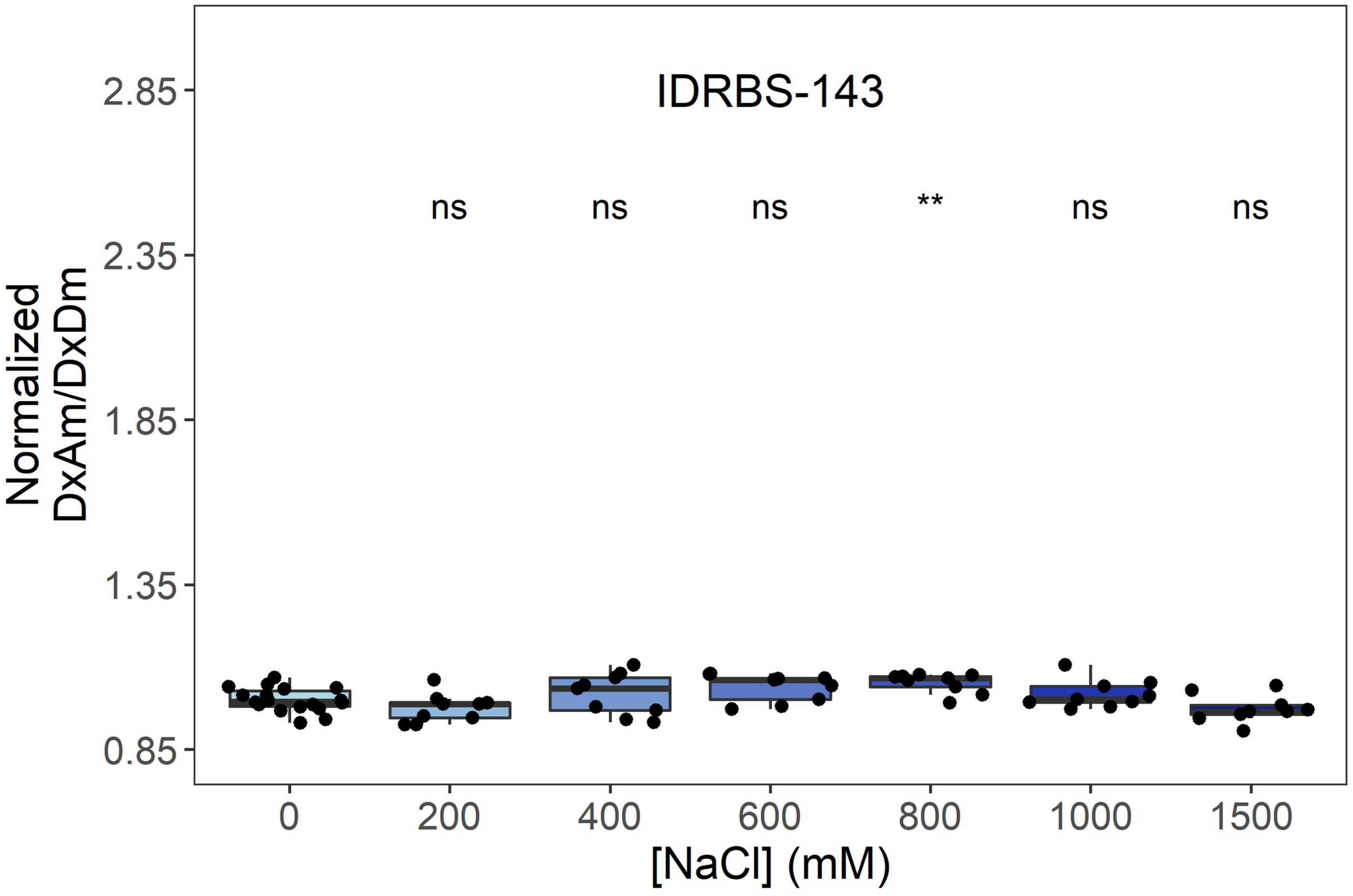
IDRBS-143
ns
ns
ns
**
ns
ns
Normalized DxAm/DxDm
2.85
2.35
1.85
1.35
0.85
0
200
400
600
800
1000
1500
[NaCl] (mM)

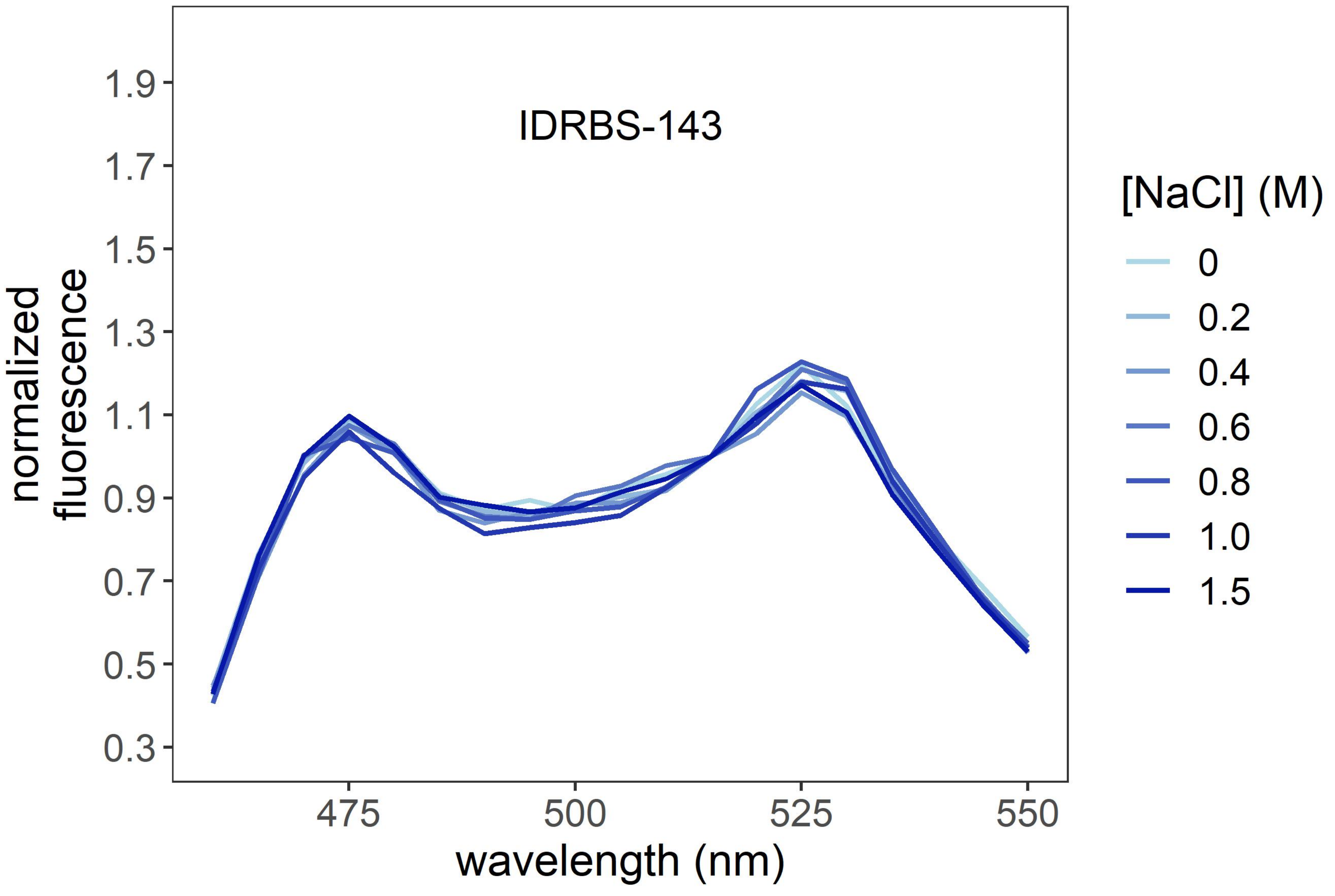
IDRBS-143
normalized fluorescence
1.9
1.7
1.5
1.3
1.1
0.9
0.7
0.5
0.3
475
500
525
550
wavelength (nm)
[NaCl] (M)
0
0.2
0.4
0.6
0.8
1.0
1.5

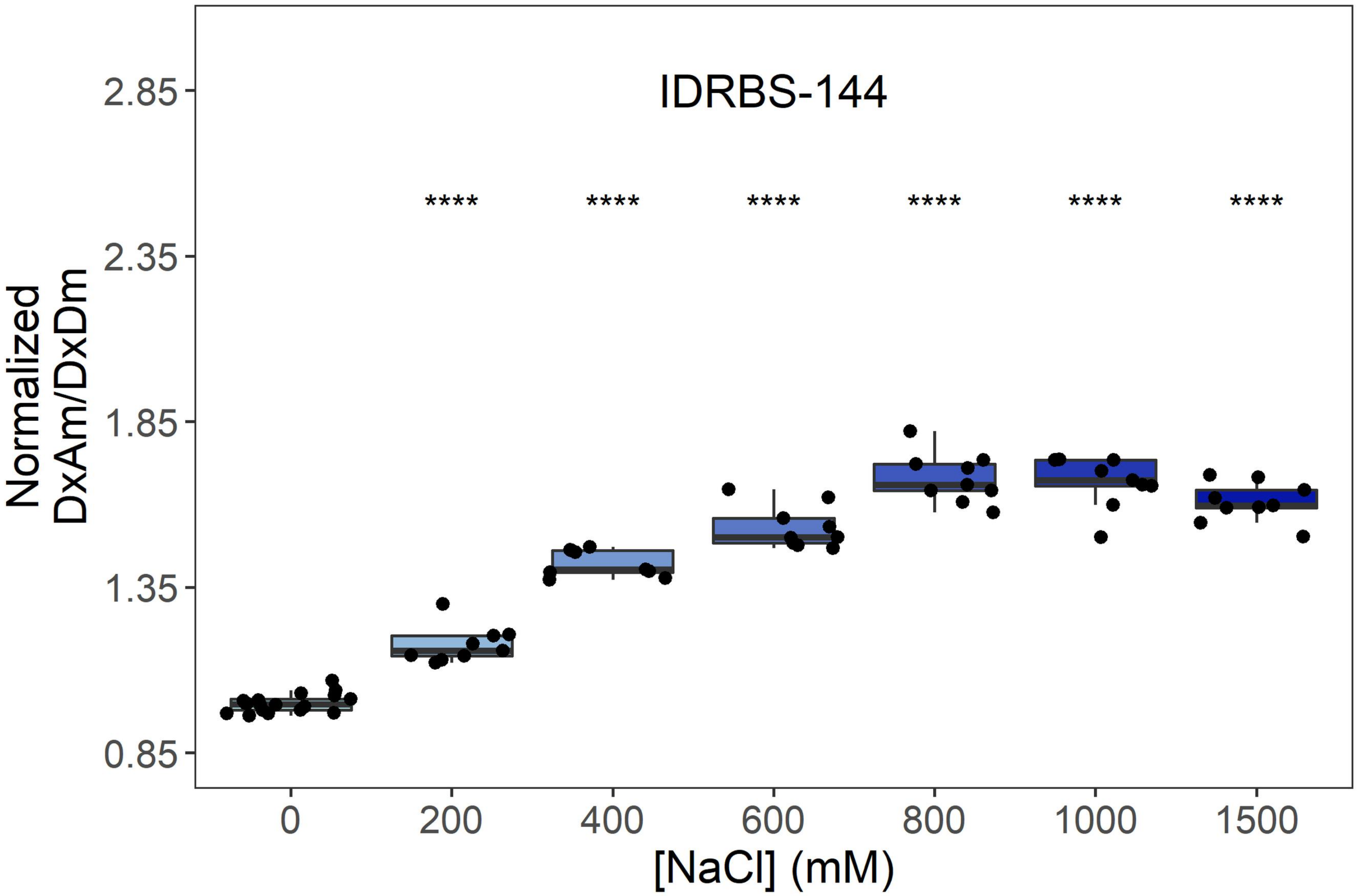
IDRBS-144
Normalized DxAm/DxDm
2.85
2.35
1.85
1.35
0.85
**** **** **** **** **** ****
0
200
400
600
800
1000
1500
[NaCl] (mM)

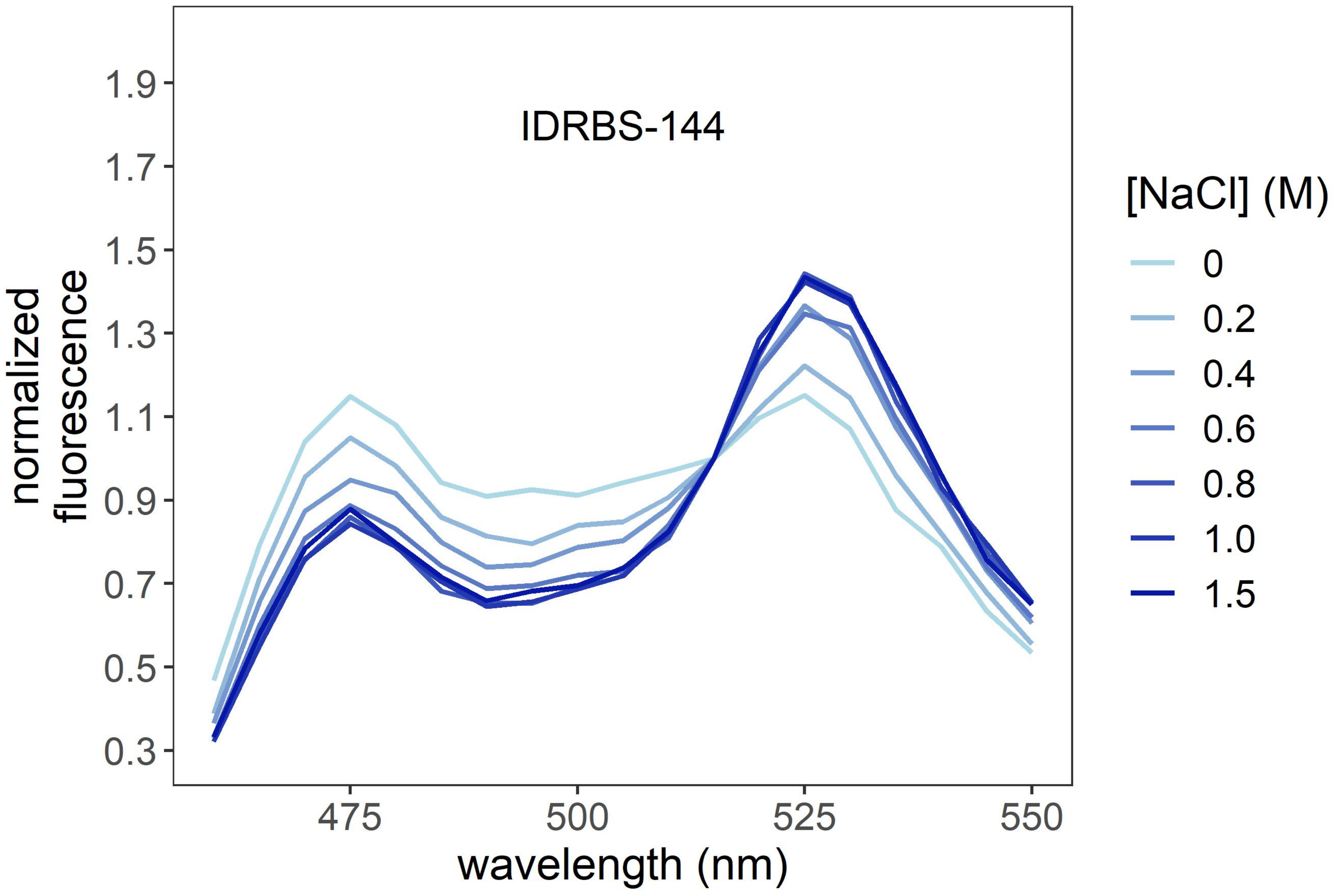
IDRBS-144
normalized fluorescence
1.9
1.7
1.5
1.3
1.1
0.9
0.7
0.5
0.3
475
500
525
550
wavelength (nm)
[NaCl] (M)
0
0.2
0.4
0.6
0.8
1.0
1.5

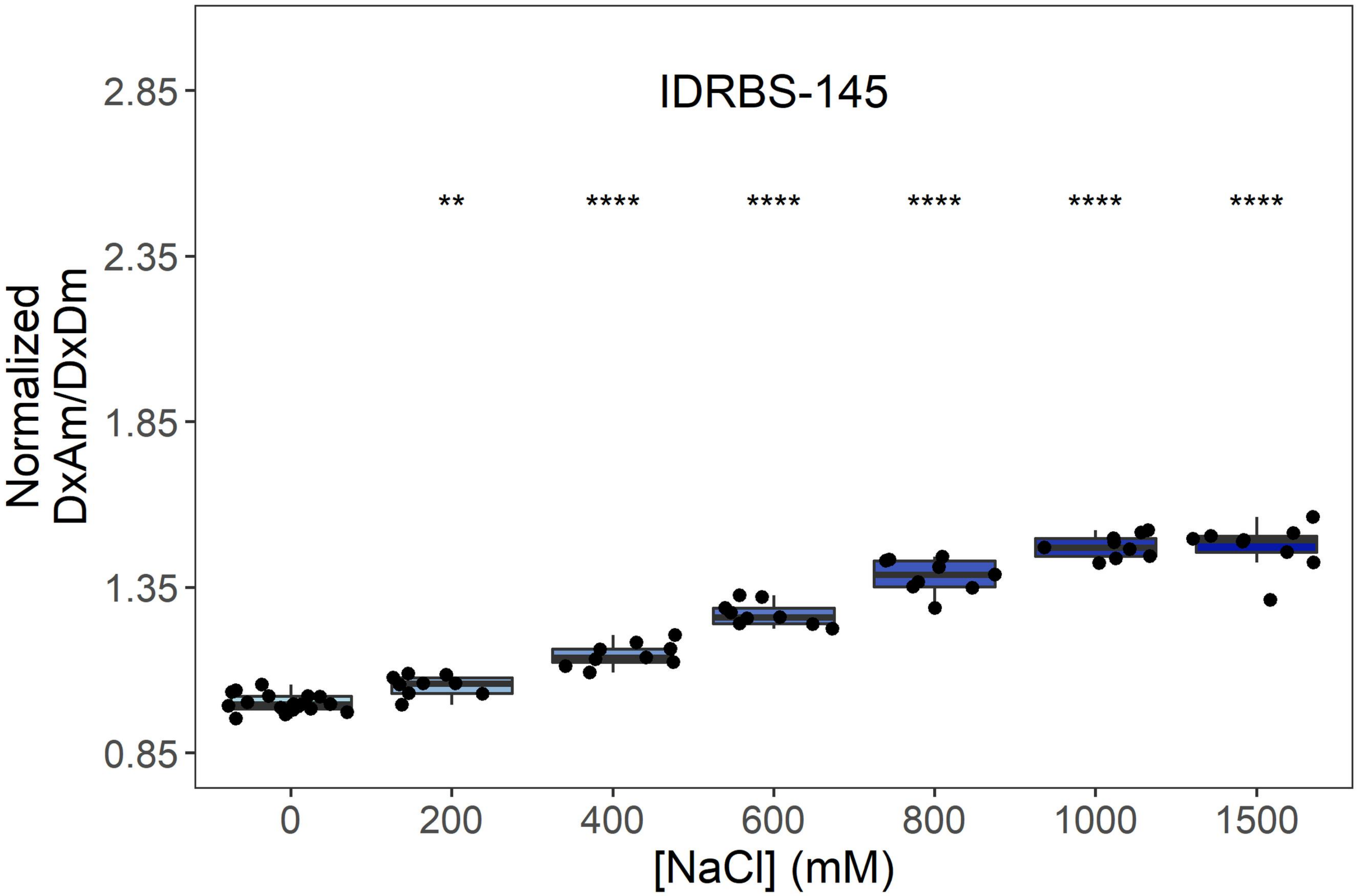
IDRBS-145
Normalized DxAm/DxDm
2.85
2.35
1.85
1.35
0.85
**
****
****
****
****
****
0
200
400
600
800
1000
1500
[NaCl] (mM)

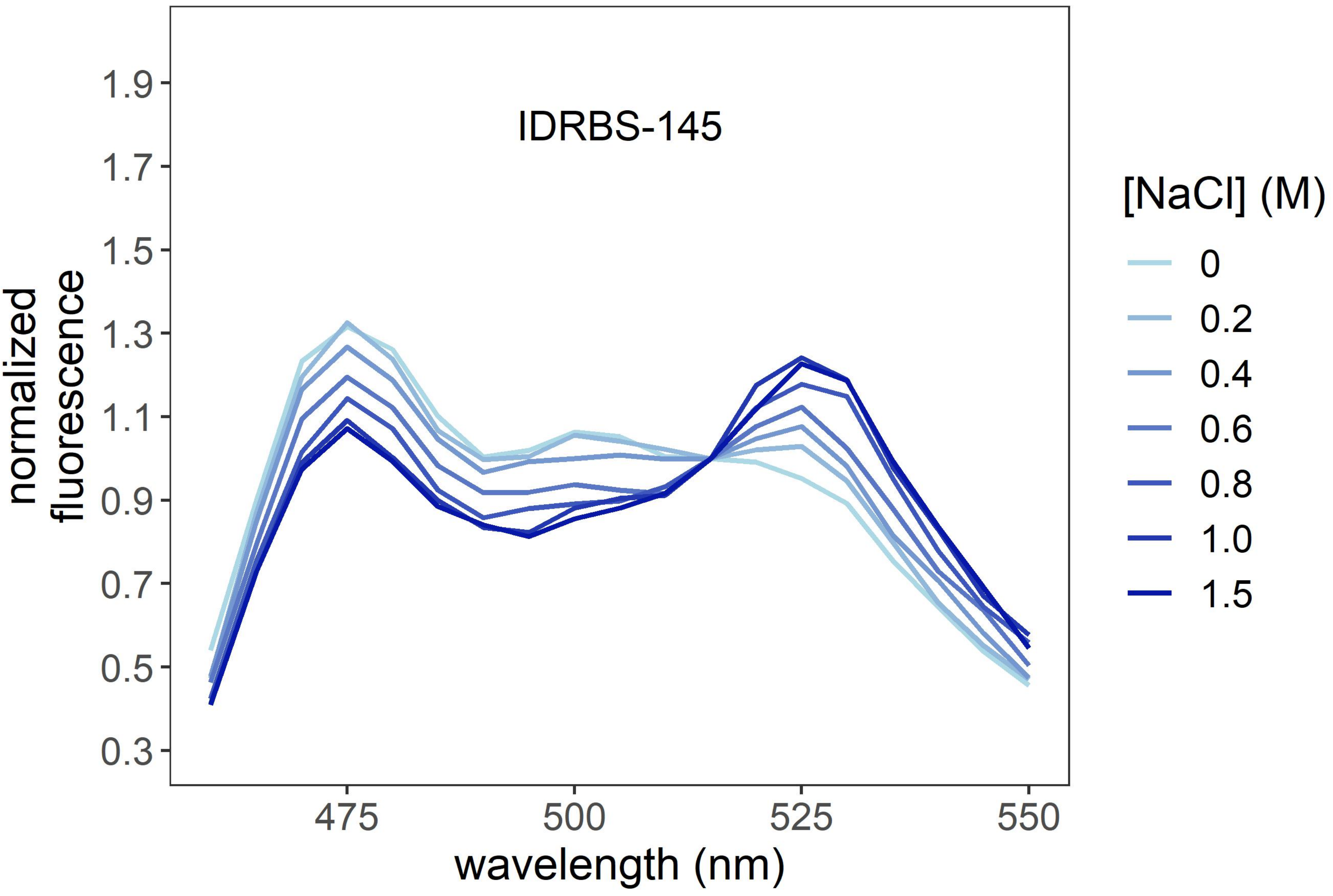
IDRBS-145
normalized fluorescence
1.9
1.7
1.5
1.3
1.1
0.9
0.7
0.5
0.3
475
500
525
550
wavelength (nm)
[NaCl] (M)
0
0.2
0.4
0.6
0.8
1.0
1.5

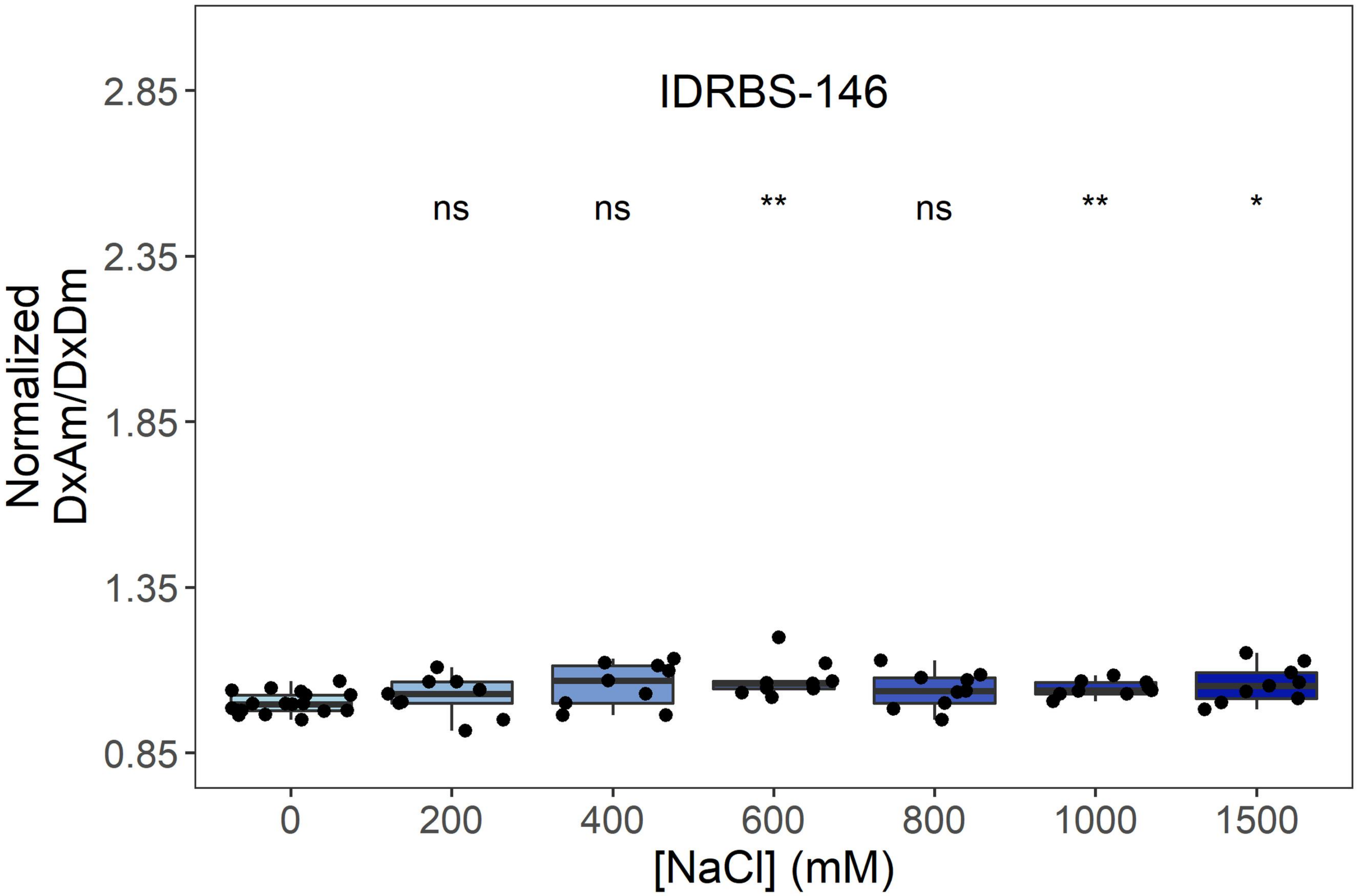
IDRBS-146
ns
ns
**
ns
**
*
Normalized DxAm/DxDm
2.85
2.35
1.85
1.35
0.85
0
200
400
600
800
1000
1500
[NaCl] (mM)

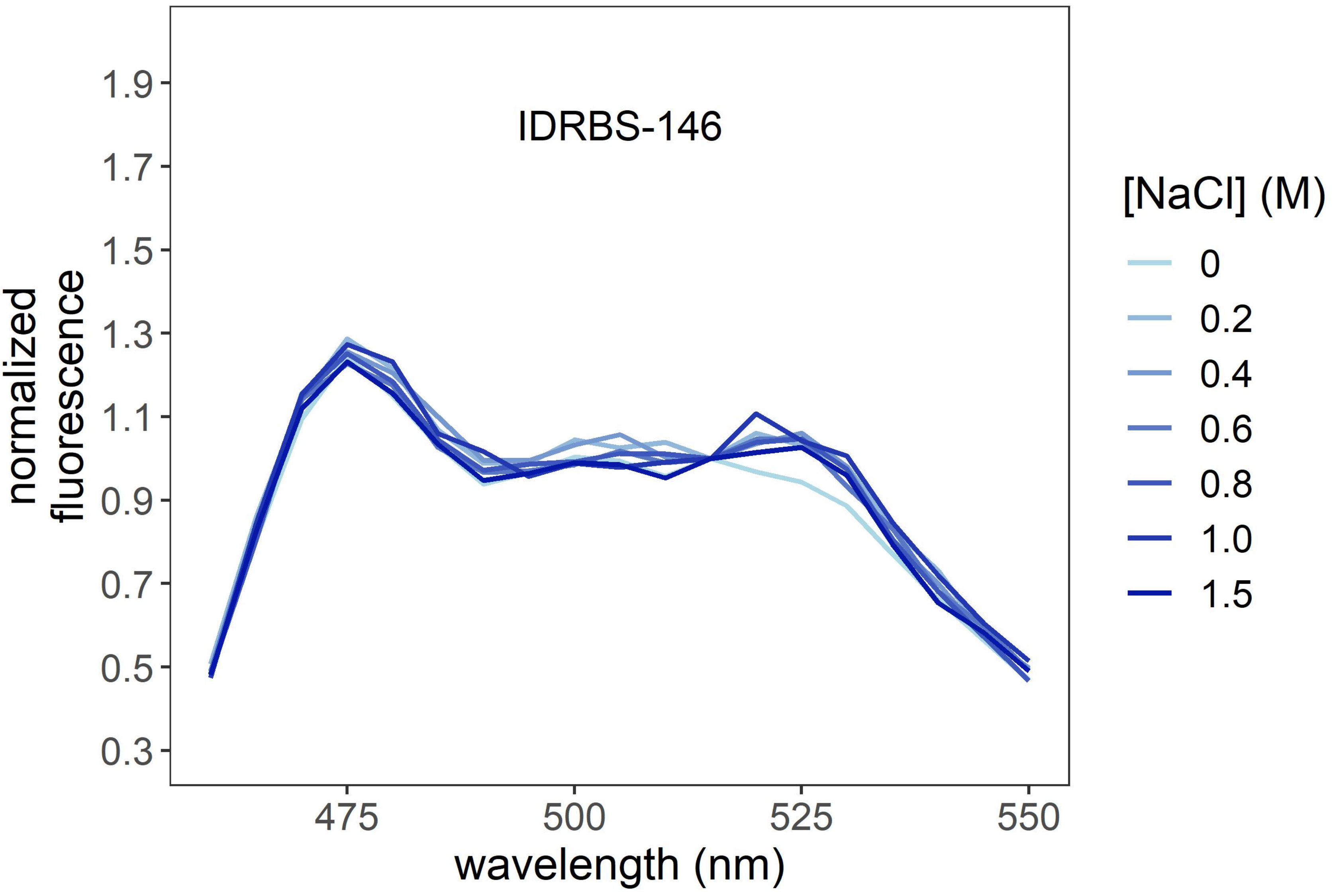
IDRBS-146
normalized fluorescence
1.9
1.7
1.5
1.3
1.1
0.9
0.7
0.5
0.3
475
500
525
550
wavelength (nm)
[NaCl] (M)
0
0.2
0.4
0.6
0.8
1.0
1.5

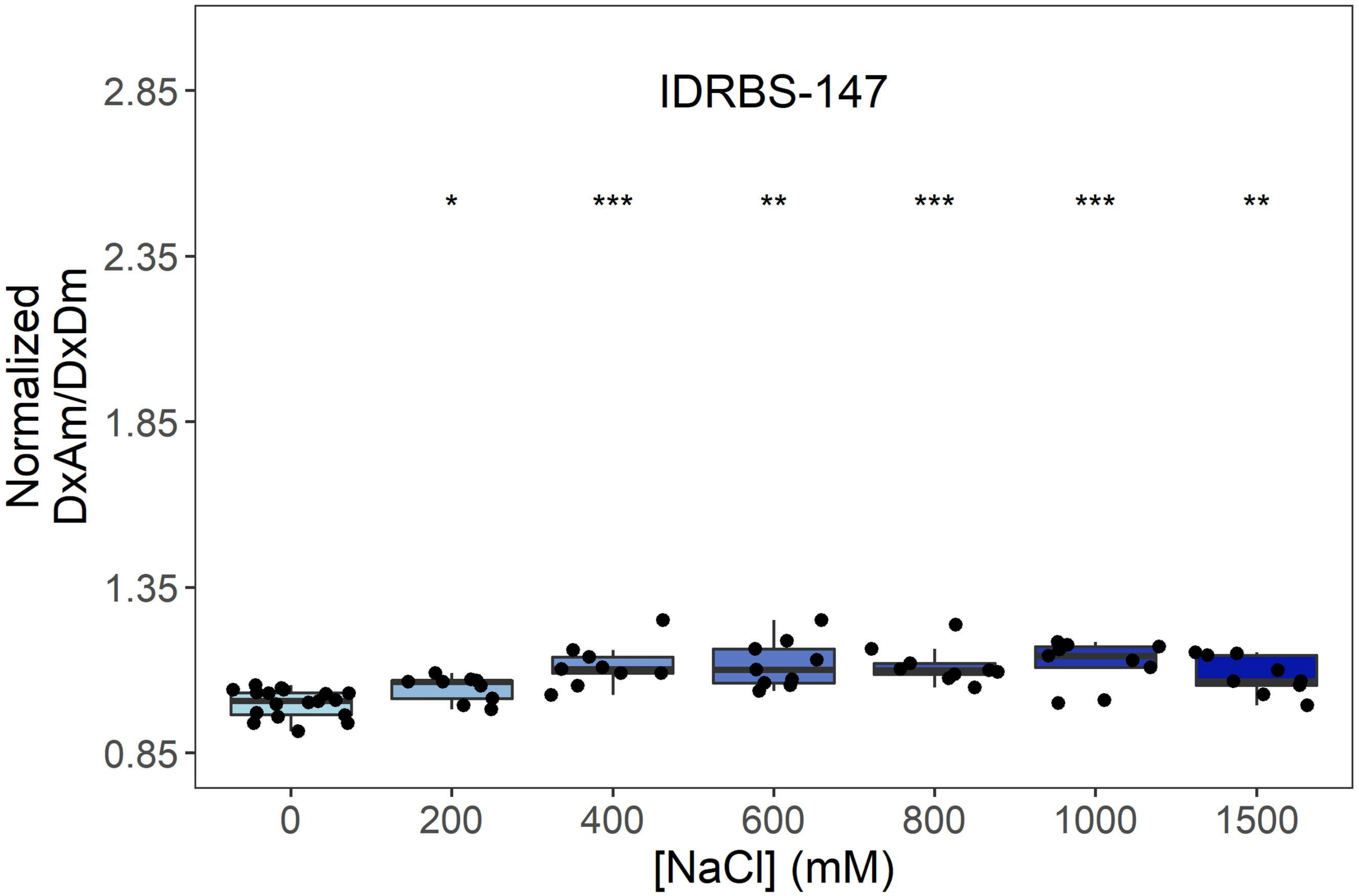
IDRBS-147
*
***
**
***
***
**
Normalized
DxAm/DxDm
2.85
2.35
1.85
1.35
0.85
0
200
400
600
800
1000
1500
[NaCl] (mM)

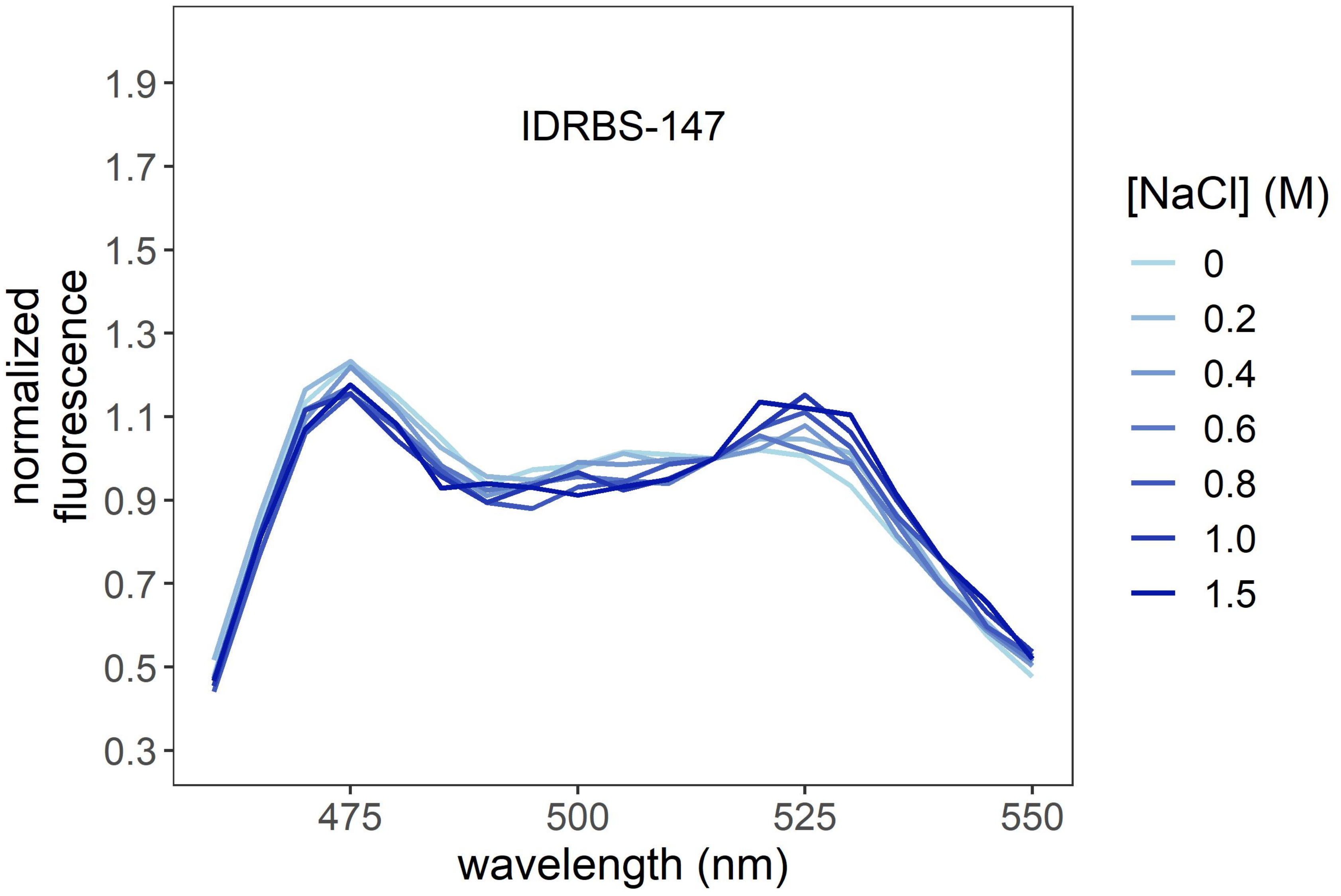
IDRBS-147
normalized
fluorescence
1.9
1.7
1.5
1.3
1.1
0.9
0.7
0.5
0.3
475
500
525
550
wavelength (nm)
[NaCl] (M)
0
0.2
0.4
0.6
0.8
1.0
1.5

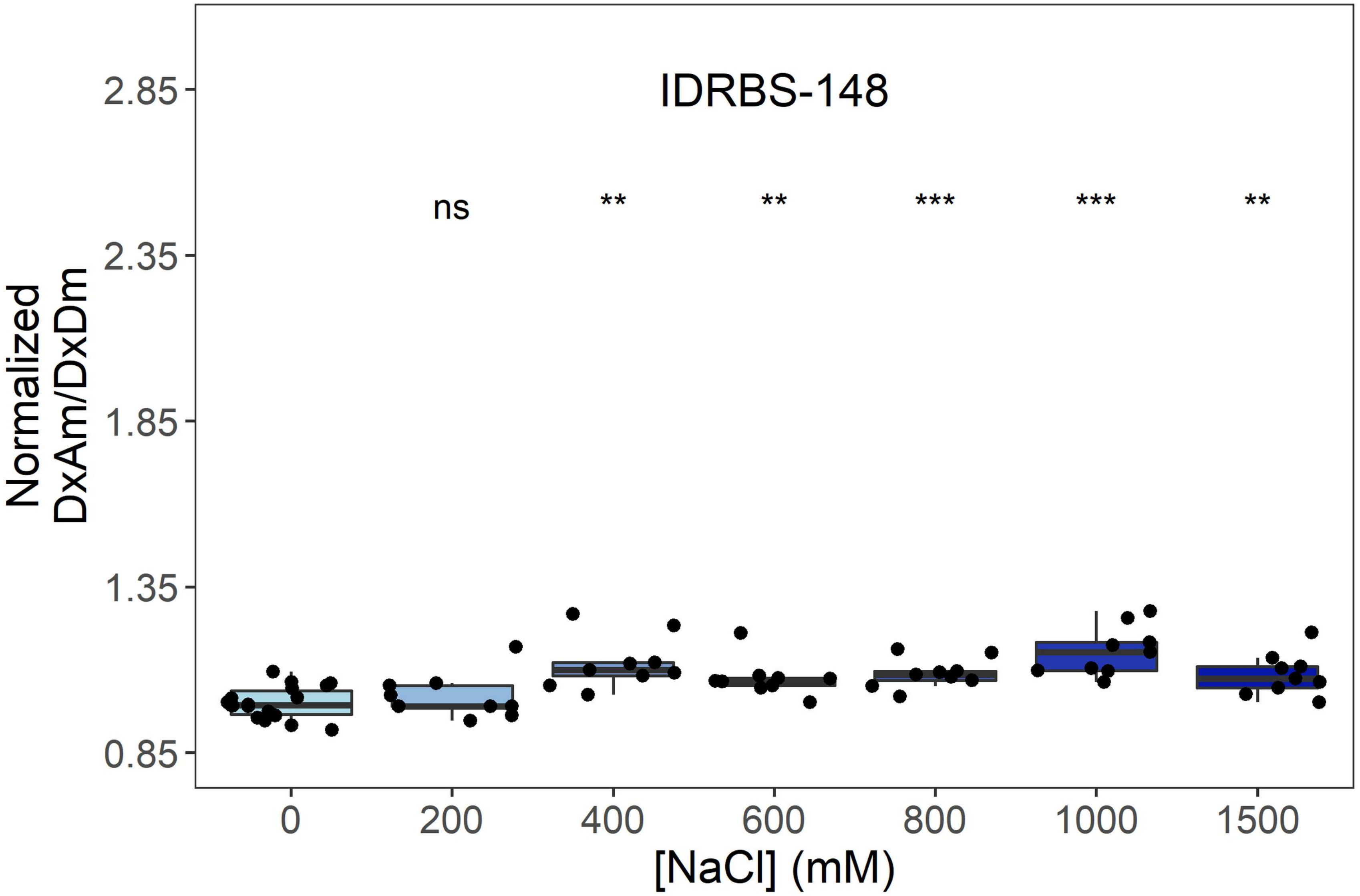
IDRBS-148
ns
**
**
***
***
**
Normalized DxAm/DxDm
2.85
2.35
1.85
1.35
0.85
0
200
400
600
800
1000
1500
[NaCl] (mM)

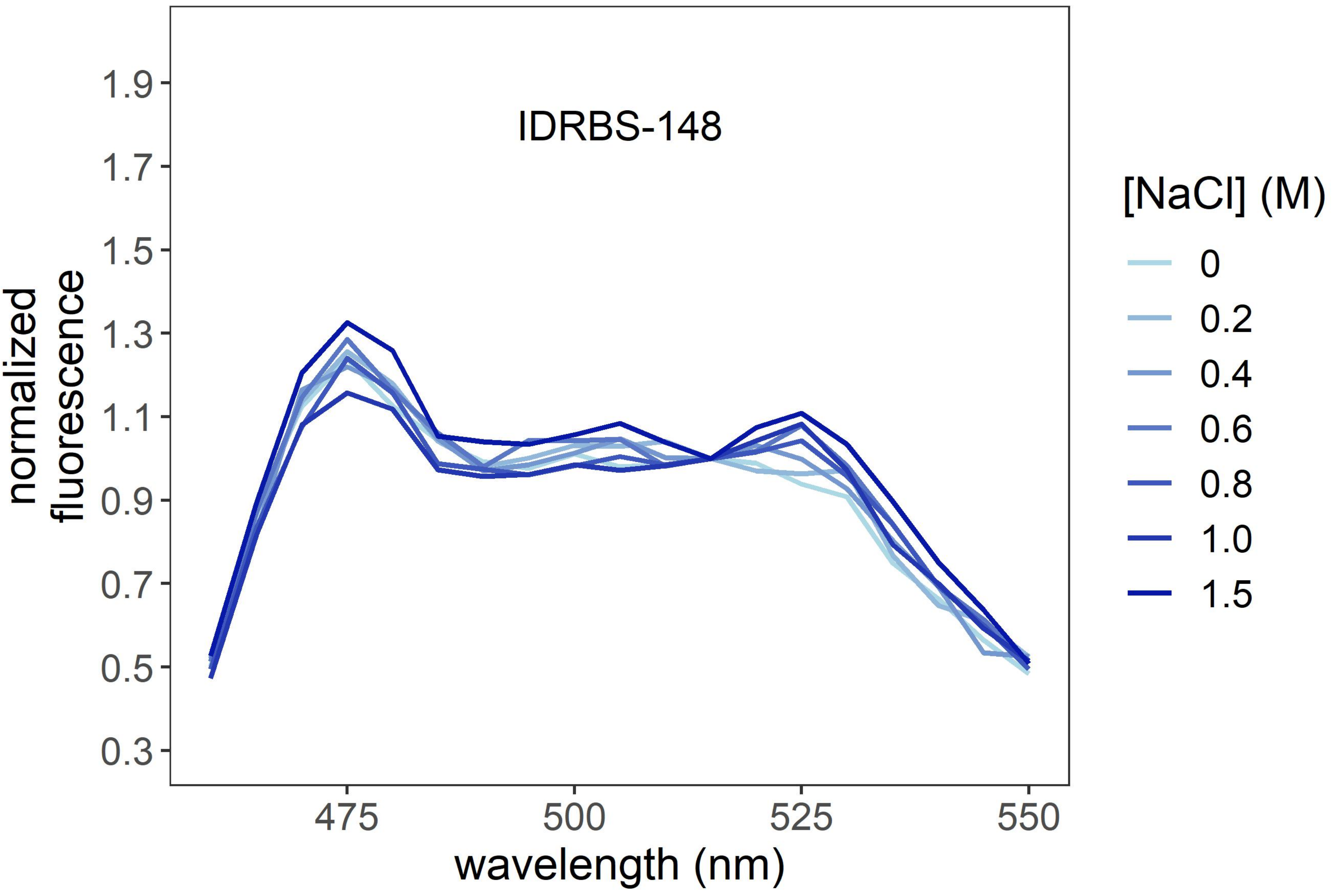
IDRBS-148
normalized fluorescence
1.9
1.7
1.5
1.3
1.1
0.9
0.7
0.5
0.3
475
500
525
550
wavelength (nm)
[NaCl] (M)
0
0.2
0.4
0.6
0.8
1.0
1.5

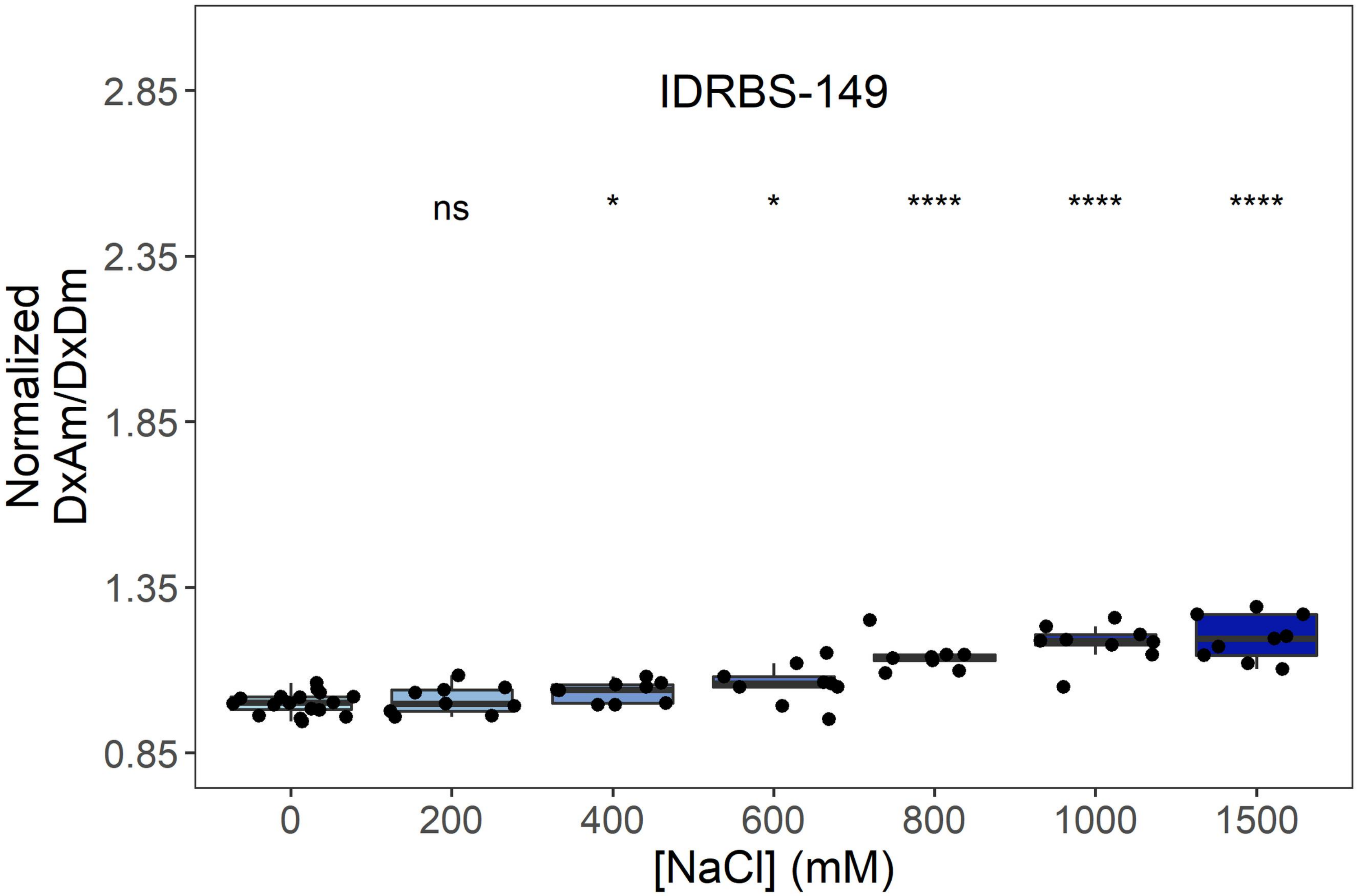
IDRBS-149
ns
*
*
****
****
****
Normalized DxAm/DxDm
2.85
2.35
1.85
1.35
0.85
0
200
400
600
800
1000
1500
[NaCl] (mM)

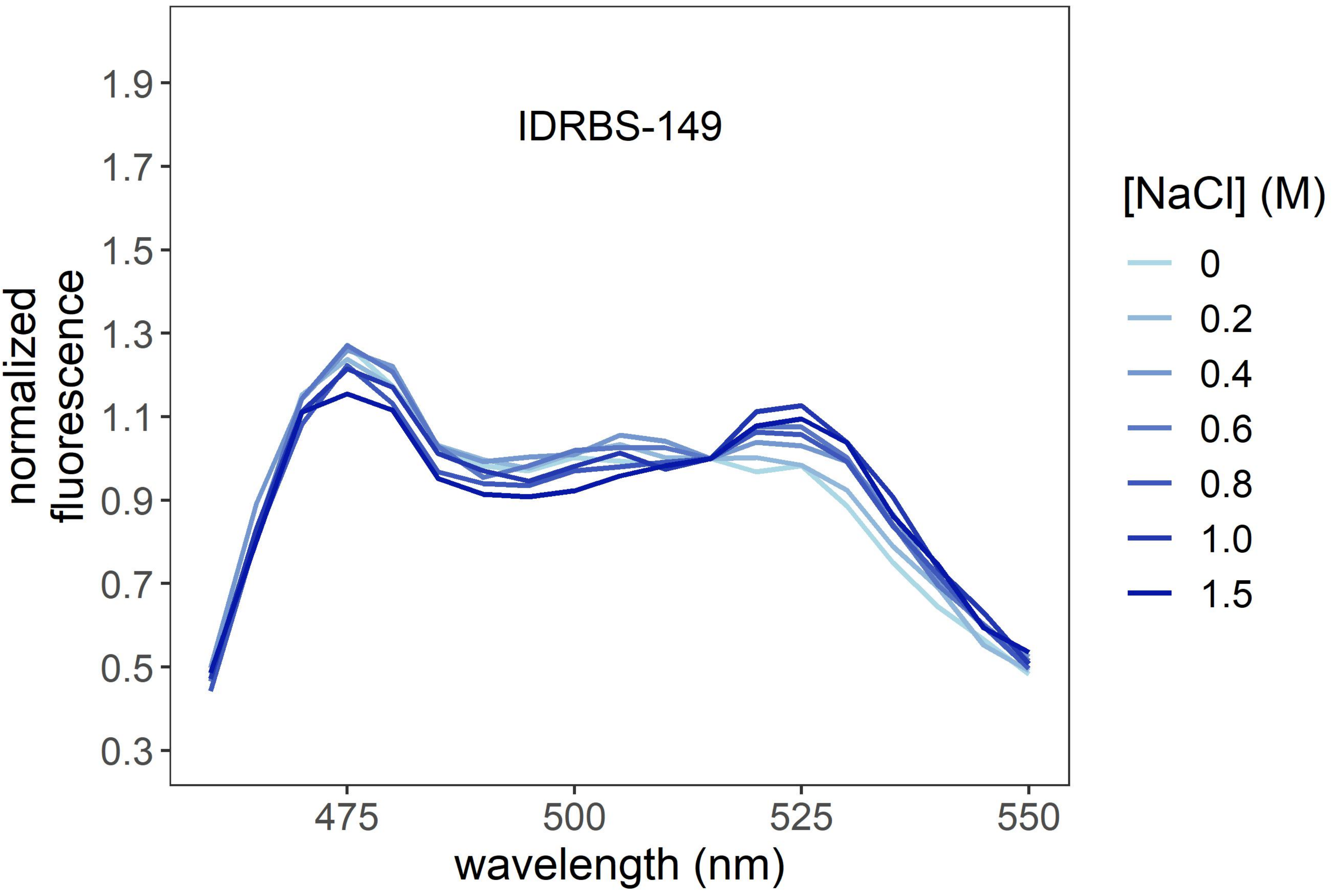
IDRBS-149
normalized fluorescence
1.9
1.7
1.5
1.3
1.1
0.9
0.7
0.5
0.3
475
500
525
550
wavelength (nm)
[NaCl] (M)
0
0.2
0.4
0.6
0.8
1.0
1.5

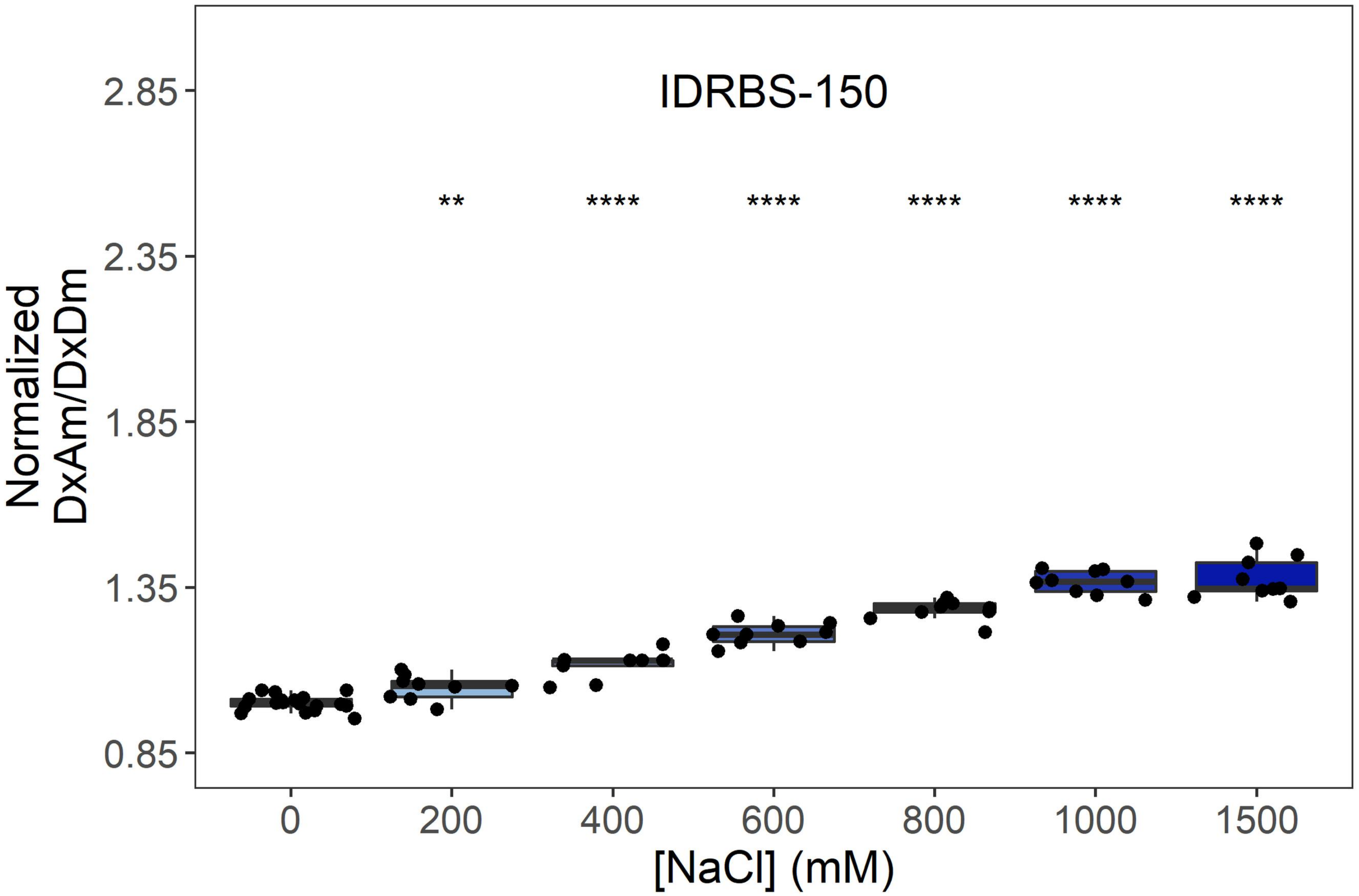
IDRBS-150
Normalized
DxAm/DxDm
2.85
2.35
1.85
1.35
0.85
**
****
****
****
****
****
0
200
400
600
800
1000
1500
[NaCl] (mM)

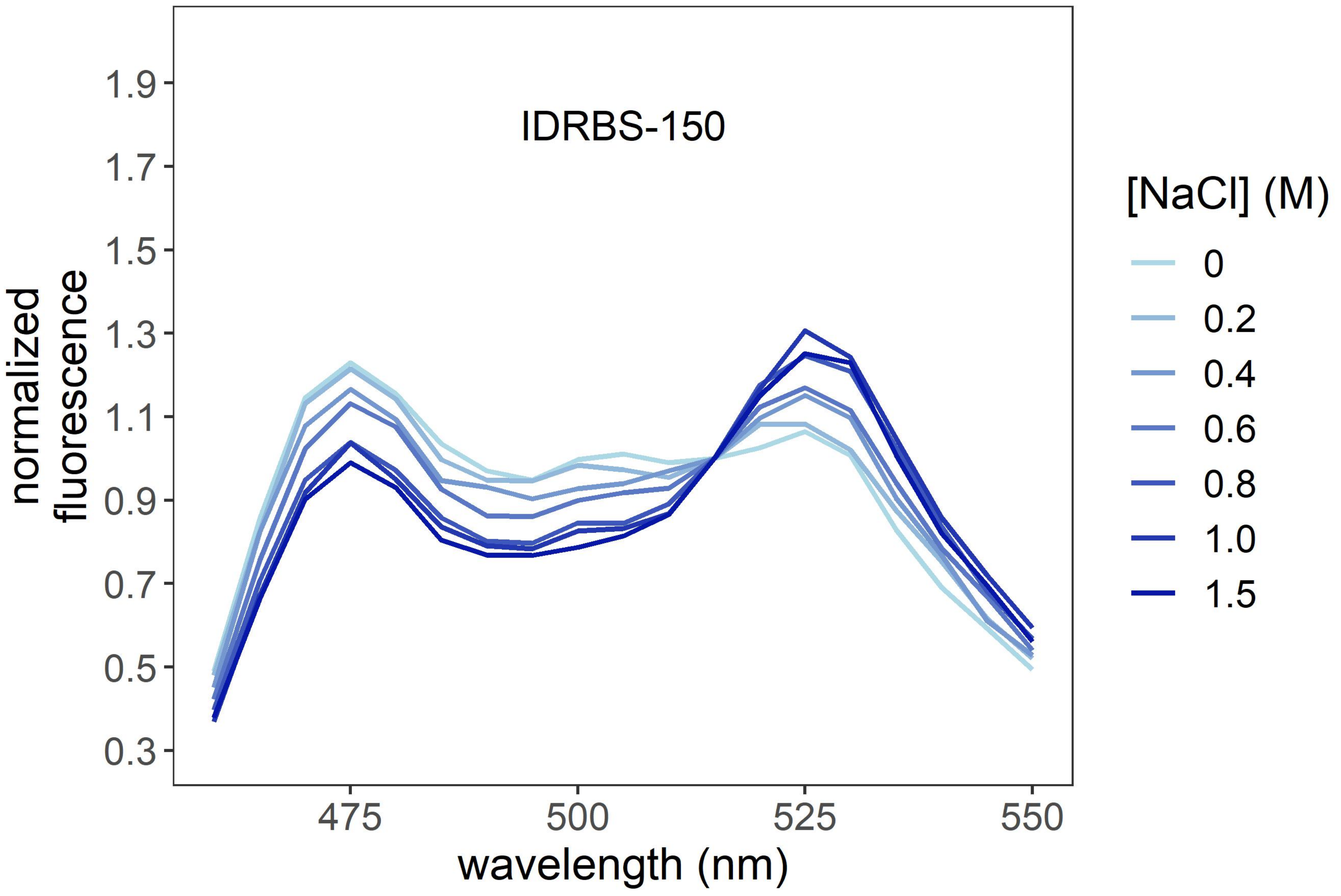
IDRBS-150
normalized
fluorescence
1.9
1.7
1.5
1.3
1.1
0.9
0.7
0.5
0.3
475
500
525
550
wavelength (nm)
[NaCl] (M)
0
0.2
0.4
0.6
0.8
1.0
1.5

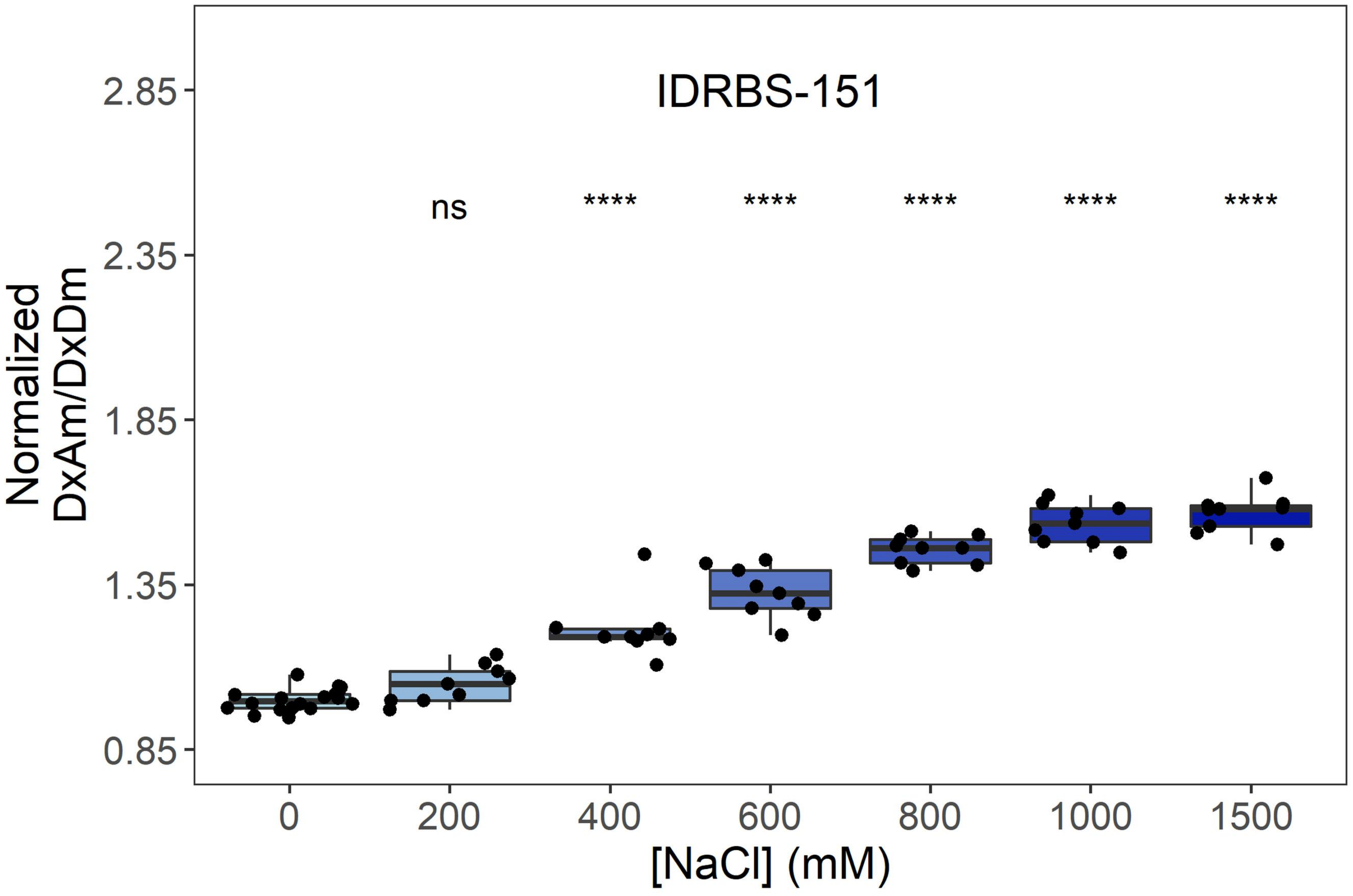
IDRBS-151
ns
****
****
****
****
****
2.85
2.35
1.85
1.35
0.85
Normalized
DxAm/DxDm
0
200
400
600
800
1000
1500
[NaCl] (mM)

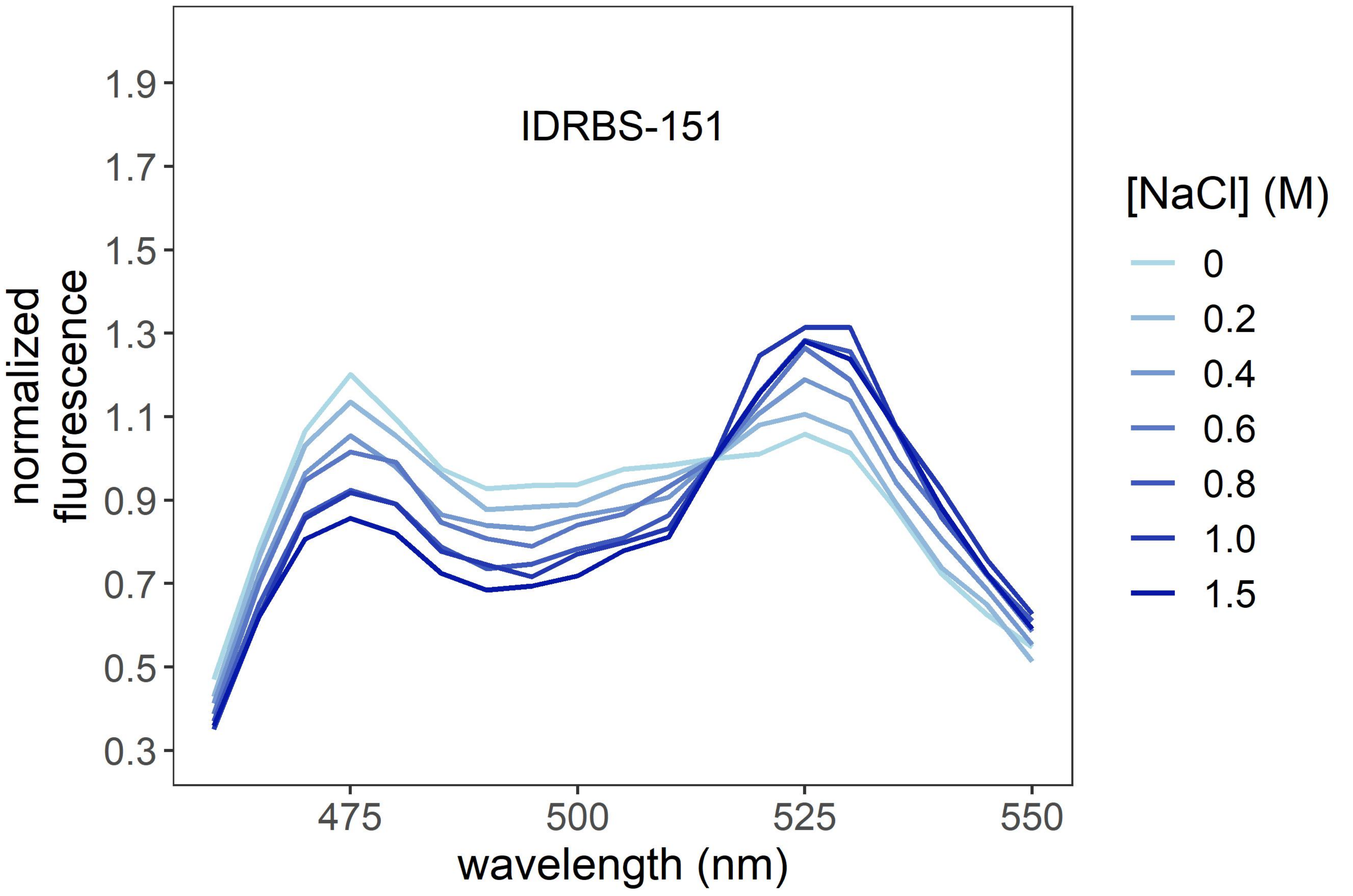
IDRBS-151
1.9
1.7
1.5
1.3
1.1
0.9
0.7
0.5
0.3
normalized
fluorescence
475
500
525
550
wavelength (nm)
[NaCl] (M)
0
0.2
0.4
0.6
0.8
1.0
1.5

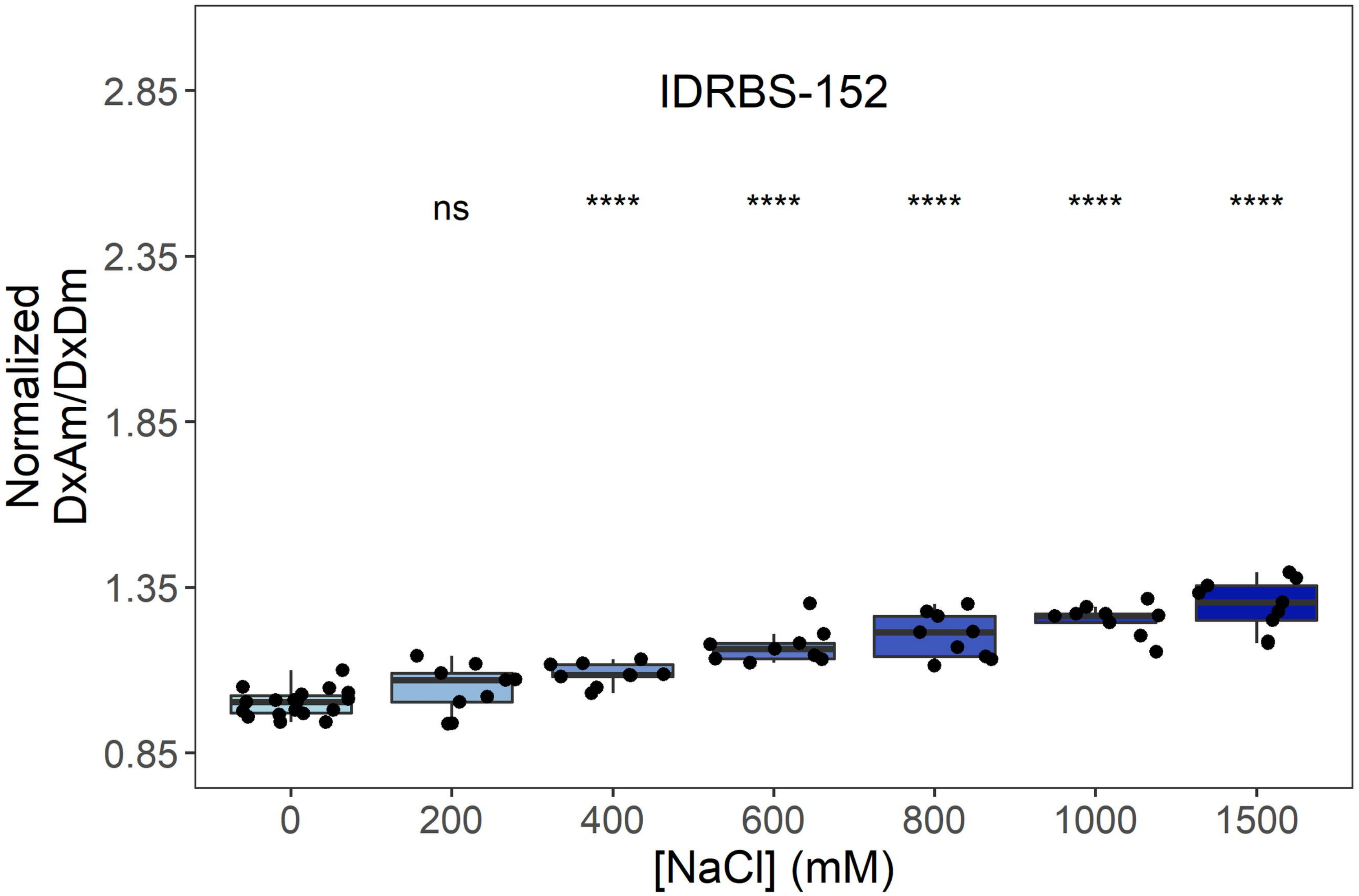
IDRBS-152
ns
****
****
****
****
****
Normalized
DxAm/DxDm
2.85
2.35
1.85
1.35
0.85
0
200
400
600
800
1000
1500
[NaCl] (mM)

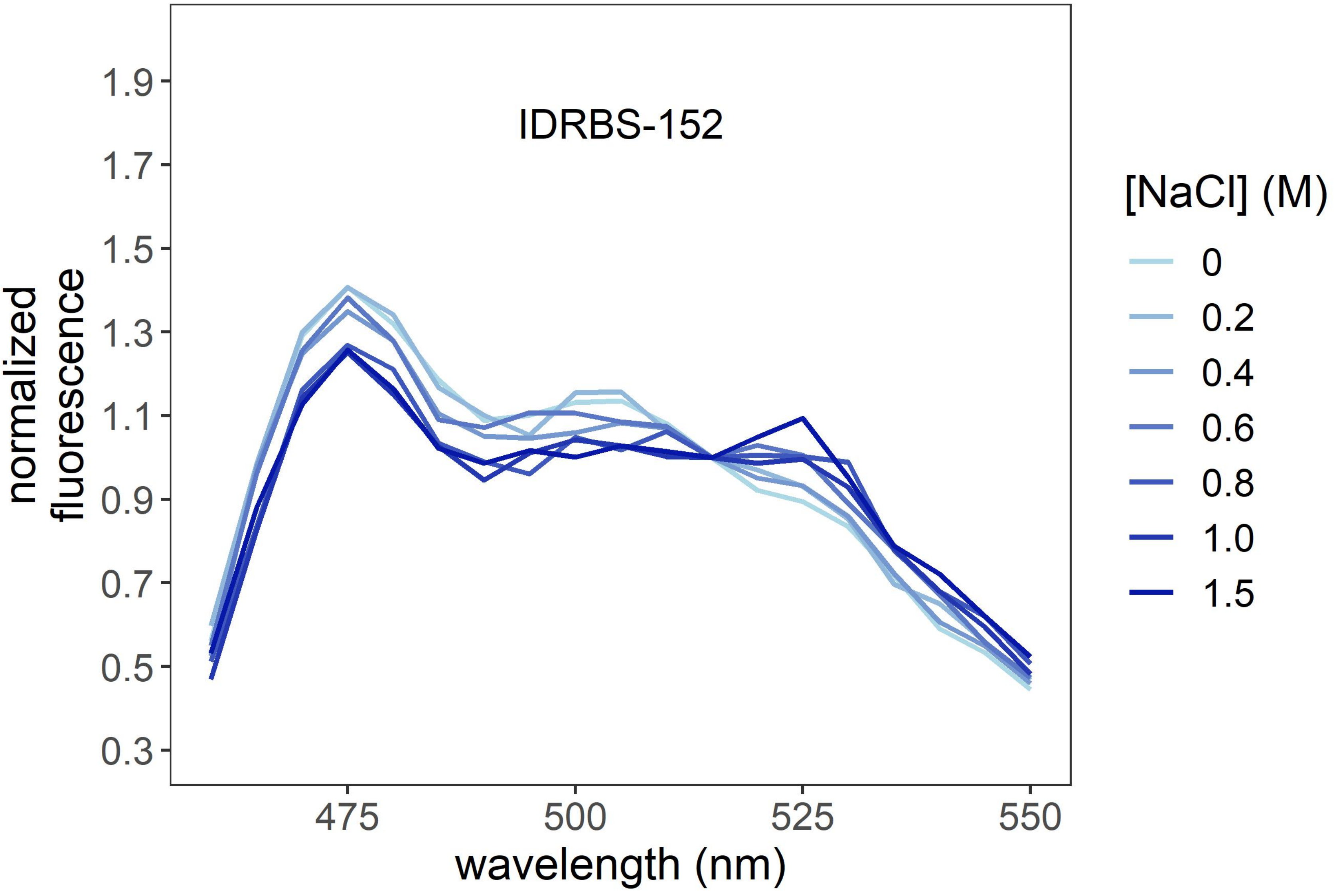
IDRBS-152
normalized
fluorescence
1.9
1.7
1.5
1.3
1.1
0.9
0.7
0.5
0.3
475
500
525
550
wavelength (nm)
[NaCl] (M)
0
0.2
0.4
0.6
0.8
1.0
1.5

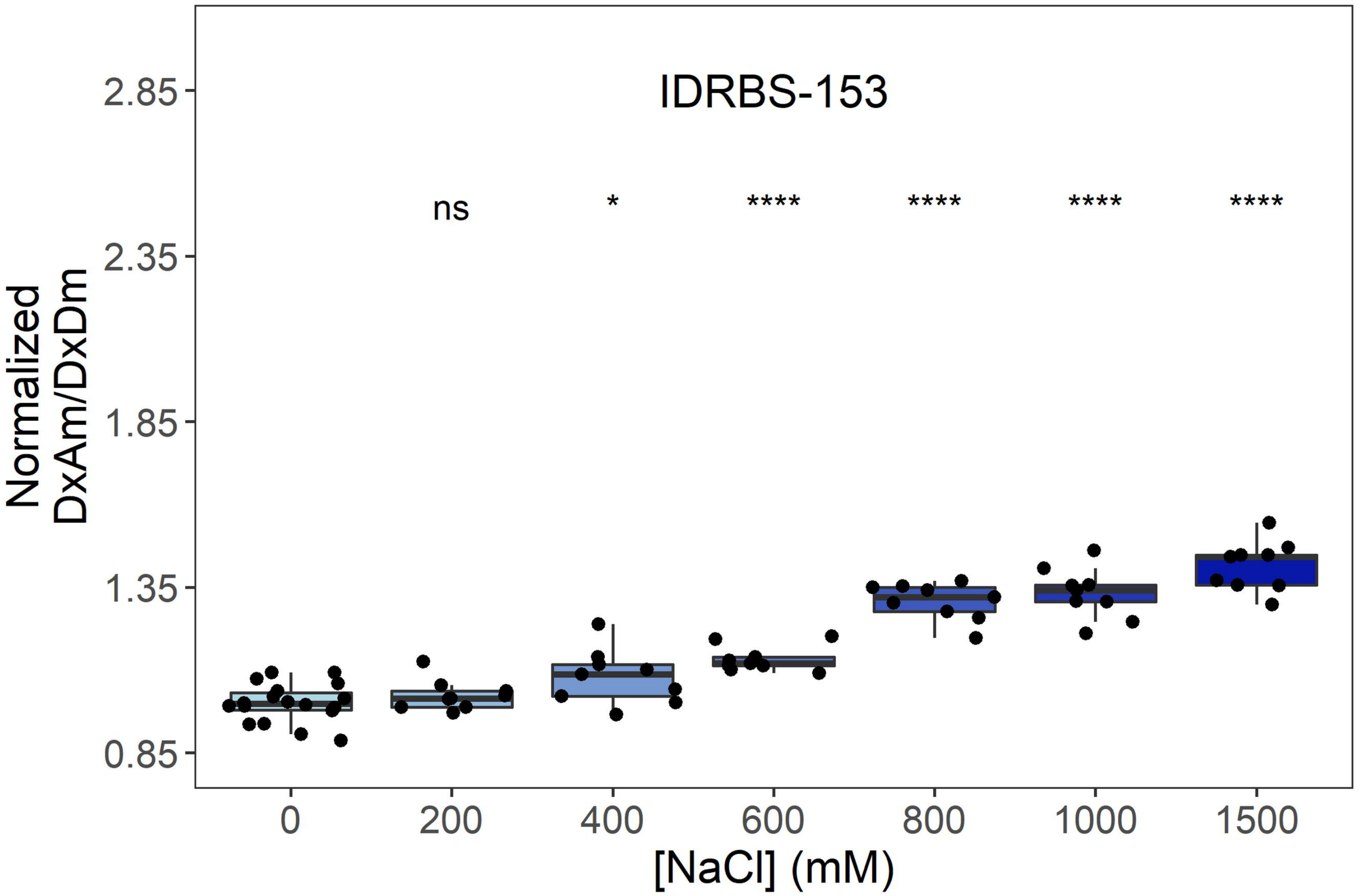
IDRBS-153
ns
*
****
****
****
****
2.85
2.35
1.85
1.35
0.85
Normalized
DxAm/DxDm
0
200
400
600
800
1000
1500
[NaCl] (mM)

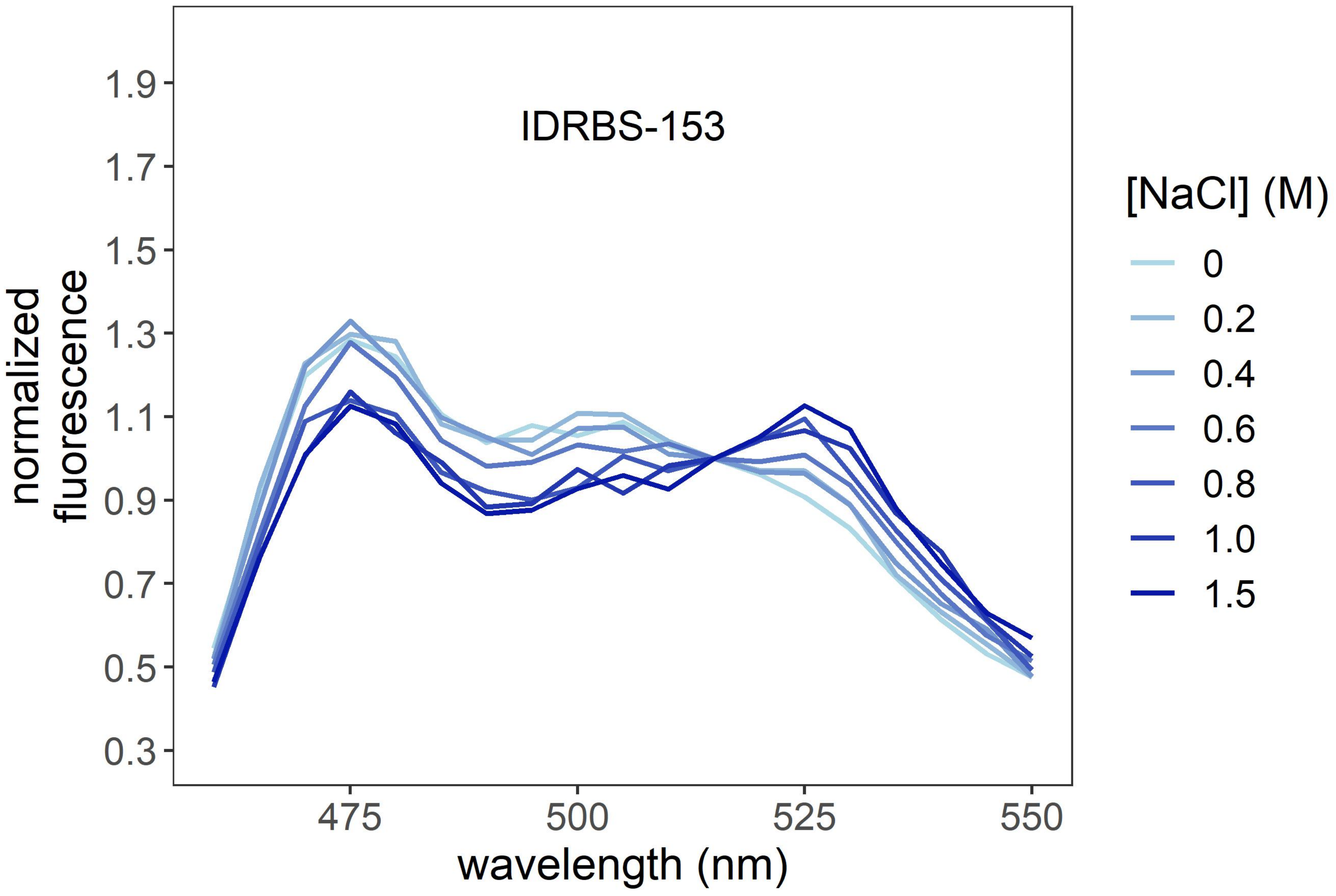
IDRBS-153
1.9
1.7
1.5
1.3
1.1
0.9
0.7
0.5
0.3
normalized
fluorescence
475
500
525
550
wavelength (nm)
[NaCl] (M)
0
0.2
0.4
0.6
0.8
1.0
1.5

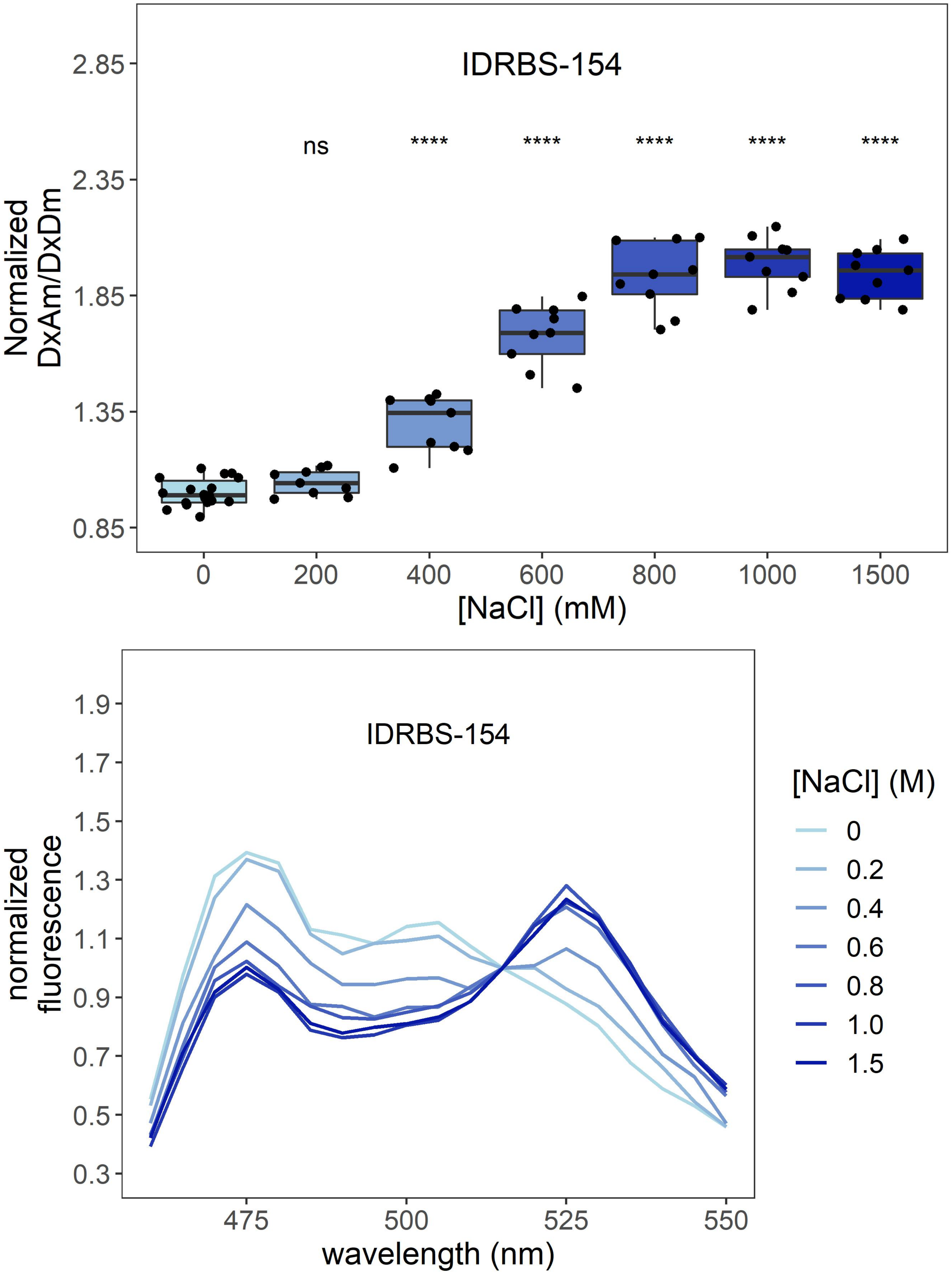
IDRBS-154
ns
****
****
****
****
****
Normalized
DxAm/DxDm
2.85
2.35
1.85
1.35
0.85
0
200
400
600
800
1000
1500
[NaCl] (mM)
IDRBS-154
normalized
fluorescence
1.9
1.7
1.5
1.3
1.1
0.9
0.7
0.5
0.3
475
500
525
550
wavelength (nm)
[NaCl] (M)
0
0.2
0.4
0.6
0.8
1.0
1.5

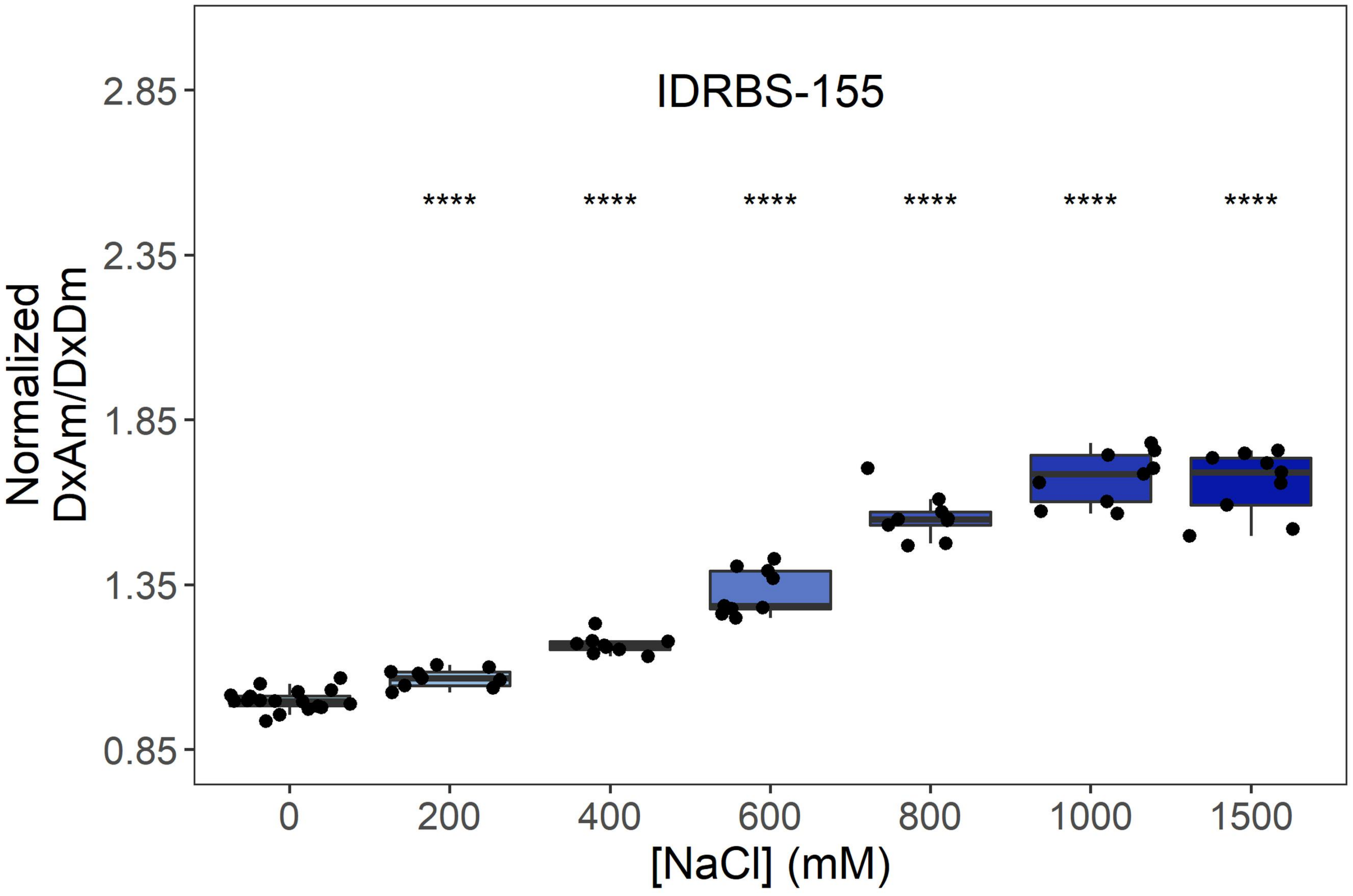

IDRBS-155
Normalized DxAm/DxDm
2.85
2.35
1.85
1.35
0.85
****
0
200
400
600
800
1000
1500
[NaCl] (mM)


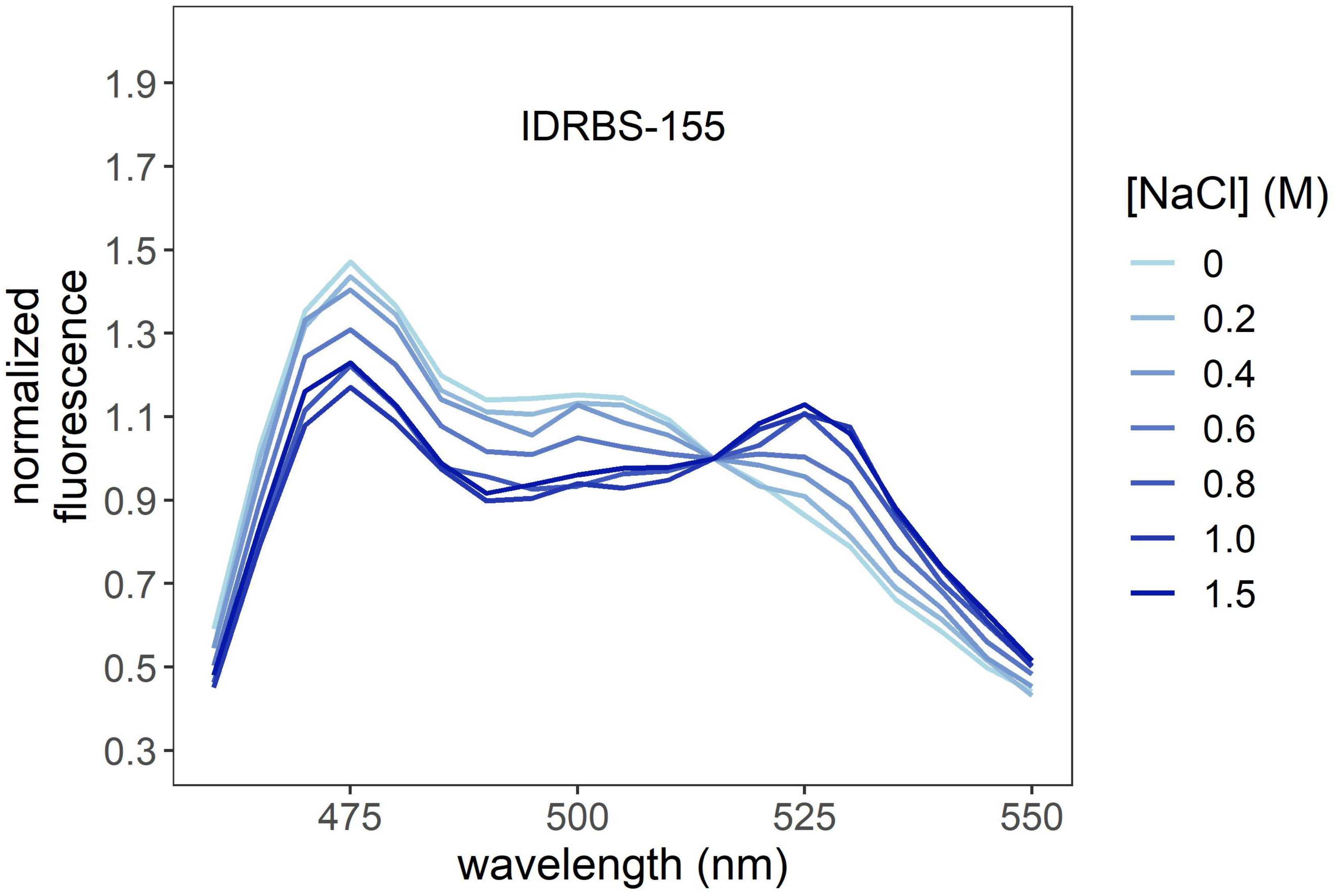

IDRBS-155
normalized fluorescence
1.9
1.7
1.5
1.3
1.1
0.9
0.7
0.5
0.3
475
500
525
550
wavelength (nm)
[NaCl] (M)
0
0.2
0.4
0.6
0.8
1.0
1.5

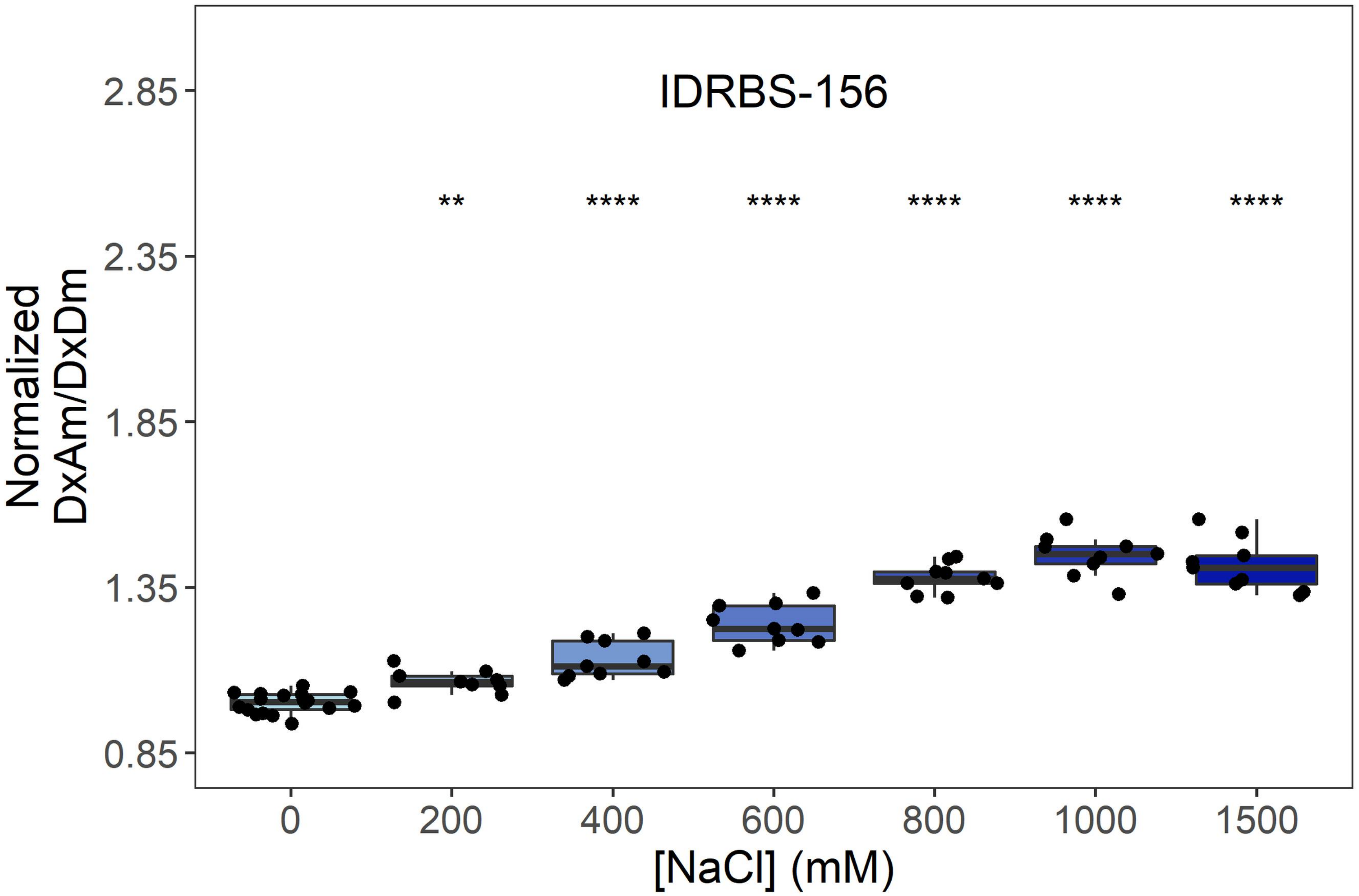
IDRBS-156
**
****
****
****
****
****
Normalized DxAm/DxDm
0.85
1.35
1.85
2.35
2.85
0
200
400
600
800
1000
1500
[NaCl] (mM)

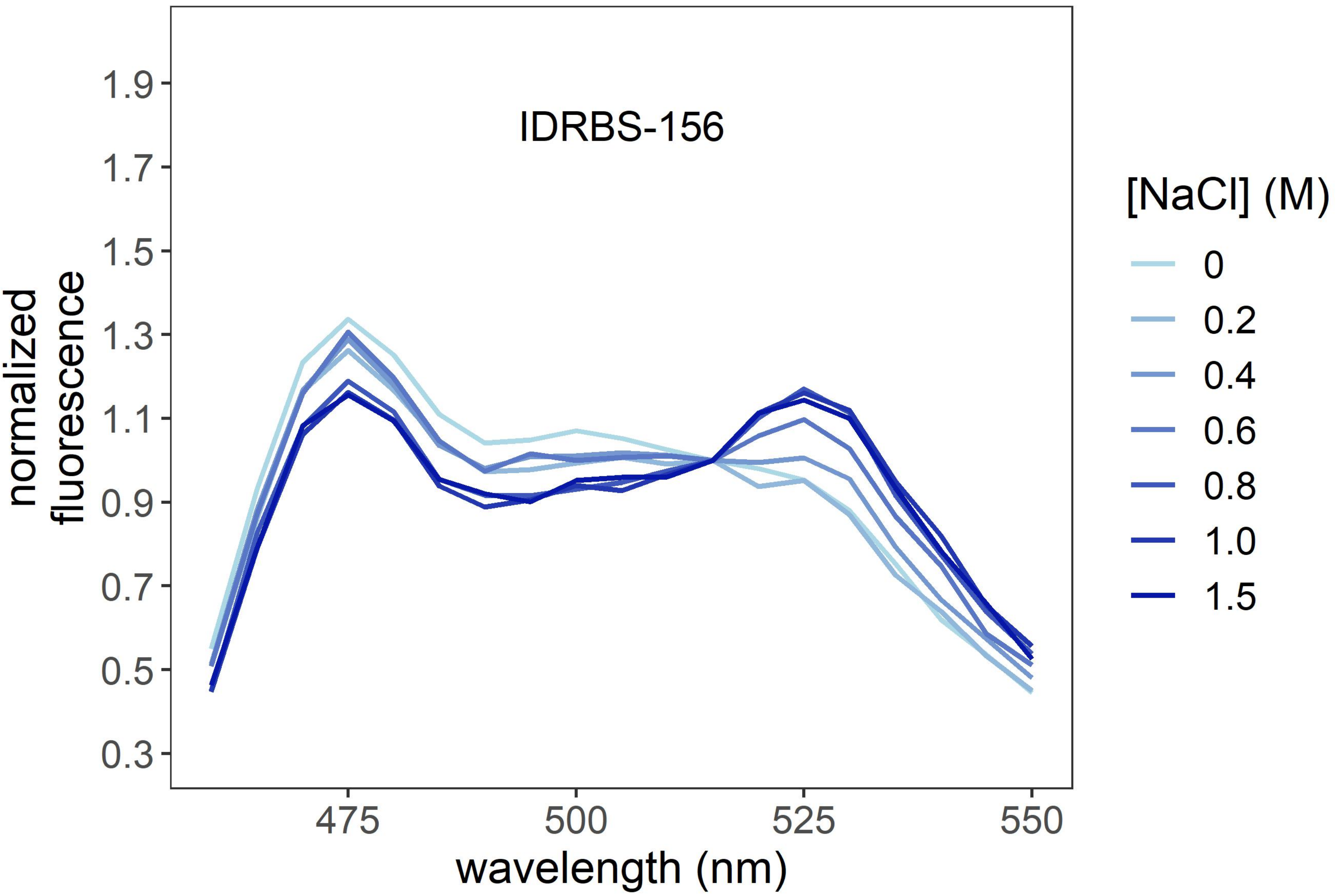
IDRBS-156
normalized fluorescence
0.3
0.5
0.7
0.9
1.1
1.3
1.5
1.7
1.9
475
500
525
550
wavelength (nm)
[NaCl] (M)
0
0.2
0.4
0.6
0.8
1.0
1.5

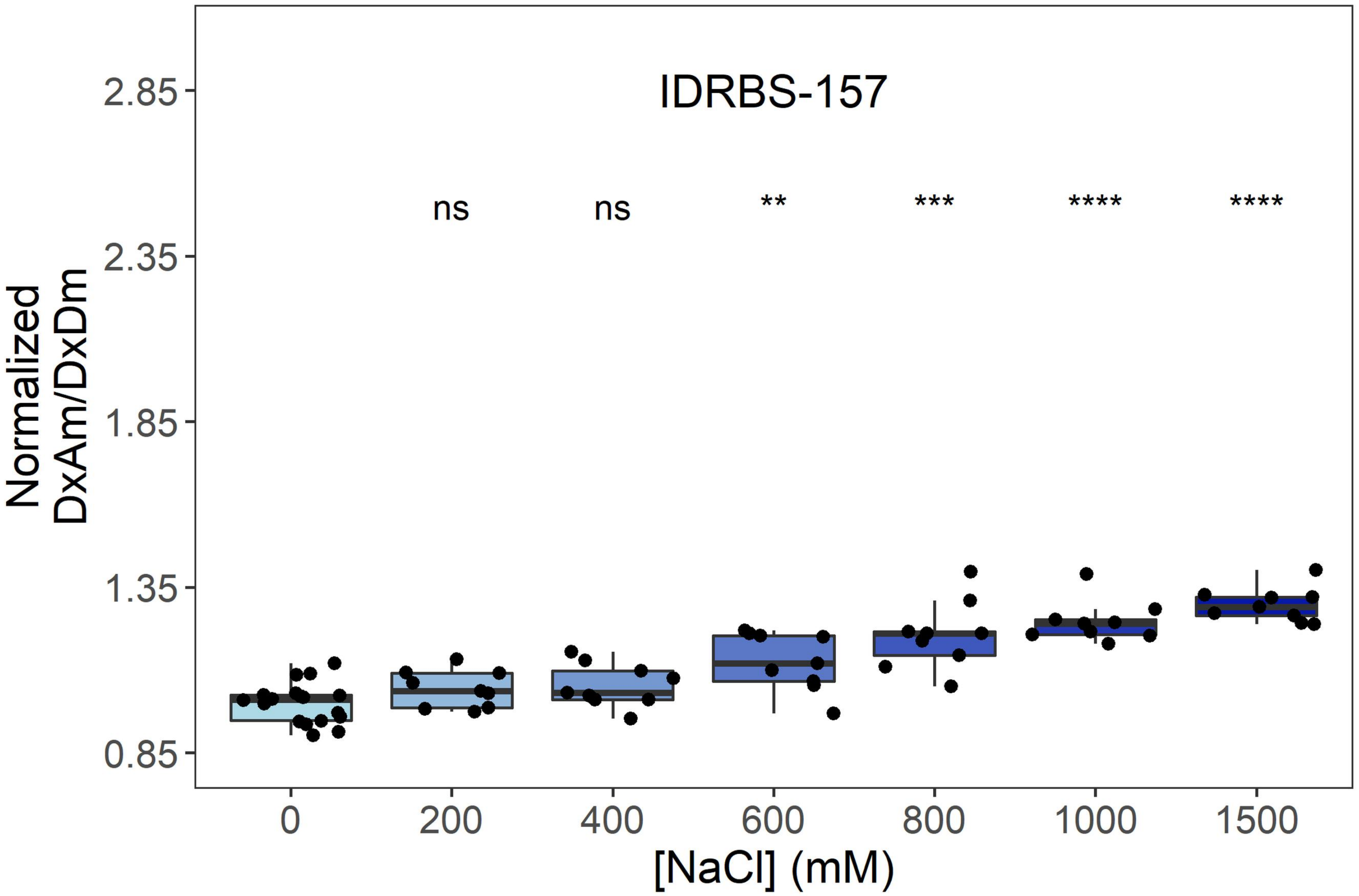
IDRBS-157
ns
ns
**
***
****
****
2.85
2.35
1.85
1.35
0.85
Normalized
DxAm/DxDm
0
200
400
600
800
1000
1500
[NaCl] (mM)

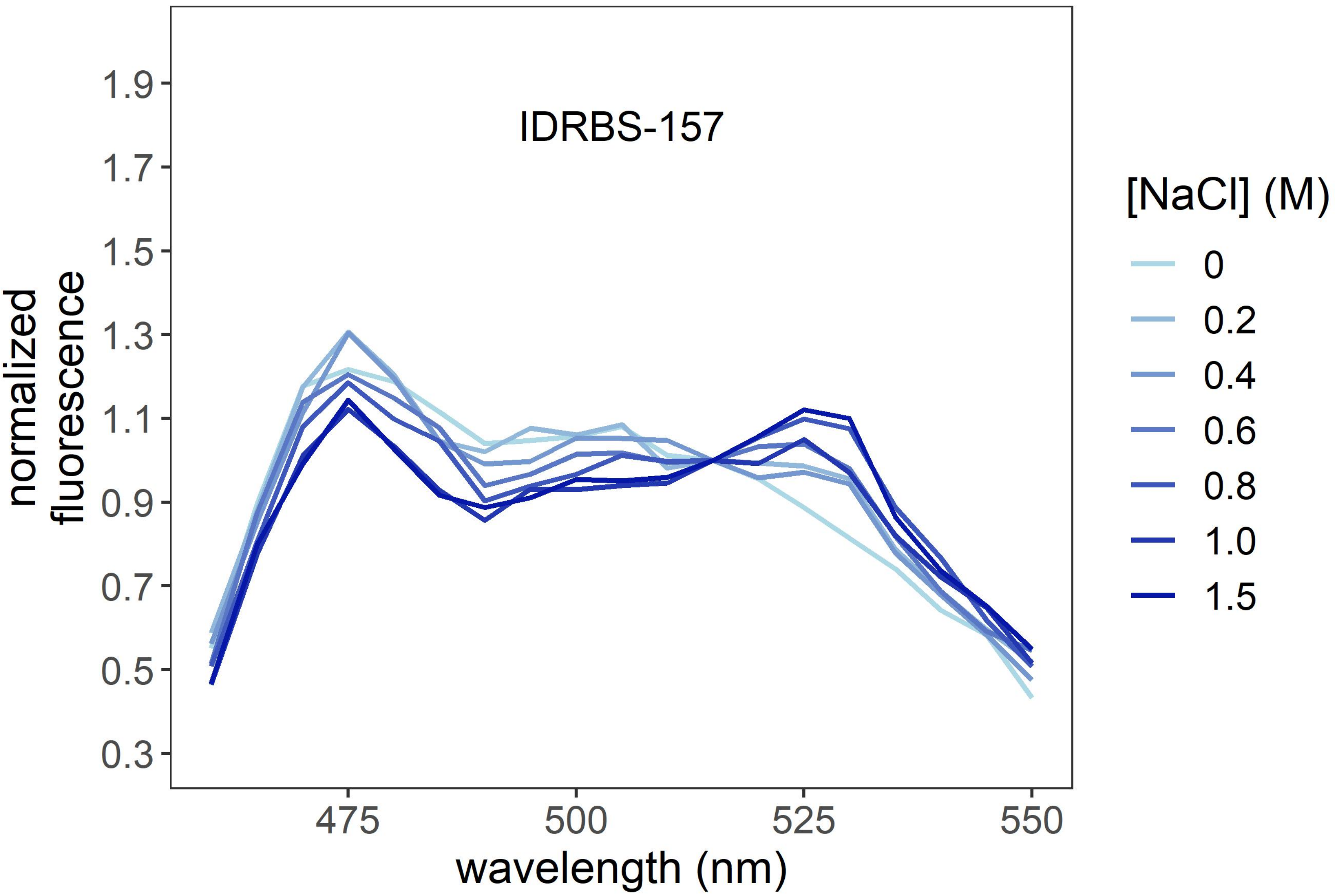
IDRBS-157
1.9
1.7
1.5
1.3
1.1
0.9
0.7
0.5
0.3
normalized
fluorescence
475
500
525
550
wavelength (nm)
[NaCl] (M)
0
0.2
0.4
0.6
0.8
1.0
1.5

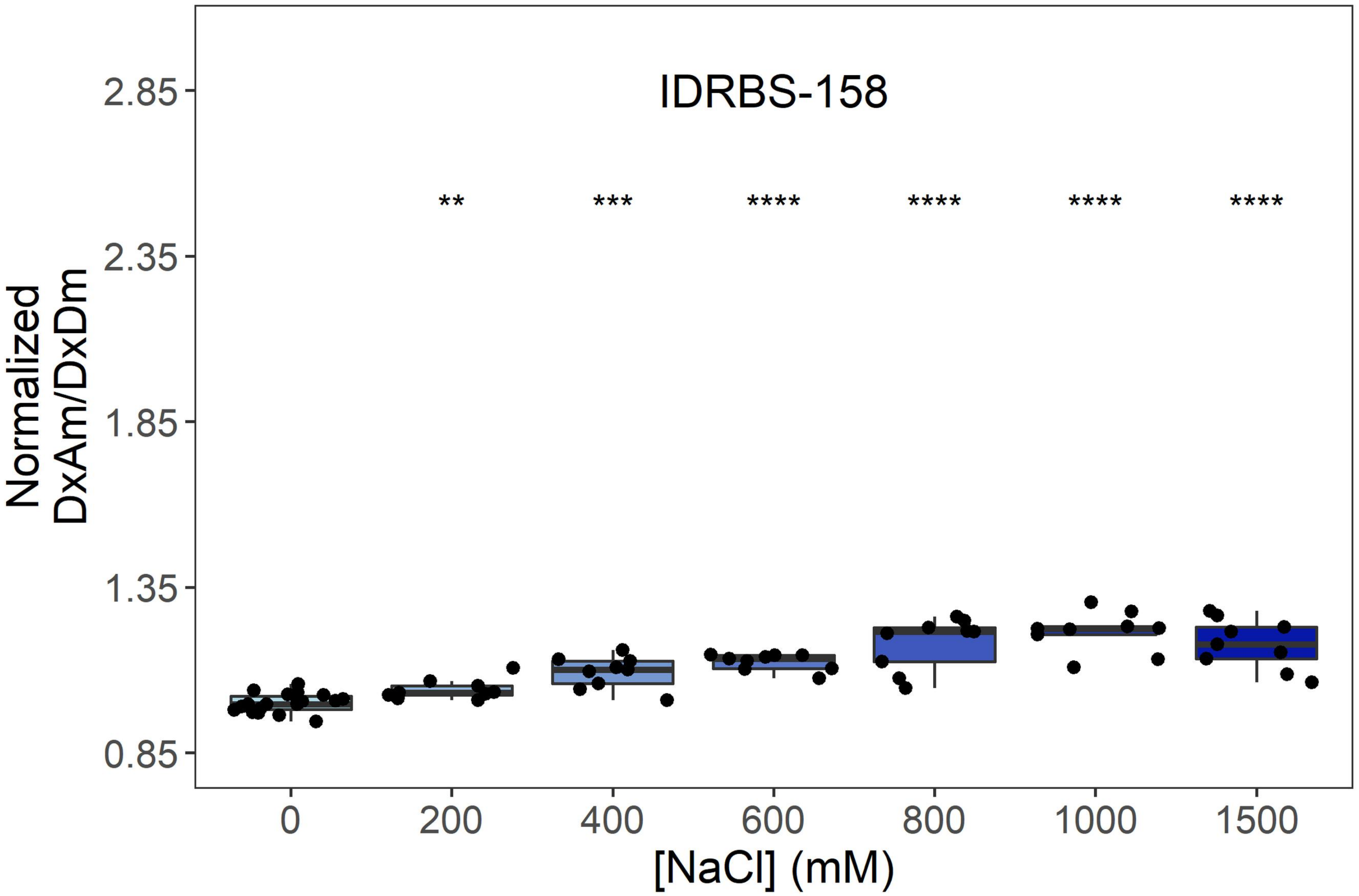
IDRBS-158
**
***
****
****
****
****
Normalized
DxAm/DxDm
2.85
2.35
1.85
1.35
0.85
0
200
400
600
800
1000
1500
[NaCl] (mM)

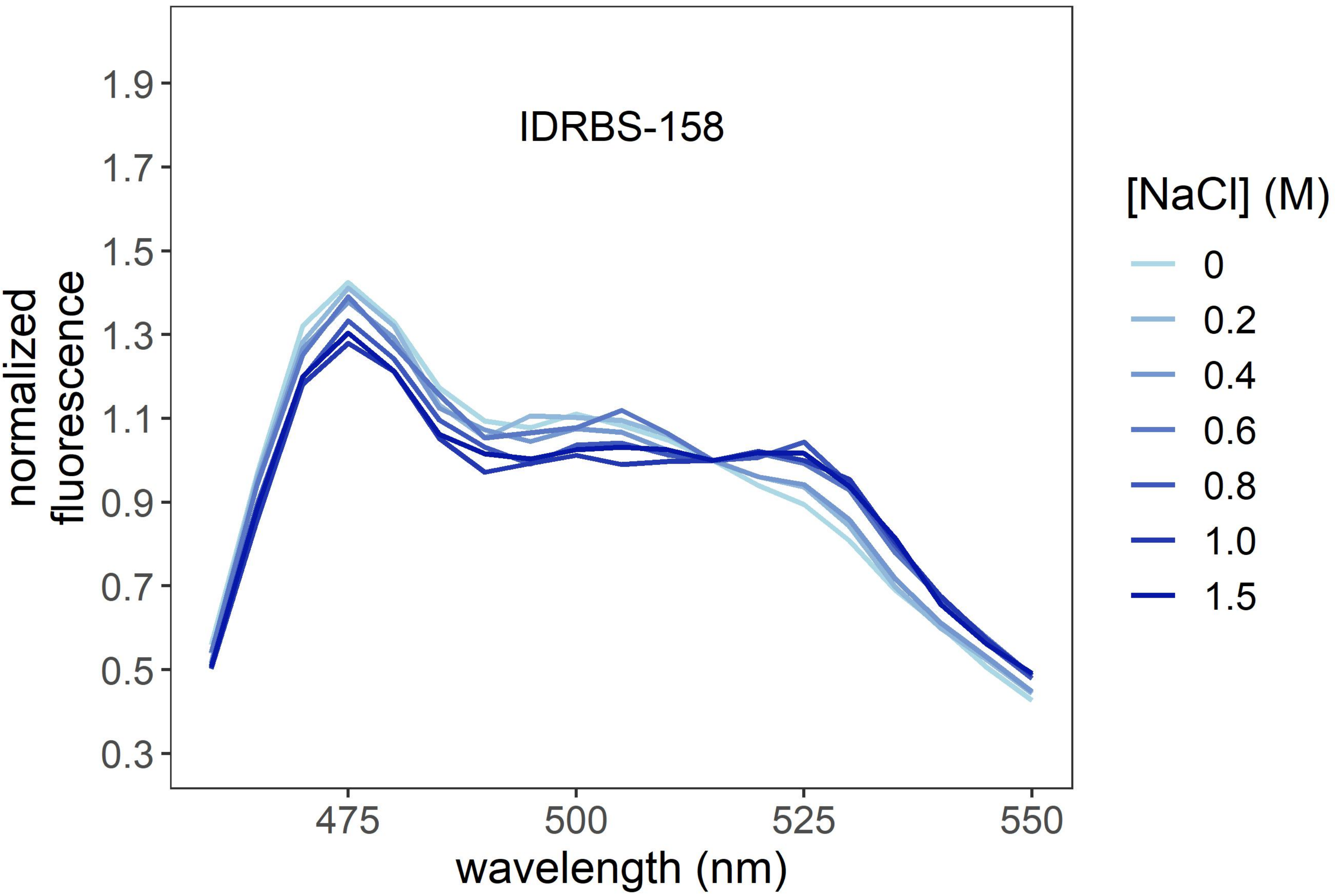
IDRBS-158
normalized
fluorescence
1.9
1.7
1.5
1.3
1.1
0.9
0.7
0.5
0.3
475
500
525
550
wavelength (nm)
[NaCl] (M)
0
0.2
0.4
0.6
0.8
1.0
1.5

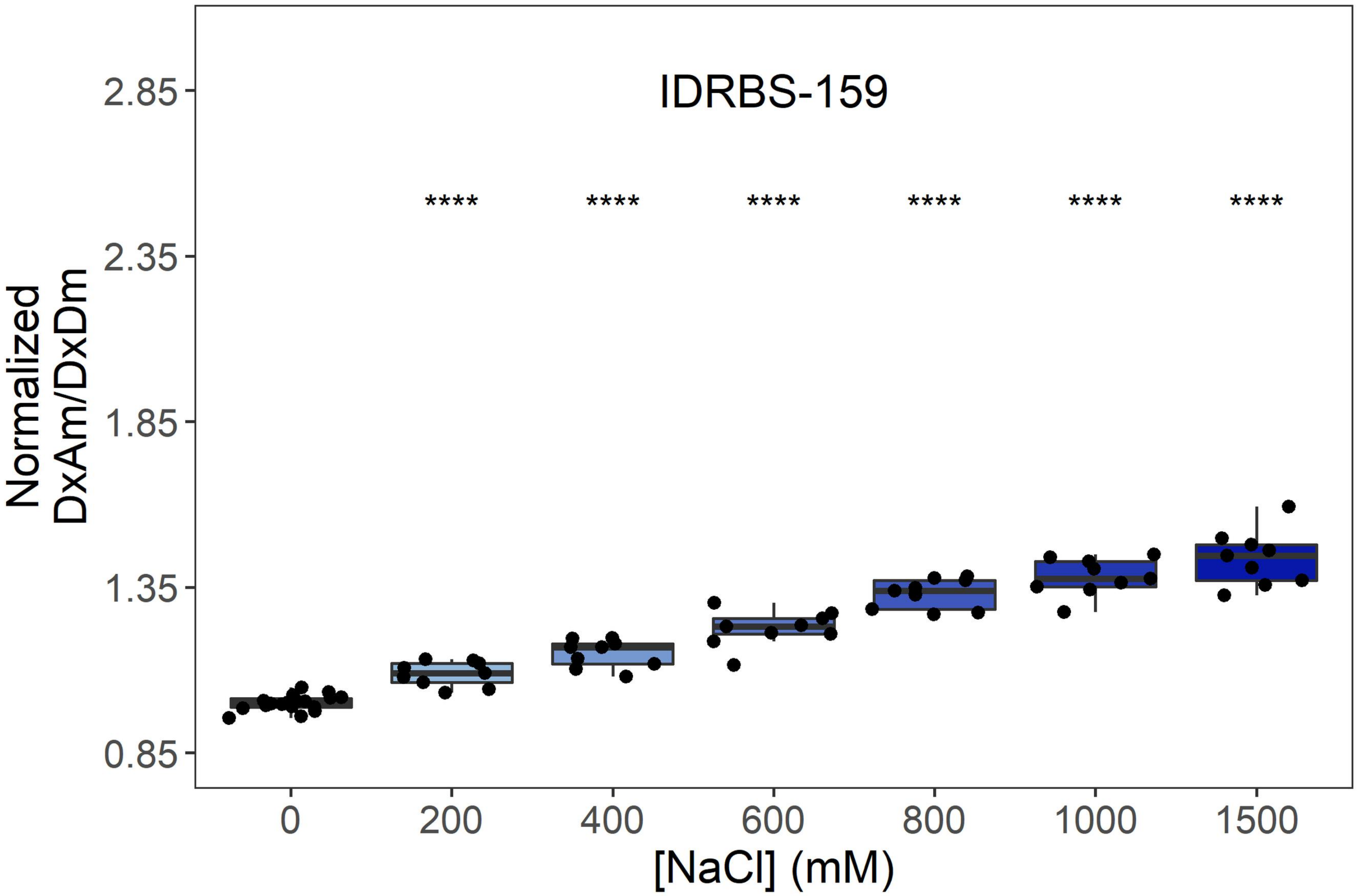
IDRBS-159
****
Normalized
DxAm/DxDm
[NaCl] (mM)
0
200
400
600
800
1000
1500
0.85
1.35
1.85
2.35
2.85

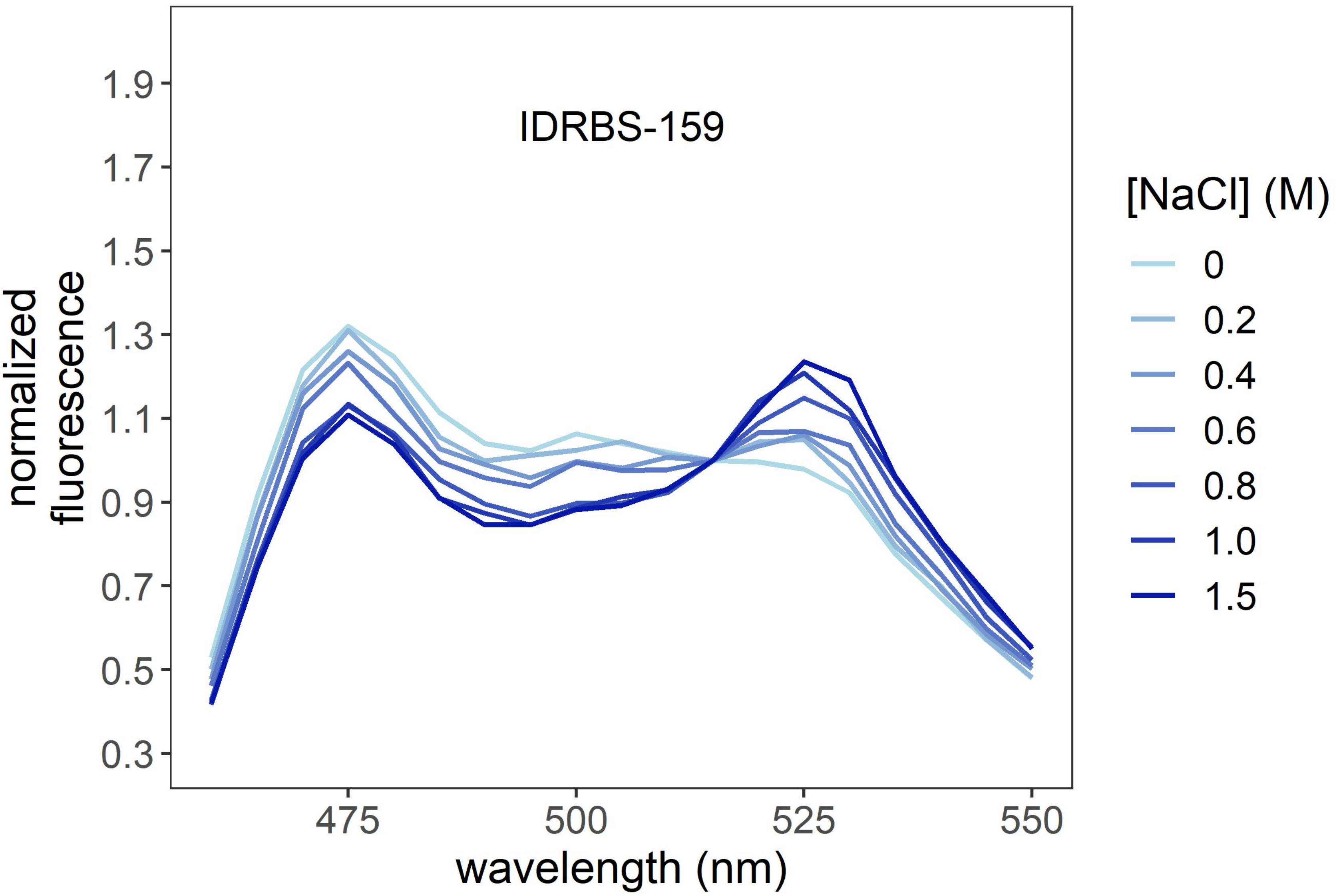
IDRBS-159
normalized
fluorescence
wavelength (nm)
475
500
525
550
0.3
0.5
0.7
0.9
1.1
1.3
1.5
1.7
1.9
[NaCl] (M)
0
0.2
0.4
0.6
0.8
1.0
1.5

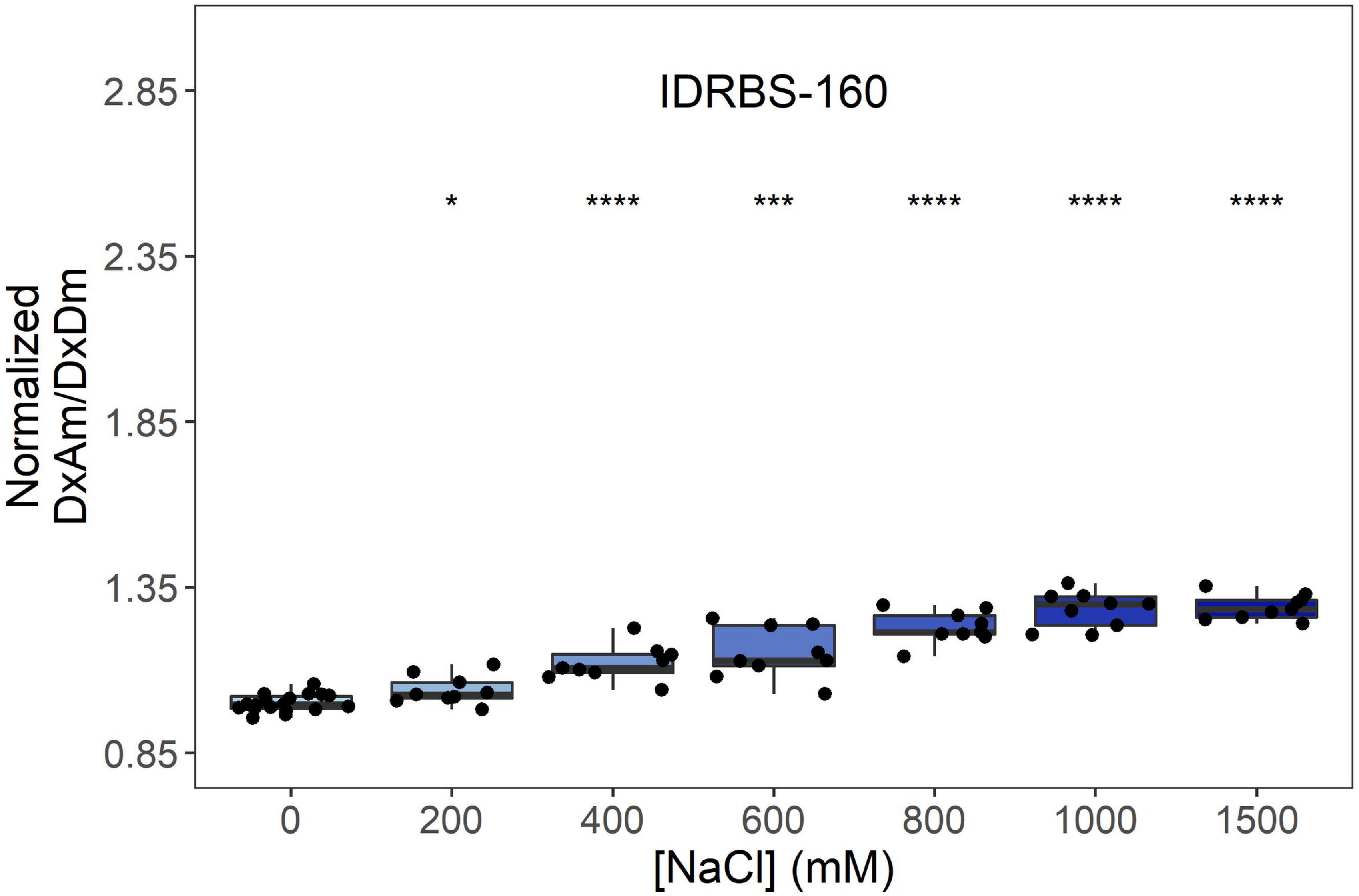
IDRBS-160
Normalized DxAm/DxDm
2.85
2.35
1.85
1.35
0.85
*
****
***
****
****
****
0
200
400
600
800
1000
1500
[NaCl] (mM)

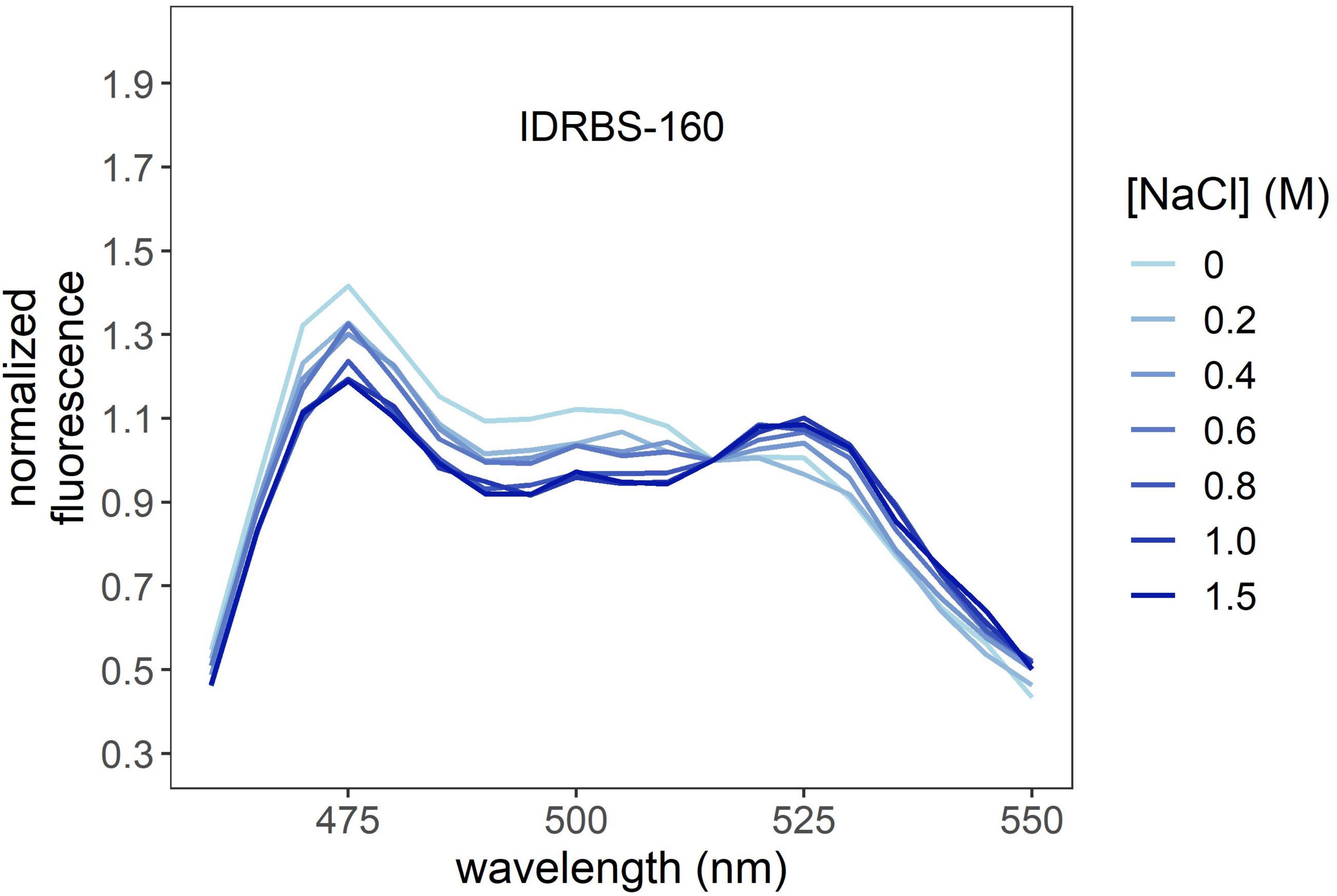
IDRBS-160
normalized fluorescence
1.9
1.7
1.5
1.3
1.1
0.9
0.7
0.5
0.3
475
500
525
550
wavelength (nm)
[NaCl] (M)
0
0.2
0.4
0.6
0.8
1.0
1.5

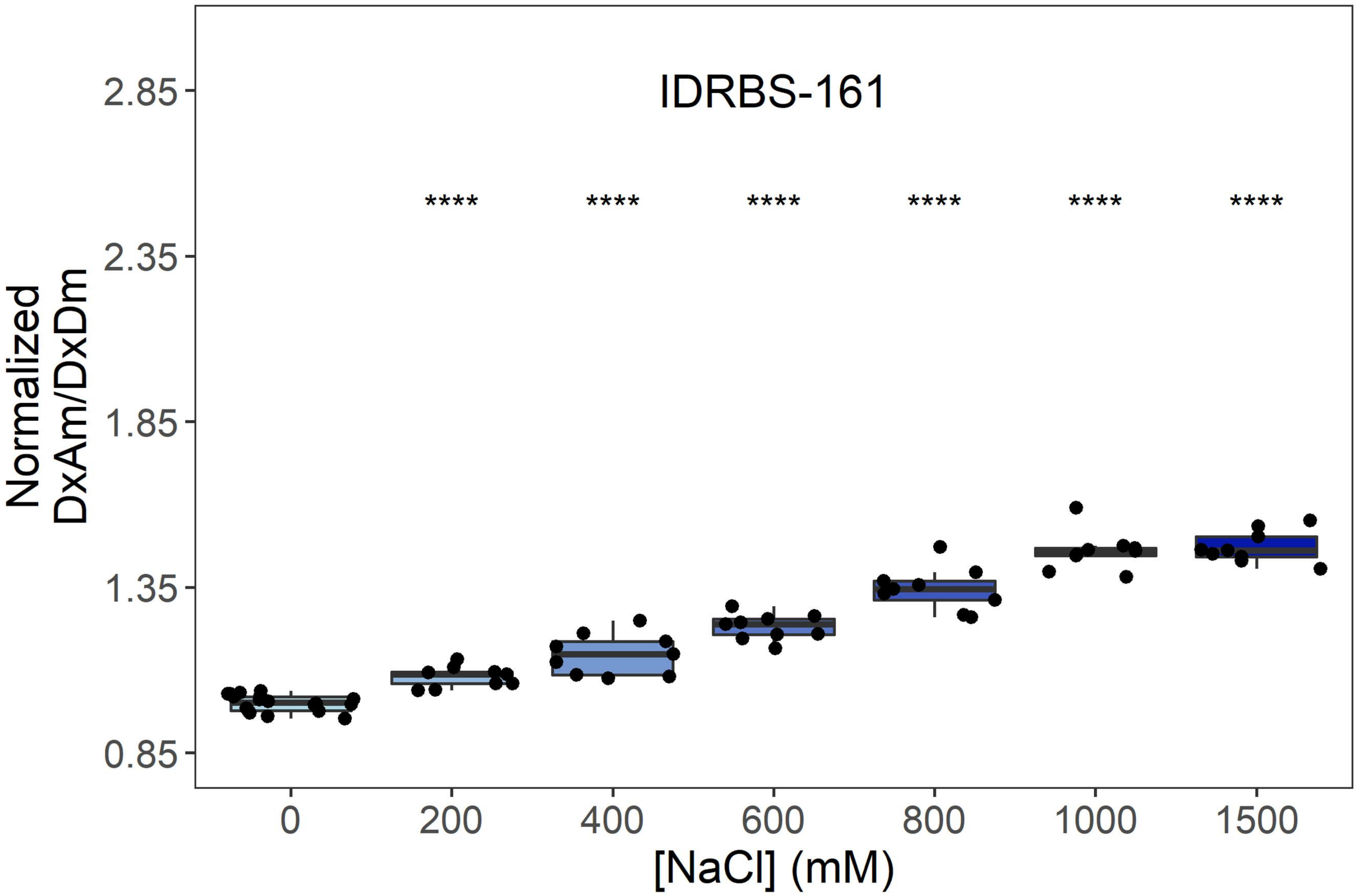
IDRBS-161
Normalized DxAm/DxDm
[NaCl] (mM)
0
200
400
600
800
1000
1500
0.85
1.35
1.85
2.35
2.85
****

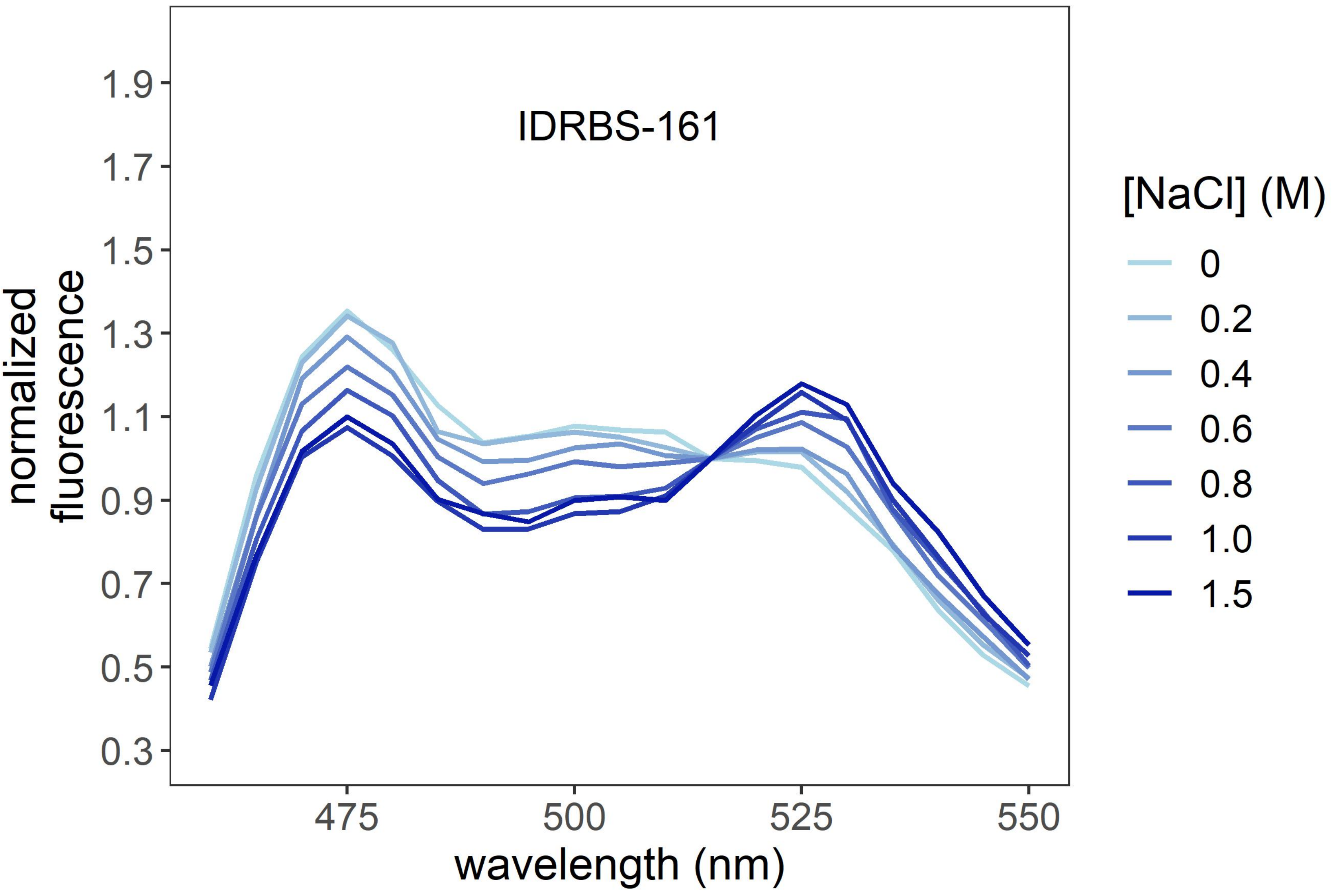
IDRBS-161
normalized fluorescence
wavelength (nm)
475
500
525
550
0.3
0.5
0.7
0.9
1.1
1.3
1.5
1.7
1.9
[NaCl] (M)
0
0.2
0.4
0.6
0.8
1.0
1.5

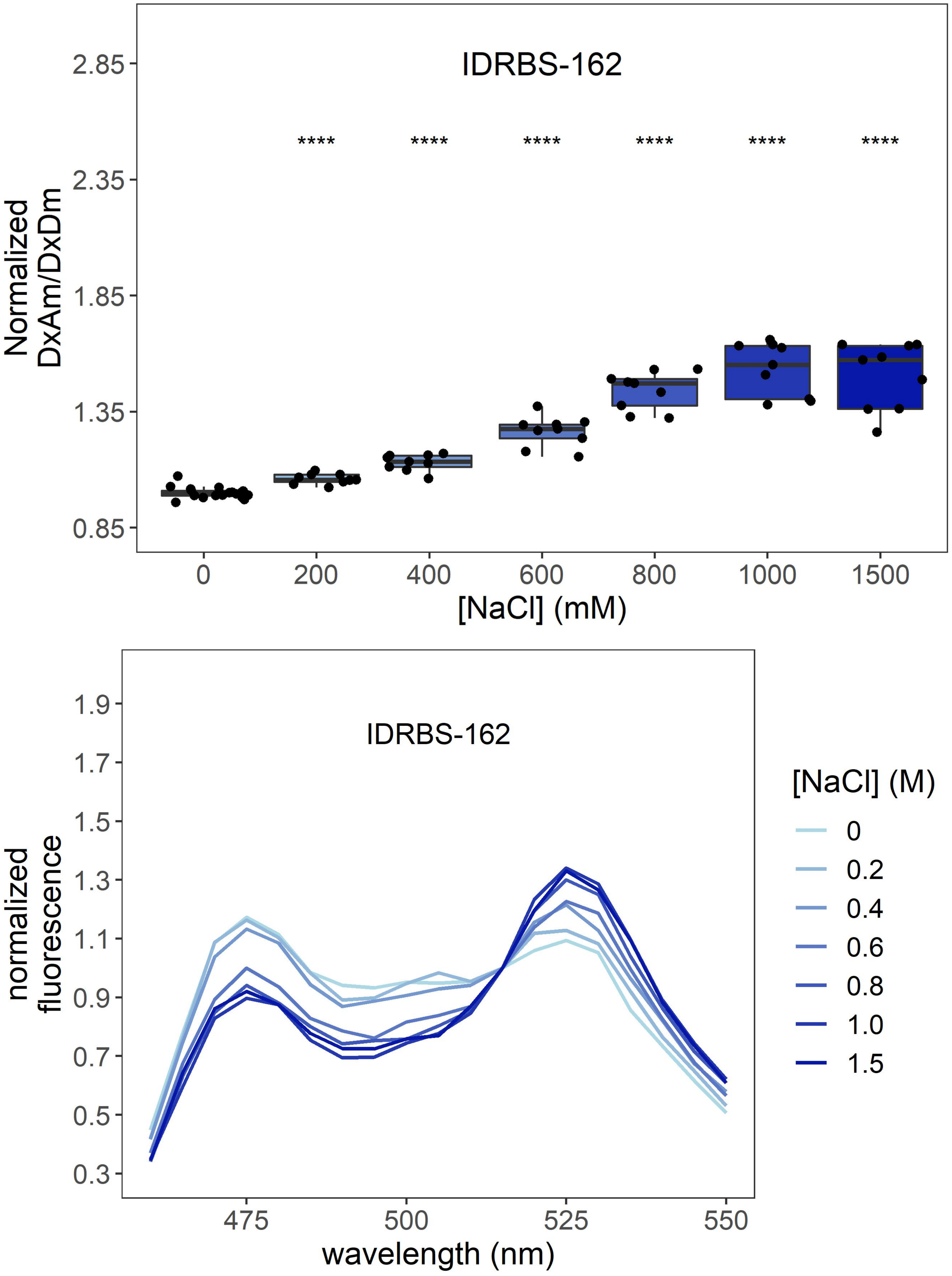

IDRBS-162
****
Normalized
DxAm/DxDm
[NaCl] (mM)
0
200
400
600
800
1000
1500
2.85
2.35
1.85
1.35
0.85
IDRBS-162
normalized
fluorescence
wavelength (nm)
475
500
525
550
[NaCl] (M)
0
0.2
0.4
0.6
0.8
1.0
1.5

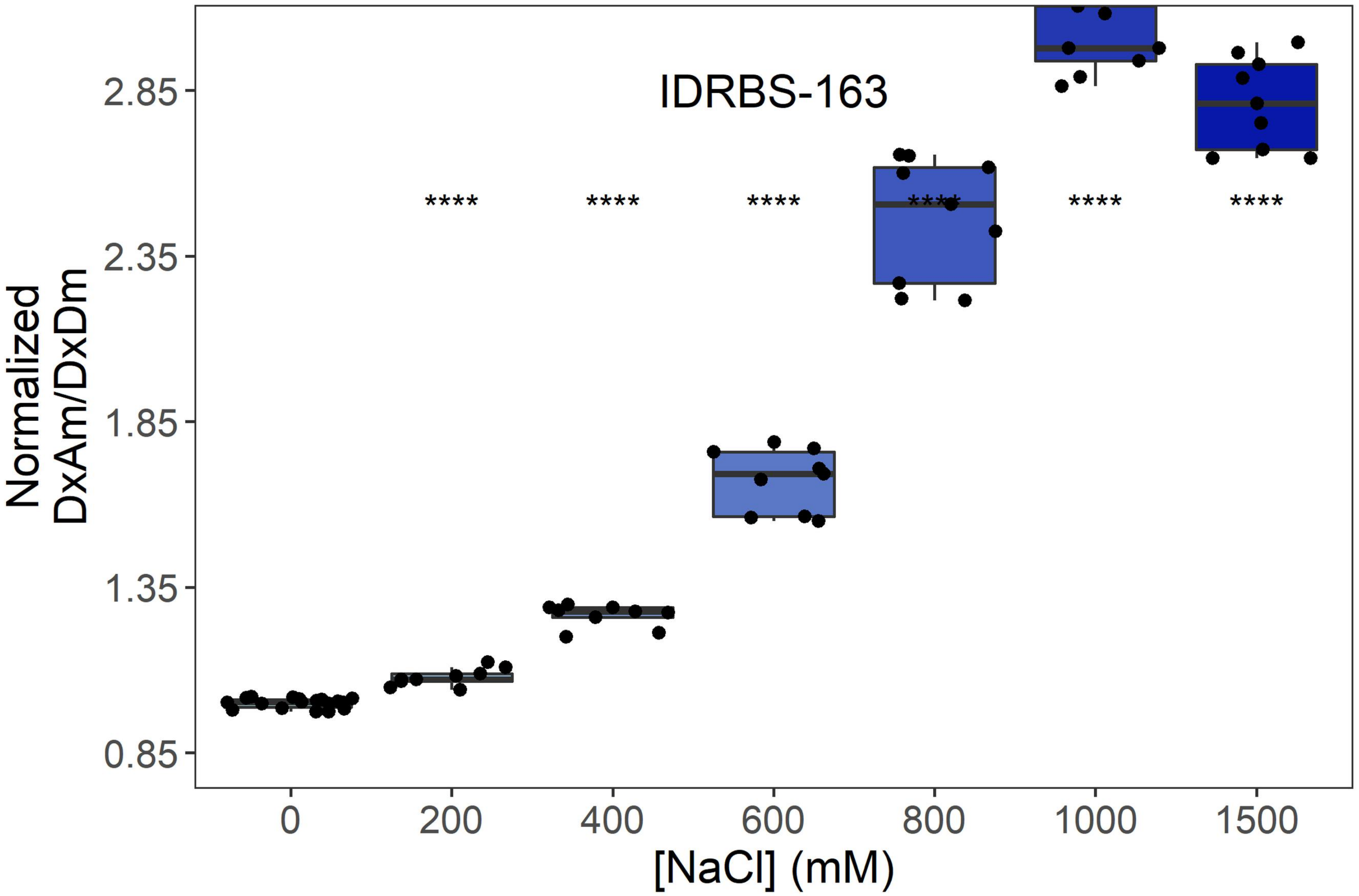
IDRBS-163
Normalized DxAm/DxDm
2.85
2.35
1.85
1.35
0.85
****
0
200
400
600
800
1000
1500
[NaCl] (mM)

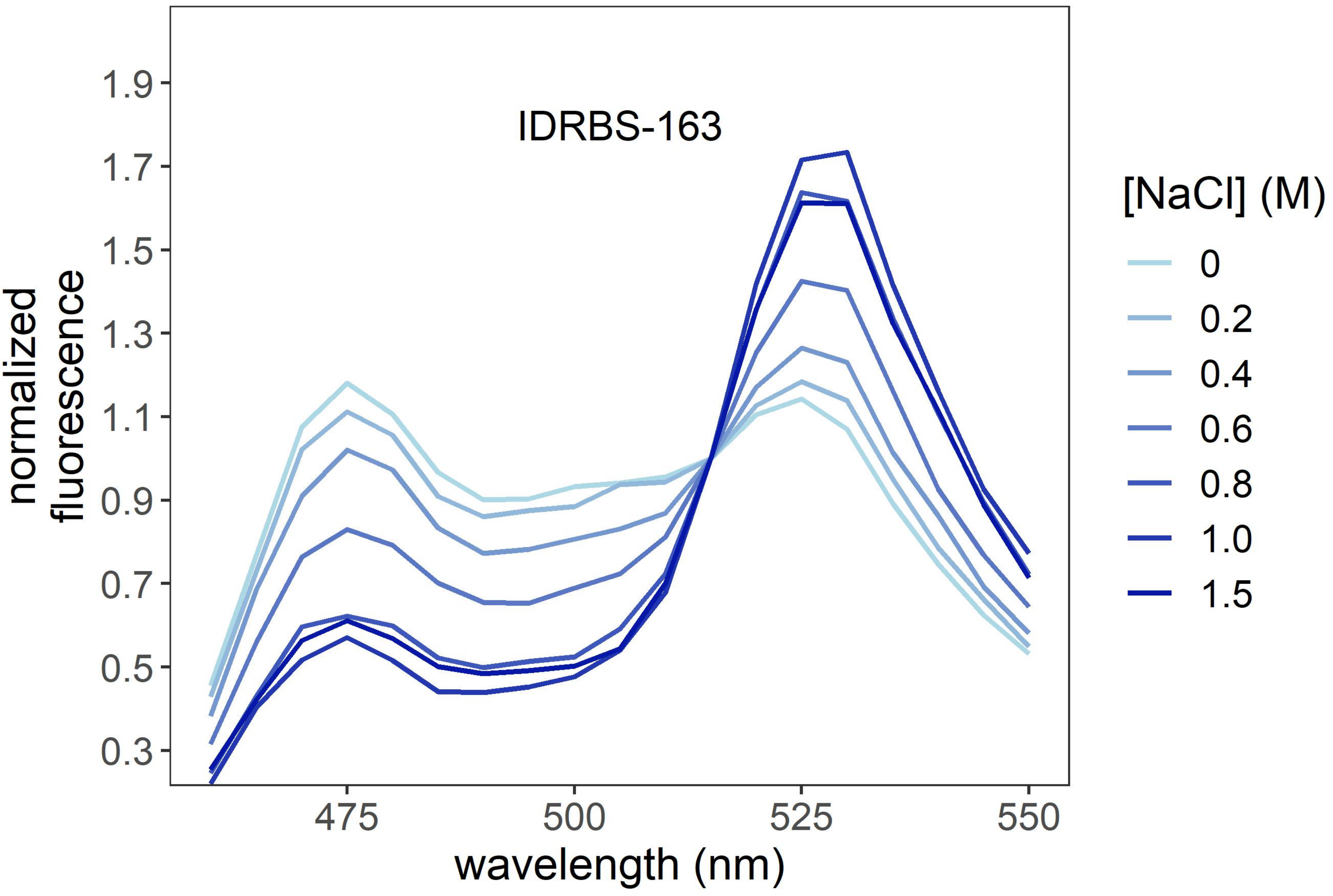
IDRBS-163
normalized fluorescence
1.9
1.7
1.5
1.3
1.1
0.9
0.7
0.5
0.3
475
500
525
550
wavelength (nm)
[NaCl] (M)
0
0.2
0.4
0.6
0.8
1.0
1.5

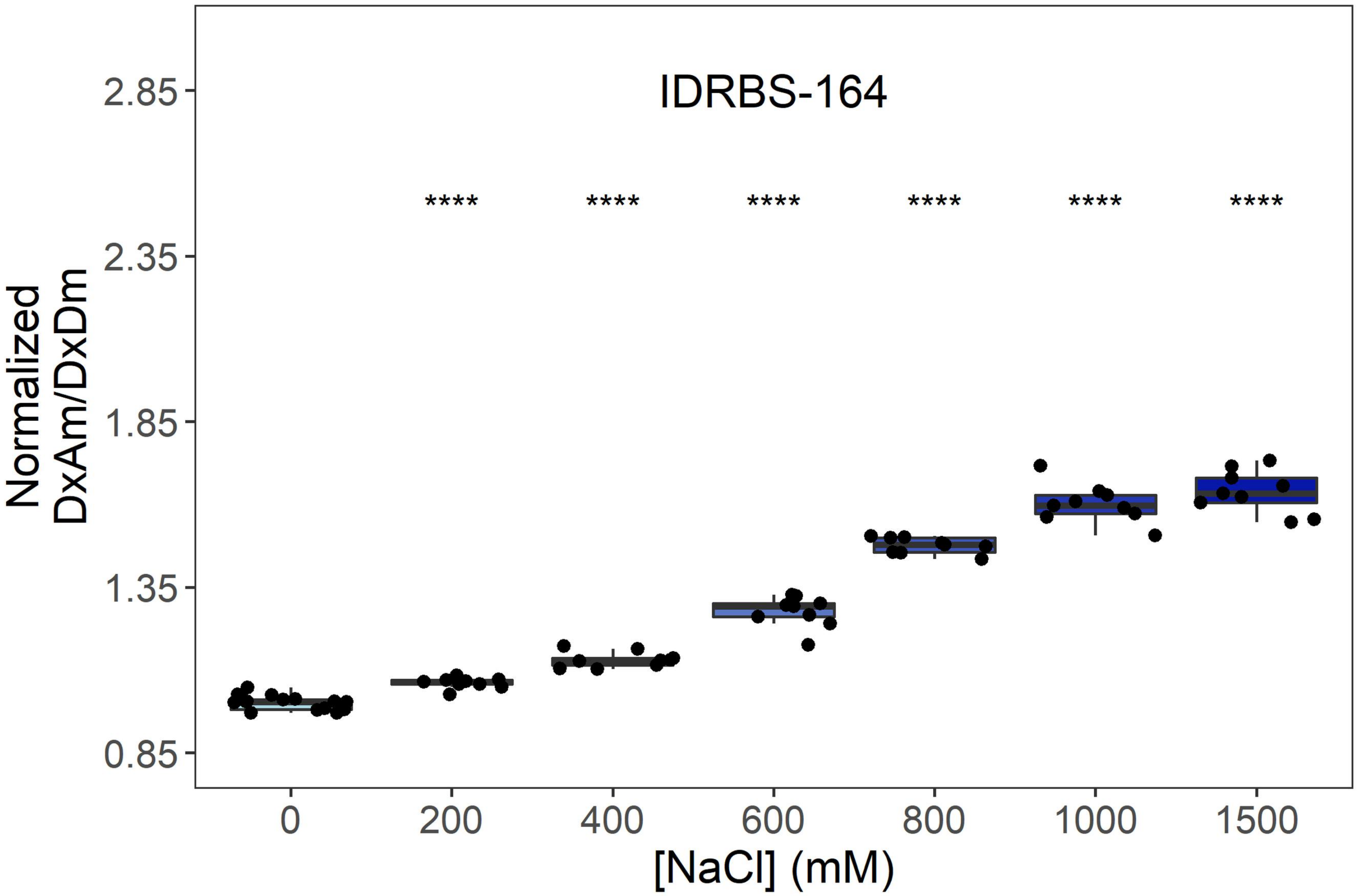

IDRBS-164
****
2.85
2.35
1.85
1.35
0.85
Normalized DxAm/DxDm
0
200
400
600
800
1000
1500
[NaCl] (mM)


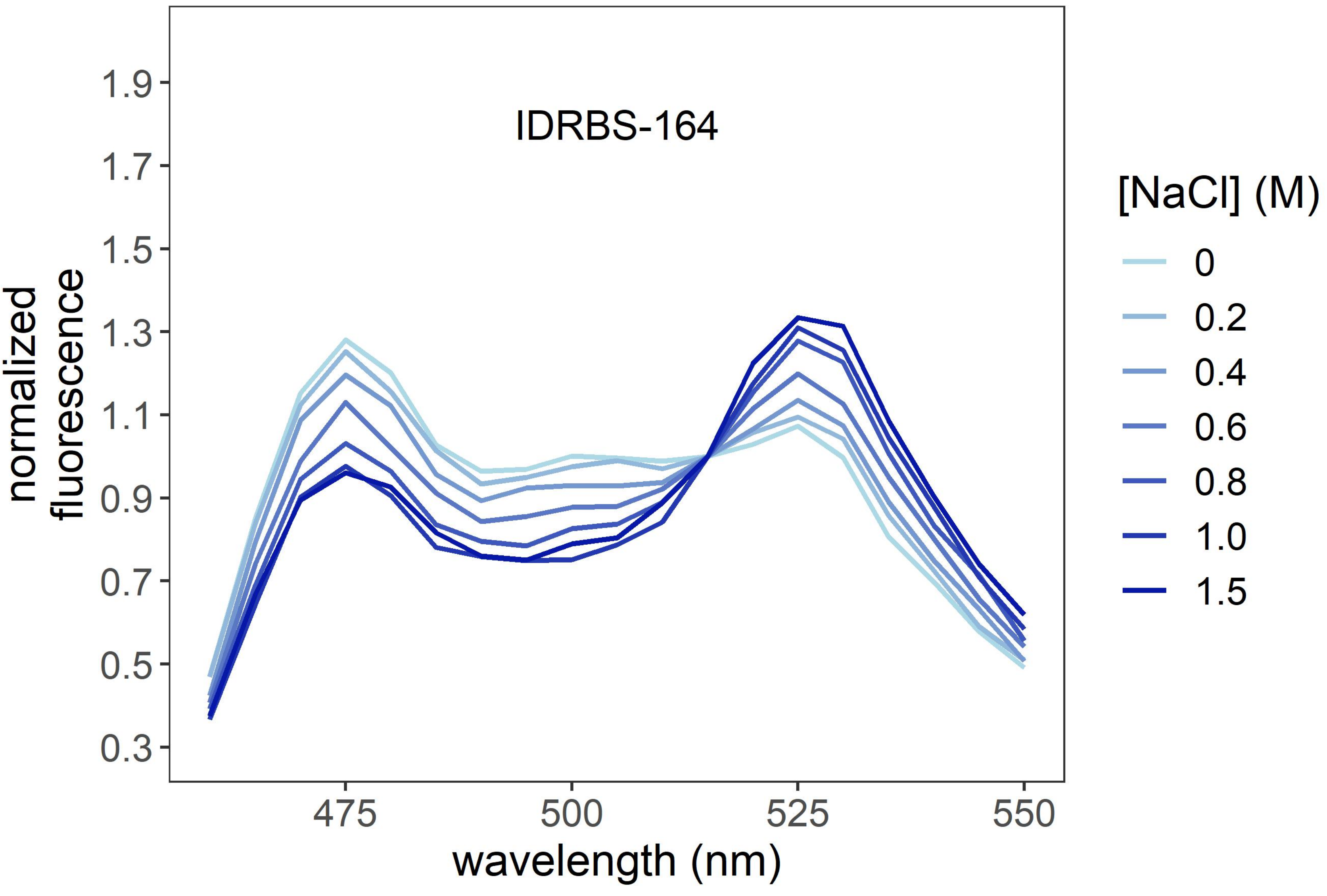

IDRBS-164
1.9
1.7
1.5
1.3
1.1
0.9
0.7
0.5
0.3
normalized fluorescence
475
500
525
550
wavelength (nm)
[NaCl] (M)
0
0.2
0.4
0.6
0.8
1.0
1.5

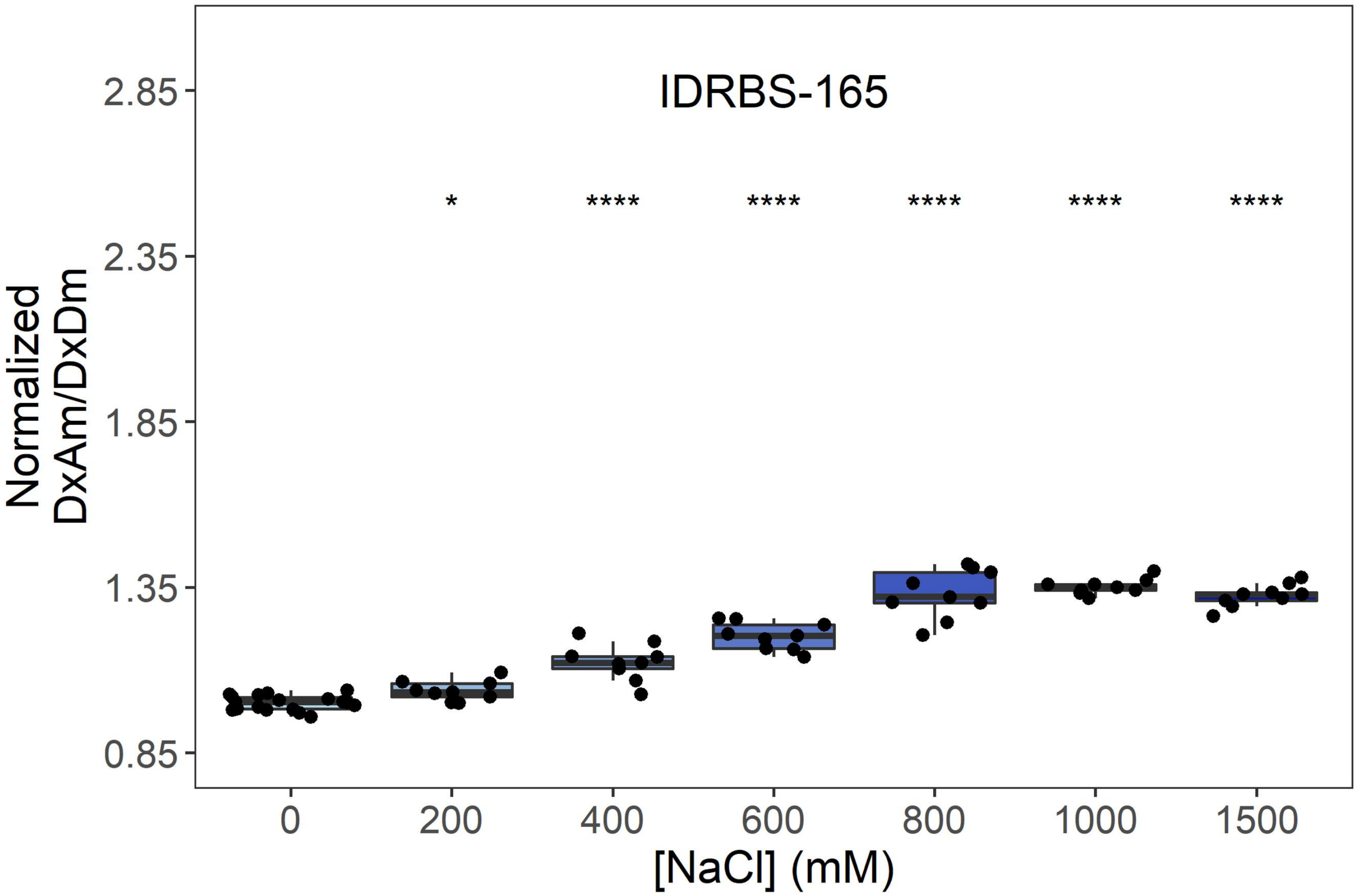
IDRBS-165
Normalized DxAm/DxDm
2.85
2.35
1.85
1.35
0.85
*
****
****
****
****
****
0
200
400
600
800
1000
1500
[NaCl] (mM)

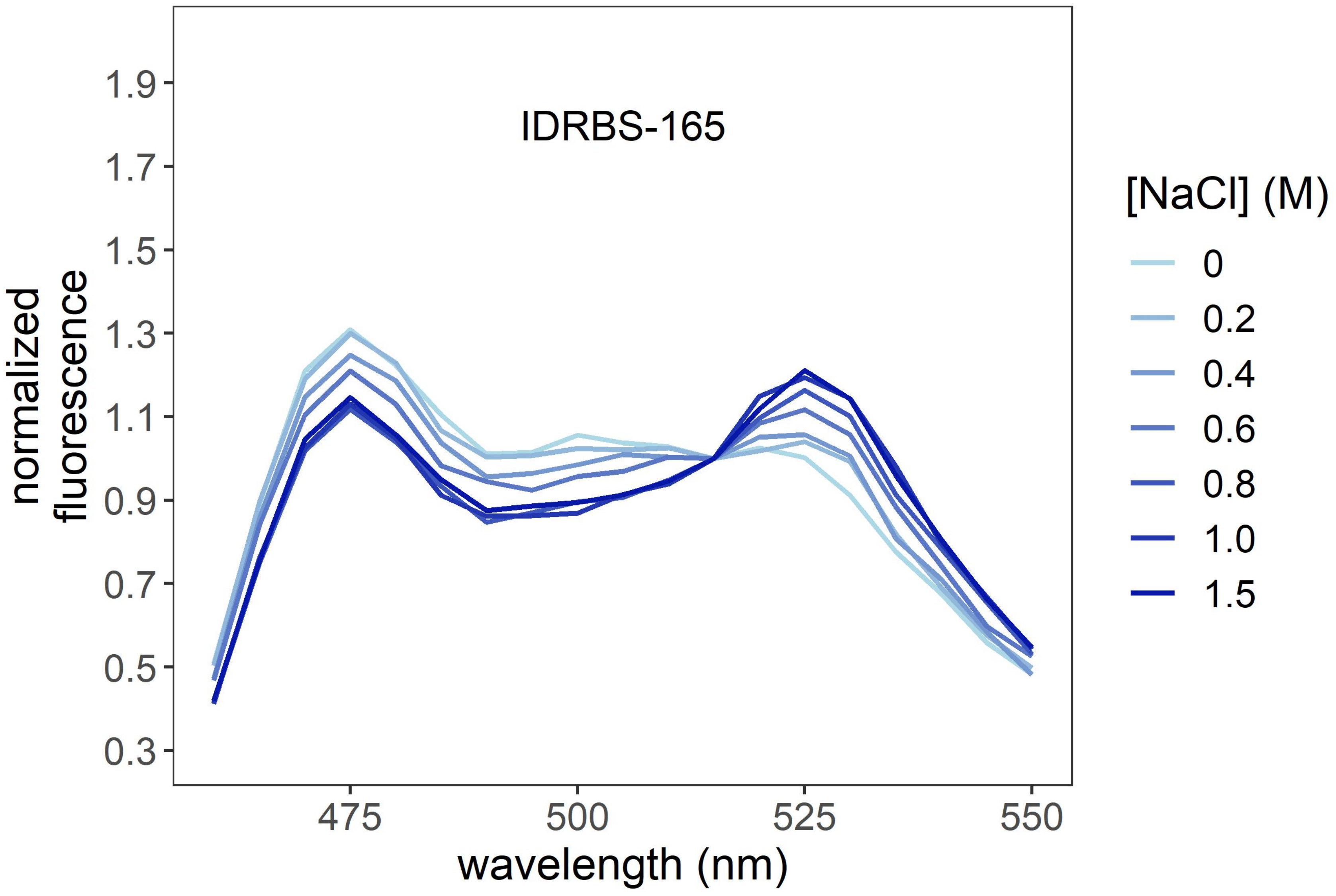
IDRBS-165
normalized fluorescence
1.9
1.7
1.5
1.3
1.1
0.9
0.7
0.5
0.3
475
500
525
550
wavelength (nm)
[NaCl] (M)
0
0.2
0.4
0.6
0.8
1.0
1.5

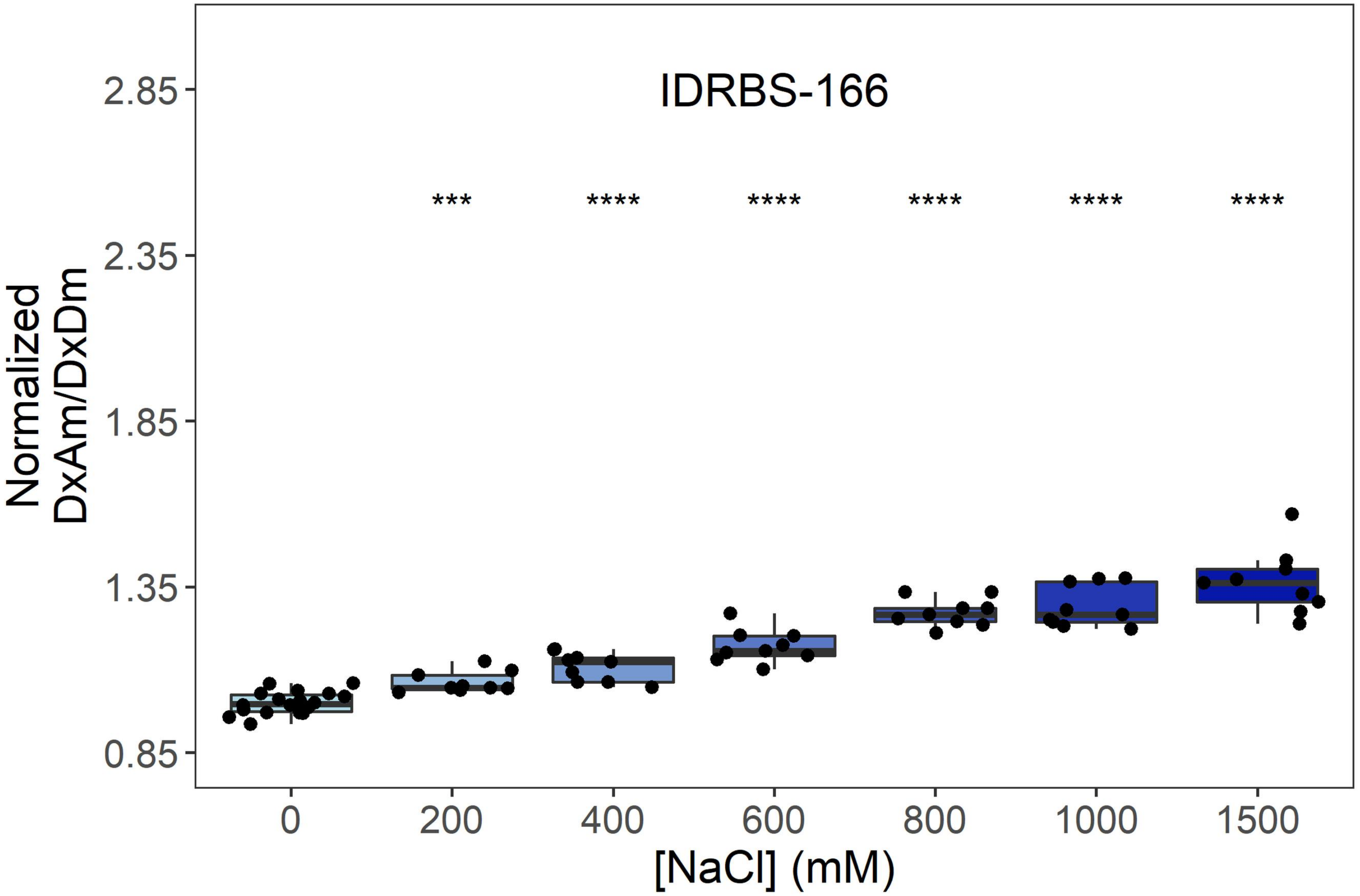

IDRBS-166
***
****
****
****
****
****
Normalized DxAm/DxDm
2.85
2.35
1.85
1.35
0.85
0
200
400
600
800
1000
1500
[NaCl] (mM)


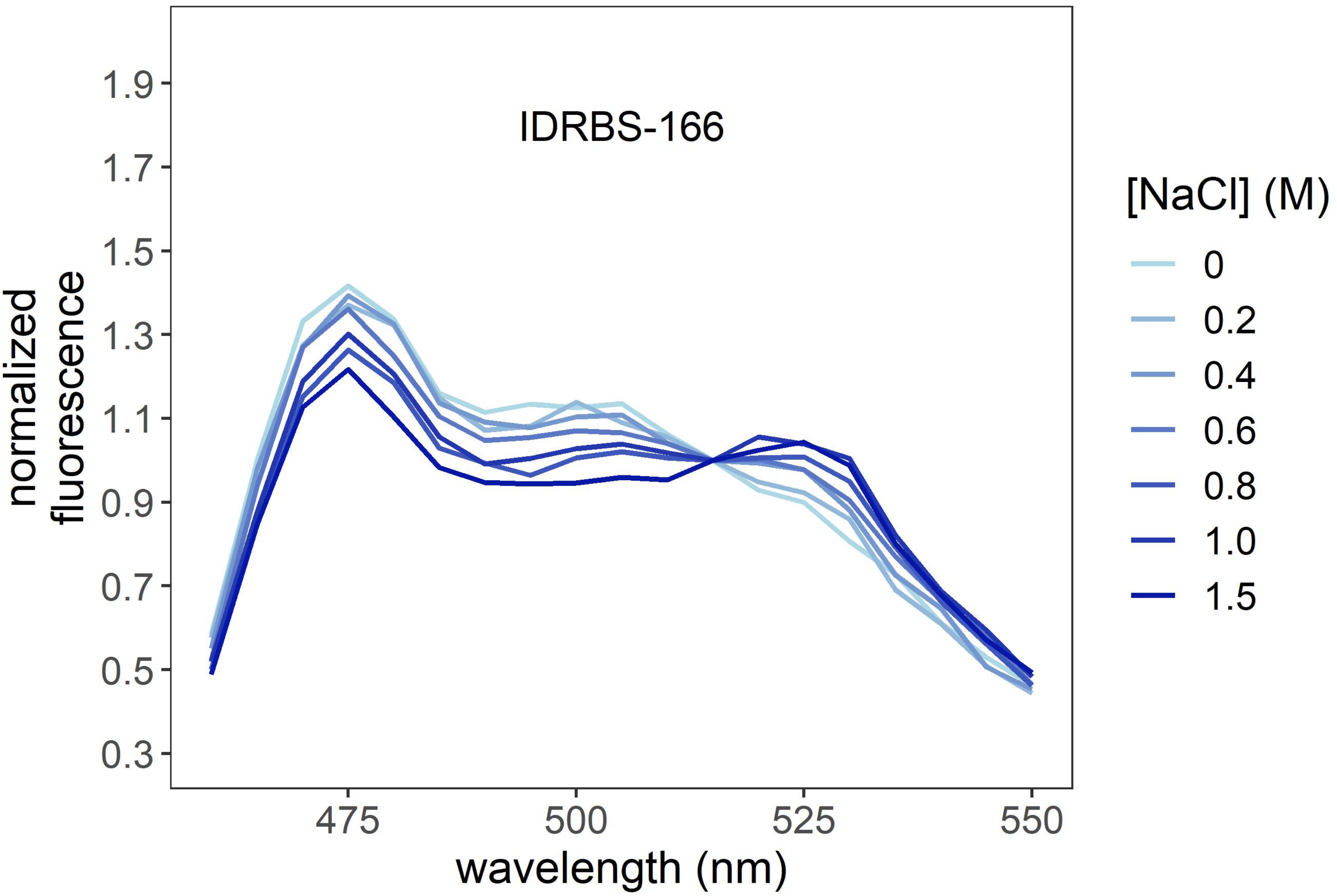

IDRBS-166
normalized fluorescence
1.9
1.7
1.5
1.3
1.1
0.9
0.7
0.5
0.3
475
500
525
550
wavelength (nm)
[NaCl] (M)
0
0.2
0.4
0.6
0.8
1.0
1.5

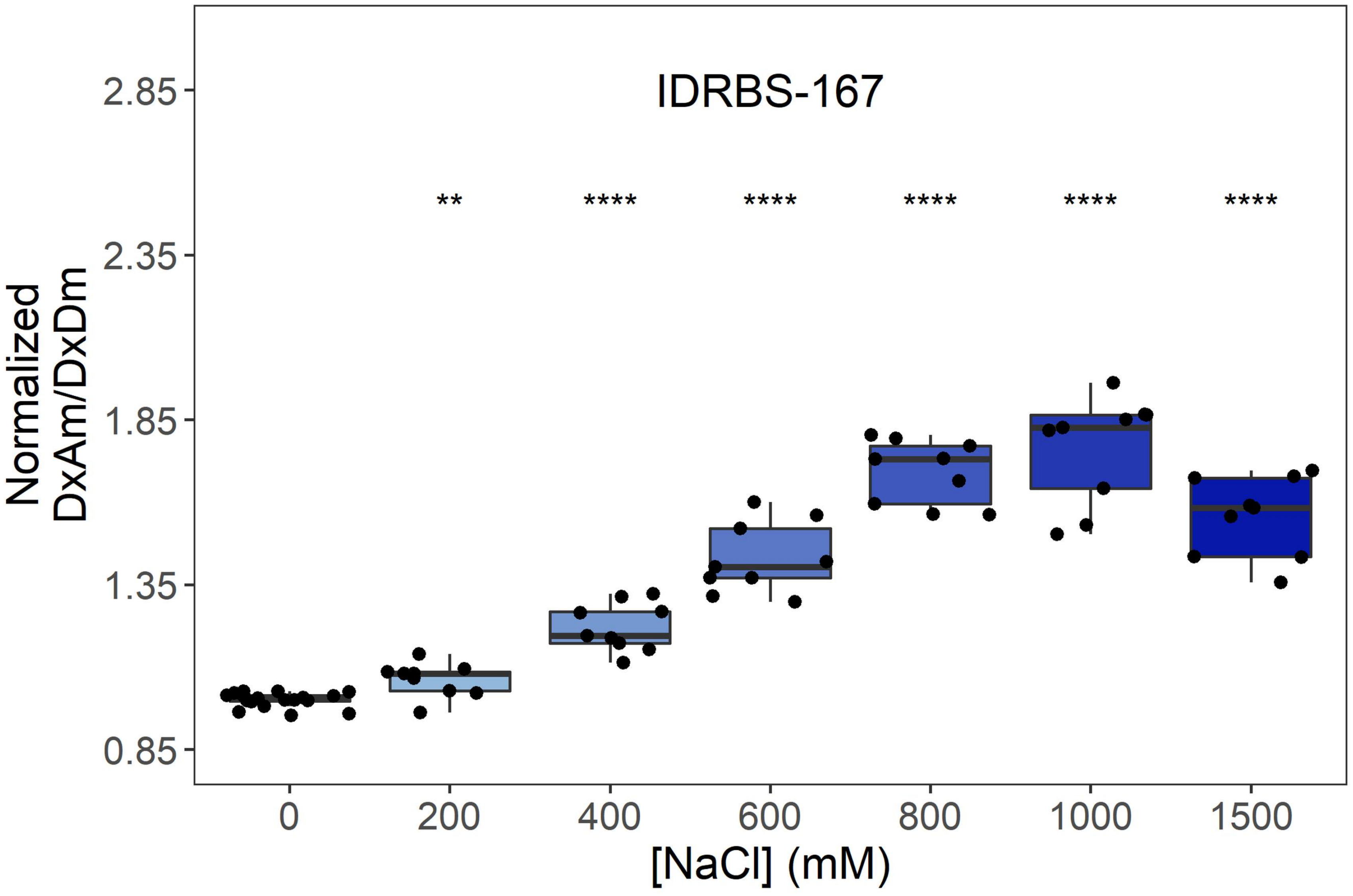
IDRBS-167
**
****
****
****
****
****
Normalized DxAm/DxDm
2.85
2.35
1.85
1.35
0.85
0
200
400
600
800
1000
1500
[NaCl] (mM)

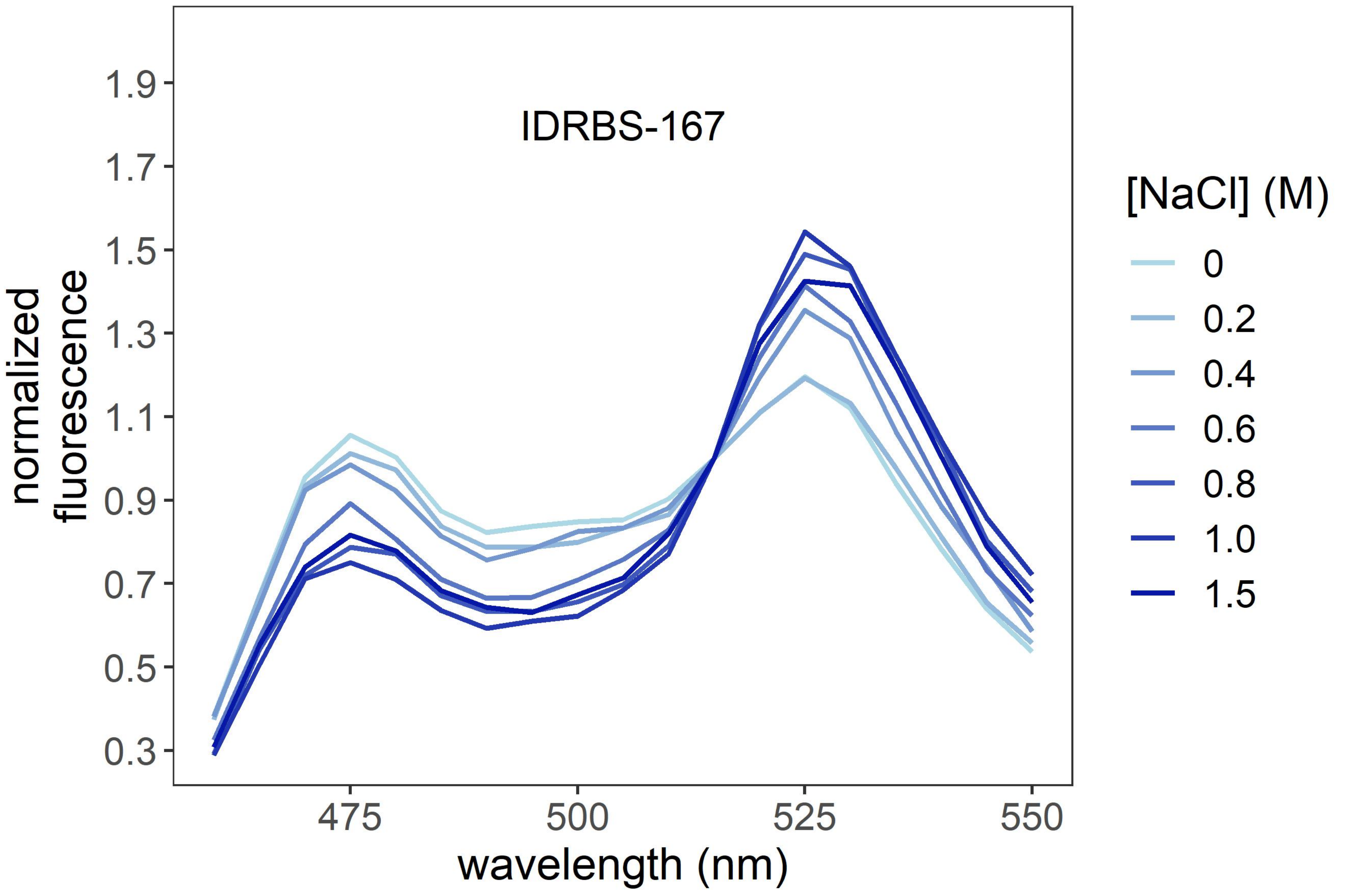
IDRBS-167
normalized fluorescence
1.9
1.7
1.5
1.3
1.1
0.9
0.7
0.5
0.3
475
500
525
550
wavelength (nm)
[NaCl] (M)
0
0.2
0.4
0.6
0.8
1.0
1.5

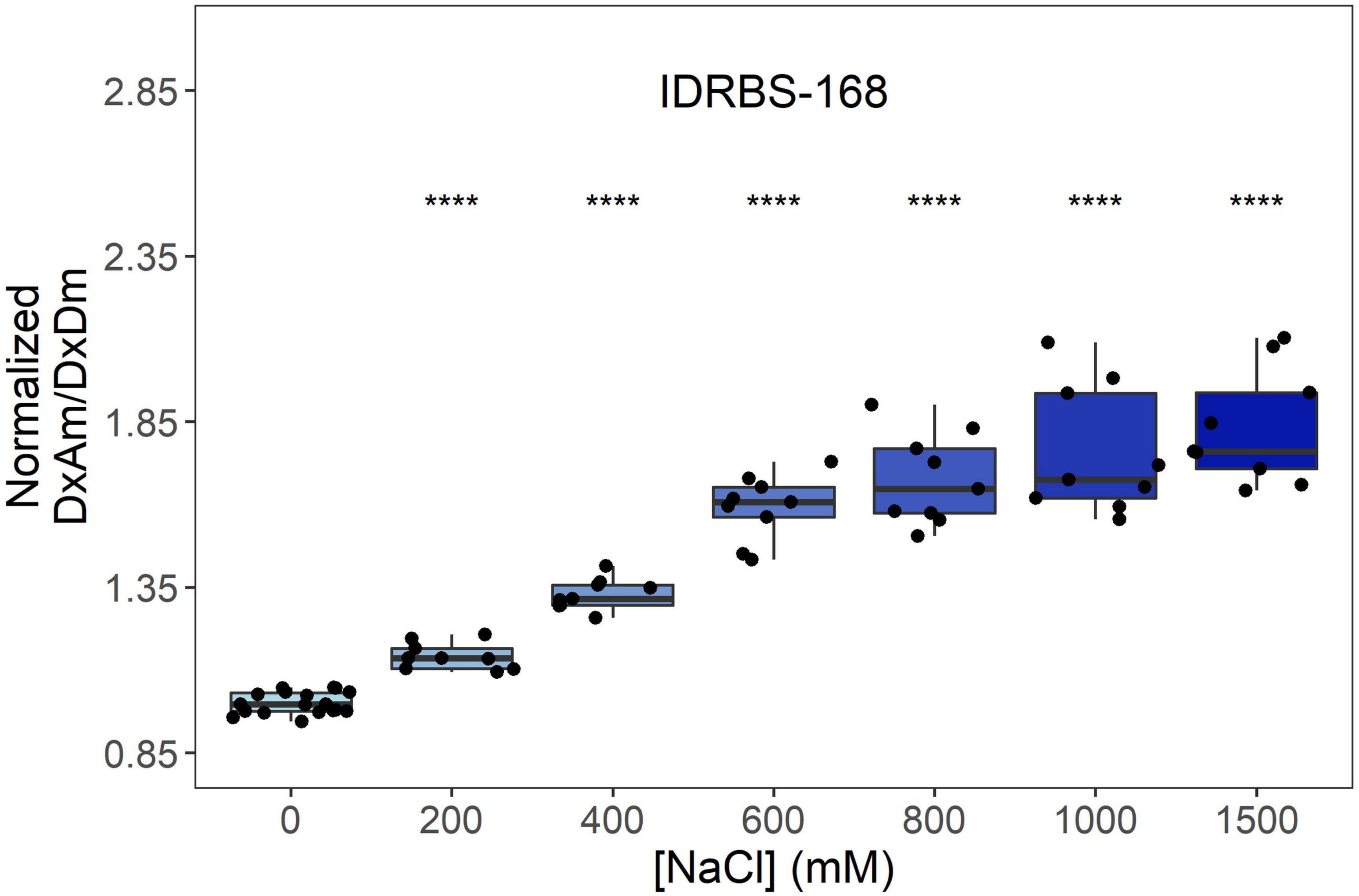
IDRBS-168
****
Normalized DxAm/DxDm
2.85
2.35
1.85
1.35
0.85
0
200
400
600
800
1000
1500
[NaCl] (mM)

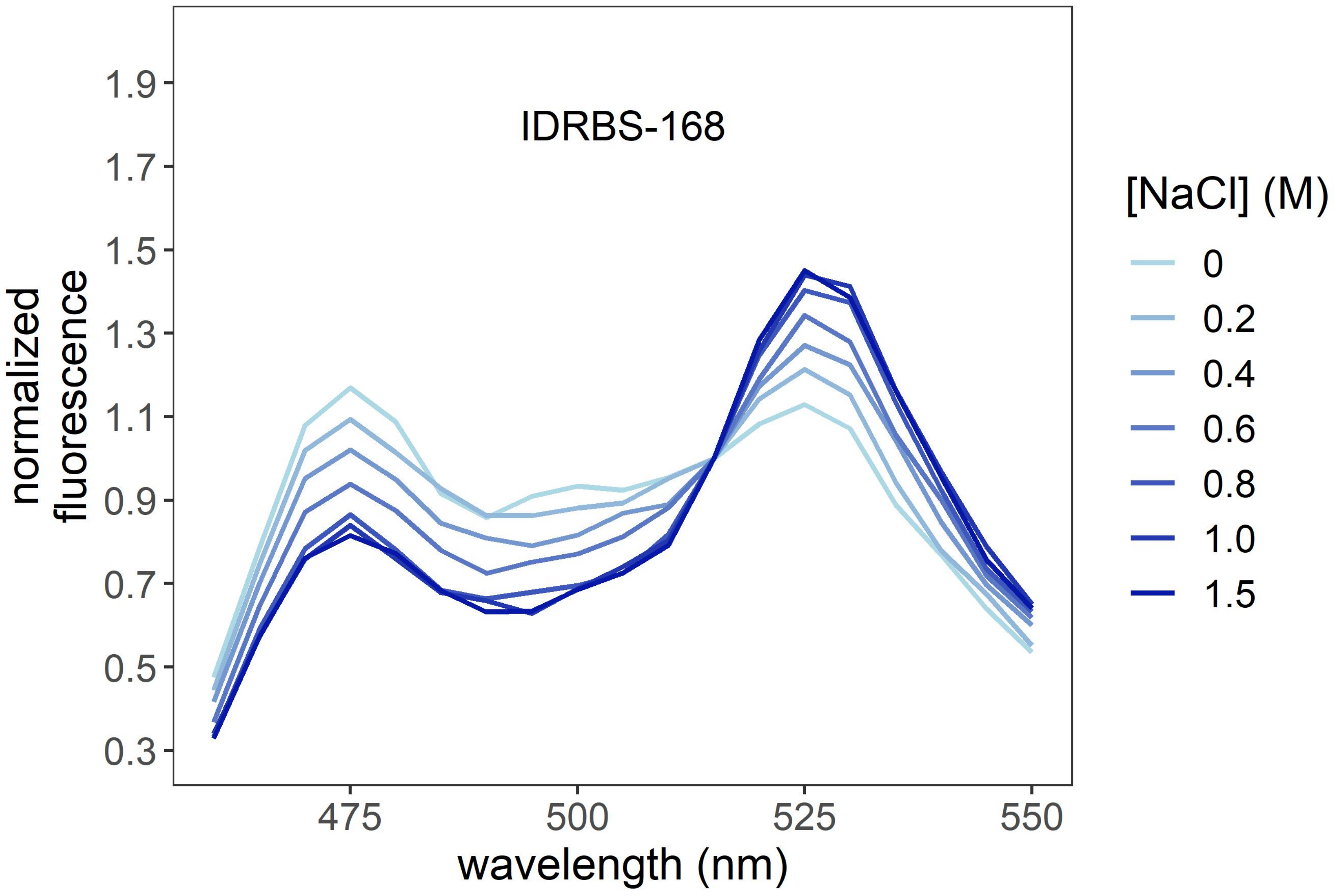
IDRBS-168
normalized fluorescence
1.9
1.7
1.5
1.3
1.1
0.9
0.7
0.5
0.3
475
500
525
550
wavelength (nm)
[NaCl] (M)
0
0.2
0.4
0.6
0.8
1.0
1.5

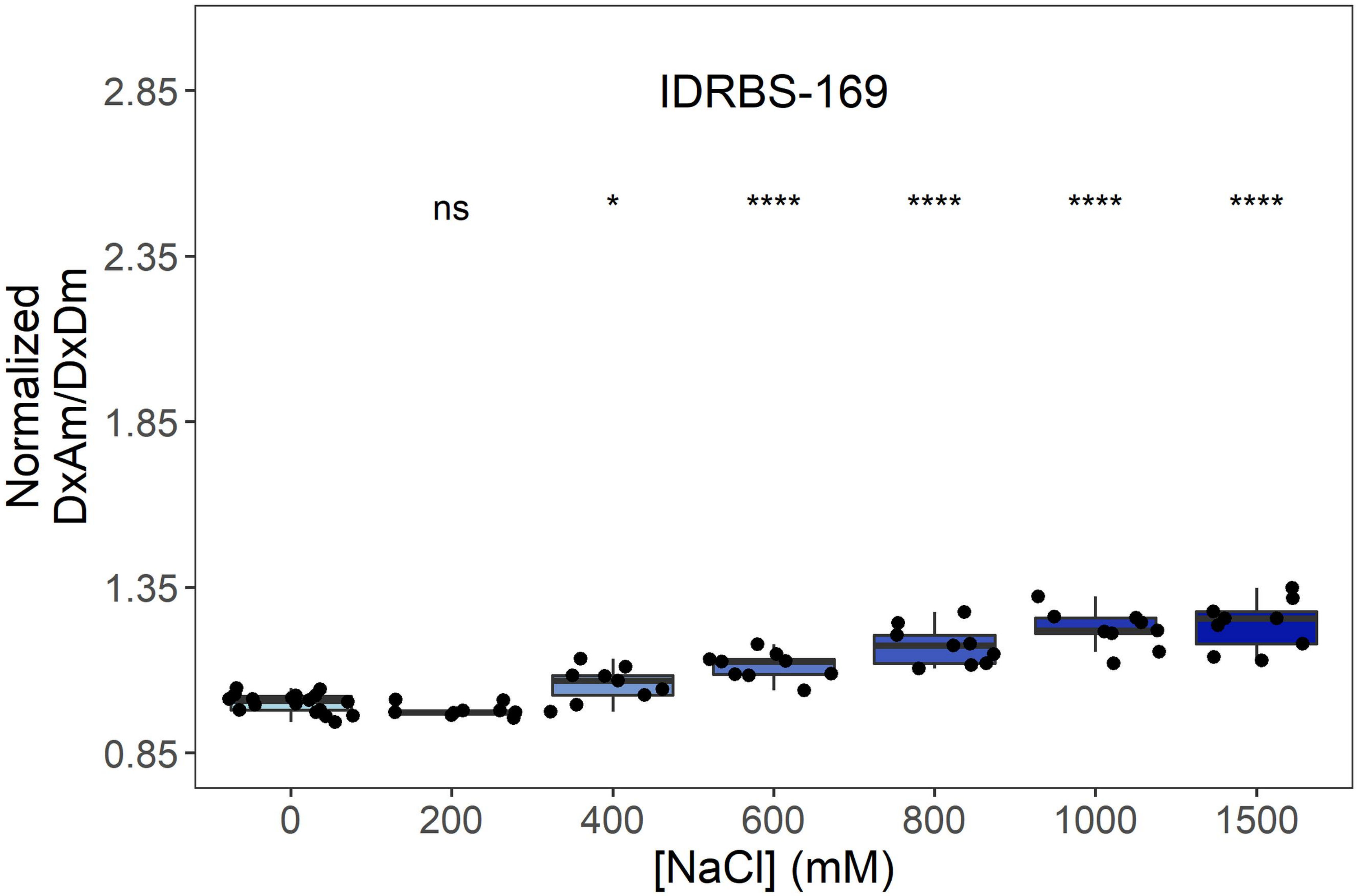
IDRBS-169
ns
*
****
****
****
****
Normalized DxAm/DxDm
2.85
2.35
1.85
1.35
0.85
0
200
400
600
800
1000
1500
[NaCl] (mM)

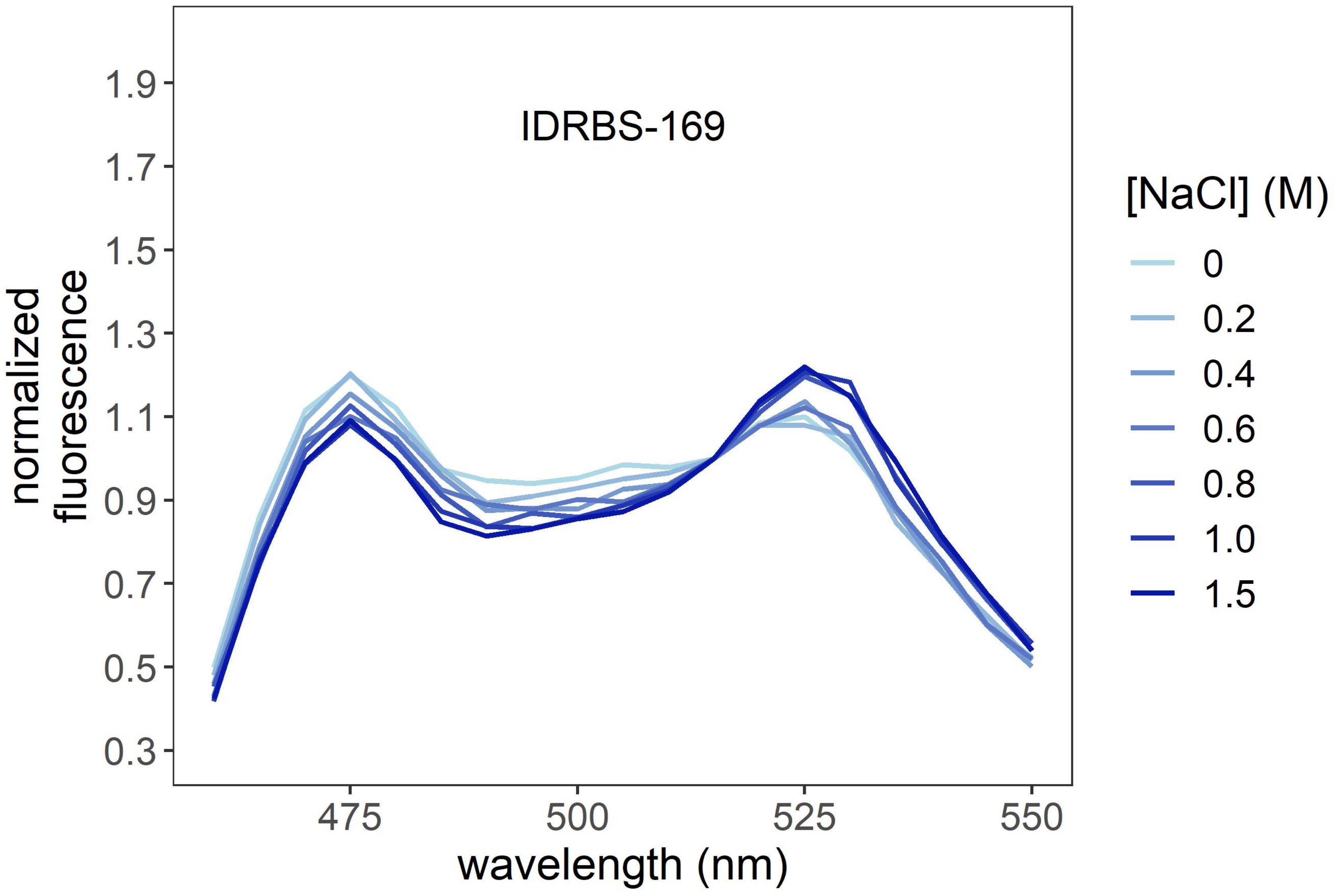
IDRBS-169
normalized fluorescence
1.9
1.7
1.5
1.3
1.1
0.9
0.7
0.5
0.3
475
500
525
550
wavelength (nm)
[NaCl] (M)
0
0.2
0.4
0.6
0.8
1.0
1.5

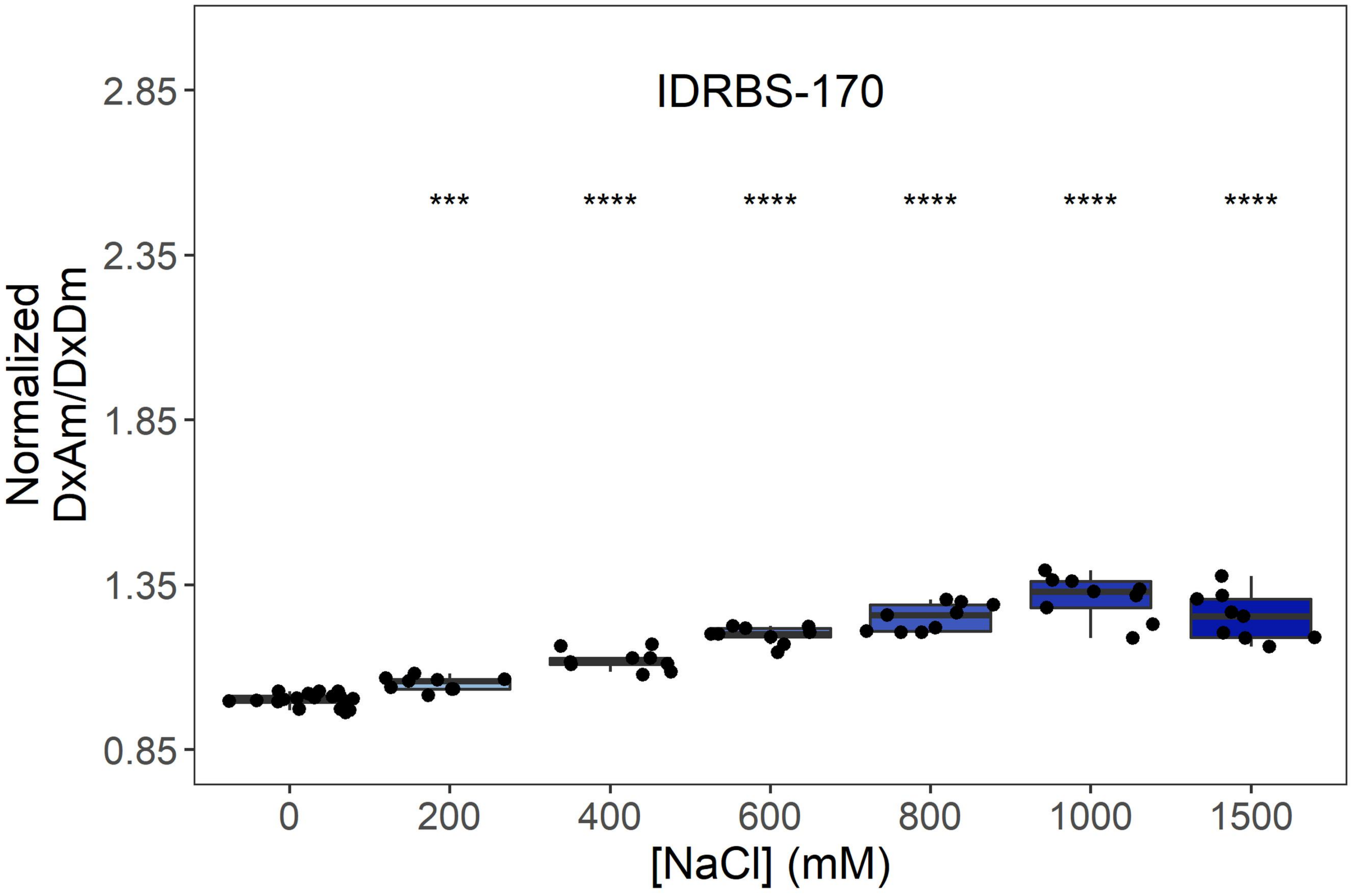

IDRBS-170
Normalized DxAm/DxDm
2.85
2.35
1.85
1.35
0.85
***
****
0
200
400
600
800
1000
1500
[NaCl] (mM)


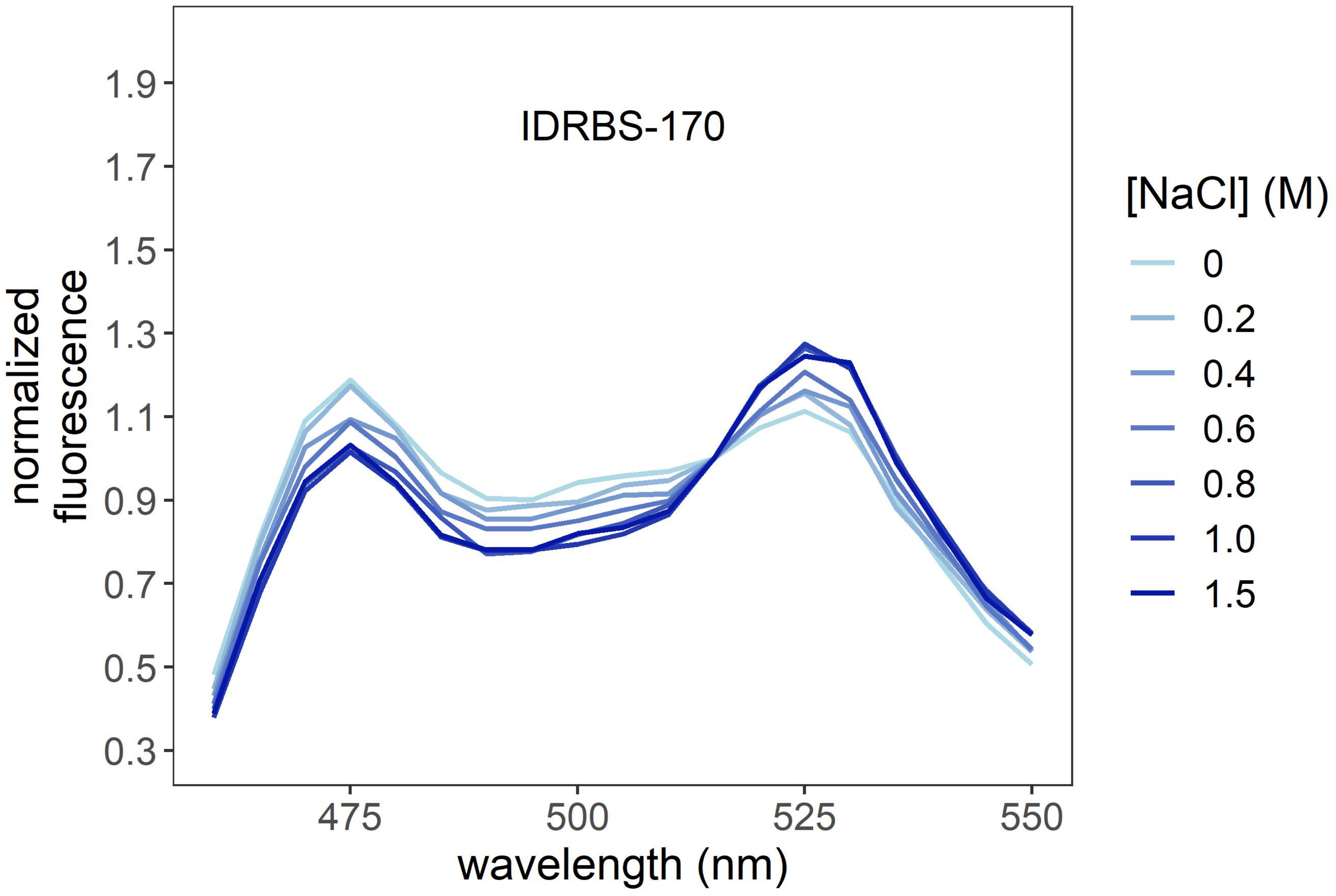

IDRBS-170
normalized fluorescence
1.9
1.7
1.5
1.3
1.1
0.9
0.7
0.5
0.3
475
500
525
550
wavelength (nm)
[NaCl] (M)
0
0.2
0.4
0.6
0.8
1.0
1.5

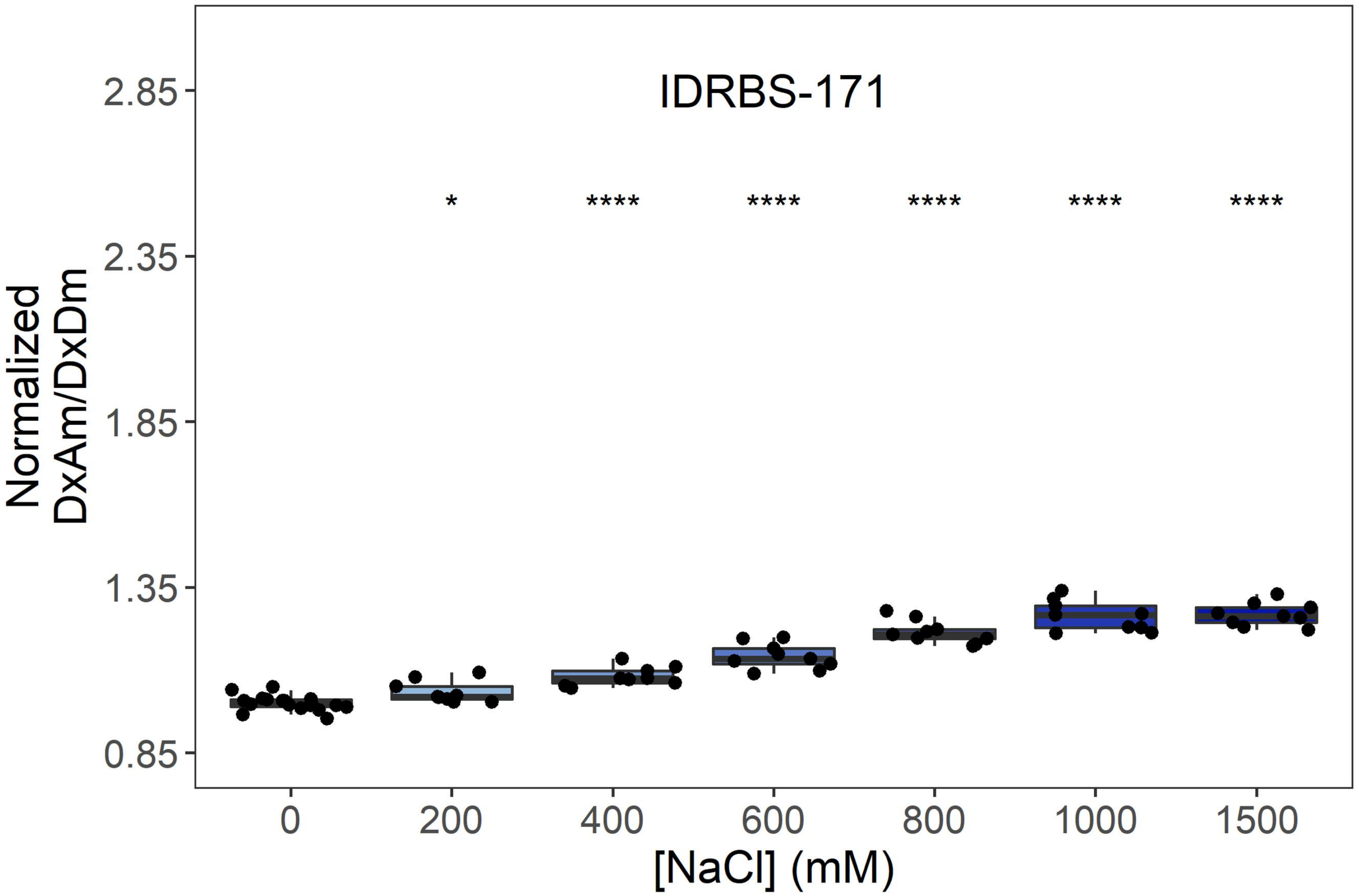
IDRBS-171
Normalized DxAm/DxDm
2.85
2.35
1.85
1.35
0.85
*
****
****
****
****
****
0
200
400
600
800
1000
1500
[NaCl] (mM)

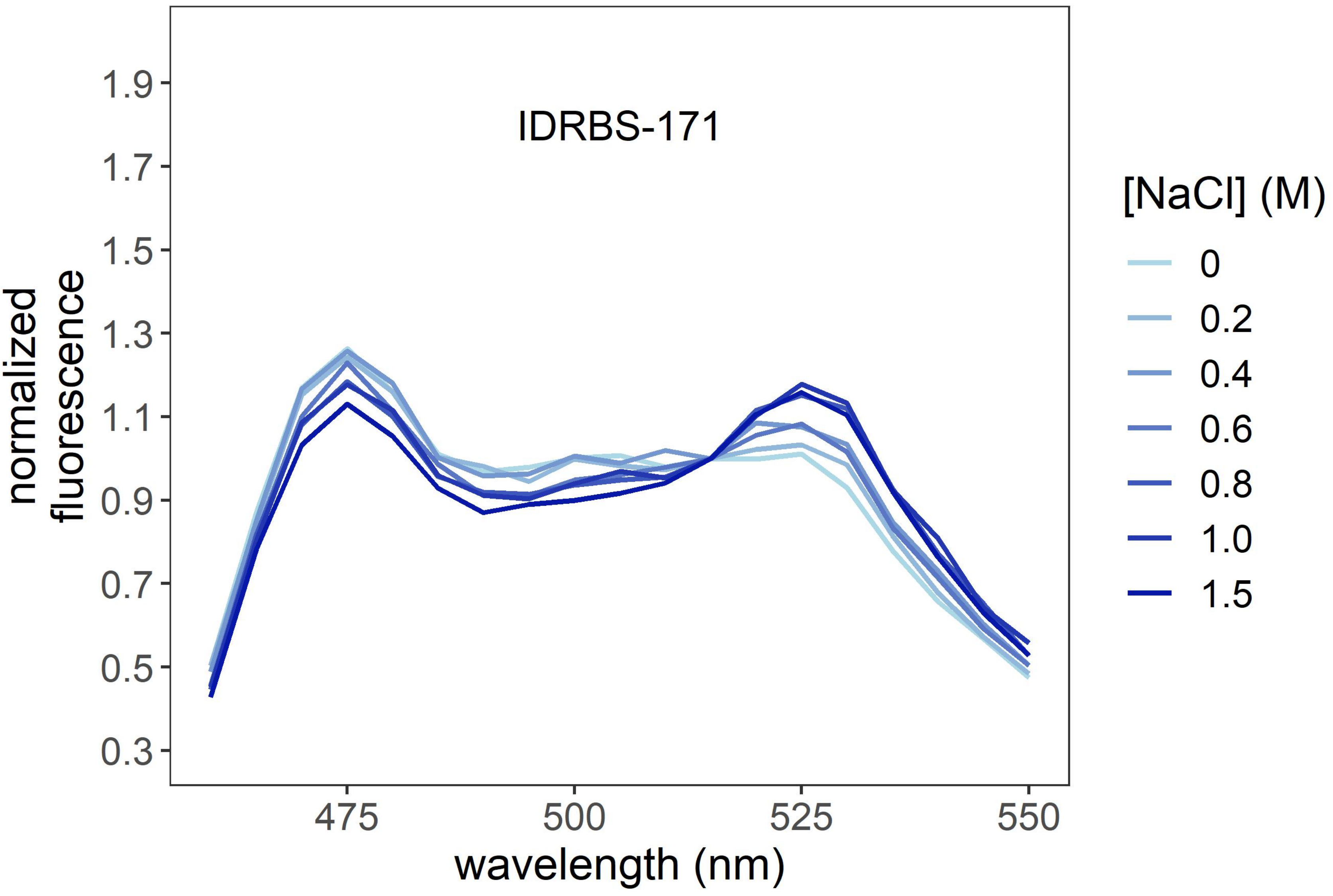
IDRBS-171
normalized fluorescence
1.9
1.7
1.5
1.3
1.1
0.9
0.7
0.5
0.3
475
500
525
550
wavelength (nm)
[NaCl] (M)
0
0.2
0.4
0.6
0.8
1.0
1.5

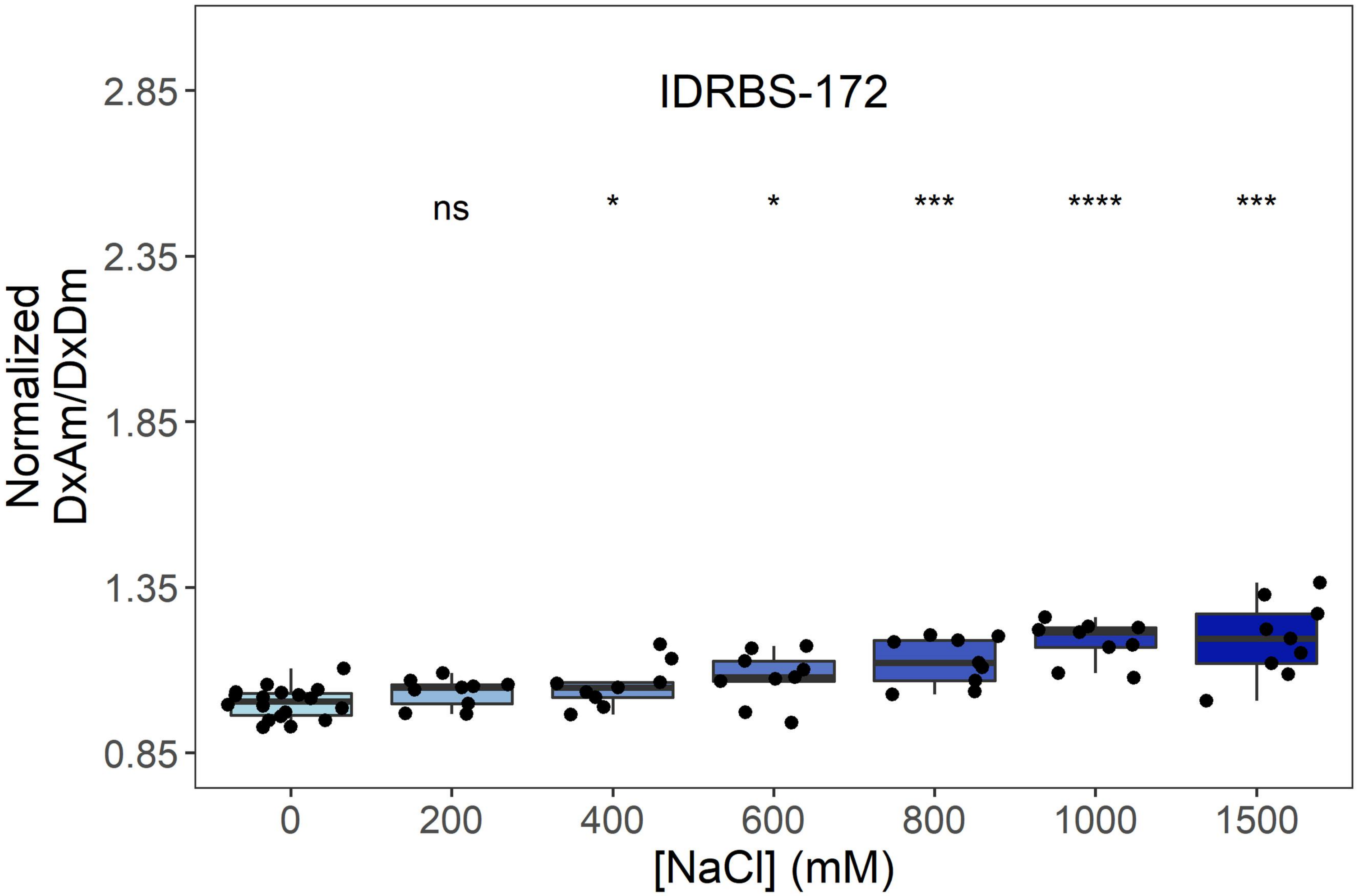
IDRBS-172
ns
*
*
***
****
***
Normalized
DxAm/DxDm
2.85
2.35
1.85
1.35
0.85
0
200
400
600
800
1000
1500
[NaCl] (mM)

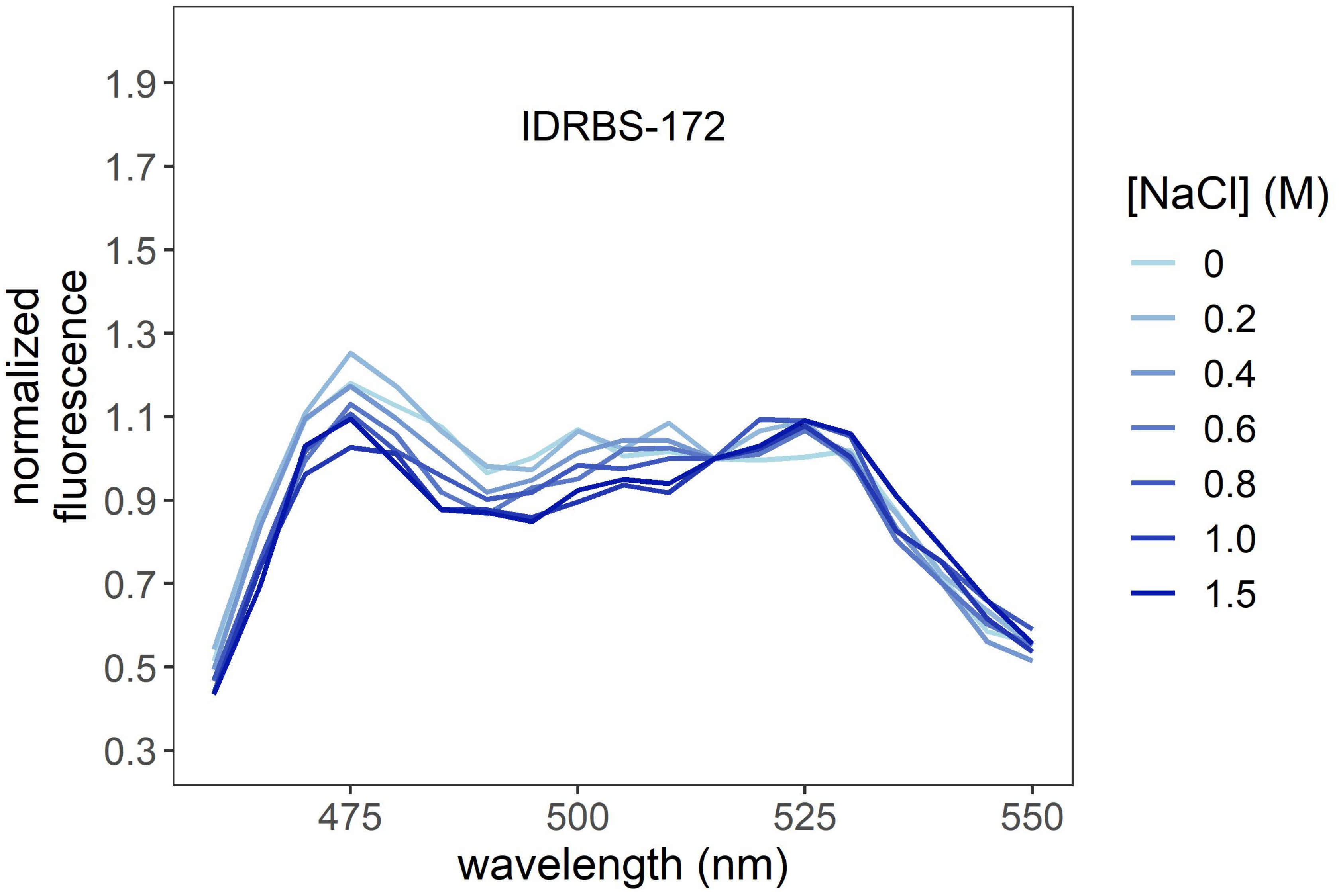
IDRBS-172
normalized
fluorescence
1.9
1.7
1.5
1.3
1.1
0.9
0.7
0.5
0.3
475
500
525
550
wavelength (nm)
[NaCl] (M)
0
0.2
0.4
0.6
0.8
1.0
1.5

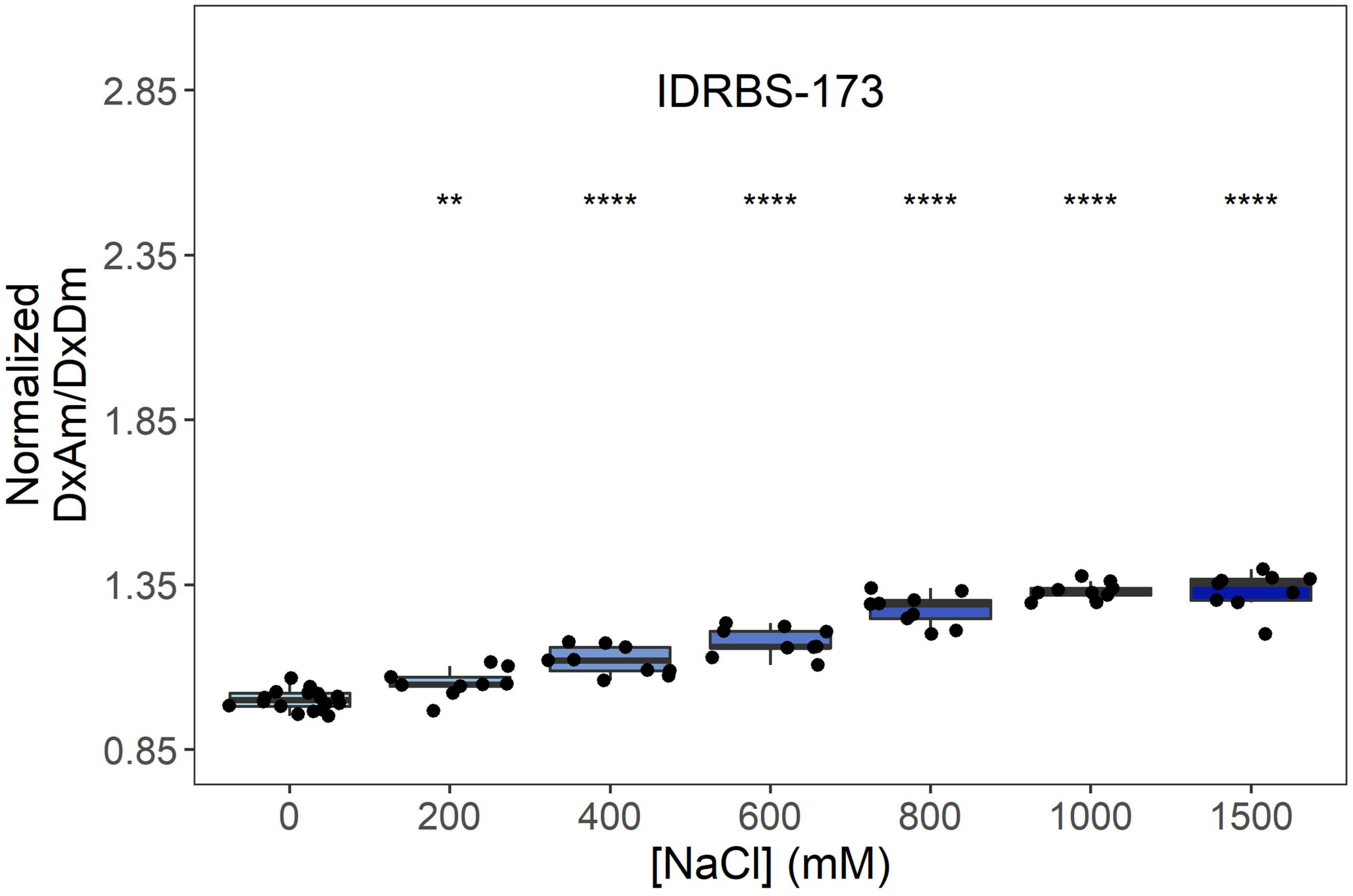
IDRBS-173
Normalized
DxAm/DxDm
2.85
2.35
1.85
1.35
0.85
**
****
****
****
****
****
0
200
400
600
800
1000
1500
[NaCl] (mM)

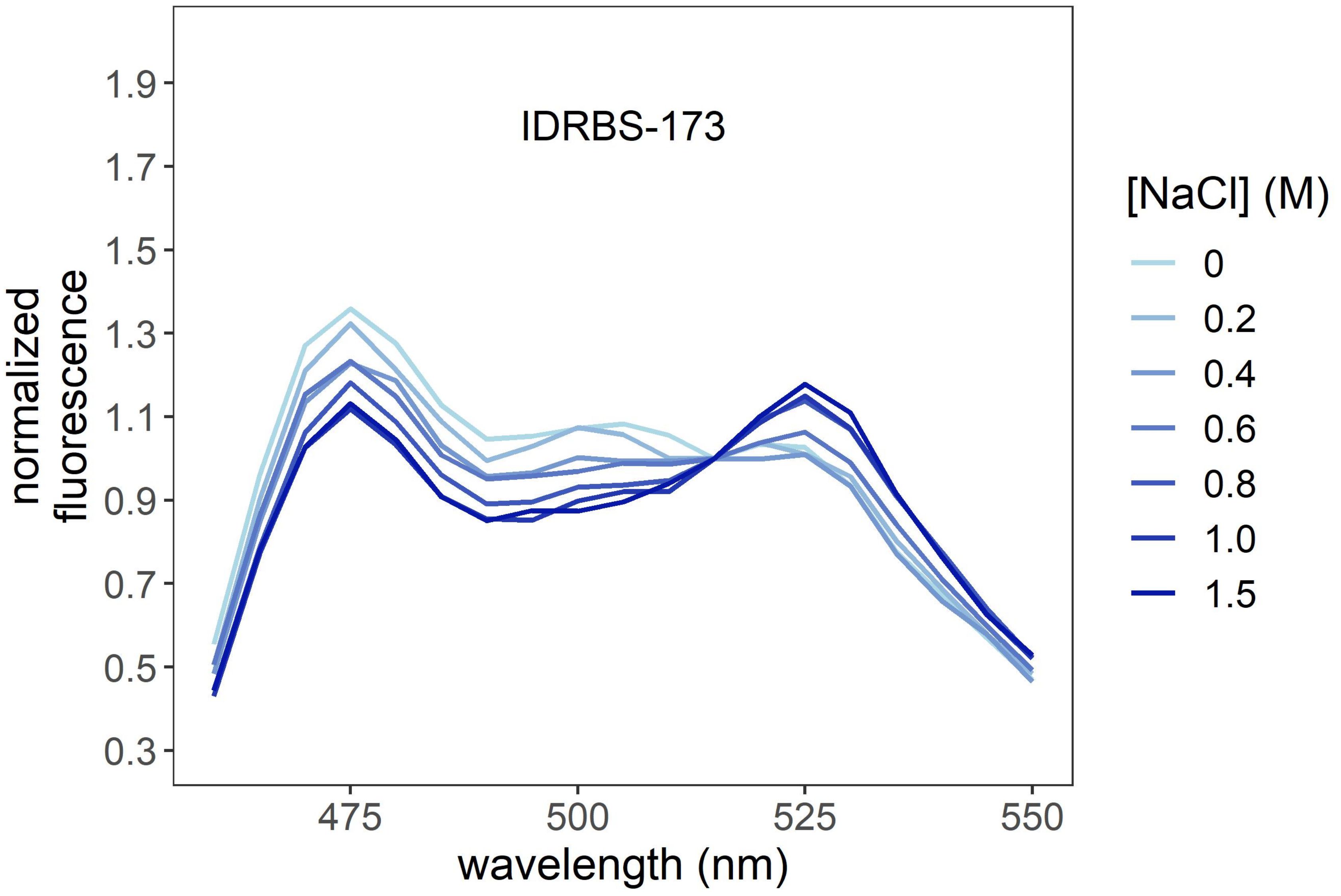
IDRBS-173
normalized
fluorescence
1.9
1.7
1.5
1.3
1.1
0.9
0.7
0.5
0.3
475
500
525
550
wavelength (nm)
[NaCl] (M)
0
0.2
0.4
0.6
0.8
1.0
1.5

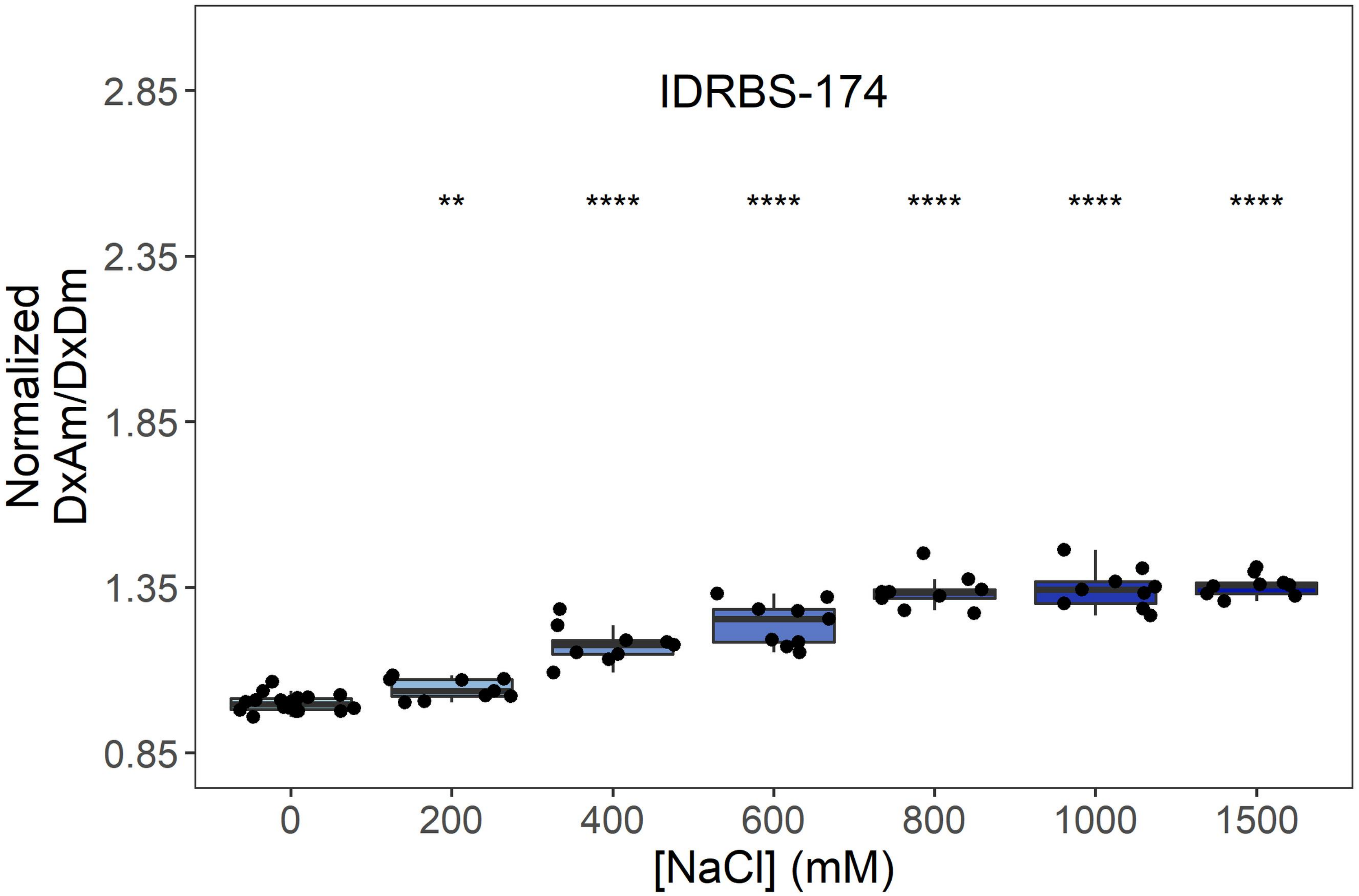
IDRBS-174
**
****
****
****
****
****
Normalized DxAm/DxDm
2.85
2.35
1.85
1.35
0.85
0
200
400
600
800
1000
1500
[NaCl] (mM)

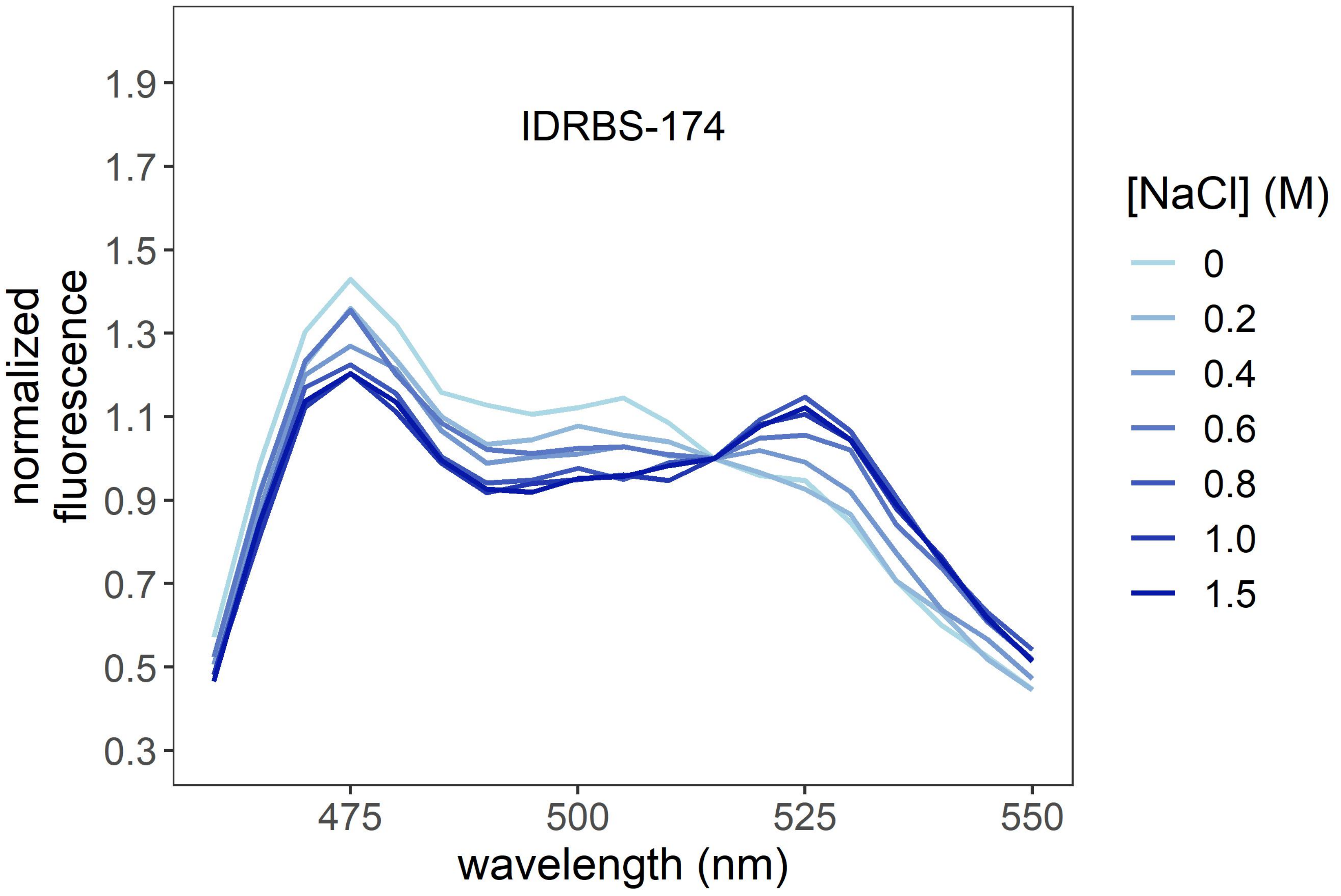
IDRBS-174
normalized fluorescence
1.9
1.7
1.5
1.3
1.1
0.9
0.7
0.5
0.3
475
500
525
550
wavelength (nm)
[NaCl] (M)
0
0.2
0.4
0.6
0.8
1.0
1.5

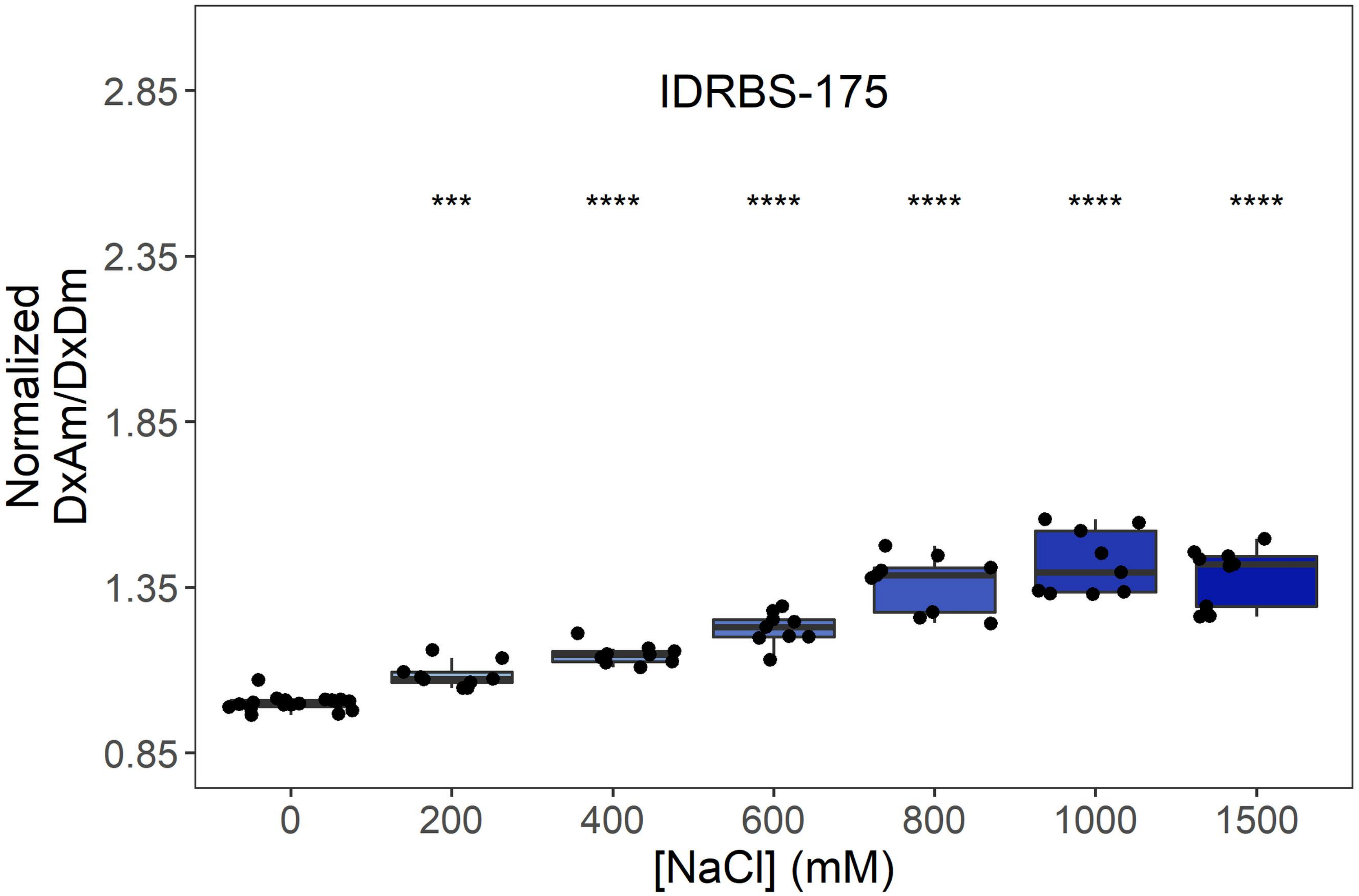

IDRBS-175
***
****
****
****
****
****
Normalized DxAm/DxDm
2.85
2.35
1.85
1.35
0.85
0
200
400
600
800
1000
1500
[NaCl] (mM)


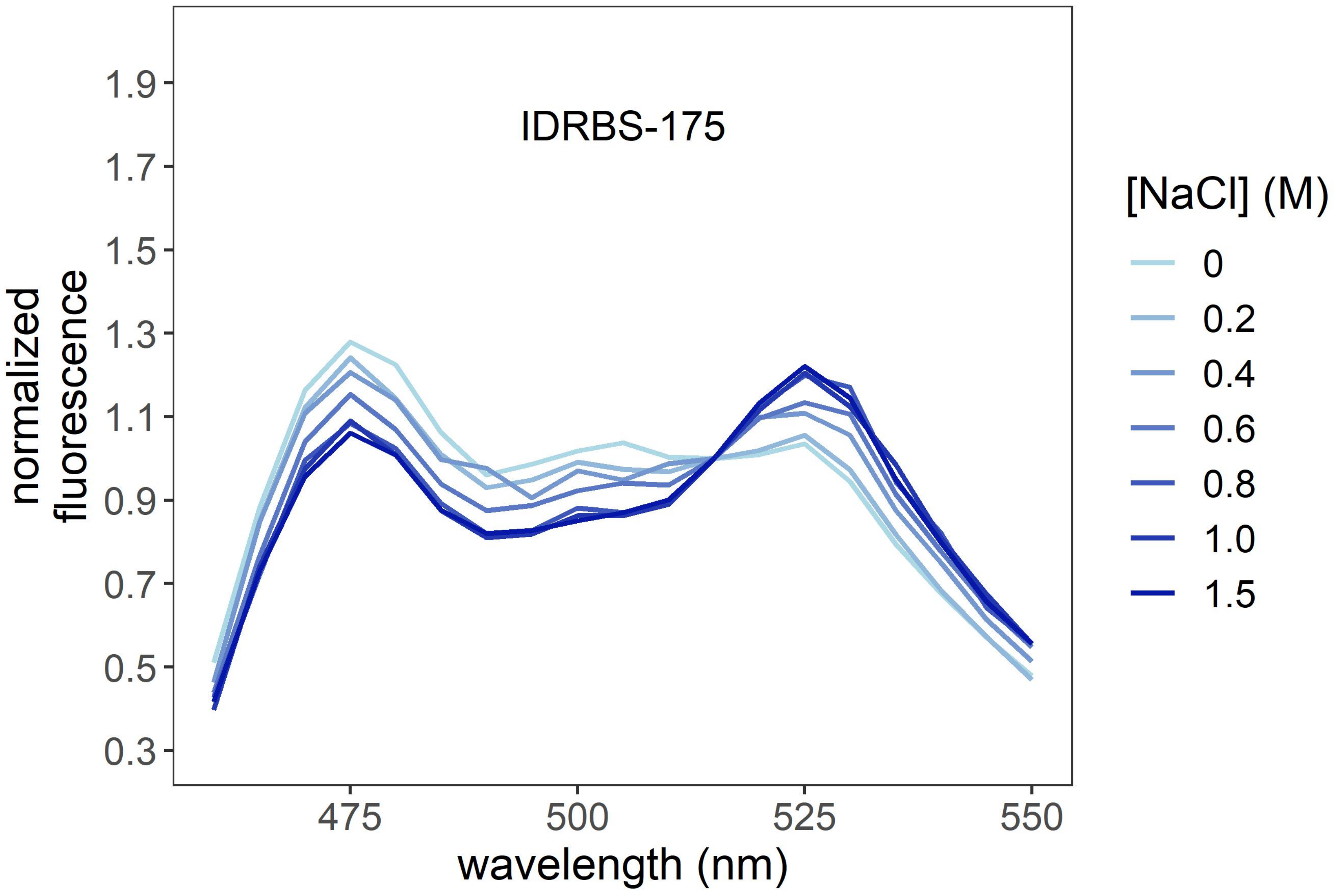

IDRBS-175
normalized fluorescence
1.9
1.7
1.5
1.3
1.1
0.9
0.7
0.5
0.3
475
500
525
550
wavelength (nm)
[NaCl] (M)
0
0.2
0.4
0.6
0.8
1.0
1.5

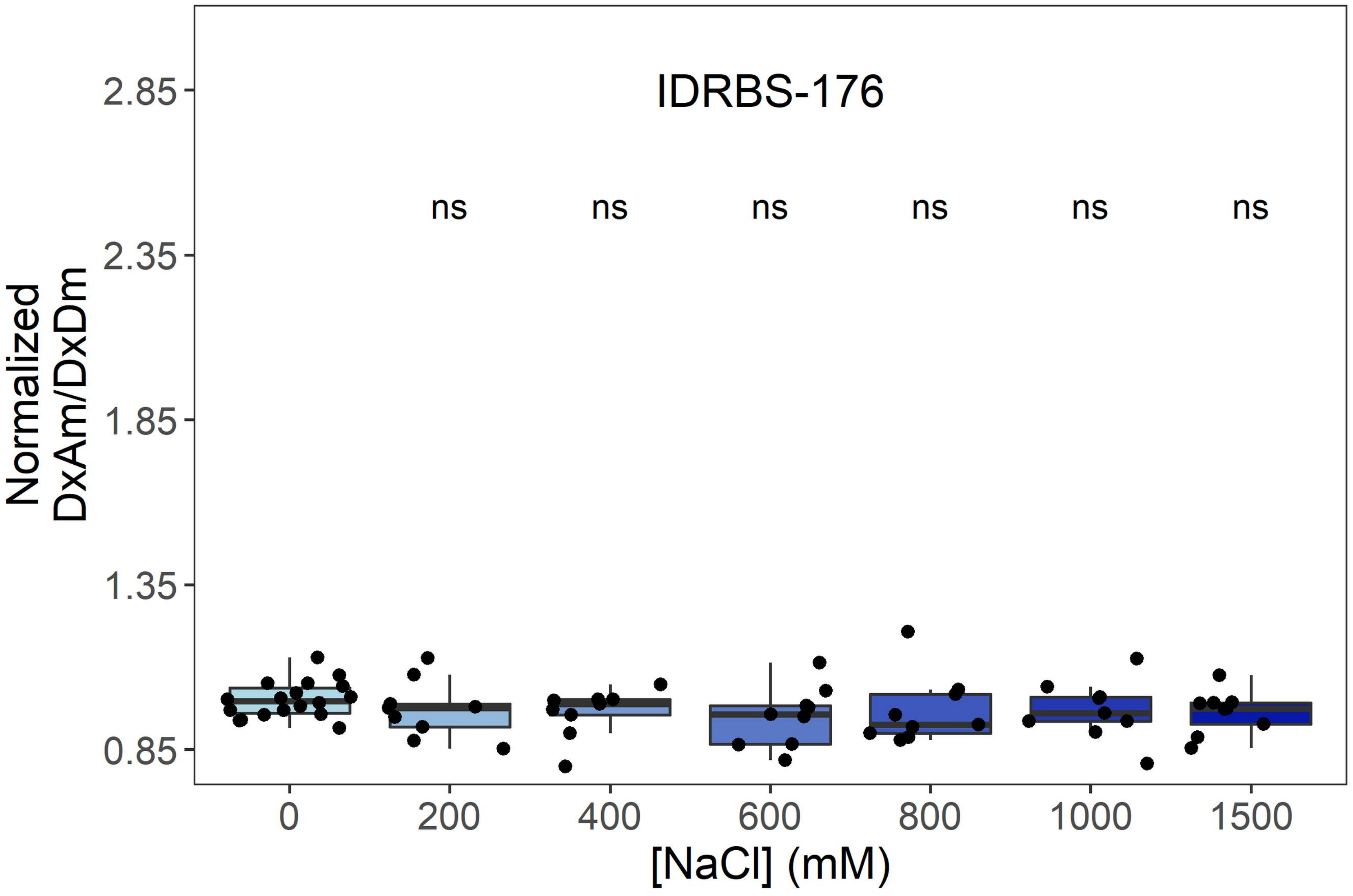

IDRBS-176
ns
ns
ns
ns
ns
ns
Normalized
DxAm/DxDm
2.85
2.35
1.85
1.35
0.85
0
200
400
600
800
1000
1500
[NaCl] (mM)


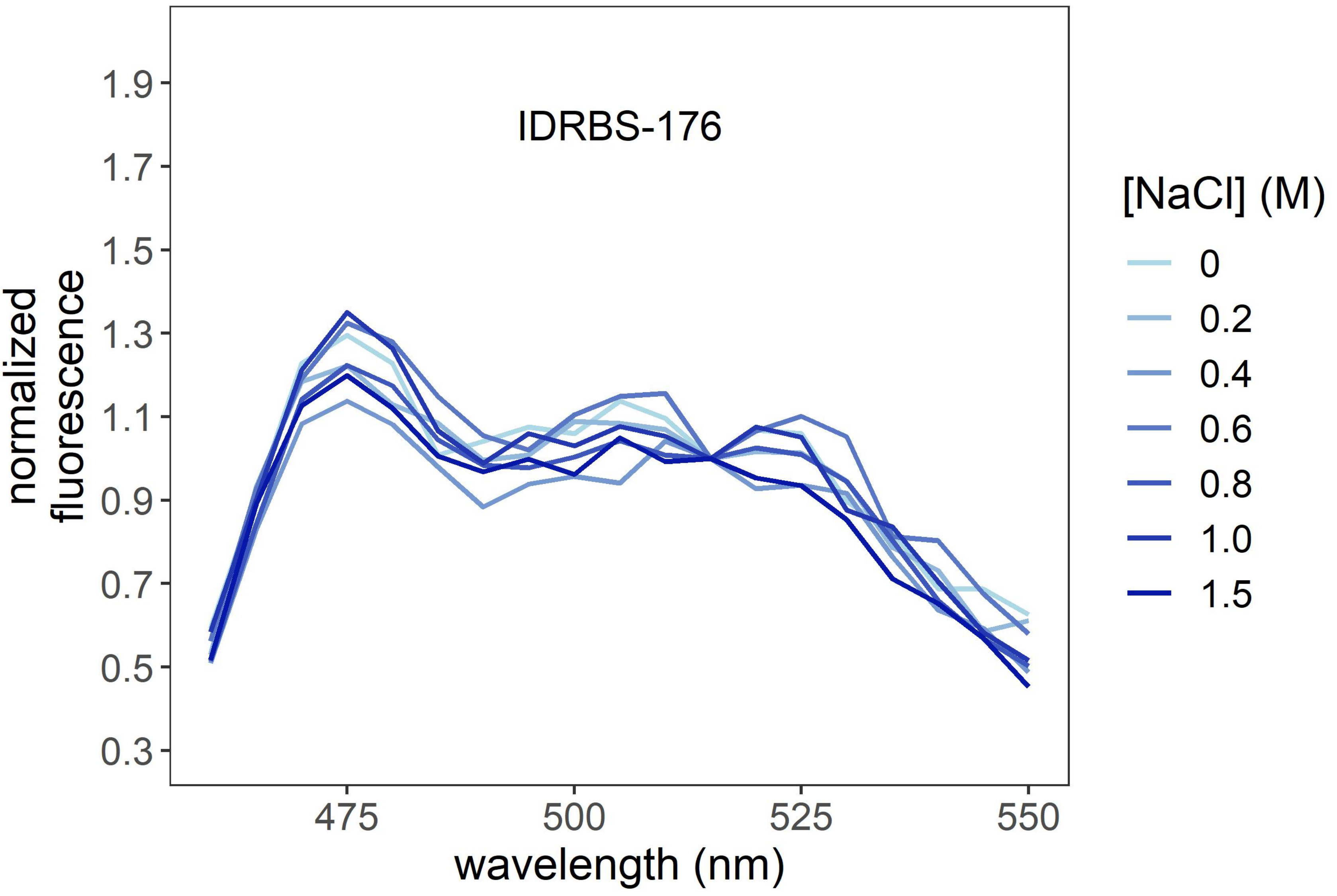

IDRBS-176
normalized
fluorescence
1.9
1.7
1.5
1.3
1.1
0.9
0.7
0.5
0.3
475
500
525
550
wavelength (nm)
[NaCl] (M)
0
0.2
0.4
0.6
0.8
1.0
1.5

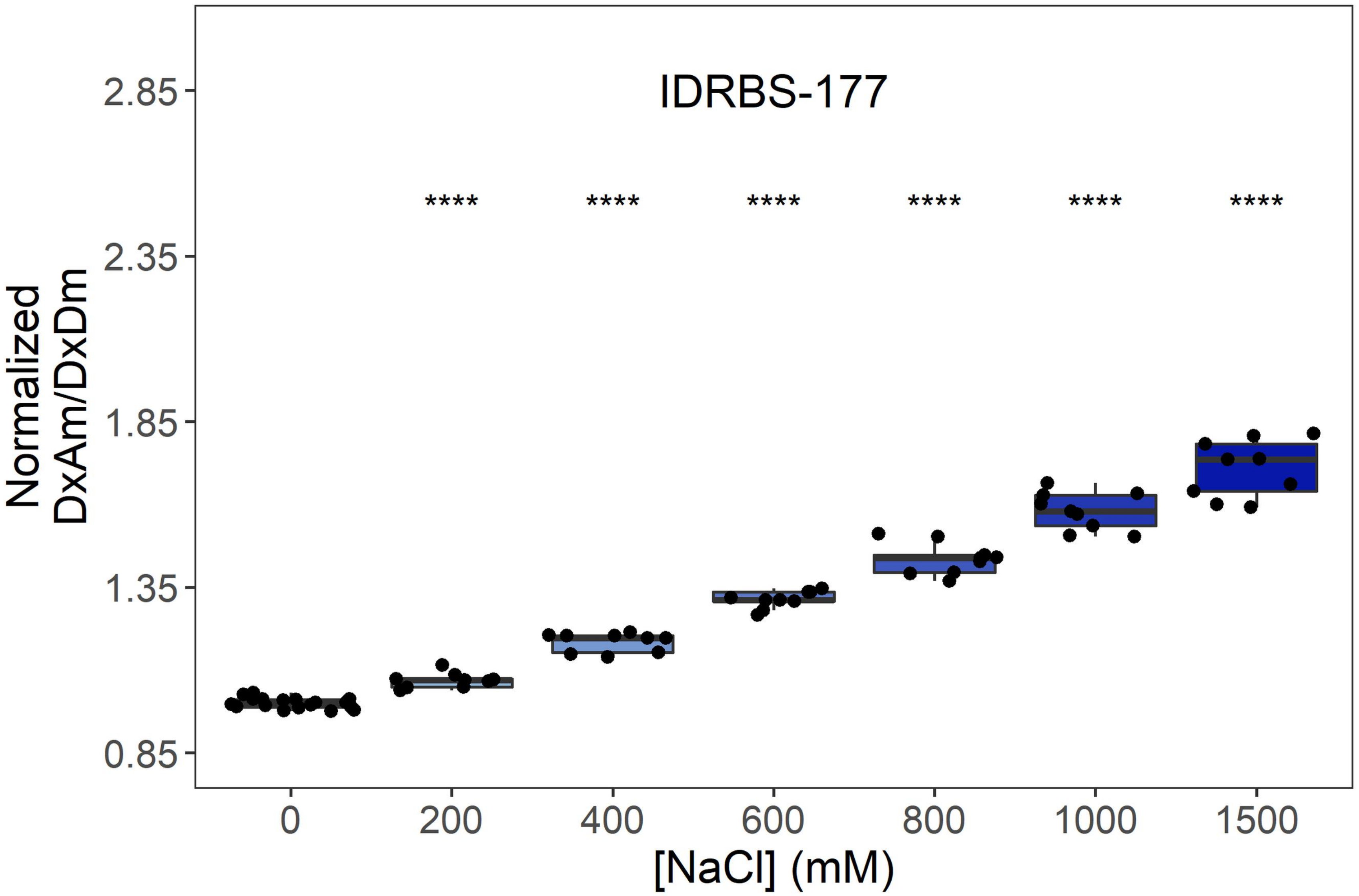

IDRBS-177
Normalized DxAm/DxDm
2.85
2.35
1.85
1.35
0.85
****
0
200
400
600
800
1000
1500
[NaCl] (mM)


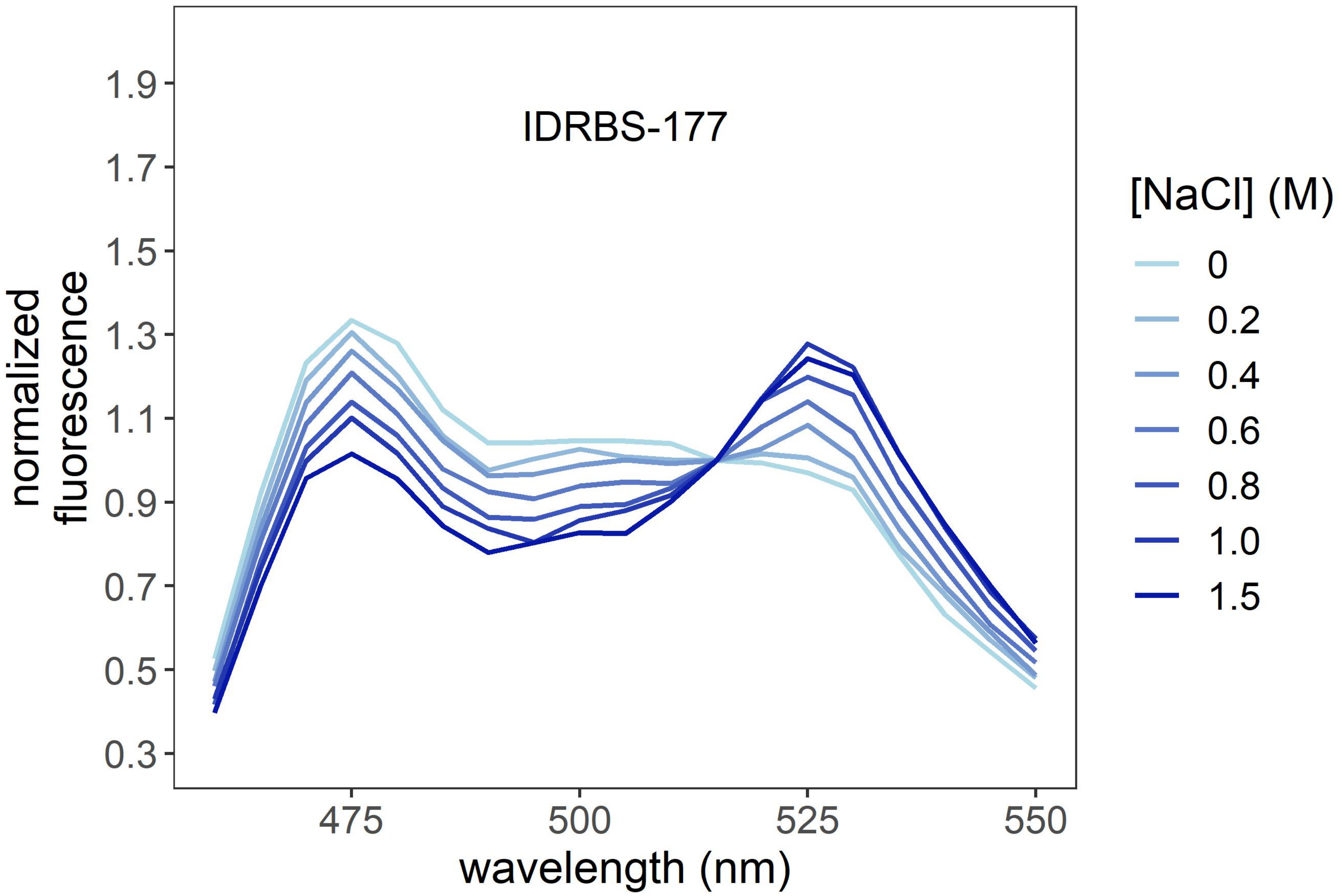

IDRBS-177
normalized fluorescence
1.9
1.7
1.5
1.3
1.1
0.9
0.7
0.5
0.3
475
500
525
550
wavelength (nm)
[NaCl] (M)
0
0.2
0.4
0.6
0.8
1.0
1.5

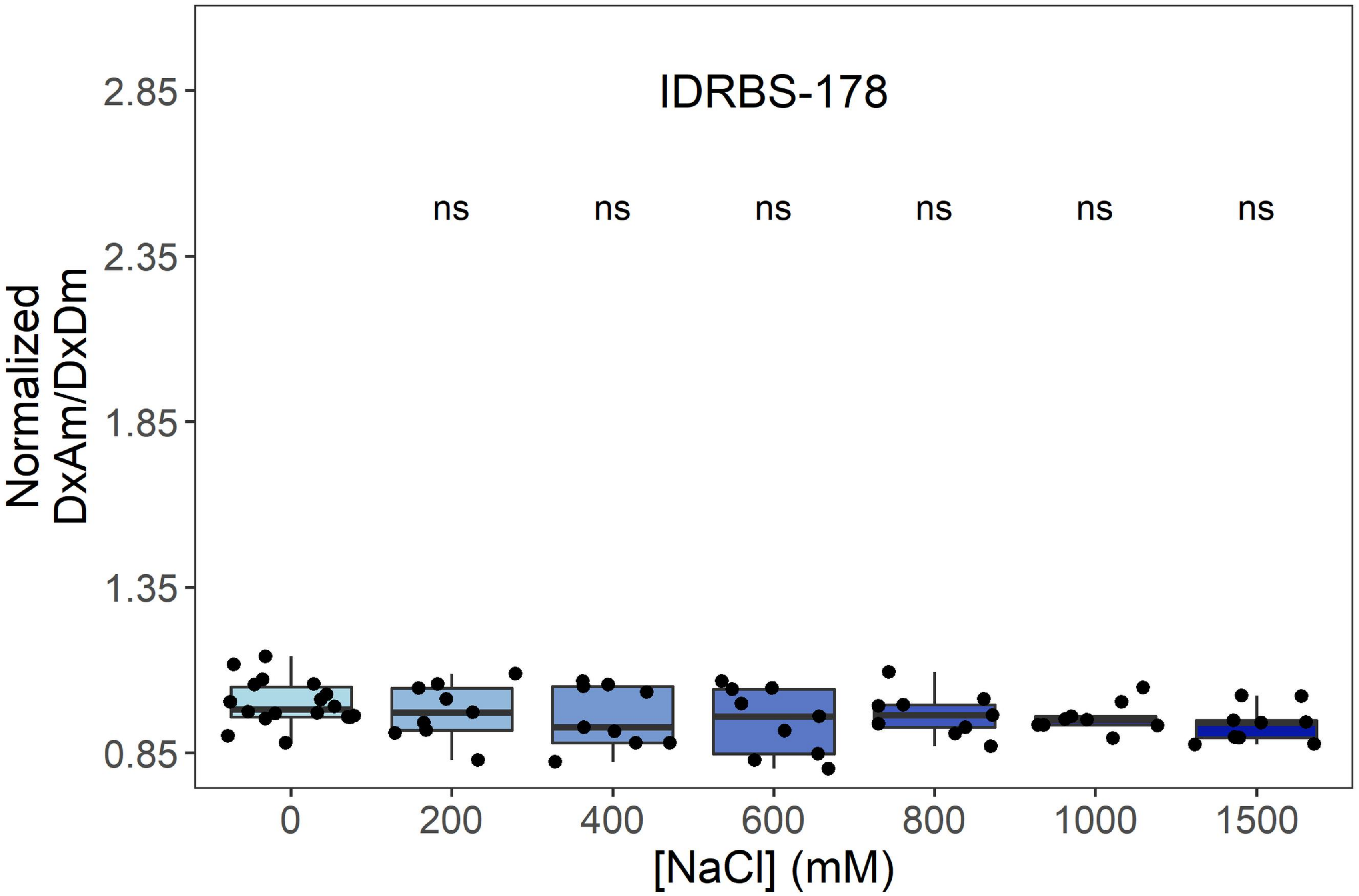
IDRBS-178
Normalized
DxAm/DxDm
2.85
2.35
1.85
1.35
0.85
ns
ns
ns
ns
ns
ns
0
200
400
600
800
1000
1500
[NaCl] (mM)

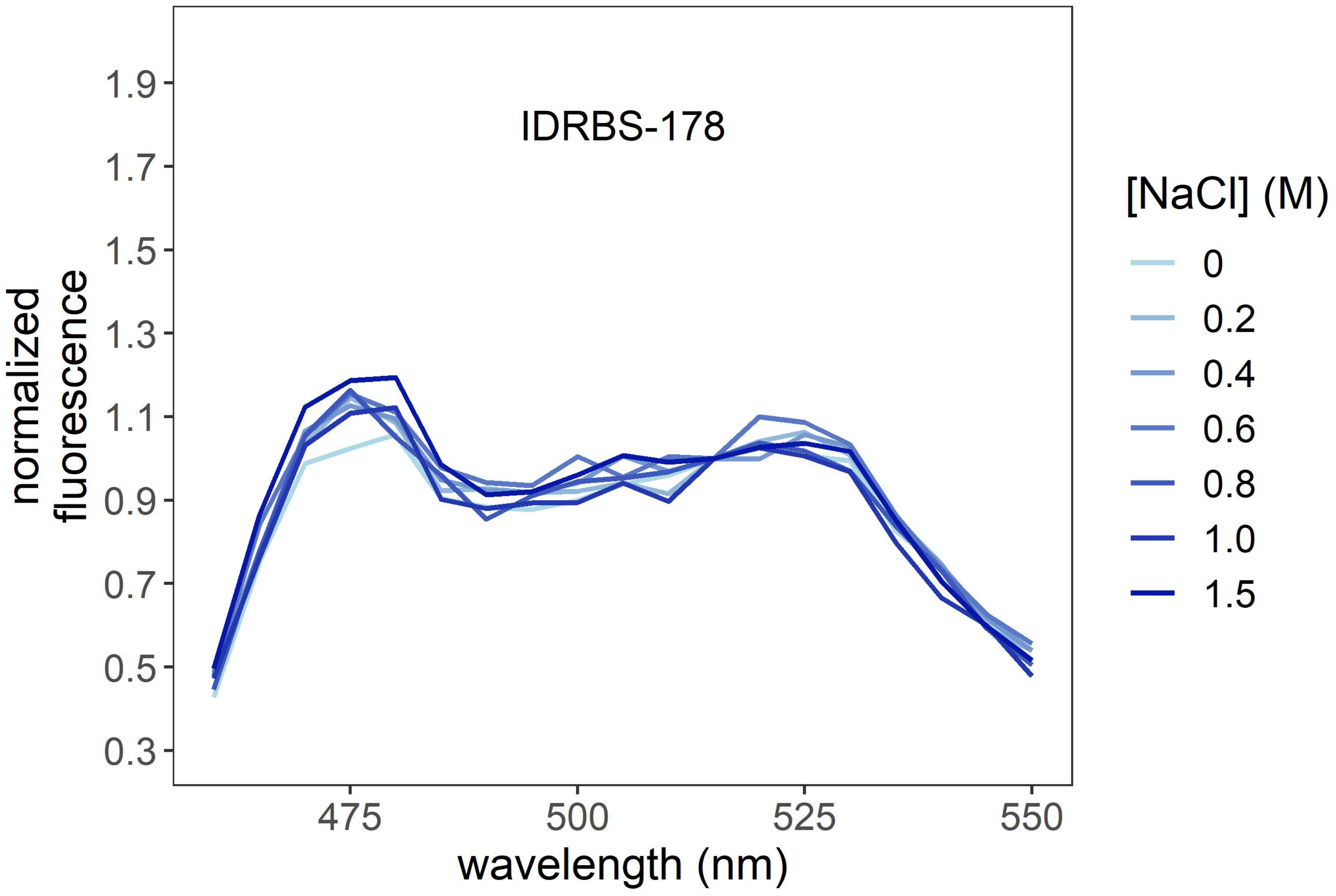
IDRBS-178
normalized
fluorescence
1.9
1.7
1.5
1.3
1.1
0.9
0.7
0.5
0.3
475
500
525
550
wavelength (nm)
[NaCl] (M)
0
0.2
0.4
0.6
0.8
1.0
1.5

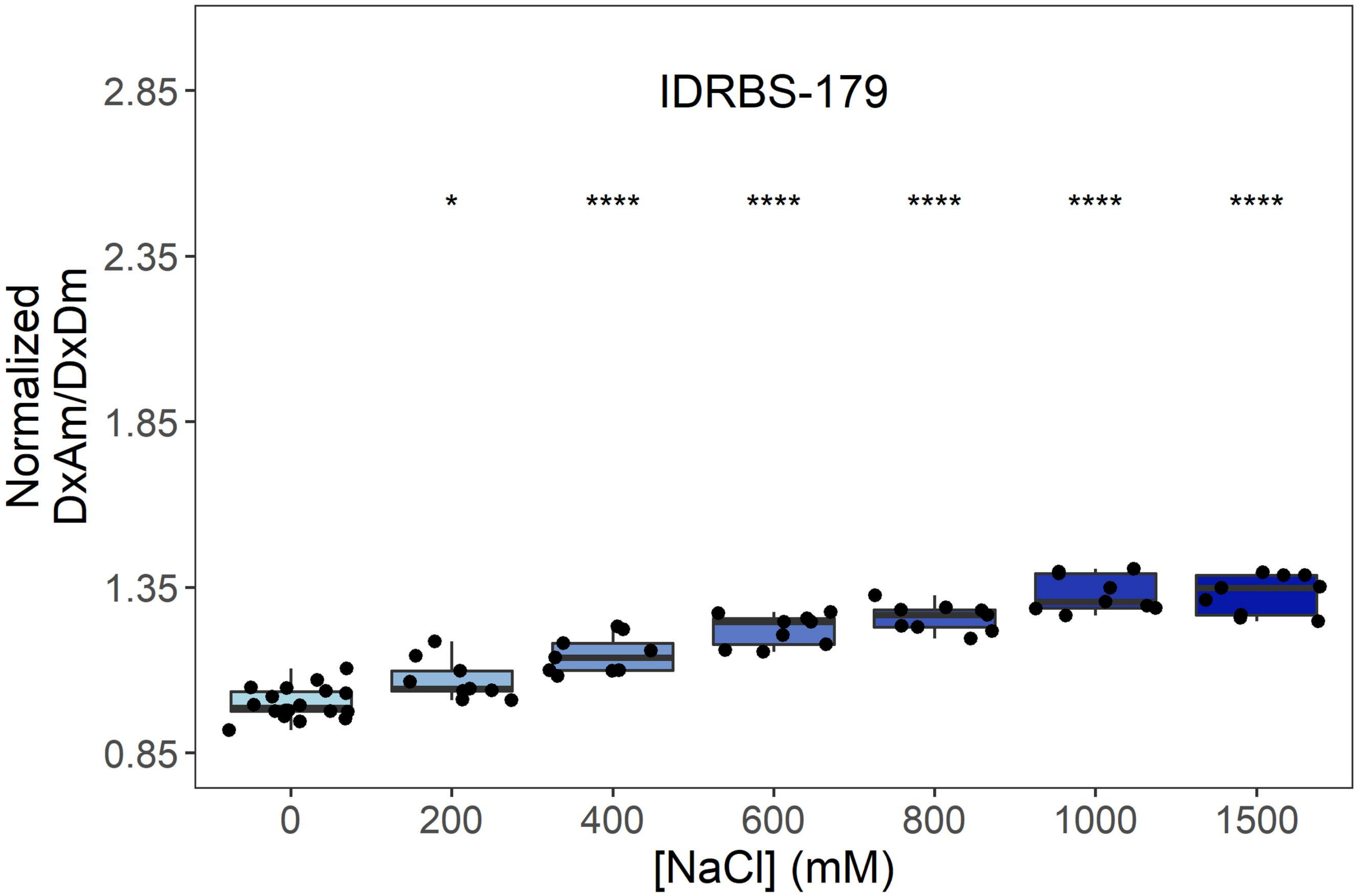
IDRBS-179
*
****
****
****
****
****
2.85
2.35
1.85
1.35
0.85
Normalized
DxAm/DxDm
0
200
400
600
800
1000
1500
[NaCl] (mM)

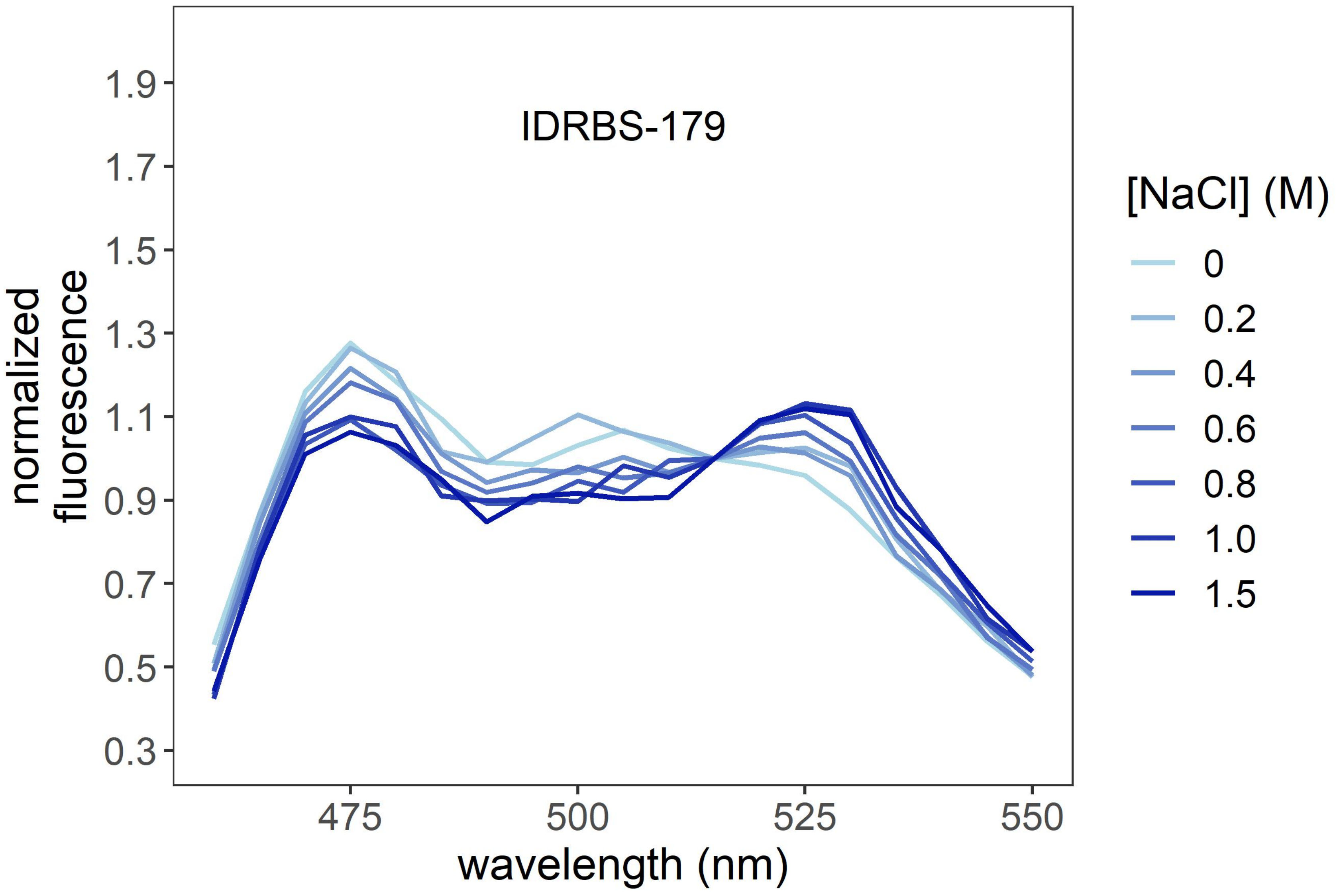
IDRBS-179
1.9
1.7
1.5
1.3
1.1
0.9
0.7
0.5
0.3
normalized
fluorescence
475
500
525
550
wavelength (nm)
[NaCl] (M)
0
0.2
0.4
0.6
0.8
1.0
1.5

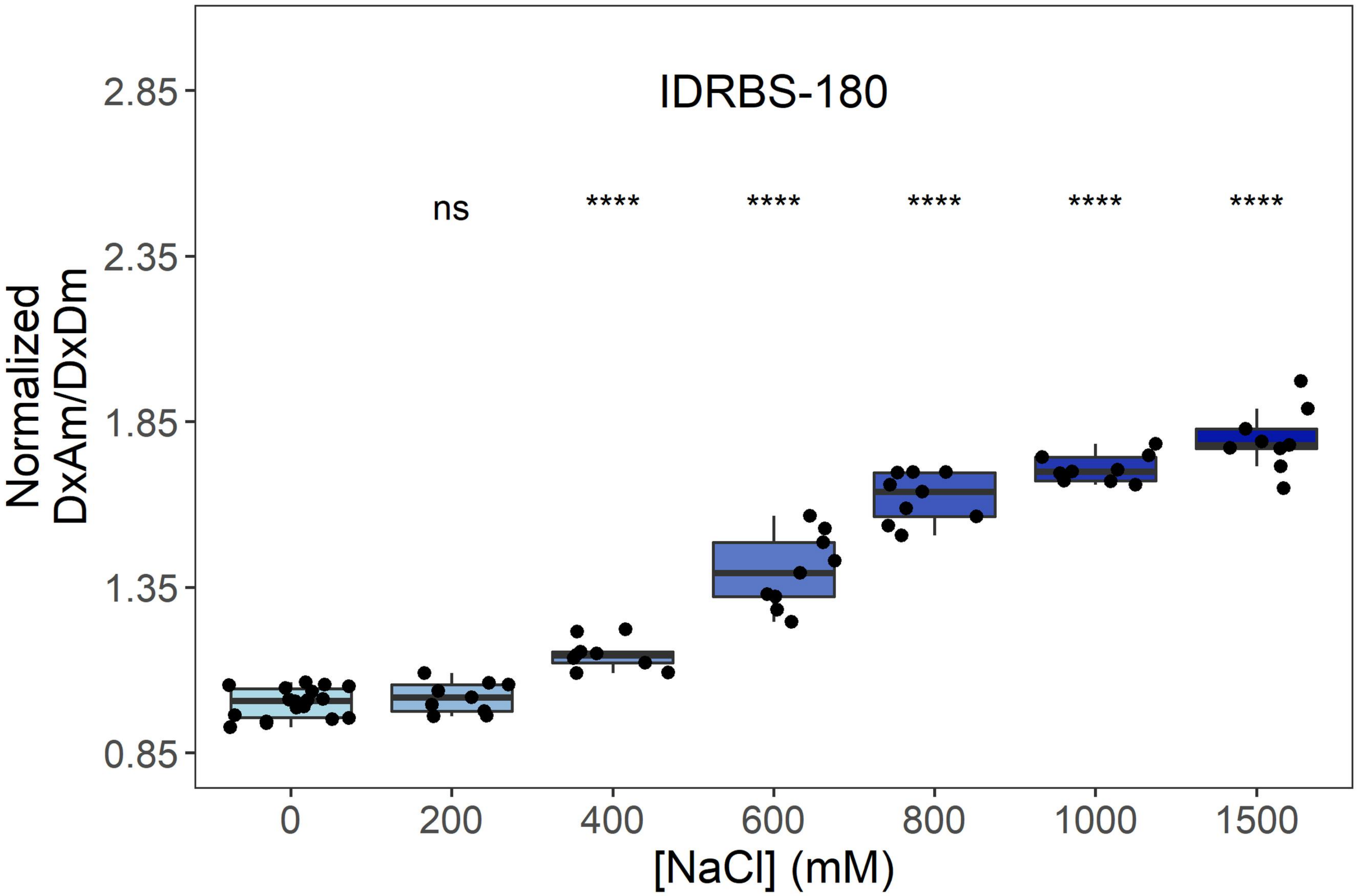
IDRBS-180
ns
****
****
****
****
****
2.85
2.35
1.85
1.35
0.85
Normalized
DxAm/DxDm
0
200
400
600
800
1000
1500
[NaCl] (mM)

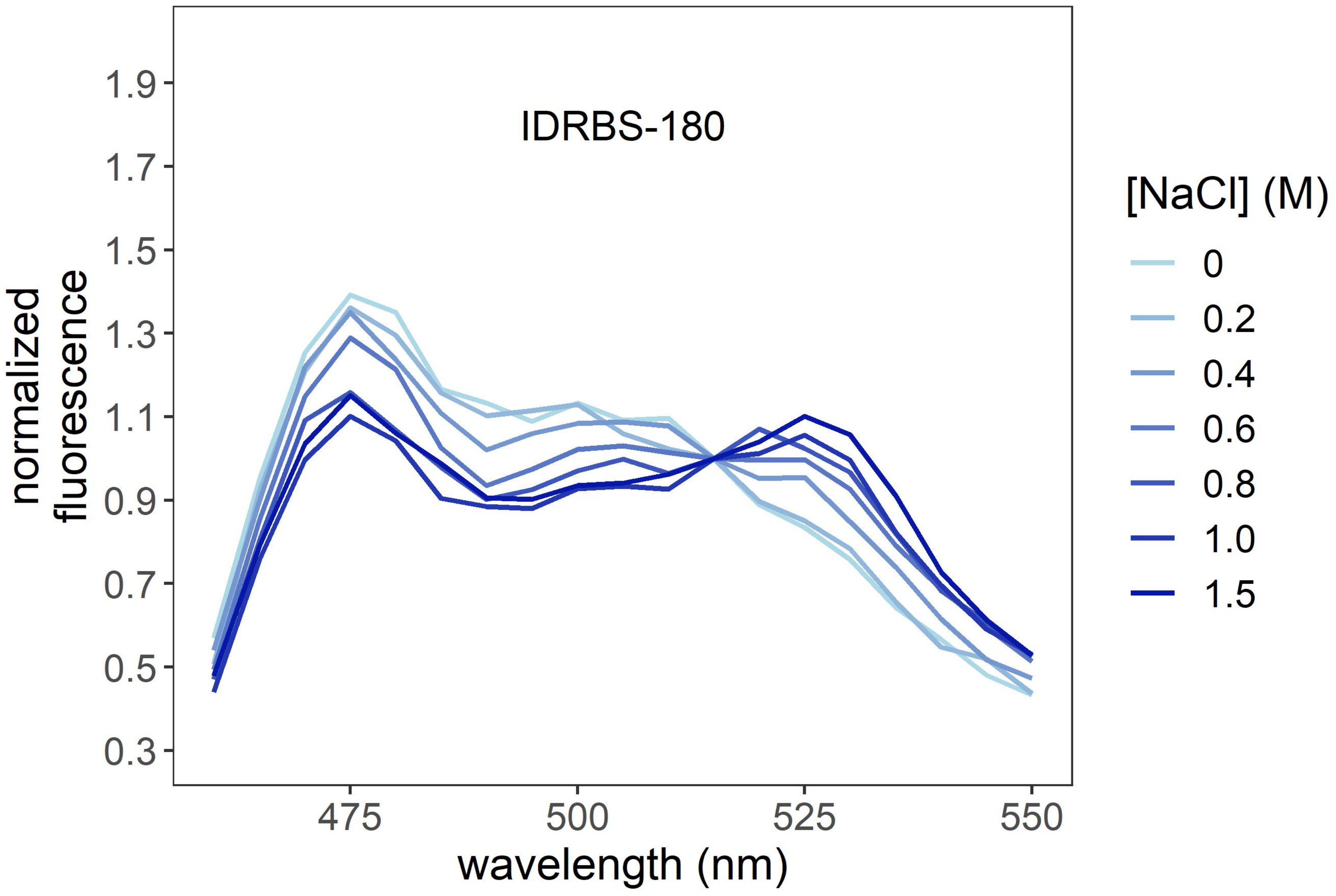
IDRBS-180
1.9
1.7
1.5
1.3
1.1
0.9
0.7
0.5
0.3
normalized
fluorescence
475
500
525
550
wavelength (nm)
[NaCl] (M)
0
0.2
0.4
0.6
0.8
1.0
1.5

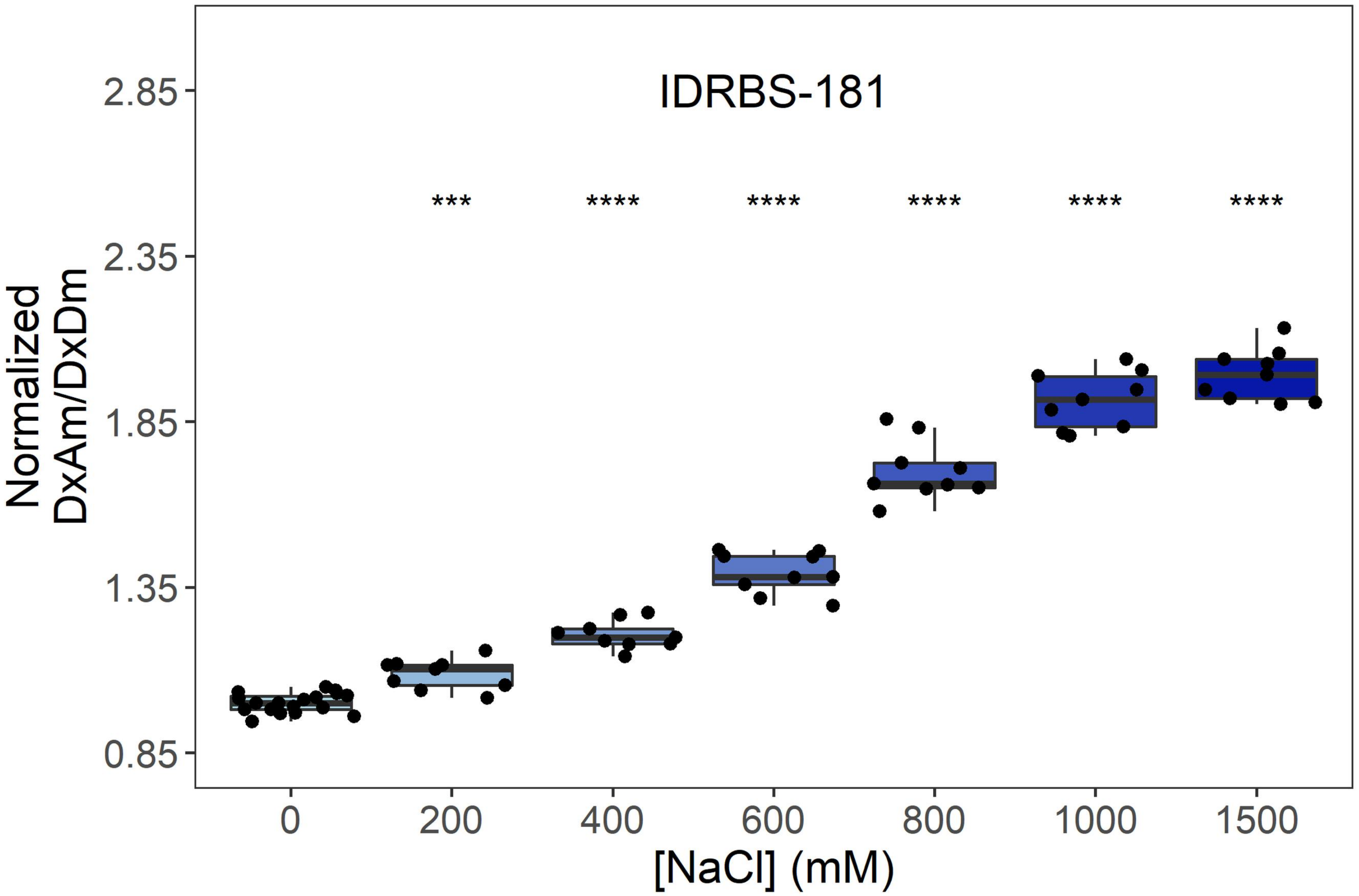

IDRBS-181
***
****
****
****
****
****
Normalized DxAm/DxDm
2.85
2.35
1.85
1.35
0.85
0
200
400
600
800
1000
1500
[NaCl] (mM)


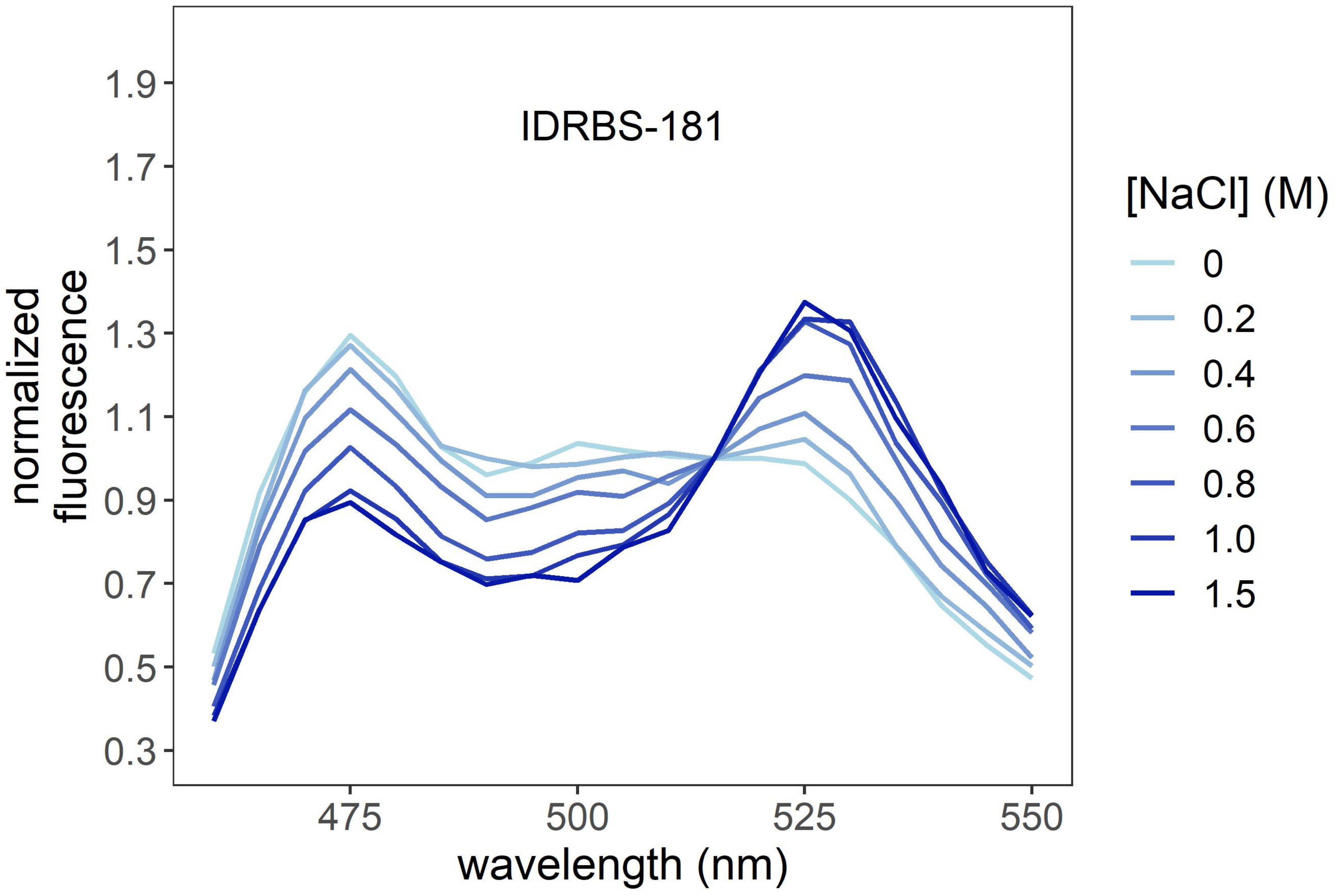

IDRBS-181
normalized fluorescence
1.9
1.7
1.5
1.3
1.1
0.9
0.7
0.5
0.3
475
500
525
550
wavelength (nm)
[NaCl] (M)
0
0.2
0.4
0.6
0.8
1.0
1.5

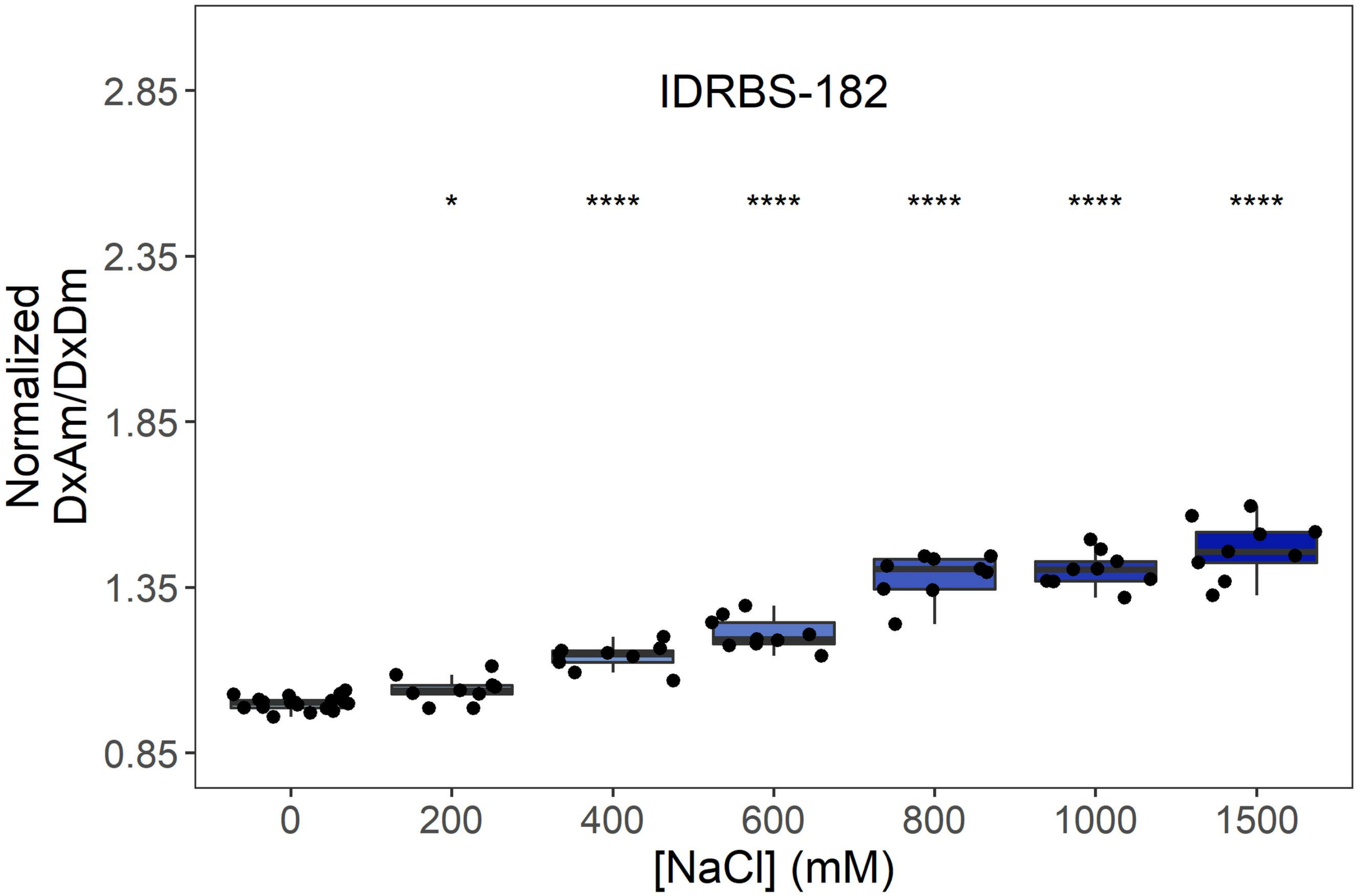
IDRBS-182
*
****
****
****
****
****
2.85
2.35
1.85
1.35
0.85
Normalized
DxAm/DxDm
0
200
400
600
800
1000
1500
[NaCl] (mM)

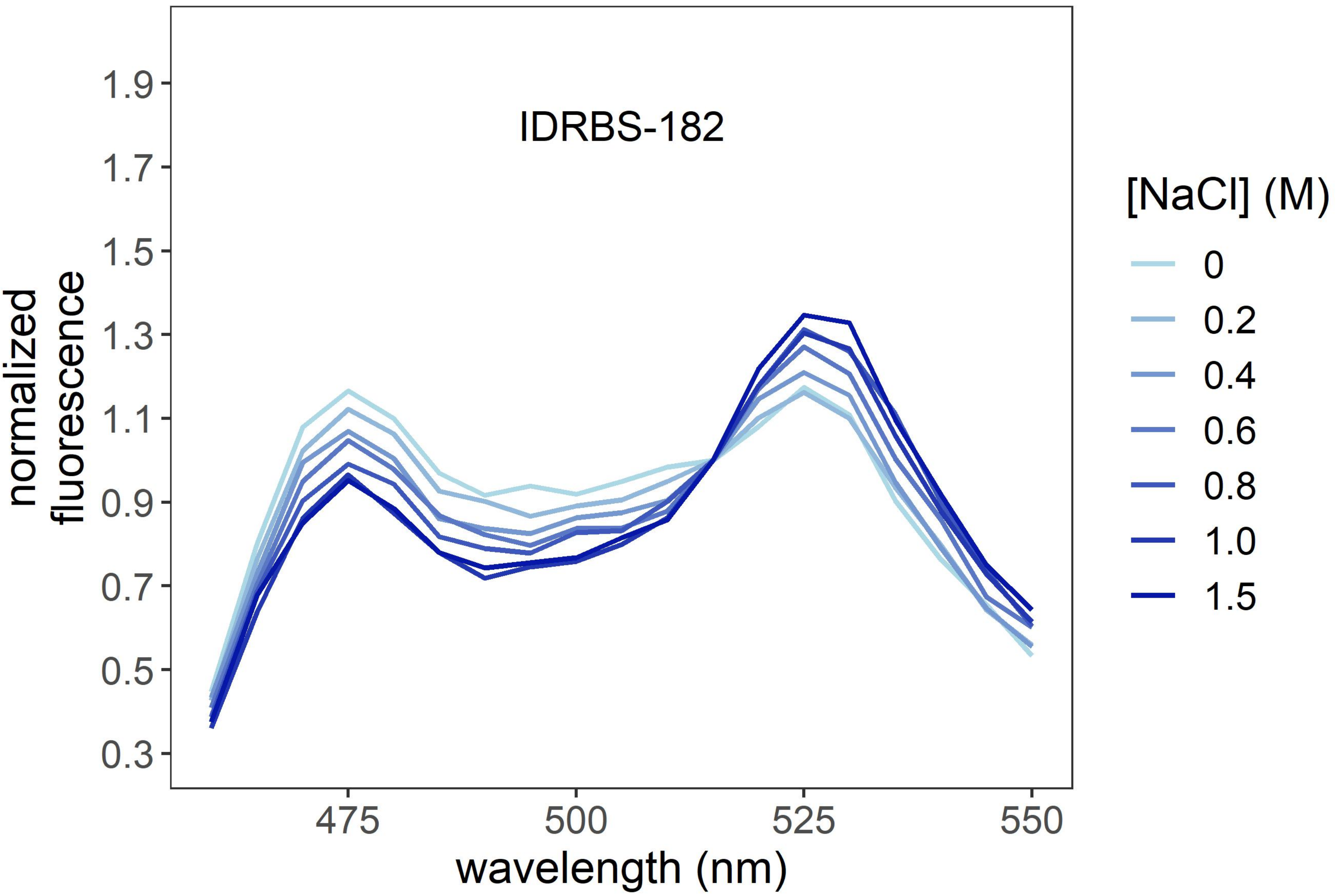
IDRBS-182
1.9
1.7
1.5
1.3
1.1
0.9
0.7
0.5
0.3
normalized
fluorescence
475
500
525
550
wavelength (nm)
[NaCl] (M)
0
0.2
0.4
0.6
0.8
1.0
1.5

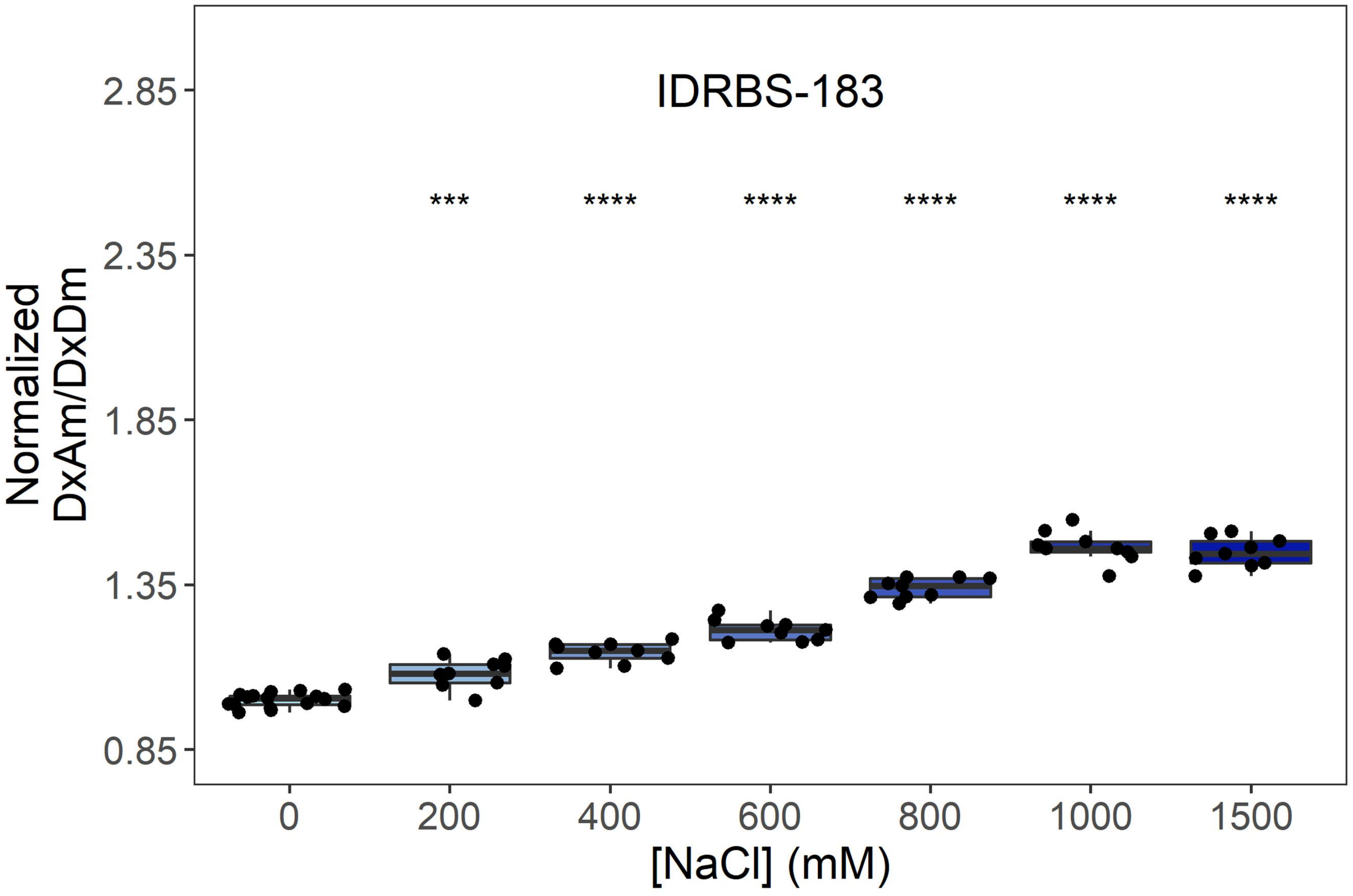
IDRBS-183
***
****
****
****
****
****
Normalized
DxAm/DxDm
2.85
2.35
1.85
1.35
0.85
0
200
400
600
800
1000
1500
[NaCl] (mM)

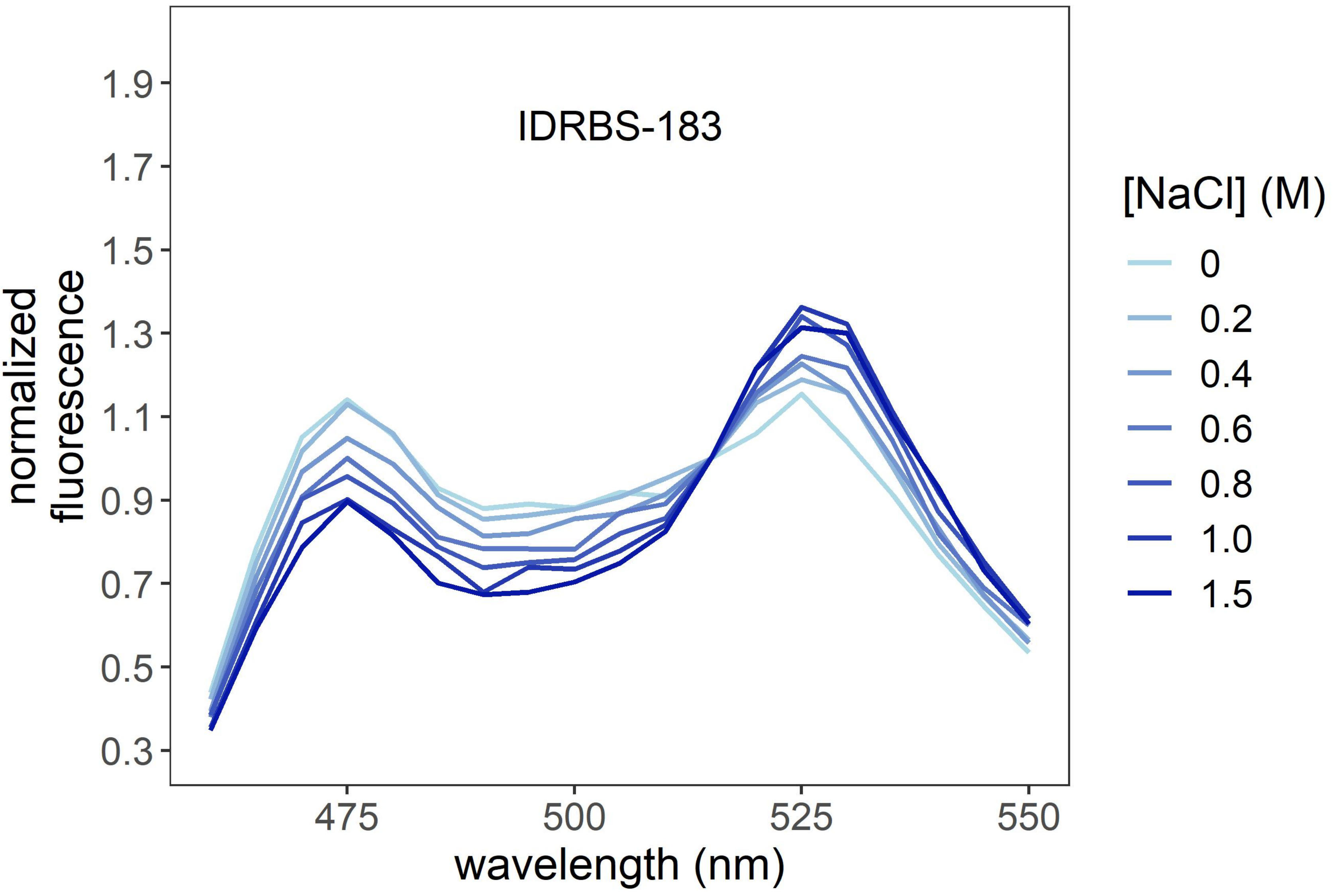
IDRBS-183
normalized
fluorescence
1.9
1.7
1.5
1.3
1.1
0.9
0.7
0.5
0.3
475
500
525
550
wavelength (nm)
[NaCl] (M)
0
0.2
0.4
0.6
0.8
1.0
1.5

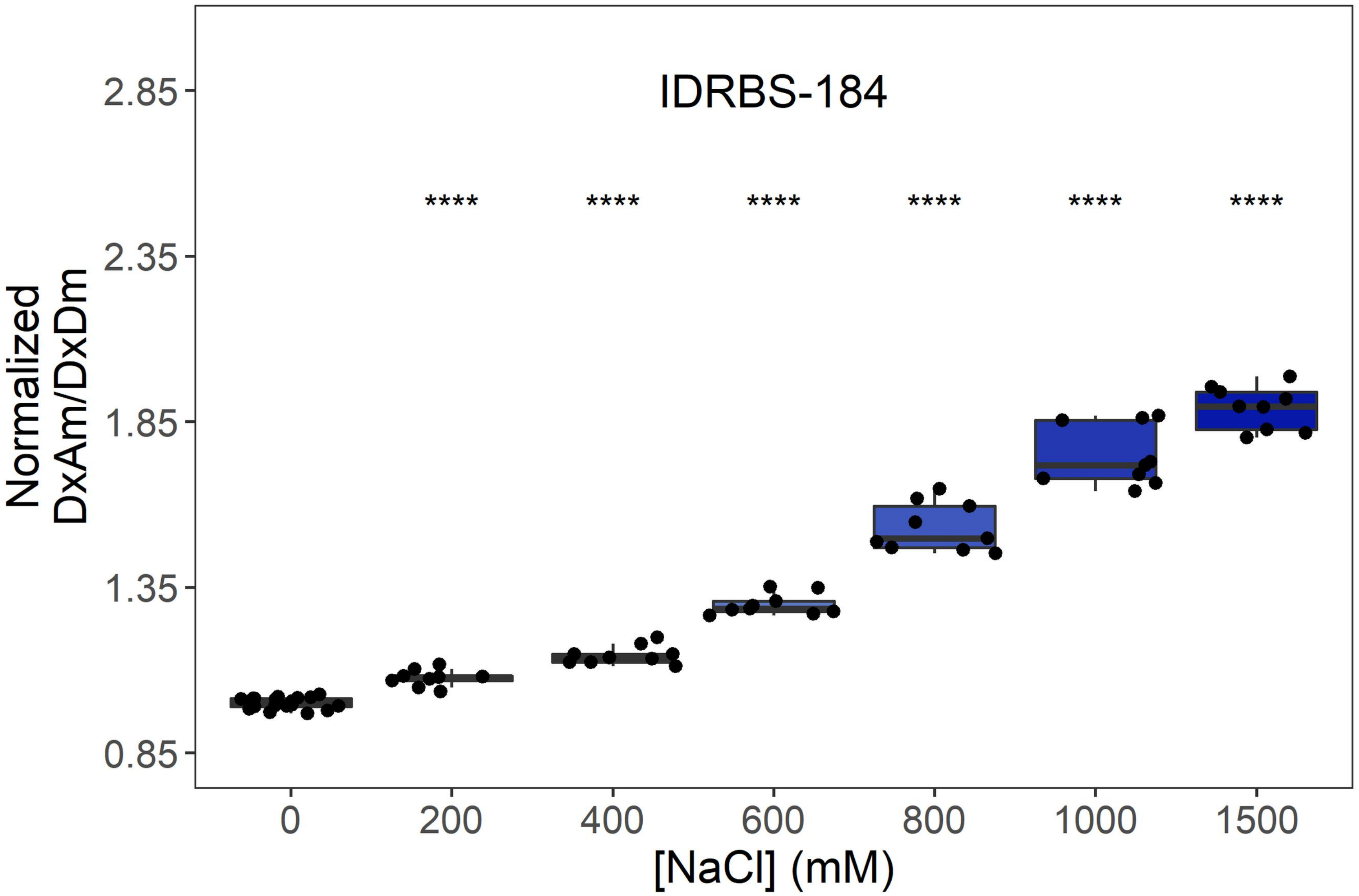

IDRBS-184
****
2.85
2.35
1.85
1.35
0.85
Normalized DxAm/DxDm
0
200
400
600
800
1000
1500
[NaCl] (mM)


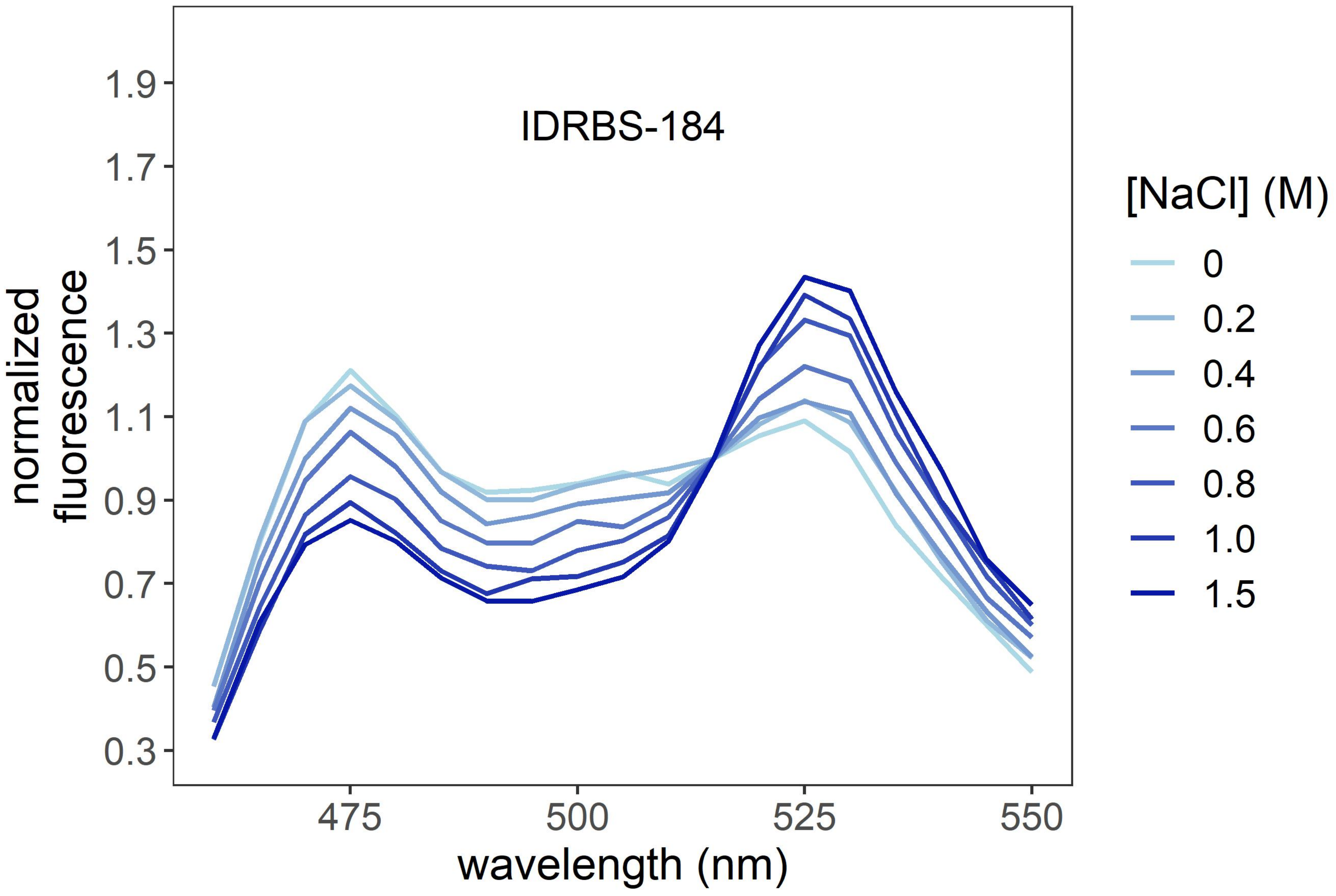

IDRBS-184
1.9
1.7
1.5
1.3
1.1
0.9
0.7
0.5
0.3
normalized fluorescence
475
500
525
550
wavelength (nm)
[NaCl] (M)
0
0.2
0.4
0.6
0.8
1.0
1.5

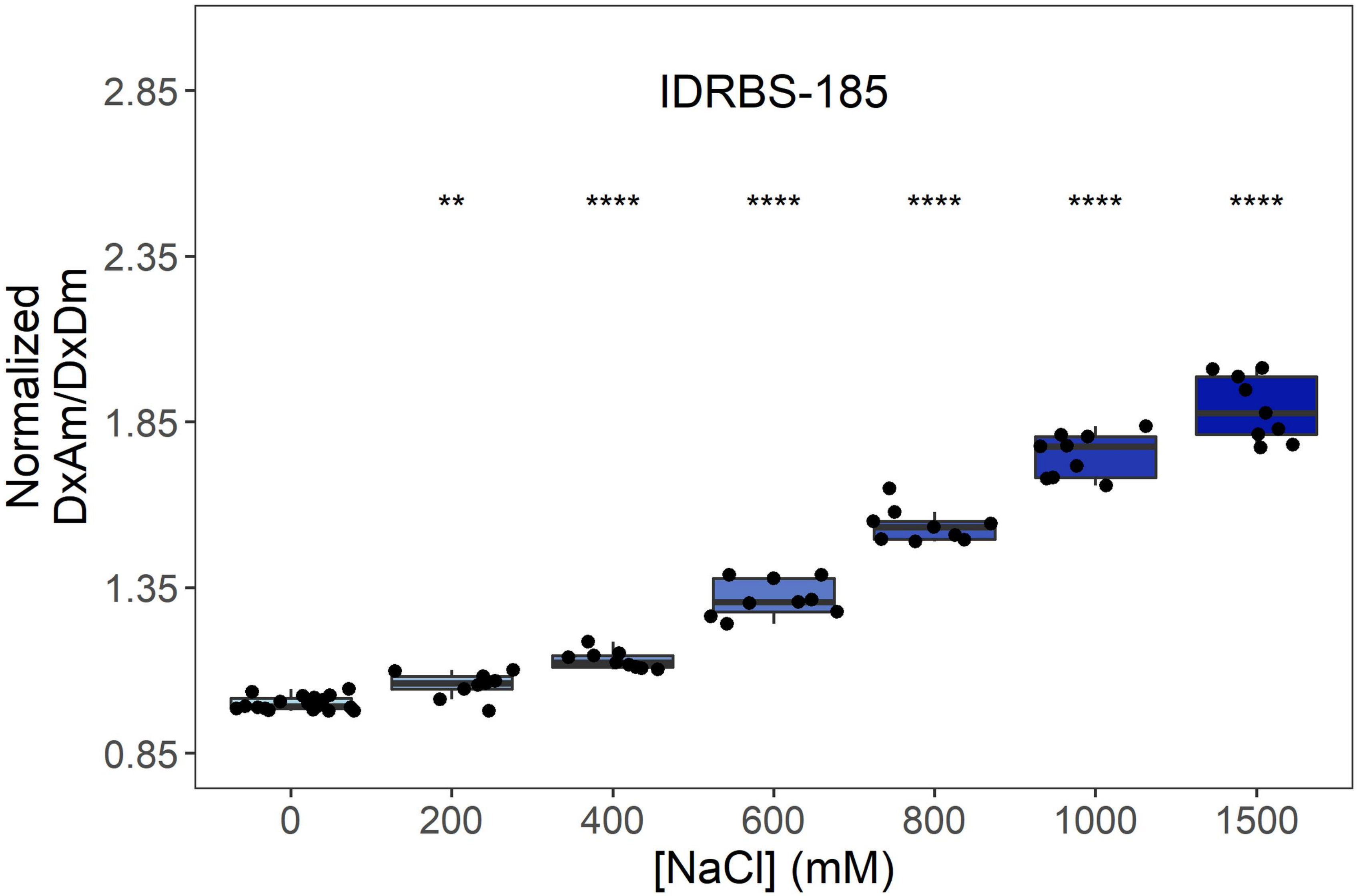

IDRBS-185
Normalized DxAm/DxDm
2.85
2.35
1.85
1.35
0.85
**
****
****
****
****
****
0
200
400
600
800
1000
1500
[NaCl] (mM)


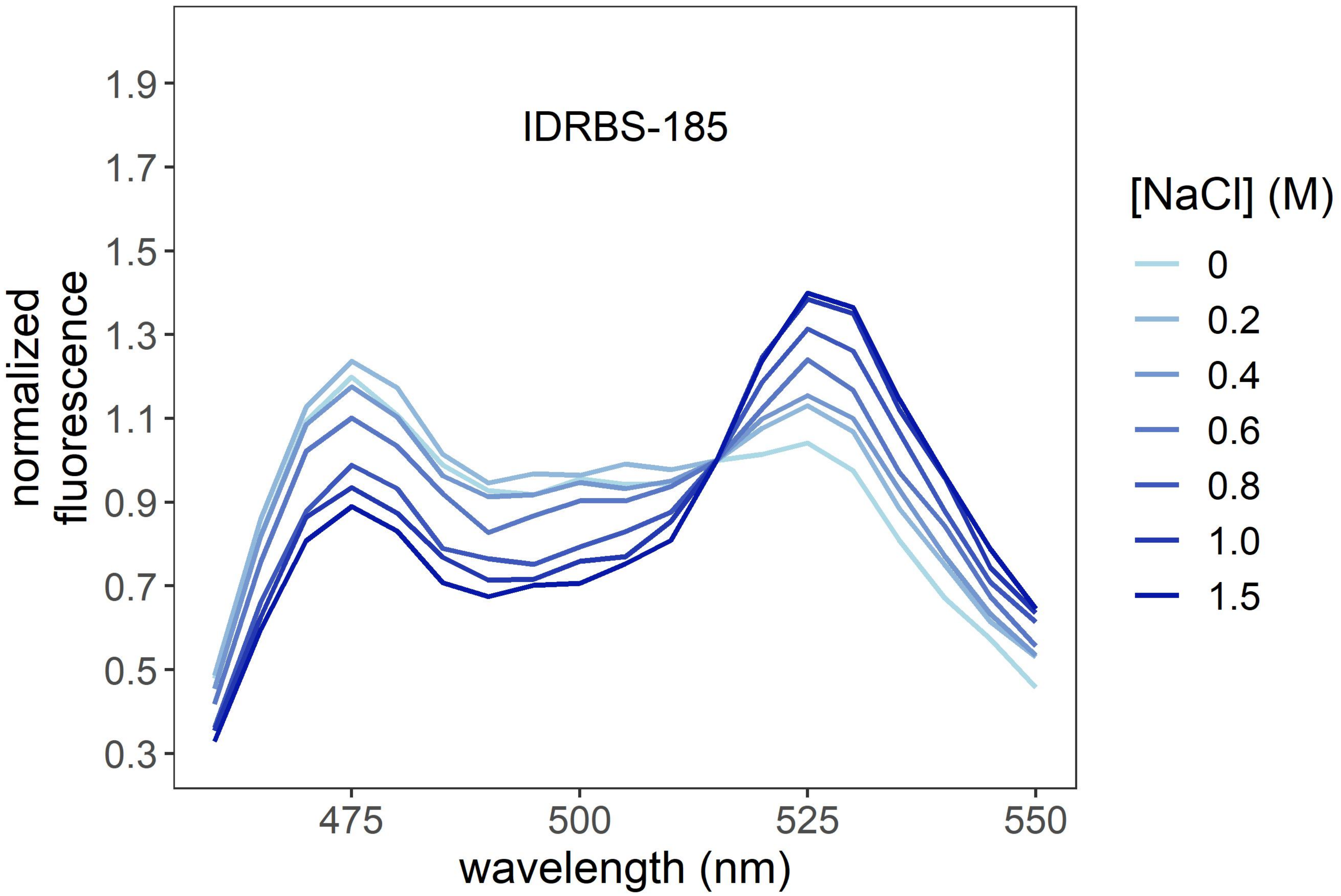

IDRBS-185
normalized fluorescence
1.9
1.7
1.5
1.3
1.1
0.9
0.7
0.5
0.3
475
500
525
550
wavelength (nm)
[NaCl] (M)
0
0.2
0.4
0.6
0.8
1.0
1.5

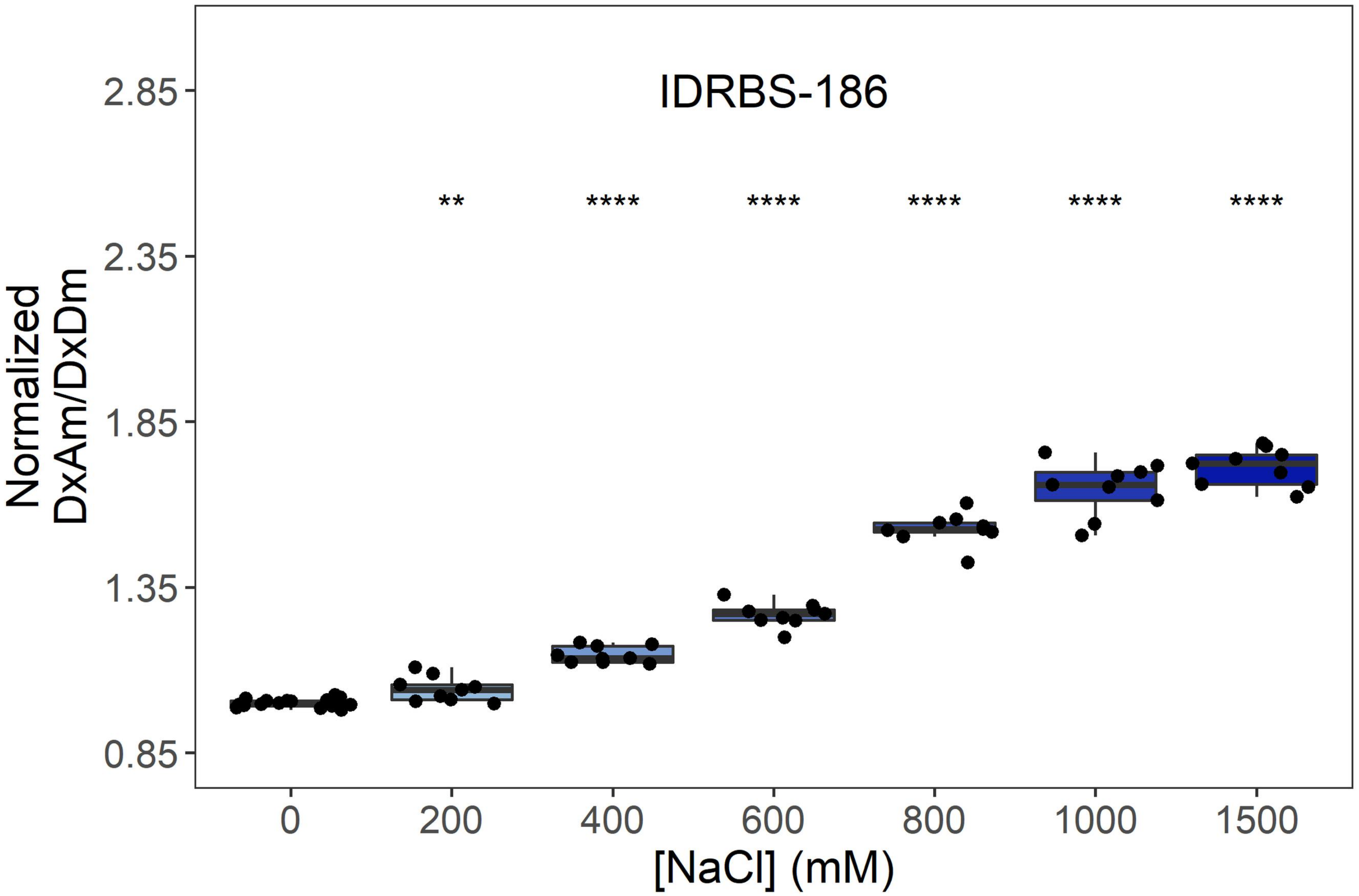
IDRBS-186
**
****
****
****
****
****
Normalized DxAm/DxDm
2.85
2.35
1.85
1.35
0.85
0
200
400
600
800
1000
1500
[NaCl] (mM)

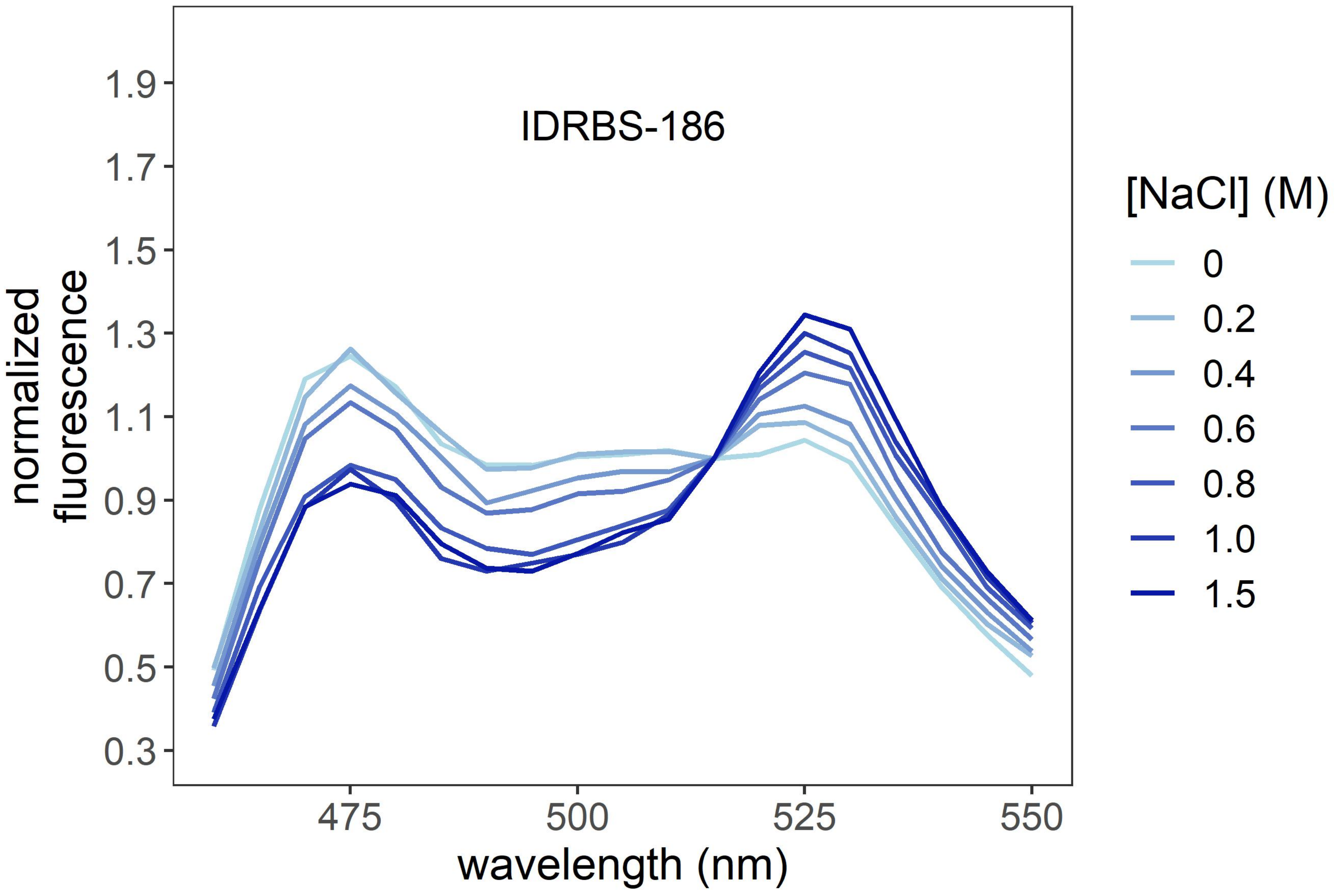
IDRBS-186
normalized fluorescence
1.9
1.7
1.5
1.3
1.1
0.9
0.7
0.5
0.3
475
500
525
550
wavelength (nm)
[NaCl] (M)
0
0.2
0.4
0.6
0.8
1.0
1.5

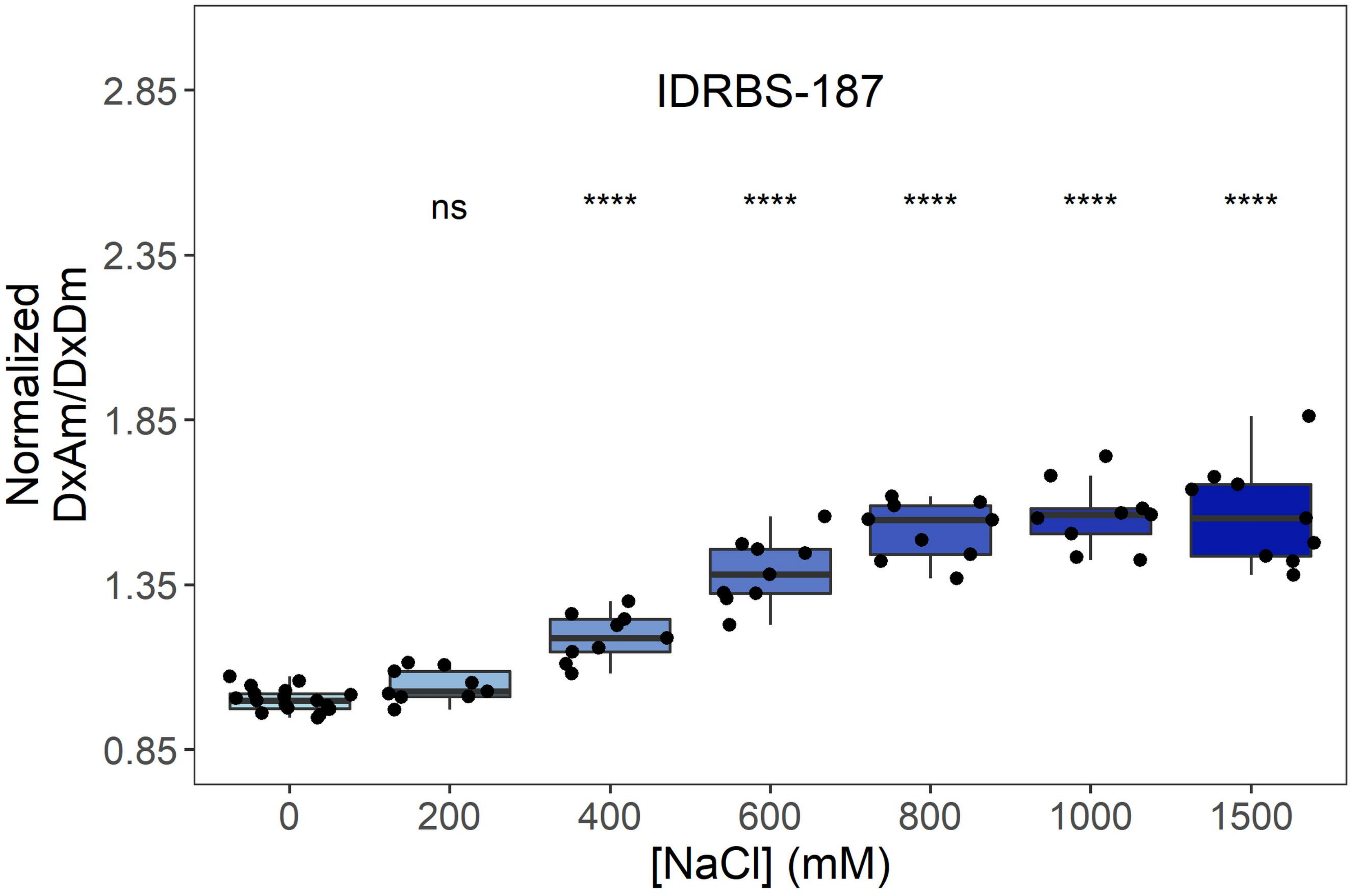

IDRBS-187
ns
****
****
****
****
****
2.85
2.35
1.85
1.35
0.85
Normalized
DxAm/DxDm
0
200
400
600
800
1000
1500
[NaCl] (mM)


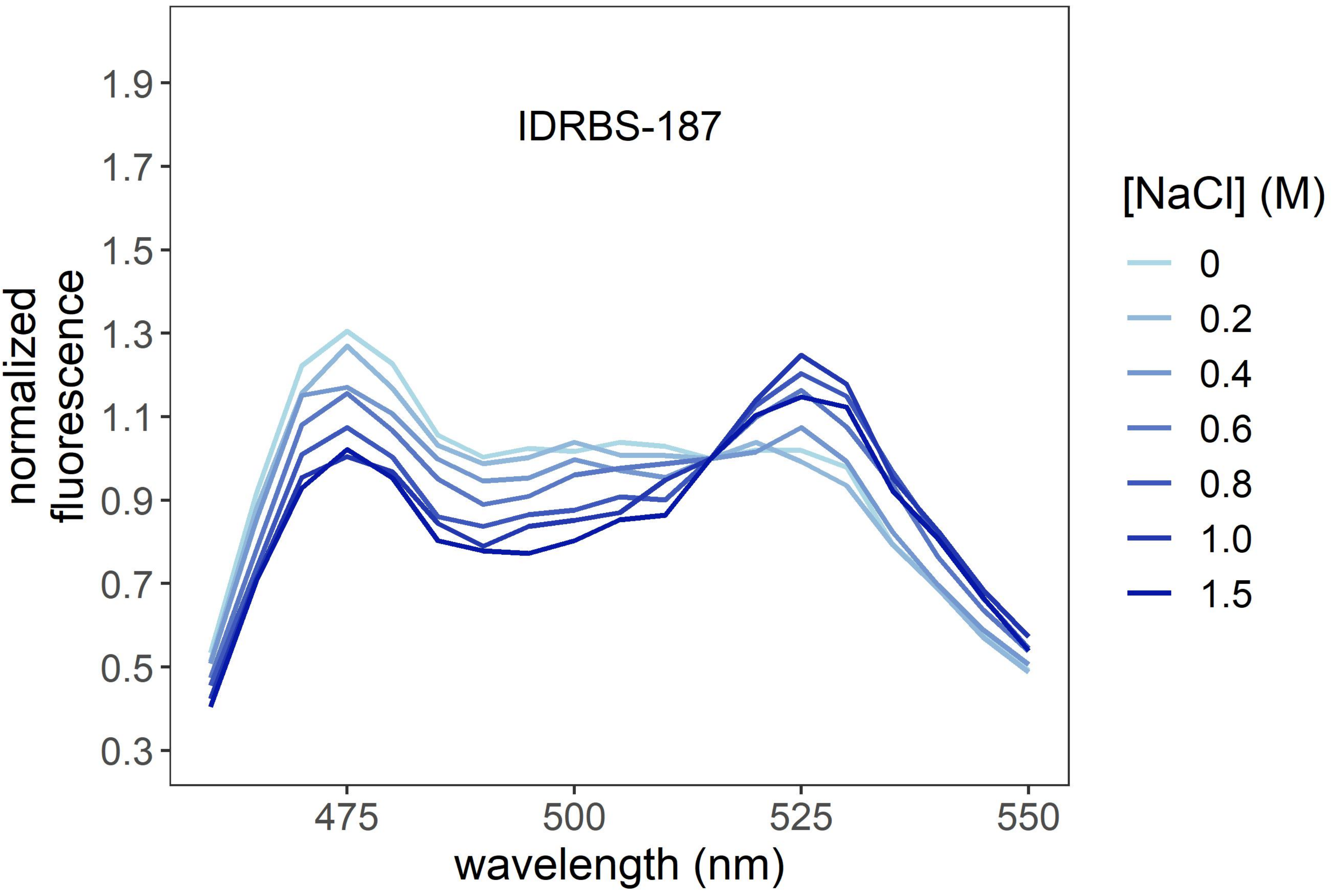

IDRBS-187
1.9
1.7
1.5
1.3
1.1
0.9
0.7
0.5
0.3
normalized
fluorescence
475
500
525
550
wavelength (nm)
[NaCl] (M)
0
0.2
0.4
0.6
0.8
1.0
1.5

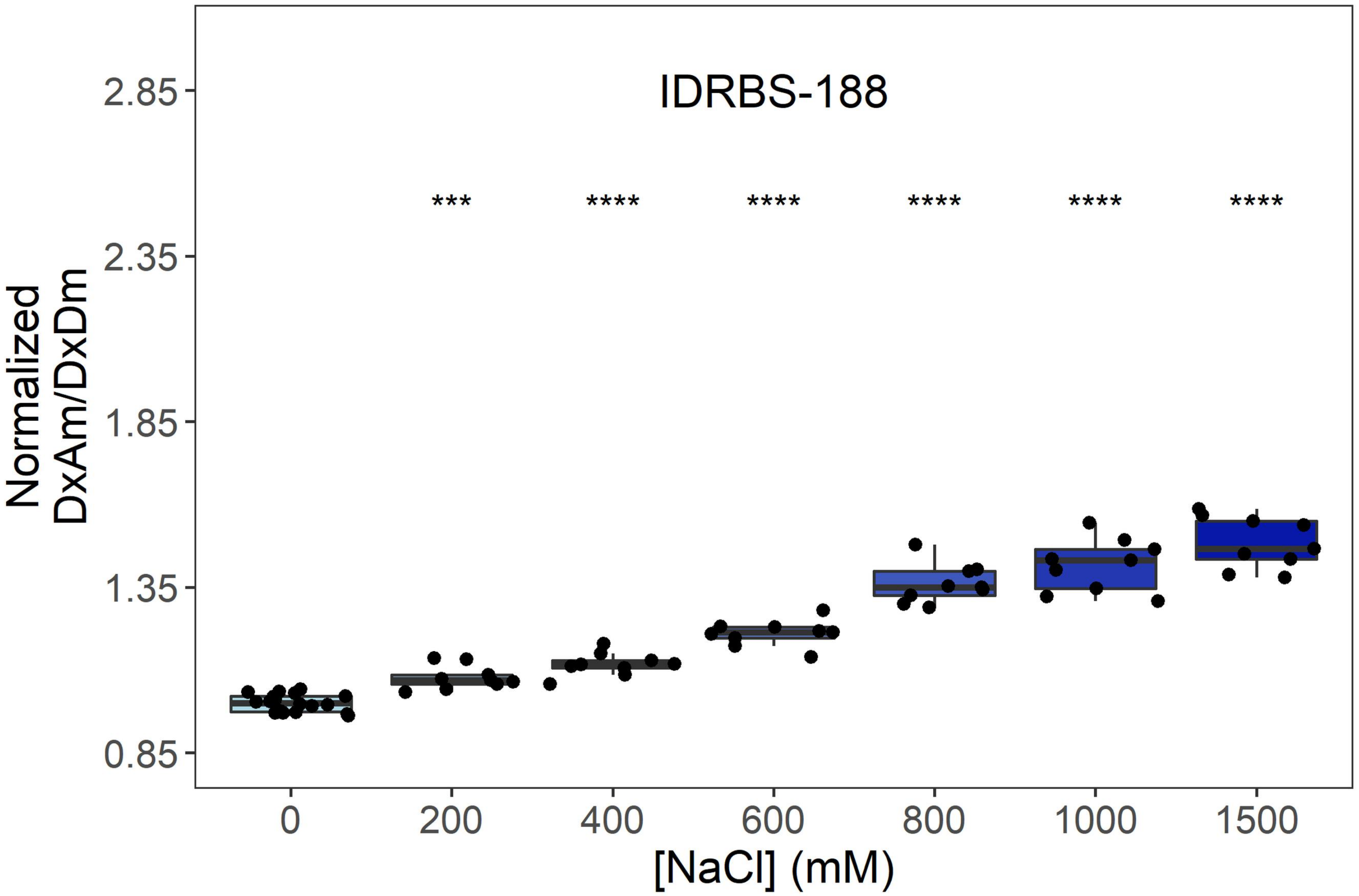

IDRBS-188
***
****
****
****
****
****
Normalized
DxAm/DxDm
2.85
2.35
1.85
1.35
0.85
0
200
400
600
800
1000
1500
[NaCl] (mM)


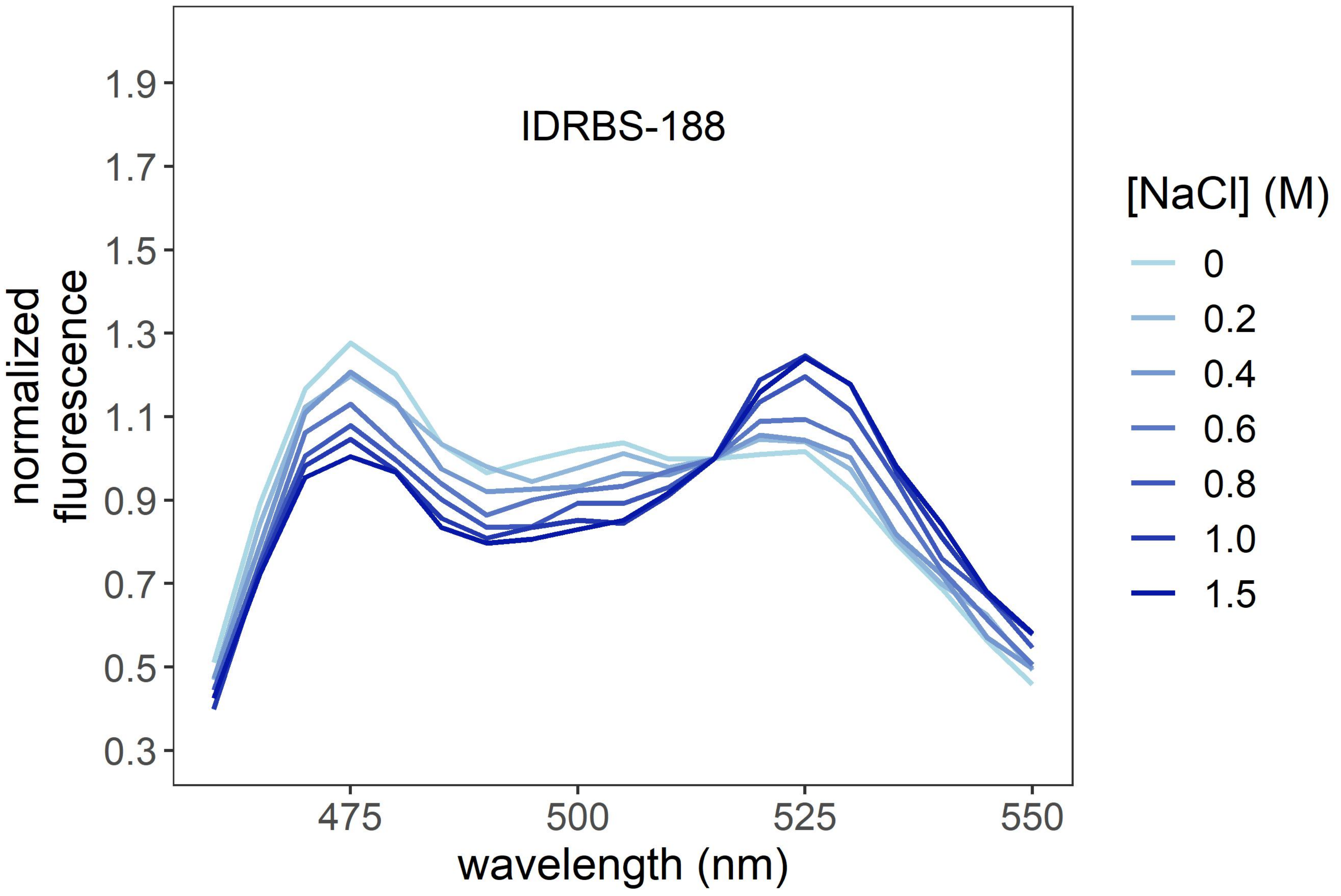

IDRBS-188
normalized
fluorescence
1.9
1.7
1.5
1.3
1.1
0.9
0.7
0.5
0.3
475
500
525
550
wavelength (nm)
[NaCl] (M)
0
0.2
0.4
0.6
0.8
1.0
1.5

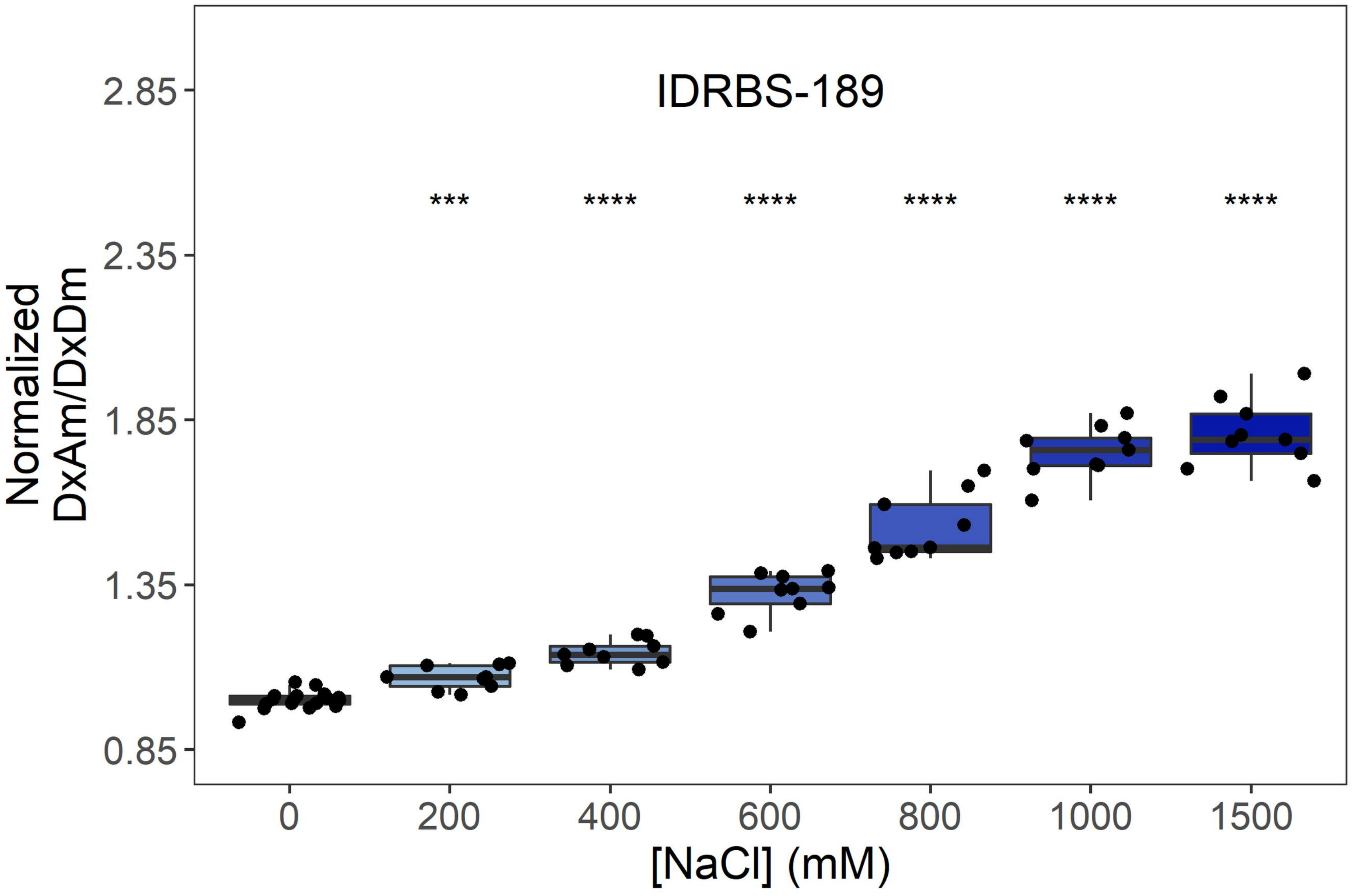
IDRBS-189
***
****
****
****
****
****
2.85
2.35
1.85
1.35
0.85
Normalized DxAm/DxDm
0
200
400
600
800
1000
1500
[NaCl] (mM)

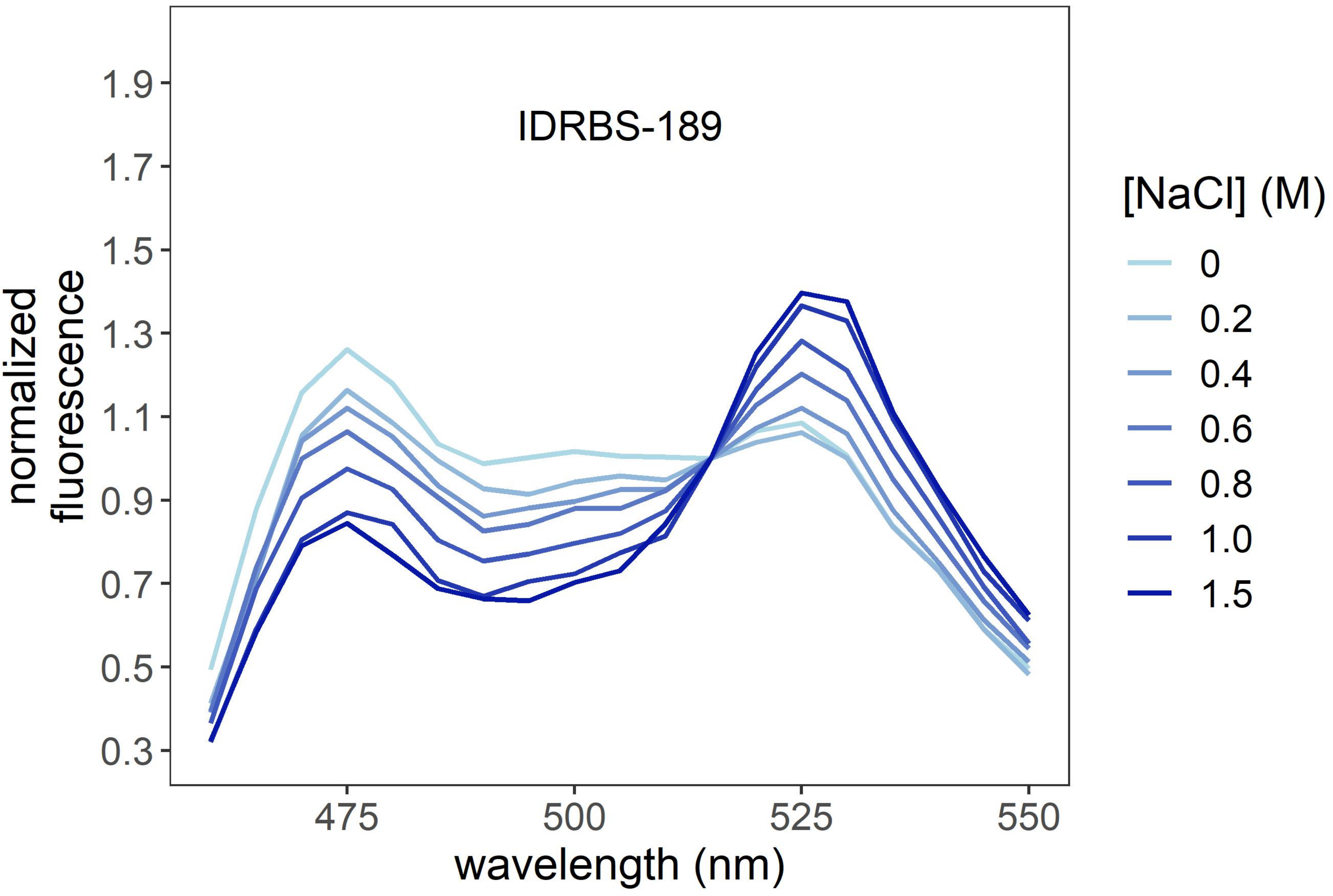
IDRBS-189
1.9
1.7
1.5
1.3
1.1
0.9
0.7
0.5
0.3
normalized fluorescence
475
500
525
550
wavelength (nm)
[NaCl] (M)
0
0.2
0.4
0.6
0.8
1.0
1.5

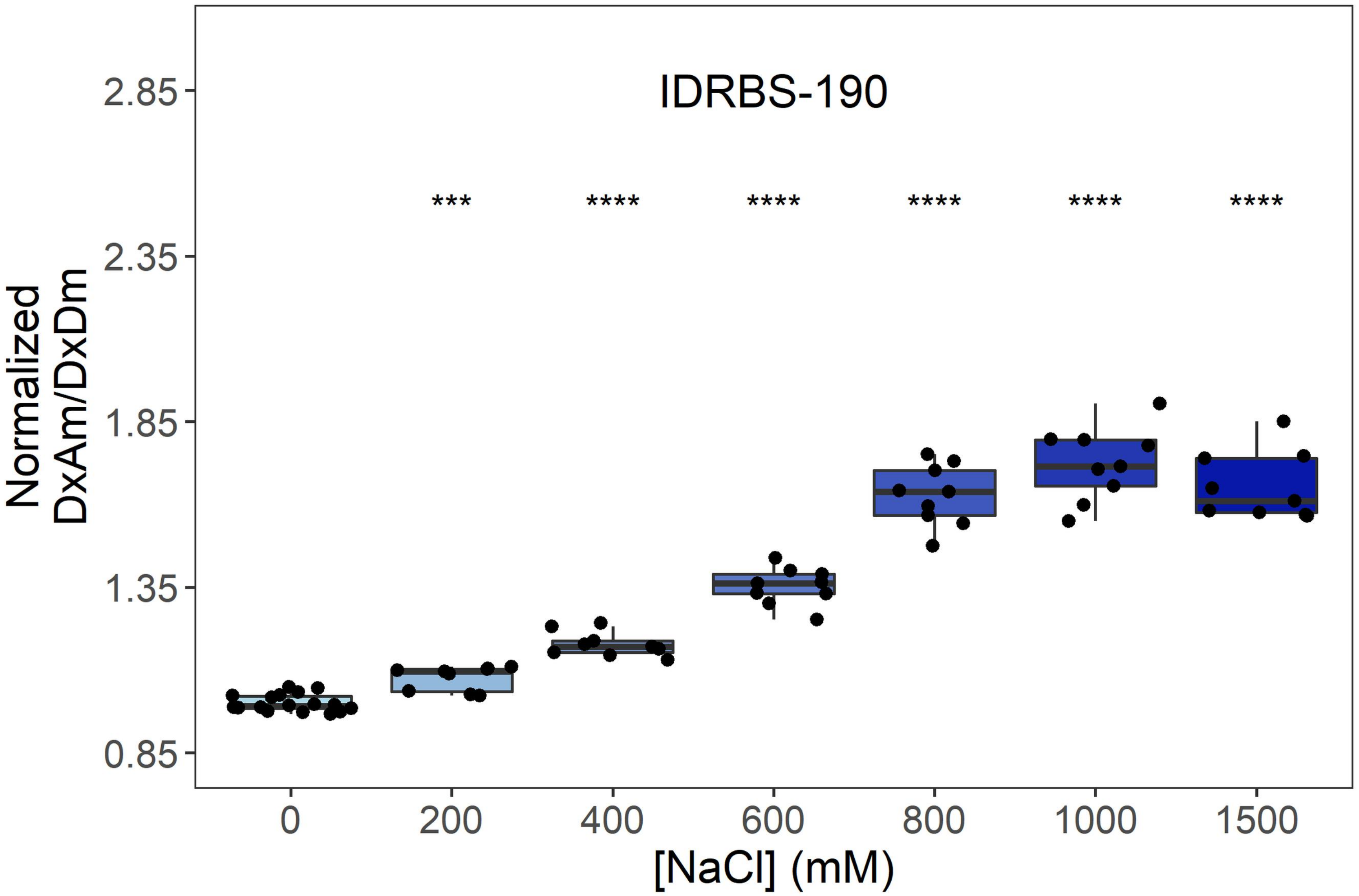
IDRBS-190
Normalized DxAm/DxDm
2.85
2.35
1.85
1.35
0.85
***
****
****
****
****
****
0
200
400
600
800
1000
1500
[NaCl] (mM)

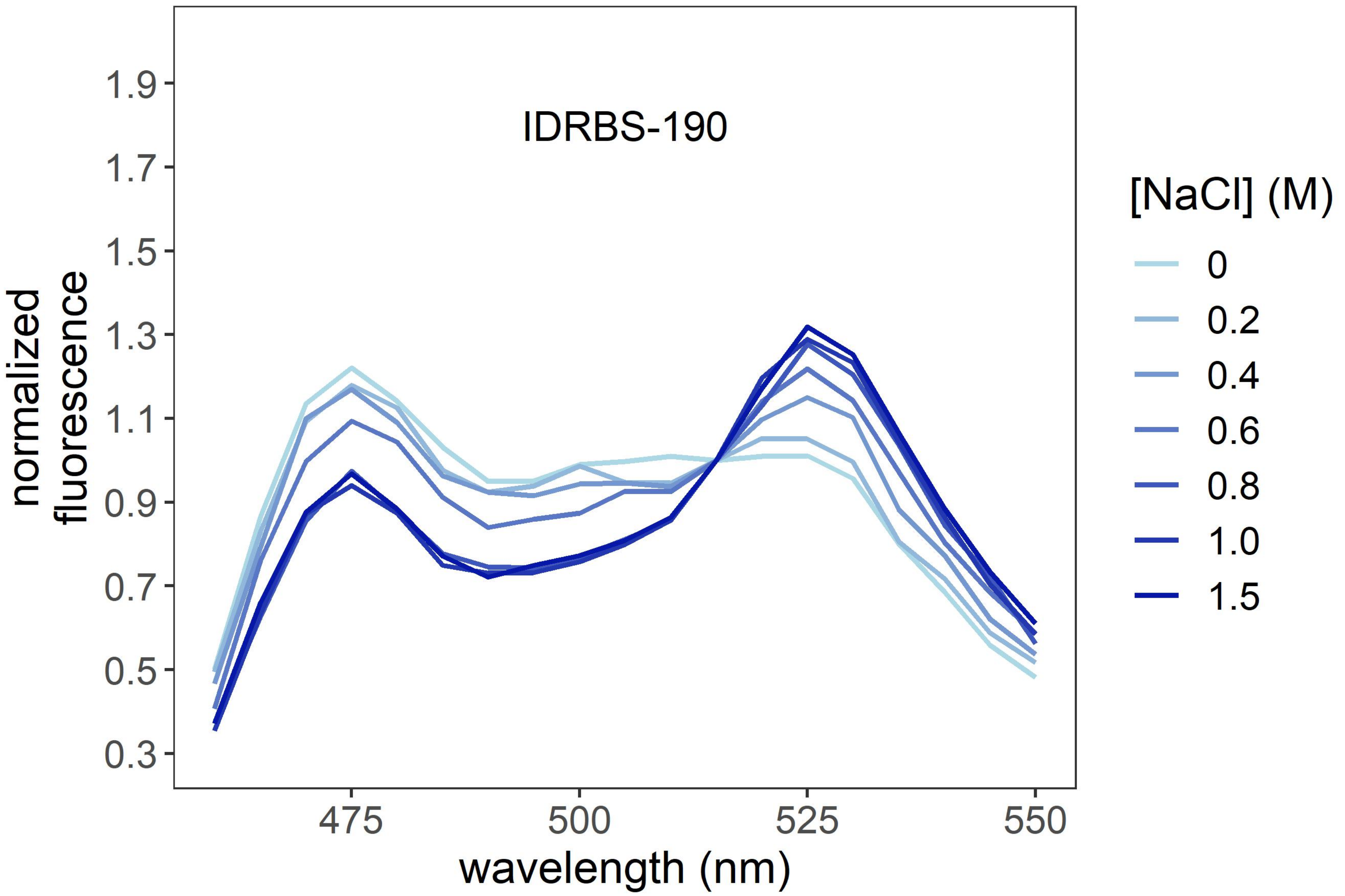
IDRBS-190
normalized fluorescence
1.9
1.7
1.5
1.3
1.1
0.9
0.7
0.5
0.3
475
500
525
550
wavelength (nm)
[NaCl] (M)
0
0.2
0.4
0.6
0.8
1.0
1.5

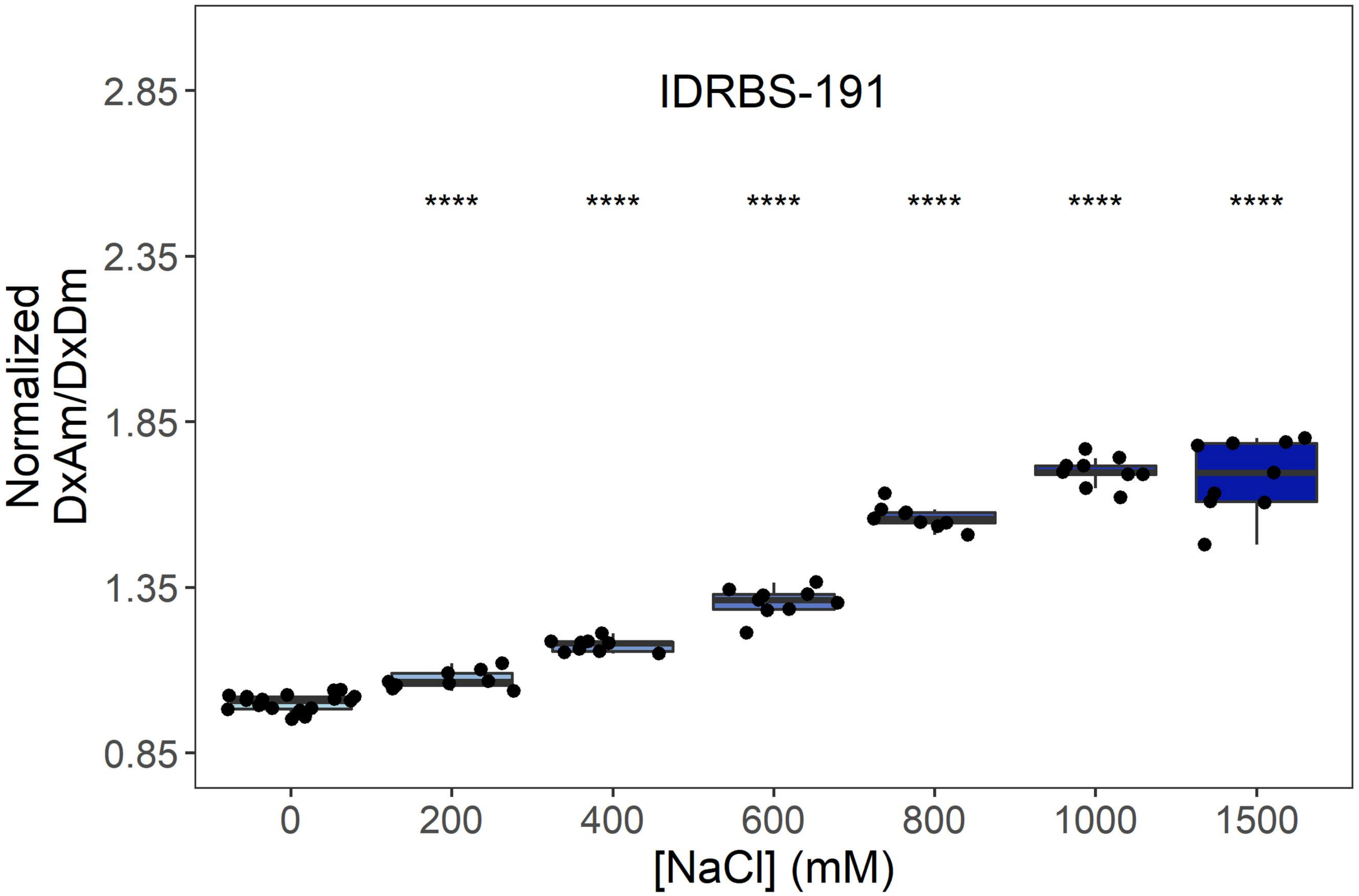
IDRBS-191
****
Normalized
DxAm/DxDm
2.85
2.35
1.85
1.35
0.85
0
200
400
600
800
1000
1500
[NaCl] (mM)

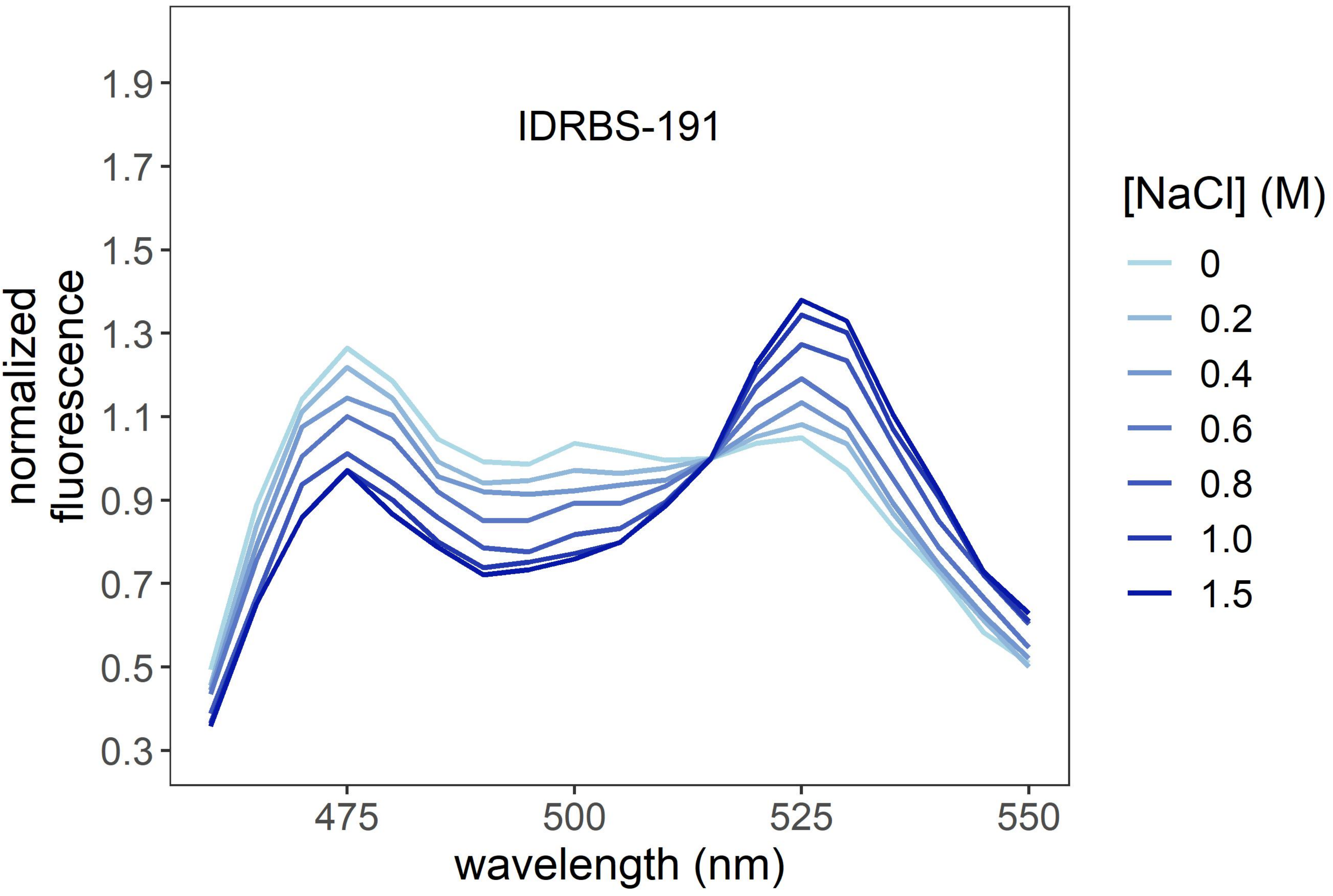
IDRBS-191
normalized
fluorescence
1.9
1.7
1.5
1.3
1.1
0.9
0.7
0.5
0.3
475
500
525
550
wavelength (nm)
[NaCl] (M)
0
0.2
0.4
0.6
0.8
1.0
1.5

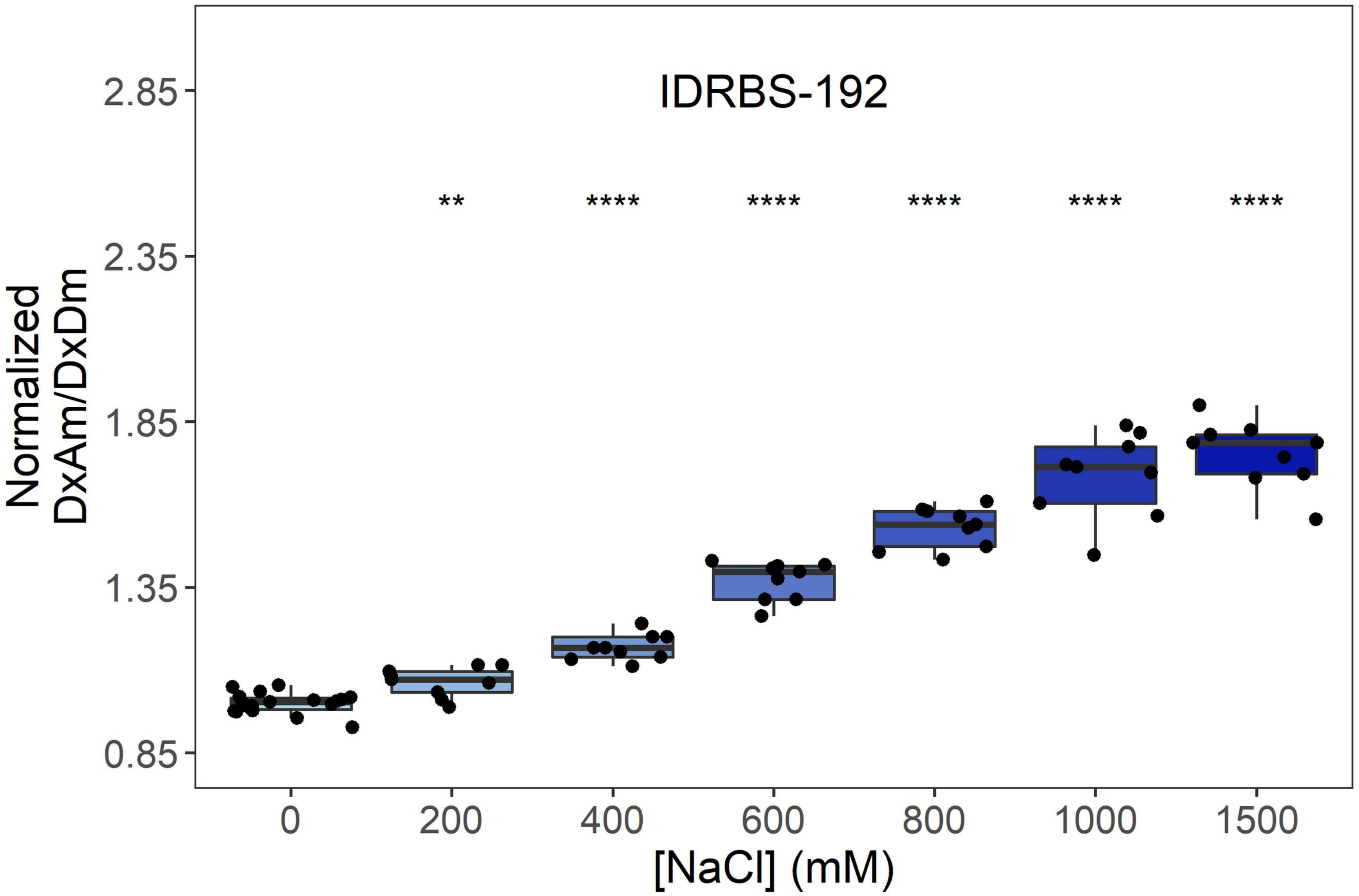
IDRBS-192
Normalized DxAm/DxDm
2.85
2.35
1.85
1.35
0.85
**
****
****
****
****
****
0
200
400
600
800
1000
1500
[NaCl] (mM)

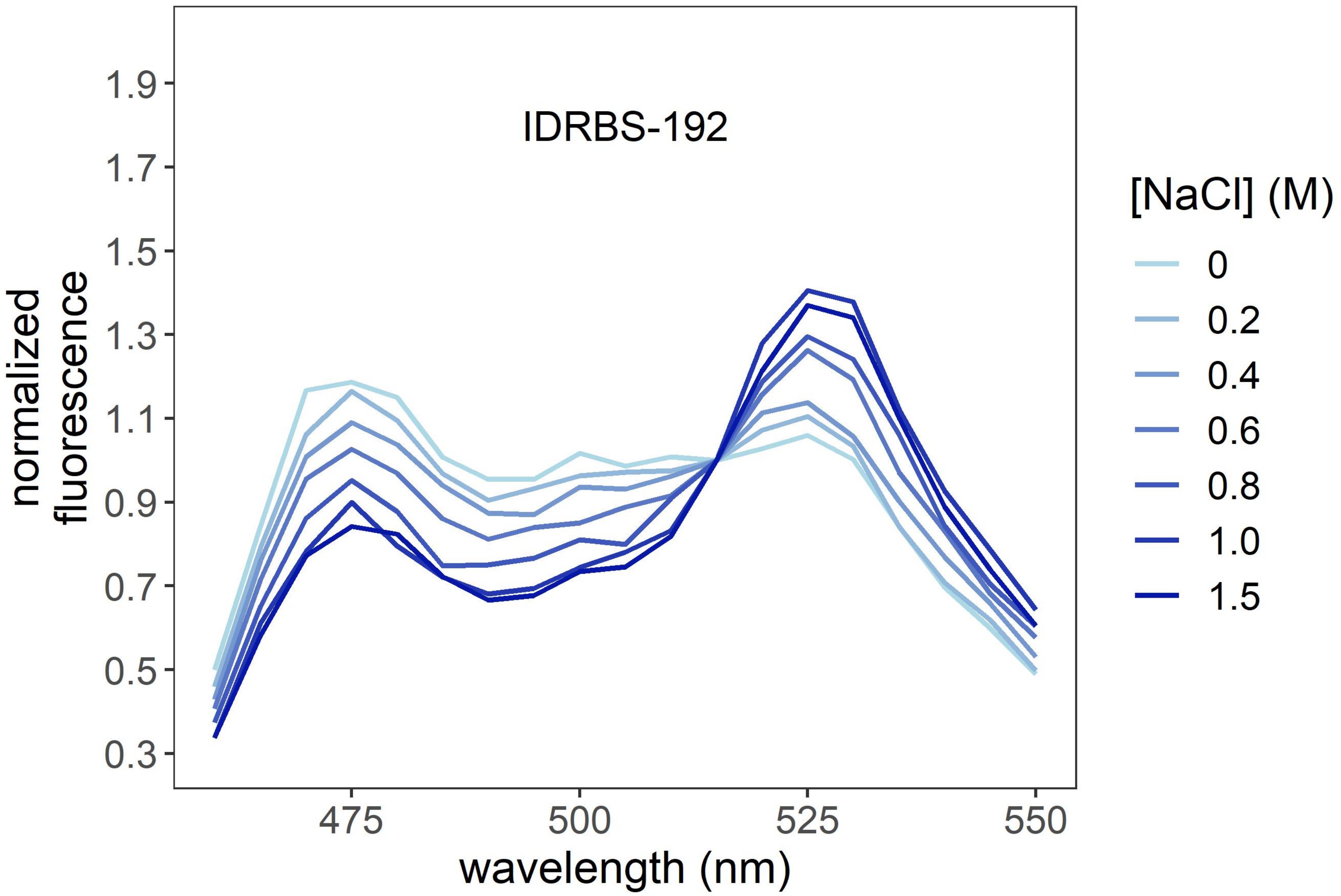
IDRBS-192
normalized fluorescence
1.9
1.7
1.5
1.3
1.1
0.9
0.7
0.5
0.3
475
500
525
550
wavelength (nm)
[NaCl] (M)
0
0.2
0.4
0.6
0.8
1.0
1.5

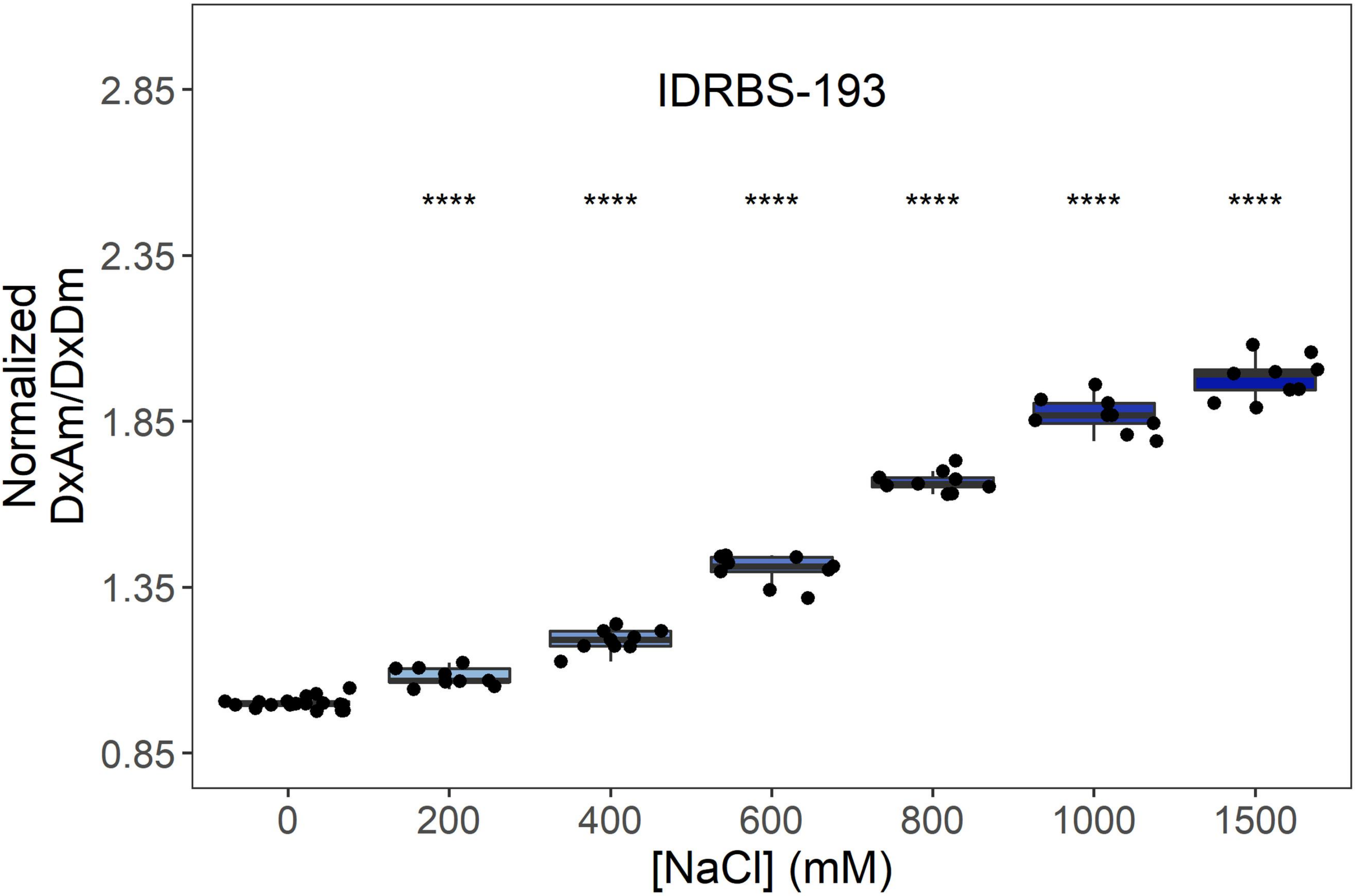
IDRBS-193
Normalized
DxAm/DxDm
2.85
2.35
1.85
1.35
0.85
****
0
200
400
600
800
1000
1500
[NaCl] (mM)

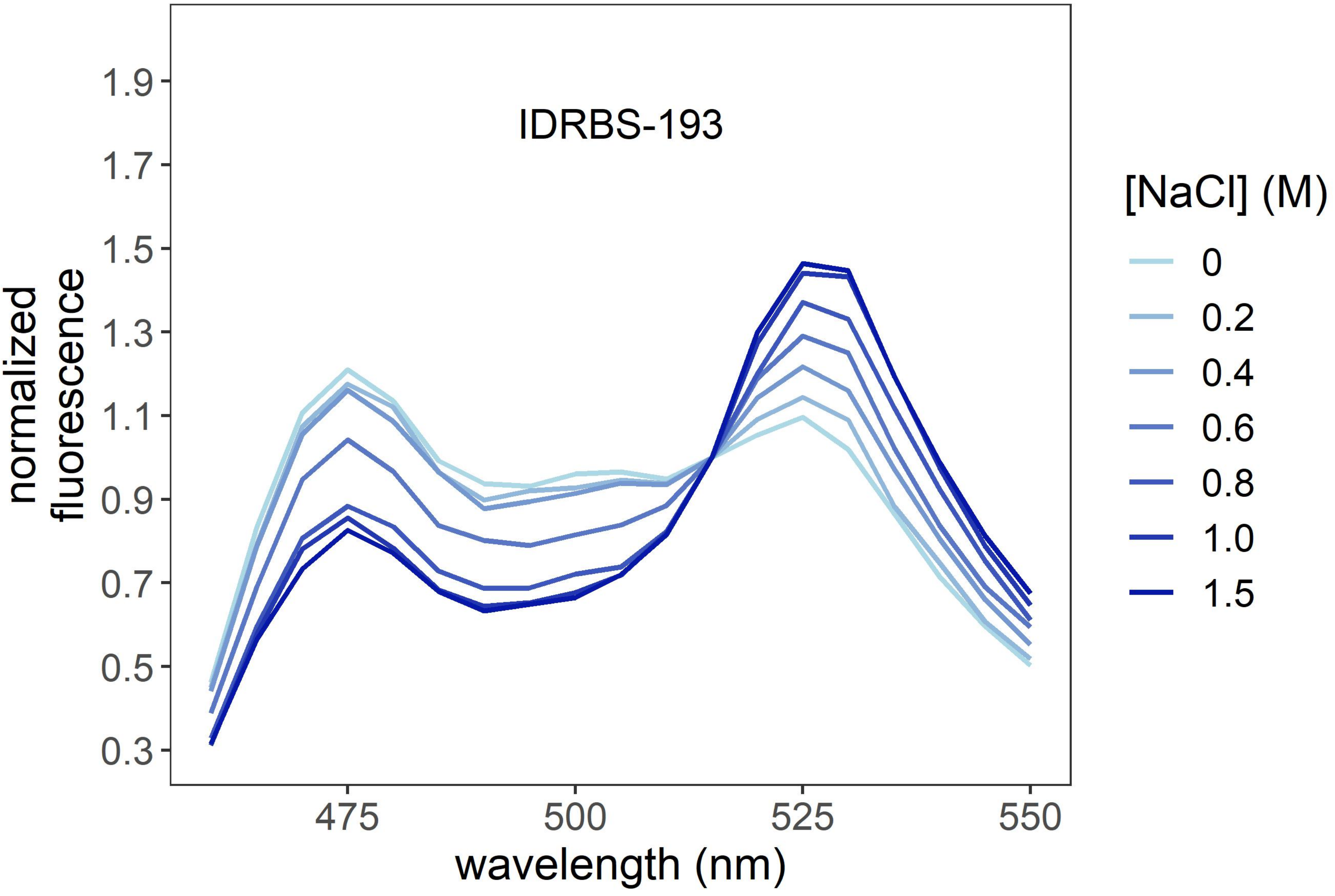
IDRBS-193
normalized
fluorescence
1.9
1.7
1.5
1.3
1.1
0.9
0.7
0.5
0.3
475
500
525
550
wavelength (nm)
[NaCl] (M)
0
0.2
0.4
0.6
0.8
1.0
1.5

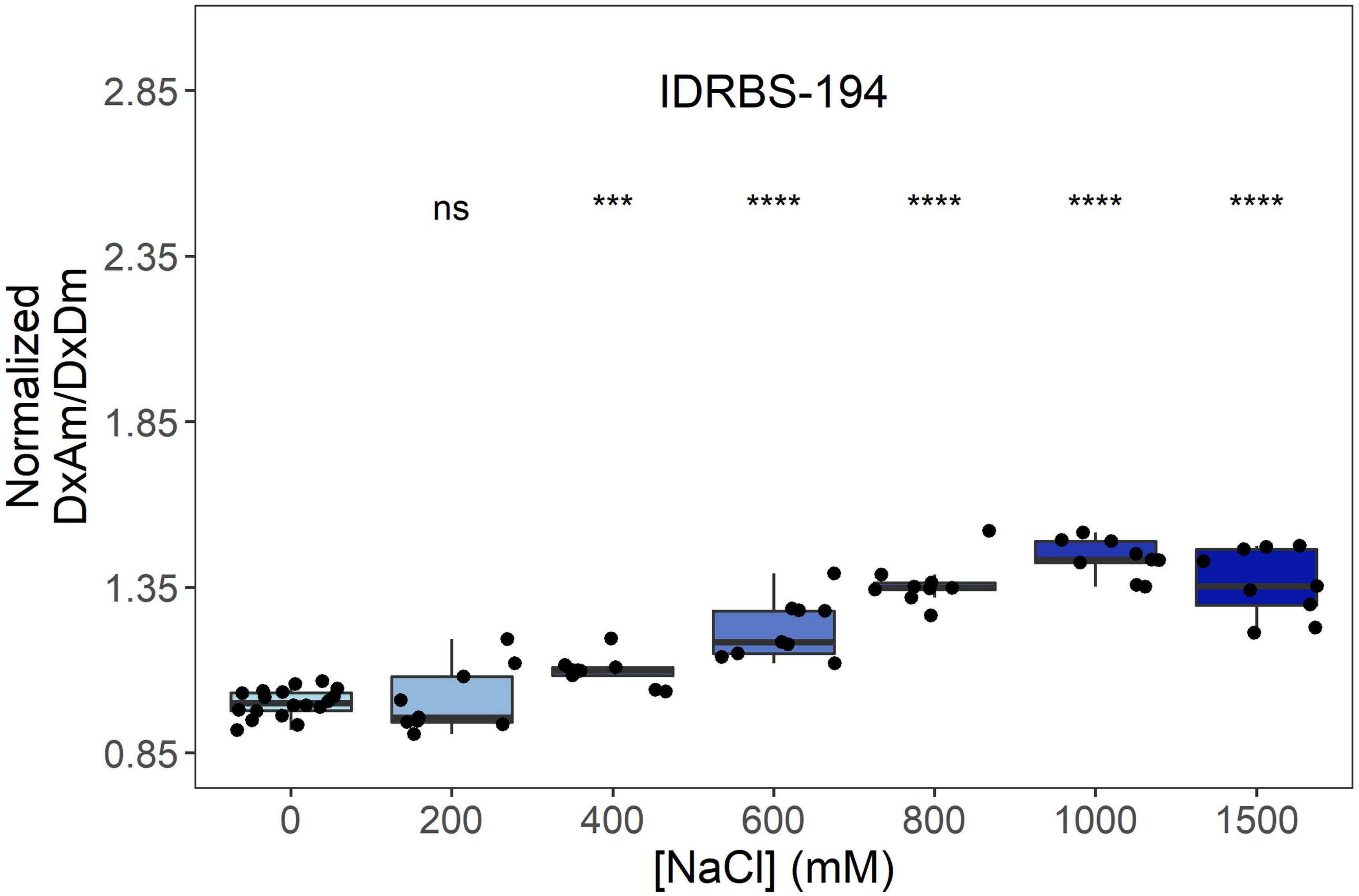
IDRBS-194
ns
***
****
****
****
****
Normalized
DxAm/DxDm
2.85
2.35
1.85
1.35
0.85
0
200
400
600
800
1000
1500
[NaCl] (mM)

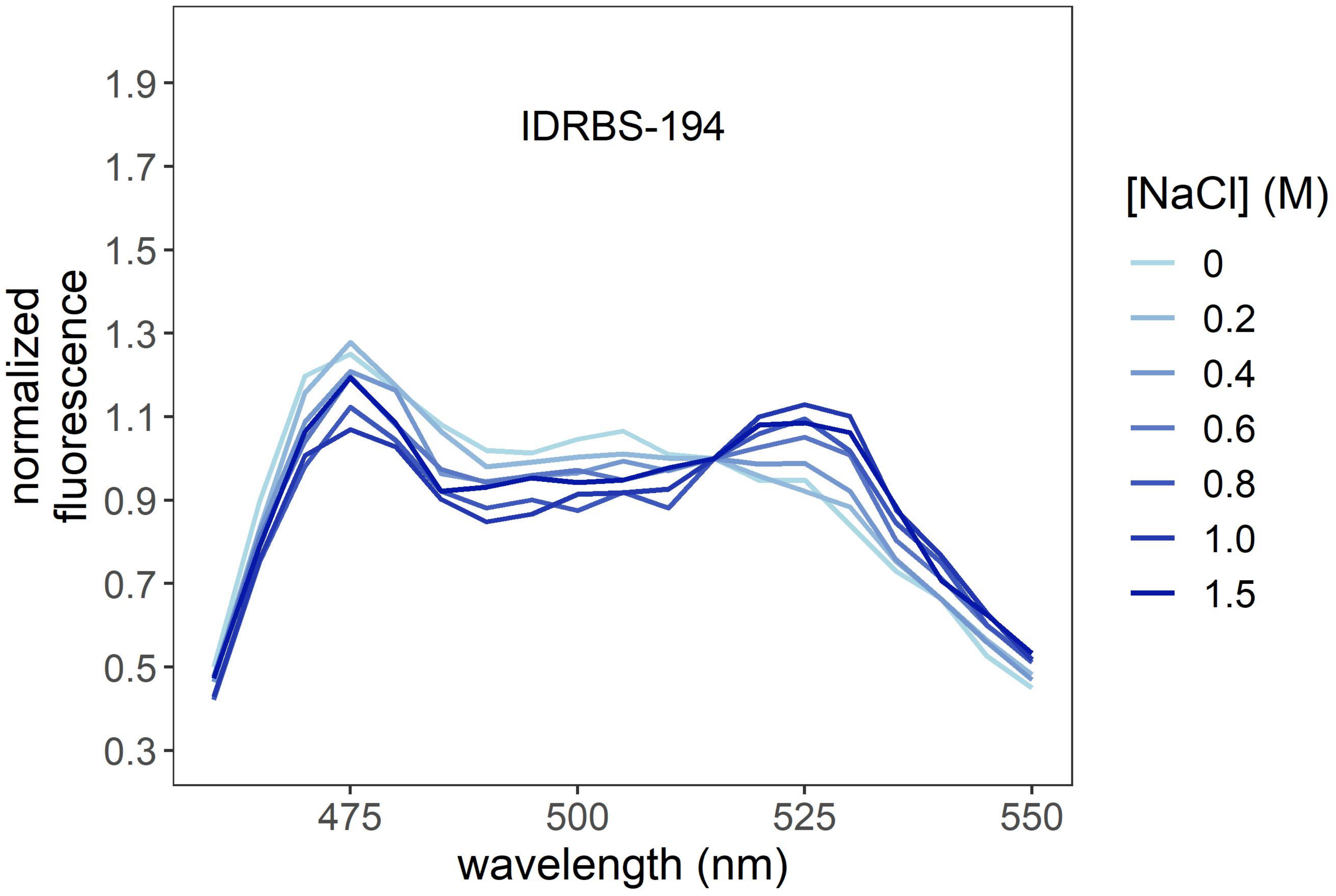
IDRBS-194
normalized
fluorescence
1.9
1.7
1.5
1.3
1.1
0.9
0.7
0.5
0.3
475
500
525
550
wavelength (nm)
[NaCl] (M)
0
0.2
0.4
0.6
0.8
1.0
1.5

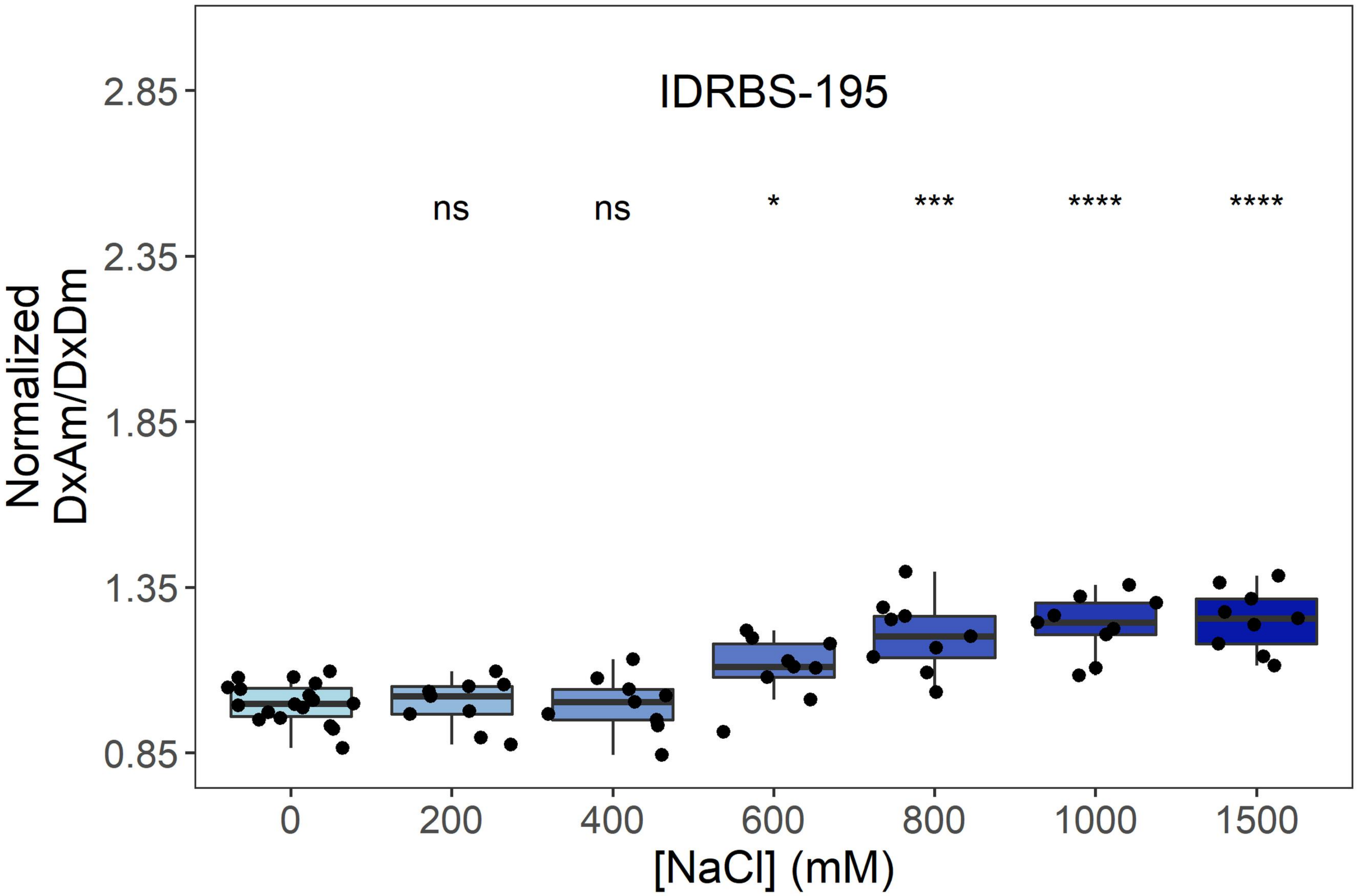
IDRBS-195
ns
ns
*
***
****
****
2.85
2.35
1.85
1.35
0.85
Normalized
DxAm/DxDm
0
200
400
600
800
1000
1500
[NaCl] (mM)

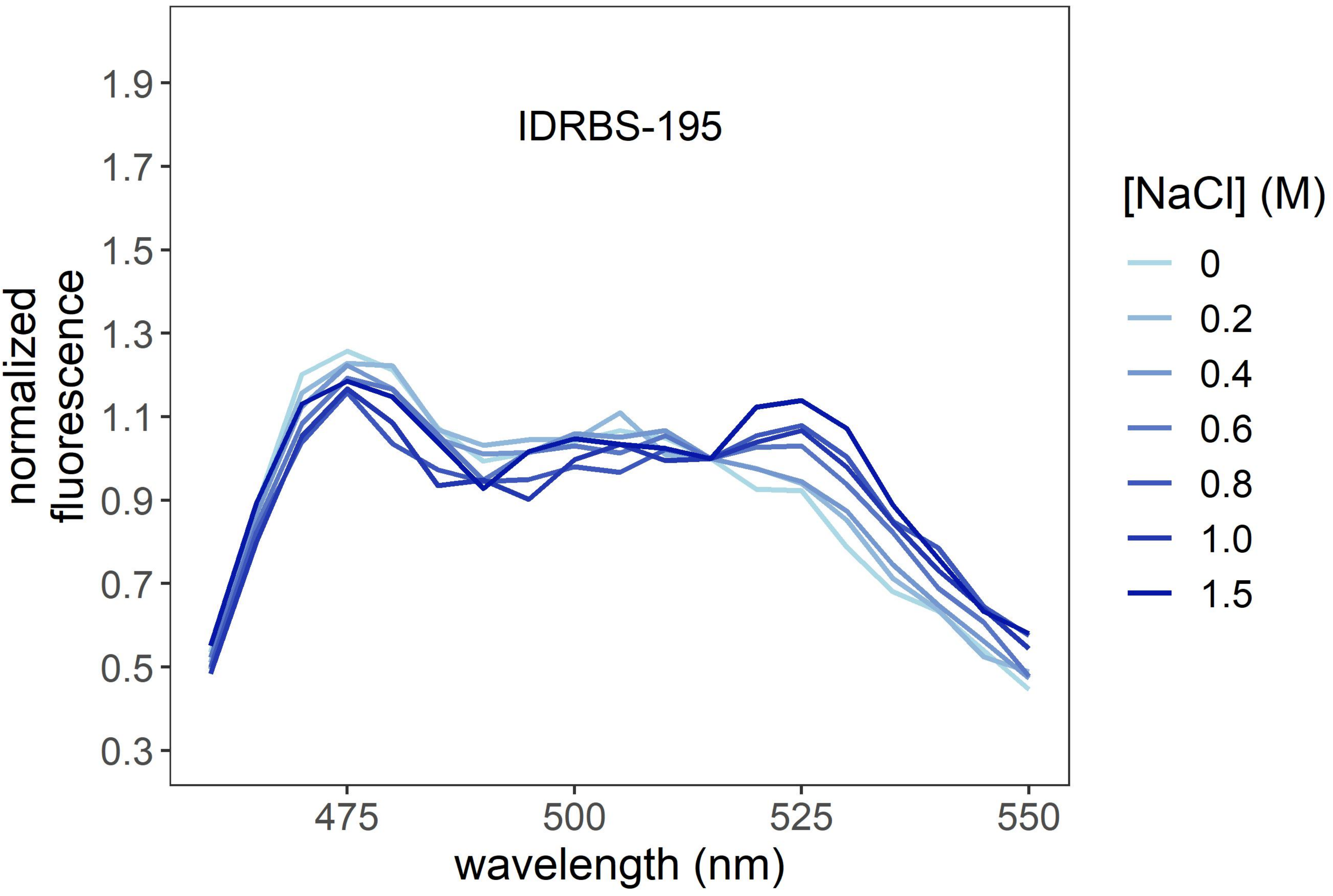
IDRBS-195
1.9
1.7
1.5
1.3
1.1
0.9
0.7
0.5
0.3
normalized
fluorescence
475
500
525
550
wavelength (nm)
[NaCl] (M)
0
0.2
0.4
0.6
0.8
1.0
1.5

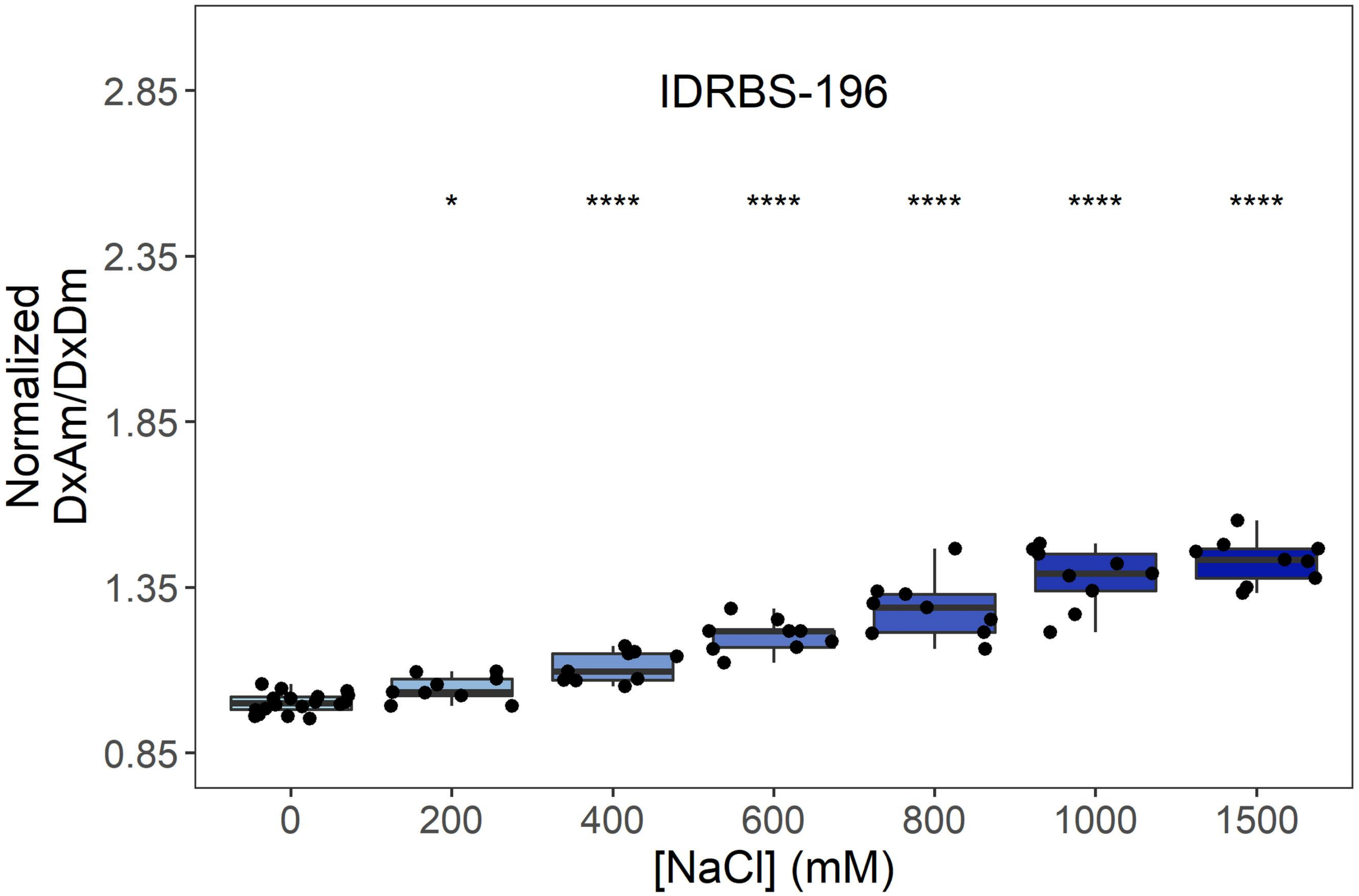

IDRBS-196
Normalized
DxAm/DxDm
2.85
2.35
1.85
1.35
0.85
*
****
****
****
****
****
0
200
400
600
800
1000
1500
[NaCl] (mM)


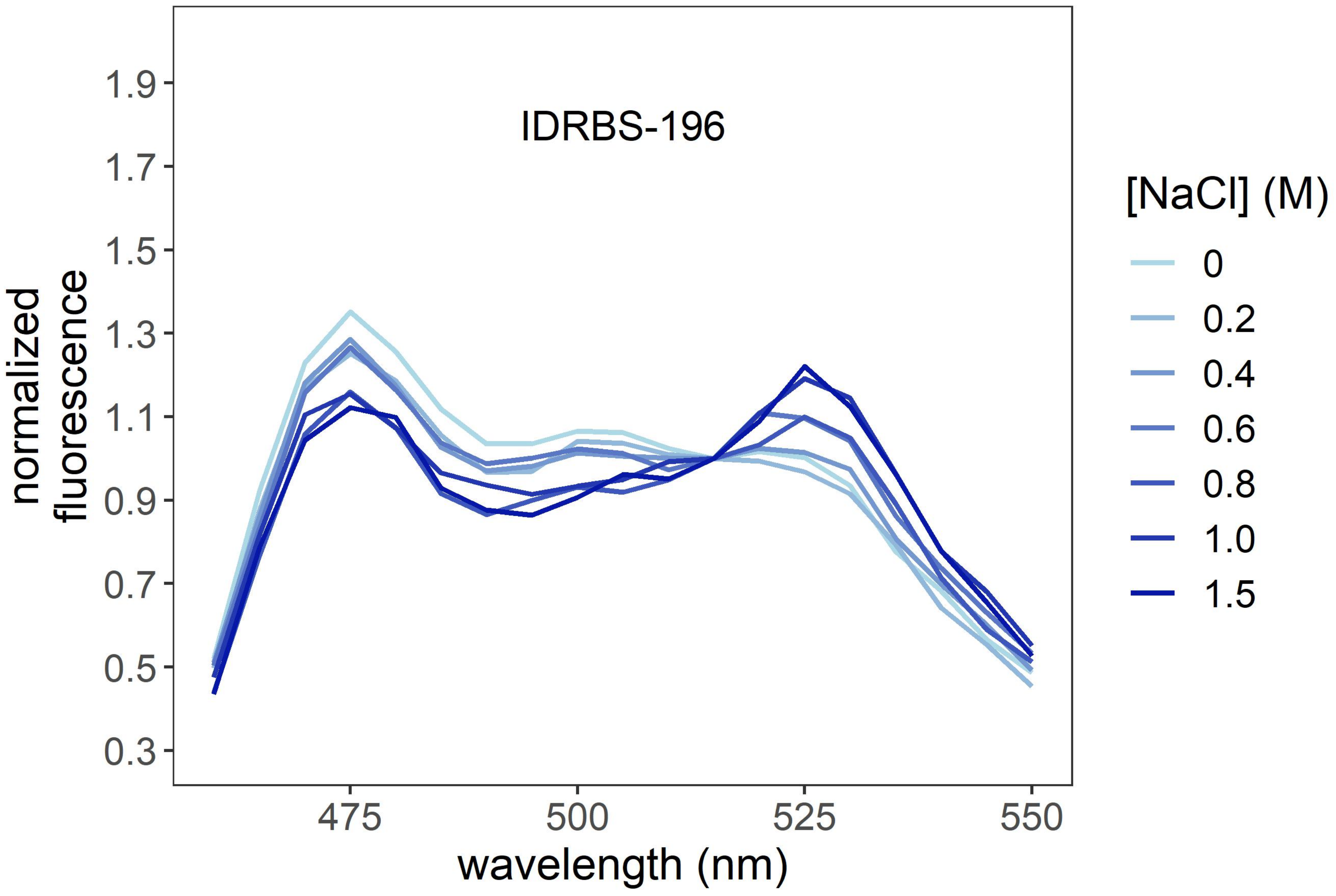

IDRBS-196
normalized
fluorescence
1.9
1.7
1.5
1.3
1.1
0.9
0.7
0.5
0.3
475
500
525
550
wavelength (nm)
[NaCl] (M)
0
0.2
0.4
0.6
0.8
1.0
1.5

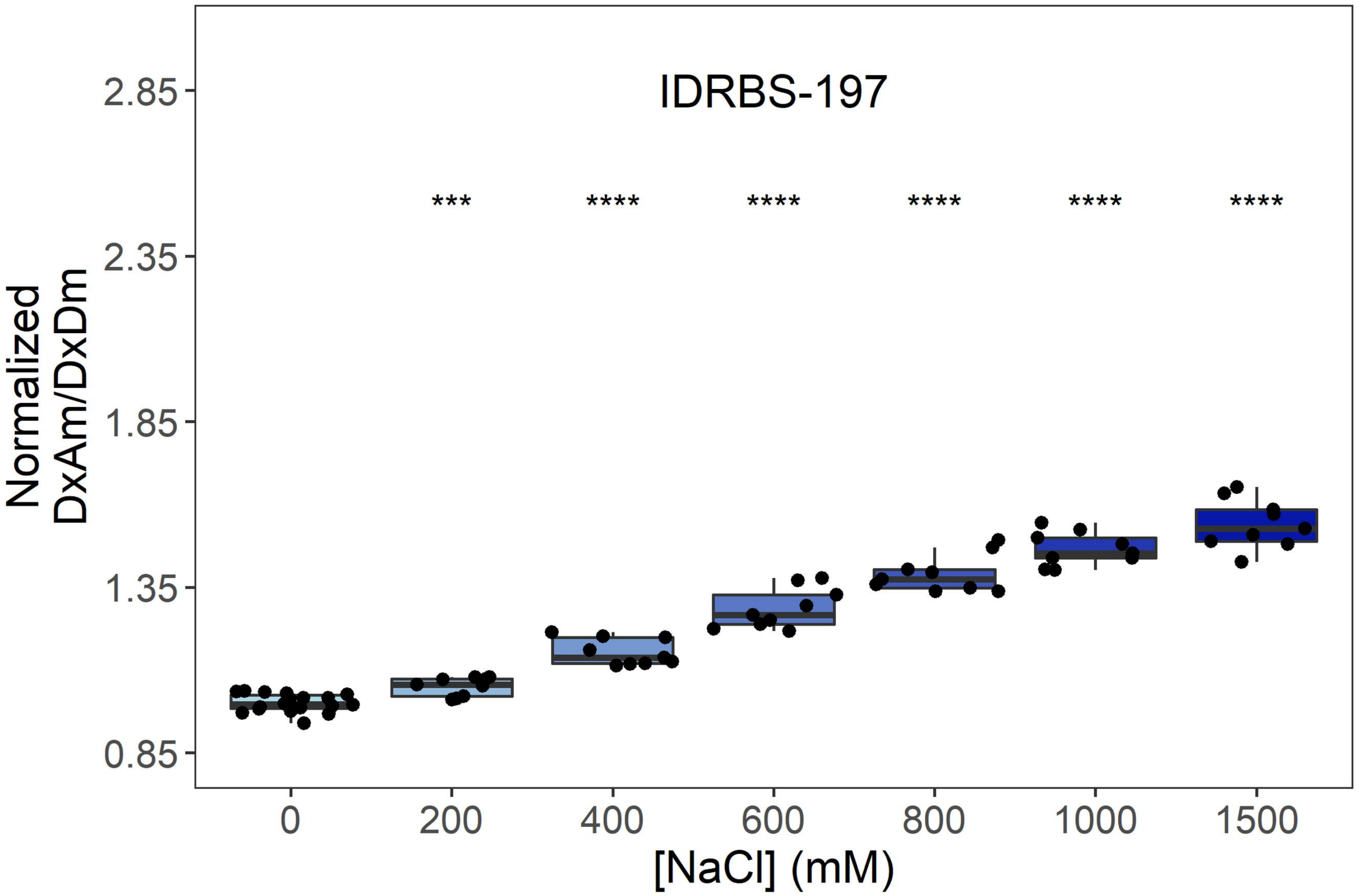
IDRBS-197
*** **** **** **** **** ****
Normalized DxAm/DxDm
2.85
2.35
1.85
1.35
0.85
0 200 400 600 800 1000 1500
[NaCl] (mM)

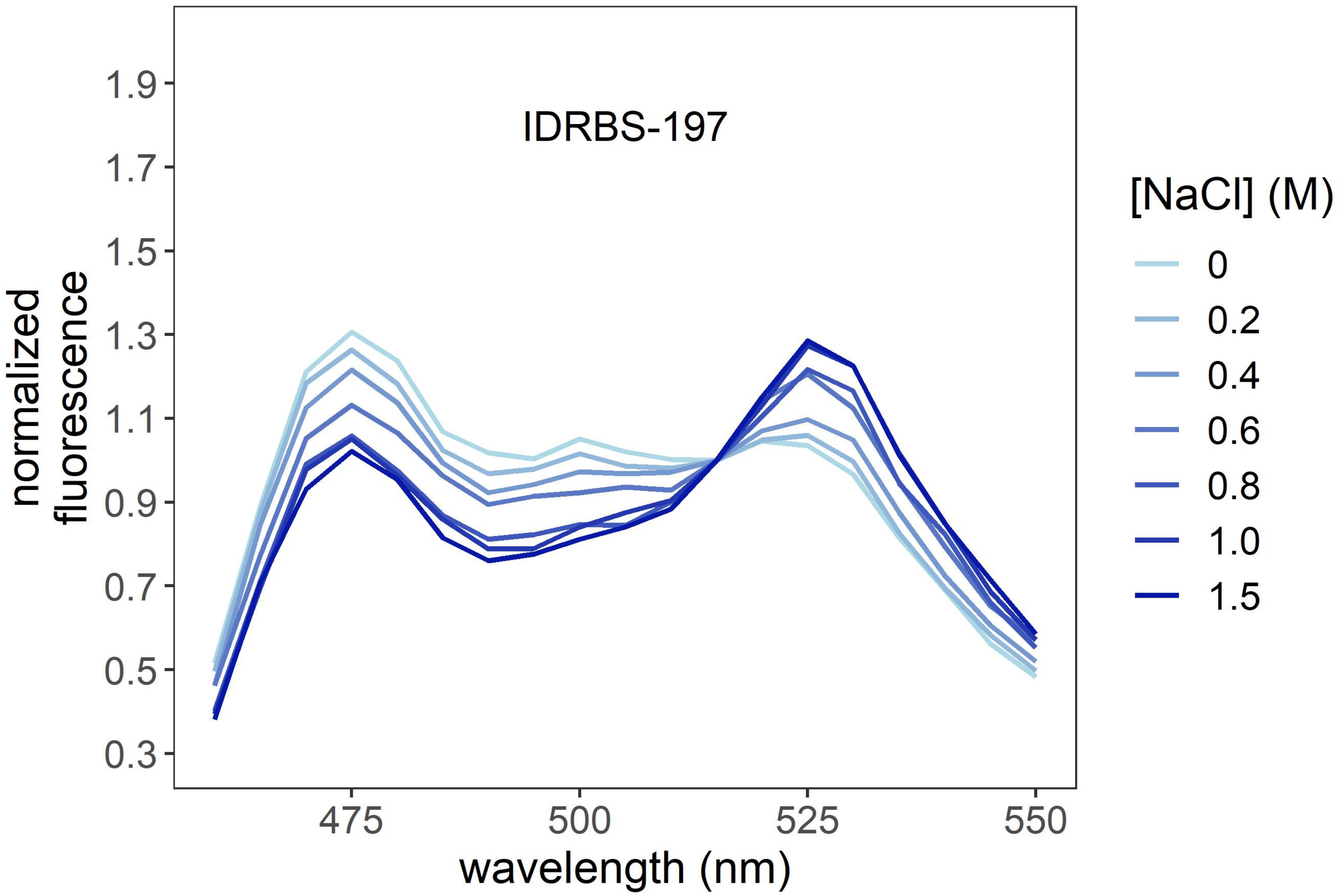
IDRBS-197
normalized fluorescence
1.9
1.7
1.5
1.3
1.1
0.9
0.7
0.5
0.3
475 500 525 550
wavelength (nm)
[NaCl] (M)
0
0.2
0.4
0.6
0.8
1.0
1.5

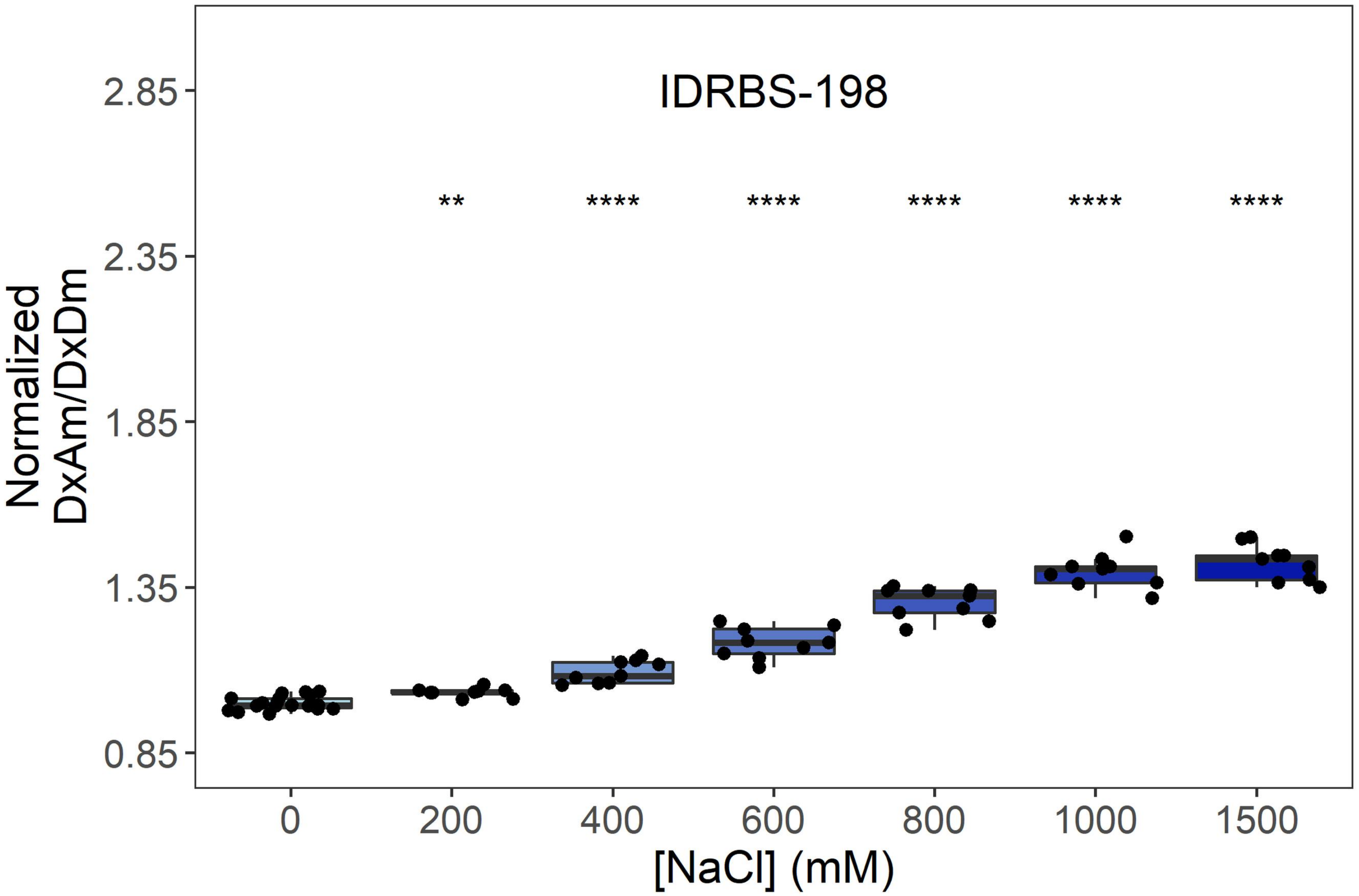
IDRBS-198
**
****
****
****
****
****
2.85
2.35
1.85
1.35
0.85
Normalized
DxAm/DxDm
0
200
400
600
800
1000
1500
[NaCl] (mM)

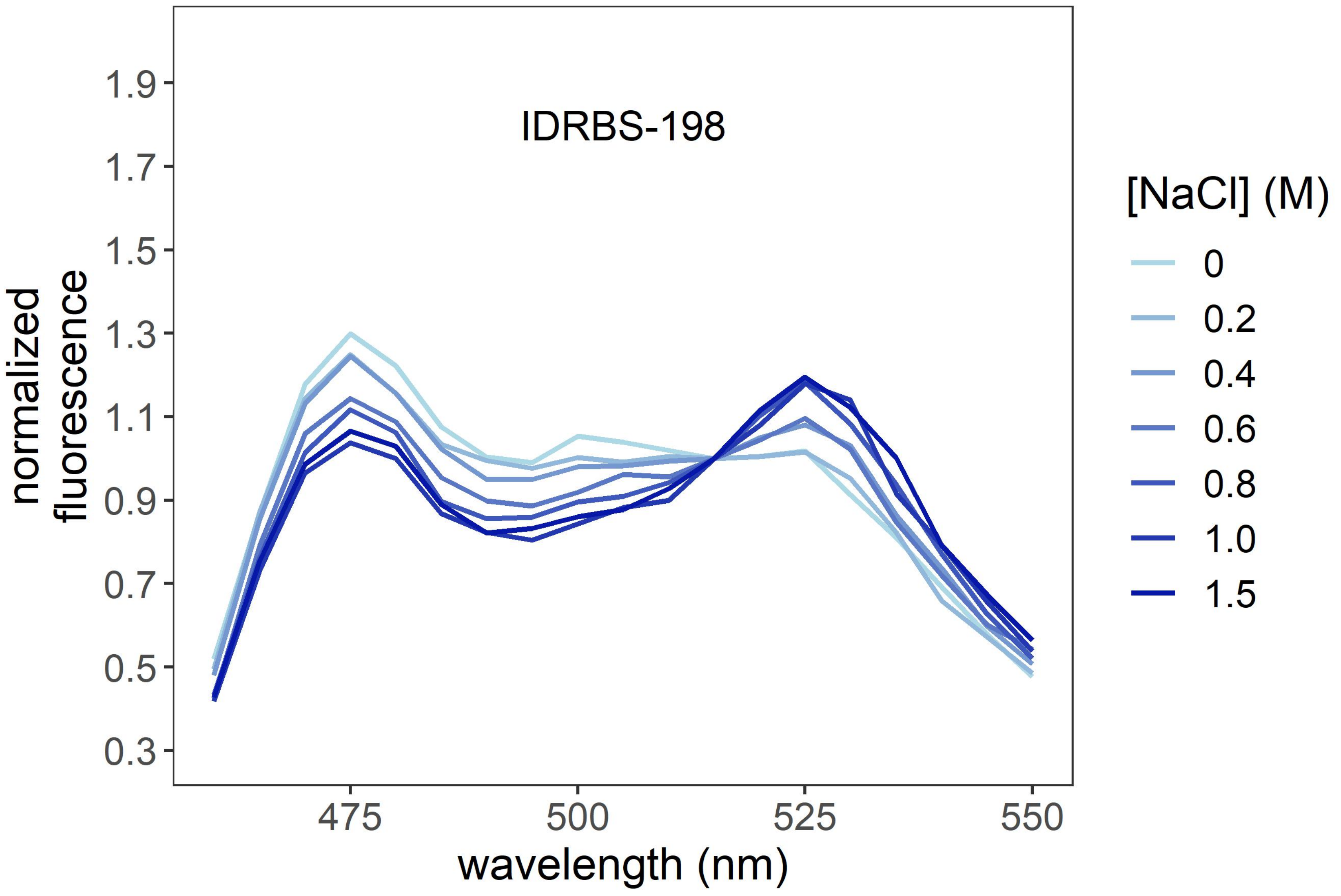
IDRBS-198
1.9
1.7
1.5
1.3
1.1
0.9
0.7
0.5
0.3
normalized
fluorescence
475
500
525
550
wavelength (nm)
[NaCl] (M)
0
0.2
0.4
0.6
0.8
1.0
1.5

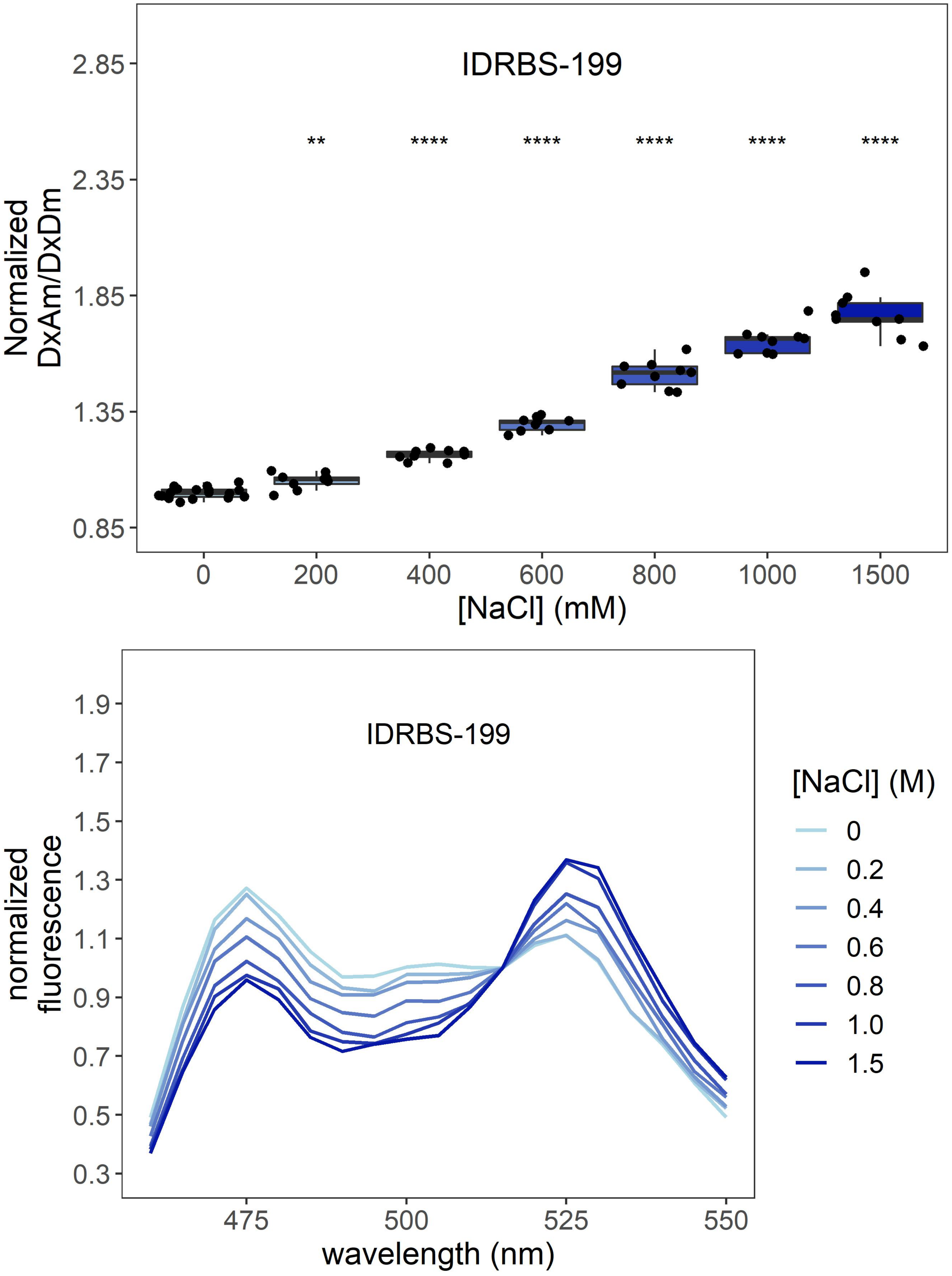
IDRBS-199
**
****
****
****
****
****
Normalized DxAm/DxDm
2.85
2.35
1.85
1.35
0.85
0
200
400
600
800
1000
1500
[NaCl] (mM)
IDRBS-199
normalized fluorescence
1.9
1.7
1.5
1.3
1.1
0.9
0.7
0.5
0.3
475
500
525
550
wavelength (nm)
[NaCl] (M)
0
0.2
0.4
0.6
0.8
1.0
1.5

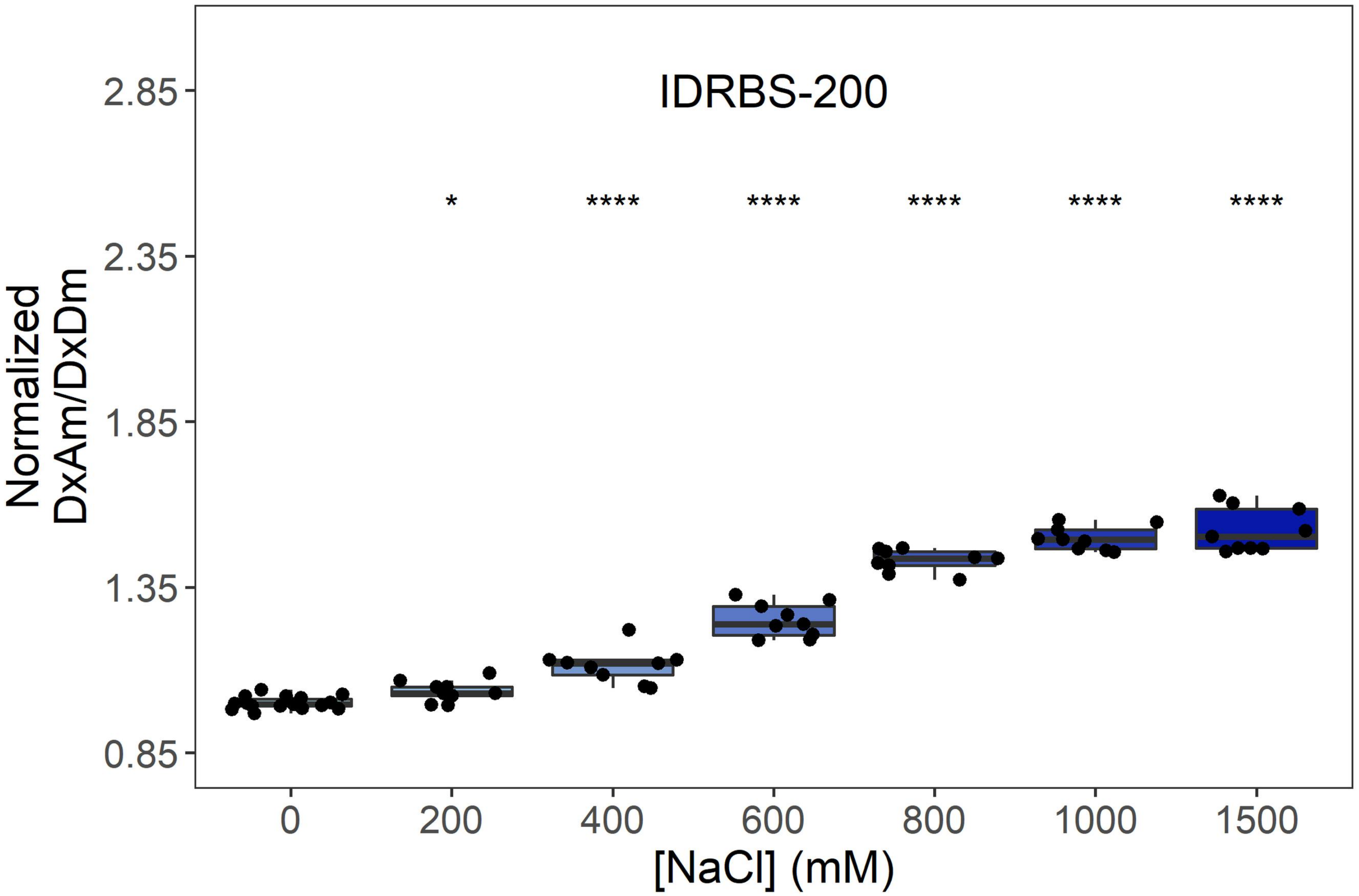
IDRBS-200
Normalized DxAm/DxDm
2.85
2.35
1.85
1.35
0.85
*
****
****
****
****
****
0
200
400
600
800
1000
1500
[NaCl] (mM)

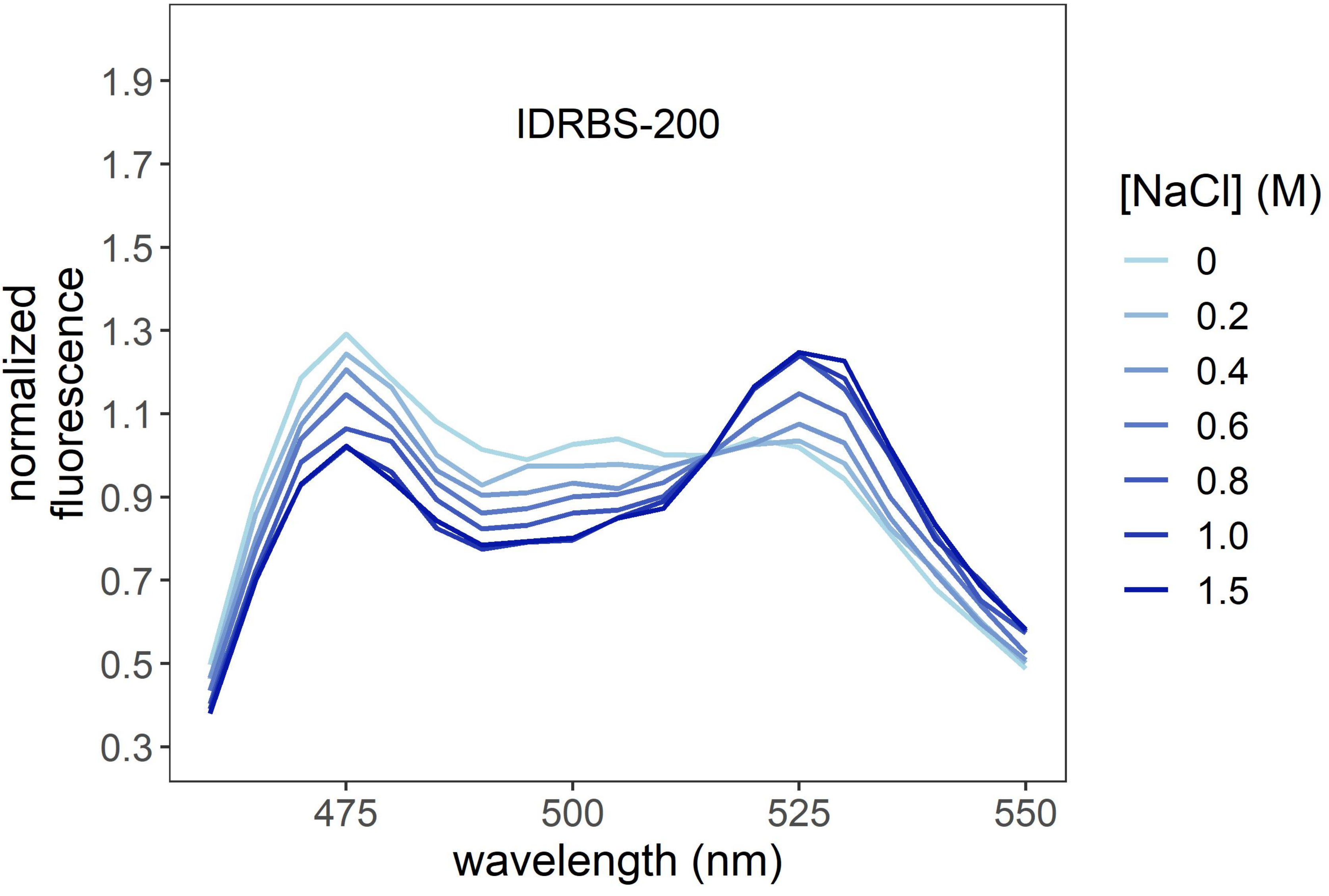
IDRBS-200
normalized fluorescence
1.9
1.7
1.5
1.3
1.1
0.9
0.7
0.5
0.3
475
500
525
550
wavelength (nm)
[NaCl] (M)
0
0.2
0.4
0.6
0.8
1.0
1.5
